## Supplementary Information for "In vitro demonstration of broad host range mobile genetic elements transferring antibiotic resistance from the human microbiome"

```
DEFINITION     Integrative and conjugative element ICE_1
   CDS           complement(1..1772)
             /color=9
             /locus_tag="B40_04892"
             /note="Transposon Tn1000 resolvase, similar to AA
             sequence:UniProtKB:P03012 protein motif:CLUSTERS:PRK13413
             protein motif:Pfam:PF00239.15"
             /product="Transposon Tn1000 resolvase [DDE recombinase]"
             /"ID=B40_04892
             /gene="tnpR"""
             /protein_id="gnl|SC|B40_04892"
             /codon_start=1
   fasta_record    1..61464
             /ID=B40|SC|contig000003
             /color=5
   CDS           complement(2360..2587)
             /color=7
             /locus_tag="B40_04893"
             /note="Helix-turn-helix domain, protein
             motif:Pfam:PF12645.1"
             /product="Helix-turn-helix domain [transcriptional
             regulator]"
             /"ID=B40_04893
             /protein_id="gnl|SC|B40_04893"""
             /codon_start=1
   CDS           complement(2580..3008)
             /color=7
             /locus_tag="B40_04894"
             /note="RNA polymerase sigma factor Sigma-70 region 4,
             protein motif:TIGRFAMs:TIGR02937 protein
             motif:Pfam:PF08281.6"
             /product="RNA polymerase sigma factor  Sigma-70  region 4
             [transcriptional regulator]"
             /"ID=B40_04894
             /protein_id="gnl|SC|B40_04894"""
             /codon_start=1
   CDS           complement(3377..3544)
             /ID=B40_04895
             /color=1
             /locus_tag="B40_04895"
             /note="Hypothetical protein"
             /product="hypothetical protein"
             /protein_id="gnl|SC|B40_04895"
             /codon_start=1
```

CDS complement(3492..4472)
/color=7
/locus_tag="B40_04896"
/note="AraC family transcriptional regulator
Helix-turn-helix domain, similar to AA
sequence:UniProtKB:P28816 protein motif:CLUSTERS:PRK15121
protein motif:Cdd:COG3449 protein motif:TIGRFAMs:TIGR04094
protein motif:Pfam:PF12833.1"
/product="AraC family transcriptional regulator
Helix-turn-helix domain [transcriptional regulator]"
/"ID=B40_04896
/protein_id="gnl|SC|B40_04896"""
/codon_start=1
CDS complement(4838..5029)
/ID=B40_04897
/color=1
/locus_tag="B40_04897"
/note="Hypothetical protein"
/product="hypothetical protein  DUF4177 containing
protein"
/protein_id="gnl|SC|B40_04897"
/codon_start=1
CDS complement(5100..5393)
/color=8
/locus_tag="B40_04898"
/note="Predicted ester cyclase  SnoaL-like polyketide
cyclase, protein motif:Cdd:COG5485 protein
motif:TIGRFAMs:TIGR02096 protein motif:Pfam:PF07366.6"
/product="Predicted ester cyclase  SnoaL-like polyketide
cyclase [ester cyclase]"
/"ID=B40_04898
/protein_id="gnl|SC|B40_04898"""
/codon_start=1
CDS complement(5412..5510)
/ID=B40_04899
/color=1
/locus_tag="B40_04899"
/note="Hypothetical protein"
/product="hypothetical protein"
/protein_id="gnl|SC|B40_04899"
/codon_start=1
CDS complement(5580..6194)
/color=7
/locus_tag="B40_04900"

/note="y4mF family Helix-turn-helix, protein motif:TIGRFAMs:TIGR03070 protein motif:Pfam:PF01381.16"
/product="y4mF family Helix-turn-helix [transcriptional regulator]"
/"ID=B40_04900
/protein_id="gnl|SC|B40_04900"""
/codon_start=1
CDS complement(6218..6877)
/color=8
/locus_tag="B40_04901"
/note="PAP2 superfamily, protein motif:Pfam:PF01569.15"
/product="PAP2 superfamily [phosphatase]"
/"ID=B40_04901
/protein_id="gnl|SC|B40_04901"""
/codon_start=1
CDS complement(7160..7876)
/color=8
/locus_tag="B40_04902"
/eC_number=2.1.1.157
/note="Sarcosine/dimethylglycine N-methyltransferase, similar to AA sequence:UniProtKB:Q83WC3 protein motif:TIGRFAMs:TIGR02021 protein motif:Pfam:PF08241.6"
/product="Sarcosine/dimethylglycine N-methyltransferase [methyltransferase]"
/"ID=B40_04902
/protein_id="gnl|SC|B40_04902"""
/codon_start=1
CDS complement(7998..8624)
/color=7
/locus_tag="B40_04903"
/note="Potential acrAB operon repressor, similar to AA sequence:UniProtKB:P0ACS9 protein motif:CLUSTERS:PRK10668 protein motif:TIGRFAMs:TIGR03968 protein motif:Pfam:PF00440.17"
/product="Potential acrAB operon repressor [transcriptional regulator]"
/"ID=B40_04903
/gene="acrR"""
/protein_id="gnl|SC|B40_04903"
/codon_start=1
CDS complement(8914..9054)
/ID=B40_04904
/color=1
/locus_tag="B40_04904"

/note="Hypothetical protein"
/product="hypothetical protein"
/protein_id="gnl|SC|B40_04904"
/codon_start=1
CDS complement(9105..10016)
/color=8
/locus_tag="B40_04905"
/note="Cadmium  cobalt and zinc/H-K antiporter, similar to
AA sequence:UniProtKB:O07084 protein
motif:CLUSTERS:PRK09509 protein motif:Cdd:COG0053 protein
motif:TIGRFAMs:TIGR01297 protein motif:Pfam:PF01545.15"
/product="Cadmium  cobalt and zinc/H%28 %29-K%28 %29
antiporter [heavy metal defense]"
/"ID=B40_04905
/gene="czcD"""
/protein_id="gnl|SC|B40_04905"
/codon_start=1
CDS complement(10028..10471)
/ID=B40_04906
/color=8
/locus_tag="B40_04906"
/note="Hypothetical protein"
/product="RibD %28riboflavin biosynthesis%29 family
protein [pyrimidine reductase]"
/protein_id="gnl|SC|B40_04906"
/codon_start=1
CDS complement(10582..10938)
/ID=B40_04907
/color=7
/locus_tag="B40_04907"
/note="Hypothetical protein"
/product="WYL domain-containing protein [transcriptional
regulator]"
/protein_id="gnl|SC|B40_04907"
/codon_start=1
CDS complement(10962..11534)
/color=7
/locus_tag="B40_04908"
/note="Helix-turn-helix domain, protein
motif:CLUSTERS:PRK11886 protein motif:Pfam:PF08279.6"
/product="Helix-turn-helix domain [transcriptional
regulator]"
/"ID=B40_04908
/protein_id="gnl|SC|B40_04908""""

/codon_start=1
CDS complement(11810..12604)
/color=2
/locus_tag="B40_04909"
/eC_number=2.7.1.95
/note="Aminoglycoside 3'-phosphotransferase, similar to AA sequence:UniProtKB:P0A3Y5 protein motif:Cdd:COG3231 protein motif:Pfam:PF01636.17"
/product="Aminoglycoside 3'-phosphotransferase [antibiotic resistance]"
/"ID=B40_04909
/gene="aphA"""
/protein_id="gnl|SC|B40_04909"
/codon_start=1
CDS complement(12697..13239)
/color=2
/locus_tag="B40_04910"
/note="streptothricin N-acetyltransferase Sat4, protein motif:CLUSTERS:PRK10975 protein motif:TIGRFAMs:TIGR01575 protein motif:Pfam:PF00583.18"
/product="streptothricin N-acetyltransferase Sat4 [antibiotic resistance]"
/"ID=B40_04910
/gene="sat4"""
/protein_id="gnl|SC|B40_04910"
/codon_start=1
CDS complement(13236..14144)
/color=2
/locus_tag="B40_04911"
/eC_number=2.7.7.-
/note="Aminoglycoside 6-adenylyl transferase Streptomycin adenylyl transferase, similar to AA sequence:UniProtKB:P17585 protein motif:Pfam:PF04439.6"
/product="Aminoglycoside 6-adenylyltransferase Streptomycin adenylyltransferase [antibiotic resistance]"
/"ID=B40_04911
/gene="aadK"""
/protein_id="gnl|SC|B40_04911"
/codon_start=1
CDS complement(14177..14785)
/color=8
/locus_tag="B40_04912"
/eC_number=2.1.1.-
/note="Uncharacterized methyltransferase ycgJ , similar to

AA sequence:UniProtKB:O31474 protein
motif:CLUSTERS:PRK08317 protein motif:Cdd:COG4976 protein
motif:TIGRFAMs:TIGR02072 protein motif:Pfam:PF08241.6"
/product="Uncharacterized methyltransferase ycgJ
[methyltransferase]"
/"ID=B40_04912
/gene="ycgJ"""
/protein_id="gnl|SC|B40_04912"
/codon_start=1
CDS complement(14769..15524)
/ID=B40_04913
/color=10
/locus_tag="B40_04913"
/note="Hypothetical protein"
/product="Replication protein repR [replication
initiation]"
/gene="repR"
/protein_id="gnl|SC|B40_04913"
/codon_start=1
ncRNA 15682..15812
/ID=B40_04914
/color=10
/locus_tag="B40_04914"
/product="Plasmid_RNAIII [repR repressor]"
CDS complement(15919..16158)
/color=10
/locus_tag="B40_04915"
/note="Replication initiation protein RepA, protein
motif:Pfam:PF01381.16"
/product="Replication initiation protein RepA
[replication]"
/"ID=B40_04915
/gene="repA"""
/protein_id="gnl|SC|B40_04915"
/codon_start=1
CDS complement(16402..16515)
/ID=B40_04916
/color=1
/locus_tag="B40_04916"
/note="Hypothetical protein"
/product="hypothetical protein"
/protein_id="gnl|SC|B40_04916"
/codon_start=1
CDS complement(16520..17257)

/color=2
/locus_tag="B40_04917"
/eC_number=2.1.1.184
/note="rRNA adenine N-6-methyltransferase, similar to AA sequence:UniProtKB:P21236 protein motif:CLUSTERS:PRK00274 protein motif:Cdd:COG0030 protein motif:TIGRFAMs:TIGR00755 protein motif:Pfam:PF00398.14"
/product="rRNA adenine N-6-methyltransferase [antibiotic resistanc]"
/"ID=B40_04917
/gene="ermB"""
/protein_id="gnl|SC|B40_04917"
/codon_start=1
CDS complement(17886..18251)
/color=1
/locus_tag="B40_04918"
/note="Predicted outer membrane protein LPXTG cell wall anchor domain, protein motif:Cdd:COG4932 protein motif:TIGRFAMs:TIGR01167 protein motif:Pfam:PF00746.15"
/product="Predicted outer membrane protein LPXTG cell wall anchor domain [membrane protein]"
/"ID=B40_04918
/protein_id="gnl|SC|B40_04918"""
/codon_start=1
CDS complement(18251..19165)
/color=8
/locus_tag="B40_04919"
/note="ABC transporter permease, protein motif:CLUSTERS:PRK05625 protein motif:TIGRFAMs:TIGR00227 protein motif:Pfam:PF01872.11"
/product="ABC transporter permease [transporter]"
/"ID=B40_04919
/protein_id="gnl|SC|B40_04919"""
/codon_start=1
CDS complement(19162..19845)
/ID=B40_04920
/color=8
/locus_tag="B40_04920"
/note="Hypothetical protein"
/product="ABC transporter permease [transporter]"
/protein_id="gnl|SC|B40_04920"
/codon_start=1
CDS complement(19842..20687)
/color=8

/locus_tag="B40_04921"
/eC_number=3.6.3.-
/note="ABC transporter ATP-binding protein, similar to AA sequence:UniProtKB:O07190 protein motif:CLUSTERS:PRK13536 protein motif:Cdd:COG4555 protein motif:TIGRFAMs:TIGR01188 protein motif:Pfam:PF00005.21"
/product="ABC transporter ATP-binding protein [ATPase]"
/"ID=B40_04921
/protein_id="gnl|SC|B40_04921"""
/codon_start=1
CDS complement(20791..21378)
/color=7
/locus_tag="B40_04922"
/note="tetR/arcR family transcriptional regulator, protein motif:CLUSTERS:PRK09975 protein motif:TIGRFAMs:TIGR03968 protein motif:Pfam:PF00440.17"
/product="tetR/arcR family transcriptional regulator [transcriptional regulator]"
/"ID=B40_04922
/protein_id="gnl|SC|B40_04922"""
/codon_start=1
CDS complement(21450..22307)
/color=8
/locus_tag="B40_04923"
/note="Alpha/beta hydrolase family, protein motif:Cdd:COG1647 protein motif:Pfam:PF12697.1"
/product="Alpha/beta hydrolase family [hydrolase]"
/"ID=B40_04923
/protein_id="gnl|SC|B40_04923"""
/codon_start=1
CDS complement(22520..23413)
/color=4
/locus_tag="B40_04924"
/note="Conjugative transposon protein TcpC, protein motif:Pfam:PF12642.1"
/product="Conjugative transposon protein TcpC [mating pair formation]"
/"ID=B40_04924
/gene="tcpC"""
/protein_id="gnl|SC|B40_04924"
/codon_start=1
CDS complement(23429..24436)
/color=4
/locus_tag="B40_04925"

/note="Peptidase P60 bifunctional lytic
transglycosylase/NlpC/P60 family protein, similar to AA
sequence:UniProtKB:P21171 protein motif:CLUSTERS:PRK13914
protein motif:Cdd:COG3584 protein motif:Pfam:PF00877.13"
/product="Peptidase P60 bifunctional lytic
transglycosylase/NlpC/P60 family protein [pepidase]"
/"ID=B40_04925
/protein_id="gnl|SC|B40_04925"""
/codon_start=1
CDS complement(24433..26565)
/ID=B40_04926
/color=1
/locus_tag="B40_04926"
/note="Hypothetical protein"
/product="YtxH domain-containing protein  MFS transporter
membrane protein  conjugative transposon protein"
/protein_id="gnl|SC|B40_04926"
/codon_start=1
CDS complement(26565..29015)
/color=4
/locus_tag="B40_04927"
/note="Type IV secretion system protein TraC  VirB4
components, protein motif:Cdd:COG3451 protein
motif:TIGRFAMs:TIGR02746 protein motif:Pfam:PF12846.1"
/product="Type IV secretion system protein TraC  VirB4
components [ATPase  mating pair formation]"
/"ID=B40_04927
/gene="traC"""
/protein_id="gnl|SC|B40_04927"
/codon_start=1
CDS complement(29002..29391)
/color=4
/locus_tag="B40_04928"
/note="TcpE family, protein motif:Pfam:PF12648.1"
/product="TcpE family [mating pair formation]"
/"ID=B40_04928
/gene="tcpE"""
/protein_id="gnl|SC|B40_04928"
/codon_start=1
CDS complement(29556..30059)
/color=4
/locus_tag="B40_04929"
/note="Antirestriction protein ArdA, protein
motif:Cdd:COG4734 protein motif:Pfam:PF07275.5"

/product="Antirestriction protein ArdA [anti-ristriction
modification]"
/"ID=B40_04929
/gene="ardA"""
/protein_id="gnl|SC|B40_04929"
/codon_start=1
CDS complement(30075..30572)
/color=4
/locus_tag="B40_04930"
/note="Antirestriction protein ArdA, protein
motif:Cdd:COG4734 protein motif:Pfam:PF07275.5"
/product="Antirestriction protein ArdA [anti-ristriction
modification]"
/"ID=B40_04930
/gene="ardA"""
/protein_id="gnl|SC|B40_04930"
/codon_start=1
CDS complement(30855..31871)
/ID=B40_04931
/color=1
/locus_tag="B40_04931"
/note="Hypothetical protein"
/product="hypothetical protein"
/protein_id="gnl|SC|B40_04931"
/codon_start=1
CDS complement(31942..32163)
/ID=B40_04932
/color=1
/locus_tag="B40_04932"
/note="Hypothetical protein"
/product="hypothetical protein"
/protein_id="gnl|SC|B40_04932"
/codon_start=1
CDS complement(32164..32304)
/color=1
/locus_tag="B40_04933"
/note="Protein of unknown function DUF3789, protein
motif:Pfam:PF12664.1"
/product="Protein of unknown function DUF3789"
/"ID=B40_04933
/protein_id="gnl|SC|B40_04933""""
/codon_start=1
CDS complement(32308..33429)
/color=4

/locus_tag="B40_04934"
/note="Putative replication initiation factor Tn916-like CTn1-Orf26, protein motif:Pfam:PF02486.13"
/product="Putative replication initiation factor Tn916-like  CTn1-Orf26  Replication initiation factor domain-containing protein [relaxase]"
/"ID=B40_04934
/protein_id="gnl|SC|B40_04934""""
/codon_start=1
CDS complement(33709..35103)
/color=4
/locus_tag="B40_04935"
/note="DNA translocase FtsK DNA segregation ATPase FtsK/SpoIIIE and related proteins, similar to AA sequence:UniProtKB:P64165 protein motif:Cdd:COG1674 protein motif:TIGRFAMs:TIGR03924 protein motif:Pfam:PF01580.12"
/product="DNA translocase FtsK DNA segregation ATPase FtsK/SpoIIIE and related proteins [ATPase coupling protein]"
/"ID=B40_04935
/gene="ftsK""""
/protein_id="gnl|SC|B40_04935"
/codon_start=1
CDS complement(35214..35780)
/ID=B40_04936
/color=1
/locus_tag="B40_04936"
/note="Hypothetical protein"
/product="YcxB family protein [transmembrane protein]"
/protein_id="gnl|SC|B40_04936"
/codon_start=1
CDS complement(35777..36319)
/color=1
/locus_tag="B40_04937"
/note="Protein of unknown function  DUF3795, protein motif:Pfam:PF12675.1"
/product="Protein of unknown function  DUF3795"
/"ID=B40_04937
/protein_id="gnl|SC|B40_04937""""
/codon_start=1
CDS complement(36380..36817)
/ID=B40_04938
/color=1

/locus_tag="B40_04938"
/note="Hypothetical protein"
/product="hypothetical protein"
/protein_id="gnl|SC|B40_04938"
/codon_start=1
CDS complement(36882..37202)
/color=7
/locus_tag="B40_04939"
/note="Regulator of competence-specific genes TfoX N-terminal domain, protein motif:Cdd:COG3070 protein motif:Pfam:PF04993.7"
/product="Regulator of competence-specific genes TfoX N-terminal domain [transcriptional regulator]"
/"ID=B40_04939
/protein_id="gnl|SC|B40_04939"""
/codon_start=1
CDS 37448..37732
/ID=B40_04940
/color=9
/locus_tag="B40_04940"
/note="Hypothetical protein"
/product="Transposase [DDE recombinase]"
/protein_id="gnl|SC|B40_04940"
/codon_start=1
CDS 37729..38643
/color=9
/locus_tag="B40_04941"
/note="IS3 family transposase, protein motif:Pfam:PF00665.20"
/product="IS3 family transposase [DDE recombinase]"
/"ID=B40_04941
/protein_id="gnl|SC|B40_04941"""
/codon_start=1
CDS complement(38662..39009)
/color=1
/locus_tag="B40_04942"
/note="Protein of unknown function DUF3788, protein motif:Pfam:PF12663.1"
/product="Protein of unknown function DUF3788"
/"ID=B40_04942
/protein_id="gnl|SC|B40_04942"""
/codon_start=1
CDS complement(39073..39531)
/color=8

/locus_tag="B40_04943"
/note="Glyoxalase, protein motif:CLUSTERS:PRK04101 protein motif:Pfam:PF12681.1"
/product="Glyoxalase [glyoxalase]"
/"ID=B40_04943
/protein_id="gnl|SC|B40_04943"""
/codon_start=1
CDS complement(39627..40013)
/color=1
/locus_tag="B40_04944"
/note="Protein of unknown function DUF961, protein motif:Pfam:PF06125.5"
/product="Protein of unknown function DUF961"
/"ID=B40_04944
/protein_id="gnl|SC|B40_04944"""
/codon_start=1
CDS complement(40027..40350)
/color=1
/locus_tag="B40_04945"
/note="Protein of unknown function DUF961, protein motif:Pfam:PF06125.5"
/product="Protein of unknown function DUF961"
/"ID=B40_04945
/protein_id="gnl|SC|B40_04945"""
/codon_start=1
CDS complement(40557..43607)
/color=1
/locus_tag="B40_04946"
/note="Predicted outer membrane protein LPXTG cell wall anchor domain, protein motif:Cdd:COG4932 protein motif:Pfam:PF05738.7"
/product="Predicted outer membrane protein LPXTG cell wall anchor domain [membrane protein]"
/"ID=B40_04946
/protein_id="gnl|SC|B40_04946"""
/codon_start=1
CDS complement(43851..44219)
/ID=B40_04947
/color=1
/locus_tag="B40_04947"
/note="Hypothetical protein"
/product="hypothetical protein"
/protein_id="gnl|SC|B40_04947"
/codon_start=1

CDS complement(44462..45010)
/ID=B40_04948
/color=1
/locus_tag="B40_04948"
/note="Hypothetical protein"
/product="hypothetical protein"
/protein_id="gnl|SC|B40_04948"
/codon_start=1
CDS complement(44997..45311)
/ID=B40_04949
/color=1
/locus_tag="B40_04949"
/note="Hypothetical protein"
/product="hypothetical protein"
/protein_id="gnl|SC|B40_04949"
/codon_start=1
CDS complement(45326..49063)
/ID=B40_04950
/color=1
/locus_tag="B40_04950"
/note="Hypothetical protein"
/product="hypothetical protein"
/protein_id="gnl|SC|B40_04950"
/codon_start=1
CDS complement(49350..50210)
/color=9
/locus_tag="B40_04951"
/note="Tyrosine recombinase, similar to AA
sequence:UniProtKB:P0A8P8 protein motif:CLUSTERS:PRK00236
protein motif:Cdd:COG4974 protein motif:TIGRFAMs:TIGR02225
protein motif:Pfam:PF00589.16"
/product="Tyrosine recombinase [recombinase]"
/"ID=B40_04951
/protein_id="gnl|SC|B40_04951"""
/codon_start=1
CDS complement(50454..51641)
/color=8
/locus_tag="B40_04952"
/eC_number=1.1.99.28
/note="Glucose--fructose oxidoreductase precursor, similar
to AA sequence:UniProtKB:Q07982 protein
motif:CLUSTERS:PRK11579 protein motif:Pfam:PF01408.16"
/product="Glucose--fructose oxidoreductase precursor
[oxidoreductaes]"

/"ID=B40_04952
/gene="gfo"""
/protein_id="gnl|SC|B40_04952"
/codon_start=1
CDS complement(51655..52845)
/color=1
/locus_tag="B40_04953"
/note="Protein of unknown function DUF2961, protein motif:Pfam:PF11175.2"
/product="Protein of unknown function DUF2961"
/"ID=B40_04953
/protein_id="gnl|SC|B40_04953"""
/codon_start=1
CDS complement(52865..53683)
/color=8
/locus_tag="B40_04954"
/note="Inner membrane ABC transporter permease protein ycjP glycerol-3-phosphate transporter membrane protein, similar to AA sequence:UniProtKB:P77716 protein motif:CLUSTERS:PRK10973 protein motif:Cdd:COG3833 protein motif:Pfam:PF00528.16"
/product="Inner membrane ABC transporter permease protein ycjP glycerol-3-phosphate transporter membrane protein [transporter]"
/"ID=B40_04954
/gene="ycjP"""
/protein_id="gnl|SC|B40_04954"
/codon_start=1
CDS complement(53692..54573)
/color=8
/locus_tag="B40_04955"
/note="sn-glycerol-3-phosphate transport system permease protein ugpA, similar to AA sequence:UniProtKB:P10905 protein motif:CLUSTERS:PRK10561 protein motif:Cdd:COG4209 protein motif:TIGRFAMs:TIGR01581 protein motif:Pfam:PF00528.16"
/product="sn-glycerol-3-phosphate transport system permease protein ugpA [transporter]"
/"ID=B40_04955
/gene="ugpA"""
/protein_id="gnl|SC|B40_04955"
/codon_start=1
CDS complement(54616..55935)
/color=8

/locus_tag="B40_04956"
/note="Maltodextrin-binding protein maltose ABC
transporter periplasmic protein, similar to AA
sequence:UniProtKB:P0AEX9 protein motif:CLUSTERS:PRK09474
protein motif:Cdd:COG2182 protein motif:TIGRFAMs:TIGR03850
protein motif:Pfam:PF01547.19"
/product="Maltodextrin-binding protein maltose ABC
transporter periplasmic protein [transporter]"
/"ID=B40_04956
/gene="malE"""
/protein_id="gnl|SC|B40_04956"
/codon_start=1
CDS complement(56041..57960)
/color=8
/locus_tag="B40_04957"
/note="glycoside hydrolase family 127 protein, protein
motif:Cdd:COG3533 protein motif:Pfam:PF07944.6"
/product="glycoside hydrolase family 127 protein
[hydrolase]"
/"ID=B40_04957
/protein_id="gnl|SC|B40_04957"""
/codon_start=1
CDS 58135..58569
/ID=B40_04958
/color=1
/locus_tag="B40_04958"
/note="Hypothetical protein"
/product="hypothetical protein"
/protein_id="gnl|SC|B40_04958"
/codon_start=1
CDS 58950..59882
/color=8
/locus_tag="B40_04959"
/eC_number=2.7.13.3
/note="Probable sensor-like histidine kinase YehU, similar
to AA sequence:UniProtKB:P0AD14 protein motif:Cdd:COG2972
protein motif:Pfam:PF06580.7"
/product="Probable sensor-like histidine kinase YehU
[kinase  two-component system]"
/"ID=B40_04959
/gene="yehU"""
/protein_id="gnl|SC|B40_04959"
/codon_start=1
CDS 59892..61421

/color=7
/locus_tag="B40_04960"
/note="Response regulator, similar to AA
sequence:UniProtKB:P40408 protein motif:CLUSTERS:PRK10572
protein motif:Cdd:COG3449 protein motif:TIGRFAMs:TIGR04094
protein motif:Pfam:PF00072.18"
/product="Response regulator [transcriptional regulator
two-component system]"
/"ID=B40_04960
/gene="yehT"""
/protein_id="gnl|SC|B40_04960"
/codon_start=1
BASE COUNT 17394 a 13503 c 10216 g 20351 t
ORIGIN
1 ccgtattttc tacgccgtcg atgtgtgtcg ttactactat cggaacacat tttatcaacc
61 tcgttgctat catgtgaaca caattcttca gttctatgtc gtcctcttgc attttttctt
121 tgtggtttaa atttagcacc atccacgtta ttcttcaagc ccgttttata aacaaatgtc
181 aattgtgcag ggtcaacatt tccattttca tcaagtccac ctacaattac tttttcaaca
241 atgctttcaa acacatagcg gtcgaattta tcaagaactt cattctgttc caaagttttc
301 ttaaactctt tcagacgttt ttttatgtct ttttcattat cagaagtttc ttgtaatttt
361 tttctttctt caacaagctg ttcttgtttg cttactaaat cagcatattt actttcgtag
421 gtttccttat caataatttc ttccagacgc atatccacaa gtttactttt tttcttctct
481 aatgcgccta tctctttttc tgcttttgca agttgttttg acaccacact actactaagc
541 gtatcgtcca ttctctgcaa aaattcttcc agaacatctt tattatcatc acacaataaa
601 cgataacttt ctacaaacgc ttcttcaata gctgtttctg gaatgccttt gctatgctta
661 cagaattttt ttcctttctt cgtagccgtt acgcattgcc atataacctt gctgtactgt
721 gaactgctgt gccaattacg ccttgttaaa gtgccaccgc agaaaccaca ttccagcata
781 caactgaaag catatttcct actatatttt tctctctttg tgctaccctc tccaatcttt
841 cctctattaa catttcttct ggcaagaatt tcttgtgctt tttcaaaaac ttcttcgcta
901 ataatcggtt cgtgatgatt acgaatataa aatttatctt cctctccgaa attctccaat
961 cttcgctttg aaatcgggtc tactgtaaag gtttttccta gcaataaatc gcctttatat
1021 ttctcgttct tgataatacc aatgactgtt gattgtaccc atgtagaact tccgtacttt
1081 gttttatatc ccaaattttc aagctcatgt gcaataacag aaccgccagc accctcaata
1141 taccgattaa aaatataacg tacaatttct gcttcttctt cattcacggt aattgttttc
1201 gtgtctttgt gataatcata tccaagacaa ccttgaaaac caaccagctc tccccgtttc
1261 attttcattt ttaaaccttt ttttacatta gcagatatat tttcaacttc ctgctgtgcc
1321 acagaactaa gtacaactaa aagcaattcc ccatccatag ttaaagtgtt gattttctca
1381 tcttcaaaat atactgctat attgcgttcc tttaacatac gaacatactt taaagtatct
1441 agcgtatttc tggcaaatct ggaaatagac tttgtaataa ccatatcaat ttccccattc
1501 atacaatcat ttatcatacg ctggaaatct tctctttttg ccacttgtgt acctgtgatt
1561 gcttcatcag catatatgtc cgcaagtacc cattctttat tcttcttaat catatctgta
1621 taataggcta cctgcgactt ataactgttt agctggtctg cactatctgt actaaccctg
1681 caatatgcac caacacgcat aagttcaatc gtttttcccc ttgtacgagt ttgtaaatca
1741 tttcttgctt tgattacttc tacttctttc acagaaccac tccttttata tttgcatttt

1801 gtattccttt tgtagtgtta ttatacaata ttcaaatata caattcaagc ggttatatct
1861 gaaacaatat tataatcttg cataagtttt gattttacta aagcatattc tttttcagaa
1921 atcattttac gggcaagaag ttggttcaac atggcaatct gcatactata tcgtattagt
1981 tttttctcca tataaattcc cccttgcttg taatttccaa gcagacagac ggctttggaa
2041 agctccgtta gtatctcaac tattcccatt cttgtgggca gaaccgcacc ttgcggacgt
2101 atcattattc tgtgagcata ggtcatggca aaacgacttt tccaacagtc gctctcggat
2161 cactgtgtgg gtcgctcgct ttcttttgga aaggtcatgg cgtacagttc ccgtggctcg
2221 ctatctcaca aacgtgtctg tctgctttat tcagttttca aagagctgtg tcgaaaacat
2281 attgctttca ttctttaagc agtagcagag caggaaagag ggggaattgg acaaatcatg
2341 ctctgctggc aactgctgat tattcctcta tctcaaattc acaaatcatc ttcaatagag
2401 cttctctgat acgtcctttt aattccatat ctacgacaac acagacattt ccgtattcat
2461 catagaacgg acgtaaacaa cattttgata tgtacggttc atagaactgc aatagcttct
2521 caatagaatt tatattgccg tccacagccg acgaaataag ataaaatgac gggtgtgttt
2581 tattcatttt tcgtgttctc ctttaacttt tttccgattt tatcaagtga acttttccta
2641 atacgaaatg atgttgaacg gtcaattttc atcttgtcac cgatttctgc atcgctcatg
2701 ccaagaaaga aatacatcaa tacaatgttt ctgtcttttt ctgaaagaac tttgagagcg
2761 tccgctaatt catcatcaag tacacggatt tcaataccac atacatcaaa aactatgtag
2821 tcgctttcat actcgtccca gacagccagc ttttcaacaa cgatttctgg aagttcacag
2881 aatgagattt ccttattcct gcgtcgcctt aattcctttc ggtagtttcg tacaatgttc
2941 ttgactacac gttttagctt gcagtcgaat tgacattgta ctgtcttttg gaagtccgac
3001 ggtttcataa ccacaccccc tttccgttat gacgtgaaaa gtgtttatcc ctctttccgc
3061 accatatctg tacggcaagg tgtgatttgt tgcaggattt tagaaaaaag ttaaaaatat
3121 tttcagcaat aaaaaaaccc gcacaaaaca tagtgtttgt gcgagctaaa gctctggcat
3181 attcagttgt aattgtacgt attgtactgc taaattgacc ttgatatttt tgtaaatttc
3241 ttaaaatatt ccataatgca agccccctca aagcaatgcg ttaaatatct aaaatcttgg
3301 ttttctctaa ttatacggga cacttgttgc aacaataata ttgttggtat tcaaaagtag
3361 gtgtgatata ttttcattag gaaattatac cttttagggc ataacaaata tcaataattc
3421 tcaatacatc ttgaaatggg ttattcttag aatttatatc tatcataagc cacttttgtc
3481 cattcccctc tctagtattg tgataaattc tctgcttttt ttccattgac attggataat
3541 tcattttcag catttcctgt ttccacagga atcacccata tttccatata actaatatct
3601 gcacttcctt tgtaatattt ttctgccatg aacggctttg taatcaattt atgatttgga
3661 agccatgttt caaacagata cttatttgct tgatttaaag cagtggtaac aagttcttca
3721 aaagattctg cctcaattct gcatacaaca tactcacctg ctggtaatgt gtgttgtacc
3781 atatttttag gacatacatc tattgatgaa gccatacctc ctacaaagta aaggaacgtt
3841 cctttatctg gacttgctga atgagataca ccaagctcaa tatcattgtt aaaaatggca
3901 ctttcggttt gcagtttctt tgcatgaaat tcacgccata gctgacctgg aacatcaatt
3961 cccgtacttt ctccaacagg aatttgttca gtaacattta ctgttttttc aaaaccaaaa
4021 tacatttccc ttgttcttaa tgttttcctt tgaatctcca aaacaatatt ccctacaatc
4081 aatggaacat tttcatcaat catgatataa cgtgaagcaa gttctggctt gtcaaatgta
4141 tttaacatag gaacattctt tctgtattca tctggagtta tatggtaaac ttctttaaat
4201 gctcttgtga aattagcatg gtctgaaaat ccatatttta aagctatgtc taatattctt
4261 tcctccgaat tattcaattc tgctattacc attgctaaac gacgaagttt gatatattct
4321 tgaactgatc tcttaactaa tcttttaaac aaacgttggt aataaaatgt agaaagacag
4381 gcaatttccg ccagtttctc tgtttcgatt ttctcactta aatgttgttc aatatagttt

4441 aatgacattt caatattttc atatgcgtac atatcagtac ctccttgtat aatttataat
4501 caaagtatac atgaaatttt tttgtatagc ttgtcgcaca atgccattat cttataagaa
4561 atcaaaagat ataaaaaaac aattttgtct atatggtaac ttttatttct ttggtacttt
4621 ttaatgagcc atatctaata aattcatcac atatctttta atgtttttcg tatcatagac
4681 cattggaact gtattgttca ttgcttcgtc aatcgtcgct tcaatatcag ctcgtacatc
4741 agcatacgat aaaccgccag cttttgcacc agctttcaat gctttttaat atctcgtttt
4801 ttaagtccta tcattagaac caactcccta aaataagtta atcttctttt tcaaaaatta
4861 aatctatctt tcgcatacag ccttttgcgt caatttctgt gggtacaaat cctgcgtagt
4921 gataccccgt ttttgcgtaa ctgtcaataa tttctttgtg tttatcagta caaaggaata
4981 caactccttt cgcattatag ttaatttcta catactcata ttttttcatt cttcaaaact
5041 cctttcaaat acaaatttat ctttggattt attgactttt acaaactgga atttatattt
5101 catacagact taatcaaatg gtgttcaaaa aaagtttcaa aagtatttgt agcaccacca
5161 tgttccacga tttttccatc tataatttta tcaatatcta ctcctgtgat ttccaaaacc
5221 ttatttgtag gtgttattcc gaaaaaatct ccaacatgag tccctttcat aataaattct
5281 gaaataacat aatcatctgc ttcgtattga cgaataattt tcatagtgta gtcgggatat
5341 gtttttttga cagcctcaag gtgttgcctc attccatcta tgccgataaa catctcgttt
5401 tcgccaacct tttacacgca atcctttgaa atatattgtg gaagctcatt aagcaagttt
5461 tctgaaacaa ccacctcata aaagtatttt ataatctctt tattgttcat ggtacacctc
5521 caaaataaat ttcgattttc ttaattctat aaactagaat ttatagaaac actactttat
5581 caacctgcct tggtttgttt ataaagtaac ccacctttgg caagtaatag caaaaatgca
5641 actactattg catatggttc tctcaccatc gccaaaaaca aaacagcaaa aacactgaat
5701 ccaacggaca atgaattaac taccctttttg cccttttcta atttgaaata gaccattagt
5761 atttttacaa ttccaacgat tgttaagccg ataaacaatg cccaatagat tgtaataatc
5821 aaaacatcat tatctacata atcaaagaga tttactgcat agatatatcc attgaccggt
5881 tttggataca gcggtagaaa aataagcatc aaagaaaata agtcaacaaa ccccaataat
5941 aagtcgcata tagtattgag attggatcta ttttcttttt ctgcaattgt aatcaactgt
6001 tctcctgata acagatcatc aatagacact gcaaaatatc ttgatatttc tttcaatgaa
6061 tctatgcttg gatatcctct tcttgattcc cattttgaaa tagcagttct ggaaacaaac
6121 aatgcctctg ccagttcttc ttgtgtcaat ccccggctct ttcttaattc ctgaagtttt
6181 tcatggaatt ccaaaacctg cacctccctt gcttattcta ttgattcttt gtcgatagca
6241 taactactgc tttataaata gtaaacaatc ccatacttaa aattactcca cctacaatat
6301 ccgtgaacca gtgtactcct gaaatcaatc ttccgataac cataaatact gaaaaagcaa
6361 ttactgtaat cgtaataatt cgtttagttg taacacaggt cagtctacgc tgaatctgtt
6421 cgattaaagt aggcatcaca cacaatatca ataaagtagt agaagatgga taagaagcct
6481 ccatcactcc ttcaattaag attggtctgt aattaattgg gataatttca aaaattgaat
6541 atgccagtat cacgatacca taatatacac ccaaaatgat gatatctgaa tccaccttaa
6601 aaagacttct tctcttaatt aactgaacca gtccaattcc tgcaaaaatc agacatacaa
6661 caattggtac aagccccatc cagtcggtaa tcgtataaag cgtcatatga acacctgtca
6721 aacgatgaaa ccagcagttg aatgttgcaa atccaatatt ggttccattc tgcccaagag
6781 gctgcacatc aacaatctga atcaaagctg tccatattac aaatgttgca aggagtatca
6841 ctccaaataa taaattcttt tttccgttct ttttcattat tctacgtcct ttcaaaagtt
6901 aatctgtctt ttcgacaatt taacgttatc aaaaaacgta aaataacaac agaatctctc
6961 tgtcacatgt atgatacgat tcgtatcatt ccttcaaata cttgttttcc tgattcatca
7021 gacctacaat ttctaattta tcaatttcgt atcttcttgt attcttctat tttcaaatca

7081 ttaccctcaa agataaaatt attgatgtca atcggtaaac atattagaac gcaaattata
7141 gtgtctggtt ctaaagaatt tatttttttt gtagaattat agcaataaaa ttcagataat
7201 ctccaccgcc agcttctatt gctttacggt catttattgc atattcatta ttttgcttta
7261 tccaatccga ccaaacttcc gaattacttt ccatttcata tattgaaatt aaatctgcct
7321 tcttacaatg actaattatg ttagaccaat atgttatatc atgtatataa tcaagttctt
7381 ctggtgtcca tgataacaat aactcttctg gaaggttgtc atggcaatct ttcttcatgc
7441 caggaactgt gatataaaca aacccacctt tcttaacaaa tggaagcaat ttttcatcta
7501 aatatatagg gtcacgtcca aaataattat atgaatctgt acttactaca gcatcaaaaa
7561 actctttctc aaatttaagt tcattggcat cagcttttac agatttaatt tgtgcagaag
7621 ataatcccat ttttttgaaa aattcttcat tttcttttgg ctcactccat aagtcaactg
7681 caaaaacaac aaatccatat tcttttgcaa gaaatacaga agttaaacct tgtccacttc
7741 caaggtcaca gacatagctg ccagacggga ttttatgatt ttgtaatagt tcctcttgta
7801 attttatagg attcggtccc ataatttttg tctgtaagtc gggtgtaaaa tattttttac
7861 tgcgaatata ttccattttc tcatctccat tattgtaata ttttaatagt taatatttgt
7921 gacttataac aatattcatt ttgagtttct ctgctgaaat taaaaagtta tgtttgcatt
7981 caggacagaa taaaggatta ttttactttt tcatttgtaa tatatttctt gacaaattga
8041 ttaacatgtg tggttaattg agattttaaa tcccaacaat tttcaactgc gtctgctaca
8101 cttatcaata gaaataatgg cgcataatat tcaatagcaa gctgttttgt atcacactct
8161 tgaagaacac ctttttttaat catttcatta aatatatctt ctatataact gataggacca
8221 tttacaagac atttttgata aagcatattc atttttaaat ttctatattg ctctagtatt
8281 aacattttgc gaaaattaga cgcaaattca tcagtcaccc aaaaatcgta ttgtgcaaaa
8341 gtaaatgttg ttagactttc aatagttgta tttctgtatg cattaggggt taccactatt
8401 gtgctttctg gtacttgata ttccttagct cgcttattat caatttgata catacgttct
8461 acgatacagt caaaaatatc ttgcttattt ttataatgtt tatataatgc tccttttgtc
8521 attcccaagg cttccgcaat catactaaca gaaacagcct cataaccatt cactgcaaat
8581 aagtgtagtg caatttttaa tatattttct tttgtgtcag acatttacaa cctccttctt
8641 tagtaaacga acgtttactt ctgtaaacca ttatagtaaa cgctcgttta cttgtcaagt
8701 agtataaaat gtagtgtgcc tttggttcat aggttgatct ctcgatcgct ggtgctttat
8761 gccttgcacc atatttccta catacatttc tctatgcaaa atactgttta ttggaaagaa
8821 ttgtactcaa agctttccat aacgttagcc cccttaaagc actttctagt atctaaaaac
8881 atcattctac aacataacag atttcaataa accttaatac atcttgtcgt cctcgctatc
8941 ttttggacta ttgttagaat ttctcctttg gttatacttc ttcataatta caattttaag
9001 taagccaacg acaatcaaaa ccacaacaat gataatccac cagtattgtt ccataaccta
9061 tttcccctt tccctttcaa tagtcttaaa tcatcttccc tgtcctatgc agaatagaga
9121 agttcattcg tattaacatg taccatacaa tgtttacatt gtggaaagtt gttttcaatc
9181 gtttgatgga ctccctcagc aattctatga gcgtcataaa gagctaaatt accatcagct
9241 aatatttcaa tatctacata cattttcgct ccaaacatgc gggtttttaa ttcatcaata
9301 ccatcaaccc cttgctgctg aagcacaact tccctaatct ttgtttcagt agcattgtca
9361 caactatggt ctaccatttt gttaatgctt tctttgaaaa tgtccaaagc agcttttaag
9421 atacaaccgc aaatcacaac gcttgcaata gggtcgaaaa tagcatagcc taaacgcact
9481 cctaaaatgc caattaagga accaaccgac gagagtgcat cagagcggtg atgccatgcg
9541 tcagccataa gtgcgtcaga atttgtatgc tttgcagcac ttcttgtaaa ccagtacatc
9601 cactctttta taacaatgga taagactgcc gccgtaagtg cgattccact agagacagaa
9661 agcgtattgt actgtcctga aaagattttc ttgatacctg aatatcctat tccagcaccc

9721 gttatagcca atgcaacaga gagaataatg gaagccacac attccatacg ttcatgccca
9781 taaggatgtt ctctatcagg ctgtttttct gatattttaa ttccaaccat cacaacgatt
9841 gtgcttaata catcagaagc cgagtgtaca gcgtccgaca gcatagcact tgactttcct
9901 atatatccag cagcaaattt aagtagcgat aaactcagat tgatgattag tgtaattttg
9961 gaaacctgaa tagccttatt catatttttg ttttgcagaa gtctaccttt ttccataata
10021 tgcctcctta tcctgccagc ttccgaaaca gacgttttgt aaggaagtag taccagacta
10081 aagctaccac caatcctatg gagaaataaa gaaccgcttc attctgcatg gctttcgcta
10141 cccaaaatga tggaaggaag ccaaccgcga actggtagta actatcaata aaaaaggggg
10201 ctggtattcc gagcaatgta agtgcagcta gttttgtaac tgccatgact cctattggtc
10261 tggggtcctc cgggcttgtt atgattttct caaaatcagc ctctgtttca gcaggatatt
10321 tttcaaatgc agtaaaaaca gtattggacc caaacagcac cgcatccgaa tgaattcccc
10381 cggcaacctg gtagtaaact tccggattat caaagccttt aatgcggcca tcaaggctaa
10441 tttgtgtgtg cataattatc tttggtttca tatttttcaa tcctcctttt tgatagtaga
10501 atatcataat acactgacag ctgtatgtca gcaggggata tgaaacctta caaaaaatac
10561 ataacagacc aatcgctttc tctaaatcaa gtttaggcag tgattcgtaa tttgtgccag
10621 gtgtttacgg gttatgcatt ttaattcatc aggttggact atttcaatgc ctgatccata
10681 ttcaatcagc tgcttgctca atatctcaaa ggaatctgtc tggaattcga tttccatttt
10741 acttcccaaa tcggtttctt caaggaagcc gagagaatac ctgttcttga tagaattgta
10801 tgtctcccct ttttcagctc taagaattac attatatagt tttttatatg aaagcatttg
10861 ttttaatatt tgttccaaag agccatcatg gctttgacag agctcttctt ctatgcaaat
10921 gcttttaatc cgatccacac gaaaattcct atattcattt cttagcctgc aaaaccgatt
10981 aagtaccagt tctgttcata aaaccccagt gatataggct ctatcattct gctttccggt
11041 tcttgccctc ctgatgcagg atattgaatt gatattaccc ttttattgca gattgcagtc
11101 tgaattgctg caataacgtt atccgccaat gtatttttct gtccagatgt gctgtacacg
11161 ttaatactgt tctccaactc ctgtgcatag tttttttcac catgcttcaa aatagatttg
11221 attttataca tagctgaatc aaaatccttt ataaatgatt catctccatg gcaatttaaa
11281 aatttccctg ctgtaatcaa tgcacccact tccgccgctg taaacattac cggcggaagt
11341 aggaagccct ctacaagaaa atatcctttt ccggcctcag atccaattgg tattccggct
11401 tcttcaagtg tccggatatc tctgtaaact gtccttaagc ttatattaaa tcgctgtgca
11461 atttctttgg caggaattat tttttttgat tgcagctgaa taagaataga ggttactctg
11521 ttaatcctgt tcaaatttta ttcctcctct gtgacttcca taaaaagtat atcctatgcg
11581 ttcttaagct agctgctgtt agtgcttctc attgcaccct ggttataaaa gccatactca
11641 caaacctctt caccaaaact ctttacatct ctaatataga actctatcta gcgaactttt
11701 agaaaagata taaaacatca gagtatggac agttgcggat gtacttcaga aaagattaga
11761 tgtctaaaaa gcttgtagtt aaagcttttt agacatctaa atctaggtac taaaacaatt
11821 catccagtaa aatataatat tttattttct cccaatcagg cttgatcccc agtaagtcaa
11881 aaaatagctc gacatactgt tcttccccga tatcctccct gatcgaccgg acgcagaagg
11941 caatgtcata ccacttgtcc gccctgccgc ttctcccaag atcaataaag ccacttactt
12001 tgccatcttt cacaaagatg ttgctgtctc ccaggtcgcc gtgggaaaag acaagttcct
12061 cttcgggctt ttccgtcttt aaaaaatcat acagctcgcg cggatcttta aatggagtgt
12121 cttcttccca gttttcgcaa tccacatcgg ccagatcgtt attcagtaag taatccaatt
12181 cggctaagcg gctgtctaag ctattcgtat agggacaatc cgatatgtcg atggagtgaa
12241 agagcctgat gcactccgca tacagctcga taatcttttc agggctttgt tcatcttcat
12301 actcttccga gcaaaggacg ccatcggcct cactcatgag cagattgctc cagccatcat

12361 gccgttcaaa gtgcaggacc tttggaacag gcagctttcc ttccagccat agcatcatgt
12421 ccttttcccg ttccacatca taggtggtcc ctttataccg gctgtccgtc atttttaaat
12481 ataggttttc attttctccc accagcttat ataccttagc aggagacatt ccttccgtat
12541 cttttacgca gcggtatttt tcgatcagtt ttttcaattc cggtgatatt ctcattttag
12601 ccatttatta tttccttcct cttttctaca gtatttaaag ataccccaag aagctaatta
12661 taacaagacg aactccaatt cactgttcct tgcattctaa aaccttaaat accagaaaac
12721 agctttttca aagttgtttt caaagttggc gtataacata gtatcgacgg agccgatttt
12781 gaaaccacaa ttatgataga atttacaagc tataaggtta ttgtcctggg tttcaagcat
12841 tagtccatgc aagtttttat gctttgccca ttctatagat atattgataa gcgcgctgcc
12901 tatgccttgc cccctgaaat ccttacatac ggcgatatct tctatataag cgtaccggtt
12961 ccaatttttt cgcagtttaa cttttccgac gcatttatcg tcttggtagt aaagatatat
13021 tatcttatca gtattgtcaa tatattcaag gcaatctgcc tcctcatcct cttcatcctc
13081 ttcgtcttgg tagcttttta aatatggcgc ttcatagagt aattctgtaa aggtccaatt
13141 ctcgttttca tacctcggta taatcttacc tatcacctca aatggttcgc tgggtttatc
13201 gatatctttc aggtgccctg ctttcatttc tgtaatcact gttcccgcct ctcttctata
13261 tcagcggcat atgtgctatc caggcagccg gttttaccag tgtatttttt atacatgtcc
13321 ctggtatatt ttgttatatt cctatcatac tccggatagg cataatgaag cctttccgcc
13381 acctcaccgg ataccgccct gaacaattga tggcatagaa ataatgcttc ccatatgttt
13441 tcataggaat ccatccggta ggtggacaaa agtttctccc acaaatcctc ggatatatac
13501 ctttcaataa acttatagtt cttgcctaca cttaatttaa agcctgtttc gatgccgacc
13561 ttccatgata tcattctcag cagctcatgg cgaacaatct gattaaaatg atcaatagca
13621 aataaaattt ccttacggca caatccttta ataacataag gtgttacatt ccaaaattca
13681 ttgcagcaat catcatactc ccttgcgctt ggctttctta catgataatc tatatcagtc
13741 ggaactatgt cccttttaat tctacaatct ttatcaatta gaacctttat taatttatcg
13801 ccctttaggt aattatctaa ctcttccaag ggcaataagg taagatcaat tttattgtaa
13861 tcatcaaata gcataagata ggaaaatccc ttttcttcag gtgggaataa ttccatatcc
13921 tccggctttt gcatcattat tatattccca aattgattaa gccagtcatc attagatata
13981 aacggttcta tatcacttac aaaatatgta atatcataat cctgaaattc atctttaggt
14041 atattaatat ttgcgcgtga cccctcaagg gtcacaattc gaatacgttc atcctgttct
14101 gctaaagaaa gtactaaatc catcatttct ttttctgatc tcatttacat actcctcgtt
14161 tatttttct atattattac atcttctttt ttgccgatac aatcagcatc attgggcgtc
14221 gcatttcatc cgccatcccc ggaatatcca tcatgttctc tggcggctgt ggctccacaa
14281 tctgatttat tataaaacta tttgaaagca gtgtatttag atatgtggtc agtgttctat
14341 gatattttgt aaccttttct tccaaaaaca tagctgtccg tttgccctca taataataat
14401 tgtccaccgg gaaatgcagt atttctcctt tttcgttata ataccagtct tgtgttccat
14461 gagcagtaaa aacaggatgt tcaactgtaa aaactaaatt gccaccagcc ttcagcatcc
14521 tatatatctt ttttattaaa ttctcatagt ctgctacata atgaaacgca agcgaactta
14581 gtattacatc aaagctctcc tctgggaaat ccacatcttc tatggcacag cattcatatt
14641 caatctgtgg aaaatgggtt tttccttttg ctacttcgag cattttatga gaaatatcaa
14701 cacctactac agaggaagca ccgttttcca tcgcatatat acagtgccat ccatagccgc
14761 atcctaaatc aagcacacgc ttcatctact tgttttttgc cttctgtacc gcttaaaacc
14821 gttagacttg aacgagtaaa gcccttatta tctgtttgtt tgaaagacca atcttgccat
14881 tctttgaaag aataacggta attgggatca aaaaattcta cattgtccgt tcttggtata
14941 cgagcaatcc caaaatgatt gcacgttaga tcaactggca aagactttcc aaaatattct

15001 cggatatttt gcgagattat tttggctgct ttgacagatt taaattctga ttttgaagtc
15061 acataaactg gcgtttctaa aacaaaatat gcttgataac ctttatcaga tttgataatt
15121 aacgtaggca taaaacctaa atcaatagct gttgttaaaa tatcgcttgc tgaaatagtt
15181 tcttttgccg tgtgaatatc aaaatcaata aagaaggtat tgatttgtct taaattgttt
15241 tcagaatgtc ctttcgtgta tgaacggttt tcgtctgcat acgtaccata acgataaacg
15301 tttggtgtcc aatgcgtaaa tgtatcttga ttttcgtgaa tcgcttcttc ggaagtcaga
15361 acaacgccac gtccgccaat catgcttttt tttgagcgat acgcaaaaat agcccctttg
15421 cttttacctg gcttggtagt gattgagcga attttactat ttttaaattt gtactttaac
15481 aagccgtcat gaagcacagt ttctacaaca aaagggatat tcattcagct gttctccttt
15541 cttacgaaaa ttaattagtt agaagctacg atcaaagttg aatcacaaca aaaaaggcaa
15601 tcaactaagt ttttcttaat tgattgcctg gtatcttctt aaagacttga aattccctca
15661 aaaacccgat ataatgggtt tacagatatt taagtatctg attaataaag taattaaata
15721 ctttaccaaa tttcgggtct cgacttcttt aattgattgg tggtaatcaa ttaaggctcg
15781 caacttattt tcttggcgga aaataagttg gtcgtggctc ttttttgta ttctttattc
15841 agttcgttgt ttcgttatat ctagtatacc gcttttaaaa aaaataagca acaatttcgt
15901 taaatatatt aacgacaatc attgcttctc ttcttctatt tgcaactcta cttgttctaa
15961 ttcattagga attaagtcaa taactaaggt tgagtttgtc aattctgcaa ttcgttccaa
16021 tgtttttaat gtcggatctg attttcctga ttcaatcaat gaataatttg atctactcat
16081 ttctaattct tgggagaatt tttcttttga tttggtacct ctcatctttt ttaagctttc
16141 tctaaatgct agttccaagt ttctccctcc tgtttatgta aatattttgc acgttttcca
16201 tttaattata tcgttacacc ttgtaaatca caagtgatta atcacaaatc acttgtgatt
16261 agtgattgtt attggtgttg cctagaacga ttctaagcac atgtttaatt ttaaagtggg
16321 caattatctt taaaatgtcc tacaatgatt caataaaagc atggctgcta gtgaaacttg
16381 cgaccatgct tttaaaaata cttaatgggg tcccgagcgc tttagctcct tggaagctgt
16441 cagtagtata cctaataatt tatctacatt ccctttagta acgtgtaact ttccaaattt
16501 acaaaagcga ctcatagaat tatttcctcc cgttaaataa tagataacta ttaaaaatag
16561 acaatacttg ctcataagta acggtactta aattgtttac tttggcgtgt ttcattgctt
16621 gatgaaactg atttttagta aacagttgac gatattctcg attgacccat tttgaaacaa
16681 agtacgtata tagcttccaa tatttatctg gaacatctgt ggtatggcgg gtaagtttta
16741 ttaagacact gtttactttt ggtttaggat gaaagcattc cgctggcagc ttaagcaatt
16801 gctgaatcga gacttgagtg tgcaagagca accctagtgt tcggtgaata tccaaggtac
16861 gcttgtagaa tccttcttca acaatcagat agatgtcaga cgcatggctt tcaaaaacca
16921 ctttttaat aatttgtgtg cttaaatggt aaggaatatt cccaacaatt ttatacctct
16981 gtttgttagg gaattgaaac tgtagaatat cttggtgaat taaagtgaca cgaatgttca
17041 gttttaattt ttctgacgat aagttgaata gatgactgtc taattcaata gacgttacct
17101 gtttacttat tttagccagt ttcgtcgtta aatgcccttt acctgttcca atttcgtaaa
17161 cggtatcggt ttcttttaaa ttcaattgtt ttattatttg gttgagtact ttttcactcg
17221 ttaaaaagtt ttgagaatat tttatatttt tgttcatgta atcactcctg aagtgattac
17281 atctataaat aaatacagaa gttaaacgat ttgtttgtaa ttttagttat ctgtttaaaa
17341 agtcataaga ttagtcactg gtaggaatta atctaacgta tttatttatc tgcgtaatca
17401 ctgtttttag tctgtttcaa aacagtagat gttttatcta cattacgcat ttggaatacc
17461 aacatgacga atccctcctt cttaattaca aatttttagc atctaattta acttcaattc
17521 ctattataca aaattttaag ataatgcact atcaacacac tcttaagttt gcttctaagt
17581 cttatttcca taactttagg gttaaccata cgcaagacca atcactctcg gacaatactc

17641 atgattttaa tgataaaatc catgttaaac ccatagataa gaaatacacc tgcaataacc
17701 gttacctgtt tgtgccaatt taaaaatgca ccactttttt atgattaaaa cggttgaaaa
17761 ctgtaccact aataactcac aatagagaga tgtcaccgtc aagttaaatg tacaaaataa
17821 cagcgaaatt tttaaatcta tttcttatcg atacaaattc ctcgtaggcg ctcgggaccc
17881 cttaattaga ttctcttttt aaaaatgaat gctgatccta ataccattaa tgaacctagg
17941 acagacaatg caatattttc tgtttctcct gtttttggta aaactttcat ttttccgtct
18001 gaacctttaa ccgttactgt tccgtctttt tgaacatctg cgccaatttc tttagcttca
18061 acttttttttg ttgttccgtc tgctaattga atgattggtt tactatcttc aactgcaaca
18121 atcacaacgc ctgtatctgt attaactggg ttttcgattg ggtttgtgtc tactggttta
18181 cttggttctg ttggctctgt cggcttgctt aggacactgt cccaacagac actttttaaa
18241 gtatgctgca ctaatcctta tttttcacag catagcgcaa tgacagacag ccatgttctg
18301 cattctctgt tccctgcaat tccaaattaa ctttttcaga taacagaagg ttatcaaata
18361 gttgtttttc tttattacct acaaaacatg gtgatatcac aagactgatt tcatcaataa
18421 gtccgttttc taataatttg ttggttaatc caccgccact gtctgtccgc atgtatttac
18481 acccatattg ctcatgcagg atctgaaagg ctttttcata gtccacatga tcatttcctg
18541 acacaatata tgggtaatgc ctttcctcca aataattcaa atattccttc ggagtggcag
18601 ttgataccag aatgacaatt tccttcagat atccaagatt tcttaagcaa tgaagacttc
18661 tcaagatacc tctgctgtca ggtataccct ctaatttatt gcttgataat gtgataatca
18721 taagtgaaac aataattgcc tgtactgagc cgagaagcgc caaacatatt gtcattccaa
18781 tggacaactt ttctaatgaa aaaagtaaca gtacaataaa agcgatgacc gccgaaataa
18841 ttgagggtac tcccaacctt gtaaagagat accccgactt accaagaggg gtaatcgaaa
18901 agtaccgtga cactttatcg tcgatttcct ctaatgtaat cattgcaaat gcaaagcaga
18961 gcaatacagg agccatgata gaaagtaata aatcaaagat cggatagtat gcttgcaaag
19021 aaaaatctgt tattctttca agaaagggaa taccgaattt aatcaaagct cccatgacaa
19081 aaggcgtaaa acaagctgca aacatcatag ggtcgctctt aatttgctta aatacctgta
19141 tggtactgga taatatcctt ttcatagctt tacacctccc aaacttctcc acatatgctt
19201 aactgtatgt cgcgcaacta tgtagaggat aatcaacgta gcaattacaa gtaccatatc
19261 aaacgatagt aaaatgcttt ttccagttat gaggttcatg caagctgtga acggaaaaaa
19321 gcggaatata tctggtaact ttacaaacag accaacgata ggtggaacaa agcaaacaat
19381 ttctatcgac ataatcacaa tgagaaattg gttcaagttt gaaatttttg tagccgcaat
19441 aatgccaagc atagtaaaga ttgctgatgt caaagctgta ccaactgcga tactaagtaa
19501 ctgattgcta ccggaaacga atcctaatat cattgcaatt accgtagata tggcaattaa
19561 cgcaactgtt tttgaaagta tgtactccga tacttttaca ggcgatacta ccatagcatt
19621 taacaccttt tgactttttt ccaagaggac aatcgctccc ataaaaaaca atcccattgc
19681 tgctgggtca gagtatatca tgatagaagc aatatccgtt ttccaatgtc ccgtcaaagc
19741 ggcaatccca cacacatata aaactgttaa tataaagtag ataaaataga aaccatattt
19801 ccattggaaa tgtatatcac cacaaattaa acgtcccaat ctcattgaag tgtcctccct
19861 gtaatatcaa caaaaatatc atttaatgta ggctcactac tatgaataga aagcaaacgg
19921 ttttcggaaa ttaaactttt taacaactta tcttctgtgg ttctatcaag cagacattct
19981 ccggttttct cgcctttatc atagtaggta tagtgtattt tggcagctcc ccttgacata
20041 atcaaattat gcgggctatc caatgcactg acttttccgg caacaataaa ggctacttgg
20101 tcacatagct ctgttgcgtc aaacatattg tgagtagtaa tgataatcgt ttttccgcgt
20161 ttcttttccg caagaataat atctttcatt aaacgattat ttgtagggtc aagaccgctt
20221 gtaggttcat caaggaataa tatatcaggg tcatggacta acgctttaat gaaattcaag

20281 cgagaacgca tacctttgga aaagtcagca acttttttgt ccccatcatt ttctaagcct
20341 accatatgca ataattcatc aatagaccga ggttgcttgt catatagaga agcaaaatag
20401 gcgagatttt gtcgtgctgt gaatttctcg tagcaggttg aaaattcaaa gtctactcca
20461 atattttcat aaaaccgatt tgtatgctcc cttatttctg tacctgcaac ttttacgctg
20521 cctttataac ttgtattaag cccagttaaa attttttgaa tagttgattt cccagcacca
20581 gagggtccta aaaaaccgaa aatttctcct ctcccaacat ggaaacttac attctcaaca
20641 aacggtttgt cagtatagct aaaatacagt ttttttacat caatcattat tctttcctcc
20701 tttaacttta tgacgtttat agatatatag tcatataaac tgtaaaaaat ttgtttggcc
20761 acttgcttgt caaataacca aacaaaatta ttactctgct actacttgct gaataatacc
20821 attcactaaa agattttgta tcatattcaa ttcttcgtcc tctttgaatt gttccaaaca
20881 ggataaaaca gaatacaggg attttatgat aaccgtatgc ggaaccaaaa atacaacccc
20941 taaaccttcc agtatctcca atatacgaat atcactgtac atatgattct caaaaatatc
21001 tgacgatatt tttctttgca gatagtccat taaagataaa tcaatctgtg aaagaatagg
21061 cgaagtaatc acccccgtta aacttagcat gataacgtcc tttgctaaat ctctatggtt
21121 cttatacttt ttcttttttaa cggttgtttc tatacttgta aataactctt tttgtaagcg
21181 aaaattgatt tccacatata gatgttcttt actgggataa aaagcataga atgaaccttt
21241 tgatatgttt acagcagctg ctaaatcatc aactgttact ttttttaagc cgtgtaatga
21301 aaacagcttt tctccctcta tcaaaagttt actgtttatg atttcttttt cttgttccgt
21361 aaatcgtggc aaactcataa cccccttatg actaaaatgt ttttatcgtc atgaaaagta
21421 taacatttac tgattatata atcaatatac tatccaagat atttatgcaa ggtagcttcc
21481 aatgcgttta ttgtagtgtc cgtgtcctca ttaaaaatca tgtgacgatt aagctctaaa
21541 gtaagtaatt cctttttcgg tacatgaatt aaatcaaata ctttgttggt ataatcaaag
21601 ctaaaaagcg gatcgccttt ggcagtaatt aaaataacag gacattgtat agccccattc
21661 tccaagcacg ataagtcagc attaaataag ctggcgacaa atcttaaagg ataatagggt
21721 aaaagcatag ggtcatttaa gcaatcttgt tttgcctgtt cactataaaa tactctgtca
21781 tagtctaaat aagcgtctcc tggaacctta tagttaggca gcaattttcc acctacacga
21841 aagcccaact gtatcagccc cataaaacgg tgtgaccatg cgggatattt tgttaagctc
21901 atagtttctt ttaatgtagt cagcatgata ttatgtggaa aaacggcttt taaccttgtg
21961 tccctacccg ccgccattgc agctaaaatc ccaccttgac tggaacccat gacagctatg
22021 ttttcaccaa aattttctat tccgtaagtc gttgcgtcat atacattatg cagcaaatct
22081 tccattgtat aatttttttt gattttaggg ctttttccgt gtgaaagata gtgaatgcct
22141 attatattaa atcctctttc actcaaacca catagaaaat tctcatacat gagagggtgt
22201 gacatagtac ccggcaaaaa aacaatggta ggactacttc tttctgtata ccagatagaa
22261 agtgttatcg gaacattatt tgaagaaata ccaacttctt tatacatacc acatacctcc
22321 tatatgactg atttggtttt atagtcatat aataacttat tgtaagcaga aagtcaatat
22381 ccataaatat atgatttaag ataaaattgt caagaaagat ttaattgctt aaaatacaat
22441 gaaatcgctt gttagtgcag cacatttgct acagaaaagg cagactaaat ccgaaaatag
22501 agcctgccaa caaaatatct tacataataa ttttccagtt gctatcctta tgtagcacaa
22561 gttcaaactg tgatacctgc attgcctttg tctgattgtc tatgaatttc acggcaacct
22621 tgactttcac attatccccg tcagccgtaa acacgggatt gactaattca gaatagaggt
22681 agtccctgcc gataggttca agcacattgc cagctacata gtaggaaagt tctttttccg
22741 ttgctgtcgg gtatagctta aagaatgttt ctaaaaatgc cgtagcgtca ttcacagtat
22801 cagcgtccac gcttgcgtct gcttcgtgtg cctttggttc ataatcggat ttctcaaccg
22861 ctggtgcaag tgtcgggttc tgaatgatta ccatatcccc gtcagcgtcc acatggactt

22921 ttacggtata ggttgctttg acattcgttg tctgttctcc ctcttttatc tgctgatcta
22981 cttcgtaggt agcgataaaa tcgtccgttc cagactgttc tatatcccaa atcagcacat
23041 ttgtaacaac ggagctggtc ggtatatcgg ttcttatggt atcaaggttt aagtcctgca
23101 actcctttgt cagatagttg ctgattgcct gcatccttgc ttcgatagct tccttgctgt
23161 tatcccatgt atagtaggac ttacagaaat ttttcacgaa attttcaatc ccgttggtgt
23221 catttagtct tagctgtatt gtttctgttt catgtaccgt gtgcatatca atagcggtaa
23281 agttcttata cacaccaaat gatacgctgg caataagtac cacccataat gcaatcacag
23341 atttcttatg tgtgcctacc ttgacagtac gcaccttttt ttcttttact ggtttttctt
23401 tctttttaaa catggtttcc aatcctttct actgttttat tcttccagca ccaatcaagt
23461 gctgttgcca atacgaatta ttaaggtcgg tatatccgat aggatcgcct gcatgataca
23521 tctgattatt tcctacatag ataccgacat gggttacata gcttcctgca ttgtaggtac
23581 tgtggaaaaa gaccaaatcg ccagcctttg cctgcgatag tgggatatgc tgtgttgcgt
23641 catactgcat ttgtgcggtt ctcggtaaag agataccagc cttgccataa caccactggg
23701 tcagtccgct acaatcaaaa gaagtgtttg ggttactgcc cccatacacg tatttccaac
23761 cttgatactt caacgcttca ttcattatcc tttgtgccag ttcgcccgat acttgattga
23821 caacaagata ctggttcaca agctccacat aaaacatatt tccatagcca taacgccagc
23881 ccccgttctt tgccacagcg atagggttcg tatagattac tttctgaccg cccgacttat
23941 cccttgcgaa gttttccgca agattgaatg tatgcttttt tccatttcct gccacatatc
24001 ccacatagcc accgccatag ttataggact gtatcacgac atttatatca ctgatacctt
24061 gatttttgca ggaagaaagc agggacgcaa aatatttaca cccctgcttg attgagcttt
24121 ctgtatctaa tgagttgggc ggtagtccca gactttcaga actttgcata acgtcctcgg
24181 ctgtaccgcc actttccact tgtatgattg ccagcaacac attgacatac tcggaaatgc
24241 cgttttctct ggcatacttt tctaccatag gctgatgttt caagacttcg gcagataggt
24301 tcataccaac gtaattagag gaataaccgt tgcccccgtc gtcgtcctcg gcactaatca
24361 gaaccccaaa gaataataca atagaaaaga ggataggaaa cagactgcca atgatagcga
24421 tatgtcttag tttcatttct tcttttgtcc tttcttgaat aaggttcgtt tgttcttttc
24481 ggtattctgc tgtgtatgct tctgaacggt tgttttcgta acacggtcag ccttgtaatt
24541 ctgtcttatg gtttcttgat tgacggtctt tgctgatgaa ccaccagaag atagcggacg
24601 ttccttgaca acatttgttt tgacttcatt attcttaacc gtagaagctg tggcagggcg
24661 tttgacttcc tgctgttttt ccacattgac cttagaagta gtcgctggtc tttcatgggg
24721 acgggtagct ccgcttgtcg ctgatccgtc cgttttctta cctgcctgcc ttgcttcttg
24781 tgccttttga agctccatac gcttgtcagc gatattctgt ctgtgtagct gttgtttctc
24841 gaaacgtcca gactggcggt tttcttgttc ctgcaccaca ccacgcttga agtcggacac
24901 actggacttt gccttttctt tggcagaatg aaccgcataa gcggtctgtg tcggcatatc
24961 tttgatattt tctttgacag cagttgcctt gtccttgacc ttattctttg tatcaaggac
25021 agctcccacg gctgaacctg ctttctgccc catgctggag cttgtatttc cacgattgtt
25081 ctttgttcca ttctggcgtg tagattttga accgacaacg acaccaccag ccaccgcacc
25141 agcgacacca ccagcactaa cagctcttgc aagcctatgt tccatacgtc tagccctatg
25201 tcgcatgaac agatacggtc tgcggaatat tctccgtccc atgctttgac tgtcgccagc
25261 gtttaaagag aacatactca ttaaatcgcc cagcttcatg tagatacctg caaaacacac
25321 tatctgcaag aacgccacca taaagaacgg gtagtccgtc gatatgttgt agaacatact
25381 ggaaatacta aaggctaccg ttacaatcag cgtaatacca gcccttgtca taatggtatt
25441 gaacacacgg acgattgcct gctttgccat actttcatag ctcggtatca tggaaagtag
25501 aaaactgata ggcaaaaaca ttgcgaaaat gataaaaagt atctggctaa ataacatcat

25561 gcccgtaagc aagaatacaa atatggttat tcctaagttg aagataagca ggaagaacac
25621 catacctaaa cggttcacga cctgcggtat cgtcagattg ttgttgtcat tgtcctcaat
25681 ctcggtcttg acgacatttt cccttatttc gccgtcctcg tcctctgggc ttgccgatac
25741 cagagcttcc acacggtcag aaccgatttc ctctatatcg gaattaccga attgaagcag
25801 tagccacggc tgttgtactt gtatggaaaa taggctgtcc cttattaagt ccacgctatc
25861 cttgcccttg ctgtcgctgt cgggaagcat gatttttgtt cccaagtcca gtgaagcggt
25921 gctgatgtct gatgaaaagt catttatttt cttgatgtag tcaggagcgt aggcaataaa
25981 ggaagcagat aaaagaaaca ccactacaaa gttgataacg gcgtgaagtg ccttgctggt
26041 ttcccttttt atcagtcccg tataggcaac ataaataccc actactaaga taatcagaag
26101 tagaaatccc acataaaatc cagaagtaga aaagccgttt tcggtaaccc ctgcaagggt
26161 ctgtatgctt ttgccgatac tgtctgccat atcgttgata aaatccagct tgtaggcttc
26221 ctgcaccaca taccccgtag cattgcttaa atacaagctg atagtccaga cgaaatttgt
26281 aatgcagtaa agcccatatt gcacactttt tccaatcccg tccagccagt tccacggaag
26341 ccacgaccag ctattatcca catagaaatc aagctgataa ttcgacagcg gatattttga
26401 ataaaggttt tcggtattga cagtgtcgtc cacaagtccc gttgcgtgta ccaccgttcc
26461 caagagtgaa agcaggataa gggacagtgc cacgacgaac agagccattt tgagaacgtg
26521 aagtatcttt tttcttgtga acgcactttt tatcctttct ttcatcactc cacctcattt
26581 ctctgtatcg gcggtctggt atcaaaggcg tgcagtaatt cttcaaagac tgggtgtatc
26641 tgcaccacgc ccacacgtcc gtataaatct tgtaataggc attgcccgtt ttccaaatca
26701 cgaagtcgct tctggttgtt ctcgtcctct ttgtcaatgc tgaaaaattc aagggtctgc
26761 ttaatctcgt tgatgtcggt actacggaaa gcgaatttta aaccgatatt gtttttcaga
26821 ctttcctttg acacgtcccc cgaagactgt gttacaaaat atacgcctgc ctgcatggca
26881 cgaccagcac gaaccagctt gttagagagc gtttcgcctt gtgccacatt taagaacgcc
26941 cacgcttcgt caaggtctac aatcttgaaa atacttctgt ccgaatggat aaaatctaag
27001 gcaaaggtac taattacaat cagcatggaa acagacaata gttcaatggt cgtgtattcc
27061 tcaaaggtgg tgtctttatc gggtagcact aagtcagcta cttgaatgat attgagctgg
27121 ttatctaaac tgattgcatt ttccaccgta ccgtctgaaa acagcagatg tgcaaaatcg
27181 tagtccgtga agctgtcgat atggtcggca atgttgcgtg agataggcgt atcttccttt
27241 cttaattcct caatcacatg gagcaatcct cgctggtcgc tttgggtaac ggcacgcacc
27301 gcctttctaa gtacggggaa cttctcgccg tccctcgatg agatccccgt aaggaatgcg
27361 agaatgtcga ttgccagact ttcagcgtct ttgacattct tcataatcac gaatgggtca
27421 agaagtcctg cgttttctct gtcgctggta aggtttacga tattgatttc atgggcgatt
27481 tctggcaagg tttctttcca gttgccacgt tcacttttcg ggtctaagat gaccgcctga
27541 ccgccaaaca gcacagaata ataaacaagc agattgttac agaatgattt tccaccgcca
27601 agacttccaa caaaggcaga agccagagca tttgtaaccg ttccttttac accttgactg
27661 gcaagggacg gctgtaaata tacattgcgt cccgtatcaa cggaatagcc tatatagata
27721 cccgtatttt cccctaactg ctgtgtcgct ccaaagccaa gtcctgctaa gaagtcggat
27781 tttacatact gcacatagtc attgatgtat cgcttgctgg ctggtaaaaa ttcagaatga
27841 agccccaaca tatcgccagc aggacgtacc agcttgacgt tcaaatcgtc gtagaagtcc
27901 ttgacttcat cacagcgtcg tttcagttcg tcaaggtcgg gagcagatac acggataaca
27961 taagagagct tatacatact ttctttggac tggtctaaat ccatttccag ctcgtccaca
28021 ttgtctagtg cgtccactac atttgaactg gtttcactgc cagactgata ggcgtgattg
28081 tctaaatctt ttaactcttt ctttttattt cttactgtcg ataacgcttt tcggttctcc
28141 acaatttcta cattcatgct ggtatcaacg ggaaaggtaa actgctgttg ctgaaaatag

28201 aagatttcgc tggacggaaa atcaagctcg ccgacaatcg cattgacggt aaagcaggaa
28261 acgtagcttt cgctgtcctc atgttccaag cgtaaatgtc gctggttttc ttccacaagg
28321 caacgggtcg gacgtatcaa gtcataatgc tttatcagcg tttccttttt cagtttcttc
28381 tttggtagcg aatactcata atcttcatag gcgacaccgt ccctgccgta gatgtgttca
28441 atcagatagc cgaagtcgtt tttatctaag cgtcttactt tgaagcgtcg gctgattttg
28501 ttttctaaca gcttttccat tttcatgtaa cggttgattt catcattcgg catagataca
28561 aagtcattca tcagcgtatg gttcacttca tggataaact ccttgaatgt cagccacgcc
28621 gattttttaa tgttcttcaa attaaattgt tcttccgtta ccatgagctt gaagccaaga
28681 aaaaagcggt agtctacttg attatcccca atcatgctta ctaacgcttc ggtctgctcg
28741 tctatcttct gataggcaac gtctttcaat cgccccatta ccagcttttt tgactgttct
28801 tgtatgcttt ggattgagct ttccgtggca atctgcaagg cgtgtatctt gccctcacgg
28861 gactgtgcga taagctgtcg gaagctgtca tgcacaatga atttctgttc agcagataga
28921 aaactgtaat tgtaaggtat cagctcatag taggcgaata cctcattgtc cttgttccag
28981 acaaggttat tgtcgatata tttaatcggg aacatagctc acactcctta ctgccgtgat
29041 agtttcatgt aatacctgct tatccagttt cacggctttt cctgcataag tggtctttgg
29101 tcgaagcccg taggtaatca tggactttag aaaactgtaa ggcttcttgc cgtcaaaggt
29161 cttttgcgac atgaaccatg taagagctac gggaataccg aaatatttca gaaatgcccc
29221 ctcaatcatg gataagggcg gtaagtcccc aaacagaatg atgataaact cggttatcac
29281 aaaccatgta atctgtgtaa aggtaacggg aaagggcagg ttaaaatcat tgattgcata
29341 caagactttt tccacgttcc agatacccgt atagctttta atctttttca tgttttcgtt
29401 tcaactcctt tctttattcc tgccgtgttc ttcacggcat agtggtatct ctttatgaga
29461 tttcacataa tagaaaagga cagccgtttt tcagactgtc cccaatagaa aaagctgtca
29521 gaagcaacaa tgcttgtcgc tgatcttcca gcttcttacc acagtatttc acacataccg
29581 tagcttgttt caataaaagt tccctcaaag gataaatccc tgccgtatgc ttcatagtcg
29641 atatagttct gtaactgtgg ggggatttcc cctaacatct gcatttcatc tatgaagtaa
29701 taagcaacgt cggtcatggt ttcacaatcg gaatagcaat acacgtcgtc cttatggtcg
29761 tacacttctt ccagactgcc gtaatgagag ataaactcgt ccagctcgtc cgtgatgtag
29821 tcgggtaatt cttcaatcat ttcaaacatt tcattgagtt cttcaatcga gatatattcg
29881 ccaatctcaa tgggaaagtt ttccgtatca tgtacggcgt attcttcgta tcgttcattc
29941 aagccgatac gctcggcaac atcttcttca tcaatcggaa aagagaacca gtcgccgacc
30001 agatacccct cattgtattt cccaagatta gcgatatata cacgcatatc ctctaccata
30061 gcaacacccc atttctacgg cagttcatag ataccgtggt ctgtttctat aaattttcca
30121 ttatcatcaa gatactgtcc gtaggcatcg taatcaaagt aacgggtcag gcgttcctca
30181 atggcagaaa aagcagggtc attgtccaac aaatgccttg ccacatcagc catgttttta
30241 caccatgaat aatgtatcac gtcatggcgg tgctggtgca gttcgtcaag gctggtaaag
30301 taacacatca gttccccgta gtcctcttgt aaatcaagtg gcagactttc aaaggtatgg
30361 taaaggtcat tcagcttttc aatagtcgta tcttcgccga cttcatcagc aaaagacagt
30421 tccatttctg tgatacggta atcttcactg tcgataggaa cgcctatctg ttcttccagt
30481 tcttcctcgt caatgggaag tgtgaaccat gcggaacgga tttcttcaga agtgtgtagt
30541 cggctttcga tacagacacg catttcattc ataggttttc acgtcctttc tttcatgctt
30601 cagctatgct agtgtaaggg acaccaaaga catagcctgc aaaatcttct aattgtttct
30661 gcggataatc ggcaaggttc agttgcttca tttcctgcca caccatacgt ttatgatgtg
30721 gtaaatctga aatgtaggta caaccaatcc ttgcctgcat atccgaaaat atatctttca
30781 agcaatcact ctaatcttaa atttgagtat agaaaaagcg gttaatctcc aatggagtaa

30841 ccgcttaatt atgtttattc ataatctcca aatttttcca aatcattgtt ccaaagaggt
30901 ggagcttctg gaaaaacttc atcatgcgtt ttaaaatata tttctcgaac gacatcaatc
30961 caattattta tattttttat tgcgacaatg ccatctccat aaggaaaaat atttaaaaac
31021 atacttgtaa tctgtgctaa agatatggct gaatgttctg caggtgttgt aattccgtaa
31081 tcacttctgc caacgggaat aatattttct gtcttcatat tacctaaacg tgcaaatgtt
31141 ccctgtggtg aagcatgaac agttttattc gctaagtcat attgttttct ccagactttg
31201 gtattcatat cgcaatgagt ttgaatatca ttaaaactta tatatctttt tttagaagaa
31261 aacaaatcac ttgctcgtgc ccattcatat ctatcgtccg tttcgctagc ttcaataaat
31321 gcttttgcaa cattttctcc attttctgtt ataaatgaag ctattatgga gagttcatac
31381 atagaacgcc acctagcata tgcaccatct gcaaatccat ttttcataag ggtaataatt
31441 tcaagaaatt cttgtaaagc tcgtccatgg atatgtagca tagccaaaaa agtgttatta
31501 ttttgaattt tttcttcaat acttaaatcg ttgacatact ggacatagga ctcagcaact
31561 tctaatgtca taatatacat tgcttctgaa gcgacaaagc ctctatacca tttttgttct
31621 tgtctgacta taaattcctc ttctcctgcc ctaaaagcca tgacctcttc atacattgtt
31681 tttttcatat attgagtagt atcatcagcc attttgaaaa gcatttcttt aaataagtca
31741 ctcaaatcaa ttttttctaa tttttctatg acttctttag cttcagcttc cgtttccgct
31801 tcttcaattc tatttttttac agttttcata aacgtatctt ttaggaaaat tgaagcatta
31861 tcttttgcca tatttttaac ccacccttcc aatatcttat atgtaaatta taacattgtt
31921 aaagctggtt agcactattt attatgctcc gataatctta ttgaacagct ctaagagtac
31981 atctttgaca ccgcctgcat tgaataccaa tccaacagca actaaggcaa ccaccaagaa
32041 gccgataagt tttgaaaact cacgtttaaa ccctaagtaa ataccaataa ccacaatcgc
32101 cataagcact aagctctggg cgttgcttaa aaaccatgta tataaatttt gcccgaaatt
32161 catttatttt tcctcgcttt cctttaattc aataatgtgt cggtcagctt cctttccgac
32221 actcataata cacatcagaa ccacgccgat acccatgcca agtgatacca gcagtaagtc
32281 cttaatcaat tcccatatca tcatcaatca ctccttttct tgttttatca gttcatccac
32341 cgaagccgtc tgttgcttca cgatctgctt gtgcttgtcc gtcagttttg cctgctgttc
32401 aatagttttg agatagtcgg tctgattgcc tgcgtctatc tgtttcagca ttttcaaagt
32461 cggggcaacc tgcctttgta gccagttcag cgttctttgc agagtataag gctctggctt
32521 tgtcgtcagc ttcacgggcg gtcgcccctc accaatgaac caagaccagc gtgcatttgt
32581 tttccagtca ctttttcgtt tgcctacttc cttatcggca aagcggatat actggttgat
32641 aatagaaaat gccgtcagct ctgcgtcgta gtgggtcagc agttctctca cggcatagta
32701 ggcacgttca tttttcagac gtatctcaaa tcggttttta atggggactt cctcaatcgg
32761 tattcccaat tttgaggatt gttcataatt cttttcatag acacagaagt agacttcact
32821 ggtaaatgaa ccaatataaa gcgtgtgtcc catgccgtcc ttgttgactt cgttgtgctt
32881 taccagctcc ccagaaccat agttcttgaa agaacgaaat ttagaaatac attcctcgtt
32941 cttgcatttc tctatcagtt ctggaatatc cagtaagccc gttctgtcgt tgatagctag
33001 gtcgatacgc ttcatcacac cgccattaac taatgcgtcc atgaggaagt cataccagct
33061 tcgctcctgt gccagcaggt agctttcaaa ctggcgacag cctttgcctt tcagttccag
33121 caggacacct ttttccttgt cttgtgaagt ataaacgaac acgtccccga tatagtaatg
33181 ctcggtgtac ttgtaatgcc cgaaatcttc gtgaagcata tagtctacat tcagctttag
33241 aatgtccttg ataacgtgct gtatatccag tgtgggaaat cggatacgca cataatcaaa
33301 gagcaggaac atgggagctt cgggattgaa tttttccaca gcttccagta gtttgtgctt
33361 catgtcctcg gtcagtgcta ccttgccagc ttcgatacgg cttagatgtt cacggctgat
33421 accagccatg attgcaagcc ttgtctgcga tacgccataa tcattgcgtt ttgctttcaa

33481 ggcgttgaga aaatcagtat cattcagttg tcataccccc aatcgtgata tttttgtgat
33541 tttgtgataa tttcaagcga ttatcacaaa tggctgaaaa tagagaaaac cctaataact
33601 agcggacttt caaaggtttt tgaaccagtt tccagcgtaa ttgtgcccct ctgttagata
33661 atgaggggct tgtatctgct ggcgtggcta ccgccacacc agcaacggct agtccgtgcc
33721 gtcgcctttc gcttcgcacg tcgccgtccc gtcctgcctt gcatatgtaa gagaacctat
33781 gttctgcaaa aaatcatatc ctttcgggac taaaggcgtg taaaattcac ttatgacgct
33841 tgttcccaca tcgcaatagc cacgcccttt gatacgcttc tggaaaaact gtttttttcac
33901 atcagaacca aacagcatac cgtaacctaa ttcgctgata cgtcccaacc ctacacggaa
33961 attgaaatta tctctgatac cgtccgagaa atattttgcg tctggacgct ggcaggcaac
34021 aatcagaaaa tagcctgctt gtcgtcctaa cataacgatt ttctttaact ggctaagtaa
34081 acttacgctt tcctttgtcc ccagcatttc aaaaaacgcc acatattcat caaagataag
34141 aaaacagggg ggcagtccca gataggcgta gttttcgccc gtcttatagt tcgggtgttg
34201 cttcatttct tcgcttcttt gtaccatgcc ctcataaaag gcattgacgc aatcaatcat
34261 ttcttctttg gtgtgatata catttcccat aactgtacct aagtctgcaa ggtcagcgtt
34321 cttcgggtct aagatgtaaa gaatagcgtt cgtatgtagc agagcttcaa taagcgtcag
34381 cagaaaatac gttttaccgc caccagttcc accagcaatc agagcgtgag gaagtgcgtc
34441 atattcccat acaagatttt tcatcagtct aagacagccg ttttctgctt tcacttcgtc
34501 aatggtaatg cggttcgcta tcatatcata aagcagtgta tattcgatat agccgtcatg
34561 taaggtcttg tcggtcagtt cacaatacaa gccactttcc aatttatcct ctaatcgtaa
34621 gagctggtct tgatatttcc ccagcgtaat ttcacagcgg atatgaagca atcctttttc
34681 catttgatag taaatctttg gaaaccagac gattttttcc cttgatctgc tttgcaggtc
34741 agtaaaaaaa ccgccgtctt gtacagtatc ggcttcatac cacttgtttt ccagtatcat
34801 tcttgccagc ttttgacggt gcagtagctt cttgaaactg tcatggcaga agcggtaata
34861 ccaaaaagcg accaatgcac aaacaccagt cgctatcagt atggttatga agttgtaagg
34921 ggaaagcgtc aagccgttct ctaacaagct gatatgctcc caatcagtac gcatgagctg
34981 tttcatattc agtagcagaa gaactaccac gaatacaaac agaagcgtcc ctatggaaaa
35041 atggtagaca aggtgcttgt cgctggctct gatacggtgt cctttattca aaatcattcg
35101 cataaatcaa tcactccaat cttaaattta agtatgaaaa aagcagtcaa tcttaaaaga
35161 ctaaccgctt tgtgtgtatg aatatgaagt tgttacattg tagggtatgt tcattatacg
35221 tgctttgttg ttttaactct tgattgtgca atttcttcaa atagctgttt ttcgtttagg
35281 tctttaaaaa ctcttttggg aattaacgta tataatcgtg gtgcttgaat taagtaataa
35341 aaatcatggc tttcccatag tgcagagtag atgtcccact tcacttctga gtcaatgctt
35401 tgtgttttaa atttaatgtt ttccttagaa aataccagcg tatattcctc atgatatttt
35461 gcagtttgtt tgaatttcag tataggcatt aggatatata aatagcttcc caatgcagtt
35521 acaacaatta tcaatccaaa tattaaaata ctaaatacgc taaaagagga gaacagcata
35581 taaatgactg acagcaacaa aaatacagcg actaaagcaa tatcatattt atgtatagtt
35641 ttactagaaa ttaaatattg cctttctgct tttatatatt cactttgagt atattgaaat
35701 cttaactcta ctgtttccaa agttatacct ccgttcgtta tataaataat ctttgttaca
35761 atttcaaatt tgaaaatcac tttttttga taagtttcag ttttatattt ctcctttcag
35821 caatcttccg aaacacctct accacataca agcattgtgc cttaaatggt agtaaggaca
35881 attcgtcatt ggattgtatt tgtttcattg cttcttgcaa ttctgaaaga tctaataaat
35941 taactgccac acaaaagaaa gtctttctac gcccgtcatt atagttagaa agtaagtaag
36001 aaagaatttg tgttttttcc tgctgttcaa gattatattg ctcaatgcct atgttttttg
36061 ctctttctaa atctgctttt cgccgtttat gcgtgatgaa agaatcataa tcatcaatgt

36121 gttgatattt ttcgcatgga tattgtctgc attcatagca atattcaatc tttccatgtt
36181 caagactaca tttcgcaatt ctgcatgatt gattaccatt gccacaaccg ccacagtaat
36241 ttcccaaaag catagggcat aatccgcaat tcaatccgca aagagaaaat aattggtttt
36301 tacgttcaaa tcctttcata gtaccagtct cctaacttca aatttatctt tggatttatt
36361 gacttttaca aactggaagt tatcgctctg acatatattt ctgtatttcg ttagaagtca
36421 attttcgtat ttgtgtattt gcatattctt tatattcttc aacagcaaaa ataggctttg
36481 tatctttaca ggttttacct ttattctttt ttaggtataa ctgtaagtcc tcattatttg
36541 caaaaatttt gtaggaacgt ctcccatttt cacttactat ttcataaata ccacatgatt
36601 ttttcatact gtaatcagca gtacgcaaat acatttttgc aatttctagt gacttgaaag
36661 gaaaattcca tctttgcaga ccacgcttag ggtcatgttc aatacagatg caccgtccac
36721 aagtcccaca gatatactgt gtatgatttt gtaaacattc tatcggattt aacaacggtg
36781 taattctatt ttcatcaata tagcattcaa tgcacattat ggtctcatct ccttacaaat
36841 tatatttttt atcttctata gtcatataaa ctgggatttg cttacttctt tttcttcggt
36901 tttggtgctg gtagttcttc atacatagca tgaaacaagc ctgtcaaaaa ttccttatta
36961 tcaacttctt ctaccaacaa catttctttt gctccctcat agggtaactc atgcagagct
37021 gttgacatat aactaattgc tgattttact ggtttgacaa gcaatctatc atcatagata
37081 ccgccgacaa tcttaccacg ataatagata ataaattctc ccatcatagc acgatatgta
37141 atttcttcta attcggataa ctgccctaaa ataaactcca aatattcttt gcttgatgcc
37201 atacttccac ctcaaattcc tatttatata ctaaattgaa gtataaaatc atcatttcct
37261 gttgggcttt ctttttattc ccaataattt aacaaaatca ttaaataaga ttgtatctgt
37321 tggctcaaac atcaaccatt ttccgtcatg atggtgctag aatataaata gtacaagtca
37381 aaatgagaat tcctaatcta atatttcatg gtacaatgta ggtaccgcac tgaaggagga
37441 tctcattatg gcaaagcact atgacaaaca gtttaaactg gatgcagtcc agtattatca
37501 tgatcacaga gacttaggac tgcagggctg tgcatcaaat ttaggaatca gtcaacaaac
37561 attatcacgt tggcaaaagg agctacgtga gacaggtgat attgaatgcc gtggttccgg
37621 taattatgct tctgatgaag caaaagaaat cgcacgatta aaacgtgaac ttcgtgatgc
37681 acaggacgct cttgatgtac taaaaaaagc aatcaacatt ctgggaaaat gacggaagcc
37741 atttatctcg aagttgctga gaaggcagaa actgcccata aggctggacg ccgggtttct
37801 gtctccggaa tgttgaaata tctcggcgtt tcccgttctg gttaccgtgc atggctaaaa
37861 catgttcctt caaacgctga acagcgtcgt gaaacggtaa aagccaaaat caaggatatt
37921 tatgatgaat caaaacagaa ttacggtgcc ccaaagatta ccaaagaact gcgaaaatca
37981 ggggaaataa tctctgaacg tacagttggt aaatacatga ggcaaatggg aatcaaagca
38041 caatgggtaa aaccatggac tattaccaca aaagattctg acttcagtag cgaattgcaa
38101 aatatccttg atgaacagtt taatccggaa cgcccaaatg ctgtctggtg ttctgatatc
38161 acatatatct ggacgataga cggatttgtt tatctaacca gtattatgga tttatattcc
38221 agaaaaatca ttgcctggac actttctaaa acactggaag tatcctgcgt gattgaaaca
38281 ataaataaag ctaaagcaag acgtaaaatc gaagaaccat tgattctgca ttccgatcgt
38341 ggcagccagt atgtatccaa agaatacaag aggataactg ccggtataca gtgtagctat
38401 tcgaaaaaag cttatccttg ggacaacgca tgtattgaat cattccattc attgatcaaa
38461 cgtgaatggc tgaaccgctt caaaatccgc gattatgatc atgcgtaccg attggtcttc
38521 gaatatctgg aagcattcta caatacaaaa cgtattcata gccattgtga ctatatgtct
38581 ccaaatgatt atgaagaact gtatcgtaga atccagaaag acgaattgct gatggcaagt
38641 taagatgagg aaaacctcat tttaacttgt actaaatctt gacataggac catgataggt
38701 ttttgtttca tcataaattt tttgcacttc ttttgaataa ctatttctat ctgtatcaaa

38761 ttttaatcgt tcatctttcc ctaagataat cataaatcct acacagtttt cccttgcgta
38821 caatgcacaa agcgtttttc cgccacgacg atatttatac tcataagtcc acgctttacc
38881 acctttgttc cataaacatt ccatgtcata cttttcatca attaaggcac acagtttatt
38941 ccatatatca tataatgatt gcccgactaa agcagtcatt tcttctgcat taggtatttt
39001 atccagcata ctcatttctc ctttataact tctaatttat ctttgtattt attgacttct
39061 acaaacaaga atttatctca cgcaggcatt tataaatttc ataggcacgt ccatgcgttc
39121 tgcaacttgt tctggtgtca ttccgctatt taagaaacgc ttgcatacta ttttcatgtt
39181 ttctccaact tcgataatat gtttatctgg gtcataaaag cggactacac gttgtcccca
39241 tgaatgttct ataatagggt ggacatattc aatatcatat tctttcagcc tatcagcaaa
39301 tctatcaaaa tcatcttctt caaaatagac ttcaaaattg ttaccaccaa aagaaatatt
39361 acttgtaccg ataaactctt tgtatgtttc acttgtctgt aaggctaagc cacctgttaa
39421 agttttattt gctccaaaat ccataattac acgaagtcca aaaacttttt tataaaactc
39481 aactgattta tctatatcag tcactaccaa cataggattt ttaaatttca ttttcgttac
39541 ctccgcaaat tcatattatc ctgttttcca tatcaatatt ataacagtca cttatatctg
39601 ctacaatgaa aatctagttt gctagtttat ttttttgatt gtccctgtag tgtatggttc
39661 tgtggatttc ctactggctg gttgcctttg ttcttcaaaa taatatcgtc agccttgata
39721 taccagtcca cgtcagcccc tctaaagttg gcatttgcta ctgtatcggc tacgggattg
39781 ataagttcca cttcggcatt atacggatag tccttgactg ccacatcagc aggaatagac
39841 acttgtatca tgcgttcctg cacactgcat tttaaatcat aagttctgcg tttgatttct
39901 tcacttgtcg ttccgtcctc attttccata cgcacctcat gtctaagggc agagaacttc
39961 aatacaccaa atgtcttttc tttgtctaat acaattccat ttgctaatct cataatgaat
40021 acccccttat ttactttcca ctggcaacat atcatcagcc agtaagatat agtttgtaaa
40081 gccacgccca cggatattgt agccctctgc ggtaatgcgt gggttaatca gtttgatttt
40141 ctgctctggc ttgaaatgtt tctcgccagc tttggcagga atggttacga taatatcatc
40201 agctcgttgt acgtcggaat acaagttgta gcttctggtt acgggaacat agcgaccatt
40261 cacacgctgt tgttcaatct tgctttcgcc tgcaaattcc aaattgccga atgtcttttc
40321 catgttcggc acaacatatt ttaattccat aatgttttac ctcatttctt tctttgaatt
40381 tttgtttagt aatagagatt ttggggagct acactcccca agcccctagt ggactgtttt
40441 ttctcatcat tccataaatg gattgatgaa aaaagaacag tttcaatgtt aggtttccag
40501 tctgccatag gtaatggcgg agagcgtgat ttttcttttc tcatacagtt accgtcctat
40561 gatttttttca tgcgacggcg tttgtagaag tatgtacctg caagaccgcc agcagaagca
40621 agcaacatca cgataaatgc cataaggttt gtactgtcgc ccgttttggg actgtccgtt
40681 ttagttggtg tatcgggtgt tgttggtgtt tctggtttct ctggttcttc gggaacttct
40741 ggtacttcct tgattgttac cgtctgcccc tcgtcctcaa tatccttatg ctccgtaacc
40801 tttttcggtt cgtctggatt gctgaaatca tacaattctt caaaggttac aagctgtttt
40861 ccgccaagct cggaagcgtc gaatgtaaag gtaatctgta cttgcatgtt ttcactgtca
40921 gcggtaaagg tatagtcatt ctctacacgc ttgccgtcaa tgataagttc ggcgttttcg
40981 tccttaatca tctgccagcc tacaagctga tacgttgtgc cgacttccaa accctctaaa
41041 gtaaccgtat caacaatcgt tacgtctttg cctgcttcga tttctttctt gccgtcctca
41101 aaagtagctg tggtatgtat cttgatgata cgctcggtaa tcaataccgt ttgcccgtcg
41161 tcctcaatat ccttgtgttc tgccaccttg ataggctcgt ccttatcgga aaggtcgtac
41221 aattcttcaa aggttacaag gttcttgcca cctagagcag aagcgttgaa tgtataggca
41281 atttcgactt tcattgtttc atcatcagca ataaaggtat agtcgttctc tacacgtttg
41341 ccgtcaatga taagttcggc gttttcgtcc ttaatcatct gccagccttt caactgatac

41401 tttgttcctt tcacaagccc gtctaatttg acggtatcaa tgattgttac ctctttgcca
41461 gcaatgattg actttttgcc gtccttgctg gtcgctgtgg tatggattga gatttctttt
41521 tcgtattcat cagtcaatgt gcctaaatca acggttactt tgtttctgga aatcacaatc
41581 tcaaaaggtg ggataagttc aaagcccttg ttgctgtcag aacgtaattc ttcaataatg
41641 taagtgtcat ataataacgc tcctttgctg tcgtctggtt cagaagttcc aaaccatacg
41701 ccgtcctcac tggtcttgcc ctcgttggtg ttgttcttgt gggaagccca gtcagaagaa
41761 gtagagaact gcccgttatc gtccgttacg acaatatggc tttctcccgt cgtcttgctt
41821 gtgatcctaa aaagaacatc agcaagacgc ttgtgcgtac ctgctccaat cttcacgccc
41881 tctaaatcgc cacgcttaat ttgattgtag acagaatggg cttcgttggt taaatccaca
41941 atttttccat tctctgtaat ggtaaactca atcgcttttg caccgtccgt caagtagccc
42001 tctggtgctt cgctttcttc gatacgatat ttcccatgtg gtaataaatc agcagaagta
42061 gaagccacgc cctcaatgtc cgtatgaatg gtctttacga cttcattttt cttgtataat
42121 tttccctcaa ccaatacggc actgtcattt agggaaataa tttcaaaggc agtatctttc
42181 aaggtagctc cgccttgtgc tttggtgtct ttcgtttcta aatcacgttt ctggattttg
42241 acaccgccac ggataacctt gtctgataca gaaaattgat tgcttccagt cagtatggca
42301 aggtcgccgt cctcgctaat ctgtgttaaa tacattcctt tgatttgttc ttcactgccg
42361 tttgcctgca tatatgcacc atctaacaaa tagccgtctg gtgctttgac ttcctctacg
42421 gttaatgtac ctaatggaag tacggcgttg ccgtcctgca tatagaagct gtcgccagcg
42481 actttgtaag cgtcggctaa acgggtaacg taatggattg taccgtcgct gtctttttca
42541 gcgattgtct ttgtcgtcca tgtacgagtt ggttcagcag gaagattatc tttgttgtag
42601 tagccagcgt aatacttcca tgtgaactcc gcaccctcta aggtagcgtc gccctgcgga
42661 ttgcttttct gtgtttccat atcaattttg aaaagctcaa tcaaggtatc ggttactttt
42721 ggcgtgtcag ctactttcaa agtcgctgtt tttcctgctt caacctttag agaatacaca
42781 gtattatcaa tcttgtagcc agtcggtgcg gataattcct tgatatagac tatgccagcc
42841 ttgacttcta cggtttcggt attgccgttg ttatctgtcg taagggtggc aagctggttt
42901 gtgcagtcct tatcagaaaa gacaccatac gtcgcaccag cgatactgta atttccgtta
42961 ccgtctgtaa tgctggtgtt ggtggaactc ttttgtagtg tagcgttacc tacattaagg
43021 ttcgcccaga actgtcccaa ttcctgcccc tcgccagagt agatataacc gccacattca
43081 tagcgtcctt tatttgcttt gacaaaggtt ttcgctccta cgaatacttc gtcctgcgtt
43141 gcctttggaa tttcattata ggaagctcgc acgttatcac actgccagcc aagatgtacg
43201 ctcaatctct gccagacaac caactgtctt aaaaggtagg cgtgattact gctcaaaccg
43261 ccgtggctgt ccgtgtattg cttgacgtat tcaatggata aggcaacgtc ggctatctgg
43321 tcagcactca tgcgtgtgct tgcgtcagct ctggtcttat atccatttct aaagtccgtg
43381 ttaatgtcaa tacaataagc gtcctcaccc tcaacggtca aatgtccctc gttaaaggtt
43441 gaaccgatag aaccgtcatt cattactttt tcaacgatac cgacacgctc atcagattgt
43501 gtccagtatt gcttgctttc ggcatgaaca atggtactcg gcaaggcggt aacgatagtc
43561 gcaaaggcta agaagcccgt acataatcgc ttgaatatct ttttcataat tcttaatgct
43621 cctttcattt tgggtaaaaa aatagacgct cttttctgaa cgccctacat tgaaaaggga
43681 tactctttta caagtacccc tcaaatttaa tatttaattt gttgtgttcc cacactctat
43741 atcaactact actgaaacat tgatatagtt tttaatctct ttatttcgga ttttatcacc
43801 ctcacttgta aacacgatat gcttgaaaaa gtgcctaaaa ttaaggattt ctacatagtt
43861 tttcgttgta gcaacacaca agtctctata tggtaagaca ctgcagtttt atccagccct
43921 gccacctcat ccttctgtgc gaaccatacc ccgccaagga agagggcggc gcagacgagc
43981 accgctaata ccggcagtaa ccggcggtac catgcattgc caatgatccg tatgaagcct

44041 tcctcccctt cctcgtccgt aaccggtata tatcccgcgg cccgcttcga ccacgatagt
44101 ttctgccggt aggggtaaag ccctgtctcc gaccggtctg tgtcaaggtg gttttccgcc
44161 gatgcgtttc cccacgtgtg catgtcagaa aagaccctga agcccccgtt gggatacatc
44221 ttttgtaatt cctcttctgt tagatagttc cagatatcgc cgcccttatc tttgaatgtc
44281 cttttcgtct ctgacagcgg ccgtacctta tacttgataa gtcattcctc ctttatcctc
44341 ggaagtactc ccgtgacttc tgctcaaagc tacgggcagg ggtacgccgg gaaccctgtt
44401 attcaaaaag ggagccagcc tttgaaacga ctgactccat tttgtattat cagaatggtt
44461 tctatatggc aggggcatca tccgtaaagc tgaaggttac cgttttggaa tatgtccctt
44521 ctgctttatt tcctgccgcg gggagattca ggtaagcata gccagtgctt atattttcag
44581 tacctccgtt cacttttttc atcgttccta tggtatacgg atcagacact tccgctcccc
44641 ccgcggtggc tgcaattccc attcttacgt caccgccggt ccctgcttta tccattagaa
44701 ttccgccggg agcataaccc tgaacggcta ctgtgactgt cctatcccca ttgtaaggat
44761 ttgaagtcag cttatccttt agctcaaact ttaatgctgc accgttagct gatgaattat
44821 tatagtcagt caaaacaatt ttcttgggat aagccaccag ccagtctgcc gcatccggag
44881 ccgcggaatt tgccgtatat gttacttctg tccccgccgt tgcattgttt gccgcataaa
44941 ccggtactgc cattgtacta acaactactg ctgtcagtgc cagcatactc ataaatttag
45001 ttttcttcat aatctgattc tccttttttct ttttctttat agtatataat caacatcagc
45061 agtgctgccg atgcaattac caagactttc aggcctagca ctacgcctga atcacctgtt
45121 ttgagggctg tggcgtgcac accctcccct ctctcaattt caaccaccgg aggcctatta
45181 tctgcaatat agatgaattt tgtttgtccg ctgtccccgc cgcctccggt attcccgttt
45241 gccgagacgg tgcctccgct gaatagaaga agcatcaata aaatgcctgc acttattatt
45301 tttcttgtca tgatgcacct cccgtttatt gtacagcgga gatataaaac aaggtcccct
45361 gccctgtatc cgttggatcg gtactgatct ggaagactcc tttgtatacc ccgttgccct
45421 cctgggactt cgtgttgaac cagatatcct gagactggtt tgccgtgcct aatacaccgg
45481 ctcttgcata gccctcactg atggccggtg catctaacag ctgcccctga ctgtcataag
45541 aagaaaccaa tagtttttta ttctggtcat tagtctggat aagttcaaag tttttcagga
45601 cgtctacatt cacctgttta ttgcttccac tcacgctctt gtaagcaacc gttaccctcg
45661 cccccgcatg atccttgtcc ggattgaggt tcgttccatt ttcagtcagc aggatatttt
45721 tcgggatcga gatgtaggaa tcagggctga tgtccttcca cactgctctc aggcgtatgc
45781 ttttattatt cgcattctga ccggatgcaa atgtataaac agctgtatcg ccttccttat
45841 atgtaagtcc cgtttttttct gtccatgctc ctgtatccga catttcctca accgtccagt
45901 ggtcgaacac cttcccggca ggatactgtg atttatcggg gaaattaata gcagcgctgt
45961 tttttccagc ataagttcta tagctattct catcagtgag gctggtaagt ggacttccgt
46021 ctgcatctat atagctgaca ttaagtcccc tgtcatattt cacagaagcg acttcctgat
46081 tatccgctgg tactctatcc agagaagttg tgtgctccat atttagataa tatgtataag
46141 cgtccccttg accgtcatac gtttccgcaa aatttttgaa gataatattc gaaccgctta
46201 tcatctttac atactttgta tcggaaggtg cagcattgta agtaaggtct gtttcagaaa
46261 cattttgtaa atacttattt atgtgtccgg ttacatactc gccaggtaaa attccacctt
46321 gttgataaac aggcactatt tctacttcga atacggaagt atacataatt aacgcaagac
46381 gttttttttc atcgtcattc aaggggccac tatccgatgc agccagcagt gcggctgtat
46441 cacggcggaa atagtcggaa atgtcaatcg aatcccccgg cctgaaatat tccttacctg
46501 ctatatcctt atacaatgtc cttacatagt ctgccggcag caaggggttt tccactgcac
46561 gaatctcagc cttaaagcca acgaatgtcc cattccctcc acttggatca tctggggctt
46621 gaaaactatc tgagaaaaca atcttgttct gctccagaag gtcaaaattg accccgtttt

46681 ttacaatcag attatcaggc ttttccgttg taacatactg ttcattattc gtaacagtcc
46741 cgctgattcg ttccgcagta atatcaaccg cacgctggct atgccgtccc gttttattag
46801 caacaggcat aaacagtaca tattggaaag gactgatttc ctccctgctt tcggcaccgg
46861 cattaaacca tgaataattc ccaagctcaa gtaaactttt tttttctctc tcaactgaat
46921 attgcgtact gatgggatct gtccccgttt catcaaagct taatattttc ggattagtat
46981 acccccactt tggtgtgacg gcaaacttaa aaaaatgatc cgtatctgcc gcccccgggt
47041 atcctgtgcc cgcagcataa ttcttatttg cgatcgactg cttatccaga aacattttcc
47101 cggcagaaat ccctactttt gtctgatttc ctgtggtagg attacctgag ttatcctttg
47161 tgtcccgtat catataggaa tccagcgcat cgccatttga acccgtggaa atattggtaa
47221 gtttatcaaa ataatgcggc ccctgtgcac tgctgcttga ttgtgtctgt aaataaaagc
47281 tgttctttat attgctgacc gttatgagcg ccctgaactc ataggaatca cggcctccgc
47341 cggacggcgg ggtatattga gccttcataa gcctgacttt aatctttgcc ccgtttttcg
47401 ctgtatattc tctcataatt tgcgacgcac ctgtgttgct tgtatccctc cagctaggtt
47461 cccaaagtgt aaaagtttcg ctgatcggca tgtaagttga gaaatcttcc ccattgggga
47521 aataattcag tgtcagtgtg ggggagggcc agtaaaaata aaatggagat cctcccacgc
47581 ctactcctcc tgtgcggtac atataggact caaatattaa ttcaccgccg ctgctgaatg
47641 tcccggttga aatatcttta tcttctatac ttccagcatc agtaatcact ttaacttttt
47701 tcggacccgt gcgggctatt gaagcctttt cgttactatc catatagttt gtaccataat
47761 acggatctgg atcctgatac ctacccgagg catccgtctt tctcagccag ctgttcccgc
47821 cttcaccgct gctctgctta tcacctgttc tctcatacac cccataaata agttgatccc
47881 gttcttcgtc tcccactacc gtcatcgtga cgggataatt cggatatgta aaagaatagg
47941 taaccgtcct gttgtcctta tcctcttctg ctctaatatc cggtaccggc acattggttt
48001 cagaatgctt aaactcaatg ccggcaggcg tgatatttgc tgacggcgct gttgcataat
48061 atcccggtgg atacgttagc ttaactttca cttctgtctt gtatcttgcg cttacactcc
48121 catcactgtt ctgaataaag cctctgagat actccagttc aaagcctgca ctttctttat
48181 cattcacgac tgtgactgaa aactcggctg tatttagatc aaatataaag gaaataactg
48241 ccccgctagg tacttcatac gcgatatcga agtcactgtt ctccccctgg tctaatgcat
48301 agaagtaaac tccatcgatg aaatcgtagt aatgtatagg atatgctttt gtaactccgc
48361 cttcagacat ttgaatttgt gcacgaacaa acttataagg ctctccgtca aattcctggt
48421 cttctattcc ggtgatctct cctcctgcaa aattataatc tcccatctct tttaaaccat
48481 tttctgtata cattgggaca gagattgtag atgtattact tgctattttc atagccttta
48541 attcacccac tgtctttgta tccgcagact ccggtgaacc ttccggtgtt tccaatgtgt
48601 ctccgggtgt ttcaggtgct tcaggtattt caggtaatgc aggtatttca ggtgatgcag
48661 gtgtttcagg tgatgcaggt gcttcagccg gttccgtgag cgaatctacc gccacatcaa
48721 tattctctga atgctctcct gttggtgtcc tgaaaaacac ttgaaatgag aacacttcat
48781 tttgggttac atgataagaa ccattctcca aatccatgct gcccccatcc gggtcaagga
48841 tatctactat ctcaactccc tcatgtatgt ttgtcccatc tcctataata tctgcttccg
48901 taccctcttc attcggaata atttggaaac gatatccgct cggcgcctca taagccttta
48961 ccttcagttt tccggtatca gataaaacag ccgataaaac cataagtaaa gcaagaaata
49021 aagctggcat tctcccccaa taatttttaa ttctttttcc cattacgaac tccccctgtt
49081 ttacaacata atatgtatta tgtcgtaaaa caggctcatc ccgtaatgta tattaaagcg
49141 taaagtcttt ttgattaggg aaatttaacc aattatagct ttcagaattt tttgtattgg
49201 attttattag atttttcaga cattttttaa gattatatat tgtaaatttt ttccaattga
49261 tatatactaa tacaaaacaa gtaggaatta ctattttgtt ttaaaaagtt ttattccata

49321 aacaacataa tacatattat gttgtttttt tatgtatagg aagaaacaac caggtccatc
49381 tttgaaagcc accttttaca ctcttttcca gttgtcgtcg tataggtcct cgtcgtttct
49441 atacttgcgt gtccaagtat atctgccagt ctgacaacat ccttttgcat acgatagtaa
49501 gttaatgcga acagatgtct tagattatgg ggaaacacct tattctcaca gactttagct
49561 tccttacata gtttcttcat ggcattccat atattgctcc gatccattgg ttttccattc
49621 ctggtaataa agaccggccc tctgttaata ttctctttca tgcagtaatc cttcagcatt
49681 tttatcagct tcattgggat gaaaataact cttactttcc ctttattgtg aacaattgct
49741 tttccgtttt tcaaactttc aacagtaata tacctgtgtt ctgataccct gatcccagtc
49801 ccgcaaatcg tctgcaaaag catatagaga cgtatattcc cacctctcaa cgctgtatcg
49861 agcagtctct tatattcatt tttggtcagt tccttatctt cttcatagaa taccgaactc
49921 tggactctat aaagctttaa ttttagattg cttaacttca taaactcaag cagcccattc
49981 aatgcaacaa tcatactatt gatactggaa atcttatatt gatcagaaag gctgttttta
50041 tatctaacca gaacttcctt tgtaatctgt ttatttcctt ccggcaaata tttataaaaa
50101 tgcgtaatat ccctcatata cttttcgatt gttgctttac tgcgttcttc atcatacaaa
50161 tatgctttgt attcttccat aattttcaag tctaatatca ttcgcttcat aaatatgcac
50221 tcctttatcg taacagtttt gtccttatca taacatgccc cagcatcaag ttgactggta
50281 ttttttttcca tatttagttt atccgagaag ctatattctg atccggacca aaagacataa
50341 agcggccccg gaaacgtcca ttttatctac atttccgggg ttctgtataa gttccgtcat
50401 attttcctga ttttgatcct cttactatta catctgccgc atgtttgtga catttataca
50461 agcttttcaa attcaggaat tttgactgtc actccgcctt tcatagctga cttgtgtgcg
50521 catattccag ttcctgtcca ataagcacct gctatatcgt ctggtacagc ttgtctgttt
50581 tcgataatcg cccgaacaaa ttcatgaatc atatgcgggt gcgatccacc atgtcctccg
50641 ccctgtttaa aagagagatg cgtattttca tcattataca ctgtttctgt cgtaaatgaa
50701 ccaatactct ccggaagtcg tgaggcatag tccggtactt ttattctatc ttctgtaatt
50761 tcagagcctc tgctatatct ctgaggatcg ccgtcaatat cgaccatttc ctgttgaatt
50821 ccccctgtcc gtttataaat aaccggatct tccgaagcaa gctgctgcca ttcaaaactc
50881 atttggcttc catatatcct gaagcattct gaataacttc gggcaactcc atataaaaaa
50941 cgttccattt caatcgcaac atcgctgttt ttcattgtga ccaaagcgct ctcaaacgca
51001 aaaggcgaat tgtactgccc aatcaattca tctctgatcc gcccggatcc tttcgcatat
51061 acttcctttg gatagtcatt taaaagcata aaacaaggtc caatagcatg cgtcggatgc
51121 ataagcggcg gaaatccttc ccaatatgac ggccatcctt ccatatcctg gtagtgggca
51181 catgtcatat attgtatttt tccgaattca cctttattta taagatcttt cacataaaaa
51241 tactctcttc cgaatactgt tgtctccata aacatatact gttttttgct ttcttttctt
51301 gcccgaatga catcttctaa ttcctttaga ttcattccca taggtatcgt acagccacaa
51361 tgctttcccg ccattaaggc acgtatagaa aaatctgcat gtgtcgccgg aggcgttgta
51421 atatgtatcg cgtctattgt ctcatctttc aataattctt caaaattatt ataacaatcc
51481 cggtctgcga ttccaaattt ttcatttgcc aattttatca aatcgtcatt cgtatctgct
51541 actgcaacac tatatacatc aggatgcctt aaataaattg gaataaattc catgccaaaa
51601 cttatcccta caatcccaac cttaatttta tgatcagtca tacagatcct ccctttaatt
51661 aaaaagcggc atttcctgag gataaatccc catatttcca ggacggggta caaggtcttc
51721 gcttcgttta gatacatcca tcagctcagg aagatcattt accggactgt caagataaaa
51781 atatgtacag caggaccaat catcttctct ctcaaataat agcggccggc cgtctaacat
51841 agggatatca gcaaaatcta cctcttctcc atctacagaa ataatccggt tatcagccgc
51901 ctgattcttg taaagcagct gggcatgatg gaaataatcg tctctgtcaa cgccgccgat

51961 ctgctgaatc gtcgctctga aatctgtatt gaaataaatc gggtcaggga tatggaggcg
52021 ataacaacac agttccatat attttttatc atacaccgga cagccacagt ataaatcata
52081 aaaataatcc tgtccccacg ctgtcccaat gtaatcttcg acacccgttc cgcatatcgt
52141 cggatactta tcatctccgt caatatacgc cttaaattcg ccttctcccc accatgtatc
52201 agaatataat tttgtattgc accgaactcc catatttgtc cccaggaacc tcccccggcc
52261 cttaacacac ggcaaaattt caaaatcctt tttctggcag gttaaattct gcctattaaa
52321 atatgagtgg aaataaagta tatcctctcc cagcgaatcc cctaacgtgt aatcaatatc
52381 atagtaaaaa tgtctgacag tggtcttaga ttcatttgtt atcacaattt tcattcctgt
52441 tttgaacggc atcgggatgt atgtactgaa acttctgccc tcaggatttg aaaaaagtgc
52501 gtttttaaat acagctgttt ttcctagtcc tacacagaaa aaatcaccaa gcggaacgct
52561 taccgccgcc ttatcattca tatcccaaaa catttcaagc ttcaggcccc gaagcgtctc
52621 cggcgtctga tcgcttattg taatccaaat gcgccgtatg ataccgcttg tattttccgc
52681 gtgtgcaagg actctgcttt ctccggttcc aagagcaaac catggcgatc cctttcttcc
52741 atttcttgct tttcctccgg ctcctttttc tccatccgtg ttttcaaaac ttgcccacct
52801 tgaaacaaca cccttcggga atgtaaacat ttcttctcca aacataataa atcctccgtt
52861 atttttagtc ttttatacct gtcatcttaa ttccttcaag gtaatatctt tgaaggaaga
52921 taaaaataat aacagttgga atgaccgcca caaccgatgc tgccaaaata agattccatg
52981 agattctcaa cgaccccatc aatgttgcga tctgtatctg tatcgtatat ttactttgat
53041 cctgtgcaac tatagtcggc cacaaatagg aattccaatt tgtcacaaaa aaggaaactc
53101 ccagagtaat aaacattggt ttcgcaagag gaacatatat ttttctgaat atctgaaaaa
53161 tagtacatcc atcaattctg gctgaatcat acaatgcatc cggcatctct ataaaatact
53221 gccggaacag gaacacacca tatgcgtgcg gcaccatggg aagtatcatc ccccacaaag
53281 aattgacgag ccccataccc ttgattataa aaaacaatgg aatgattata accgggaacg
53341 gaatcatcat tgtacttaag acccacataa acaaggctct ttttcccgga aatcttagtc
53401 ttgcaaaggc aaatgccgcc attgaatgga tgatcaaagc caaaacggta accattaccg
53461 caacaaaaag tgagtttccg ataccggtaa gaacatccag tgattcaaat acctctttaa
53521 atgcatcaag cgtaatatgc tgtggaatga tcctataatc aaatacttcc gtttcttttt
53581 tcaatgaact actgataaac agaataatcg gaaataatgc gaagaccaaa aaagtacagg
53641 caactaaagt ttttaaaatt aatattctcg tctttttgc cactgtatca cctacttaaa
53701 ttttaacagt tttaactgta ttactgtaat aatcattata atcagcgcca gagatactgc
53761 aatcgcagat gcatagccaa acctgtactg ttcaaatgca agctgataaa cataatatac
53821 tgtgacctct gttgacttcg ccggtccgcc tgctgtcatg accctaatct gatcaaatac
53881 ttgaaatgat gagataacag ttgtgatcaa tataaatatg cttgtccctt ttattaaggg
53941 gaacgtgatt ttgaaaaatc tctgtacagc attagcgccg tctatttttg ccgcctcata
54001 ataatctgcg ggaacgtctt ttaaggccga aagaaaaata atcatattgt acccaaaaac
54061 aatccatatt cctactatag ctacacaaac caacgcctgt gttttcgatc caagaaaaag
54121 ctgcctgctt attcctaatt gttccaatat tttattaatg taaccgtttc tggaattgaa
54181 taaaaaagtc cacaatgttg ccgctacaat tgttgaaatc gtattcggaa gatagatccc
54241 gatcctcaca gcctgctgcc tcttatttcc acggatcgaa tgaacaaaca cggcagctcc
54301 aagagacagc agatatacag caattacata tattccggca tatttgcctg ttataagtag
54361 actgtcccag aatgttgaat ctcttaatac atcgatataa ttgctgattc ctgtaaatat
54421 ctttgtgccg aacgcgttcc agtcaaaaag gctgatatat aatgcgtata atatcggcac
54481 aaagatgaat aaagccatca atatcacttc cggagcaatc ataagatatg cggtcaccgt
54541 ctcgtgtttt atacgcttat gttttctttt catcatgatc ttactccttc gtctaaatgc

54601 tcctcttcct tcccattatc cggtaaatgt agatccgctt ttttcctgat attgcaacat
54661 gctttcctgt gcggcattag ctgcttcctc cgcactcatg ctgtaaagtg catcctggat
54721 catagtagac acaatttcct ttgcttctcc tgaaagggcg atttccggct gcccggtcgg
54781 aagcacttca tccgagaata cggcagcatt atcgtttaca taataaatat ctttacccgc
54841 ttcaattaca gaatttctca tacagtaagt aaatttagca tccgtatcaa attcggtcac
54901 acggtcgacg tttaaccaca tccatgagat gaaatccttc gcaccggcaa ttttttcctc
54961 actgctctga ttgcttacca aaacattcca tcctccaact acactggctg tttctccgcc
55021 cttttcgact gccgggagag gtactactcc atacttttcc ggatctgcac gaaatacgga
55081 ccaactgcca ttgatctgca ttgctgtttc tccatttgtc aaaatcgaga tgtctgatac
55141 ggatctgccc ggcgtctcat tgcagttccc tgaattcaga aggtctttcc acaattggag
55201 tgcatttact acctcatctg aattcaggtt tgccattcca tcttccgtaa ataaacttcc
55261 cccctgcatc catacaaaag gatagaaggt aaaacactgc cagtcgccct tatcgacttc
55321 gaaagtaaaa cctgaatatt cgtccgtatt caatgctgca gccgcttctt ttaattcatc
55381 ccatgtagtt ggcggcttta ccccctgctc ttctaatatt tctttgttat agtatagtgc
55441 aaccagatct ccatccaagg gaattccgta aatatgtcca tcccttgata cacggtcaaa
55501 agcagcttct gtaaaatcat tttttatctc atcggtaaag atatcatcca gtggttcaca
55561 aaccccggca tcaagatact gattaataat tgcaggacta cattcaaaga tatccggcgc
55621 ttcatttgtg gcaaatgctg ttgcaagctt tgtcgtagtg tattcctcaa acgggattgc
55681 aagaaatgtt acatgatatt tgtcctgaga attattatat tcctcgatat atttttccag
55741 gaattccatt tccgcttcct gagtccactg attccaataa acaagttcaa tagggccatc
55801 accttgcgcc tctttcgttg tctccgtact tcctgtggtt tttccgcacc ctgacgctat
55861 tccgctcatt cccagcacta acgctcccac cagaacgaca gatattcttc tctttaattt
55921 gtaaagtttt ctcattgttc tcctccttta tacgttttta tgagctcata tcttttatag
55981 ccttattcta taaattcttt ttggtaaata aaatactaat ttttttaaa agtggaattt
56041 ttatttttcc ggcatccata ctctcatatc cccttttgcc ctatttcccc atgagaagta
56101 tgggattgct gtcagcaata cattacttgc ctgtggcatc tgagtcatat ataagtcatc
56161 cgattcttcc agacacatcc ctgaaacttt aagcaagacc agatctttca aaattccttc
56221 tttacatacc atagtctccg cctgaacatc cttcggaatt ctgtatttat gcaagctttc
56281 tctttgatcc gcctgttcaa aacaatacac aataggccct cttataaatg caacacatcc
56341 agcatcggcg cgcacctgag gattggcata aactttttt atattcagtc caaagtgaat
56401 ttcaagtatg tcctctcccc agtttctttc gagatagatg tagcctttgc taatcgtaga
56461 tttataattt acttcctctc cattacaaat aacctggaat ccttccgcat attctggaat
56521 tctgactgca agagtccact tttcctctgt ttttgatgct attttaaatg tcacatctcc
56581 atcccaaggc aatttactgt caacttgtat tttagctttc tccagttctg ctgtctgact
56641 gataaacaaa tgtgaataaa taatctcttt tttctcagtc caaatatatt ttccgactga
56701 cataaaaagc ctcacaagat taggcgggca gcatgcacat gcataccact tcggtcttcg
56761 ggtcagtaca tgctcaaatc caaaatgttc tccggatatt ccgcggactg tttccagcgg
56821 gttgacataa aaaaaacctt tcccatccaa ctgcattcca cccattaccc cgttatatag
56881 ttccctctcc ataatatctg cataagatgc atccgggtca agatccagca tctgtttcgc
56941 gaacataacc atggcaactg atgcacatgt ttctgcataa gctgtatcat tgggaagatc
57001 ataagcttga gtgaaggctt ctccacgggc tgttgaaccg attccgcctg tgatatacat
57061 ctttttttcgg acaatattgt tccacaattt tttacaggcc tcataaagcg ttacatcctc
57121 cgtttccata gccagtcctg ccatggcggt ataaagatag actgccctta cacaatgtcc
57181 taccgcctcg tcctgttccc gcacaggttt atgacattgt gcattttctc gatcataagg

57241 atccatatga aataaagtcc atcctctgtt ttgtttttct ttcataaaaa aatcaggatc
57301 ttgccctcgt tgttctacaa aatatgacgc taatttcttg tatttttcat tccctgtaat
57361 aaaatacaat ttcatcagcc ctatttccac ctctggatgc cccggaatcc ccgaaatttt
57421 atcgtcccca aaccgtcggt caatgtggtc tgccaatcgg attgccacat caagcagctc
57481 tcttttccca gtctcttcat aatatgccgc agctgcttca aacaaatgcc ctgcacaata
57541 taattcatga ccttcctgga gattctgcca tttatgctcc ggctctttga taataaaata
57601 tgtattcaaa tagccgtctt cctgctgcgc cgcttttata taactgatta ttttatctgt
57661 tctttcttcc agttccggat cagatttgac tgacaatgaa tatgcaacgc cttcaatcca
57721 ttttgaaacg tcgctatcct gaaaaacagt cccataaaat tcaccattag aaatacctgc
57781 tgcaatctta aaattttcta ttgcatggct tttttctgca cctgcaactt catctttaag
57841 tatcttttcc tgatagggta tcgcctcttt tataatcaca ttttgtatat cgccccaaaa
57901 accttctgtt atttttatat ttttaacttt aactcttttg gcaggtataa atgctttcat
57961 actcattatt ctccttttgc ggttttttatt ttattttatt tctatacact attagaaaaa
58021 aatactaatt ttttttaata gtgtattttt tttctttcat tatattatac tgtacatgat
58081 acaatctttt taacagccaa tactcttttg aattaggatc ggaattgtaa tttaatgaaa
58141 gcgaaaaaga tatcactaaa aatccagtta ttagaaatat ttatttttgc tgctatcatt
58201 cccatgataa ttacggcatt gtttctttat cgtttcgaag taaacagaac cctgaacgct
58261 gaaattttat cccagcggca gatattagaa ttaaccgcat cgaatttgct gcaaaaaatc
58321 agccacttca acaaccttac ttcatcggtt tactacaatg atgctatatt aaaatgtttg
58381 gaaaatggaa aagacaggac gttgtcttct tacgagaaaa tgcagataca gccggatatt
58441 aattccatga tgcgtacgga taaccatgtt ctttcttttt cgcttacaac gatgtccggc
58501 gatacattat actggaatga cggatacagc aatgtggtaa acgccacatc actcaaccca
58561 gttttctgaa gaaaatttat gacagcaacg gcagcggcgt cctgtcagac cccataacac
58621 atattaaaaa acgaaaattt ccttcttctg ctgcttccat ctactatgga cggttaatcc
58681 gcagtaccga gcataatttc tgtcccatcg gcattttact gattgaaata gatatctcat
58741 cttttgaaga tattatctca acttcaggat ttaaatctga agccatactg cttattacct
58801 ccgaaaacaa cactttaata tggggatacc ttgaagatct ggtaaaatgt caaatcggtg
58861 aaattatacc aaaagaaacg ttatctgctc cgtcaggcga ttattctgtg agaactatca
58921 acggaaataa atactattgc ttttactcca tgacggactc gctaaactac aattgctact
58981 cgctgctccc cgttcccatc ataaaaagtg ctgcacagta tgttcggatt gtccttttgc
59041 tgatatcagt agtaaatttt atctgtttgt tacttatttt ttcactatgt accaaacaga
59101 ttattacgcc aatacaacat ttgtctatta ttatgcataa ccagaattat atgaattact
59161 ttcccatacc agactatcct attcaaaatg aaataggtta cttatatgag tcttttcagg
59221 aaatgagtac gcgaactctg catcttgttg aacaattaaa agagtcatcc aaaaaagaag
59281 cccggcagca gctcgccctt cttcatgcac agctaaaccc tcattttcta tataatactc
59341 tggactctat aaactggatg gctcagaatg aaaatgcaaa aaaaattccg gaagtaataa
59401 atgccttgtc acaaatctta agatacagca ttattaaaga caatgatttt gtatctgttt
59461 ctgatgaact actttggtta aaaaattata tatatataca gaatatgcgg tttcaggatt
59521 cgttctctat tcaatataat attgcagaaa atattatgaa ttataaaatt ccccggttca
59581 ttctccagcc atttattgag aattccctgc tgcatggttt ttctgaaatc catcagggcg
59641 gaatcataaa actgaatatc tactctaaac aaaatgattt atttttgatt atagaagata
59701 atggaaaggg tatgaccgaa cctgccatat acacagttct ttatggtaac agcagcggaa
59761 ttggcattgg aaatacaaat gcctatatta agaagaaatt tggaagacct tatggaatac
59821 agatcatttc aaaattagca gaaggaacaa aaatcataat ccgtctgcct gtacaaaagt

59881 gagggatgta tatgaatcat gtaatcatta ttgacgatga aattcaaagc cgtaccggaa
59941 ttgccgccat ttttgaaaat cactctccca aatggtttat atcaggttta tttgaagacg
60001 gcgaccaggt tcttcaatac cttgaaacac atacagacat tcagctgatt gtcacagata
60061 tccgtatgcc aaaatttaat ggccttgaac taatcagtaa aatcaggaaa accaatagta
60121 ctatccccat tattattatc agtggatact ccgaattttc ttatgctgct aaagctgtcg
60181 agttccatgt atttcgttat gtgctcaagc ctgttcttcc atctgaattt gataaaatca
60241 ttacaagtgt agaaaatttt ctgcagttgg aaaaggcacc ctataaatat aattttaaag
60301 attcagaata tgaacagttt ttggatttat tattctcaga agctgcaaca caaaataaaa
60361 aagagaccgt cacaatgctt aaacgccaat gcgggtttag ttttcagaat atctggtttt
60421 tatttctgga aggcaattca catttttcta atatctctgc cagctttcgt ccgcagttaa
60481 aggagttttc aacagccctc tcccaggaca gtcatgtttt tctgttccgt cagaagcttt
60541 attgtatttt aatcaaaaaa cctgatatag acacagattt tctttataag cagctttcga
60601 aactcctttg gtcttttccg cgtgaagcag gtgtacatgc aagcatttat cagagcacct
60661 ttaccgacac attcaagaat gccttttta gcggaatttc tgctttaaaa cagtactttt
60721 atgaaaaaaa atcaatctgc ctatatgctg atcaaggtat atgtacgttt ccgcacatga
60781 tttataacaa tctgcagctc caggttgatc aaaacaatct gttggatgtc cgggattcta
60841 tctatgaatt tatggattat ataagagaaa aaagaccttc ctattttgaa cttcaatcat
60901 gggtgaataa gatttcactt gtgatcgtca aatactgcaa cgacaaaaat atgtccacag
60961 catgctatat tgattatgcg gaaagcctgg aattccttta taactattat agtttagaag
61021 aaattgaaga aacattgctg tcaatgattg acagtgtctt caataaagct tttgaacaaa
61081 taaacagtac gcatgcttat ctcattgatc aggtgcagaa ctatctgacg aatcatttaa
61141 atgaaaaaat caccctatca gatttggcag atatattttg tatcaattat acaagcctca
61201 gtaatattta tagttccgcg accggccaga ctatcattga gtatttaaca tttttgaaaa
61261 tctcaaaagc caaagaacta ttaattaaca ctgataaaaa aatttacaca atcagtgaag
61321 aattagggta tagtgacaca aaatatttta ttaagcgatt tcggaaaatt gtcggaatta
61381 ctccaaaaga atttagaaag atatatagga taaagggata attagagcca aaggacataa
61441 agcgaccccg gaaacgctga tttt

//

DEFINITION Integrative and conjugative element ICE_2

fasta_record 1..18181
/ID=B79|SC|contig000001
/colour=5

CDS complement(280..1497)
/color=9
/locus_tag="B79_00399"
/note="Tyrosine recombinase, similar to AA
sequence:UniProtKB:P22886 protein motif:CLUSTERS:PRK00236
protein motif:Cdd:COG4974 protein motif:TIGRFAMs:TIGR02224
protein motif:Pfam:PF00589.16"
/product="Tyrosine recombinase [recombinase]"
/"ID=B79_00399
/gene="Int-Tn"""
/protein_id="gnl|SC|B79_00399"
/codon_start=1

CDS complement(1579..1782)
/color=9
/locus_tag="B79_00400"
/note="Excisionase from transposon Tn916, protein motif:Pfam:PF09035.4"
/product="Excisionase from transposon Tn916 [excisionase]"
/"ID=B79_00400
/gene="xis"""
/protein_id="gnl|SC|B79_00400"
/codon_start=1
CDS complement(2243..2473)
/color=7
/locus_tag="B79_00401"
/note="helix-turn-helix domain-containing protein , protein motif:Pfam:PF12645.1"
/product="helix-turn-helix domain-containing protein [transcriptional regulator]"
/"ID=B79_00401
/protein_id="gnl|SC|B79_00401"""
/codon_start=1
CDS complement(2470..2892)
/color=7
/locus_tag="B79_00402"
/note="RNA polymerase sigma factor Sigma-70 region, protein motif:CLUSTERS:PRK12519 protein motif:TIGRFAMs:TIGR02937 protein motif:Pfam:PF08281.6"
/product="RNA polymerase sigma factor Sigma-70 region 4 [transcriptional regulator]"
/"ID=B79_00402
/protein_id="gnl|SC|B79_00402"""
/codon_start=1
CDS 3397..3750
/color=7
/locus_tag="B79_00403"
/note="Anaerobic benzoate catabolism transcriptional regulator Helix-turn-helix , protein motif:CLUSTERS:PRK08154 protein motif:Pfam:PF01381.16"
/product="Anaerobic benzoate catabolism transcriptional regulator Helix-turn-helix [transcriptional regulator]"
/"ID=B79_00403
/protein_id="gnl|SC|B79_00403"""
/codon_start=1
CDS complement(4096..6015)
/color=2

/locus_tag="B79_00404"
/note="Tetracycline resistance protein tetM from
transposon Tn916, similar to AA sequence:UniProtKB:P21598
protein motif:CLUSTERS:PRK13351 protein motif:Cdd:COG0050
protein motif:TIGRFAMs:TIGR00484 protein
motif:Pfam:PF00009.21"
/product="Tetracycline resistance protein tetM from
transposon Tn916 [antibiotic resistance]"
/"ID=B79_00404
/gene="tetM"""
/protein_id="gnl|SC|B79_00404"
/codon_start=1
CDS complement(6392..7306)
/color=4
/locus_tag="B79_00405"
/note="Conjugative transposon protein TcpC, protein
motif:Pfam:PF12642.1"
/product="Conjugative transposon protein TcpC [mating pair
formation]"
/"ID=B79_00405
/gene="tcpC"""
/protein_id="gnl|SC|B79_00405"
/codon_start=1
CDS complement(7321..8322)
/color=4
/locus_tag="B79_00406"
/eC_number=3.4.-.-
/note="Peptidase p60 Lysozyme_like and NLPC/P60
domain-containing protein, similar to AA
sequence:UniProtKB:P21171 protein motif:CLUSTERS:PRK13914
protein motif:Cdd:COG3584 protein motif:Pfam:PF00877.13"
/product="Peptidase p60 Lysozyme_like and NLPC/P60
domain-containing protein [pepidase]"
/"ID=B79_00406
/protein_id="gnl|SC|B79_00406""""
/codon_start=1
CDS complement(8319..10430)
/color=4
/locus_tag="B79_00407"
/note="Tn916 ORF15 signal peptide containing protein , ab
initio prediction:Prodigal:2.60"
/product="Tn916 ORF15 signal peptide containing protein
[membrane protein]"
/"ID=B79_00407

/protein_id="gnl|SC|B79_00407"""
/codon_start=1
CDS complement(10499..12946)
/color=4
/locus_tag="B79_00408"
/note="Type-IV secretion system protein TraC VirB4 components, protein motif:Cdd:COG3451 protein motif:TIGRFAMs:TIGR02746 protein motif:Pfam:PF12846.1"
/product="Type-IV secretion system protein TraC VirB4 components [ATPase mating pair formation]"
/"ID=B79_00408
/gene="traC"""
/protein_id="gnl|SC|B79_00408"
/codon_start=1
CDS complement(12930..13322)
/color=4
/locus_tag="B79_00409"
/note="TcpE family [mating pair formation], protein motif:Pfam:PF12648.1"
/product="TcpE family [mating pair formation]"
/"ID=B79_00409
/gene="tcpE"""
/protein_id="gnl|SC|B79_00409"
/codon_start=1
CDS complement(13411..13908)
/color=4
/locus_tag="B79_00410"
/note="Antirestriction protein ArdA, protein motif:Cdd:COG4734 protein motif:Pfam:PF07275.5"
/product="Antirestriction protein ArdA [anti-ristriction modification]"
/"ID=B79_00410
/gene="ardA"""
/protein_id="gnl|SC|B79_00410"
/codon_start=1
CDS complement(14025..14246)
/ID=B79_00411
/color=1
/locus_tag="B79_00411"
/note="hypothetical protein"
/product="hypothetical protein"
/protein_id="gnl|SC|B79_00411"
/codon_start=1
CDS complement(14289..15494)

/color=10
/locus_tag="B79_00412"
/note="Replication initiation protein RepA, protein motif:TIGRFAMs:TIGR03070 protein motif:Pfam:PF02486.13"
/product="Replication initiation protein RepA [replication]"
/"ID=B79_00412
/gene="repA"""
/protein_id="gnl|SC|B79_00412"
/codon_start=1

CDS complement(15672..17057)
/color=4
/locus_tag="B79_00413"
/note="DNA translocase FtsK DNA segregation ATPase FtsK/SpoIIIE and related proteins, similar to AA sequence:UniProtKB:P64165 protein motif:Cdd:COG1674 protein motif:TIGRFAMs:TIGR03924 protein motif:Pfam:PF01580.12"
/product="DNA translocase FtsK DNA segregation ATPase FtsK/SpoIIIE and related proteins [ATPase coupling protein]"
/"ID=B79_00413
/gene="ftsK"""
/protein_id="gnl|SC|B79_00413"
/codon_start=1

CDS complement(17086..17469)
/color=1
/locus_tag="B79_00414"
/note="YdcP family protein Bacterial protein of unknown function DUF961, protein motif:Pfam:PF06125.5"
/product="YdcP family protein Bacterial protein of unknown function DUF961"
/"ID=B79_00414
/protein_id="gnl|SC|B79_00414"""
/codon_start=1

CDS complement(17488..17802)
/color=1
/locus_tag="B79_00415"
/note="YdcP family protein Bacterial protein of unknown function DUF961, protein motif:Pfam:PF06125.5"
/product="YdcP family protein Bacterial protein of unknown function DUF961"
/"ID=B79_00415
/protein_id="gnl|SC|B79_00415""""

/codon_start=1
CDS complement(18155..18181)
/color=1
/locus_tag="B79_00416"
/note="Partial multidrug transporter MatE MATE efflux
family protein, similar to AA sequence:UniProtKB:Q2G140
protein motif:CLUSTERS:PRK09575 protein
motif:TIGRFAMs:TIGR00797 protein motif:Pfam:PF01554.12"
/product="Partial multidrug transporter MatE MATE efflux
family protein [transporter]"
/"ID=B79_00416
/protein_id="gnl|SC|B79_00416"""
/codon_start=1
BASE COUNT 5171 a 3776 c 3249 g 5985 t
ORIGIN
1 gcgtttttta atagtgaaac gtatactctc ttcctttctt tttctttcac tgattgaacc
61 ttccatgatt acccagaaaa gctgattttt gataattttt catcaaaaca aagtataaat
121 ttctaattat ctttttatat tttcttaaat gctcgtaaag ccttattcta tgtgctttcg
181 agtattttta ctgtaggaag atacttcacg tttctttgca tatttcctca tgtcttagct
241 gtcagaagtg gtaaataagt agtaaattca tttgtactac taagcaacaa gacgctcctg
301 ttgcttctct ttattcaagc gtttcatttc tgccattgca gaatcgaatg ttgcatgtgc
361 gtaatagttc agcgtcatgg ctatattagc atgtcccata atgtactgta atgcctttgg
421 attcattcct gcatttgcat agttggtaca gaatgtatgt cgcaaactat gtggagtgat
481 gtgtggcaat ttatcctcgt tatacttatt gtatttctta acaagacctt tcatcatgcc
541 gttgtaatca cttgccactt ttggatagtt ctttctatta agaaagagga aatcactata
601 tccatcaatc tcaacacgct tatcattctt tcgattcgct aacactcgct taaatgcttg
661 ataggcttct tcaaccatag gaacttgacg ttcgccactt ttggtctttg gtgtttcaat
721 gtagtaccca atttcagtat ctctcaatag ctgatggtct atattgacaa gacgattctc
781 aaaatctaaa tctggaagtg tcaaaccacc aaactctgaa atacgaagac ctgtttttaa
841 gagtatcaga atttcatcat aatttttgct gtaggtttta tcagcttttg caaaggctaa
901 cagtttttct tcctgttctt ctgttagtac ggtcttaggg acagtatcat catcaagaac
961 tgctttcagt tgaaagtcaa atggattctt ccgaacacaa tcatcttgta tagcaatata
1021 gaatgaagcc tttaaagaac gtttgtagtt attgatggtt tgataagcat aaccattttc
1081 actcattcta atagcccatt ctttagcgtc tgatggctta atactgtcaa tacttcttac
1141 acctaacttg tctttcttca aaatatccat aagatatttg cgtccagttt cagtgttttt
1201 tctaaccttt ggtctttgag cgttctgttt tgcgtaaagc tggcagagtg tcattttctt
1261 tcctacaaca tcaataccat catgaatgtc tttctgtaac tctgcgattt tctctctaag
1321 tgagatacaa tcacgctttc ctgctggtac tcggtctgta gccacaagtt tccacgagta
1381 aacaaattgc ggttctccaa atgaatctat atatttgtat aagtatcttc cgtcttttcg
1441 ttggctctct ccagtcttta agattcgacc tttattgtca cgtctttttt ctgacatggc
1501 atttgctcct ttcctttatg gaaagagcct tgatacgact taatactatt ttatcatata
1561 caagaccctt tggcgacgct agattgcgtc caatgtatct ataatttttt caaattgttt
1621 tcgtttaatc tgaatacgat tgccattcat aatcagccaa tttgcatttt tattttcctc
1681 tgccaagcgt cgtagcttgt tttcgccaat acgaaaatat tttgacgctt cttcaatggt

1741 tagggtataa cgttcccaaa taggaatgtc agtctgcttc ataaaatcct cctttccaaa
1801 tcacttattt ggatttcata aaagttgttt taccagcaat cgaacagctt tagcaaagct
1861 cacgggagtt ccacccctgc atggttctca tgtagccata ctcattgcct gcgacggttt
1921 tatcacgctc ggactattga ctgtatggga gtatcattat cacgataaga atgtcgttgc
1981 aggcaatcct gctaaagatt gcttctcgga tcactaacat gaatcgctcg ctatctttat
2041 aagataggtc atggcggtta gttccgttgg ctcttttctt atcgaaacgt attcgattac
2101 ttttattcag ttttcaaaga acaatggctc gttagcctat caaaacacat tgaaagctca
2161 atatgctttg gtggaataac aaacctccct gttcgggaag cgtggaatgg tttagcacgc
2221 ttccacgaaa ggagagagga tattacttaa tttcaaatga caaaatcttt gtaatcagtc
2281 tggtttccat tcttccacgt aagacttcat caacgaccat actttgattg ccatattcat
2341 ctttcataag tcgtagggaa cgcttcgtta tgtaccctct gtaatgatgt agaatctggt
2401 taatcgcttc ggtatcgcca tctgttgcct ttacaatgag aggaaaggga atcataggat
2461 attgtgtttt cattcttcaa attcctccat aaacttttta attaaggcta gtccactggt
2521 tctatgccga tagacagtag aacggttcaa tttcaacagg tctgcaattt ctgaatcgct
2581 catgtccata aagtaaaaca gcagtagaat ttcacgtttc ttgtctggca actcacgtaa
2641 tgcttcactc aacaaatcat tttcaacgcc tactgataac ccattgagtg taaaaatctg
2701 aaagtcagtt gaatagttat ctgttgtcgc aaactggcta acaagataat cgccaacatc
2761 cgaaaaggac acctcacgct ttgcaatcct tgaaagataa agcatataat tctttcgctc
2821 gtcttccata gcacgtttac agatatagtc aaactgattt tctattgtgg tctgaaaaga
2881 agatggtttc atgtttctca cccccttttct gtctaggaaa ggaagtgagc cttgctcgtt
2941 tatctccttt cactcttagt cccaatgtga aagggggatt tgttgcatta ctgataaata
3001 aactttgtaa aaaagttctg aatagccaaa aaagcatata aacagattta tttctctgtt
3061 tacatgcttc tgttattcta tctatatgat ttataaaacc acattggtgg acgtacttat
3121 ctattgcaga tagacgactt tttttgacaa gaacccaatg taaggaaatt tattgtatat
3181 gatgtacttc atggcgacgt tgacctccaa caaaccgcca tttggaagta atatacaata
3241 ttttaacagc gtaaatagca ctaccatata acggtttttt ttattggcgt ttagtagtgc
3301 tttttattaa atataaacct ataaaccata taacacgttt ttctatacct gtttttaatt
3361 cagtaggaac aataaaatgt atagaggtgg tctactatgc gtaaaaaaga agataaatat
3421 gattttagag cctttggttt agccattaaa gaagctcgat tgaaacgagg tttaactcgt
3481 gaacaagtgg gagcattgat tgaaattgac ccacggtact taactaatat tgaaaataaa
3541 gggcaacacc ccagcataca agttctttat gaccttgtat cgttacttca tgtttccgtt
3601 gatgaatttt tcttacctgc taataacttg gtaaaaagca cccgacgatt acagatagag
3661 aaatacatgg atagctttac agacaaagaa ctatccttaa tggaatcttt agccagcggt
3721 atcaacgaag caagaaacat cgaagactaa ttaaaagaat ccatacataa cggaaagagc
3781 cgataaaatg agattgtatt aatctcattt tatcggctct gcgtctttgc gtctggctct
3841 gtaatcacag ttactttgaa ctgctttatt tcaattaaat tttcttgtct gcatttcgga
3901 caatagaggg ggaatttttt taattcagta tcttccctta tctttaatcg tgttttattt
3961 ccacatacag gacacaatat ccacttgtag tttataataa ctatctcctc ctttacactt
4021 taattcaaat ctttattaaa aaatatttca tcttatttaa caagaaacca tatttatata
4081 acaacataaa atacactaag ttattttatt gaacatatat cgtactttat ctatccgact
4141 atttggacga cggggctggc aaacaggttc accggtagta acatggtacc cttttaactc
4201 tgttaaacaa acactacgtc catttgtaaa gaaagttaaa tcactacgat attcttgaat
4261 acaccgagca gggatttctc cactaagaat gacctcatta tttttcaatt gagtgtctac
4321 gatgttcgca caatatttag gagcatcgtt gtatgctcgt gaaagatatt cctgtggcgc

4381 ataaatttta aaactaagat atggctctaa caattctgtt ccagcttttt ttaagacttg
4441 ttccaataca ataggagcaa gcatccgaaa atctgctggg gtactaacag ggctatagta
4501 taagccatac ttaaaacaga ttttacaatc cgtcacattc caaccatata atccttgttc
4561 gcaaccatag cgtatccctt ccataactgc attttgaaat gattgattta agtatccaag
4621 agaaaccgag ctctcatact gcattccact tcccaacgga agcggtgata cagataaacc
4681 aatggaagcc cagaaaggat ttggcggcac ttcgatgtga atggtatatt ctgcattttt
4741 taacggtctc tccatataaa tgactgtagg ctcttttagt tctatctcca catgatactt
4801 ttcttgcaac agtgcactaa tcacttccat ttgtactttc cctaagaaag aaagtataat
4861 ttcatgtgtc gtagaatcca cgtaatatcg tagaagcgga tcactatctg agatttccaa
4921 aagggcatca agcaacattt ctctctgttc aggtttactc ggttcaacag ttgtttgtag
4981 tagagggtgc ggattttcaa tcttttttct ctgtggcaat agttttgtat ctccaagaac
5041 actatttaac ttcaaaaact cattttgcaa aataacaatt tctccagaat aagctctatc
5101 aatcttacat aattcaccat ttattgaagt atacatttct gtaactttta tttttctttt
5161 ttctgatact ctaaccgaat ctcgtaaatg tagtactcca ctataaaggc gtatatatgc
5221 aagacgttgt ctttttttttg tatattcaat tttgaaaaca tttccgcaaa gttcagacgg
5281 acctcgatgt gttgatgaat aaaatttatt cgtaatcact tctataaggt tatcaatccc
5341 tatattgttt tttgcacttc cgtgataaac agggaacagg gaacaattat gaaatcttat
5401 gctttcctct tgttcgagtt ccaatgcttc taatgattta ccggacatat atttctctaa
5461 aaggtcatcg tttccctcta ttaccgtatc ccattgttca gattcggtaa agttcgtcac
5521 acacatatta ggatacagtt ctaccttctg tttgattaca atttcggcag aaagtttctc
5581 tttaatatcc tgataaaccg ttgataaatc aattccattt tggtcaatct tattgataaa
5641 aaagattgtg ggaatcccca ttttcctaag tgcatgaaat aatatacgag tttgtgcttg
5701 tacgccatct tttgcagaaa tcagtagaat tgccccatct aaaactgata atgaacgata
5761 tacttctgct aagaaatcca tatgtcctgg cgtgtctatg atgttcacct tcgtattttc
5821 ccactgaaaa gaggttattc ctgtctgaat tgtaattcct ctctgacgtt ctaaaagcgt
5881 attatccgtc ctcgttgtac ctttgtccac gcttcctaat tctgtaatcg ctccactgtt
5941 atataataag ctttctgtta aggtagtttt tcctgcatca acatgagcta aaactccaat
6001 attaataatt ttcatgtgat tttcctccat tcaaaaaccc aaaagggcat aaaaatccca
6061 gtgataaata cttttatcac tgggattttt atgcataacc ataggtatac aaagcataca
6121 gatattctct ggatacttta gaatcacatg ataaaggtat tcttaaactg ggtgcaaaaa
6181 actaagccct cctaaaaaag gacatctaat tatttgttcc cgctatcaaa ttgacagttt
6241 atttaagaat accttgccgc atatttatta actcctttta aatagtcact taaataatag
6301 cacgtaagag catatttgta aaggaatctc caattttttta tcaaaaagag tacatgatta
6361 caaagtatct gtaatcatgt accaatattt gttattttac aatcttccaa ttactcccgt
6421 tcttttcaag taccaaatca aattgagata cctgcgttgc tttggtctgc tggtcgatat
6481 actccactgt cagcgatacc gtgacttgat tatccttacg attgtgaata ggatttacca
6541 gttcttgaaa gatgtactct tttccgattg gttttaatat cccgtcattc acatagtagg
6601 aaagttcact ggctgtcgct gtaggataga gcttgaagaa cgtcgttaaa aactcattga
6661 tttcattggt tgtaatggaa tcaaccgtcc cctcactttc aatggctttt ggtttataac
6721 ttgatttctt aggtatgttg gtaatggtcg gattcttaac cagtaccata tttccagaac
6781 catctacata gacactcact atataagcag agtggacggt ctttgtattt tctccctctg
6841 taatgagctg gtctacactg taggttacat taaactcatt gtcgccagtt ggctctaccg
6901 tccatatctg aaatcctctt acagaagacg atacaggaat atctttgcgt actgtatcaa
6961 cattgagagc ttgaagttca tctgtcagat agccttttag actttccatt cgattatcaa

7021 tggacttatc ggattgctcc catgaatagt agactttcgc aaagttctct acaaaatttt
7081 ctacatgatg agtatcaacg tattcctttt ctatgatagt tgtttcgtga atagtatgag
7141 tatctatagc tgtaaagtgc ttgaatatcg caaagctgaa actaagccct aaaagtaccc
7201 acaaggcaat cacaaccttt ttatgaggat tgaccttata gacacgaggt ttcttttcct
7261 ttggtatctg tttttcttta ttctgatttt ttctaaattt catcattaaa tcttcctttc
7321 tcattgtttg attcgtcctg ctcccactaa atgctgttgc cagtaggggc ttgttaagtc
7381 ggcataaccg attgggtcgc ctgcatgaaa catacggtta ttgccaaggt atatcccaac
7441 atgagtaata taagagccag cgttataggt agaatgaaag aaaaccaaat cgccagcttg
7501 tgcttccgat agtgggatat gctgggtcac atcatattgc tgttgtgcgg ttcgtggtaa
7561 gttaattcca gcttttccat acgtccattg tgtcagtccg ctacaatcaa aagaagtagt
7621 cggggaagct ccaccgtaaa cgtatcgcca gccctcatat ttcagtgctt cgtccatgat
7681 ggcttgtacc gtatcatcat caaactctgt tgtgacaaga tactgcgtta ccagttgcac
7741 ataaaacata ttgccatagt tgtatcgcca gcccccattg ataggtatgg ctatgggatt
7801 ggggtaagac actttttcgc cacctgaata ctcttttgag aaactttgag ccagttcaaa
7861 ggtatattta tttccacgat tagccacata ccctaagaaa ccaccaccat aattgtagga
7921 ctggataacc gattctaaat ctacactgag cctttcgcta ctggctaata attcactgaa
7981 atacttcaca ccttgcttaa tggattcttc tgtactcaat gaattaggtg gaagaccgag
8041 ggattccgag gactgcataa catcttccgc agtaccgccc gattccacct gtataatcgc
8101 aagaagtatg ttgacatatt cttcaacgcc atattctttg gcatattttt ctaccatagg
8161 cttatgagcc agcacttctg cggaaacatt cacacctcca taatgaatat tggaaattcc
8221 gctgtcctgt tcatctgaaa ataaaatggc aacaaacaga agcagtgaga agaccatcaa
8281 gaataatcca gaaccaccaa tcactaaagt tttcaacttc atggtttctt accgactttc
8341 ttaatggtgg cggttttgat tggtggtcta cttcttgtat tttgtagtgg tactctttga
8401 acagtagacg gacgttcttt tgtgattgga cgttgtgaag ttctatctgc tgtagtggtt
8461 gaagttgctg gcttttgaac ggtttttttct tgaacggtat tgccttggcg ttccactttt
8521 ggacttgaaa aatcggactt aactgctgga cgctcttgtt tggcttgttg agattcctta
8581 tatgaagtct gaatattaga ctgttttgag gtctgttcat catgatattg ttcttgtctt
8641 gtagtcggtc tttcatgaac agaagaagca ggctgttttt tctgtttgac ctgttccatt
8701 tcagagcgac gcttcgcaat ggtttttcgc ctttgttcct gctgttcctt gcgtccactg
8761 gctctgtccg ctttggtttg agaaatacta ctggttaaat cacggacatt ctcttttact
8821 ttggattttc cttgatatac tgcatatctt gcattggtcg gcaaatcttt aacctgttct
8881 ttcaaaccac tagcagtgtc taccattctg tctttggtat cagctactgt accgatggtt
8941 tgaccgatac gttttccaag tgttgatttt tcctttccgt ctggtcggga gtgatctgct
9001 tgtgtccttg cagaactccc cgaacccgac tgtccttttt tacctgtaac aatggcagac
9061 ccagccccta gagtagtcat ggaacgtcca agtttccgct gtagacggtg catgtgagcg
9121 tgcataagca tacgaggttt tctcatcaca cgacttccca cactttgaga atcgttactc
9181 tgtagagaaa acatactcat taaatcgccc agcttgaagt agattcctgc aaaggtcaca
9241 atctgtagaa aagcaatcaa aaagaacgga taaccagccg ataaggtata gagcatggtt
9301 gaaatactaa atgctgtcgt aataatcaat gtgattccag ctcgtgtcaa aatggtatta
9361 aagagctttg ttatggctcg ttttgacata ccatcaaatg atggaatcat gcttaaaata
9421 aagctcacag gcagaaacat agcatagatg ataaaaagta cctgcgagaa aatcatgatt
9481 cctgttaata ggaatacaaa tatggaaatc ccaatattga agacaaatag gaagaagact
9541 gtacctaaac ggttaatggt ctttgtaatg gttagattgg tattgcttct gtcttcaatt
9601 tcttccgcaa caattttttc tctgtcttcg ccattgttgg aatctgggct ggtggagagc

9661 aggctttcca cacggtcaat accgatactt tcaatgtctg aactgttgta ttgaagcagt
9721 agccacggtt gctgaacctg tatggaaaac aggctatctc tgattaagtc cacgctgtcc
9781 ttgccttgac tatcggaatg gggcatgaca atcttcgtgc caagtgataa actggcatta
9841 ctgatgtctg atgaaaagtc attgattttt ttaatgtagt cgggagcgta ggcaataaag
9901 gaagccgata ggataaacac cagcacaaaa ttcataatgg catgaattgc ctttgtggtt
9961 tctctcttta tcagtcccgt ataggcaaca taaaccccaa gaaccaaaat caagagtaag
10021 aggaatccaa catagaaacc ctctgttgaa aatccgtttg cactcacacc agctaaggtc
10081 tgcatattct taccaatgga atctgctgta gcggaaatga agtctaagga ataggcttcc
10141 tgtactaagt aacctgtcgc attggaaaca tacaaactga ttgtccaaat aaaattggta
10201 atggcatata gtccatacat gacctgtttt ccaatcccgt ccgaccagtt ccacggaagc
10261 cagccccagc tattatccac ataaaaatcc agttgatagt tttcaagtgg gtatcggctg
10321 tattcatttg ccacattgac cgtatcatct accaagcccg cagcttgaac caccgttccc
10381 agcatggcta aaagaaaaat ggcaatcaca agtgtgaaag ccactgtcat tgccacttta
10441 cctagacgtt tcagcgtcca gtttgatttt attctgttta ctattgatgg tttcacattt
10501 acacctcttt tcgcacaggt ggtctggtat caaaggcatg gagcagttct tcaaatacag
10561 ggtggaactg tatcacaccg acacgaccat ataaatcact gataaggcat tgcccgtttt
10621 ccaaatcacg caatcgcttc tgattgtttt cgtcctctgg gtctacacca aaaaaggcta
10681 aggtcttttt aatctcgtta aggtcagtgg aacgaaatgc aaattttaag ccgaggttat
10741 ttttcagttt ttcatctaag aggtcgtctg tattttgggt cacgaaatat accccagcgt
10801 tcatagcacg accagcccga accagcttca tagatagtgt ttttccttgt gctacctgta
10861 aaaagctcca tgcttcgtct aaatctacaa tcttgaaaat gcttcggtct gtatggataa
10921 agtctaaagc aaaggtacta atgacaatca gcatagcaac ggataaaagc tccatagtgg
10981 tatattcctc aaaggaagtt tccttgtcgg gaagtaccaa gtccgcaacc tgtataatgt
11041 tcagttgttt ttctaagctg atagactgct ccacataacc attactgaat aataaatgtg
11101 caaagtcata gtctgtaaaa ctttcgatat ggtcggctat actggtactt agtggcgtat
11161 tctcaacccg taattcctca atcactttca tcaaccctcg tacttcacta ttggttactg
11221 cacgaatggc ttttctaagg attgggaagc gttccccatc acgagaggaa atccccgtaa
11281 ggaatgtcag aatatcaata gccagtgatt cagaatcttt gggatttttc ataatcacat
11341 aagggtcaag taagcctttg tttttctcat cagaagtcag agtgacgata ttgatttcat
11401 gggaaatctc tggcaaggtt tctttccatc tgccacgttc tgcttttggg tctacaatca
11461 ctgcttgtgc cccataaagc accgcataat agacgataag gttattcgca aaggatttac
11521 caccacccag cgaaccaaca aaagccgacg ctaacgcatt ggttactgaa cccttaaccc
11581 cttgactggc aagagcaggt ttcagataga cattgcgtcc agtatctaag ctgtagccaa
11641 cataaatccc ctcattttcc cccagcattt gagtagcacc aaaacctaaa ccagcgagga
11701 aatcagaggt cacgtattga atataatcat tcatataacg cttgctggca ggtaaaaatt
11761 cttcatgtaa gccgagcata tccccaaatg gtcgtaccag ttttacgctt aaatcgtcat
11821 aaaaatcttt cacttcatta caacgacgtt tgagttcgtc aagatcattt gctgataccc
11881 ttaccacata agacagcttg tacatagatt ccttgctttg gtctaaattg gtttccagct
11941 cattcacact ttccagagct tccgccacat tggagctggt ttcattatca ctttgccaag
12001 cgtggttatc caagtctttc agttctttct ttttattgcg gacagtagat agggctttac
12061 gattcgctac aatttccaca ttcattgacg tatcaatcgg gaatgtaaat tgctgttgct
12121 ggtagtagaa gatttcagag gacgggaagt ccagttctcc gacaatgctg ttaatggtaa
12181 agtaagctac atagacggtt tcatcttcct gctggatttt caaatatcgc tgtttttctt
12241 ccaccaaaca gcgagtaggc ttaatcaagt catagtattt aatcagcgtt tcattatcca

12301 gctttttctt tgatagatgg tactcatact cttcataggc agtgcctgtc tgtccgtaaa
12361 ggtgttcaat cagatagccg aagtcgtcct tatctaacct gcggattttg aaacgacgag
12421 agattttatt ttctaagagc ttttccatct tctgaaaacg caggatttca tcattactca
12481 tactaacaaa atcgcccatc agcttatggt tcacatcata gacaaaatca gacaaagcat
12541 tttttgcttc aacggtaaga cttttcatag aaaactcctg atcgttgaga agcaacttaa
12601 agccgataaa gaaacggtag ttcacttgat tttcgccaat catggatatt aaagcgtctg
12661 tctgttggtc gattttgtca taggcaaccg ctttgagctt gccagtgact tcatttttgg
12721 aacgctcttg tgcagaacgt atgctggatt ctgtactgat ttgtaaagca tgaattttgc
12781 catcacgatt ttgtgcgata agctgtctga aagaatcatg cacttgtatt ttctgttctg
12841 gacttagaaa tgagtaattg taaggaacaa gctcatagta agcataacat tccccgtctt
12901 tattccagac gagattgttt tcaatgtatt taattggata tgccataaaa ttcactccta
12961 actgctgtaa tggcttcttg tggctggttt ctgccaagcg ttactttttt tcctgcatag
13021 gtcagctttg gtcgcagtgc ataagcaatg acagacttca aaaatccata aggcttttta
13081 ccatcaaaag tttttgtaga cataaaccat gtgaaagcca caggaatccc aaagtatttg
13141 agaaatgctc cctctatcat ggaaagaggg ggcaagttgc caagtatcat cactgcaaag
13201 agtgacacga caaaccatgt catttgcgta aaggttatgg gaaacggaag tctaaaatca
13261 ttgatagaat acagtacctt ttccacagac cagatactgg tatagcttcg tattttcttc
13321 atgtaatcaa tcctttcata aaaaataggg gtagctgatt gagccacccc gtaaaataga
13381 aaatctgcca gtagtaatgt accgacagat ttaatagacg atttcaaaaa tcccatgatt
13441 ggttgagata aacgttcctg aaaggtctaa atcccgacca taggcttgat aatcaatata
13501 gttttgaaga ctagctggta cttcgcctaa agcacccgtt tcttcaatgt agtagcgtgc
13561 cacgtcatac atatcatcac aatcggaatg aatgataata tcctcttgat gttcgcttag
13621 ttcttcaatg cttgaaaaat gagtgagcag agcagatagc tccgattgta attcttcggg
13681 taattccgat accatttccc atagtcgatt gagttcgcca atggaagtgt attcgtcaac
13741 cgtaaagggt aactcgtagt catgaatggc gtattcctca tattcatcat tcaagccgat
13801 tttctctttg acttcctcaa agtcaatggg aaaggtaaac cacgcaccga ccaattcgcc
13861 ctcattgtat ttgcctaaat tcgcaatata gacttgcata tcgtccatat attcacgtcc
13921 tttctttgta gagattcaaa aatccctacc gcacttcgtt tggtgtacca ttcctttgcg
13981 gaacataaga aaaccactta tattccacaa aagaacggtt ttatttaagc accaataatg
14041 cgattgaata gctctagtaa aatgtctttt actccagcag cgttgaagac taagccaacc
14101 gcaataatcg caataattaa aaagccaatc agtttgctaa actcacgctt gaagccaaga
14161 tacaagccaa tcacaacgat tgctaaaagc accagtgatt gagcgtttga tagaaaccag
14221 ttataaaggt tttgtccaaa attcataaaa atgttctcct ctctatattc aatgaatttg
14281 tatttgagtt atttttttgt tgttatcacg tcctgttctt ttactgactg ttgcttcaaa
14341 atctgcttgt gtcggtctgt cagtttcgca tggtcgagaa tgtcttttac aacctgcgtc
14401 tggttgattt catcaagttt aatcgcaacc tttaaggtcg gggcaacttg atgagatagc
14461 cagttcagcg tcctttggaa ggagtaaggc tctggttttg tggttagttt taatcgttca
14521 cgattgttcc caataaacca agcccattct tcattcagtt tccaatcaga acgaggtttg
14581 gaatcgtctt tatctacaaa acggatatac cgattgataa ttttaaaggc ggtatgctct
14641 ggattgtcat agacgagtaa atcacggact gcataatagg cacgctcatt tttcaatcga
14701 atctcaaaac ggttttttac ttctgcgtct tcaatgggaa tatcattttt cttgtactgc
14761 tcgtagtcct tttcatagat acagaaataa acttcacttt gtaatgaacc gatatagagg
14821 gtgtttccca tacattcctt ttcctctttg cgtaccagtt cgccactgcg atagctttta
14881 aaactgcgga agacggagat acattcttcc tgttggcact tttcagtgag tacagggata

14941 tttaaaatcc ctgtcttatc gttaatggca aggtcaaggc gtttcatcac accgccagcc
15001 accaaaacgt ccataaagaa ctcataccag cttctttgtt gtgccagaag atagctttca
15061 aattgtctgc acccacgacc tttcaattcc accagaactc ctttgtccag ttcatgggag
15121 caaaggacga atatgtcgcc taaagcataa tgctctgaat aagaatagaa accatagtcc
15181 tcatgaagaa aataggacag tttcagttgt aagatgtttt cgaccacctg ctgtacgtct
15241 gttgtcggaa agcgaattct tacataatca aacagcattt caaggggagc gtcgggattg
15301 aagcgttcca gagcttccca aagggactgc tgtaaatcct ctgatggctt gacttttcct
15361 gtttcaatat cgcttagata ctgccttgta ataccagtcg caacagctaa acggttttga
15421 gatagtccat aagccaagcg tttttctttt aaatgctgta accaagtttg ttcattcagt
15481 aaaaatccct ccaatcaaaa aggcgtatgt caacttttaa agcccatttg acatacgctg
15541 aaatttgta aatcccttgt aaccaaagga ttttctaatg tttttttgac tgtttcctgt
15601 cgatttgtac ccccctgtta gatacggggg gttaagtgct ggcgtggcta ttgccacacc
15661 agccagcaag atcagtccac acctgcgact tccgcttcgc acgtcgcctg cgtggactgt
15721 ctgctgttgg ataacttttt aatttcctcc aagaaatcat atccttttgg tacaagggga
15781 gtataaaact ctgatatgac acttgttcct acatcaacat agccacgacc tttgattcgc
15841 tttaagaaga aatccttttg tacgtcactg ccaaacatca tgccatagcc catttcagac
15901 atacgaccta aagccactct gaaattaaac tgatcacgga ttccgtcgcc taaatatttt
15961 gcgtctggac gttgacaagc cagtattaga aagaagccag cttgacgacc taacatgaca
16021 atctgtttca gcttattcat aactgcggtg ttttcttttg ttcccagcat ttccatgaaa
16081 gcgacgtatt catcaaagat taagaagtgt gccgggagac ctaagtaagc ataatttttg
16141 ccagtcttat agttcttcat ctgcttcatt tcctcactac gtttcatcat ttcttcatag
16201 aatgtttcaa tgcaagaaag caagtcttct tttctatagt agacatttgc catcacagaa
16261 cctaagtccg caagatcagc atttttcggg tcaagaatat acagttttga atctgtatga
16321 agcaaggctt caatcagtgt cagtataaag taagttttac cgccacctgt accaccagca
16381 atcaacatat gagggagctt atcatattcc caccatacgt ttttcattaa gcgaagttta
16441 ccatctttag cttctacttc atcaatagaa atacgactgg ctatggtgtc atagagcaaa
16501 gtatattcca cataggaatc ctttaactct ttatccgtca gctcacagta caagccactc
16561 tctaatttct tttccaagtg taagagttgg tcttgatatt ttcccagcgt gatttccacc
16621 cgtatctgta tcaagccatt tttaagtcga taatacattt tagggaagta ggttatcttt
16681 tcctttgtac gaccagcact atctttaaag aaaccctctg ttttgacctg ttcagattca
16741 taccacttgt tttcaagtat catctttgcc agtttttgac ggtggtaaag ttgtttaacc
16801 gtatcatagc gaacccgttt gaatacaaac gctaccagca agcagataag aattgcgaca
16861 ctgaaactga taattaaata gggaatgtca atcttatctg cttgtgatag gttaaaatcc
16921 tgccagttga tctgctggat tgtcttcaca tgaaacagtc cgacaaccag caggaaaaca
16981 ggcaggagtg acgctatcgt aaaatgaaag actaaatctt taccagatgg gcgaatcctt
17041 ttaccacgct gtttcatgcg aaaaagtctc ctttctacct agcgactatt tgtcttgtgt
17101 cggttctttc tttgcttgtg gttgagcttt gaatgaacta gaatcctttg tcagcacaat
17161 atcgtctgcc ttgatatacc agtcaacatc tgctccttga taggtggcag tagcaacggt
17221 gtccgcaatg ggattgataa gttccacccg tgcgttataa tcaaactctt tcaaaggcac
17281 gctggcagga atacttactt gaatcatgcg tccttgtcct ttggatttta agtcataggt
17341 acgttccttg atttcatctg aaaccgaccc gtcttcattt tggattctca cttcacgacg
17401 tagagcagag aatttcaatt ctccaaaagt cgtgtcttta tctaatacaa tgccatttgc
17461 taatctcatc atttttcctc tctttcttta ttcttttatc atgtcgtcag catgtaaaag
17521 gtaatttgta aaaccacgag tgccgatttt gtagccctct gcggtaatac gtggattgac

```
   17581 taacttcaca cgttcctcaa agccgaaatg tttttcgcca gcttcagcag gaagcaccac
   17641 cacaatatca tctgctcttt gaacatcaga atagagatta tagcttcttg ataagacagt
   17701 tagccgtccg ttgattcttc gctgaacgac tttatcctcg ccagcaaatt ctaaattgcc
   17761 gaatgttttt tccatgttgg gaatcacaaa tttaagttcc atatttttac ctatcctttc
   17821 tttttattg gctgaatgaa tgtttgatgg tcttaaagag tggggaacga ccttttgatt
   17881 cttgattttt tgttttcata agttcacttc ctttcaaaat cgggtaaaaa aatagacacc
   17941 tcattttttg aagtgtctac ctattaaata ttcaaatttt attggaagta tctttatatc
   18001 ttcacttttc aaggataaat cgtcgtatca aagctcattc ataagtagta aattagtagt
   18061 aaattgagtg gttttgacct tgataaagtg tgataagtcc agtttttatg cggataacta
   18121 gatttttatg ctatttttct atataaaaat aattttatgc gctcttcttt tcccttacaa
   18181 t
//
DEFINITION     Integrative and conjugative element ICE_3
     fasta_record    1..80990
                     /ID=D90|SC|contig000002
                     /colour=5
     CDS             complement(366..1829)
                     /color=9
                     /locus_tag="D90_02917"
                     /note="Tyrosine recombinase, protein
                     motif:CLUSTERS:PRK00236protein motif:Cdd:COG4974protein
                     motif:TIGRFAMs:TIGR02224protein motif:Pfam:PF00589.16"
                     /product="Tyrosine recombinase [recombinase]"
                     /"ID=D90_02917
                     /protein_id="gnl|SC|D90_02917"""
                     /codon_start=1
     CDS             complement(1842..2048)
                     /color=9
                     /locus_tag="D90_02918"
                     /note="Excisionase family DNA binding domain-containing
                     protein, protein motif:TIGRFAMs:TIGR01764protein
                     motif:Pfam:PF12728.1"
                     /product="Excisionase family DNA binding domain-containing
                     protein [excisionase]"
                     /"ID=D90_02918
                     /protein_id="gnl|SC|D90_02918"""
                     /codon_start=1
     CDS             complement(2038..2739)
                     /color=9
                     /locus_tag="D90_02919"
                     /note="Excisionase family DNA binding domain-containing
                     protein, protein motif:TIGRFAMs:TIGR01764 protein
                     motif:Pfam:PF12728.1"
                     /product="Excisionase family DNA binding domain-containing
```

protein [excisionase]"
/"ID=D90_02919
/protein_id="gnl|SC|D90_02919""""
/codon_start=1
CDS complement(2726..2977)
/color=9
/locus_tag="D90_02920"
/note="Excisionase family DNA binding domain-containing protein, protein motif:TIGRFAMs:TIGR01764protein motif:Pfam:PF12728.1"
/product="Excisionase family DNA binding domain-containing protein [excisionase]"
/"ID=D90_02920
/protein_id="gnl|SC|D90_02920""""
/codon_start=1
CDS complement(3084..3344)
/color=9
/locus_tag="D90_02921"
/note="Excisionase family DNA binding domain-containing protein, protein motif:TIGRFAMs:TIGR01764protein motif:Pfam:PF12728.1"
/product="Excisionase family DNA binding domain-containing protein [excisionase]"
/"ID=D90_02921
/protein_id="gnl|SC|D90_02921""""
/codon_start=1
CDS complement(3337..3915)
/color=8
/locus_tag="D90_02922"
/note="type II toxin-antitoxin system PemK/MazF family toxin MazF, similar to AA sequence:UniProtKB:P96622protein motif:Pfam:PF02452.11"
/product="type II toxin-antitoxin system PemK/MazF family toxin MazF [toxin-antitoxin]"
/"ID=D90_02922
/protein_id="gnl|SC|D90_02922""""
/gene="mazF"
/codon_start=1
CDS complement(4016..4555)
/color=7
/locus_tag="D90_02923"
/note="RNA polymerase sigma factor Sigma-70, protein motif:CLUSTERS:PRK06759protein motif:TIGRFAMs:TIGR02937"
/product="RNA polymerase sigma factor Sigma-70"

/"ID=D90_02923
/protein_id="gnl|SC|D90_02923"""
/codon_start=1
CDS complement(4800..4997)
/ID=D90_02924
/color=1
/locus_tag="D90_02924"
/note="hypothetical protein"
/product="hypothetical protein"
/protein_id="gnl|SC|D90_02924"
/codon_start=1
CDS complement(4994..5725)
/ID=D90_02925
/color=8
/locus_tag="D90_02925"
/note="ImmA/IrrE family metallo-endopeptidase ImmA"
/product="ImmA/IrrE family metallo-endopeptidase ImmA
[pepidase]"
/gene="immA"
/protein_id="gnl|SC|D90_02925"
/codon_start=1
CDS complement(5745..5996)
/ID=D90_02926
/color=1
/locus_tag="D90_02926"
/note="hypothetical protein"
/product="hypothetical protein"
/protein_id="gnl|SC|D90_02926"
/codon_start=1
CDS complement(5999..6298)
/ID=D90_02927
/color=1
/locus_tag="D90_02927"
/note="hypothetical protein"
/product="hypothetical protein"
/protein_id="gnl|SC|D90_02927"
/codon_start=1
CDS complement(6311..7546)
/color=10
/locus_tag="D90_02928"
/eC_number=2.7.7.7
/note="DNA polymerase IV, similar to AA
sequence:UniProtKB:Q47155protein
motif:CLUSTERS:PRK03103protein motif:Cdd:COG0389protein

motif:Pfam:PF00817.14"
/product="DNA polymerase IV [replication]"
/"ID=D90_02928
/protein_id="gnl|SC|D90_02928""""
/gene="dinB"
/codon_start=1
CDS 7697..8107
/color=7
/locus_tag="D90_02929"
/note="helix-turn-helix transcriptional regulator ImmR, similar to AA sequence:UniProtKB:P96631protein motif:CLUSTERS:PRK09706protein motif:TIGRFAMs:TIGR03070protein motif:Pfam:PF12844.1"
/product="helix-turn-helix transcriptional regulator ImmR [transcriptional regulator]"
/"ID=D90_02929
/gene="immR""""
/protein_id="gnl|SC|D90_02929"
/codon_start=1
CDS 8208..8741
/color=8
/locus_tag="D90_02930"
/note="ImmA/IrrE family metallo-endopeptidase ImmA, protein motif:Pfam:PF06114.7"
/product="ImmA/IrrE family metallo-endopeptidase ImmA [pepidase]"
/"ID=D90_02930
/gene="immA""""
/protein_id="gnl|SC|D90_02930"
/codon_start=1
CDS complement(8783..9613)
/color=8
/locus_tag="D90_02931"
/note="GIY-YIG nuclease family protein, protein motif:TIGRFAMs:TIGR01453"
/product="GIY-YIG nuclease family protein [nuclease]"
/"ID=D90_02931
/protein_id="gnl|SC|D90_02931""""
/codon_start=1
CDS complement(9806..10192)
/color=10
/locus_tag="D90_02932"
/note="ParB/RepB/Spo0J family partition protein ParB-like nuclease domain, similar to AA

sequence:UniProtKB:Q83AH2protein
motif:CLUSTERS:PRK13832protein
motif:TIGRFAMs:TIGR00180protein motif:Pfam:PF02195.12"
/product="ParB/RepB/Spo0J family partition protein
ParB-like nuclease domain [partitioning system]"
/"ID=D90_02932
/protein_id="gnl|SC|D90_02932"""
/gene="parB"
/codon_start=1
CDS complement(10201..11166)
/color=9
/locus_tag="D90_02933"
/note="Plasmid recombination enzyme Mob_pre, protein
motif:Pfam:PF01076.13"
/product="Plasmid recombination enzyme Mob_pre"
/"ID=D90_02933
/protein_id="gnl|SC|D90_02933"""
/codon_start=1
CDS complement(11691..12716)
/ID=D90_02934
/color=10
/locus_tag="D90_02934"
/note="Replication initiation protein RepA"
/product="Replication initiation protein RepA
[replication]"
/protein_id="gnl|SC|D90_02934"
/gene="repA"
/codon_start=1
CDS 13022..13477
/ID=D90_02935
/color=1
/locus_tag="D90_02935"
/note="hypothetical protein"
/product="hypothetical protein"
/protein_id="gnl|SC|D90_02935"
/codon_start=1
CDS complement(13429..13854)
/color=7
/locus_tag="D90_02936"
/note="RNA polymerase sigma factor Sigma-70 region 4,
protein motif:CLUSTERS:PRK12522protein
motif:TIGRFAMs:TIGR02937"
/product="RNA polymerase sigma factor Sigma-70 region 4
[transcriptional regulator]"

/"ID=D90_02936
/protein_id="gnl|SC|D90_02936"""
/codon_start=1
CDS complement(13949..14461)
/color=7
/locus_tag="D90_02937"
/note="RNA polymerase sigma factor Sigma-70 region 4, protein motif:CLUSTERS:PRK08583protein motif:TIGRFAMs:TIGR02937protein motif:Pfam:PF04545.10"
/product="RNA polymerase sigma factor Sigma-70 region 4 [transcriptional regulator]"
/"ID=D90_02937
/protein_id="gnl|SC|D90_02937"""
/codon_start=1
CDS complement(15174..17093)
/color=2
/locus_tag="D90_02938"
/note="Tetracycline resistance protein tetM from transposon Tn916, similar to AA sequence:UniProtKB:P21598protein motif:CLUSTERS:PRK13351protein motif:Cdd:COG0050protein motif:TIGRFAMs:TIGR00484protein motif:Pfam:PF00009.21"
/product="Tetracycline resistance protein tetM from transposon Tn916 [antibiotic resistance]"
/"ID=D90_02938
/protein_id="gnl|SC|D90_02938"""
/gene="tetM"
/codon_start=1
CDS complement(17460..17834)
/ID=D90_02939
/color=9
/locus_tag="D90_02939"
/note="Transposon-encoded TnpV family protein"
/product="Transposon-encoded TnpV family protein"
/gene="tnpV"
/protein_id="gnl|SC|D90_02939"
/codon_start=1
CDS complement(17815..18825)
/ID=D90_02940
/color=10
/locus_tag="D90_02940"
/note="DEAD/DEAH box helicase family protein"
/product="DEAD/DEAH box helicase family protein [helicase]"

/protein_id="gnl|SC|D90_02940"
/codon_start=1
CDS 18851..19309
/ID=D90_02941
/color=1
/locus_tag="D90_02941"
/note="hypothetical protein"
/product="hypothetical protein"
/protein_id="gnl|SC|D90_02941"
/codon_start=1
CDS complement(19348..21057)
/ID=D90_02942
/color=8
/locus_tag="D90_02942"
/note="Cysteine histidine-dependent amidohydrolase/peptidase CHAP domain-containing protein conjugative transposon protein"
/product="Cysteine histidine-dependent amidohydrolase/peptidase %28CHAP%29 domain-containing protein conjugative transposon protein"
/protein_id="gnl|SC|D90_02942"
/codon_start=1
CDS complement(21098..21868)
/color=8
/locus_tag="D90_02943"
/note="Class B sortase, protein motif:Cdd:COG3764protein motif:TIGRFAMs:TIGR03064protein motif:Pfam:PF04203.7"
/product="Class B sortase [transpeptidase]"
/"ID=D90_02943
/protein_id="gnl|SC|D90_02943""""
/codon_start=1
CDS complement(21906..24239)
/color=4
/locus_tag="D90_02944"
/note="Type IV secretion system protein TraC VirB4 components, protein motif:Cdd:COG3451protein motif:TIGRFAMs:TIGR02746protein motif:Pfam:PF12846.1"
/product="Type IV secretion system protein TraC VirB4 components [ATPase mating pair formation]"
/"ID=D90_02944
/gene="traC""""
/protein_id="gnl|SC|D90_02944"
/codon_start=1
CDS complement(24232..24597)

/color=4
/locus_tag="D90_02945"
/note="PrgI family protein Type IV secretion system,
protein motif:Pfam:PF12666.1"
/product="PrgI family protein Type IV secretion system"
/"ID=D90_02945
/protein_id="gnl|SC|D90_02945""""
/codon_start=1
CDS complement(24597..24782)
/ID=D90_02946
/color=1
/locus_tag="D90_02946"
/note="hypothetical protein"
/product="hypothetical protein"
/protein_id="gnl|SC|D90_02946"
/codon_start=1
CDS complement(24797..25633)
/ID=D90_02947
/color=1
/locus_tag="D90_02947"
/note="hypothetical protein"
/product="hypothetical protein"
/protein_id="gnl|SC|D90_02947"
/codon_start=1
CDS complement(25707..26066)
/ID=D90_02948
/color=8
/locus_tag="D90_02948"
/note="Glutamyl-tRNA amidotransferase subunit A"
/product="Glutamyl-tRNA amidotransferase subunit A
[amidase]"
/protein_id="gnl|SC|D90_02948"
/codon_start=1
CDS complement(26081..27877)
/color=4
/locus_tag="D90_02949"
/note="Type IV secretion system protein TraG VirD4
components, protein motif:CLUSTERS:PRK13897protein
motif:Cdd:COG3505protein motif:TIGRFAMs:TIGR02767protein
motif:Pfam:PF02534.8"
/product="Type IV secretion system protein TraG VirD4
components [coupling protein]"
/"ID=D90_02949
/gene="traG""""

/protein_id="gnl|SC|D90_02949"
/codon_start=1
CDS complement(27864..28067)
/ID=D90_02950
/color=1
/locus_tag="D90_02950"
/note="hypothetical protein"
/product="hypothetical protein"
/protein_id="gnl|SC|D90_02950"
/codon_start=1
CDS complement(28077..28961)
/ID=D90_02951
/color=7
/locus_tag="D90_02951"
/note="LtrC-like late transcriptional regulators protein"
/product="LtrC-like late transcriptional regulators
protein [transcriptional regulator]"
/protein_id="gnl|SC|D90_02951"
/codon_start=1
CDS complement(28967..29761)
/color=1
/locus_tag="D90_02952"
/note="PcfB family protein, protein motif:Pfam:PF12687.1"
/product="PcfB family protein"
/"ID=D90_02952
/protein_id="gnl|SC|D90_02952""""
/codon_start=1
CDS complement(29745..31115)
/color=4
/locus_tag="D90_02953"
/note="Relaxase/Mobilisation nuclease domain, protein
motif:Pfam:PF03432.8"
/product="Relaxase/Mobilisation nuclease domain
[relaxase]"
/"ID=D90_02953
/protein_id="gnl|SC|D90_02953""""
/codon_start=1
CDS complement(31106..31447)
/color=4
/locus_tag="D90_02954"
/note="Bacterial mobilisation relaxosome protein MobC,
protein motif:Pfam:PF05713.5"
/product="Bacterial mobilisation relaxosome protein MobC
[relaxosome]"

/"ID=D90_02954
/gene="mobC"""
/protein_id="gnl|SC|D90_02954"
/codon_start=1
CDS complement(31705..32622)
/color=1
/locus_tag="D90_02955"
/note="DUF3846 and DUF4314 domain-containing protein, protein motif:Pfam:PF12957.1"
/product="DUF3846 and DUF4314 domain-containing protein"
/"ID=D90_02955
/protein_id="gnl|SC|D90_02955"""
/codon_start=1
CDS complement(32619..33572)
/color=10
/locus_tag="D90_02956"
/note="DUF3991 and TOPRIM domain-containing protein, protein motif:Cdd:COG0358"
/product="DUF3991 and TOPRIM domain-containing protein [replication]"
/"ID=D90_02956
/protein_id="gnl|SC|D90_02956"""
/codon_start=1
CDS complement(33588..34019)
/ID=D90_02957
/color=1
/locus_tag="D90_02957"
/note="hypothetical protein"
/product="hypothetical protein"
/protein_id="gnl|SC|D90_02957"
/codon_start=1
CDS complement(33970..41772)
/color=8
/locus_tag="D90_02958"
/note="N-6 DNA methylase, protein motif:Cdd:COG4646protein motif:Pfam:PF00176.17"
/product="N-6 DNA methylase [methyltransferase]"
/"ID=D90_02958
/protein_id="gnl|SC|D90_02958"""
/codon_start=1
CDS complement(41788..42471)
/ID=D90_02959
/color=1
/locus_tag="D90_02959"

/note="hypothetical protein"
/product="hypothetical protein"
/protein_id="gnl|SC|D90_02959"
/codon_start=1
CDS complement(42477..43034)
/ID=D90_02960
/color=1
/locus_tag="D90_02960"
/note="hypothetical protein"
/product="hypothetical protein"
/protein_id="gnl|SC|D90_02960"
/codon_start=1
CDS complement(43012..43245)
/ID=D90_02961
/color=9
/locus_tag="D90_02961"
/note="Tyrosine recombinase"
/product="Tyrosine recombinase [recombinase]"
/protein_id="gnl|SC|D90_02961"
/codon_start=1
CDS complement(43259..43852)
/ID=D90_02962
/color=1
/locus_tag="D90_02962"
/note="hypothetical protein"
/product="hypothetical protein"
/protein_id="gnl|SC|D90_02962"
/codon_start=1
CDS complement(44215..44304)
/ID=D90_02963
/color=1
/locus_tag="D90_02963"
/note="hypothetical protein"
/product="hypothetical protein"
/protein_id="gnl|SC|D90_02963"
/codon_start=1
CDS complement(44304..48554)
/color=8
/locus_tag="D90_02964"
/note="Cys-Gln thioester bond-forming surface protein,
protein motif:Cdd:COG4932protein motif:Pfam:PF05738.7"
/product="Cys-Gln thioester bond-forming surface protein"
/"ID=D90_02964
/protein_id="gnl|SC|D90_02964""""

/codon_start=1
CDS complement(48625..49329)
/ID=D90_02965
/color=8
/locus_tag="D90_02965"
/note="Prokaryotic membrane lipoprotein lipid attachment site profile"
/product="Prokaryotic membrane lipoprotein lipid attachment site profile"
/protein_id="gnl|SC|D90_02965"
/codon_start=1
CDS complement(49561..50505)
/color=10
/locus_tag="D90_02966"
/note="ParB/RepB/Spo0J family partition proteinParB-like nuclease domain, similar to AA sequence:UniProtKB:P0CAV8protein motif:CLUSTERS:PRK13832protein motif:TIGRFAMs:TIGR00180protein motif:Pfam:PF02195.12"
/product="ParB/RepB/Spo0J family partition proteinParB-like nuclease domain [partitioning system]"
/"ID=D90_02966
/protein_id="gnl|SC|D90_02966"""
/gene="parB"
/codon_start=1
CDS complement(50462..51286)
/color=10
/locus_tag="D90_02967"
/note="CobQ/CobB/MinD/ParA nucleotide binding domain plasmid partitioning protein ParA, similar to AA sequence:UniProtKB:P37522protein motif:CLUSTERS:PRK13869protein motif:Cdd:COG2894protein motif:TIGRFAMs:TIGR03453protein motif:Pfam:PF01656.17"
/product="CobQ/CobB/MinD/ParA nucleotide binding domain plasmid partitioning protein ParA [partitioning system]"
/"ID=D90_02967
/protein_id="gnl|SC|D90_02967"""
/gene="parA"
/codon_start=1
CDS 51893..52417
/color=8
/locus_tag="D90_02968"
/note="Putative NADPH-dependent FMN-containing oxidoreductase ywqN Flavodoxin family protein, similar to

AA sequence:UniProtKB:P96726protein
motif:Cdd:COG0431protein motif:Pfam:PF03358.9"
/product="Putative NAD%28P%29H-dependent FMN-containing
oxidoreductase ywqN Flavodoxin family protein [electron
transfer]"
/"ID=D90_02968
/protein_id="gnl|SC|D90_02968""""
/gene="ywqN"
/codon_start=1
CDS 53433..56477
/color=8
/locus_tag="D90_02969"
/note="Sortase B-anchored collagen-binding adhesin LPXTG
cell wall anchor domain-containing protein, protein
motif:Cdd:COG4932protein motif:Pfam:PF05738.7"
/product="Sortase B-anchored collagen-binding adhesin
LPXTG cell wall anchor domain-containing protein
[virulence]"
/"ID=D90_02969
/protein_id="gnl|SC|D90_02969""""
/codon_start=1
CDS 56493..56762
/ID=D90_02970
/color=1
/locus_tag="D90_02970"
/note="hypothetical protein"
/product="hypothetical protein"
/protein_id="gnl|SC|D90_02970"
/codon_start=1
CDS 56710..57036
/color=1
/locus_tag="D90_02971"
/note="YdcP family protein DUF961 domain-containing
protein, protein motif:Pfam:PF06125.5"
/product="YdcP family protein DUF961 domain-containing
protein"
/"ID=D90_02971
/protein_id="gnl|SC|D90_02971""""
/codon_start=1
CDS 57052..57435
/color=1
/locus_tag="D90_02972"
/note="YdcP family protein DUF961 domain-containing
protein, protein motif:Pfam:PF06125.5"

/product="YdcP family protein DUF961 domain-containing
protein"
/"ID=D90_02972
/protein_id="gnl|SC|D90_02972"""
/codon_start=1
CDS 57603..58010
/color=7
/locus_tag="D90_02973"
/note="Helix-turn-helix transcriptional regulator ImmR,
similar to AA sequence:UniProtKB:P96631protein
motif:CLUSTERS:PRK09706protein motif:Cdd:COG1426protein
motif:TIGRFAMs:TIGR03070protein motif:Pfam:PF12844.1"
/product="Helix-turn-helix transcriptional regulator ImmR
[transcriptional regulator]"
/"ID=D90_02973
/protein_id="gnl|SC|D90_02973"""
/gene="immR"
/codon_start=1
CDS 58026..58352
/ID=D90_02974
/color=1
/locus_tag="D90_02974"
/note="hypothetical protein"
/product="hypothetical protein"
/protein_id="gnl|SC|D90_02974"
/codon_start=1
CDS 58453..59847
/color=4
/locus_tag="D90_02975"
/note="DNA translocase FtsKDNA segregation ATPase
FtsK/SpoIIIE and related proteins, similar to AA
sequence:UniProtKB:P64165protein motif:Cdd:COG1674protein
motif:TIGRFAMs:TIGR03924protein motif:Pfam:PF01580.12"
/product="DNA translocase FtsKDNA segregation ATPase
FtsK/SpoIIIE and related proteins [ATPase coupling
protein]"
/"ID=D90_02975
/protein_id="gnl|SC|D90_02975"""
/gene="ftsK"
/codon_start=1
CDS 60031..61227
/color=4
/locus_tag="D90_02976"
/note="XRE family transcriptional regulator Replication

initiation factor domain, protein
motif:TIGRFAMs:TIGR03070protein motif:Pfam:PF02486.13"
/product="XRE family transcriptional regulator Replication
initiation factor domain [relaxase]"
/"ID=D90_02976
/protein_id="gnl|SC|D90_02976"""
/codon_start=1
CDS 61240..61374
/color=1
/locus_tag="D90_02977"
/note="DUF3789 domain-containing protein Putative
conjugative transposon protein Tn916-like CTn1-Orf25,
protein motif:Pfam:PF12664.1"
/product="DUF3789 domain-containing protein Putative
conjugative transposon protein Tn916-like CTn1-Orf25"
/"ID=D90_02977
/protein_id="gnl|SC|D90_02977"""
/codon_start=1
CDS 61375..61596
/ID=D90_02978
/color=4
/locus_tag="D90_02978"
/note="Conjugation related protein"
/product="Conjugation related protein"
/protein_id="gnl|SC|D90_02978"
/codon_start=1
CDS 61800..62576
/color=7
/locus_tag="D90_02979"
/note="Helix-turn-helix transcriptional regulator immR,
similar to AA sequence:UniProtKB:P96631protein
motif:CLUSTERS:PRK09706protein
motif:TIGRFAMs:TIGR03070protein motif:Pfam:PF12844.1"
/product="Helix-turn-helix transcriptional regulator immR
[transcriptional regulator]"
/"ID=D90_02979
/protein_id="gnl|SC|D90_02979"""
/gene="immR"
/codon_start=1
CDS 62876..63379
/color=4
/locus_tag="D90_02980"
/note="Antirestriction protein ArdA, protein
motif:Pfam:PF07275.5"

/product="Antirestriction protein ArdA [anti-resitriction modification]"
/"ID=D90_02980
/gene="ardA"""
/protein_id="gnl|SC|D90_02980"
/codon_start=1
CDS 63397..63900
/color=4
/locus_tag="D90_02981"
/note="Antirestriction protein ArdA , protein motif:Cdd:COG4734protein motif:Pfam:PF07275.5"
/product="Antirestriction protein ArdA [anti-resitriction modification]"
/"ID=D90_02981
/gene="ardA"""
/protein_id="gnl|SC|D90_02981"
/codon_start=1
CDS 64018..64416
/color=4
/locus_tag="D90_02982"
/note="TcpE family, protein motif:Pfam:PF12648.1"
/product="TcpE family [mating pair formation]"
/"ID=D90_02982
/gene="tcpE"""
/protein_id="gnl|SC|D90_02982"
/codon_start=1
CDS 64394..66844
/color=4
/locus_tag="D90_02983"
/note="Type IV secretion system protein TraC VirB4 components, protein motif:Cdd:COG3451protein motif:TIGRFAMs:TIGR02746protein motif:Pfam:PF12846.1"
/product="Type IV secretion system protein TraC VirB4 components [ATPase mating pair formation]"
/"ID=D90_02983
/gene="traC"""
/protein_id="gnl|SC|D90_02983"
/codon_start=1
CDS 66844..69039
/color=4
/locus_tag="D90_02984"
/note="Conjugative transposon protein Tn916 orf15-like MFS transporter YtxH domain-containing protein, protein motif:Cdd:COG0349"

/product="Conjugative transposon protein Tn916 orf15-like MFS transporter YtxH domain-containing protein"
/"ID=D90_02984
/protein_id="gnl|SC|D90_02984"""
/codon_start=1
CDS 69036..70043
/color=4
/locus_tag="D90_02985"
/note="Endopepidase NLPC/P60, similar to AA sequence:UniProtKB:P21171protein motif:CLUSTERS:PRK13914protein motif:Cdd:COG3584protein motif:Pfam:PF00877.13"
/product="Endopepidase NLPC/P60 [pepidase]"
/"ID=D90_02985
/protein_id="gnl|SC|D90_02985"""
/codon_start=1
CDS 70059..70970
/color=4
/locus_tag="D90_02986"
/note="Conjugative transposon protein TcpC, protein motif:Pfam:PF12642.1"
/product="Conjugative transposon protein TcpC [mating pair formation]"
/"ID=D90_02986
/gene="tcpC"""
/protein_id="gnl|SC|D90_02986"
/codon_start=1
CDS 71091..71468
/color=7
/locus_tag="D90_02987"
/note="GntR family transcriptional regulator Helix-turn-helix domain-containing protein, similar to AA sequence:UniProtKB:O34817protein motif:CLUSTERS:PRK09990protein motif:Cdd:COG1802protein motif:TIGRFAMs:TIGR02325protein motif:Pfam:PF00392.15"
/product="GntR family transcriptional regulator Helix-turn-helix domain-containing protein [transcriptional regulator]"
/"ID=D90_02987
/protein_id="gnl|SC|D90_02987"""
/codon_start=1
CDS 71469..72332
/color=8
/locus_tag="D90_02988"

/note="ABC transporter ATP-binding protein, similar to AA sequence:UniProtKB:P0A9U1protein motif:CLUSTERS:PRK13536 protein motif:Cdd:COG4555 protein motif:TIGRFAMs:TIGR01188protein motif:Pfam:PF00005.21"
/product="ABC transporter ATP-binding protein [transporter]"
/"ID=D90_02988
/protein_id="gnl|SC|D90_02988"""
/codon_start=1
CDS 72329..72973
/ID=D90_02989
/color=8
/locus_tag="D90_02989"
/note="ABC-2 transporter permease"
/product="ABC-2 transporter permease [transporter]"
/protein_id="gnl|SC|D90_02989"
/codon_start=1
CDS 72986..73177
/ID=D90_02990
/color=1
/locus_tag="D90_02990"
/note="DNA-binding protein"
/product="DNA-binding protein"
/protein_id="gnl|SC|D90_02990"
/codon_start=1
CDS 73382..73570
/ID=D90_02991
/color=4
/locus_tag="D90_02991"
/note="Cystein-rich KTR conjugal transfer protein"
/product="Cystein-rich KTR conjugal transfer protein"
/protein_id="gnl|SC|D90_02991"
/codon_start=1
CDS complement(73678..74037)
/color=7
/locus_tag="D90_02992"
/note="helix-turn-helix transcriptional regulator, protein motif:TIGRFAMs:TIGR00270protein motif:Pfam:PF12844.1"
/product="helix-turn-helix transcriptional regulator [transcriptional regulator]"
/"ID=D90_02992
/protein_id="gnl|SC|D90_02992"""
/codon_start=1
CDS 74209..74418

/color=7
/locus_tag="D90_02993"
/note="Peptide-binding protein Omega Transcriptional Repressor, protein motif:Pfam:PF07764.5"
/product="Peptide-binding protein Omega Transcriptional Repressor [transcriptional regulator]"
/"ID=D90_02993
/protein_id="gnl|SC|D90_02993"""
/codon_start=1
CDS 74435..74707
/color=8
/locus_tag="D90_02994"
/note="Plasmid epsilon-zeta postsegregational killing system antitoxin, similar to AA sequence:UniProtKB:Q57231protein motif:Pfam:PF08998.5"
/product="Plasmid epsilon-zeta postsegregational killing system antitoxin [toxin-antitoxin]"
/"ID=D90_02994
/protein_id="gnl|SC|D90_02994"""
/codon_start=1
CDS 74919..75338
/color=8
/locus_tag="D90_02995"
/note="Plasmid epsilon-zeta postsegregational killing system Zeta toxin AAA family ATPase, protein motif:Pfam:PF06414.6"
/product="Plasmid epsilon-zeta postsegregational killing system Zeta toxin AAA family ATPase [toxin-antitoxin]"
/"ID=D90_02995
/protein_id="gnl|SC|D90_02995"""
/codon_start=1
CDS 75385..76122
/color=2
/locus_tag="D90_02996"
/eC_number=2.1.1.184
/note="23S rRNA adenine(2058)-N-methyltransferase ErmB, similar to AA sequence:UniProtKB:P21236protein motif:CLUSTERS:PRK14896protein motif:Cdd:COG0030protein motif:TIGRFAMs:TIGR00755 protein motif:Pfam:PF00398.14"
/product="23S rRNA %28adenine%282058%29-N%286%29%29-methyltransferase Erm%28B%29 [antibiotic resistance]"
/"ID=D90_02996
/protein_id="gnl|SC|D90_02996"""

/gene="erm"
/codon_start=1
CDS 76127..76258
/ID=D90_02997
/color=1
/locus_tag="D90_02997"
/note="hypothetical protein"
/product="hypothetical protein"
/protein_id="gnl|SC|D90_02997"
/codon_start=1
CDS 76655..77533
/color=10
/locus_tag="D90_02998"
/note="ParA family protein CobQ/CobB/MinD/ParA nucleotide binding domain, similar to AA sequence:UniProtKB:P37522protein motif:Cdd:COG4963protein motif:TIGRFAMs:TIGR03453protein motif:Pfam:PF01656.17"
/product="ParA family protein CobQ/CobB/MinD/ParA nucleotide binding domain [partitioning system]"
/"ID=D90_02998
/protein_id="gnl|SC|D90_02998""""
/gene="parA"
/codon_start=1
CDS 77981..78397
/color=7
/locus_tag="D90_02999"
/note="RNA polymerase sigma factor Sigma-70 region 4, protein motif:Cdd:COG2739protein motif:TIGRFAMs:TIGR02937protein motif:Pfam:PF08281.6"
/product="RNA polymerase sigma factor Sigma-70 region 4 [transcriptional regulator]"
/"ID=D90_02999
/protein_id="gnl|SC|D90_02999""""
/codon_start=1
CDS 78404..78652
/color=7
/locus_tag="D90_03000"
/note="helix-turn-helix domain-containing protein, protein motif:Pfam:PF12645.1"
/product="helix-turn-helix domain-containing protein [transcriptional regulator]"
/"ID=D90_03000
/protein_id="gnl|SC|D90_03000""""
/codon_start=1

CDS 79047..79304
/color=9
/locus_tag="D90_03001"
/note="DNA binding domain excisionase family, protein motif:TIGRFAMs:TIGR01764protein motif:Pfam:PF12728.1"
/product="DNA binding domain excisionase family [excisionase]"
/"ID=D90_03001
/protein_id="gnl|SC|D90_03001"""
/codon_start=1
CDS 79323..80768
/color=9
/locus_tag="D90_03002"
/note="Tyrosine recombinase, protein motif:CLUSTERS:PRK00236protein motif:Cdd:COG4973protein motif:TIGRFAMs:TIGR02224protein motif:Pfam:PF00589.16"
/product="Tyrosine recombinase [recombinase]"
/"ID=D90_03002
/protein_id="gnl|SC|D90_03002"""
/codon_start=1
BASE COUNT 22329 a 17547 c 17143 g 23971 t
ORIGIN
1 tccggccttc cccttaggag aagcccccga ggggcatttt tgactcccta caaatccccg
61 aaaatccttg attttactgg cttttctggg atttcactgt tagtaaattc tcgtccaaaa
121 cctgctaatt tttcgtggtt ttttgccttc cttttagcag ggtattagca aatgcaaaga
181 aatatgggaa acatcattga gttaatctgt agcccataaa tcacaggaaa cgcatttgaa
241 gtttacgcat tctcttatgc aaatcaataa tctgtgttca aatattataa tggtttttct
301 tcttgatttc aagatttttc aggaattaca aaaggcaacc ctgacggatt gccttttttg
361 tatgactaaa ttgttttcgc taaagccttt aataatgctg ctgtttccgg atttgataac
421 agttttgtga gcaaggcggc gttttcatct gtactgttct cagcaggctt ttctttttca
481 acttcgctct ctttttgaac ctctgccatt gggtcaagga gttcaaccga agtttcaaac
541 ttgggcattg gagcagtttt accttcttcc gcatttttga gacctttagc gttgtagaac
601 tgttcctcaa acttctgtgc gttatatcgc ctgtcctcgt cgattatatg gctgtaaaca
661 tcggcaacca tatccattcg ggcgtgtcct gagtcgcctt gaaccgattt catatcgccg
721 ccgttccatt tcagcttgta ggttatactt gcgtgacgga agctgtgaaa taccacatcg
781 ggtaagtcgt tgtcctgtat cagctttttc aaagcacggt tgattacctg accttccatc
841 ggtctgcctg acgaatggca gaataccaag tcataatcca aatactcatc accgaaaagc
901 tctttcattt cgtcaatctg cttttttctc tctaacagca tttgagctac ggttgacggc
961 aagaagattt ttctcacact tgttttagtt tttggcgttt tcaaaacaag ggaggtattt
1021 gtatttgaca gagttctcgg aaagacacgg ataatgtcct tattatcaag cacctccatt
1081 acatctctgt ttactcgctg aagctcttta tctacaaaaa tggaagcgtt gttttctttc
1141 aggctttcct ccgaaatatc aatgcaatcc caagtaagac cgagcatttc acccatacgg
1201 agtgaacagg caaaggataa attgatggct aatgcgagaa tatcatcatc gcacacttca
1261 agagcgtgca taagagtttc tgctgtccat atttctcgtt tctgatgttc ctctttcggg

1321 agcgttgcat tgagaaccgg gtttcttgtc atcaactccc attttaccgc ctgattaaag
1381 gcactcctta aaaacttatg gatttctctt acggtatgta cggtcagata ctcattttta
1441 ggctttttat tctgtaccac tttggtctta actttcagaa ggctttgata gaatttatcc
1501 ataagtctcg gagtcagttc atcaagtttc acatcgccga taagcggaat gatgtaatta
1561 tacataagcc ctttctttga acggtaggtt gacattgccc agttatttac tccgtacacg
1621 gagcagtatt cttcaagcaa atccgcaact gtcgtagcat taggaatgat gaatgtaccg
1681 ctttgctgtt caaactcaat ttgagttttt ctctttttag cgtctgcgtt tgtatcaaag
1741 gtttcccatt tttgatgttg gtttccgtca tcgtctttgt aggtataaac gacggaatat
1801 cgattctttc tctttacaat agaagccata gtcgttcctc cttattcatt atccagccat
1861 ttatcaaagc tggtctttat tatgcggtat ccggtttcga gcattactgc ctgaaaacac
1921 ccctgtttga tgagcttata tacggtctgt cgggtaatac ccaagattga ctgaacttcc
1981 tctacggaat agctcttttt tgtatatgta ccgccgttta attcacgaac tctatcctca
2041 atcgacatat ccctctacct cctcgatttc cttaacttta gtgtatttac tttgcgaagc
2101 gtaccactta tcaaagcttt ctctttttat tcggcgttta ccgtcaaccc atataaactc
2161 aacaatgcca tcgttcataa ggtcatttgc ggtgcttctc ggaatgccga gtacctttgc
2221 aacagtaaga ggagagagcg ttttgccgta tttcttaccg ggtctttcgc cgtttacttt
2281 ctcataatgg aattgtcctg cataccaatc ttcaaagcta tccacatcaa ctctcatttt
2341 gcccagtacc aaataggtct taaatcggca ctgattgatg agtcggtatg tagccgtctt
2401 tccaagaccg agtattctca ttacgtcggg tacagacatt gtctttttgt cctgatatgt
2461 aagaactgcc ttgacgattt tcttattttc ttcaacgctt ttctttgaag ctctgtattc
2521 gttgccgact cggtatgttt tgaataagcc gcaattcatc agtttcagtg cttcttcttc
2581 ggaaatggaa aacaaccgag atatttcctt gacagaatag gcttgccact gtgcttcctc
2641 ctgaagcttt tcatcgctga acaattctgc caagcgttca ttttcctctc tgagattaaa
2701 ctgctccact cggtcattga acatattaac aaatgccatc ttcagaacct cctttctcgc
2761 cattgagcca attttcaaag ctctttttag atattcgata ttgacctccg acacgaacgc
2821 tatggaattt cttgctttta agcagctcat agacggtagt tctgctcact ttcaagatgt
2881 gctgtatatc atctaccgtg tatgctcgta tatcaggttc agtctgcctg aatgtattag
2941 gaagtcgagt atcgctcttt tcaattttct ttcgcattac cgtttacccc cgatttctta
3001 tttgccactc tgactaccat atagccaagt gacttggcta tatggtactg aaaatcggag
3061 tttttgttaa ttatgcgatt ggtttaatcg cataattgac aaccggagct ttcgctgtcc
3121 atattatcaa gccatcggtc aaagctcttc ttggaaattc tgtaatgtcc gccgaccttt
3181 acaaaatgga aaacgcctga gttgacgaga gcataagcag tgttctttcc aatgctgaga
3241 atatccatta tttcttctac cgtataaact cttttctcaa agccggaact atcatcaaca
3301 gcgttgggta gtttgttcat tgcttcaatc ttatcttcaa acattgtcaa cacctccctt
3361 agcgtgatga tggttctttt tgaaaatcat atagtcatag ccgtatccta cctgacattt
3421 atcgcacatc tctttatctg cctgaaaggg gtcaagtctt tttacaatgt tttcagggat
3481 agactgaaac tcgcttctgc aacgagggca aagacttagt acaatgcctt tacgctctgt
3541 cttgggttta acgggaatac cgactgcaaa tttcaatcct cgcttgattt cgctgatttt
3601 atctgcgtcg gtaattcttc cgacaaaatt atcaagctct gtggctttat ccacggtgcg
3661 tagctgctca agcattacca tcgacggctc ttttaagcca caatccgtgt ttagctcaat
3721 atgagtattc attgccgttt tctttttaac agccgttata gcggctacaa cgaccgtagg
3781 actgctttca tttaagcgat ttgtctgaac cacaagaacc ggtctcattc ccgattgcac
3841 actgccgttt gtaaccccca agtcgcagta ataaatatcg ccccatctga cattcttaaa
3901 ttctttttct ttcatatgta aatcctcgct ttcgattatt ctgaggtaat tcatatccgg

3961 attagaatta accccttcac ttgtttgaaa tgggagagcg tttttggggg gtgttttaga
4021 aaaatttcct caactttttt attgcactgt ttaccgattc ccttacagaa gtaaaatcga
4081 ctccttcttc atctgcgatt ttttcaagtg ttttgttgta gaagaaatac ttaaaaactc
4141 tgcgtttctg tgtttcgctt aagtaacaca aaacctcccg aagtcgctta atattgtctt
4201 tcagcattat ttcttcgatt gggtctgtat cgactgccaa ttccggatga tgcacttcaa
4261 attctccgtc attgatgtca aatgcgtgtc cgaaaagaag ccttctcatt ctctctcttg
4321 cttcgtagtt ctgatactcc gttatcactt cgccctgagc catagaaagc agaatgaacg
4381 gagtgtagcg acggataaca tcagggtatt taacccacaa ttcttcttcc gccagttctg
4441 ttacaatcgc ccatttctcc gtgccggtgt agccgtgata ttcataccgc aggttaatga
4501 gcttgcagtc gttcgcaaat aataattcct gttgttctaa tgtaagtttt gtcattgtcg
4561 taatctcctt aaatttgaat ttttgaattt ggttgaaatc aaaaattcgg agattacgga
4621 tacttggcga tatacgccag ccacttatat gtcacggtct tgctcctttc gggagcgaca
4681 aaaaagccgg acaccaagaa cattaggata tttctatccc ttaattctca gcgtccggca
4741 atttgatgac tcaagagctt tcgctcaagg actcgtggct cggtacatct gtccgatatt
4801 taattgttcg ctctgttact acttcttgtt aatctgtact tgactgttcc cttgcgtagg
4861 gataggtcaa cctctactac tctgtggcag ttcggacatt tcagttcaat cacgcctgtt
4921 gtcattgtta ccttgtcaaa aatacgccag ccacagtttt tgcaccttac gattatcttt
4981 tgttccaacg atttcatagc tcgctcacct ccggataaac cgtaaccaaa tcaaaaggat
5041 tatcaagata ctcctttttc aaaagtccga gctgtttcat tcgtatagca agtgcttttt
5101 tggaacaccc taaaaaatcc gcaagtgcat caaatctctt atatacgctc ggatagtata
5161 ttttgttcag acactcaatc ttttctccaa gactgaacag atacattccc tgctttataa
5221 ggttttcggg aagaaggata gcagctccaa gcgtatttgc ctgccattcc tcccaatcgg
5281 atataggctt atttctctcg gaatttgcct tgtaataatg tactccggca gatttctgtg
5341 tcacaccgta atcattcggg aacaacatct taaatatctg atggctgccc tcgtgcatta
5401 aggtgaaatt ctttctgcct ttcagcttgc tgtcaaagtt caagtccttt tccacgagaa
5461 cggtttttcc atccaaaaag aaaaaggctt cgttatcatc atcttcaaat acctgaacac
5521 ccatttccgt aaatgaggtc attccgagaa tactaccgtc ataagatagg tgctggtatt
5581 cgacattcaa tccgagaacc ttctccaaga gaagctccgg gtcaattctg tatatttgtg
5641 tgttacggac atcgggaagt gccatatatg cctgaacata cttgtcggca atcgcttcga
5701 tgtcaaatct cgataaatgt ttcaagttac aaaacctcct accattagga ttttgcttct
5761 acaaaccact ttccgttttc ttcgtcgaac agaaatgtgg gcttgccgca aatctgtacc
5821 gtgtagcgga taccggttcc tccgactttg gtggaagccg cacggcattt ctgaattaca
5881 cggtctatct catatttttc atcattctca aacttgatga agttcggacg agccgttcca
5941 tcaggtctat gtgttacatt tacttcaaca taaacttttc ttctttcgga ttccattttt
6001 attcctcgta atttctcaaa gtcaaaacta atcttcccaa gtgcttgtat ccgtcgatgt
6061 tcttatccga caccctgtac ttgggtcggt catctccggc agtattgatg taacaggaaa
6121 gccttccttt cttaaagcgt tttttccggt caaagaaaag aaacattccc tcatatatac
6181 cgtagtcggt aaaatggttt ccgccgtccg cacgaaagat aacaatatct tcgctgcacc
6241 cctgcaaaaa gcagtcgggc atcgtaacat aagccgtaac ctctttttcg tcaatcacat
6301 ttagacctcc ttaaccaacc atacctgtcg gcatagtgat ttcggctttt tccggtggta
6361 gccgcaagtt ctgacacaga accccattcc ttatgctgtc ctttccgaaa cgccgtctga
6421 tttcttcaac acacttttcc attcgctcca atttgtcctg ctttgcggcg tccataaaca
6481 tatctatttg acggggcgtg tcttgtggaa taagatttat ggcttgtact gtcacagaac
6541 ggatagggtt attccaccca tatctttttt caaacaactg aaaagcggtc tttgcaataa

6601 tcatcggaga ctgtgtagga agtgcaatct tcgtctgcca ttgccttgta ttgagcgtgt
6661 tgtctctaat atgtattgcg acaccttctg cactcagtcc gtgaacacgg agcttgtgtc
6721 cgatgtcttg ggttaattcg agaaacacgg gccacacctg ctccggtttc tctaaatctg
6781 ctactgttgt tataccgtgt cctacacttt ttataggaga aacgaagtct ttcttggcaa
6841 caagggaaag gtcgttgccg ttggcatagt tccataaaac tactccgttc tttcctaatc
6901 ttctgcgtaa gaactcaggg tcgttattcg ctaaatcccc gatggtacga atacaatagc
6961 tgtcaagcac tcgctgggtt gcccgtccta caccgagcag gtcagcggta ggaagtcccc
7021 aaatcttttc tttgaatgta tcctttggta taacggttac tgcgtccggt ttcttcatat
7081 ccgagccgag cttcgcaaat atcttattaa aggaaacacc cacagaaatc gtcaagccga
7141 gttcaaactt catcgtttca cgaatttcat ttgccacctt ttccggcgaa ccgaaaaggc
7201 tttctgtccc gctaatatcc agccaacact cgtccatacc atacggttca acttggtcgg
7261 tatatcgctc ataaacgctt ctcgcaagct ttgagtattt gatatactcc tcataatggg
7321 gcggcactac aaccaagtcc ttgcacttct gtttagcctg ccatactgcg tccccggttt
7381 tcacatcaaa agcctttgct ttgtagttct ttgcaagcac gataccgtgt cgttcctcca
7441 cagagccgca gacagcaatc ggatatttct tcaatgcggg gtcgagcata cattcgacgc
7501 tggcatagaa gttgttcata tcacaatgaa gtatgcttcg taacattttt tgaacccccc
7561 ttgacatttt gggaagttta gtgtaaacta tgagtagttc aacttcctgt ttcttattat
7621 acgcagttta agtgtacttg tcaagagtaa atgagaagtt aaactgtaaa tattctttta
7681 aggagtgaaa tcaaagatga cattcggtga aaaagtcaaa gcagagagga ctaaactcgg
7741 tcttaatcag gatgaattgg ctgaaaaaat aggcgtaacc cgacgtgtaa tttgttctta
7801 tgaaaatgat aaatcccgtc cgagaggaac ggaaagatac aagaagctgg cagaagcatt
7861 aaatgtaaat gtgaattatc tgctgtctga ggatgacgct ttcatcgccg atgtagagga
7921 taaatacgga cgcagagggg caagacaggc tcaggagttg cttgcagaag ttaccggtct
7981 gtttgccggc ggcgaaatgg ctgacgaaga tatgcgtgaa atggttgacg ctattcagga
8041 agcatatctt attgcaaaga agaacaataa aaaatacacc ccgaagaaat accgcaaaga
8101 cgagtaatcc tcacagtgaa ctaagtccaa tatatgggac acacaatagt ttataattta
8161 ttatagggta aatatccgat atactataat atgggaggtg agccggattg gggtattaca
8221 ctacggctga aattgtgaaa atcgtcaaca agctcattga ccgttgcggt actcgtgacc
8281 cgtacaaggt cgcaaaagag cttggtatca atattatcta ccgagatttt gaaaagcagc
8341 gtggagcata caaggtcatt ctgaagaacc gctttgtttt ccttaaaaac ggtatgcacc
8401 cggtcgtgga acagatagtg ctatggcacg aaatcggaca cgatgttctc cacaggcaag
8461 aagcggttgc tgtcggcggg ttcaaagagt tcaacatctt tgatatgaga gagaaccgta
8521 tggagtatga agcaaatatc ttcgcttctc aggcttcgct cccggacgac accattttgg
8581 aatacattga aaacggctat gacattcagc agattgcacg ggcaatgtgt tccgacataa
8641 acctaattgc tctgaaagta gatacgctga ttgcacaagg ctatcaactt cgcaaacagg
8701 aacaccaaaa cgattttctt aaatacagta aaaaaatata agaccgtcaa gtctcgaatg
8761 agatctgacg gtctaatttt tatcagtttt tattatatcc gaattgtctt gtgcaaaggg
8821 tatctttcca ccatttttca cgactcagta tgtaattatc atccattcgt gcgttatagt
8881 tttctaatac tgaatactga aaatttgctt ttatataatc aaaccctttg gtgtcaacaa
8941 tatgtttaag ttcaacattc ccaccgtgtc cattgtctat atagttagtc cacctttgaa
9001 gcaacattcc atactgagca gttgccgagc caacatacat ttttccgttg ctggtatcag
9061 ttataagata aactgctttt tgatttctca aagcatcaag ccaaccgctg cgtttattcc
9121 gaataatcgt ttccagttgt gaaaaactca atcttatgtt ctcataaccc ggaaattcgt
9181 caccatcata agcagttgaa agaatttcta ttacttccag ttcctccatc atagactcat

9241 aggttcttcc cataccacgc ttcgtattgt gatacttcaa tactaatctg ccgtaatact
9301 tacgatattc ttcaatttcc tcagcttcaa agccgacatc atcaacgaca tcaatctctt
9361 ttgtaatcct cttaatcgtt gttagcaacc atttatcatc atagaggtaa agcagacaaa
9421 ttgcaatctg tcccgtatgg aaatatcttc tttgattgtg ccacaaaaac cactctatgt
9481 taattttatc agggtctctt ttgtattcct ctaaagggtc tgtatttcca ttgaaaacat
9541 tcaacttgag tctggtttta ccaatatcac aatccggtat cttaagtaag tcgtttatca
9601 gaattctatc cattacatat ctccttattc gtgcaggaaa accttaatat aaatcttgac
9661 ccacgagcct ttcttttcct ttgaggtaat cttaccgaaa aggagcttcc cggcgtccat
9721 caggcgagca aatattacat tatcctgctt cggcacataa ccgattttta cgccgtcaac
9781 agttttaata acaatcgctt gcttatcata agtctgtgcg gaaaccctgc cgcttcatgg
9841 cttcggattt catcttgtaa gcaaacgccc gctccgatgg caggatattc tcacgctgca
9901 aattgctgtc tacaagggtg atgatggctc ggtcacggtc taagggcagg acaaacgcag
9961 gcacggtatt tatccctgca agttcagaag cacggacacg ccgctgtcct gcaatcactt
10021 cataaccgtc cccgtcctct ttcgggcgtg tgattatcgg cgtgacaatg ccaaattcct
10081 tgatgctttc cgctaactct gacagtgttt catcttctgc cacatgaaac ggattgtcgg
10141 ggaacgggta caagtctttg gtctttaaca ccttaaaatc ctgtttcttc attcacatat
10201 ctacctttct ttcgctctgc ttctatgatt tttctgcaat tcttccaaca cttcgggcgg
10261 tatttttgac agcagccgcc ccatctgctc gttggttctc tgcaattcaa atatcttctg
10321 attcgctttc tgcaccttta gttcctgctc gtacttttca tcacgcatac gccccgcata
10381 gtctgattcc tgcccaattc tctctttcaa actgtcgata tacgcctgct gtttcccgat
10441 ttccttagag aatttctcca cgtctggcag ccatgcagag agtaaatcta acgctttatc
10501 ccttttcttc cctgcgttaa aggcgttgat gtcggatagg gcagacacga tttcttcata
10561 ctgtttatca agcctgccgc ctaatttata gagccatgtg gggacgtgtt tccgcttggt
10621 ttccattgag gactgccccc gttcaagctg attccaccgt gaggacatcc gctcatggta
10681 ggcggtctgc cactcggata atgatttctg gttgcctaag atagctttcg ctgacagctt
10741 attgtctggc gtaatcggca caaagcagag gtgcatatgg ggcgttctct cgtccatatg
10801 gacgacagcg gagaggatat tctgctttcc aacacgctcc gaaatgaagt caagagccgt
10861 ctggaaatac gctttttgtt cttcgggcgg taactggttc ataaattctg gtgaagctgt
10921 gatgagcgtt tccaccatca tcacgctgtc tttccttgtc ctgcaccccg cttcggctac
10981 catgcggtta atctctttct tgtaggtgta ctttggtggt gctatgagat ggtaattgtt
11041 tttagagcgt tccatatcta tatctgggtt gcttttgtag gcttctttct tccgctcgtt
11101 gtggcgttca caagccgcaa cgccgcccgc ttttcgtttc tggaaacgca ggattgcata
11161 aggcataggc ggattcctcc tttctttcgg cgggtgaccg tcctttgggg tgacagcggt
11221 gacggtggtg acagcagttt tgggataccc cctgccgact gccgcaactg tcaccctcac
11281 cccctgcatg acggtcatgt cgatgatgac ggtgtttttg ggataccccc ctcgcaagag
11341 ccgtcaccgt catgccgcca ttcttttatc cgaagccgta aggcgtagga tagcagggca
11401 cagccctgcc ttaagggagt ccagagggaa cgtctggcac acgactttgc agggcaaagt
11461 gtagtgtgtt acaccctgta aacgcagtcg gaaaaatcgg aatgattttt ctgaccgcag
11521 gggtggtttt acacgaccga aaagggcgag taaatctcac gcctgcattt tgcgtttacg
11581 gggtgttctc ccgtagggat gacagcggca gggtatccta aaatcactgt caccaccgtc
11641 actgctgtca cccgcagggc agagccgcca gctttacatg gctttaagcg tcaatgacag
11701 taacaggggg gtatcctaat atcgctgtca ccaccatcac cgctgtcacc cgccctcctt
11761 gcaagggcaa tccgcctgcc gtctttcctg cggctgtact ggtagcagat acggttctcg
11821 cttaaaaatg tggtgcggta ttcattcagc catttcgtaa tcaccgtggg tatggtttcc

11881 gtttccccca tagcggctaa cagttccgtt gccgtgccta tccattcttc cttatccctc
11941 ataaaatcca ccaaccgaaa aaggacatct ggtatcgttt ctttcgcaag ctgctcctgc
12001 gttttccgct ccacaagctc ccaacggcaa tcacggaaac gcagcgtgta ttcctgataa
12061 ggcgtgtccc tgcccgtcac atacagcttg gcggtgtcag acgcacgttt ctccttttcc
12121 agaacaaagg tagcgtccgc actccccgtt aatcctgtcg tcccagacac cttgttgaac
12181 acgtcgctgt cattctgctt tcggatgtgg tgtacgacaa tgaccgccag agagtgcctg
12241 tcggcaaagt ctttgatgag ggagatgtcc ccatagtcgc ttgcataggc attgtctttt
12301 gaagctgtac ggactttctg caaggtatca atgacaatga gcctgctgtc tgggtaatct
12361 ttcagataat cttcaagctg cacgataaga ccgtctgaca gcttgcagct tgccacggca
12421 aagtggagcc gcccgcttgc ttcgtccgtc aaacgaaata acctgtcctg tatgcggcag
12481 aacgtgtcct caaggcagag gtaaagcaca tcgccctcca ttgtcggcat atcccataaa
12541 gggattccct gcgacacgca taagcatagc ttcagcatga gccagctttt gcctatcttc
12601 tgtgagccgc agaacagcga taagcctgtc gggataaggc tgtccaccac aaaggatggt
12661 ttctcaagcg gttcataaag gagcgtttcg gcgttgactg tctgtaactt ctgcatggct
12721 ttcctccttt cgggcggtgt ctttgttttc gttgcacata ggcgttgacc tccttaaaaa
12781 tagatttact cccacggaaa aaagtgagag tatgtaatgc cgccaatccc acacaaataa
12841 aaaacagatt tctttttcgg gcggtgtcgg tcacggttgg agatatttct tcaatttccg
12901 cctttacttt ctcctttgca atcgcaacag atttctgtat tgccgaagcg gtgcaatgct
12961 ccatagcggc gatttggtaa aaattgaact catactcgta gtagagcagg aaacgccgcc
13021 tttgtatctc tggaaggctc ccaaccgcct tatagagcgt ttcattccgt tcttcctcaa
13081 ccatgcgttc atcaaggctc ttaggcacac gcaacgcccg tctgtaaagg gtttcgtccc
13141 atacctcgtt aaactccctg tgccgctcgt cccattagaa aagattcctg ttcctgcgct
13201 ccatctgccg aaactccata aagaactgtt ccgacacttc caactcgtgg gatttgccct
13261 gcccgtcctt aaagctgata aaataccttg tgccgctttc cgtggattcc tcccgaagcg
13321 tgtatgcctt aaccctgtat gccatcctgt tgtcctcctg caaaaaatga gtgaggggat
13381 ttccatccct cacccagtag cccgcaggat ggcaggctgt tttagaagct ataaaatttt
13441 ccgcagcttc ttccgcagat ggtctaacct cgcatagatg gcaccagtcg tcaaatgcac
13501 aagcggggca atctcctttg tggaataccc ctgcattttc agcaggacga ttttcaaggt
13561 acgcccgtcc accgtgacta atacttgata gagattttcg ctctcaatct cttccagtaa
13621 ctccgcaacc gtacccactt ccgcctgccg ctccctgcct gccatgtcct caagatattc
13681 cgcaacgtca ttcgtccatc ggtaaaaccg cctgttggaa ttgaagtctg ccctgtccgc
13741 catgcgtatc tgctcaatgg tcgcttcatc aacgccgcac tcacgcagca gcttttcctc
13801 cgcttctttc cagatacgcc atttcctgtc ctcccgtccg tggttgtatg ccattcttac
13861 ttcctccaat cagaattgat tgagagggca gagaaaagcc ccctcatctt cccaaaggaa
13921 aaaacaaggg ggctgaacgc cttaaatttt aattttctat tatctttctt agttttatta
13981 ggattctggc tttgcgtttg tttacaacgc tctgggttat gcccaacctc gccccgactt
14041 cacgctcgga aagtccgtca aaaaagattg cctgtatcag ctcctgttca ctatccgaca
14101 gcaaaggcag ggcggctttc agcctgtcca ccatgaccgc attgacaacg gtttctgcaa
14161 tgtccaccgc ttcatcagtg ataaagtcca gaggattccc ctcgctgtcc gtaaatccgt
14221 cgagagatag cagtctattc tttgtatcta atttttgcag ataacgccac cgttccttgt
14281 cacggtagaa gtctgtgtat tgctccctca cgacttcaag cagacagcct tgaatgggga
14341 taaacagctt gtccatatag gtctggtcgg attccctgca acggcagaac tccgtgtagg
14401 ataattccac atagccgcca ctttctctga tatatacctt tcttggtgca tatttcacca
14461 taaatacctc ccatctgaat ttttgaaatg taaaaaatcc agatggagag gcggagaacg

14521 acaccgcata tcagaaacag ccctacggca ctttccaaca aaaatcgaca aaagaaaaac
14581 cgcaaaggct ctgtgacctt tacggttata ggaaatagta attctattgt atggagattg
14641 tacttctact ttcaaggtga gaagtaccgt tgcactggca gaaatttttt taactggttt
14701 cctgccgttc tctgatatgc tatgtattga taaattttgc ttcgtatgag cagatagata
14761 actgcccgaa aaaaggacat aaaaaaatcc ctccaaattt aaaaacaaat tcagagggta
14821 atttagggta taacaaaata acccacccga atatcgtttt ttttatttgg tgagtttgtt
14881 ctttcaaata cgaaacaaaa agagccgatg attcgtgaat tgttccacaa tttcatcggc
14941 tctgcgtctt aagcgtctgg ctctttgatg acagttatct tcaagttgtc aactttaatc
15001 aatctttctt gtctgcattt tggacaataa aggggataat tctccaaaac agtgtccttc
15061 ctaattttat tacgggtttt gctcccacaa acaggacaca atatccattc gcatttcatc
15121 ataattagtc tctaatcctt tcaaatctca ttttatatga cttttgcaag ctgttaagct
15181 aacttgtgga acatatgccg aaccttatct atacggctat tcgggcggcg gggttggcaa
15241 ataaatttac caatagctgg ctggtatcct tttaactctg tcaagcagac tccctgccca
15301 tttgtgaaat aagttaaatc gttcctgtat tcttgaatac atctagcagg gatttctcct
15361 ttcagaatga cctcgtcatt ctttatctga gtacttacaa tatctgcaca ataccttgga
15421 gcatcatgat acgcccgtga gagatattcc tgcggtgcat aaatttcaaa gtggagatat
15481 ggctctaata gttctgtccc tgcttttttt aaagcctgct ccaatacgat aggggaaagc
15541 agccgaaagt ctgcgggggt acttacagga ctataataca atccatattc aaaacagatt
15601 ttacagtctg tcactttcca tccatacagc ccctgctcgc agccataaag aaccccctcc
15661 ataaccgcat tttggaacga ttgatttaaa tatccaagtg aaactctgct ttcatactgc
15721 actccgcttc caatagggag cggctctatg gacaacccga cagaagccca gaaaggattt
15781 ggcgggactt ctatgtggat ggtatattct gcttttctaa gcggtctttc catatatata
15841 acagtaggct cttttatttc tgcctccaca tgatattttt cctcaaggat ggcacaaatg
15901 acttccatct gcacattccc caaaaaagaa agtataatct catgcgttgt agtatccaca
15961 taatatttta aaagagggtc gccatctgaa atttctttaa gtgccccaag caatatttcc
16021 cgctgttcag atttctttac tgcaatcgtt gtttggagca tagggagagg attttcaata
16081 aattttctct gcggcaacag catttcgttc cccaaaatac tgtttagctg caaaacatca
16141 tttggtaaaa ttacaatatc accagagcag gctgtatcgg atgaatataa ttcaccgttt
16201 gtcggaacac acatctctgt gatttttatt ttctcttttt cagatatttt aataacatcc
16261 ctcaaatgca atgttccgct atatatacgc acataaacaa aacgccgcct tttctctgaa
16321 tattcaatct taaaaacctg cccgcatagt tcagattgac cttcaggcgt tgatgaataa
16381 aacttactgg caatcacttc tataagctgc cgaatcccca gattgttttt agcacttccg
16441 tgataaacgg gaaataacgt tccgttttgg aatctcctgt tttcttcctg ttccagttct
16501 gacattttaa acggtttccc tgacatatat ttctctaata gttcatcgtt tcccataatt
16561 accgcatccc actgttccat atcgtcattg tccgttacat ttatatgggg atgctgccca
16621 accttttgct tcactataat ttccgaagaa agctttgctt tcatttcttg atataccatt
16681 ggcaaatcaa tcccctcttg gtcaatttta ttgatgaaaa aaattgtcgg aatcttcatt
16741 gtctgtagtg catgaaacag tatacgggtc tgtgcctgta tgccatcctt tgcagaaact
16801 aataatactg ctccgtctaa tacggataaa gaacggtata cttccgccaa aaaatccata
16861 tggcctggcg tatctataat gttgactttt acatcctccc actgaaaaga tgtcactgct
16921 gtctggatag tgattcccct ttgacgctcc aaattcattg tatctgtcct tgttgtgcct
16981 ttatctacgc tccctggttc tgcaattgca ccactggtat acaataaact ctccgttaat
17041 gttgtctttc ctgcgtcaac gtgagccaga atgcctaagt taattatttt catgtgattt
17101 tcctcctatc aacacccaaa aaagggcata aaaataccca gtgataaata ctcctatcac

17161 tgggtaaata actccaatag ccccaaaaca cttatatgtt ttcgggcata taaaattaca
17221 tgataaaagt attcttaaac tgggtacaaa aaactaagcc ccatattaaa agtgaaacgg
17281 gactgctact ttttgttccc actatcaaat tgacagttta tttaagaata ccttgccgca
17341 tatttattaa ctccttgtat aatactgaat ctaattatac tccttaaccc tttatttgtc
17401 aagctgacaa actaaagcag aaaaagcggc aggatttccc cctgccacta atcatctgtt
17461 tatgcaaaaa taatttcctt ttccacaatc tccctcgcac aagcccttat gttattgagg
17521 catcctgtcc attctaaggc gttttctgcc tttagctgtt ccgttatgcc ctgtgcctgt
17581 ttcataccct ctatgagcct ttcaaagcgt tcctgtgcct gtctgttgat gtcggcaagg
17641 taggcgttta gcctgccgct tgtaagaaga ttggtgtatg taactttacg gtactgtttt
17701 agataatcta aatgccgttg cccccagatg cctattgcct gttcttcttc ggcgggtaca
17761 gttaagcacg gtatcaaata atccccttgc ctttcgtatt tgccgcccag ttcctcaaat
17821 aatgattttg ccattgtctg ttacctccac attctttttt attttgaatg tccgcaaaat
17881 ccgtcctaca tcaatcggat ttgatggcag attccaatgc acaattcttc ggcaataatt
17941 gtggaaatcc aatccttctt gtcctatgga agtagatgct aaaacaaatg gtctaaatgg
18001 agaattgaaa gaatttctaa cgctcttttt tctgtctgcg tccttttctt ttccatctcc
18061 tttagtaaag gcaactgcaa agtgagaacg aatagcagtt ggtttttcct ttgttccatt
18121 tgctcttgcc ttaaacgcat tgaatgtatc aatagtgtat attgtggttc tcacatccat
18181 agaacttgct attgtatagt gtaagttatc aaccaaatta ttgtctgcgt ctaaaccgtt
18241 tgtaattaaa tgagcatatt catcaaacat agcctgaatg tttccctgct tgcaataggt
18301 aagcagattt tcccaatgtg cgtcctcgct ttttttacca caagcaagct cgactgtggc
18361 tgtagactcg gctgtattca tacggtttat aaagattttt gcaatctgac ttggcaaata
18421 acttggaaaa gtagttcctc gtttgcagta agattgatag gttctgttta tagttattgc
18481 aggagaagct attgccatat ctgcaagaac atctaacaaa tcatcaggtt ttttcccaag
18541 cgtacaaatc tttccgtagt ttgtttccaa atacaactcc cgtaatgtct gaagatgggt
18601 taaaaatccc ttttgtcgct tggatttctc ttcttcttca tcaaacgaag caagagcttc
18661 accactattc agccaagttg taacatatcc cgcaccgtcc aataacagag gtgcaagata
18721 ataccatcgt ttatcctgcg ttcccgatgt cggagcagta tatttatcta atttctcagc
18781 aatcttgcca agacttatga gagagccaag aagaaaaaac tccaaataat caacttagga
18841 gtaaaagagg atggggaaag aaagagtgat attcagtaat cggtttgacc gaccgacgct
18901 tgaacaaatt gaagaagaaa tgaaaaaaga gaattacgat gataacgaaa acttgatgtc
18961 agcagaagaa tttcttgatt ttttgcagaa gaacagaagg actgtcccca acgaaaagcg
19021 tttagtcaat aaagaaaaat ttatatgggc agtccgtgag ttatcagaaa cctatgaaat
19081 tgacgcagat ttaagagaaa gcgatgacgg atataccgca tctctctata tgaactatgc
19141 ttcatataat ggatatataa agaaactgct gggacttatt tttatcctgt ccgacgactt
19201 ttctatgttc aaagcaaaag ataatataga cagcgatttg ttgatgtgtt ttacttatca
19261 cacgcacaat gtctatctga aagatagaga aataactgat tttgaataat gtacttgatg
19321 gctactcaaa cgagcagcca tctttttta atatgccgga ataccgtatc cgtagataac
19381 tgagctacca accgtgtatg ttttggttct gcaagtgtcg cctgagttgc cctcaacggt
19441 atagactgta ccgttctcgc atttctgaac gataccggtg tggtctgtca caccgtcgcc
19501 ctcccagtca aagaagatga ttgtaccggg cgaaggttca tacgaaccgt ctgcccattg
19561 tcctcggtct ctaaaccatt gaacgccgtt tacacagccg gcatacttcg gaataatacc
19621 ggcgtcgata tatccgcact cgttggcaca ccaagaaacg aagcaggcac accactctac
19681 acgggagtca aatccgtacc aagaccagta aggctgaccg cctacattac ctacctgaga
19741 cagagcaacc gttacgattt ggtcgtcgct gtatccgata ccgtaaagca cctgagacca

19801 tagttgattg ttttcgtcct ttagcagctc ggcaagatac tctttctgct cctgattaaa
19861 gccgtacata gccgccattt catcaacggt cttgtgcgat accgtgatat acaggaaagt
19921 ttctgttact gtggtttcgg tctgtacgat gttgccgtgt ccgtcgtcgc tttcttcgat
19981 ttcggtgtgg gttttcgatt ccgttctgga agaaatatcg ttcatctccc aaaaaatatc
20041 cgacagtagc tgtttcttgg tttcatcaac cgtagcaacc tccatcggat tatcggggtc
20101 ggtatttact ttaacggaat agaccgcaag aacttctttc cacacagctc ggcttccgct
20161 catttcaagc acatcatagc ttacggagtt cttttcctgc tccaatcggt catcatattc
20221 ttgattgatt tcctgaacga ctgtctgcat agacattccc gtgccggaat cctctcctga
20281 aaagaagata ccaaacacag aaccgacaat taatccgaca agacacagaa caataataac
20341 caccaccgca acccagccgc cggcagcaat ggcggcaaca agacctttta ccgccgcaat
20401 gattgctttt actgcggcaa cggtggcttt ggctgcggct ttaatcgtag cggcagttgc
20461 tttggctgtg gcttttgccg cctgtgccgc tttctgcgaa gccttaactg tcgcctgagc
20521 tgttttctgt gccgcttttg ccgcttgctg actggttttt atcgcagtct ttgcagtctg
20581 ctcagccgtc ttaaccgatt tctgtgcggt ctttgttgct tcttttccgg caaacttaat
20641 ggttttcttt ccggaggaag tggcagactg ctttatggtc ttttccgcct tatcgacagt
20701 cttaatcgca gagcttgtgt tttgcgaagc ctgttttctc aaagactgct cggctcgctg
20761 ctgcttgaaa cgctgaatac cgtcctttgc ggtgcggata ttctcttttg tggtttcaag
20821 tccttttcgt cccgctttgt caaacgcata agcaccatcg tgaaccacct cgtcaatacc
20881 gctttccagt ttatcggaag catattcctc ggcagaactg ctgttggcat ttaccgaccg
20941 ctccgccttt tcttttgttg cgatatacgc ctgcttcatt cgttgggttg caacggcagc
21001 cttatctatg gttcttatcg taccctttgc attgtccttc gttttaatat cagacataca
21061 atccctcctt cctatgcagg ctgaaccacc cgttgcgtta ttcgttaatg agcttcgcca
21121 ctacaacaag tctgccgttt gtccgggaat actcacaggt ggaaagggtc aggagcttgt
21181 cgccgtattc ggcggttacg cccgtatcgt aaagcgacag ttccttacac ctctcaacat
21241 aggctgtgaa atcatcggcg gtctccgcat tgacgaactg ataatatttg aagctgtcgg
21301 gactatccgt atataccacg gttttgaaca cggcaattac ttcataggtc tgtctgtcgg
21361 tcagcgtatc aaaggatact gttttgtgct tttcccaaaa gctcttgtcc ttgaatttca
21421 taagaccggc gaacatagag ccgtcgttca tatgatgacc gtagataatg atattgtcgc
21481 ttggcagttc catatcgcag ttttcctgaa cataaggaca gccgtagtcg gtgtatgcct
21541 tatcaaagcc gtgcttcaga tagaagttcg ggttatcctt tgactgcata acgggataat
21601 tgatttttgt gtcctcaacc ttaatccagc cgaccatatc gttattttgc aaataaagct
21661 cctgatactc cgggataaaa gtcttttctt cggagtagtt catcggctct tgggtttctt
21721 caggctcggt ctccacaacc tcaatcagat tgtcgtacat ctccgcctgc cttgcagagt
21781 cgtaatagtg cttgatgaga aaacctgaac tgacggcaaa tacaatgctg aaagcacaga
21841 ttatggcaat atagattttg tttttcatag gtgtttcctt tctcatttac agaatagggc
21901 aagggttata caccctcgcc cggtttggtt gtcatcagct tatacagctt cgtattcttc
21961 gggaacttgt ttgcaaacgg cacaagcgaa ctgccgacct taatcagtcc gtgtcctgcg
22021 tcaacattcg tgatatatga aagctggagg tcggaaatgt taaggagctt ggcaagctca
22081 attctgtcgg ttgatgcctg attgagcata atcaaaaact cgctgtttgc aagcatagtt
22141 cttgccgtat ggctctgcaa aaggtcgtcc acattctgcg taatgccggt ggcgtaagca
22201 ccgtactttc tcacacgctt ccaaagagta aacaggaagt ttgcggagta ttcgtgctgg
22261 aacagaagat agatttcatc aatgaagatg aatgtctgct taccсttagc acggttctgt
22321 gtaatacggt tcagaataga gtcgaggacg acaagcatac cgataggcat aagctgcttg
22381 ccgaggtcaa gaatgtcata acagataagt cggttatccg tatcaacatt ggttggcttt

22441 gcaaaggtat tcaaagaacc gtgagtaaac agctcaatag caagggcgat ttcctttgct
22501 tccggctcgt cctgtttcaa aagctccgca cgaaaatcct gcaaggtagg aacagtaccg
22561 gtatatcctc tctgctgata gtctcggtaa acacttgccg tacaacggtc gatgatagat
22621 ttctgctttg ctccgaggtt tgtaccgccg atgagctgtt cgcagaggga cataatgaac
22681 tcggatttca ggataaccgg gtttgcaccg tcaccgtatt ccttgttcat atccatcgca
22741 ttgatatggc tcggcgaagt ggcggaaata ttgatgattt cgccaccgag agcctttaca
22801 agctgcgaat actctcgctc agggtcgatg ataataacat ctgcattgga ggaaagaatg
22861 atgttttcaa tctctccctt tgccgcaaag gatttaccgc cgccggatac gccgagaata
22921 aaagagttgc cgttgagtag ctgtctgcgg tcggcgataa tcatattttt actgatgaca
22981 ttctgaccgt agtaaatgcc gttttcgtgg taaatgtcct gaaccctgaa gggaataaag
23041 accgcaaggc tttctgtggt aagggttctg aatgcgtcaa tcttgcgtgt tccaaagggc
23101 atagccgtat tcagtccgtc catctgctga tatttgagaa cggcaaactg gcacaaatgc
23161 tttcttgccg tagtaagcag agcttcggtg tcgttatcaa gctgttcttt ggagtccgca
23221 gtatgaacca tagtgagtac agcgaacatc attcgctgat cacgagtggt aaggtcatca
23281 aggaactcct tcatttcctt tttctgctgt tccatatcat acggaacggt agcggaaaag
23341 ttattgttct gattctgctt tctctgccag ttggttatat tcgtctcaac cccaagcagt
23401 ctgctctctg cttctcgcac cgcttcatcg gtcggaacgg gtacaatgtc aatagacatc
23461 atcaggttgc ggttcaggtc ggtaagctcg gctaccatac tgtccttgat ataggaagcg
23521 tactcacgaa gaaaaaggac tctcccgtaa cggtttccta ttttgaagta gtccttttca
23581 agttccattg tatccgggca aacataatcc ttgaaatcgt gtcccttttt tctcgtttcc
23641 ttaatattga agtggtatgc ggtttcctca ccgacacgga agaaatcgtg gacaattctc
23701 agcctctcat cggtttcaag ctccacacat ttgcttccga gtcttgcaaa gtgggcaatc
23761 aggtcagcac cgacacgggc aaaataggtt cttgcgtcct ccacgctctt tttgttgatg
23821 gagatagtaa catacttatc ctgaacgatt gcatttgcac cggtcgcctt gtcgagaagc
23881 attttgttat attcctctcg gtaaacatca agctcatcgc cggtttcggg gataagaatg
23941 gttttctcaa agtccaaacg gttaagccgt ctgttgttga tagtcagttt tgtagtagct
24001 ccgctgtcga gagagttgag aagctcggag tattcaagga acattgcttc cttgtcctca
24061 cggcttgcta cggcaaagtt aatatccgtg aacttgaagc tcttggaata cttgtccttg
24121 cccacacgga aaatgccgtc gtcataaata gcggtgatgg gaatacaatc ctgtacgccc
24181 tttggtacta caaacttttc tttgtcctgc tttaacaggt ttgtaagggt tttaatcatt
24241 gttcttataa gcctccttct gtttttgttc gattgtcggt ttcatcaatt cgtaataggt
24301 gttggtggaa tgaaatacca gctttttcgg cataaggaac tcggacttta tccacgccca
24361 tataaatttc tcagccgtca ttccgttgta tttcacaaag ccgagagccg caaaaggggc
24421 agctcccaaa atgcacaccc acgatacggt ttctgttccg aggtacggtt ttaggaggaa
24481 gtaaagtcct accgctaccg cacaggcgag gacagaaaaa atgaactgtc tgagcgacag
24541 tccaaaaaac attgactctg tataatttcg tatctctcgg ttaattttga cttccatcaa
24601 aagctccttt ctctgcgata tgcttgtttt gcgtgacgga tttcatcaga gcagacgcag
24661 gtgtccgcaa ggcaaatcac attgccgttt ttctttcctc cgtaatacac acaatatttg
24721 cagtcgcagt cggcaggcgt atagccctgc catctatcgt tattattctt gtttttcttc
24781 atcagagaac ctcccgttat aagcccatca tttcacggat aattcggtcg ctcatcttaa
24841 ccgcaccgac aagcacaagc atattgaata ccagttcgcc gatatatgcc catacctgag
24901 taacggctgc tgcgtcaggg ttcactaccg gcggcgtaga agcaaatagg gagaagatga
24961 tacaagcaag tacaattact gctccctcaa gcaaaacggc acagtagctc tttataaagc
25021 tcttacccac attctgcgac ggttctccgg caaatgtgga aagcggtatc ggagcgagag

25081 cagtgtacat atacagcttg aaaaatcgtc cgtacaccgt cattatcata atgaacgaca
25141 agaccgtgat aaacagtccg ccgataagag taaccgccca caatgggata ctttcaaaaa
25201 atccgcagct ttcaatggtt gttaccattt cagcaggtaa agttgtttga ttgggcgtac
25261 cgaatcccgc agatgtcata atggttgaaa tcgtaccttg tacgatgtcg aacagggcaa
25321 gcataagctc cagcccgtag gtaatcacac ctttggcaat ggcaaagcgg ataaacagct
25381 ttaatgcgtg ttccggtttc tttacatcgg taaaacttcc gcaggtgcga accacaccga
25441 caacaaaaaa cagaacaagc agggcaagac cgatagcctg aactgctccg ttgatattca
25501 ccattaccga ccaaatactg ccgcctttga actgttgcgg cgaagtggtt agcaactgcc
25561 atatttccga aagcttgctg ttccaagttt caagggcatt ttcgaggttc tgaactaccc
25621 agttatcact catctgtttg cacctccttt aagattaggg gcatacctga aatgccaggt
25681 atgccccgtt tgataactgt gttctcttaa ccggtaataa gggtaaggat ttcctttgca
25741 aaggtaatga tgataccgcc tgcaagagtc aggaaaccgt tggctctctg agacgggtcg
25801 tgagacttca aagagagacc gacctgaacg ataccgaagc caagcaggat aagaccgatg
25861 gcacggataa gaccgaagat gaaatcggac agattgttga ctacggtaat cgggtcgctg
25921 gcggcaaaag cggtaacgct cataccgcaa acaagagttg ctgctgcgat tacggcgcag
25981 taagcacgga agcctttctg aacctttccg gtcataggca gcttctgagt agttttcttt
26041 tcgttgttct tcttaaaaat gttcattgtg ttttcctccg ttaaaataag tttttattca
26101 cttcaaaatc ctcatcggac agaagttcgt aattggtttc tccgtagctt tcctccggaa
26161 gttcttccgg gtcaatagaa agaacggaaa ttgatgagtg cttgatggtc acttcgccgt
26221 gtttatatgc agaagccttt ccatcggttg taagtgccac attcgggtgc ttcaagatgt
26281 cgtacttgaa gtccataacg gggcgttctc cacgaataaa cagaattgca tattggttat
26341 caagcatacg aacctcatca ggagtaagaa gctctcgacc gctaatctga taatttgtgg
26401 agtagttgcc gcttcgaccg gtactttttc cgaaagtgtt ggtatcaatg gtttccttgc
26461 cgagaagctc ggacacatat ttgtgggtgc tttgctcgtt gccgccgaga tataaaaact
26521 cgtcgcagtt acctacaatg ctttcccatt gcttttcaaa caaggcttta agctgtgcca
26581 agttctgcaa gataatactt accgataccc cacgggaacg cataaccgaa agaattttgt
26641 cgaagtcatc agggagtgaa acattcgcaa actcgtccat caaaaaatga acgggaacgg
26701 gaagcgagcc gccgtgaata tggtcggcag cataaaacaa ctgctgaaaa agctgtgtgt
26761 aaaggataga taccaagaag ttaaagctgg aatcgttatc cggtatcaaa gcgaacagag
26821 cgaccttttt ctctccgagt gacgggaggt caagctcgtc tgtggtggtt aaagaagcaa
26881 gactttcaag gttgaacttt tccagtcttg ccgcaagggt tatctgtatg gactttaagg
26941 ttttcgccgc acctgaatga taggaacggt aatacttcaa agcaatgtgg tcgggattcg
27001 acatctccaa ttccgcaaac agttcatcaa gaggagaagg tcttgtgtcc tcctcgtcct
27061 caatcgcacc ggctctgagc atttccatta tcattgcaaa gttctgttca tcttcgggtg
27121 cttcataatg aagataaaat acaagtgcaa gcaaaagcat tgaagccgag gtatcccaaa
27181 aggggtcttg tgattgactt cctttcggag tggtactctt aaaaaggttc gtcacgagct
27241 tttgcacatc gttgtcattc tgcaaataga caaacgggtt atagcaatgg cttttctcca
27301 ttgaaatcaa atcgagaaca cggacttcat agcctttgct ttcaagcagc tttccggtat
27361 ctcggagaat ttcgcccttt gggtcaagaa tgacaaatga tgtgttgcac tgcataagat
27421 tcggcttgca gtaaaacctt gtcttaccgg ctcccgaacc gccgcacaca agggtgttta
27481 agtttcttcg gtgtttcttc gcattgagtc cgatacagac attttgcgtc atcagcttat
27541 ttgcagaagc cggagactgt ctgtattttt tgtttacggc tcttgcattg ccccattttg
27601 cagaaccgtg ttcttcacgc cttcggtaat tcttctgtgt tgagaagtaa atcccaatac
27661 ccattgcata gcagagaagc aaaacgagga cagtttttag gctgtcctcg catagctcaa

27721 tatgaaatgg ttcgttgaac gctgttaata ggctcggcag tatttccggc aatcctcctt
27781 ttacagaggg tgcaatcaga agtgccagcc atacgaccgg gataataccg atgacggaaa
27841 gaataatagc agactgcctc tcgttatctc gattcacagg agcgttcctt ttcgataact
27901 ttgaactttt tctttggggc agttccgacc ttgatgataa ggtcgtccac cgcatcggta
27961 gatttgccct cgttcatcaa atctgttttc tcgtcgttga gagaacggat aataatgccg
28021 tgttcataat cgtccaatgc gatatgatac atttcttcct tcgccatagt ctcaccttat
28081 ctttcgtagt ctttgcttcg ttcctgcttt ttcttgtaaa gctcatcgga aacacgggag
28141 tgaatatccg ccattgcgtt tctcgttttg gaaagctcat agcgaatgtc cttcggctct
28201 ttgccggcga gcttttcagg gatacggttg atagcaaaat tctgtgtatc gacgccgtac
28261 tttttgcaaa gcatatagcc gatacatact gcctgaaagc ccatatccgc tctgctgtaa
28321 ctgtcgctgt tgatagagag ctgtgcgtgg ccaagctcct gtgctacgca ctgagcgacc
28381 gctacgctgt ctcccacatc acgctttaca tacaaggtct gttcctcgtt gttataaaac
28441 gccgccatat tgggatacgg aagctcattt gttgcttcca cttcaaccgg ggaagaatcc
28501 agcataacgg taatgagtgc tttcgggtca cggttgacag tcggtgcagg cggtcttttg
28561 ccgttggtct gcgaaacatc aaagaccttt ttcacattgt aggaaatgcc ggtcgtaccg
28621 tctttctttg catactcgac aggttcaaga atggaaatgc ttttcgcacc tttctgaatt
28681 ttcacattat ctttgctcca gtcatcaaag tttttaagct ggcttgcttc gggaagctgc
28741 ttgaaaatca gaagtgcgtt gaccgcagag tatctgtcaa gacggctctg cgtatcaagg
28801 aaggctttga acttttccgg gtcggaaaca atcgcctttg cggtgtcgcc cgccatctga
28861 tatacggctt ctttttcagc cttcatcttc tcggcatatt cctctttgga aagtcgtttg
28921 ctctgctgtc ctgccttagc aggggtaaaa ttgtttgtca ttgtgattac ctctctttca
28981 catttttgct tttaggtttc ggctgctggt gtacggtctg tgccggggcg ttcttgactt
29041 ccggctttga tacctccggc tctttgcgtt ctgcttcttt ctgctgcttg gactgtgcct
29101 tatatcggtc aagctttgct tttaccgacg gctttctctg tctgtcgctt acagtaccct
29161 tatcagtttg catatctacc ggctcggagt cgtgccttga cggagggttt ttgtccgttt
29221 tggcgactgt ggggtttgac tgggaatacc cctctttctg aataggtttt ctgacggctt
29281 cctcagtgat gatttcgttt ttggttttcg tcggcttgtc tttctccaaa gcttcacgct
29341 tttcgacagc cttctgagac tcggtgacaa tcgcagcctt atcaacttta ccgagttcaa
29401 atctctcaac gattctctga attttggaag cgtcctctgc acgggcaata atatcaatcg
29461 ggacattatc gccctttgtg tttttatccc tgagtacgca gtagagaacg ccgtagcgtt
29521 ttgcctgctc ggtaaacttc ttcaagtcct tgttggggat agaaaacacc ttcagctcct
29581 tgcccgactt aatcatatta gtaagtcgag ccttgccttt tgttttctgt tcctgtttga
29641 gaacagagac taaaagaacg gcaatattct ttgctgccgc tccgcttaat tttgcggcta
29701 cttcaaagcc ttcaagcgat agcctgacga cctgttctgc cgcatcacct ccgtttgtca
29761 ttacggttct gctcctttcg ttcttgtttt tcttcgtaaa ctcttaccgt ttcaaggttg
29821 gcttcaataa cctttgacct ttcggctatg tcatcacaaa gaccgacctc tttgcggaga
29881 gtccacagct tgctcgtgat tgccgtaatc tgctctttga catttgatag ctcatcgtca
29941 ctcaaatttc ttcttaactg ttttcgcagg tgcgttctat cgtcagtcaa ggatttgatt
30001 tgcgacaaca cggactcttt atatgagaaa agctgctcgt cggtagaaat gttctcccgt
30061 ccaagcagcc ttgtctcctg agtgattttg tcgagcttca gcaaatcttc tttcaaaaga
30121 tagtgaagcc ttgcggtatt ctgtttttg tacttcggca gataacccag cttatagcag
30181 taatgcagat acagcccata cagtccgccg accttttga ctttgtgttc tcttgtgggc
30241 attcggtact gcctgactct taccgtttcc tttgcaaaag gaacgataag aaccttttga
30301 ttttccgtca gcctacgctt gattgcgtcc tcggtataat tctctccgag tgatttaagt

30361 cgtatcggct tcttgtatcc tttcgggaca atcgtccaat actttcgact gtcgctgaga
30421 caatgttcat agcccatctg agtcaggata taatcaaagg tctttaagtt ggtgctgtgt
30481 gcaatcgctt catcaatggc gtcacgggtc attgaatatc ggctcggcat accggcttgt
30541 tcctgcttgt agtaataaga ggattgctta ctgcggtttg ggttcggatt gtaatacagt
30601 ccgtactttt cacaaaggca gtcggcaacc aaacgaaact tgaaccacgc tttttcgtcg
30661 ccccataacc gcttgccgtc cacaaaggaa acggagttta ttacataatg acagtgcagg
30721 tgtttcgtgt tcagatgagt agtaaccacc acctgaaatc ttttgcccca cacttcgttt
30781 gcaaattcca ttccgatttt ctgtgcaagc tccggggata tgtcctgttc cttaaagctc
30841 aggtatccgt gataggcttg aatgccgtcg tctttgccgt atgctttttt tacagatata
30901 aactggtctc ttgcggtcgc agggttgcag ttgataccct ctacgaaaaa ctcacgctcg
30961 gtttttttctt cgtcctttgc atagctgaga acatcacgga gagcctgata ctgttcctcg
31021 gtgtaaatgt ctgccgccgt cttttcggga ttggcggcat agtcgattac ctgaccgagc
31081 cttgatgtga ccgaccacaa tttacttact gccatttcat atcgctctta tcgggtctta
31141 aaaaagtacg ctccacatca gcctgaaact tgtgccaacg ctcggcttcc tttttgagct
31201 gcggtgcgtc cacaaatccg agggtgtttg ctttcgccgc aagctgattg atgttgttgc
31261 cgatagcgga aagctcacgc ataacatcgt aaaatctttc atcgggcttt tctctcggct
31321 cgtatcctct gagaagcaaa cggataagtc cgccttcgga aagacacgcc cgctttgctt
31381 ttttctgcaa atcctgagct tctgccttgt tcatcagaat gtgcttttca atgtttctct
31441 tgctcatttt tatcgctgtc ctttctgtaa atatagtgtc tgtcccggtt tatctgaaac
31501 aggcacattg ttgttactcc gaaaaagccg ccgatgagta acccggcggc gaccaatact
31561 atctcccgca aagcttatgc tcctttcttg cttggggtat taggggcaac ccctaacgag
31621 tgtaccgttt tggacattcc tgtccataaa tggtctgctc gctaagatta tgcagacact
31681 ctgggttgcc ccatagattg actcttatct ctccatatcc tttgagacag gcttatacgg
31741 ctttgcttca ataccgtcca aggacttata aaatctctcc gggtatttga acttttcctc
31801 gtacttacga gccagttctt tcggcatact gagaaacttt tccgactcaa tcggtgcgta
31861 cgcaatgaaa aacttgccgg caataatgtc gagcatttca cggttgctgt cgtcttttac
31921 ataacatttt tcaatcttcc agccgtcctt gccgccgtgt tcatcagcca tatacgcttc
31981 cagtcgcaca catttatccg acccgtcaag ggaagaagca tagatagaat atccgcccat
32041 accctcaatg aaagccttat tgttactgct gacaacatag gttctttgtt cctctgtgta
32101 aggcttgtca aagctgtcag ccgtaaaaac aatatatccg gtcgtgtggt ttctttcttt
32161 ttctgcctgc cggaaatgag ctttcaactg cggataactc atctcaacat tctcaggctc
32221 ggaataaact gccctgttca gcggcaagcc gttcatcttt ccttcatcgt tgcagatgat
32281 ggcaacctcg tcctcaaagg gcatatattc ttcaatatct ccctccacaa gagactgcat
32341 tgcttccagc gtatcctcaa tttcaatgag cttcggatac ctgcccggct caacaagaat
32401 tacggaaatc ttctcaggtg cttcgacctt ttcaaaggta tcctcgtaaa tattcaaggc
32461 gagggagctt ccgttttccc aactcatatg aatctgtccg gcgtcgtcaa caaacttaac
32521 cgttcctttc agaccgtatg gcatttgcct ttcgccagcc atttcagtga gtacaatctg
32581 cgtacccttc gggtacattt ctctgatttc ttttagtttc atcgtgcaaa actcctttcg
32641 gttttatcct ttatccctaa gcgtttacaa agaaaatcgt tgacatcttt cccgtattgg
32701 ggagggtcgt caacgacttc atacttatct gagagaatag tctgcaacgc tttggtagct
32761 tttcttccgg caatatcatt gtccaaatga agaacaattc gcctgattgt tttgtccttt
32821 tcaagtagcc tgccgagtgt aacaggcact ttcgagtcct cgattttctg cttgggggag
32881 tacactcccg caagagaaat aaggttaagc tgtcgccagt ctttgccctc cagcttcatc
32941 aaagttgcat aggaaagcag gtcaatggca cactcaaaaa gatgaacttc tccggtgttt

33001 tcggcagtca gacgaaaaga atattgcttt ttactgccag tgcagtcgcc cataatcctt
33061 gattgatttg ttgctctgta agcggcgtat ttaggttctt tgttctcgtc atagccgata
33121 aaaacagcat tgtgatacgg aagcgactca atgattagct cctgttccaa gcagtagttg
33181 atgatttcaa aatcaatgcc acgcccgaaa aggtaatcaa ataccacatc ggcagtagga
33241 ctttttcag gcagaagcaa gggttgttcc tcatagcttt tgatgttttc acaagtcgga
33301 aagctgactg aaccgtttcc cataatggtc tgaaccgcct ccacaaaatc cattccacgg
33361 actttgataa ggtaatcgag agcggatttg cccccgatac ctctcgacca ccacatccat
33421 ttgccgttgg aaatcttcaa gctgtcgtgg gttcttgtgg tgtaggtgtt gcttgaaaat
33481 ttaacaagct cgcccggttc atactctctg agataggtta ataagtccat tcgtttggct
33541 tcggttatga cttccggtgc aatatacggc atacaaacca cctcgcttta ctcaaggcaa
33601 tcaaattcct tatcacttat ctcatcgcaa caagcgagtc ttgcaagcaa taacagaaca
33661 tcgtcaacag gacactttat gtcgttccag aaaagccaag cgacttctgc aaggtctttc
33721 tcctcataaa gagcaacgat ttcttcaggg gagaactccg tctgaatact tctgtcaata
33781 agttctttat agaaattgag ctttgcaatt ttaagtgaaa attctgtttc ccttgcgggc
33841 ggtgtgcttc tttttaattc gtgatacgca ttgctcaatc tacggacaac gcacatctgc
33901 ggtgcatcaa acggttcgat ttcttcattg gcaaaaaaga aatttgcctt aaaatctgtt
33961 tctttcttat tatctttcat aggttttctc ctttcgtgtg ttcggcactt cgccctcatc
34021 gggttcgccg ccgacaattt cgttatctcg tttatccata ttgagcaaga tattcagctc
34081 gtccagtcgg gcaagctttg ttttcaattc ttcctcttgg ataaatggtt tttggacttc
34141 tttctttgcg gtttccaact gatttttggt gttctcaagg tcattttccg caagcgataa
34201 gctctcggca aatctctcaa ttccgttatc gaggcggata atattgccgt atgcgtcatc
34261 gccgagagga acatttctgc ttgttgcacc tttcagtcga acctcataat ttcgctctgc
34321 ggtattaaaa agcagctccg tttcaaaccc tctgtacttg ccgagaggaa tagggtcggg
34381 gctgttcatc tgcttgcagg cttcaatgat tgctttaccg gcgtcggctt tttcggaata
34441 aaaaacaccc ttaacctcca taccgacaaa gcggtcgtcc gttggtttcg ggtgctgttt
34501 cgcagtttct acatcttgct tcaagccttc aatgcggttt tccaaagctg taattctctg
34561 cggataatat ttgattatct tatcttccaa agcgtatctt tcagagagaa agttggattt
34621 taagagcttc agcttttgga cttgaatgtc caaatccatc ttctccttaa tatacgggtt
34681 gccggtggca agcattttaa tctctgcata ggacagagcg gtttcatcaa tatcctcggc
34741 agaacgcacc ggggatttgc tcgtcataat ctgactggcg aacttttgct ttccctctac
34801 aagctgatag agatatgcgt cgaaggtctg ctctgttaca tagcggtaaa tatctacatc
34861 ggggtttgag ttaccctgac ggataattcg accggaacgc tgctccaagt ctgacggtct
34921 ccacgggcaa tccacatcgt gaagtgcgat aagtctgtcc tgaacattcg tgccggctcc
34981 catcttctgc gtagagccga gaagcactct gacctcgccc ttgcgggtct tttggaacag
35041 ttctttcttc ttcacatcgg tatctgcttc gtggataaag cggacttcgc tttcggggac
35101 accacgctct atgagtttct ttctgatgtc gttatatacc gagaatgtgc cgtcgttctt
35161 tggtgttgag aggtcgcaga acacaagctg tgccgatttt ttatcggctg tttcctccca
35221 aatacggtag atgttatcga cgcaggcgtt gattttactg ccctcaaaat caggcagcat
35281 atcgttgagc atacgctggt caagagccaa ctttctaccg tcgtttgtga ttttgagcat
35341 attatctaca ctcgaatcta caccgccgcc acggactttc tcggctcgct ccgcaaggga
35401 agcaaccatc tctttctgca tttccgaagg cttgaccgca acattgtgga agtgtgcatc
35461 cggcacaggc agctccagca tatccgccgt cttaatgtcg gcgacctcct taaacatcgc
35521 cataagctcc ggaaggttat agaattttgc aaagcgggtt ttcgctctgt aacccgttcc
35581 ctccggggta agctcaacgg ctgttacagt ctctccaaag gtacttgccc acgagtcgaa

35641 gtgttgcaag ttgtttttta caagcgtgtt atactgcaag tatcgctgaa tggtatagag
35701 ttccaccata ctgtttgaga taggagtacc ggtcgcaaat accgttcctc gaccgcctgt
35761 aagctcgtca aggtatcggc acttcataaa caggtcggag gatttctgtg cttctgtctg
35821 agcgataccg cccacattac gcattttggt gtaaagatag aggttcttgt aataatgact
35881 ctcgtcaata aagagcctgt ccacaccgag ttcttcaaag gtaacaacat cgtcctttcg
35941 gctttggtcg ttgagctttt ccagcttttg cttcacggac tttttagact tttcaagctg
36001 cttaatggag aaattttctc cacggtttcg cttcaaatcc ataataccgt ctgtcagttc
36061 ttcaagctgc tgctcaagaa ttgccctctg tctttcaatg gacatcggga ttttctcaaa
36121 ctgagaatga ccgataatta ctgcgtcata atctcccgta gcaattcgac cgcagaactt
36181 tttgcggttc ttcgtctcga aatctcgctt tgtcgccaca agaatattcg ctgccggata
36241 tagctgcaag aattcagccg cccattgttc ggttaagtgg ttcgggacta cgaacagcga
36301 cttattgcaa agtcccagtc gtttggactc catagcggcg gctaccattt caaaggtctt
36361 acccgcaccg actgcgtgtg ccaacagcgt attgccaccg tacagaatgt gtgcaaccgc
36421 atttttctga tgttcacgga gttcgatttc tgggttcata ccgttgaaaa caatgtggct
36481 tccgtcgtac tcacggggtc ttatgctgtt gaacttttcg ttatacatct ttgtgagttt
36541 ttctcggcgg gtagggtcag cccatatcca gtcctgaaag ccctgcttta tgagttcctg
36601 ctttgcctgt gcgattgccg tctctttttt attgagaacg gctttctttc tgccttcctc
36661 atcttcgata tagtcaaaga ttcggacatc tttcaggttc agtgtttcct caattatctt
36721 atatgcgttg atgcgggaag taccgtatgt gctgtacgct tttacattgg aacggtcgta
36781 gctctttccc tcaatgttcc attcgccggt aaactgagag tagtgaacct tgatattcca
36841 ttgtgcatag cgtggggtgt cgagaaactc aaacataaac tgttggaata tttcggcgg
36901 tatccaagtc gcaccgagcc ttacggaaat ctcgcttgcc gtcaaatcct ttggctgcac
36961 cttttcaagt gcctgcacat tgacggtgta atcttcggga tacagtgccg cagacctttt
37021 ggctgtcgcc agcttctcac gaacattgcc cgacagatat tcgtctgcca tcagatactt
37081 cggctcatag gaattaccgt actcataaag gggatttaag aagattacac cctttaagtc
37141 agcgaataat tcctcctcac ttttgccgga caactccatc atatattcca tatcaatggc
37201 tgctttttcg cccatagaga cagcgagagc ctcgtctgcg gtgtcaacct ccgttaccgg
37261 cttgtgcggt ttaatcgtcc ttttgaagaa catatccgcc ttgcgttcca gttcgccatc
37321 ttcattgata acctcaagag cggagagcag agcataggag ctgtcgtcgg aaaaggcaga
37381 agtattggca cggctgttaa tcagtccgta cttctttgta aagctgtcat acagtgtgtt
37441 cagcttatcc tgtgcctgtt tgatttcgct gtccgggtaa tcctcggttt gcagttcaat
37501 caagttacgg acgcaatccc ttatcccaat catacccttg atacggttct cggcggttgc
37561 cgatacctct accggagaca tacgggagtt ctcacggaaa tagattttat cgtccaaaac
37621 cgtgtaagag aagttcctta ccgtagggtc tgccgggata gaattatctt cttcctccag
37681 ctcgtcggca acctcgtatt cggagatttc tccgtgaatg tttgagactg cttcgttcaa
37741 aagctccgaa aggtcggcat tttcaaacgg ctcacaggtt gcttccggtc cgaaacgacc
37801 gcttaccatt ttcatttcgc cgagtatcat ttccgggtgc tgaacaaaat atgagttcat
37861 ccttataccg ttttcgtcgg tgtcaagatg aacccagtca ggctcaatat ctattagcct
37921 gtctcgcttt tgcaaaaaca gaatatcgga tacaacctct gttccggcat tgcctttgaa
37981 agtgtcattc ggaagtctga ttgctccgag cagctccgcc ctctgtgcga tatattttcg
38041 gacattggag ttttccttgt ccatagttcc tttgcttgtc acaagagcca tcacaccgcc
38101 ggggcgtaat ttatcaagcg attttgcaaa gaaataatcg tggattaaga agttgtactt
38161 gtcatatcgt ttatcggaga ccttgaaatc tccgaaaggc acgttgccga taacaccgtc
38221 aaagaagctg tcggggacat ttacttcctc aaagccctgt gccgcaatgg aagatttctg

38281 ataaagctgc tgagcgatac cggctgaaat ggtatcaagc tcaacaccgt aaacctttgc
38341 gttttccatt gacttaggca gcataccgat gaagttaccg ataccgcagg acggctcaag
38401 cagattgccc tcctgaaatc ccatctgttc caacaccttg taaatggcgg tagatacttc
38461 gggcggagtg taaaaggcgg tcaaggtgct ttctcttgcc gacgcatatt cctccggcgt
38521 caggatattt ttcagcattg catactctgt atgccaagct ccggctcgtt cgtcaaaggc
38581 ttcgggaata ccgccccaac cgacatatct cgacaagatt ttctgctcgt caggtgtggc
38641 aaagcggttt tcttgctgac accttttcaa gaccgtaatc gccgcatagt tgcggtggaa
38701 acgctctttt ttattgactt cttcaacctg attattggca aggtcaaagt tgagtcgctc
38761 tgacatagga atatccgggt ggaggtcaaa gctctttacc tttgactttt tcttttgctc
38821 ccaagcggga actatcaact caggttcttt atctcggtcg tccttaactt caacgccgtt
38881 gtgttcggat aaagccttaa agtcatcaag caagtcaggt ctgataccaa gcacagaacc
38941 gtcatcttca aacacaaagg cttcatcaac aagaaaatga taaaactctg caccgtgcca
39001 ttcgttatcg ctgtcccttg caacaagacc gttttcatca taactgagtt caatgtcgtc
39061 aatctttaat ctgccaagca cggaataaag aatttctgtt tctcggtcat aaaccgggac
39121 ttcggcggca gcagccacaa ggctttgcat tgtcgctttg gtacagtctg tgaaatcagg
39181 ctgctgtgaa aactgacttt ctgcttcctc aaaccattcc gtacctacat cggcaagctc
39241 ctgatttgaa atgctgccga ggtaatcaaa gaaatcctca gtcgagtcat attcccgtgc
39301 agcttgctgt atctcgtcaa aagaaagcgt atttgtaaca tactgacctc cggcattgct
39361 gtcggggtta tagtacatcc aaattacact ttcattgtcc cggtcgataa agaaggcgtc
39421 gtcataaccg gttttcggac tgaaatcctg tacggtttca gtatcgtttt gagcttcatt
39481 ttcgcctgta acggcttttt cttccgccgg taatacactg accttcaagt ggtcgttcat
39541 tggattttca cgcacttttc ggtcaaattc ttcccttgga agctccttgt tgaaaagggg
39601 catatcaaag tcatagagca ttacacggtt ttcgtcaaag gataaaatct catactgaga
39661 agccccgata tacacgctgt cgccgagatg atattcatac tgtgcttccg tttcgtccac
39721 tttgttttca ggctcaggcg gtgcggtaga gagtatctct cggttttttc tctgctctgc
39781 aagctcggct cgccgttctt cctgcttttc aagccattcc gggtactttt ctttttcctt
39841 tgggtttagg tatctgtcca tacggataag gtctccgata cgcttttcca cctgagacca
39901 agaaagagta atgtgcggtt tgtcgcttcc gataccttttt gaaattgtga taccttttcc
39961 gtcgtgctgt tcgtcaatgc ctgcaccgat gatgaccgga taagagccgc cccaaccgta
40021 ttcgtttttc aagaagtcgg cgttctcttt tgcggaaagg ctcttttcaa actgttcata
40081 gatacgcatt ttgccctcgg aaactccgct tccacgggtc agtacagcgt ctatgatttc
40141 ctgagaaaaa gtaaaggcag aggtttttg ttcctctgcc tgatttccca taatgagagt
40201 caactgaccg tccatagacg gaagcggctt cattgtatcg gtcaattctc ccaaaagtct
40261 cattgcttca aaatgctgtg cgataacgcc ccagtcgtag aagctctgag cggttctgtt
40321 ttcgtattcg ccctcccaca agtggaggac attctgaaat gttttgtagc ccacaagtct
40381 gccgtcgttt aaggtaagct gggtgtaatc gttgttgaaa ataccccttga tatattcggt
40441 acgggtcgca ttatccgtat ttctctcata gaagtctttg atttcttcct ttgaagcgga
40501 caaatgcggg gtagtgccga gtatctctcg gatagtatca tcgccaccga aaaacggcag
40561 gctcttgtcc tcgtggtttc tgtcgtaata ctctaagcga atattacctg attcttcacg
40621 atttcctgtg ccgagttcct gatgttgttc atcagacgaa cccacttcat catatcttcc
40681 gctttcatct gctccgtcag cccggctttc tgtgccatct gcttcacgag agtttcctcc
40741 agttcgctgg ctctctgctg aacctccgcc aagtgactgt tcagaatccc tttcgtcaga
40801 agattgtagt acaggatttt gtgatgttcc ttcagataag ttctcctcat ctgcgaccat
40861 ctgccgagag tcacttccgt ctgctccggc agcttcaggt tcggaagatt gtagtctccc

40921 accatcgtgt atgtgatttc gctcatcgct ttgactcctt tcgtctttga tgtctttatt
40981 gtattcggat tgctcctgtt catcaattat gcgattttcc ttgctgagcg tgctgattgt
41041 tctgctgaca gggagcaatg ccatttcagc aatgtcgctt gtagcaatgc ccaccgcatt
41101 gagaacttcc tgtgaattga agcttgaaat cccggcaaag tcatcgggag aatagacctt
41161 gtctgcgttc agtcccaatc gagccataac catatatgac acgctgtttc tgacaagctc
41221 caaatacaag gcatttgctt cgtcggcaga caaatactca atatcgctgc ctgcacccaa
41281 actcataaag tcggagatat agtctgcgat attgtcctcg acggcatttg caatgctctc
41341 tttgacgaca ttttcaatac tcgttgtatc gtttaccgcc ccgaaagtat tttcaagcgt
41401 ttcgataacc tctgcttcat attcgggtct catctcccaa ataggaacag ggcgggaatg
41461 tcggctttcg tgggtgtccg atatatcaaa gtaatgaatc agcctttgac ggcttctgtc
41521 tgcgtcctca aatgcggcaa taccttttgc acctcggttt acccaacgac cgaaagtgcc
41581 gttccatcgt tcgatttcaa gcacggctgt tgcatcgggt ctttgggcat aaatgagaag
41641 ctgttcatca aatcggagcc tgaaattcct gcctgctgtt ttcagaaacg attgccagtt
41701 taccggattg tctgtaaccg caaagcaggt tctttcatac agctccgaaa tcaggtcaaa
41761 tttccttgcc acctcgtctg cacctcctta tcgtgttttt tctttttcct ttgcaggttt
41821 tgtgttttct tttgcggctg gttctacggt atgaccgttt ctctcgcaaa gctcggcaaa
41881 ttcgcagata tggtagagat ttgtgccgac ctccaaatgg tagtcgtcga tatatctgca
41941 agttctctcc attgttttgc cgtcgccgag cttaattgag attttttctc cgtccgcaat
42001 gcgaaactgc tcctgatagt gcgggtcaat aaatcggata ccacgctcgg catttttcaa
42061 gtggctgttc agccactctt tctgatagca atagcagtac agattgtact cgcccctatt
42121 ggggttaaac ctcataagat agctgtactt ttccgtatca agacgaatgc cgtagtggtt
42181 gcggtcatca ttaaaggaac tgtcgggagt ggaataacag taagaggaca tagcctttct
42241 gccggacaaa acatcgccgt cacgcaatcc gtttatcaca tcgtcaaatt cagccttaaa
42301 ttcatcggtt ttcaggtctt ttctgaaatc gttccaagtc gtccaaaact cgttcccggt
42361 agaaccgaag tcggcacgca aatatccgat aagaccggtc tgcatagaga gctgctgact
42421 ttggctgaag gtgtatttac gctctgcttc ggtcaatact ctgtaatcca ttttcatcac
42481 ctcgcttgtt cgtttggttg tctttgcttt ggaatagaaa aaccatatac gccggcaata
42541 tcatcgccga gtcttttggt aactcgtgtg atgtcggctc ttgtggcttt tccgttgtag
42601 attttaactt gcagattttc aatctgcgta tcggtaacac cgtctttgaa tacacggtaa
42661 agataatggt ttgtaccgtc gtgatgaaca gcgtcggctc gaaggtctcc gtatttgtca
42721 acataccatt ctgtgtagtc ggtgtcagaa taaaggcagt ccttaatgtt gcctgaatca
42781 atcatcttgt atcccgacac tctgccgttc catcggccga ggtcgccgat tacgataatc
42841 ggctgtgaaa gctgaatgtt gagatttgct cgttcatcat aaagattttc ggcattggat
42901 tttaccataa tgtcgtaaag ctcatcatcg gaatatccgg gatacagttc ctccaaagac
42961 tctcgccaat catcaaggtc aaggctaata tcgctccaaa tgatatggcg gtcattctgt
43021 tcctgcctgc tcatcagtgc cacctcccaa ataagagaac agcttattgg ctttggtctg
43081 cttctgaaac tcgataatca ggctcatatc cggaatagtg ataccgggaa ttttcaaaat
43141 gctctccata tcaagctgct gcaatttctt ttcggtattg cagcctgcgt caaacagttt
43201 cgtcagaagc tttgttttct gttgaaaagt gaatttttcc ttcattgctt tgtcctcctt
43261 atcgttcttt gtctttgttt cgcttcttgt ttgaaagctc ctgtgccttt tcctttgccc
43321 attcaggcag acgctcaggc tttacctcgc ccataacatc cattctttca tatcgggttg
43381 atttgccggt gtgaagattg gtacaaaaca cggcgcttcc acggctgttt gccgacgcac
43441 cgaaaccacc tgtaacatag tagagctgtc tgtcggctct ctgaaattca ggctttaata
43501 ccttggggtc aatggctatg attttgccgt tgatgtcttt ggaatagtgg tcggggatac

43561 agtcggcttc tgttatgagt gtgattggaa tatccagttt ttcgacctgt tccttgaaaa
43621 gctccgcttc atttgccaca cggcttccga aaaggtgtac gatttcgata tagtcatcgc
43681 ttaccataca ttccgaacac agttcaaaca ggtcgtttcg ctctgcaaag ccgcacataa
43741 atttatcgcc gtttgtactt tgctcattgg aagccaagat aacttccttt tcgcccacat
43801 tgaccgagca aagaacagtg tattcaccga tttttctttt tccctcgttc atatccacac
43861 ctcagcttac ttcaatcaca attctgatgt ttccgcagtc gcagccactg cactgccttg
43921 acagttcggg aaatagggat tgcactctct gtcttgctct gcggatagtc tcaaagcacg
43981 gcaagccgta gcctttatag ttgttgacta tatcctcaag cggtaaatct ccgactctca
44041 tatagctgta tcggttgtaa tagcaaagga taagctgcat atcatcatac ctcgtttcgg
44101 ggcattctct gagtattgca agcaccttgc tgtcaaccgt cttttgctta ttcataatgc
44161 acctccgtaa aagaacggtg ggcgaagttt tcactccgcc cacataagcg ttccttattt
44221 ctttctgttc ttaatctgca agaagataaa gccgccgata ataaaggcga ctaatacaac
44281 tgctacgatt gttttagcgt ccattactta ttgtcctcct tcttcttttt tctgtacgca
44341 acaacggctg tgcctacaat accgagggca gaaagtcccg caagtgcagt ccacaagccg
44401 acattgctgt tatctccggt ctttggtgtg tctctcagct cattgtgcat ttccacaacg
44461 gtagtagagc cgacctttac ggttgcaatc ttatccgcag gaagcacata ggaagcagaa
44521 acagtatcct gaacctcgga aacggtgtaa tcgccgatac gaagtccctc gataatgatt
44581 tcgccgttct tatcggtcgt gaagcttcgg tcgtaaccgt ttactcctgt tactctgaaa
44641 gtgaaacctt cgaccttacc gtcagaggaa gtttttacga ttttcaggct tcctttcatt
44701 gccgtgttaa taaaaccgac gcccgctttg ttttcaacct cataggtcat ttcatcttcc
44761 tcaatgaata cggagtacac acccttatca agttcaaagc cttccggtgc tttggtttct
44821 cttacaaggt attttccgta aagaagctcc ttcatctcgt aaatacccgt agaagtttcg
44881 ttgagcgtac cgataagctc gtcgccgtcg tccaatttga cgtctccgtt tacatctttg
44941 taaacctcaa agattgcacc cgtcagtttg ttgtccgggt aatcttcatc gaccttagtc
45001 aacttgatat tgccgtgaac atattcattg acgatttcaa cctcaatgac ctgaccgacc
45061 tcggagattt taacttcata agaagtttca tcaagcacga agccctccgg ctgctcgatt
45121 tcttttacaa gccagtttcc atacggtact ttgtcaaaag aaaagctacc gtcattttca
45181 gaggtagcag tcatcaaagc attttctttt gtaaattcag tttcatcagc cttaaacaga
45241 ccgattacag cacctcccag tgcttcgccg ttttcggtaa tcttcttacc ggaaacagaa
45301 ccgtaaatga gcttgttctc aatcgctttt ccgtcgttga ctgcaatctc aacatttgcg
45361 gtatcctgac cggaatagct gaaagtgaac ggatacttgg cgtcggagag aatataatgc
45421 tcatcggtgg caagctcctt aacatagtag cttccgaaag gaatttcggt tttgataaca
45481 gccttaccgt tttcggaaag agagataatc tcaatcagac cgtctgccgg gatagaagta
45541 ccgcttgcag aaacaagttc ttcggcggca aaaagtccaa agctgatatt cttcatttca
45601 tcgccgttac cgatgttgaa gatgtcgttc ttttcaatag tcttttcaag acttaccgtt
45661 actttctgcc tgtcatttac aaaacttgtg gcggtttcgg ttacagacac ttcctgaccg
45721 gcataagtca gctcaacctc gtgagcttct ttgttgataa ccataccctc cggtgcggta
45781 atttcaacca ccgtatattt tccgagataa agctccttac tcttggcaag tccgttttca
45841 tctgtggtaa cagtatctac gacttcgccc tttgcataac ggagtgtacc gtcgggagtg
45901 ataatatctt ctgctgcggt aatctcatat accgcacctg caagaccttt cacttcatag
45961 acaggctgat aaatcacatc ggcattttcc tctccagaga tatttacccc tgagaatacc
46021 tcgccggtct tttcaatgct gacagtacct ttctgagcca tattaggctt atcgacctta
46081 atcacggtaa taccaccctc atcggaagaa tgttcctctg ccacatcaaa gtacaccggt
46141 gtgctgtcaa gcacatatcc gtaaggagcc tgaacctcaa caagagaata tcccttgccg

46201 tattccagtt tctccggtgt aacaagctgt ccgtttgcat cggtgtagaa agtatcaatc
46261 gttgtcggag taggataagt gaaggtcatt gttacctgat taccgtcagg gtcaaagatt
46321 ttgaagcctg caccggcata cggaatgttt ttaccgcttt ctgcgtccac cttaaccacc
46381 ttgatatagc tttcaaagtt tgcgttgttg atgagatagc ggtaggtctg tccgttctga
46441 gaaatgaaca catcgaagtc cttcatcagc tcacgacctt cccagcctgc ggtctgatga
46501 acggtatata cgccgtaagg catatccttt gtctgaccaa aaccgttttc atcacaaaca
46561 atcacatctc tttcatcttc ttctgctgaa ttaaagctgc cggaagattt cagatagatt
46621 tcaaaggttg ctccgttttc cggtgtttca atctttgttt cgccgtcatc ggtgtgcttg
46681 ataatggcga tattgccttt cataacctgt tcggtaacat catttgcggt ctgattgtgt
46741 tccaccgtat aaagctgcgg ttctgcaccg accttatgga ttgtaccgtc gagcagataa
46801 ccctcggacg gcgtgatttc acggattgtc cagtcattat cacagatata ttcctttgtg
46861 gtaaactgcc cgttcttatc cgtaacatat ttgtccacaa gggtttcgcc cttataaatg
46921 ccgtaaactg ctccggcaag ggtagcgtcg ccctgcggtg tgccttcctc tcggtcgctc
46981 tttgttacgg ttacgctgaa ctttttgaga atgttcgtga agtttcttgt ggttgcttct
47041 ttccagttaa tcggtgcggt ctgatttgca ggcacaacat aacggattgc ggtatccact
47101 tcctcaatgg tgtatggggt agtgccgctt ataagcacat catcgaaaga agcaacaccg
47161 ttcttatctg tcactgcata ttcatctacg gcaatacccg caagcgaagt tccgtaaagg
47221 tggaaagtta caccctcgac cagattatcc tccgaagatt tgataacctg aagattacct
47281 ctcttgagaa cattattgaa gttgaccttt gccacctgac cggcaacaac agtaactctg
47341 tgaacctcct gcggtacata tttgtcgata gactgttctg ttaccgtgta aacaccgggc
47401 ataaggttat ccagttggaa tttgccgccg tttgcggttg ttacagtctg attgacaccg
47461 ttaccgctga tggtaaactg aataccgtca acctttccgt cctcacttgt ctttacaatc
47521 tgacaagagc cgtagcttac cttcattttg acaaagccgc tgacagggtc gctgacctcc
47581 tgagcataag tgaccacatc ctgaatgccg ttgccattct gatggatacc gtcagaccat
47641 acgacaacac ctctgcggac accgttcttt tttgcggctg tgatggtaaa ttccttgctc
47701 ggagcttttt ccattgaaac agtcaatttg tttccgctga cagagaacga aacgccgtca
47761 atatttgccg agaaattgta attgccaagc acaccgtttg tatcggtgag agttgcaaca
47821 tacttgcttc cgttccattc aagctcgttc accttggcag aaccgcttga ccttgtgcag
47881 aagctcggaa ctttcgtatg gttctgaacg cttgttacca tactgttgta gtacgacaag
47941 attttgctgc ggagagggtg tgctgagctt acacattcaa gcacggcatt gtaccctccg
48001 gtagaaacgt gattgaagct tgcgtctctt tctccgacaa tcgtttccca aatcaaaatc
48061 tgagtggcat aagcgtgtgc gatacagttt gcgtcgcttt cattctgcga cttccagctt
48121 gtcgaaatac ctccacggta gccgtactgc aagatacgac cgataagcag gcgaatatca
48181 tcgccggaaa tcgtaccgtt cggactgatg ttgttgaaga agttctcgtc cttttcggtg
48241 agagtatctc ctgtccgctg acctgtaccc ggctcgatac agtaggcgat attgcccgaa
48301 tacgaaccca ttgcacgaag tcccgtatat ttcgtggaca agcctttcca accgttcata
48361 tagttgagat tgccgtgtcc ccattcgccg ttgttattgg tatctccgct tcggggataa
48421 tcgacgaggt acacatcggc tttttcgcct acggctgcaa aggctgtcgt tgccccaatg
48481 cccgtaaaag cagtaaggct cataaccgcc gccaacataa gcgagacggc tttcttgaat
48541 tttgttttaa ccattgcgtt ttcctccttg aaatgcaaaa acgcaccgat tttactcagt
48601 gcgtccgctt aaaattattt tgttttagca gtagccgata tagatgttgt aatccgtatc
48661 ggatagcttt tctgtccata tccacacagc ggtaaaaccc tcgctgttct tgtatcggtt
48721 cagtctgctt tcaatatcag acttgatacc gcttcggtta ggatttgcgg atatggggtt
48781 gtcccaacat tctgtcgccg tgatgtcaag agaaagccct atgcttaccg catactcttt

48841 tgcgtaacta acataagggc tgacatcgaa tgtctgcttg tgcggctctg tcggtttcgg
48901 ttctgccggc ttttcagtcg gtttttcaac cggcttttgt tcggcagttg tctgttcttt
48961 tggctgttcc gtttgttttt cgggttcagc ctgcgtggtt gtctgcttcg gtggctctgc
49021 ctttggcttt tcggttgcag gagtatcagc cgttttttct tccgccggtt ttgtctgtac
49081 cgtcataacc gtctgacttt cagcggttga ttggctttct tttgccgtaa tcggtgtttc
49141 ctcctgaact gccgtgcttt ccggttcttt tgccgtatct tcctgagaat ttgtttcggt
49201 attaccagcc ggaaaagaag tctccgtttt cgttatctcc tgtgcagctt ctttggctgt
49261 gttaccgtta caacctacga ggaaaatcat caggcaggca agaacacaag gtatcagcct
49321 gtttttcata tccatcgctc ctttctgctt taatcgtaat caaaaccggt caaaacatca
49381 aaggtgcgat tggctgaact ttttatcgta gcagacattt tgcttttgct ccaaatgtgc
49441 aaacccttta atagacagct atatgttatt gagatagata tactcaaggg agatagagta
49501 tattttcagc aggttaatag ccggtatata taggggttag agtgtgagaa agggggaagg
49561 ctatcggctg tcacggtcgg ctctgcttcg cagaaacttg ttgtaatgct ccaatgcctt
49621 gcacacaaaa tcctccgttt ggcttatggg aatattcttc gggataagct gtcgaaccct
49681 gtcgcccctt aaaacaattt tctccctttg gtttggtttt tcctccgaca taatcgcttg
49741 gattgcttcg gttgtgagct tgccctcctc aaagaacttt cgcatacgga ttgtttggtc
49801 gtgggacgga gttgcgtcgt tcaggtcgat ttcatcaacc acatcacgct gactgtcctc
49861 atcgagaaaa gacaattcaa ccgccggtcg cattttaatt cgcccctcgt caacataatc
49921 caaaagttcg ggaacaaggt ttgtaaggcg aatataacgc tgaatttgcg ttttgctgtc
49981 atccgattta tctgctaaca acattactga ttttgttccc accaacttgt cccccagtgg
50041 gtcacaagtt aaatccgtgc gttctccttg ccttttcatc gcttccaaac gcattttgta
50101 ggcataggct ttttcagagg gaagtatctg cgacctttgg tagttgctct caaccattaa
50161 aatcgtggct tcgtctcggt tcaggtcaac gacttcacaa cggagcgtat caaacccggc
50221 aagctcacag gctcgctttc gtctgtgtcc tgaaatgagt tcgtatctgc cgtcctcttt
50281 ctgccttact gtcgccggtg tgattacccc tcgttccttg atactctcaa tgagctgtgc
50341 catatcctcg tcatcacgca ccttaaaagg gtggtcgggg aaatcgtcaa tctcagtcaa
50401 gggaatatct cggattttgg aaagcttttc ttcatctctc tgttcctgcg ttgagaataa
50461 gtcatcgagc ttcgtgagag tgaagtcgct ctttcttcct gccatcggct aacacctcct
50521 ttgtaaattc tgtgtacgcc tttgacaccg tgctgcccgg ttcgtaggca aaaatgctct
50581 tgcctttgga agaagtctct gcggctttta cagcaatcgg gatatatgtt cgatacatct
50641 tgatttgact accgaagttc tctctcagag cttctaccgt gcttttggcg aggttggtac
50701 ggctatccac caaagtgaga agcataccgt caatcttcaa gctcggattg atacgtttct
50761 ttactcgtga aatagtcccc acgagctgcg tcattccttt tgccggtaaa tactgagcct
50821 gaacagggat aataacgctg tctgccgcag atagagcatt gattgttacc attccgaggg
50881 aaggcataca gtctataagg atatagtcgt atttgtcctt cacttcgctg agatagtttc
50941 tcaatgccgt ctctctgctc attgcgttga caaggttaaa ctccattgct gaaagctcaa
51001 ggtttgccgg aataaggtca acaccttcgt cgtggtgcag aataccgacc gtagggtcat
51061 tcatcgtttc atttatgaca tctgtgagtt tcgttgcaag cgtgataccc aagttgtctg
51121 tatcctgcca accgagacaa gtggttaagt cgccctgtgg gtctgcgtct atgagcagca
51181 ctttctttcc ctgcattgca agaccgacgc cgaggtttac cgtagtggtc gtctttccaa
51241 ctccgccttt ttgattgcat atcgcaatcg ttttgcagtt cgacactttg tgcgtcctcc
51301 tttcgtgaaa ttcatagcca tcgcagtatg gctatgtact tgaccggctc atttcctaac
51361 cggtcgggca atatttgttc tcaacacccc ttgccgagca ataagctcac gggaagtcac
51421 taaggaacag actgctaatc tgtatcatag gaatttcacc tctgccgggt actccgccag

51481 ctccgtagag aagtatcatt atccttctga ctgtcatcgc cgtaacaggg gcaacctgtt
51541 acctataccg ggagttctcg ctgttctgca atgcagaagt cacggcgtac ccgataaggt
51601 ggctaaaatc atcagaatta aagtcttagt gaatggttta ttcagttgtc aaaatacaag
51661 tgaaagggct gaccttttcg tggctcggaa ttaacccctt cacttgtttg aaatgggaga
51721 gcgtttttgg ggggtgtttt ttcaaaattt cttgaaaatt tttttgctaa caaaaaaagc
51781 cacctacttt cctcttaatt gaggtcaata gacggcgggt agaaatattc ataattttgg
51841 tgtataatag gaaattaaga gaaagacaaa tttgaagttg gaggatacac gcatgaaaat
51901 tgtaatcatt aatggaagtg ccagaaaagg aaatacgctg gcagcaatca atgcatttat
51961 aaaaggagcg tcagaaaaga ataaaattga aatcattgaa cccgacaaac tcaacattgc
52021 accatgcaag ggatgtggtg tctgtcaatg ctctaaggga tgcgtcgata aggatgatac
52081 aaattctacc attgataaaa ttgctgccgc agatatgatt ctttttgcta caccagttta
52141 ttggtgggga atgtcagcac aattgaaact tatcattgat aagtgctact gccgtggatt
52201 gcagttaaaa aacaaaaaag ttggcacaat tgttgtaggt ggttctcccg taggcagcat
52261 ccagtatgag ctgattgata agcagtttga ctgtatggca aaatatcttt catgggatat
52321 gcttttcaaa aaatcatatt atgcaacagc cagagatgaa cttgaaaaaa ataaggattc
52381 catgaacgaa cttgagtgga ttggaaaaaa tttataaagc acacaccaaa ttccagtctt
52441 gcgaagagac tgaaaacaac agaagatttc agtttgtcta acttactcta aattagcaaa
52501 gaccgcttac aaaagtgtag gcggtcttta ttcgtgttca gatatatttg attttgaact
52561 ttcttttctt ggctgaaagc accttgcaaa gaatatcatc tgctgcgtcc gtgtatctgc
52621 cttgtgctat taggctgttc ttcaaacgga taagtgattg cagcacttcg ctgtattcgt
52681 ccttggaaag gtataaatga tatttttgtt ctttcaaaac aaccacctcc tgttaccgca
52741 attatataat aagaaatggc atagaagtag tgtgcgattg tcgcttgacc acccctaaat
52801 gaacgaatta acagggtaac gaacgcctta ttcattagca ggaattttgc tcgaattttt
52861 ctgccattag cagagagggc attttcatta gcaaaatagc gaaaatcctg ctaattttta
52921 ttagcaagcc gtccacctga aaggcggacg acattagtcc taatttccat tagctgacgg
52981 agaatttgct aatgaaaatc tgtgaaattt acgatttttc tcttttcccg taaaaatgaa
53041 aaaacgccgg aaacccttga ttttactggg ttttccggcg gagtttttgg cgttcccgag
53101 gtgattcgaa cacccgacct accgcttagg agttctctgc gaaccgtggt agctagaagt
53161 tcgattatcc tcttgggagc gtttaatgcc gtaaattaaa gcataagtgc ttgttatgtt
53221 taattgttga tactctctgg tgtaccctca tacaccgtag agtgttagca ggttgttagc
53281 aacaaaggct aatttaaagc tttaatttat cagaaaacat tacaaacaaa ctataaatca
53341 tttatcacta gaaatcaaga tactcgtatg ttatagacgt tcagaaatga gcgtctattt
53401 ttataccaaa aatgaaagga gtattaaaat ttatgaaaaa gacattgaaa cgattgtgta
53461 cgggcttctt agctcttgca actatcgtta ctgctttacc gactacaccc gttcatgcag
53521 aaagtaagca atactggaca gaaagtgcag agcgtgtcgg tatcattgaa aaagtaatga
53581 atgacggttc tatcggctcg acattcaatg agggcatgat gaaagttgag ggcaaaacag
53641 cctattgtat cgacattaac acagatttta ggaatggcta caagaccaga gctgacgcaa
53701 gctcacgcat gaatgccaac cagatttcag atgtggcgtt atccttagag tatgtcaaac
53761 agtatggcga agctcacaag gaactgaact acaagcaagt ctatctgttg gaacaatgtg
53821 tagtctggca gagattgagc gtacatcttg gctggcaatg cgataacgtg cgagcttctt
53881 atgatgaaat tccaaaggca acgcaggacg aagttttctc tggtgcaaga acctttgtca
53941 aagaaaataa aggacgctat gaatgtggcg gttatatcta ctctggcgag gggcaggaat
54001 tggggcaatt ctgggcgaag ctgaatgttg gaaatgcgaa acttaaaaag acttccagta
54061 atgccagcat tacagacggt aacgggaatt actcaatcgc tggtgcgacc tatggtgtct

54121 ttactgataa ggactgcacg aaacagcttg ccacccttac gactgatgaa aacggaaata
54181 cagatgttgc agaggtaaaa gcaggcacag tctatatcaa ggaattatcc gcaccagcag
54241 gatataaagt agataaaact gtatattcct taaagattga agctggaaag acggcgactt
54301 tgaacgtatc tgatacgcca aaagtaacgg acactttgat tgagcttttc aagatagata
54361 tggaaacaaa gaaagacaat ccgcaaggga acgcttcttt agcaggtgcg gaatttacat
54421 ggaagtatta tgctggcttc tataacaaag acaatctccc tgccgaagct actcgaacat
54481 gggttacaaa gacaatcgct gaaacagaca gctacgggac aactcattac atcacaaaat
54541 tagcggacgc atacaaggtg tctggcgaca gcttctatat gcaagacgga aaagccgttc
54601 ttccacttgg gacactaacc atagaagaaa caaaagctcc aaacggctac ttgttagagg
54661 gtgcatatat gcaggctggc gataagtccg aacagataaa gggcttatat gtaacacaga
54721 ttaccgagga cgacgacctt gccgtattat ctggaagtaa ccagttttcc gtatcagaca
54781 aggttatccg tggcggtgtg aaaattcaga agcgagattt agaaacgggc gatatgaagc
54841 cacaaggaag tgcaaccttg aaagatactg cttttgacat catttcctta aatgataatt
54901 ctgttttggt tgagggcaag ctatacaaga aaaatgaagt ggtaaagaca attcgtaccg
54961 atattgaggg tgtcgcttct acttctgctg acctcttacc ttatggaaaa ttccgtattg
55021 ttgagagtga agctcccgac ggttacttaa ctgacggtgc aaagccgatt gattttacaa
55081 tcacagaaaa tggaaaaatc gtggacttaa ccgacaaagc ccattctatc tataatcaga
55141 ttaagcgtgg agatattgag ggtgtaaaaa tcggtgcagg cacacacaag cgtcttgctg
55201 atgttccctt taggatcaca agcaagacga caggagaaag tcatatcgtg gtaactgaca
55261 ataacgggca gttctccact tctgctgact gggcttctca taagcacaat accaacgcag
55321 gcaagaccag cgaggacggt gtgtggtttg gaacttctga accagacgac agcaaaggtg
55381 cgttacctta cgatacttac gtcattgagg aattacgctc tgaaagcaac aaaggctttg
55441 aacttatccc accttttgaa atcgtggtat caagaaataa tcttttgatt gatttaggaa
55501 cgctgactga tgaatacgaa aaagaaatct ctatccatac cacagcgacc agcaaggacg
55561 gcgaaaagac tatccttgca ggaaaagaga ttacgattgt tgatactgtc aaattagacg
55621 gacttacaaa aggcacaaaa tatcagctca aaggctggca gatgttaaaa gaggaaaacg
55681 ccgagcttat cattgacggg aaacgtgcag aaaatgatta tacctttgtc gctgatgagg
55741 aagaaatgaa agtggaaatt gcctatacat tcaatgcgtc tgctttaggt ggcaagaacc
55801 ttgttacctt tgaagaattg tatgatttga gcaatccaga cgaacccgtg aaagttgcgg
55861 aacacaaaga cattgaggac gacgggcaga ccgttttaat cacagagcgt atcatcaaaa
55921 ttcatacgac tgctacgaat aaggacggca acaaagagct tgaagctgga aaaaacgtta
55981 caatcattga taccgtaacc ttagagggct tagaagttgg tacacagtac aaacttgtgg
56041 gctggcagat gttgaaagaa gaaaatgcag aacttcttat caatggaaaa tgtgtggaaa
56101 atgattatac gtttactgct gacagcgaaa ctatgaaagt ggaagttgct tttacgtttg
56161 acgctacttc tcttgacggt aaacagcttg taacttttga agaattgtac gatttaagca
56221 atccagacga gccgaagaaa gttaccgagc ataaggatat tgaggataag ggacagacga
56281 ttacctttaa ggaaaagcca gaagaaccag agaaacccga aacaccacag ataccagaaa
56341 agccgagcag accgagcgac agtcccaaaa cgggcgacag tacaaatgta atggcatttc
56401 ttgtgatgtt gcttgcgtct gctggtggac tggctggaac atatctttac aaacgccgta
56461 aaatgaagaa atcataaggc ggtaacggta tgatgaaaga aaaatcacgc tctccgccga
56521 agctgtcaaa cggcagactt ttgcttattc tggcttgtct gccaccaacc tgcaagaaca
56581 cggatagtca agggtgcgaa gcattgattt taccctttac tatctggggg attgctggta
56641 cttcccaaac cgccagaata agaaattaca atcaatttat caaaatagaa agaaacgagg
56701 taaaacatta tggaaatgaa atatgtcgtg ccagatatgg cacagtcttt tggaactctt

56761 gaatttgcag gagaaagcga gcctatcttt gaaagagata aaaataaccg cagggtctta
56821 gccagacgaa gctataacct ttattctgac gtacagaaag gcgaaaatgt tgtagtggaa
56881 attcccgtgc aggctggcga aaagaagttc aagtatgaac agaaagtaaa acttgtcaat
56941 ccgaagttgt acggcagagg ttacgcaatc ggggatatgg gacataccga ttatgtatta
57001 gttgctgatg atattgtagc agttgaagaa aagtaaagga gtgaagaatt tatgagatta
57061 tcaaacgggt ttgttattga caaagagaaa acatttggcg aattaaaatt tacagcggtg
57121 cgtgatgtgt ttatgcagaa tgaggacggg acaccgagta cccagcttaa aaagcgtatc
57181 tatgacttga aatgtagtct gcatggtgga attatccccg tgtccgtacc gccagaagtg
57241 ccgttaaggg aatttccgta caatgcagtt gtagagcttg tcaatcccgt agccgatacg
57301 gtatcaagaa agacgtttgg tggtgcagat gtggactggt atgtaaaggc agaggatatt
57361 gtgttgaaga ataaaggcaa tcagaacgca ggcaatccac agaaccatgt accgcaggga
57421 cagacgaaaa aataaatcta ggtcatagat tatactctga aattacagta taatgagcca
57481 aaagagccgt atttaagcta tgtaaaataa ttttgacaag gaaaaaaagg atttgtcgaa
57541 gatattggat agacatttga gggttaatat atgacaatga caccgaaagg agcaatcgga
57601 atatggaaat aggtaagaaa cttaaaaatg cacgagtaca gtctggaatg acacaagaaa
57661 atgttgcaga aaaaatcaat gtatcaaggc agacaatttc aaactgggaa aatgaaaaat
57721 cttacccaga cattatcaat gtaatagagt tgagcaacct atattctata agtcttgacg
57781 aattgctaaa gggtgatgaa aaaatgattg aacatttaga agaaagtaca aatgttgtaa
57841 aaagcaacca aaaaatgata tgggcaatta ttgtaaatat cattgtcgta gcattgctga
57901 tagcattaaa tatgtttata ccagataacc gttatttctt agttggaata ttttgtctta
57961 tggttatcac ttcatcagca ttactttacc agataatcaa aaaattttaa ggaggatatt
58021 ttgttatgaa tttgaacata gttttagcgg taatttgtgg agcagttgca cttgtaggag
58081 cgttttgtgt tgtgtttcag atttatcata tgactgttat agacgcaaca gccagaggat
58141 taaagcaccc taaattttgg ggagtattta caatgagtgg gaataattca agtggtttgc
58201 ttatgtatct aatcggaaga agaaaatacc ctattgtaaa tatgagtgaa agcaactcaa
58261 aagaattaga aaaacgtaaa aaatcagcag gtattggact gttatttttg gctattggtg
58321 taattggcat tatatgtgcg accttaatat aactaaaaat gcagatttat gattatatct
58381 tgcaacattt acaaagcagt tagtctatat gattgactgc ttttttcgta cccagaaagg
58441 agtgaatgat ttatgcgtaa gatttggaat agaggacacc gtatcagagc cagcgacaag
58501 caccttgtct accatttttc catagggacg cttctgtttt tattcgtggc ggttctcctg
58561 ctactgaata tcaaacagct catgcgtacc gactgggagc atttcagctt gttagagaac
58621 ggcttgacgc tttcccctta caacttcata accatactga tagcgactgg tgtttgtgca
58681 ttggtcgctt ttctgtatta ccgcttctgt tacgacagtt tcaagaagct cctgcaccgt
58741 caaaagctgg cgagaatgat actggaaaat aagtggtatg aagccgatac cgtacaagac
58801 agcggttttt ttactgacct gcaaagcaga tcaagggaaa aaatcgtctg gtttccaaag
58861 atttattatc aaatggaaaa gggactgctt catatccgct gtgaaattac gctgggaaaa
58921 tatcaagacc agcttttacg gttagaggat aaattggaaa gtggattata ttgtgagctg
58981 accgacaaga ccttacatga cggttatatc gaatataccc tgctttatga tatgatagcg
59041 aaccgcatta ctattgatga agtacgggca gaaaatggct gtcttaaatt gatgaaaaat
59101 cttgtctggg aatatgacgc actccctcac gctctgattg ctggtgggac tggtggcggt
59161 aaaacctatt ttctgctgac gctcattgaa gccttactgc ataccaacgc tgtcctttac
59221 atcttagacc cgaaaaatgc ggacttagca gacttaggga cagttatgga aaatgtgtat
59281 cacaccaaag aaaaaatgat tgattgcgtc aatgcctttt atgagggtat ggtacagcga
59341 agtgaggaaa tgaagcgaca cccgaactat aagacgggcg aaaactacgc ttatctggga

59401 ctgccaccct gctttcttat ctttgatgaa tatgtggcgt tttttgaaat gctggggaca
59461 aaagaaagcg tgggcttact tagccagtta aagaaaatcg ttatgttagg acggcaagca
59521 ggatatttcc ttatcgttgc ctgccagcgt ccagacgcaa agtatttctc ggacggtatc
59581 agagataact tcaatttccg cgtgggactt gggcgtatca gcgaattagg ttacggtatg
59641 ctgtttggtt cagatgtgaa aaagcagttt ttccagaagc gtatcaaggg gcgtggctat
59701 tgtgatgtgg gaacaagcgt tatcagcgaa ttttacacgc ctttagtccc gaaaggacat
59761 gactttttgc agactatcgg ctctcttgca caagcaaggc aggacgggac ggcgacgtgc
59821 gaagcgaaag gcgacggcac ggactagccg ttgctggtgt ggcgtaagcc acgccagcag
59881 gtaaaacccc tcggtatcta acagaggggt acgtttgcaa atagacaata gacgaaaaac
59941 ataggaaaca taaggcttct ggcatgattt cccagtaaaa atcggctgtg aagtcgcctt
60001 aaaaaacttc acacttgaca gattgggggt atggttctga atgaagaaca atggataaaa
60061 gaattacggg agaaacggat tgcttacggt atctcacaag gcaggctggc ggtggcttct
60121 ggtatcacaa gggaatatct caacaagata gaaagcggaa aaatgaagcc gtcaaaggaa
60181 cttctggaaa ctctgcataa agaactggca aggttcaatc cagaagcacc gcttaccatg
60241 ctgtttgatt atgtgaaaat tcgttttccc acgctggata tacagcacat catcaaagat
60301 atattaaaac tgaatatcaa ctatatgctc catgaagatt acggacatta cagttatacg
60361 gagcattatt ctttagggga catctttatc tatacgtcgg ctgacgaaga aaaaggtgtc
60421 cttttagagt taaagggggcg tggttgccga cagtttgaga gttacctgct ggcacagcaa
60481 agaagctggt atgactttct catggacgca ctcatagacg gtggcgtgat gaagcgtatc
60541 gaccttgcta tcaacgacca tacgggcatt ttggatattc cagagcttgc ggaaaaatgc
60601 agaaaacggg aatatatcgg aaagtccaga agttataagt tttaccagtc gggcgagctt
60661 atcaagcaca gcgaggacga cagagaatat atgggacgta ccctttatct tggttcgctg
60721 aaatctgatg tatatttctg tatctatgaa aaggactatg agcagtatgt caagttaggg
60781 acacctctgg aagaagccga tattatcaac cgttttgaga tacggcttag aaatgaacgt
60841 gcctattatg cagtacggga tttgctgacg tattacgacg cagagcagac cgccttttct
60901 atcatcaatc agtatgtacg gtttgttgat gaagaaccag acaagcggaa aaatgactgg
60961 aaactcaatg accgctgggc ttggtttatc ggcgataatc ggcaaagtct gaaactgacg
61021 acaaagccag aaccttatac cttagatcgt acattgcgtt gggtacaaag gcaggtagca
61081 ccgaccttga aaatgctgaa aaagattgat aagggaaacg gtacagatta catggagaca
61141 atcgaacagc aggcaaagct cacagaaaag catgaaatga taatcaaaca gcagacgacc
61201 cctgcaaaag atttagtaga aatttaggag tgatgaaaaa tgtgggtatt attaaaagac
61261 ttcctgctgg tatctatggg aatgggtatc ggcatagtct tgatgtgtat tttgaatgtc
61321 ggcaaagaag ctgactataa aatgaaacaa ttagaagaaa gcgaggacaa ttaaatgaat
61381 ttcggacaaa atttgtatca atggttttta agtaacgcac agagcttagt gcttatggcg
61441 attgtggtta tcggtatcta cttagggttc aagcgtgagt tttccaaact tatcggcttc
61501 ttggtagttg ccttgattgc tgtcggtttg gtattcaatg ctggcggtgt gaaagatgta
61561 ttgttagagc tgttcaataa gattatcggt gcataaaatt ttattgtggt actgatatgc
61621 gaatgttata atttgaatgt tgtattacat aaaacgctgt acttaccaga aaaatctatg
61681 aattttgtat attatacaag gttgcaactt gaagttgcta atgtgcattt caaaaaatat
61741 agaatataac attatggctt gataaaattt attcatcaaa tcataggagg tatcacacaa
61801 tgagtttagg agaaaaaatc ttcaagttaa gaaaagaaaa aggaatatcg caagaagcac
61861 tagctgaaca aataggcact actaggcagg caattagtaa atgggaaaac aatcaagggt
61921 ttccagaaac agaaaagctt ttacaattat caaacatttt tgaagtatct acggattatc
61981 ttttgaaaga cgaaaagaat gtaaagaata gtgaagaacg aggatattat gtaagcaagg

62041 aaatgtcggt agggtttatc gctaatcaga aaaaattagg actttatctt gctttagggg
62101 ttatgctttg catttttgct ggaataccat ataccttgtt tttagataat atgacttggc
62161 gttatttagg tatggcattt tgctttgtgg caggaattat attttttgtc gtagcaattt
62221 tttatgaaaa acaagaatac aacattttaa ggcaagaacc tttactactt gattatgaat
62281 atctaaaaga attgaaaagc gaatacactt caaaaaagaa aaaatatgta atagttgctg
62341 ttccttgtac tgttctcttt atagcaggac taattttctt tctatttact gtgaaaggct
62401 atatttcatg gacagaatat cattcctttg catttttcgg tcttgctgtt ggaatgttgg
62461 gttttgtata tacaatcaat gttatggaaa cctatgaact attagtgaaa aacgagcaat
62521 attcaacgag tttctttttc aaactaaaaa gaaagattaa aaagagatta ggataagcta
62581 taatttaata ttcacacaaa gcagttagtc ttatgattga ctgctttttt cataccaaaa
62641 tttcagattg gagtgattac atgaacgata tttttaagga tatgcaggca aaagtcggct
62701 gtgaatacat ttctgacctg ccctcttaca agcgtaaggt gtggcgtgaa atgaaacggc
62761 tgagccctgc cgattatgaa gaaagacaat tagaagattt ttcaaagttt gtgtttggta
62821 tgccgtacca gaccttaaaa gatgtgatga aacaacagaa aggacgtgaa gaacaatgca
62881 ggaaacaagg gtgctggtgg aaacgagaga aacaactggc gaagaaacaa catcatactg
62941 gttcgacctg ccgatagatg ttgccgagtt tgaagaaaag ttaggtgtcg gtgcagaaag
63001 cacggcgtac cgcattatcg aaaaggtgtt gccttatgct gatgaagtcc acgaacatac
63061 aagcgtgtac cagcttaacg aattagagtt tatgtaccgc cagctttcaa gcgatatgca
63121 ggaagaatat gtatcactac ttacggtgtt tgagaattta gaagcacttt atatttgcag
63181 gaacgtgatt acggtttatc ccgactgcaa aagcatgata gatgttgcaa ggcaaaagct
63241 gatgaatgac tcgacgttca aacatttatc cgaggactgt caagaatact actttgactt
63301 tgaagcctac gcttctcact tgcaggaaca cgggaaattc ttagtaacgg aacacggtat
63361 ctttgaacta ccagagtagg aaatgaggtg gtgcttatga tagatgatat ggcggtttac
63421 attgctaatc tcggtaaata caatgagggt tatttagtcg gtgcttggtt cacgttcccc
63481 attgacgagg aagatgtaaa agaaaaaatc ggcttgaatg aacagtatga ggaatacgct
63541 atccatgata ccgacaactt ccccattgcg attggcgagt atgtttccat tgaagaactc
63601 aatgagatgt atgaaatgat agaggaactt cccgactata tcgtagagtg tctggacgaa
63661 tttatcagtc actacgggac gctggaagaa gtcgtggaac acaaggacga tatttattat
63721 tatcccgact gtgaaaccat gacagatgtt gcctattact acatagacga attgcaggca
63781 cttggggaca ttccacccag cttacagaac tacattgatt atgaagccta cgggcgagat
63841 ttggatatgg gcggttgctt tattgaaaca agccgaggga tatgcgagat accatattaa
63901 cgctggaaga tcagcaacgg catggttgct actgacagcg tatttctctt ttaagggaca
63961 gtctgaaaat ggctgtcctt ttctctttga agaacttacg aaaggagctg aaagaacttg
64021 aaaaagatta aatcttatac gggtatctgg aacgtggaaa aagtcttgta tgcaatcaat
64081 gactttaacc tgccttttcc cgttaccttt acacagatta catggtttgt gattacggaa
64141 tttatcatca ttctgtttgg ggatattccc ccactttcca tgattgaggg agcatttctc
64201 aaatacttcg gtattcccgt tgctctcact tggtttatgt cgcagaaaac ctttgacgga
64261 aagaagccgt acagcttttt gaaatcacag ataacctatg ccctgcgacc taaaatcact
64321 tatgcgggaa aagccgtaaa actgcataag cagaccttga atgaaacaat cacggcagta
64381 aggagtgtga actatgttcc cgataaaata tattgacaat aaccttgtct ggaacaagga
64441 caatgaggtg ttcgcttact atgagctgat accgtacaac tattcttttc tatccgcaga
64501 gcagaaattt atcgtgcatg acagtttccg acagcttatc gcacagtccc gtgagggtaa
64561 aattcatgcg ttgcagattg ctacggaaag ctcaatccat agtatgcagg agcagtcaaa
64621 gaagctggta actggaaaat taaaggaagt tgcctgccag aaaatagacg aacagaccga

64681 agcgttagta tctatgattg gtgacaatca agtggactac cgcttttttc ttggctttaa
64741 gctcatggtt acggaagaac agctcaatct gaagaacata aaaaaatcgg cgtggctgac
64801 gttcacggaa tttctccatg aagtgaacca cacgctgatg aatgattttg tttccatgcc
64861 gaatgatgaa atcaaccgtt acatgaaaat ggaaaagtta ctggaaaata aaatctcccg
64921 tcgcttcaag gtgcgtcgct tggaaatcaa tgattttggg tatctcatgg aacatcttta
64981 cggcaggaac ggtatcgcct atgaagatta tgagtatcag ttaccaaaga agaaattgaa
65041 caaagaaacg ctgataaaat actacgacct tatccgtccg acaaggtgtg tgattgagga
65101 aagccagcgg tatttacgat tggaacatga ggacaaggaa agctatgtgt cctactttac
65161 cgtcaatgcg attgtcgggg agcttgattt tccgtcaagt gaaatcttct atttccaaca
65221 acagcaattc acattccccg ttgatacttc tatgaatgta gaaatcgtgg agaatcgaaa
65281 ggctcttacc acagtaagaa ataagaaaaa ggaattgaaa gatttggata atcacgccta
65341 tcaagcagga agtgaaacca gttcaaatgt ggtggacgca ttagacagcg tggatgagct
65401 ggaaacggac ttagaccaga caaaagaaag tatgtataag ctctcttatg tagtgcgtgt
65461 atcggctgat gatttggacg aattaaaacg ccgttgtgat gaagtcaagg acttttacga
65521 cgacctcaat gtaaagctgg tgcgtcctgc tggggatatg ctggggcttc attctgaatt
65581 tttacctgcc agcaagcgat atatcaatga ttatgtgcag tatgtaaaat cagatttttt
65641 agctgggctt ggctttggtg cgacacagca gttaggagaa actacgggta tctatatggg
65701 ctattccgtt gatacgggaa gaaatgtgta cctgcaaccg tcccttgcca gtcaaggtat
65761 aaaaggtaca gttaccaacg cccttgcgtc tgcttttgtc ggttcgcttg gcggtggaaa
65821 gtcgttctgt aacaatcttc tggtgtatta ttcggttctc tttgggggac aggcggttat
65881 cttagaccca aaatcagagc gtggcaactg gaaggaaaca cttccagaaa tcgcacatga
65941 aatcaatatc gtaaacttaa ccagcgataa aggaaatgca ggacttcttg acccattcgt
66001 gattatgaag aatgtaaaag acgctgaaag tctggcaatc gacatcttaa cattccttac
66061 gggtatttcc tctagggacg gcgaaaaatt ccccgtactt cgtaaagcgg tacgctccgt
66121 tacccagagc aacagtcggg gcttgctcca tgtgatagac gagctacgcc gtgaagatac
66181 gcccatatca agaaatatcg cagaccatat cgacagcttc acggactacg actttgcaca
66241 tctgctgttt tcagacggga cggtagaaaa tgctatcagt ctggataacc agctcaacat
66301 cattcaagta gctgaccttg tactgccaga taaagatacc acttttgagg aatacacgac
66361 cattgaatta ttatcggttt ctatgctgat tgtgattagt acctttgccc ttgattttat
66421 ccattcggac agaagtattt ttaagattgt agaccttgac gaagcgtggg cgttcttaaa
66481 tgtggcacaa ggagaaacgc tctctaacaa gctggttcgt gctggacgag ctatgcaggc
66541 aggcgtttat tttgttaccc agtctgccta tgatgtttca aaggaaagtc tgaaaaacaa
66601 tatcggcttg aagtttgcct ttcgttctac tgacattaac gagataaaac agaccttaga
66661 gttttttggt atcgacaagg acgacgagaa caaccagaaa cgcttgcgtg atttagagaa
66721 cggacaatgc ttattgcagg acttatacgg acgtgtcggt gtggtgcaga tacaccccgt
66781 ctttgaagaa ctgctacacg cctttgatac cagaccgccc gtacagagaa atgaggtgga
66841 gtgatgaaag aaaggataaa aggtgcgttc acaaaaaaga agattttcca ctttctcaaa
66901 atggttctgt tcgtggtggc actctccctt atcctgcttt cacttctggg gacggtggct
66961 catgcgacgg ggcttgtgga cgataccata aatgcagaaa atctttactc aaaatatccg
67021 ctgtcgaatt accagcttga tttttatgtg gataatagct ggtcgtggct tccgtggaac
67081 tggctggacg gtatcggaaa atcggtgcag tatgggcttt actgcattac caactttgtc
67141 tggacgataa gcctttattt aagcaatgcc acgggctatg tggtgcagga agcctataaa
67201 ctggacttca ttaacgatat ggcagacagt atcggaaaaa gcatacagac cattgcaggc
67261 gtaacacaga acggtttttc ttctacgggc ttctatgttg gtttcctgct tctcattatc

67321 ttagtggtgg gactttatgt tgcctatacg ggactgataa aacgggaaac cagcaaggca
67381 cttcatgccg ttattaactt tgtggtggtg ttcgtgctgt ccgcttcgtt tattgcctac
67441 gctcccgact acatcaaaaa gataaatgaa ttttcaagtg atattagtac cgcttcgttg
67501 gacttgggaa caaaaatcat gctcccaaac tctgacagca agggcaagga cagcgtggac
67561 ttgataaggg acagcctatt ttctattcaa gtggaacagc cgtggttact cttacaattc
67621 gggaacagca acgcagaaga aatcggggca gaccgtgtag aagctctggt atcggcaagc
67681 ccagaggacg aggacggaaa gaccagagag gaagtcgtga aaacagaaat cgaggacaac
67741 gacaacaaca atctgacgat accgcaggtc gtgaaccgtt taggtatggt gttcttccta
67801 ctgttcttca acttagggat aacgatattt gtattcttgc ttacgggcat gatgttattc
67861 agccagatac tttttattat ctttgctatg tttctgccta tcagcttttt actttccatg
67921 ataccgagct atgaaagcat ggcaaagcag gcaatcgtga gggtgtttaa taccataatg
67981 acacgagcag gaataacgct cattgtaacg gtggcgttca gcatttccag tatgttttat
68041 aacatatcca cggactatcc gtttttcatg gtggcgttct tacagatagt gtgtttcgct
68101 ggtatctaca tgaagctggg tgacttaatg agtatgttct ctttgaacgc taatgatagt
68161 caaagcatgg gacgaagaat tttccgcaga ccatatctat acttggcaca tagggctaga
68221 cgtatggaaa gacggcttgc tggtgcggtt actgctggtg gtgttgctgg tgcggtggct
68281 ggtagcacga tttcaaacag aaaagcgaca aggaacacag ctacaaaaga tagtcggggt
68341 aatacatctt ccagcatggg acagcgtgta ggttcaaagg tgggtgctgt cttagataca
68401 aaaaataaag tgaaagataa ggcaagtgct gtcaaagaga atatcaagga tatgccgaca
68461 cagaccgctt atgccgtcca ctccgcaaag gaaaaggcaa agtccagcgt gtccgacttc
68521 aagcgtggca tggtgcagga acagcagtcc agacagtcgg gacgtatgga aaaacgggaa
68581 cagcataggc agaatatcgc cgataagcgt atggagcttc aaaaggcaca agaagcaagg
68641 caggcacagc gaaaggctga cggatcagcg acaacgggag ctacccgtcc ccatgagcga
68701 ccagccacag cttcaaccat tccaaagccg agtgctgaaa aaacgcagga agtcaaacgc
68761 cctgccacag cgacgacttc aaaaacaaat gagccagtca agacaacggt tatcaaagag
68821 cgtcccttat cttcgggtac ttctgatagg aaaacgagcc agcctacaca gccagtacat
68881 aggcagaatg tagaaaaagt ggtatcgcag gaaacacgcc agaatgacac caaagaccgc
68941 aggacaaaag ttcagcagac gcaaagcgtc cagaaaaacc agcagaccac agagaaaacc
69001 cgtaaccttg taatgaagaa aggacagaag aaaaaatgaa actgaaacat attgctatta
69061 tcggcagtct gtttcctatc ctctttcac tggtgctttt ctttggggtg ttgattagtg
69121 cggacagcga cgacgagaac agcaactttt cttctggcat tacgggtatg aacttatccg
69181 cagaagtctt gaaacatcag cctatggtgg aaaagtacgc cagagaatac ggcatttccg
69241 agtatgtcaa tgtgttgctg gcaatcatac aagtggaaag tggcggtaca gccgaggacg
69301 ttatgcaaag ctctgaaagt ctgggactac cgcccaattc attagataca gaaagttcaa
69361 tcaagcaggg gtgtaaatat tttgcgtccc tgctttcgtc ctgcaaaaat caaggtatgg
69421 acgatttgaa tgtagcgata cagtcttata actatggcgg tggttatgtg ggatatgtgg
69481 caggaaaagg aaataaacac actttcaacc ttgcagagag ctttgcccgt gaaaaatcgg
69541 gtggaaagaa agtaacatac accaatccaa tagctgttgc gaagaatgga ggctggcggt
69601 atagctatgg aaatatgttc tatgtggaat tagtcaatca gtatttaact gttccgcagg
69661 tgtcgggaga gctggcacaa aaggtaatga atgaagcgtt gaagtatcaa ggctggaagt
69721 acgtctatgg tggtagtaat cccaacactt cctttgactg ttcgggactg acacaatggt
69781 gttatggaaa agctggtatc tccttaccga gaacggcaca agcacagtat gatgctaccc
69841 aacatctccc actctcgcag gcaaaggctg gggacttggt atttttccat tctacctata
69901 acgctggttc gtatgtaacc cacgtcggca tttatgtagg aaataaccag atgtaccatg

69961 caggcgaccc gataggattt gcagacctaa gcagtagtta ctggcaacag cacttaatcg
70021 gtgcaggacg agtaaaacaa tagaaaggac ttgaagatat gtttaagaaa aataagaaac
70081 agacagaaac tatcaaagaa ccaaaagaaa aaaaggtgcg tactgtcaag gtaggcacac
70141 ataagaaaac tgtgattgcc ttgtgggtgg tgcttatcgc aagcgtgagc ttcggggtgt
70201 ataagaattt taccgctatc gaccagcaca ccacccatga aaaagaaatc attgagcttc
70261 gcttgcagga caccaacggg atagaaaatt tcgtgaagaa ctttgcgaag tcatattaca
70321 catggaataa cagcaaagaa gcgattgaag caaggacgca ggcaatcagc ggttatctga
70381 caaaggaatt gcaggacttg aatatagata ccattagaac cgacataccg accagctcca
70441 cagttacaga tgtgattgta tggcatatcg agcaatcggg aacggatact ttttctgcta
70501 cctacgaagt agatcagcag ataaaagagg gcgaacagac aagcaatgtg aaagcaacct
70561 atactgtaaa agttcatgta gacgctgacg gggatatggt aatcgttcag aaccctactc
70621 ttgcaccagc aatcgaaaaa tcagactatg atccaaagac accagaaaca gacgcaagtg
70681 tagacgctga tactgcaaat gacgctaccg catttctgga aacattcttt aagttgtacc
70741 caacagccac agaaaacgaa cttgcttatt atgtgtcggg gaatgtgctt gaacctatcg
70801 gcagggacta cctttattct gaactggtaa accctatctt tacaaaagac ggggagaatg
70861 taaaagttaa ggttgctgta aaattccttg ataatcagac aaaggcaacg caagtatcgc
70921 agtatgagct tgtgctacat aaggatagta actggaagat tgtaggataa ttgtaaaagg
70981 cttgtatcat atttttatgc tctgcaagcc ttttaatatt gacaactgta tttttaagta
71041 ttatgctgta cttgtataag ctatacagta tagaacaagg cggtgtttaa atggaaatct
71101 ttatcaataa tcatgcaagc atacctattt atgaacaaat ttcttcgcaa ataaaagcat
71161 tgattataaa tggggaatta aaggctggcg aagctctacc atcaataaga ggacttgcaa
71221 aatcattgca tataagtgta ttgactgttc aaaaatctta tgatgtgctt caaaaagacg
71281 gttttatcga aacaacagca ggcaaagggt gttttgtatc tgctcaaaat caagattttt
71341 atatagaaga acaacagaaa aaaatcgaag atagatttgt agaagctatt gaaatagccc
71401 gtgcaagtgg cataaacctt gataaattga ttggattatt aaaattatta tatgaggagg
71461 acgagtaaat ggctgaaaat attttagaag ttaaaaacct tacaaaagat tatggcgatt
71521 ttgtattaaa tgacttaagt ttttccattc ctagaggtgt aattatgggg cttattggcg
71581 aaaatggagc tggaaaatcg accacagtaa attgtatttt gaacgaaatt aagaaaaaca
71641 gtggaaacat tttgattttt ggaaaagacc acataattga tgaatgtgaa ataaaagata
71701 aaatcggaat tgtatttgac gaaaaccact ttccagatat ttttacacca aaagaaattg
71761 aaaaatatat gtcgggaata tataccaatt gggacagtat agcatttcgt aattatttga
71821 aacagttcga attaccagca aagaagaaaa ttaaagattt ttctaaagga atgaaagtaa
71881 aacttgcatt cgctgttgct ctttcacata aagcagaatt acttatatta gatgaagcaa
71941 caagtggatt ggaccctatt attcgagatg atattcttga tatactgatt gattttgtgc
72001 aagatgaaaa ttactctatt ttgttttcat cacatattac aagtgattta gaaaaaatag
72061 cagattatat tacttttatt cataaaggac ggttagtttt ctgctcatta aaagacgact
72121 taattgataa ctatggtgtt gtaagttgtg gagcaagtgt ctttgatacg ctagataaga
72181 cagaaattat ttcctatcaa aaagaagatt atcaatataa ggttcttgtt cgtgatagaa
72241 acaagattgc aaaagatttt ccaaaagcac ttgtagaacc tgctactatt gaaaatatta
72301 tgttgtttta cgtaaagggg gaattacaat gaaaggctta ttgatgaaag atttattagt
72361 aattcagaaa aaatatggaa ttactagaat tattatggat atattcatta tagtagcttt
72421 gatgttattt ttaaaaggga caggagctat ttacattagt ttattgttaa ttccagttga
72481 aataatgaca atgataattt ccttaacgac ttgcgacgaa caatggaaat gggggaaata
72541 tgttgttgca ctacccgttt caaaaactac gattgtaaaa agcagatata tagttgcttt

72601 gattatgact tgtattggct ttggggtagc tttgcttgtt aataccataa catattttat
72661 attccctcaa tttgcatttg gttattatat gtttattgct tgtgcttctt ttgccgtaac
72721 cttagtattt ttgtcatttg tattgccttc aaattattca cttggagtta atgcaggttt
72781 tgcggttatg tttactctag tgattattct tgttgtattg gcagtttgga caaatgtgac
72841 taacaattct attttatggt ttgtagtcga taattttgaa ttgagtatcg gcatagcttt
72901 tataatagtt atagctcttg ctttcagttc ttatcttcta tcaaaagcat tttttactag
72961 aaaatacaat tagtgaggta ttgttatgag aaaatgtctt agatgtcatg aagaaatggt
73021 cgaaaattta gatgttaaag ttgatatgca gggatatggt attagaataa ctactaaagg
73081 cgtttttgga actacggtag aaaaacctaa agttgcagtt tgtccaaaat gtggcgaagt
73141 ttccttatat attgaaaatt ttaaatcaat aaaataaatg attaaaatct gcatatagac
73201 tgagggttta ttgtttgcta acgtccgagc ggatattgaa gaatttatga atagtacata
73261 cccagagatg caagcaaatt tcaaaaggta ttttggaaat agacatttta cgccgaaaaa
73321 atatgtttat atcactacga aaaagcaaaa attaaatttc ttaaaagaat aagggggggc
73381 tatggataag caattatggg tattatgtcc tatttgcaat aataagacac gaactaaagt
73441 gcgagaagat acagaactga taaattttcc tttgttctgt ccgaagtgca aaacggaaac
73501 tttaatcaat ataaagcatg gaaaagtaac tattatcaaa gagccagacg caaagacgca
73561 gagccgataa tttgtaggtt attatgactt atagattgtt ggctcttatt tgttacctgc
73621 ggagcagatc agcaaataga taatgcagaa tagcaaaaag ggaaaacaag aagctgatta
73681 cttcttgctc tccctgctgt tcaagatacc gtcaatcgta ccttgaatga tttttaattc
73741 gttatcactc aataaatcaa gagaggtttc tatctgacgg cgagctgtac tttttacgac
73801 gttctctgct ggatagaaat attcatcgac cgatatattg aacatggtaa ctaaatcatg
73861 gaataattga aaactagggt gtgaaccaac attttcaata tctgcgatat ggcgttctcc
73921 atagaacact ttatcgccta aatcattacg agaaagccct gctctctcac gggcttctct
73981 gattgccagt cctaaaggtc tgaaatcaaa agtttgtgtt tcttttgctt ttctcatttt
74041 aaatcaccta gcatgagttt atacttcatg ctgtatattt agaatgtgtt aaaattgagc
74101 gaatcattca attagaaaac gaaaaagaat agaatcacaa atcacttgtg atttgtgatt
74161 gttgatgata aaataagagt aagaagaaat agaaagaagt gagtggttgt gggaaattta
74221 ggcgcacaaa aagcaaaacg aaatgataca ccaatcagtg caaaaaaaga tattatgggg
74281 gataagacgg ttcgtgttcg tgctgacttg caccatatta taaaaatcga aacagcaaag
74341 aatggcggaa acgtaaaaga agttatggat aaagccttag aagaatatat tcggaaatat
74401 ttacctgaca aactttaaaa ggagtgaaaa tgaaatggca gttacttatg aaaaaacatt
74461 tgaaatagag atcattaacg aattatcggc aagcgtttat aatcgagtat taaactatgt
74521 tttgaatcat gaattaaata aaaatgactc tcaattattg gaagtcaatt tattaaacca
74581 attaaagctt gcaaaacgtg taaatctttt tgattattct ttagaagaat tacaagccgt
74641 tcatgagtat tggcggtcaa tgaatcgcta ttcaaaacaa gttttgaata aagagaaagt
74701 ggcttaatat ggcaaatata gtcaatttta ctgacaaaca atttgaaaat cgcttaaatg
74761 ataatttaga agaattggtt caaggaaaaa aagcggttga atcgccaacc gcttttttac
74821 ttggtgggca accagggtca gggaaaacga gtttgcgaag aagatgattg agcaggcagg
74881 aacaaagtga taaattcaaa ttgtgaggga aaatatagat gaatcaagga agaattattg
74941 tgattacagg ctcaccgggg acaggaaaaa caacaactgc gtcaattgtc gcaaaggaat
75001 cagatatgga taaatcagtg catatgcaca ccgatgactt ttttcattac ttaagcaaag
75061 gcgcaatacc accacatttg ccagaatcca acaagcaaaa tctggttgtc attgaagcct
75121 ttttagaggc tgcaaagcga tatgctcgtg gcggatatga tgtaattgta gatgggattg
75181 tcgggccatg gtttttggag ccatggaaag ctcttgccca agaagattac gaggtacact

75241 atattgtatt aagagcgagt atggaaataa gacttagaag caaacttaag agtgtgttga
75301 tagtgcagta tcttaaaatt ttgtataata ggaattgaag ttaaattaga tgctaaaaat
75361 ttgtaattaa gaaggagtga ttacatgaac aaaaatataa aatattctca aaacttttta
75421 acgagtgaaa aagtactcaa ccaaataata aaacaattga atttaaaaga aaccgatacc
75481 gtttacgaaa ttggaacagg taaagggcat ttaacgacga aactggctaa aataagtaaa
75541 caggtaacgt ctattgaatt agacagtcat ctattcaact tatcgtcaga aaaattaaaa
75601 ctgaatactc gtgtcacttt aattcaccaa gatattctac agtttcaatt ccctaacaaa
75661 cagaggtata aaattgttgg gagtattcct taccatttaa gcacacaaat tattaaaaaa
75721 gtggtttttg aaagccatgc gtctgacatc tatctgattg ttgaagaagg attctacaag
75781 cgtaccttgg atattcaccg aacactaggg ttgctcttgc acactcaagt ctcgattcag
75841 caattgctta agctgccagc ggaatgcttt catcctaaac caaaagtaaa cagtgtctta
75901 ataaaactta cccgccatac cacagatgtt ccagataaat attggaagct atatacgtac
75961 tttgtttcaa aatgggtcaa tcgagaatat cgtcaactgt ttactaaaaa tcagtttcat
76021 caagcaatga aacacgccaa agtaaacaat ttaagtaccg ttacttatga gcaagtattg
76081 tctattttta atagttatct attatttaac gggaggaaat aattctatga gtcgcttttg
76141 taaatttgga aagttacacg ttactaaagg gaatgtagat aaattattag gtatactact
76201 gacagcttcc aaggagctaa agaggtccct agcgcctacg gggaatttgt atcgataagg
76261 ggtacaaatt cccactaagc gctcgggacc ccttgtagga aaatgtccta agtgtggcaa
76321 caatattgta ttaaaaaaat cgttttatgg ttgttcaaat tatcctgaat gtaccttcac
76381 tttagctgaa cattttagaa agaaaaaact caccaaaaca aatgtaaaag aattactaga
76441 gggaaaagaa accctggtaa aaggaatcaa aacgaaagat agaaagtcct acaatgccgt
76501 tgtaaaaatc ggagaaaagg gatatattga ttttatctct ttttcaaaat aagcataaaa
76561 gccctttaaa gagggctttt atatattaat cacaaatcac ttatcacaaa tcacaagtga
76621 ttaatcacaa atcacttgtg atttgtgatt cctaatgata caatattact atacaaaaaa
76681 agaatggggc gtagttatgg agaaggaaga actcaaaata cttgaagaat taagacgtat
76741 tttaaacagt aaaaatgaag cgattgttat cttaaacaat tattttaaag gtggtgttgg
76801 aaagtcaaaa ttatcgacta tgtttgctta cttgacagac aaatttaatt taaaagtttt
76861 aatgatcgat aaggacttac aagcaacatt gacaaaagac ttagcaaaaa catttaaggt
76921 agaattgcca cgtgttaatt tttatgaagg actgaaaaat ggaaacttgg cttcttctat
76981 tgttcatttg actgataatt tagacttgat ccctggcacg tttgatttga tgttactgcc
77041 aaaattaact cgctcatgga cttttgaaaa tgaaagtaga ttgcttgcta ctcttttagc
77101 acctttaaaa agtgactatg atctcattat tattgatact gtaccaacgc caagcgttta
77161 tacaaataat gcaatcgtgg caagtgatta cgtcatgatc cctttacaag cagaagaaga
77221 aagtacaaac aacattcaaa actatatttc ctatttgatt gatttacaag aacagtttaa
77281 ccctggacta gatatgatcg gttttgttcc ttatttagtt gatacggaca gcacaacgat
77341 aaaatcaaac ctggaagaac tgtacaagca acataaagaa gataacttgg ttttccaaaa
77401 tattatcaag cgaagtaata aagtaagtac ctggtctaaa aacggcatta cagaacacaa
77461 aggctatgac aaaaaagttt tatctatgta tgagaacgta ttttttgaaa tgcttgcata
77521 tagtatgcag taataattca aaagctgtta tcgtgtgcgg taaataaaaa aaaccgttcc
77581 tagataatac acttttgtgc gtcttgaaaa ataacgggac ggttttggat ataaccagaa
77641 ctgtcctttt ttcttaggag taaataaggc atagctctgg ttaaaggggg ctataccatg
77701 attgcatacg gaagaacgtt ctgccgttct caactgaata aatatttcag tccactatta
77761 caatcattac accctttaca agaattaaaa attcatatca atagaattta taaaacagtc
77821 tgtttgtgta attacacatt atcgggctgt ttttgttttg ttaaaaattt ttttcaactt

77881 ttttcggttt caacttagca aatcacacct gtcgttccct atatggtgcg gaaagaggga
77941 taaccacttt tcatatcgta acggaaaggg ggtgagattg atgaaaccat ctgacttcca
78001 gaaaacagtt caatgccgtt ttgaaagttg tttgaagaaa gttgtccgac atgttgttaa
78061 ggactatcag aagaaattaa agcgacgaca aaagggagaa acgctttttt gtgaacttcc
78121 agaaatcgtt gtagaaaatc tggctgtctg ggacgactat gatacggact atacgatttt
78181 caatgtatgt ggtagtgata ttcgtatttg tgatgatgaa ctggcagaag ccttgaaaca
78241 tttgtctgag cgtaatcgag aaaatttgtt gatgtattat ttcttggaaa tgagcgatac
78301 cgagattgca aaaagacaaa atatttcaag aagtggtgtt tttcaaaacc gacacaattc
78361 actagagctt atgaaaaaaa tactaaagga gaaatgatag caaatgaaac acacaatgaa
78421 gaagccgtcc tactacttgc tttcgcaggc aatggacggg gacgaaaaag cgattgaaaa
78481 gatactggca ttttatgacc cttacatatc aaagtgctgt ctacgaccac tctatgatga
78541 atacggcaat gtctatattg ttgtagatat ggaattaaag ggacttatca gagaagcact
78601 tattaaaatg attttagggt ttgatattgc cttagaaatc gaagaagaat aacaagcagt
78661 aaccgcagag catgatttgt tcaatcccct ctttcctgct ctgcccctgc ttttatatga
78721 aagcaacacg ctttcgccac agctctttga caactgaata aagcagacag atacgtttgt
78781 gagatagcga gccacgggaa ctgtacgcca cgaccttttc aaaagaaagc gagcgaccca
78841 cgcagtgatc cgagagcgac tgttggaaaa gtcgttttgc catgacctat cctcacagaa
78901 taatgatacg tccgcatggt gcggttctgc ccacaagaat gggaatagtt gagatactaa
78961 cggagctttc caaagccgtc tgcctgctta taattgatag tttcagcaac aaaatctatc
79021 tactcttttt atttaggtgt gttataatga ccttattaga cgagtgtatt ttgttgctgt
79081 cacaaatgaa aggggtctta atagtgataa gtgatacaca aaatgcacaa tctcctacaa
79141 aggaaaaaaa gacttataca gtagaggaaa ttgctgaaca actggacatt agcaaaaagg
79201 tagcttattc attagtaaaa agcggtcaat tcagttatgt tcgtgctggt aaagccatac
79261 gggtatcaaa ggtttctttc gataaatggt taaatgaaat ttaactataa ggagagaaaa
79321 ttatggcttc aatcaaacaa agaaaatcaa gtttttcagt tatttattgg tacttagata
79381 gtgcaggaga aagaaaacaa aaatgggaca cgctggaaac aagaaaagaa gcaaaacagc
79441 gaaaagcctt tatagaatac tatcaagaga aatttggata tgtgatagtt ccattagagg
79501 aacaatttgc acatcagatt gaaaattcta aaaaggagtt agatgttgta gacaaagata
79561 ttacattgcg tgatttccta gaaatttttg ttaatttata tggcacatct aaatggtcgc
79621 cgagtacatt tagtagtaag gtaggcagta ttgaaaatta tatcaatccg attattggag
79681 attggaaact gaatgaaata actacaaaaa gtctatcaaa atactacaat gatttattga
79741 gcgttccaga agtaccacgg gcaaacagaa aagcaactgg tcgttgtgtt caaccagcga
79801 acatcaaaaa aatacatgat attatccgct gtgccttaaa tcaagctatc cgctgggagt
79861 atattgatac aaacaaaaga aaccctgcgt ctttggctac gttaccaaaa attccaaaga
79921 ctaggcgaaa agtgtggagt gttcaaacat ttcgagaagc ggtaaaagcc acggaagatg
79981 atttgctttc aatttgtatg cacttagctt tttcctgctc tatgcgaatt ggagaaatta
80041 caggacttac atgggaagat gttattattg atgaagagtc cattgctaca aataatgcaa
80101 aggtaatcat caataaagaa ctttcaagag taaatgcaga agctatgcag aaattaaagg
80161 aaaaggatat tttaaagatt ttcccaacgc aaaaaccgca ttgtaccaca aggcttgtct
80221 tgaaaacgcc aaaaacagaa acaagtaatc gtgttgtctg gttacctaaa acagttgcgg
80281 aactattggt gcaatataaa aaagaccagc aggaattaaa ggaattttta ggaagtgcct
80341 acaatgatta caaccttgtt attgcgttgg ataatggcaa tcccgtggaa agccgtattg
80401 ttagagatag gtttcaaaag ctctgtgaag aaaacgatta tgaagttgtg gtatttcaca
80461 gcctacgcca tttaagtacg ggttataaac tcaaaatgac aaacggagat gttaagtccg

```
   80521 tacaaggcga tactggtcac gcagaagcag aaatggttac agatgtatat tcagaaatca
   80581 ttgacgagga cagacgtttc aatgcacaaa aaatggataa agaattttat tctacgctta
   80641 atgatgatac agacaatcca aagcaagata ccgacttaac cgatagtgat atggcgttat
   80701 tggaactgat taaatcttta tcgccagaaa tgaaagagca atttttaaag caggcacttc
   80761 aaagatagaa tttcaaaagt gttagcaaat tccaaaaaaa ccagcgattt ttcttcaaaa
   80821 ataaaaaagt gttagcaagc ctatttcctt aaaagtctta aaaaaatatc gaataaaaaa
   80881 tgacagattg aaaacaattc gtattcagcg taaatagaaa gccataaata aagaaacgct
   80941 gaaaaccttg taaaataagg ctttcagcgg agtttgtggc gtcccagaca
//
DEFINITION     Integrative and mobilisable element IME_1
     fasta_record    1..122003
                /ID=B40|SC|contig000001
                /colour=5
     CDS          164..367
                /ID=B40_02170
                /color=1
                /locus_tag="B40_02170"
                /note="hypothetical protein"
                /product="hypothetical protein"
                /protein_id="gnl|SC|B40_02170"
                /codon_start=1
     CDS          371..727
                /ID=B40_02171
                /color=1
                /locus_tag="B40_02171"
                /note="hypothetical protein"
                /product="hypothetical protein"
                /protein_id="gnl|SC|B40_02171"
                /codon_start=1
     CDS          745..2586
                /color=10
                /locus_tag="B40_02172"
                /note="Replication initiation protein RepA, protein
                motif:CLUSTERS:PRK11823 protein motif:Pfam:PF09250.5"
                /product="Replication initiation protein RepA
                [replication]"
                /"ID=B40_02172
                /gene="repA"""
                /protein_id="gnl|SC|B40_02172"
                /codon_start=1
     CDS          2779..3141
                /ID=B40_02173
                /color=1
                /locus_tag="B40_02173"
```

/note="hypothetical protein"
/product="hypothetical protein"
/protein_id="gnl|SC|B40_02173"
/codon_start=1
CDS 3110..3460
/ID=B40_02174
/color=10
/locus_tag="B40_02174"
/note="replication initiation protein RepA"
/product="Replication initiation protein RepA
[replication]"
/protein_id="gnl|SC|B40_02174"
/gene="repA"
/codon_start=1
CDS 3532..4704
/color=7
/locus_tag="B40_02175"
/note="RNA polymerase sigma factor  Sigma-70 region 2,
similar to AA sequence:UniProtKB:P77994 protein
motif:CLUSTERS:PRK05901 protein motif:TIGRFAMs:TIGR02393
protein motif:Pfam:PF04542.8"
/product="RNA polymerase sigma factor  Sigma-70 region 2
[transcriptional regulator]"
/"ID=B40_02175
/protein_id="gnl|SC|B40_02175"""
/codon_start=1
CDS 5132..5782
/ID=B40_02176
/color=1
/locus_tag="B40_02176"
/note="hypothetical protein"
/product="hypothetical protein"
/protein_id="gnl|SC|B40_02176"
/codon_start=1
CDS 5932..6417
/ID=B40_02177
/color=7
/locus_tag="B40_02177"
/note="helix-turn-helix [transcriptional regulator]"
/product="helix-turn-helix [transcriptional regulator]"
/protein_id="gnl|SC|B40_02177"
/codon_start=1
CDS complement(6735..7280)
/color=8

/locus_tag="B40_02178"
/note="Uma2 family endonuclease, protein motif:Cdd:COG4636
protein motif:Pfam:PF05685.6"
/product="Uma2 family endonuclease [restriction
modification]"
/"ID=B40_02178
/protein_id="gnl|SC|B40_02178"""
/codon_start=1
CDS 7994..8269
/ID=B40_02179
/color=8
/locus_tag="B40_02179"
/note="putative phage-type endonuclease"
/product="Putative phage-type endonuclease [restriction
modification]"
/protein_id="gnl|SC|B40_02179"
/codon_start=1
CDS 8379..8852
/ID=B40_02180
/color=1
/locus_tag="B40_02180"
/note="hypothetical protein"
/product="hypothetical protein"
/protein_id="gnl|SC|B40_02180"
/codon_start=1
CDS 8941..9234
/ID=B40_02181
/color=1
/locus_tag="B40_02181"
/note="hypothetical protein"
/product="hypothetical protein"
/protein_id="gnl|SC|B40_02181"
/codon_start=1
CDS 9270..9665
/ID=B40_02182
/color=1
/locus_tag="B40_02182"
/note="hypothetical protein"
/product="hypothetical protein"
/protein_id="gnl|SC|B40_02182"
/codon_start=1
CDS 9653..9832
/ID=B40_02183
/color=1

/locus_tag="B40_02183"
/note="hypothetical protein"
/product="hypothetical protein"
/protein_id="gnl|SC|B40_02183"
/codon_start=1
CDS 9905..10171
/ID=B40_02184
/color=1
/locus_tag="B40_02184"
/note="hypothetical protein"
/product="hypothetical protein"
/protein_id="gnl|SC|B40_02184"
/codon_start=1
CDS 10337..10582
/ID=B40_02185
/color=1
/locus_tag="B40_02185"
/note="hypothetical protein"
/product="hypothetical protein"
/protein_id="gnl|SC|B40_02185"
/codon_start=1
CDS 10594..10881
/ID=B40_02186
/color=9
/locus_tag="B40_02186"
/note="tyrosine-type recombinase"
/product="Tyrosine-type recombinase [recombinase]"
/protein_id="gnl|SC|B40_02186"
/codon_start=1
CDS 11079..12212
/ID=B40_02187
/color=7
/locus_tag="B40_02187"
/note="Helix-turn-helix domain DNA-binding protein conjugative transposon protein"
/product="Helix-turn-helix domain DNA-binding protein conjugative transposon protein [transcriptional regulator]"
/protein_id="gnl|SC|B40_02187"
/codon_start=1
CDS 12230..12442
/ID=B40_02188
/color=1
/locus_tag="B40_02188"

/DUF5348=containing
/DUF5348=protein
/DUF5348=conugative
/DUF5348=transposon
/DUF5348=protein
/note="hypothetical protein"
/product="hypothetical protein  DUF5348 containing protein conugative transposon protein"
/protein_id="gnl|SC|B40_02188"
/codon_start=1
CDS 12486..12962
/color=1
/locus_tag="B40_02189"
/note="PcfB family protein  DUF3801 domain-containing protein  conjugative transposon protein, protein motif:Pfam:PF12687.1"
/product="PcfB family protein  DUF3801 domain-containing protein  conjugative transposon protein"
/"ID=B40_02189
/protein_id="gnl|SC|B40_02189""""
/codon_start=1
CDS 12959..14506
/color=4
/locus_tag="B40_02190"
/note="Type IV secretory systeim protein TraG  VirD4 components, protein motif:CLUSTERS:PRK13876 protein motif:Cdd:COG3505 protein motif:TIGRFAMs:TIGR02759 protein motif:Pfam:PF02534.8"
/product="Type IV secretory systeim protein TraG  VirD4 components [coupling protein]"
/"ID=B40_02190
/protein_id="gnl|SC|B40_02190""""
/gene="traG"
/codon_start=1
CDS 14597..14974
/ID=B40_02191
/color=9
/locus_tag="B40_02191"
/note="transposon-encoded TnpV family protein"
/product="Transposon-encoded TnpV family protein"
/protein_id="gnl|SC|B40_02191"
/gene="tnpV"
/codon_start=1
CDS 14985..15176

/ID=B40_02192
/color=1
/locus_tag="B40_02192"
/note="hypothetical protein"
/product="hypothetical protein"
/protein_id="gnl|SC|B40_02192"
/codon_start=1
CDS complement(15196..16239)
/ID=B40_02193
/color=1
/locus_tag="B40_02193"
/note="hypothetical protein"
/product="hypothetical protein"
/protein_id="gnl|SC|B40_02193"
/codon_start=1
CDS 16571..16795
/color=7
/locus_tag="B40_02194"
/note="Helix-turn-helix domain-containing protein Excisionase family DNA-binding domain, protein motif:Pfam:PF12728.1"
/product="Helix-turn-helix domain-containing protein Excisionase family DNA-binding domain [transcriptional regulator]"
/"ID=B40_02194
/protein_id="gnl|SC|B40_02194"""
/codon_start=1
CDS 16911..17117
/ID=B40_02195
/color=1
/locus_tag="B40_02195"
/note="hypothetical protein"
/product="hypothetical protein"
/protein_id="gnl|SC|B40_02195"
/codon_start=1
CDS 17178..18686
/ID=B40_02196
/color=1
/locus_tag="B40_02196"
/note="SRRM2 serine/arginine repetitive matrix protein"
/product="SRRM2 %28serine/arginine repetitive matrix protein 2%29"
/protein_id="gnl|SC|B40_02196"
/codon_start=1

CDS 18702..19388
/color=10
/locus_tag="B40_02197"
/note="DNA primase, similar to AA
sequence:UniProtKB:P0ABS5 protein motif:CLUSTERS:PRK05667
protein motif:Cdd:COG0358 protein motif:TIGRFAMs:TIGR01391
protein motif:Pfam:PF01807.14"
/product="DNA primase [RNA polymerase]"
/"ID=B40_02197
/protein_id="gnl|SC|B40_02197""""
/codon_start=1
CDS 19348..20706
/color=8
/locus_tag="B40_02198"
/note="Virulence-associated protein E, protein
motif:Cdd:COG5545 protein motif:Pfam:PF05272.5"
/product="Virulence-associated protein E [virulence]"
/"ID=B40_02198
/protein_id="gnl|SC|B40_02198""""
/codon_start=1
CDS 20974..21201
/ID=B40_02199
/color=1
/locus_tag="B40_02199"
/note="hypothetical protein"
/product="hypothetical protein"
/protein_id="gnl|SC|B40_02199"
/codon_start=1
CDS 21214..22863
/ID=B40_02200
/color=9
/locus_tag="B40_02200"
/note="protein motif:Pfam:PF00239.15"
/product="Serine recombinase [recombinase]"
/protein_id="gnl|SC|B40_02200"
/codon_start=1
CDS complement(22934..23854)
/color=7
/locus_tag="B40_02201"
/note="AraC family transcriptional regulator
helix-turn-helix domain, protein motif:CLUSTERS:PRK10572
protein motif:Cdd:COG2169 protein motif:TIGRFAMs:TIGR04094
protein motif:Pfam:PF12833.1"
/product="AraC family transcriptional regulator

helix-turn-helix domain [transcriptional regulator]"
/"ID=B40_02201
/protein_id="gnl|SC|B40_02201"""
/codon_start=1
CDS 24070..25083
/color=8
/locus_tag="B40_02202"
/note="ABC transporter substrate-binding protein yclQ,
similar to AA sequence:UniProtKB:P94421 protein
motif:CLUSTERS:PRK10957 protein motif:Cdd:COG4594 protein
motif:Pfam:PF01497.12"
/product="ABC transporter substrate-binding protein yclQ
[transporter]"
/"ID=B40_02202
/protein_id="gnl|SC|B40_02202"""
/gene="yclQ"
/codon_start=1
CDS 25049..26122
/color=8
/locus_tag="B40_02203"
/note="Iron-uptake system permease protein FeuB iron ABC
transporter permease, similar to AA
sequence:UniProtKB:P40410 protein motif:CLUSTERS:PRK10577
protein motif:Cdd:COG0609 protein motif:TIGRFAMs:TIGR03869
protein motif:Pfam:PF01032.12"
/product="Iron-uptake system permease protein FeuB iron
ABC transporter permease [transporter]"
/"ID=B40_02203
/protein_id="gnl|SC|B40_02203"""
/gene="feuB"
/codon_start=1
CDS 26119..27114
/color=8
/locus_tag="B40_02204"
/note="Iron-uptake system permease protein FeuC iron ABC
transporter permease, similar to AA
sequence:UniProtKB:P40411 protein motif:CLUSTERS:PRK10577
protein motif:Cdd:COG0609 protein motif:TIGRFAMs:TIGR03869
protein motif:Pfam:PF01032.12"
/product="Iron-uptake system permease protein FeuC iron
ABC transporter permease [transporter]"
/"ID=B40_02204
/protein_id="gnl|SC|B40_02204"""
/gene="feuC"

/codon_start=1
CDS 27128..27940
/color=8
/locus_tag="B40_02205"
/note="Iron dicitrate  ABC transporter ATP-binding
protein, similar to AA sequence:UniProtKB:O32188 protein
motif:CLUSTERS:PRK11231 protein motif:Cdd:COG4604 protein
motif:TIGRFAMs:TIGR03873 protein motif:Pfam:PF00005.21"
/product="Iron dicitrate  ABC transporter ATP-binding
protein [transporter]"
/"ID=B40_02205
/protein_id="gnl|SC|B40_02205"""
/codon_start=1
CDS 28083..29861
/color=8
/locus_tag="B40_02206"
/eC_number=3.6.3.-
/note="Iron import ATP-binding/permease protein IrtA,
similar to AA sequence:UniProtKB:P63391 protein
motif:CLUSTERS:PRK13657 protein motif:Cdd:COG2274 protein
motif:TIGRFAMs:TIGR02204 protein motif:Pfam:PF00005.21"
/product="Iron import ATP-binding/permease protein IrtA
[transporter]"
/"ID=B40_02206
/protein_id="gnl|SC|B40_02206"""
/gene="irtA"
/codon_start=1
CDS 29858..31606
/color=8
/locus_tag="B40_02207"
/note="Putative multidrug export ATP-binding/permease
protein SAV1866, similar to AA sequence:UniProtKB:Q99T13
protein motif:CLUSTERS:PRK13657 protein motif:Cdd:COG2274
protein motif:TIGRFAMs:TIGR02203 protein
motif:Pfam:PF00664.17"
/product="Putative multidrug export ATP-binding/permease
protein SAV1866 [transporter]"
/"ID=B40_02207
/protein_id="gnl|SC|B40_02207"""
/codon_start=1
CDS 32071..32439
/color=8
/locus_tag="B40_02208"
/note="MN-binding protein  fumarate reductase flavoprotein

subunit, protein motif:Cdd:COG4659 protein
motif:TIGRFAMs:TIGR01947 protein motif:Pfam:PF04205.8"
/product="FMN-binding protein  fumarate reductase
flavoprotein subunit"
/"ID=B40_02208
/protein_id="gnl|SC|B40_02208"""
/codon_start=1
CDS 32795..33034
/ID=B40_02209
/color=4
/locus_tag="B40_02209"
/note="TraM recognition domain-containing protein  type IV
secretory system conjugative DNA transfer family protein"
/product="TraM recognition domain-containing protein  type
IV secretory system conjugative DNA transfer family
protein [relaxosome]"
/protein_id="gnl|SC|B40_02209"
/codon_start=1
CDS 33099..33239
/color=1
/locus_tag="B40_02210"
/note="Maff2 family protein, protein motif:Pfam:PF12750.1"
/product="Maff2 family protein"
/"ID=B40_02210
/protein_id="gnl|SC|B40_02210"""
/codon_start=1
CDS 33389..34429
/color=7
/locus_tag="B40_02211"
/note="Transcriptional repressor DicA  helix-turn-helix,
protein motif:CLUSTERS:PRK09706 protein motif:Cdd:COG3800
protein motif:TIGRFAMs:TIGR03070 protein
motif:Pfam:PF12844.1"
/product="Transcriptional repressor DicA  helix-turn-helix
[transcriptional regulator]"
/"ID=B40_02211
/protein_id="gnl|SC|B40_02211"""
/codon_start=1
CDS 34596..35039
/color=7
/locus_tag="B40_02212"
/note="RNA polymerase sigma factor  Sigma-70  region 4,
protein motif:CLUSTERS:PRK11924 protein
motif:TIGRFAMs:TIGR02937 protein motif:Pfam:PF08281.6"

/product="RNA polymerase sigma factor  Sigma-70  region 4
[transcriptional regulator]"
/"ID=B40_02212
/protein_id="gnl|SC|B40_02212"""
/codon_start=1
CDS 35032..35232
/color=7
/locus_tag="B40_02213"
/note="helix-turn-helix domain-containing protein, protein
motif:Pfam:PF12645.1"
/product="helix-turn-helix domain-containing protein
[transcriptional regulator]"
/"ID=B40_02213
/protein_id="gnl|SC|B40_02213"""
/codon_start=1
CDS 35288..35659
/ID=B40_02214
/color=1
/locus_tag="B40_02214"
/note="hypothetical protein"
/product="hypothetical protein"
/protein_id="gnl|SC|B40_02214"
/codon_start=1
CDS 36276..36671
/ID=B40_02215
/color=4
/locus_tag="B40_02215"
/note="plasmid mobilization relaxosome protein MobC"
/product="Plasmid mobilization relaxosome protein MobC
[relaxosome]"
/protein_id="gnl|SC|B40_02215"
/gene="mobC"
/codon_start=1
CDS 36634..38265
/color=4
/locus_tag="B40_02216"
/note="Relaxase/Mobilisation nuclease domain , protein
motif:Pfam:PF03432.8"
/product="Relaxase/Mobilisation nuclease domain
[relaxase]"
/"ID=B40_02216
/protein_id="gnl|SC|B40_02216"""
/codon_start=1
CDS 38450..38662

/ID=B40_02217
/color=1
/locus_tag="B40_02217"
/note="hypothetical protein"
/product="hypothetical protein"
/protein_id="gnl|SC|B40_02217"
/codon_start=1
CDS 38652..39395
/color=10
/locus_tag="B40_02218"
/note="phage replisome organizer N-terminal region,
protein motif:TIGRFAMs:TIGR01714 protein
motif:Pfam:PF09681.4"
/product="phage replisome organizer N-terminal region
[replication]"
/"ID=B40_02218
/protein_id="gnl|SC|B40_02218"""
/codon_start=1
CDS 39392..40249
/color=10
/locus_tag="B40_02219"
/note="Primosomal protein DnaI ATP binding protein,
similar to AA sequence:UniProtKB:P06567 protein
motif:CLUSTERS:PRK08116 protein motif:Cdd:COG1484 protein
motif:Pfam:PF01695.11"
/product="Primosomal protein DnaI ATP binding protein
[replication]"
/"ID=B40_02219
/protein_id="gnl|SC|B40_02219"""
/gene="dnaI"
/codon_start=1
CDS 40246..40434
/ID=B40_02220
/color=9
/locus_tag="B40_02220"
/note="transposon-encoded protein:"
/product="Transposon-encoded protein TnpW"
/protein_id="gnl|SC|B40_02220"
/gene="tnpW"
/codon_start=1
CDS 40546..40746
/ID=B40_02221
/color=9
/locus_tag="B40_02221"

/note="Serine recombinase TnpX"
/product="Serine recombinase TnpX [recombinase]"
/protein_id="gnl|SC|B40_02221"
/gene="tnpX"
/codon_start=1
CDS complement(40739..42412)
/color=9
/locus_tag="B40_02222"
/note="Serine recombinase, protein motif:Pfam:PF00239.15"
/product="Serine recombinase [recombinase]"
/"ID=B40_02222
/protein_id="gnl|SC|B40_02222"""
/codon_start=1
CDS complement(42490..42663)
/ID=B40_02223
/color=9
/locus_tag="B40_02223"
/note="Transposon-encoded protein TnpW"
/product="Transposon-encoded protein TnpW"
/protein_id="gnl|SC|B40_02223"
/gene="tnpW"
/codon_start=1
CDS complement(43122..43364)
/color=7
/locus_tag="B40_02224"
/note="helix-turn-helix domain-containing protein, protein motif:Pfam:PF12645.1"
/product="helix-turn-helix domain-containing protein [transcriptional regulator]"
/"ID=B40_02224
/protein_id="gnl|SC|B40_02224"""
/codon_start=1
CDS complement(43348..43785)
/color=7
/locus_tag="B40_02225"
/note="RNA polymerase sigma factor  Sigma-70  region 4, protein motif:CLUSTERS:PRK11924 protein motif:TIGRFAMs:TIGR02937 protein motif:Pfam:PF08281.6"
/product="RNA polymerase sigma factor  Sigma-70  region 4 [transcriptional regulator]"
/"ID=B40_02225
/protein_id="gnl|SC|B40_02225"""
/codon_start=1
CDS complement(44259..45242)

/color=8
/locus_tag="B40_02226"
/eC_number=2.1.1.192
/note="23S rRNA adenine 2503 methyl transferase RlmN,
similar to AA sequence:UniProtKB:Q7A600 protein
motif:CLUSTERS:PRK14469 protein motif:TIGRFAMs:TIGR00048
protein motif:Pfam:PF04055.15"
/product="23S rRNA
%28adenine%282503%29-C%282%29%29-methyltransferase RlmN
[methyltransferase]"
/"ID=B40_02226
/protein_id="gnl|SC|B40_02226"""
/gene="rlmN"
/codon_start=1
CDS 45843..46199
/color=7
/locus_tag="B40_02227"
/note="helix-turn-helix, protein motif:CLUSTERS:PRK08154
protein motif:Cdd:COG3800 protein motif:Pfam:PF12844.1"
/product="helix-turn-helix [transcriptional regulator]"
/"ID=B40_02227
/protein_id="gnl|SC|B40_02227"""
/codon_start=1
CDS 46401..46730
/color=4
/locus_tag="B40_02228"
/note="Plasmid mobilization relaxosome protein MobC,
protein motif:Pfam:PF05713.5"
/product="Plasmid mobilization relaxosome protein MobC
[relaxosome]"
/"ID=B40_02228
/protein_id="gnl|SC|B40_02228"""
/gene="mobC"
/codon_start=1
CDS 46691..48025
/color=4
/locus_tag="B40_02229"
/note="Relaxase/Mobilisation nuclease domain, protein
motif:Pfam:PF03432.8"
/product="Relaxase/Mobilisation nuclease domain
[relaxase]"
/"ID=B40_02229
/protein_id="gnl|SC|B40_02229"""
/codon_start=1

CDS complement(48227..48436)
/ID=B40_02230
/color=4
/locus_tag="B40_02230"
/note="putative tranposon-transfer assisting protein
TTRAP"
/product="Putative tranposon-transfer assisting protein
TTRAP"
/protein_id="gnl|SC|B40_02230"
/codon_start=1
CDS complement(48441..51788)
/color=4
/locus_tag="B40_02231"
/note="Antirestriction protein ArdA, protein
motif:Pfam:PF07275.5"
/product="Antirestriction protein ArdA [anti-ristriction
modification]"
/"ID=B40_02231
/protein_id="gnl|SC|B40_02231"""
/gene="ardA"
/codon_start=1
CDS complement(51914..52282)
/ID=B40_02232
/color=4
/locus_tag="B40_02232"
/note="conjugal transfer protein Cysteine-rich VLP"
/product="conjugal transfer protein  Cysteine-rich VLP"
/protein_id="gnl|SC|B40_02232"
/codon_start=1
CDS complement(52279..54354)
/color=10
/locus_tag="B40_02233"
/eC_number=5.99.1.2
/note="Type IA DNA topoisomerase, similar to AA
sequence:UniProtKB:P14294 protein motif:CLUSTERS:PRK07726
protein motif:Cdd:COG0550 protein motif:TIGRFAMs:TIGR01056
protein motif:Pfam:PF01131.14"
/product="Type IA DNA topoisomerase [topoisomerase]"
/"ID=B40_02233
/protein_id="gnl|SC|B40_02233"""
/codon_start=1
CDS complement(54351..55100)
/color=1
/locus_tag="B40_02234"

/note="hypothetical protein, DUF4366 containing protein"
/product="hypothetical protein  DUF4366 containing
protein"
/"ID=B40_02234
/protein_id="gnl|SC|B40_02234"""
/codon_start=1
CDS complement(55090..55356)
/ID=B40_02235
/color=1
/locus_tag="B40_02235"
/note="hypothetical protein DUF4315 containing protein"
/product="hypothetical protein  DUF4315 containing
protein"
/protein_id="gnl|SC|B40_02235"
/codon_start=1
CDS complement(55363..56085)
/color=8
/locus_tag="B40_02236"
/note="phosphoadenosine phosphosulfate reductase, protein
motif:CLUSTERS:PRK13794 protein motif:Cdd:COG5270"
/product="phosphoadenosine phosphosulfate reductase
[reductase]"
/"ID=B40_02236
/protein_id="gnl|SC|B40_02236"""
/codon_start=1
CDS complement(56099..58069)
/color=8
/locus_tag="B40_02237"
/eC_number=3.4.19.11
/note="D-gamma-glutamyl-meso-diaminopimelic acid
endopeptidase CwlS , similar to AA
sequence:UniProtKB:O31852 protein motif:CLUSTERS:PRK13914
protein motif:Cdd:COG3584 protein motif:TIGRFAMs:TIGR02594
protein motif:Pfam:PF00877.13"
/product="D-gamma-glutamyl-meso-diaminopimelic acid
endopeptidase CwlS [pepidase]"
/"ID=B40_02237
/protein_id="gnl|SC|B40_02237"""
/gene="cwlS"
/codon_start=1
CDS complement(58066..59022)
/color=8
/locus_tag="B40_02238"
/eC_number=2.1.1.72

/note="Adenine specific DNA methyltransferase, similar to
AA sequence:UniProtKB:P09358 protein
motif:CLUSTERS:PRK11524 protein motif:Cdd:COG2189 protein
motif:Pfam:PF01555.12"
/product="Adenine specific DNA methyltransferase
[methyltransferase]"
/"ID=B40_02238
/protein_id="gnl|SC|B40_02238"""
/codon_start=1
CDS complement(59019..61415)
/color=4
/note="Type IV secretory system protein TraC VirB4
components AAA domain, protein motif:Cdd:COG3451 protein
motif:TIGRFAMs:TIGR02746 protein
motif:Pfam:PF12846.1locus_tag=B40_02239product=Conjugal"
/product="Type IV secretory system protein TraC VirB4
components AAA domain [ATPase mating pair formation]"
/"ID=B40_02239
/gene="traC"""
/protein_id="gnl|SC|B40_02239"
/codon_start=1
CDS complement(61375..61767)
/color=4
/locus_tag="B40_02240"
/note="PrgI family protein Type IV secretion system,
protein motif:Pfam:PF12666.1"
/product="PrgI family protein Type IV secretion system"
/"ID=B40_02240
/protein_id="gnl|SC|B40_02240"""
/codon_start=1
CDS complement(61769..62317)
/color=8
/locus_tag="B40_02241"
/note="Transcriptional activator adenine-specific DNA
methyltransferase, protein motif:Cdd:COG4725 protein
motif:Pfam:PF05063.8"
/product="Transcriptional activator adenine-specific DNA
methyltransferase [methyltransferase]"
/"ID=B40_02241
/protein_id="gnl|SC|B40_02241"""
/codon_start=1
CDS complement(62328..63191)
/ID=B40_02242
/color=1

/locus_tag="B40_02242"
/note="hypothetical protein  membrane protein  putative
conjugative transposon protein"
/product="hypothetical protein  membrane protein  putative
conjugative transposon protein"
/protein_id="gnl|SC|B40_02242"
/codon_start=1
CDS complement(63448..65058)
/color=9
/locus_tag="B40_02243"
/note="Serine recombinase, protein motif:Pfam:PF07508.7"
/product="Serine recombinase [recombinase]"
/"ID=B40_02243
/protein_id="gnl|SC|B40_02243"""
/codon_start=1
CDS complement(65159..65341)
/ID=B40_02244
/color=9
/locus_tag="B40_02244"
/note="transposon-encoded protein TnpW"
/product="Transposon-encoded protein TnpW"
/protein_id="gnl|SC|B40_02244"
/gene="tnpW"
/codon_start=1
CDS complement(65338..66195)
/color=10
/locus_tag="B40_02245"
/note="Primosomal protein DnaI  ATP binding protein,
similar to AA sequence:UniProtKB:P06567 protein
motif:CLUSTERS:PRK08116 protein motif:Cdd:COG1484 protein
motif:Pfam:PF01695.11"
/product="Primosomal protein DnaI  ATP binding protein
[replication]"
/"ID=B40_02245
/protein_id="gnl|SC|B40_02245"""
/gene="dnaI"
/codon_start=1
CDS complement(66192..66950)
/color=10
/locus_tag="B40_02246"
/note="phage replisome organizer protein, protein
motif:TIGRFAMs:TIGR01714 protein motif:Pfam:PF09681.4"
/product="phage replisome organizer protein [replication]"
/"ID=B40_02246

/protein_id="gnl|SC|B40_02246"""
/codon_start=1
CDS complement(67318..68940)
/ID=B40_02247
/color=4
/locus_tag="B40_02247"
/note="Relaxase/Mobilisation nuclease domain:protein motif:Pfam:PF03432.8"
/product="Relaxase/Mobilisation nuclease domain [relaxase]"
/protein_id="gnl|SC|B40_02247"
/codon_start=1
CDS complement(68912..69310)
/ID=B40_02248
/color=4
/locus_tag="B40_02248"
/note="Plasmid mobilization relaxosome protein MobC"
/product="Plasmid mobilization relaxosome protein MobC [relaxosome]"
/protein_id="gnl|SC|B40_02248"
/gene="mobC"
/codon_start=1
CDS complement(69844..70047)
/color=7
/locus_tag="B40_02249"
/note="helix-turn-helix domain-containing protein, protein motif:Pfam:PF12645.1"
/product="helix-turn-helix domain-containing protein [transcriptional regulator]"
/"ID=B40_02249
/protein_id="gnl|SC|B40_02249"""
/codon_start=1
CDS 70742..72661
/color=2
/locus_tag="B40_02250"
/note="Tetracycline resistance protein tetM from transposon Tn916, similar to AA sequence:UniProtKB:P21598 protein motif:CLUSTERS:PRK13351 protein motif:Cdd:COG0050 protein motif:TIGRFAMs:TIGR00484 protein motif:Pfam:PF00009.21"
/product="Tetracycline resistance protein tetM from transposon Tn916 [antibiotic resistance]"
/"ID=B40_02250
/protein_id="gnl|SC|B40_02250"""

/gene="tetM"
/codon_start=1
CDS 72713..73951
/color=2
/locus_tag="B40_02251"
/note="tetracycline efflux MFS transporter Tet40, protein motif:CLUSTERS:PRK10054 protein motif:TIGRFAMs:TIGR00880 protein motif:Pfam:PF07690.10"
/product="tetracycline efflux MFS transporter Tet40 [antibiotic resistance]"
/"ID=B40_02251
/protein_id="gnl|SC|B40_02251""""
/gene="tet40"
/codon_start=1
CDS complement(74094..74234)
/color=1
/locus_tag="B40_02252"
/note="Maff2 family protein  membrane protein  conjugal transfer protein, protein motif:Pfam:PF12750.1"
/product="Maff2 family protein  membrane protein  conjugal transfer protein"
/"ID=B40_02252
/protein_id="gnl|SC|B40_02252""""
/codon_start=1
CDS complement(74302..75051)
/color=4
/locus_tag="B40_02253"
/note="type IV secretion system protein TraG  VirD4 components, protein motif:CLUSTERS:PRK13850 protein motif:Cdd:COG3505 protein motif:TIGRFAMs:TIGR02759 protein motif:Pfam:PF12696.1"
/product="type IV secretion system protein TraG  VirD4 components [coupling protein]"
/"ID=B40_02253
/protein_id="gnl|SC|B40_02253""""
/gene="traG"
/codon_start=1
CDS complement(75228..75593)
/ID=B40_02254
/color=1
/locus_tag="B40_02254"
/note="hypothetical protein"
/product="hypothetical protein"
/protein_id="gnl|SC|B40_02254"

/codon_start=1
CDS complement(75672..75863)
/ID=B40_02255
/color=9
/locus_tag="B40_02255"
/note="transposon-encoded protein TnpW"
/product="Transposon-encoded protein TnpW"
/protein_id="gnl|SC|B40_02255"
/gene="tnpW"
/
/codon_start=1
CDS complement(75907..76770)
/color=10
/locus_tag="B40_02256"
/note="DNA replication protein dnaC ATP binding protein,
similar to AA sequence:UniProtKB:P0AEF0 protein
motif:CLUSTERS:PRK08116 protein motif:Cdd:COG1484 protein
motif:TIGRFAMs:TIGR00362 protein motif:Pfam:PF01695.11"
/product="DNA replication protein dnaC ATP binding
protein [replication]"
/"ID=B40_02256
/protein_id="gnl|SC|B40_02256""""
/gene="dnaC"
/codon_start=1
CDS complement(76767..77519)
/color=10
/locus_tag="B40_02257"
/note="phage replisome organizer putative N-terminal
region, protein motif:TIGRFAMs:TIGR01714 protein
motif:Pfam:PF09681.4"
/product="phage replisome organizer putative N-terminal
region [replication]"
/"ID=B40_02257
/protein_id="gnl|SC|B40_02257""""
/codon_start=1
CDS complement(77633..78031)
/ID=B40_02258
/color=1
/locus_tag="B40_02258"
/note="Cysteine-rich VLP domain-containing protein"
/product="Cysteine-rich VLP domain-containing protein"
/protein_id="gnl|SC|B40_02258"
/codon_start=1
CDS complement(78012..78188)

/ID=B40_02259
/color=1
/locus_tag="B40_02259"
/note="hypothetical protein"
/product="hypothetical protein"
/protein_id="gnl|SC|B40_02259"
/codon_start=1
CDS complement(78288..79838)
/color=4
/locus_tag="B40_02260"
/note="Plasmid mobilization system relaxase MobA/MobL family, similar to AA sequence:UniProtKB:P07112 protein motif:CLUSTERS:PRK13889 protein motif:TIGRFAMs:TIGR02768 protein motif:Pfam:PF03389.9"
/product="DNA strand transferase Plasmid mobilization system relaxase MobA/MobL family [relaxase]"
/"ID=B40_02260
/protein_id="gnl|SC|B40_02260"""
/gene="mobA"
/codon_start=1
CDS complement(79921..80217)
/color=1
/locus_tag="B40_02261"
/note="Protein of unknown function DUF3847, protein motif:Pfam:PF12958.1"
/product="Protein of unknown function DUF3847"
/"ID=B40_02261
/protein_id="gnl|SC|B40_02261"""
/codon_start=1
CDS complement(80157..80696)
/color=7
/locus_tag="B40_02262"
/note="RNA polymerase sigma factor Sigma-70 region 4, protein motif:TIGRFAMs:TIGR02937"
/product="RNA polymerase sigma factor Sigma-70 region 4 [transcriptional regulator]"
/"ID=B40_02262
/protein_id="gnl|SC|B40_02262"""
/codon_start=1
CDS complement(80693..81154)
/color=7
/locus_tag="B40_02263"
/note="RNA polymerase sigma factor Sigma-70 region 4, protein motif:TIGRFAMs:TIGR02937 protein

motif:Pfam:PF08281.6"
/product="RNA polymerase sigma factor  Sigma-70  region 4
[transcriptional regulator]"
/"ID=B40_02263
/protein_id="gnl|SC|B40_02263"""
/codon_start=1
CDS            complement(81165..81617)
/color=7
/locus_tag="B40_02264"
/note="RNA polymerase sigma factor  Sigma-24, protein
motif:CLUSTERS:PRK11924 protein motif:TIGRFAMs:TIGR02937"
/product="RNA polymerase sigma factor  Sigma-24
[transcriptional regulator]"
/"ID=B40_02264
/protein_id="gnl|SC|B40_02264"""
/codon_start=1
CDS            complement(81721..82134)
/color=7
/locus_tag="B40_02265"
/note="RNA polymerase sigma factor  Sigma-70  region 4,
protein motif:TIGRFAMs:TIGR02937 protein
motif:Pfam:PF08281.6"
/product="RNA polymerase sigma factor  Sigma-70  region 4
[transcriptional regulator]"
/"ID=B40_02265
/protein_id="gnl|SC|B40_02265"""
/codon_start=1
CDS            complement(82624..84279)
/color=8
/locus_tag="B40_02266"
/note="ABC-F family ATP-binding cassette domain-containing
protein, similar to AA sequence:UniProtKB:P63389 protein
motif:CLUSTERS:PRK11819 protein motif:Cdd:COG4555 protein
motif:TIGRFAMs:TIGR03719 protein motif:Pfam:PF00005.21"
/product="ABC-F family ATP-binding cassette
domain-containing protein [transporter]"
/"ID=B40_02266
/protein_id="gnl|SC|B40_02266"""
/codon_start=1
CDS            complement(84598..84951)
/ID=B40_02267
/color=9
/locus_tag="B40_02267"
/note="transposon-encoded TnpV family protein"

/product="Transposon-encoded TnpV family protein"
/protein_id="gnl|SC|B40_02267"
/gene="tnpV"
/codon_start=1
CDS complement(84978..86858)
/color=9
/locus_tag="B40_02268"
/note="Serine recombinase, protein motif:Pfam:PF07508.7"
/product="Serine recombinase [recombinase]"
/"ID=B40_02268
/protein_id="gnl|SC|B40_02268"""
/codon_start=1
CDS complement(86932..87906)
/color=4
/locus_tag="B40_02269"
/note="type IV secretion system protein TraG VirD4 components, protein motif:CLUSTERS:PRK13876 protein motif:Cdd:COG3505 protein motif:Pfam:PF02534.8"
/product="type IV secretion system protein TraG VirD4 components [coupling protein]"
/"ID=B40_02269
/protein_id="gnl|SC|B40_02269"""
/gene="traG"
/codon_start=1
CDS complement(87903..88376)
/color=1
/locus_tag="B40_02270"
/note="uncharacterised PcfB family protein, protein motif:Pfam:PF12687.1"
/product="uncharacterised PcfB family protein"
/"ID=B40_02270
/protein_id="gnl|SC|B40_02270"""
/codon_start=1
CDS complement(88413..89357)
/ID=B40_02271
/color=7
/locus_tag="B40_02271"
/note="helix-turn-helix domain-containing protein"
/product="helix-turn-helix domain-containing protein [transcriptional regulator]"
/protein_id="gnl|SC|B40_02271"
/codon_start=1
CDS complement(89486..89704)
/ID=B40_02272

/color=1
/locus_tag="B40_02272"
/note="hypothetical protein"
/product="hypothetical protein"
/protein_id="gnl|SC|B40_02272"
/codon_start=1
CDS 89908..91311
/color=9
/locus_tag="B40_02273"
/note="Serine recombinase, protein motif:Pfam:PF07508.7"
/product="Serine recombinase [recombinase]"
/"ID=B40_02273
/protein_id="gnl|SC|B40_02273"""
/codon_start=1
CDS 91570..92433
/ID=B40_02274
/color=1
/locus_tag="B40_02274"
/note="hypothetical protein membrane protein putative conjugal transfer protein TrbL/VirB6"
/product="hypothetical protein membrane protein putative conjugal transfer protein TrbL/VirB6"
/protein_id="gnl|SC|B40_02274"
/codon_start=1
CDS 92743..93972
/color=9
/locus_tag="B40_02275"
/note="IS91 family transposase, protein motif:Pfam:PF04986.7"
/product="IS91 family transposase [DDE recombinase]"
/"ID=B40_02275
/protein_id="gnl|SC|B40_02275"""
/codon_start=1
CDS 94026..94211
/ID=B40_02276
/color=1
/locus_tag="B40_02276"
/note="hypothetical protein"
/product="hypothetical protein"
/protein_id="gnl|SC|B40_02276"
/codon_start=1
CDS 94168..94635
/ID=B40_02277
/color=1

/locus_tag="B40_02277"
/note="hypothetical protein"
/product="hypothetical protein"
/protein_id="gnl|SC|B40_02277"
/codon_start=1
CDS 94942..97353
/ID=B40_02278
/color=4
/note="protein motif:Cdd:COG3451 protein
motif:TIGRFAMs:TIGR02746 protein
motif:Pfam:PF12846.1locus_tag=B40_02278product=Conjugal"
/product="Type IV secretory system protein TraC VirB4
components AAA domain [ATPase mating pair formation]"
/gene="traC"
/protein_id="gnl|SC|B40_02278"
/codon_start=1
CDS 97385..98623
/ID=B40_02279
/color=4
/locus_tag="B40_02279"
/note="NlpC/P60 family protein"
/product="NlpC/P60 family protein [pepidase]"
/protein_id="gnl|SC|B40_02279"
/codon_start=1
CDS 98636..99331
/color=4
/locus_tag="B40_02280"
/note="NlpC/P60 family protein, similar to AA
sequence:UniProtKB:P21171 protein motif:CLUSTERS:PRK13914
protein motif:Cdd:COG3584 protein motif:Pfam:PF00877.13"
/product="NlpC/P60 family protein [pepidase]"
/"ID=B40_02280
/protein_id="gnl|SC|B40_02280"""
/codon_start=1
CDS 99355..99618
/ID=B40_02281
/color=1
/locus_tag="B40_02281"
/note="hypothetical protein DUF4315 containing protein
conjugative transposon"
/product="hypothetical protein DUF4315 containing protein
conjugative transposon protein"
/protein_id="gnl|SC|B40_02281"
/codon_start=1

CDS 99608..100375
/ID=B40_02282
/color=1
/locus_tag="B40_02282"
/note="hypothetical protein"
/product="hypothetical protein  DUF4366 containing protein
cell surface protein"
/protein_id="gnl|SC|B40_02282"
/codon_start=1
CDS 100372..102462
/color=10
/locus_tag="B40_02283"
/eC_number=5.99.1.2
/note="DNA topoisomerase III, similar to AA
sequence:UniProtKB:P14294 protein motif:CLUSTERS:PRK07726
protein motif:Cdd:COG0550 protein motif:TIGRFAMs:TIGR01056
protein motif:Pfam:PF01131.14"
/product="DNA topoisomerase III [replication]"
/"ID=B40_02283
/protein_id="gnl|SC|B40_02283""""
/codon_start=1
CDS 102672..102827
/ID=B40_02284
/color=1
/locus_tag="B40_02284"
/note="hypothetical protein"
/product="hypothetical protein"
/protein_id="gnl|SC|B40_02284"
/codon_start=1
CDS 102838..103314
/ID=B40_02285
/color=1
/locus_tag="B40_02285"
/note="hypothetical protein"
/product="hypothetical protein"
/protein_id="gnl|SC|B40_02285"
/codon_start=1
CDS 103364..103567
/ID=B40_02286
/color=1
/locus_tag="B40_02286"
/note="hypothetical protein"
/product="hypothetical protein"
/protein_id="gnl|SC|B40_02286"

/codon_start=1
CDS 103670..104269
/color=7
/locus_tag="B40_02287"
/note="helix-turn-helix , protein motif:CLUSTERS:PRK09706
protein motif:TIGRFAMs:TIGR03070 protein
motif:Pfam:PF01381.16"
/product="helix-turn-helix [transcriptional regulator]"
/"ID=B40_02287
/protein_id="gnl|SC|B40_02287"""
/codon_start=1
CDS 104403..105533
/ID=B40_02288
/color=4
/locus_tag="B40_02288"
/note="anti-restriction modification"
/product="DUF4316 domain-containing protein
anti-trestriction protein ArdA [anti-restriction
modification]"
/gene="ardA"
/protein_id="gnl|SC|B40_02288"
/codon_start=1
CDS 105563..109063
/ID=B40_02289
/color=4
/locus_tag="B40_02289"
/note="anti-restriction modification"
/product="DUF4316 domain-containing protein
anti-trestriction protein [anti-restriction modification]"
/protein_id="gnl|SC|B40_02289"
/codon_start=1
CDS 109067..109267
/ID=B40_02290
/color=4
/locus_tag="B40_02290"
/note="putative tranposon-transfer assisting protein
TTRAP"
/product="Putative tranposon-transfer assisting protein
TTRAP"
/protein_id="gnl|SC|B40_02290"
/codon_start=1
ncRNA 109403..109573
/ID=B40_02291
/color=1

/locus_tag="B40_02291"
/product="group-II-D1D4-3"
ncRNA 109598..109690
/ID=B40_02292
/color=1
/locus_tag="B40_02292"
/product="group-II-D1D4-1"
CDS 109796..111076
/color=8
/locus_tag="B40_02293"
/note="Group II intron-encoded protein LtrA, similar to AA sequence:UniProtKB:P0A3U1 protein motif:Cdd:COG3344 protein motif:Pfam:PF00078.21"
/product="Group II intron-encoded protein LtrA [reverse transcriptase]"
/"ID=B40_02293
/protein_id="gnl|SC|B40_02293"""
/gene="ltrA"
/codon_start=1
CDS complement(111161..112327)
/color=8
/locus_tag="B40_02294"
/eC_number=2.7.11.1
/note="Non-specific serine/threonine protein kinase, similar to AA sequence:UniProtKB:O34507 protein motif:CLUSTERS:PRK13184 protein motif:Cdd:COG3349 protein motif:TIGRFAMs:TIGR03903 protein motif:Pfam:PF00069.19"
/product="Non-specific serine/threonine protein kinase [kinase]"
/"ID=B40_02294
/protein_id="gnl|SC|B40_02294"""
/codon_start=1
ncRNA 112529..112722
/ID=B40_02295
/color=1
/locus_tag="B40_02295"
/product="group-II-D1D4-3"
ncRNA 112762..112833
/ID=B40_02296
/color=1
/locus_tag="B40_02296"
/product="group-II-D1D4-1"
CDS 112881..114272
/color=8

/locus_tag="B40_02297"
/note="Group II intron-encoded protein LtrA, similar to AA sequence:UniProtKB:P0A3U1 protein motif:Cdd:COG3344 protein motif:Pfam:PF00078.21"
/product="Group II intron-encoded protein LtrA [reverse transcriptase]"
/"ID=B40_02297
/protein_id="gnl|SC|B40_02297"""
/gene="ltrA"
/codon_start=1
CDS complement(114448..114819)
/ID=B40_02298
/color=1
/locus_tag="B40_02298"
/note="Cysteine-rich VLP domain-containing protein"
/product="Cysteine-rich VLP domain-containing protein"
/protein_id="gnl|SC|B40_02298"
/codon_start=1
CDS complement(114816..116150)
/color=4
/locus_tag="B40_02299"
/note="Relaxase/Mobilisation nuclease domain, protein motif:Pfam:PF03432.8"
/product="Relaxase/Mobilisation nuclease domain [relaxase]"
/"ID=B40_02299
/protein_id="gnl|SC|B40_02299"""
/codon_start=1
CDS complement(116111..116440)
/color=4
/locus_tag="B40_02300"
/note="Plasmid mobilization relaxosome protein MobC, protein motif:Pfam:PF05713.5"
/product="Plasmid mobilization relaxosome protein MobC [relaxosome]"
/"ID=B40_02300
/protein_id="gnl|SC|B40_02300"""
/gene="mobC"
/codon_start=1
CDS 116818..117654
/color=9
/locus_tag="B40_02301"
/note="Tyrosine recombinase, similar to AA sequence:UniProtKB:P0A2P6 protein motif:CLUSTERS:PRK00236

protein motif:Cdd:COG4974 protein motif:TIGRFAMs:TIGR02225
protein motif:Pfam:PF00589.16"
/product="Tyrosine recombinase [recombinase]"
/"ID=B40_02301
/protein_id="gnl|SC|B40_02301"""
/codon_start=1
CDS 118304..119491
/color=8
/locus_tag="B40_02302"
/eC_number=1.1.1.22
/note="UDP-glucose 6-dehydrogenase, similar to AA
sequence:UniProtKB:P76373 protein motif:CLUSTERS:PRK15057
protein motif:TIGRFAMs:TIGR03026 protein
motif:Pfam:PF03721.8"
/product="UDP-glucose 6-dehydrogenase [dehydrogenase]"
/"ID=B40_02302
/protein_id="gnl|SC|B40_02302"""
/gene="ugd"
/codon_start=1
CDS 119513..120805
/color=8
/locus_tag="B40_02303"
/note="N-glycosyltransferase, protein
motif:CLUSTERS:PRK11204 protein motif:Cdd:COG3858 protein
motif:TIGRFAMs:TIGR04242 protein motif:Pfam:PF03142.9"
/product="N-glycosyltransferase [glycosyltransferase]"
/"ID=B40_02303
/protein_id="gnl|SC|B40_02303"""
/codon_start=1
CDS 120783..121820
/color=8
/locus_tag="B40_02304"
/eC_number=3.2.1.4
/note="Beta-mannanase, similar to AA
sequence:UniProtKB:P16218, protein motif:Cdd:COG4124
protein motif:Pfam:PF02156.9"
/product="Beta-mannanase [hydrolase]"
/"ID=B40_02304
/protein_id="gnl|SC|B40_02304"""
/codon_start=1
CDS 121938..122003
/ID=B40_02305
/color=1
/locus_tag="B40_02305"

/note="hypothetical protein"
/product="hypothetical protein"
/protein_id="gnl|SC|B40_02305"
/codon_start=1
BASE COUNT 30789 a 29453 c 31666 g 30095 t
ORIGIN
1 ctccatactg atatttttga agatcaaatt tgttgcattg gtaaatatat ggtaacaaaa
61 tattgcaaat ataaaaatgt tactttataa ttgctattgt aacaaatgat tgcaatgatt
121 acaagcgttg tataaataat gataaacgga aagagaggta gttatgaaaa acagaacggt
181 aacagcagtg gtaggagaat cacgattgat ggacacagag gaattaagag cctacaccaa
241 cttggggcgt aataatgcca tgaaacttgg tgaggagatt ggcgcaaaag tccagattgg
301 acgcagggta ctatgggaca ggcgaaagat tgacgagtat ttcaattcac tgacgggggt
361 aaaataggga atgcaaaaaa ataagcataa cagagtacat atagggaatc ggcccggcaa
421 ggcagattat ccaatagcaa gtctcctgcc tgtaggcaag gaaaacgcaa tatccactgc
481 tgatctagtc aagctgagtg gatgcagttc ctcacggaaa ctacagcagc atatagccta
541 tgaaaggaac cacggtgcaa ttatctgcag tggttcaggg aatggctact ggaaaccaaa
601 ggacaggcag gagatagtag agttttgccg aataatggat gcaagggcaa gaaacacatt
661 tgcggcaacg agaagtgcta aacaggcact aaaagaacca gaggggcagc aggattttat
721 aagataacaa ggggcgggtg aataatggca gaggcaatga aggattgggc tttaaagtat
781 gctgaaatcg ggctggcagt gttcccgttg aagccaaagg ataagcgacc ggcaacagag
841 aatggatgta aggcagctac tataaataaa caacagatag aagcatggtg ggacagatac
901 ccggattata atatcggaat agcaacaggg agcgtttcag gtggcttggt ggtgattgac
961 ttggatgtgg acgaggaaaa gggcgttaat ggatatgaga gtctgacagc ctggcagaag
1021 gaaaacggag aattgccgga aacctggaca agcatcactg gtcgcggagg atatcactac
1081 atatataaag atggagccac caaccgaaac agagcaggac tttatgatgg cgttgacatc
1141 cggggagaag gtggctacat agtagcacca cccagcatac acccaaatgg tcgtcagtat
1201 gaatgggagc aggggccggg ggattgtgaa atcgcccagg cagacaatag agttatgaac
1261 ttcattatgg ggcctgcacc agaggactgg gagaaacaga ctttccagga gccggaagtg
1321 ataccaaagg gggagaggac aaacagtatg atccggctga ttgggagcct aaaggcaaag
1381 ggccttgatg atggagccat aaaggcagca gttagggccg aaaatgcaaa gaagtgtatc
1441 ccgccgatca cggatcagga acttgagaag acagtattcc cggcgttgaa acggggatgg
1501 aaaactgaga aaccttatac tgcgatacat gataatggaa aattccggca gcagaaaaac
1561 tatagtcttg aaatgttaac gatgaaagag gttgaagaaa aaagtcctga gtggcttatt
1621 acagactaca taccaagata tcagatcaca acgcttgccg gggatggggg cgcaggaaag
1681 acaactatat ggtgcgacat agcggcagcg gttagtagtg ggaatccttg ttttcttgag
1741 gccggaatcc cggaagactt ttccaaggca gatccgggaa cagttctttt cttttcatca
1801 gaagattcgg cagagtatac gttgcgtagc aggctgagaa aggctggagc cgcattggag
1861 aatatacatt ccatcagcct gaaagatgaa cggttttctg atgtaaagtt cgatagcccg
1921 tttcttgaaa gactgattga acattatcgt ccaaagctgg taatatttga tccgattcaa
1981 agttttatac cgccagacat ccagatggga caaagaaatg ccatgagatc atgcttgaat
2041 cctttgattg gtttgggcga aaaatacggg actactttcc ttgtgattgt tcatgcaaat
2101 aaacagagcg gtgtttatgg tcggaaaagg atagcagaca gcgctgatat ctgggatata
2161 tccagaagtg tattaatggt gggagatact ccggaaaagg gtgttcgcta tctgtcacat
2221 gagaagtgta attatggcat gacgggtgat acggtacttt tctccattga aaatggcagt

2281 attataaaca aaggctatac agagaagaaa gaccgggagt ttgtaagcga gaattcattt
2341 attgtccagc agaggccgca aaaggaagag gcgaaagagt ttatattgtc attcctgagt
2401 gatggtgaaa aagaggttgc ggaactggat gagatggcag cggcgcaaag tataaccaag
2461 aattcattaa aaaatgccaa agccgaatta aggaaagaag gtaagataaa ataccgtaat
2521 tttggctatg ggagtgaccg taaatatttc atttctctta tacccagtga aaaagtcaac
2581 gagtaaataa aaaacatagt atttataagg gtttgagata ctagaggggg gtcaacgagt
2641 aaagcgagta aaggctattt actggagggg ggcctctagt aaatggagcc cttgtaaaca
2701 gggcgtttgc cctttactgg ttgaccggtg gcccgggtat atgagaatca acgagtaagg
2761 agaggagcca acgagtaagt gagcgtgaac atattagcag attttaagga gaatggtatt
2821 gataaaaatg agccacatat tgtgctgtat acagataatg agtatgaggc aggcatgatt
2881 ataaaggcga aactggagga gagaggctgt aaggttgaat ctctgattgt ggttgaagga
2941 aaatggacgc tggtgcaact tcatgatatg gctaattatg gtactggaat agaaaaggta
3001 catccccggc tgttgtatgt gtctggcgac atgttgcagt atctgaacgg attaagaaac
3061 aggccggaag aagtggcaca actaaaaaat gaaatcagga gaagggcaaa tggacagaaa
3121 ggaaacaggg aagcgcagta ggaataaagg taaagcagga gaacgcgagc tggcaggagt
3181 attaagggat tatggatatg attgcaggag agggcagcag ttttgcggca gcaatgggga
3241 cgctgatgtc gttggtcttc ctggaataca tatcgagtgt aagcggacgg agcggttaag
3301 cctgtatgat gcaatgggac aggctgaaag ggattcacgt agagaagaga taccagtagt
3361 tatgcatcgt agaaaccatt gtgaatggct tgcgatcctt cctcttgata acttcatgga
3421 gatttacaga gagtgggagg caggccagat gcaaaaataa aagagcttgc cacaaaggac
3481 aagcccacag ccttaaaaac ctaacattta ttgtaaagga gtgggctgta tatgtcaaat
3541 gaacaccggg tgaaccccgg caattttcaa aacgaactac ggaggaacaa cggggaattt
3601 caaaataagc aaagaggaag cagtgagaac tttccagttg aaccattggt gaacaacaca
3661 aaattacagg ttgaaccgag agcgaaccgt gaaaaatttt ctgaaatgac tgggaggggc
3721 ggtctgaaaa gttccgataa aattcatcag accgcatgcc ctccaatcct cacacaaaat
3781 tccgaatacg aacaaaaaag tggtacatct gcggagacga tggaagatcc ccttttccga
3841 tccaatgagc agttagtcgc gttgatccgg gccagtatag atgaagctga caatatgctg
3901 cagctatggc agcagaacca gcggtttgta aaaatgatgg ctgtgaagtt tacgtgccgt
3961 gctgaattag atgatcttat gcaggaggga tatattgctt tgtgtgaagc tgtccgacac
4021 tatgacccgg agaagggagt tgtatttatc acttatgctg cgttctggat cagacaaggt
4081 atgcaacgat atattgagaa ttgtggtaat gtggtgcgta ttcctacaca tgcccgggaa
4141 tggatgcaca aatataagag gatggaagca gagtaccgga aatattacgg caccactcca
4201 acggatcagg tattgtgtgc gctcctgggg atcagccagg agaaactaaa gtccatcaga
4261 gaaagtatag gaatggggca gatttgtagc ctgaatgaac ctttagggga tgaagaaggc
4321 gaactgactg tagaggactc tgttggatca ggtgaggagc tggaaggggа tataatcggc
4381 cggctggact accaaggcat gaagaaacag ctctgggaag cggtggacag cctgccaggg
4441 attcaatcgg atgtaatacg ctgccggttc ctggatagca aaactctaca ggaaatagga
4501 gagaagtata atctttcacg ggaggtagca cggagttatc aggacaaagc tatgcgggaa
4561 ttaagaaagc ccagcaggag ccggaagttc agagggtact atgaagaata tattgcggct
4621 gcatcaattc atcatatagg gcttgaatct ttccggcgta catggacaag tgaggttgag
4681 agggatgcca ttggaatatg gtgacgttcc gaacgttcgg aaaccgacga atgcgaacgg
4741 tatttatttg ttgggatctg gcagcaggct acattgtgct gaatcccccc taaaataaaa
4801 tgcaggaaag cctatagtag accggttgtg gggattcaga taatgcacgg gccgcatata
4861 tgggggtgtg gtattgaaga caaagagaag ccccccgacc cttccagcac cgatttaaac

4921 gtgataggaa ccggagtggg taactgtttc caatggtgca gcactctgaa aatatttttt
4981 ccatctcaat ctaagaataa actcgaatac tatttgaagt gttgcaataa tagattgggg
5041 acaggatcaa gacattcgag atcccctaaa agggacgttg caatgttgta tacttagtat
5101 ttcgtatatt tacaaaacaa aggagcggag aatgagaagc tatattgatg taaaggattg
5161 cataaagctg ggcgcgagga acatgactgc cgccatagca aagataaacg gaaaagagta
5221 caacatacag ttcacgaagc agaggacagg atttggtgaa aagctgtttc ttgtttgccc
5281 gaagtgtggc agcaggcgta caaagctata tctgtatgga gatgggcttc tgtgccggga
5341 atgctatccg tttccggttt atgacagaat taagaatgta tctatcggca gctataacta
5401 tatttcctac aggatgctcg ggcttgcaag gcgtgaacga attccgatta agttcccgtt
5461 ccattatctt gaatatgaga aaccaaaata taagcaattt gataaatggc acatgacaat
5521 tacaaagcta caggcattgg agaacatgag gaaccaggct atatttttcg gtaagaagta
5581 ttcacgggcg acaattaaca gtatctttga agagcggaac ctatatctgt atgtctgtga
5641 tctgtacgat ctggaattgt atttctatga ttgggatgtt ggtttcttga aatatccggg
5701 aaatgagcca gaggtcaagc caaccggtat aatgagtcag gcaagtaaat accagagagc
5761 ggtgttggga cggcatcgtt aagcgggata gaggtaattc ggacatggtt acgttcatgt
5821 ggaaataaat cacttttagg tgaggttttg gtaaggtatg tggaagtgat catcgttatc
5881 acataatatg agctgatatg gtatggctat gtctgcataa aggaggaaaa catgagatta
5941 aatgtgaata gatataaaga actattggaa agtaagaatt tggatgaact ggacattgag
6001 aggaaaacag gactttcgat gagtacaatt aactggatat ttgaaaatga gtatctggag
6061 ataagcacgc tggagcggct ggctgatgtt gtaaagtgtg gtacaaagga gattgctctg
6121 ccagatcata acaacattga gaatgtcatt gaatggcagc gagacagtaa gaccgctacg
6181 gtgagcctaa cacagggcag aaccattaca agagtgatga agctggctga gagccgcccg
6241 gaggaatgcc ggatcattgc agagaataaa gatggcagca tttgtgccag ggttccggta
6301 ggatggataa gaataaaccc gggaatgaac ttgaatgagg aacagagaga aaaacgggct
6361 gatcggatgc gctgcaatat tttgaacaac gattatagtc gtggtgaaat gggataaaat
6421 caatttaaat gcctttaggg tagaaatata aggctaaagg tttaaaatgg ctttttaata
6481 agaattgcat ggaatagtgt tggacattga taatttgcat attaggggga tatgaaacag
6541 gctgttatat acttgataga acagcagcgg gatgcttctg cttttatatc ttaaaaatat
6601 tcttcatttt ctgagtccct aaaaggagta tcttgacaaa tatgagattc taaaaaggtt
6661 attccgacat ttccgacgcc ctaaaaggga aaataaaacc gccctcaatc ctgatggtcg
6721 ggggcggcag tgggttaaat attcaattgt ccgaagtcga ttgacaggtc ttcatagatg
6781 cctgctttta cggaatcgga gaaagtatag tcccctgtat cctcagattc aaaattataa
6841 accaaaatac ggttcttctc aggatcgact atccagtatt cccgtacacc ggcagtacgg
6901 tatttgaata atttagttaa atagtctatg cgcttactgc ttggtgatac aatttcaata
6961 acccagtcag gcgctcccgt gcaaccttta tccgttaatt tgtcagggtc acagattacg
7021 ctgatatccg gttctacgta tttcgtatcg tcttcattga gaaatacggc aaatggggca
7081 ggaaacactt cacaaggtcc acctttagag tggatatagt taccgattgt tctatcaagg
7141 tgatgtaata ttctttggtg ggtagtactt ggtggggcca tataatagat ctgcccgtcg
7201 atcagttcgg cacgtacccc atcaggcagg gcatagatat cttcgatggt gtattctttc
7261 ttctggtttg gtaatgacat gataacacat cctctcatga aatagtattg acggagtggc
7321 agtggttatt cagagatttc cgaggcccta aaaaggaatc tggtttcgct cacttttgag
7381 ctttattcag tatcgcgttc cattcgttca tcaatagcct ttttaatgta accgttgact
7441 gattcccctg tcatatctgc atgttggcgt actgtttctt tgtaaccctt tgggacaacg
7501 agattaattc tttctctatt ttcagactgc cacttatttt tatagtctgt ttttcctgcc

7561 aaaagtatca ccttctttca tggggggcgc tctgataatg gagacacctt ttctattatg
7621 gttagattat aacattaatt tcaatcgtgc acaagataca aaatagccaa atcttgcaca
7681 cgatatttgt gtataatgtc aattgaaaac atatcgtgta cacgatataa tataatcaag
7741 ataaaggtaa ggcaaacgaa cagcccgaaa agctgtgaag ctgaaaacca tatacctgtg
7801 agtgaacaat agagggatca atagccagga agatgattca gccggatgcc agaccggtat
7861 caacaataaa taaaagcaac cggggaagcc gaaagccccc acaaactttg agccatatac
7921 ttgtgaacgg tctggagatg cagtaatagt caagaagatg ctcaagggat tgcactggaa
7981 aggagatttt acaatgacag agagaatgat tgtcaaccgc gtaagaaagc tgaaagagct
8041 ggaagcgcag cagaaggaga ttgaggaaca gattgaggca ctgaaagcgg agatcaaagc
8101 agacatgcag aggaaggatc tggaggagca gaaggccggg gattatatgg taaggttcac
8161 aacggttatc tcaaacagat ttgatggtaa agcattcaaa gcggatcatg caaagctgta
8221 tgaccagtac atcaggacaa cggagagcag aagattctcc atcgcataaa aaatagccct
8281 tgccagtgct gcgaacactg acaggggcgg catagccgga gctacactat aaagcaaata
8341 catagtagca taatcccggt gggaagggca agaacaaatt gaatttggat aatttagaga
8401 aacttataaa ataccatcca taccatatct gtacttttgc tgactttgca aatgtaacgc
8461 aggatttgtt agaagtggca ttaaagggag aggaggagct tgaaccggta gaagttagga
8521 atatttcgga atatgtacaa gttccgtatc gggtcttgac ctgtaaaaag atgattatgt
8581 taagcaagga taggtaccgc caccgtatta tgtttgaaga gctctacgaa aagctatttg
8641 aaatatggga agcagcggaa aatgggagca aagaagccgc cagctataag cgatataatt
8701 ataagcattt ggtaacactc gtggcagact ttcaatatcg gggagcagtt acttattgtc
8761 gttatttggg agtcaaagaa atgatggaac agtacctgtt gtttatacga tgcgaaatgc
8821 gaaaaccaag aggaagggag attccgacat gagtaacgaa gaatataggc ggaaaagacg
8881 atataaagcc aaaggggcaa acgatgttgt tgctgatctt gatgaaagaa ggtgctgagt
8941 atgcaaaatg aaggatatag gcaatgcata gtacaaatgc tttatctaat ggataaggaa
9001 gaagataacg aggttctgga agagatatat accattgcac gctaccatta tattaaaatg
9061 gctcctgaca gaaggaagca aattgataag tgtagaggta tagtattcag gatcatagcc
9121 ggaatggatg aagagcgcgt caggcaatta tataacgtat tagttggaca ttacagattt
9181 tcgggggagc ggtgcggtga catcggcccc ctgggaagca gccgccataa ataaaaatat
9241 gattgataag ttagtgggag gattaattta tggttgaata tataaaactt aaacccggtg
9301 atgagctgtg gaaagctgaa atccttaatg aacaaacact tttcggaatg cttaccgatg
9361 agcagttttc taaaatcgaa gatgggaggt gcatggaatt aaagggagtc tgcttcatgc
9421 agaggccgcc attttacgga atatcccctc aaatagacta tgaaaagaac acgccctata
9481 tagaattgat tgagagcttt gatgtaaatt atcttatgca ggaaaggggg tgcgaagctg
9541 tatttaacag cattgattat gctgaccaga ttgtagacct acataaaatg attgggcggt
9601 ttagcaggga catagaaatt gttaagaaat tgatggacag gaggggcact acatggcgaa
9661 gctgacacca attaaggcga tcagagcaaa gtgcctggat tgctgcaatg gtcaaatgaa
9721 ggaagtcagg ctttgcactg ttgagaattg cgcgttacac gagtataggg acggacacag
9781 accgaaaggc gaagaagtta ccatcggaga tgttttcgct gaaaaatcgt gagatagggc
9841 aacaattttt ttcaagatta atgcattggg gctgttcaga ccttaaggaa agaaggtgaa
9901 ttctatgaaa gccgatatat tggaacaaat gtaccggtac tatgtacaaa tgccgcgcta
9961 taaggaactg cgaaagatat atgactatga cattcccaag gcccggcgcg atgatctgga
10021 aaacagcatg ggagcagaga actattcgaa agcccaggag atggccttct ctatatctga
10081 tcaggctgaa cgggtgggat tcatacttgg attccagatc ggatcaaagc tcatgtgcga
10141 gtctatagga gaatgggagg tggacgaatg accggaaaag caaaaacccc aagggtgcgg

10201 caacacccaa ggggcaaaaa ataagcataa cccaatacac acagtacggg ctatattgat
10261 tatataccgt catatagccc atttgcaaga caagattcac ctgctgccag gcagcagtga
10321 gagaaaggac ggttatatgt caagaattgg aaatatatac acagaatcat acagcaaatc
10381 atcggctgcc atatcagcgg atggcaaatt gtctgaaacg atccaggaag aaatcgtgga
10441 caggatcggg gagccagcca gcatgacgaa ggaatctgta gcagatttaa tttatgccgg
10501 gagcagttat ggacagaaag caggatttga gagcggtttt aactatgcaa tagccctgat
10561 atttgaaagc ctaatgagtt aggcggtggt tatatggcag tcgataaaag agtcaggaaa
10621 ttaccaaaag gaatacagca aaggtatact gactatgagg ggcgtttcat gtacaaggga
10681 gaaagatatc tggtacatgg tgcgacaatt acggaaacac aaaagaaaat gactgactta
10741 aagtacaaac tgcagcacgg cctgtttgta gaaagtagca aaaccaccct ggatgaatgg
10801 tttgaaacct ggctgaatgt ttctattaaa aactctgatt atttcaaaaa agcccggaaa
10861 tacgggcaaa tcacgggata aggacaccga atttactacc catttactac attcggaggg
10921 agccttgaca cgctgtcctt tcccatgaag cccactatcc caaacagcgc acatcgaaaa
10981 atcgaatacg acaccgcaga cggcggcgtt ctcttttccc tttggggaga tcaggacgcc
11041 gctttttgc gcccaaatag aggaaggagc gaccgcctat gtactttacg tccggcagcg
11101 accgggcctt tgaacagctc atgcagacaa ggccaggcgc agatcatttt gacagcggcg
11161 aggccggagc gccggaggac tgcggcactt gccgatttta ccgcccacac tggaagtatc
11221 agttttgcat ctacgcagaa tgtccttacc agccgggcaa gctcaccgtc tttgacacgg
11281 tgaaatttca agtgaaagga gtgtgcgaca agatggcagt ttttcgagtg gagaaaaacc
11341 gaggatacac ggtgatgtcc aaccaccacc tgcggaataa ggacttgtcc ctgaaagcca
11401 agggcctgtt atcgcaaatg ctctctctgc cggagggctg ggactacacg ctaaagggcc
11461 tgtctttgat aaatcgggaa agcatcgacg ccatacggac ggcggtgtgg gagctggaac
11521 gggccggata tatcacccga cagcagaacc gggacggcaa gggaaagatg gcagatatgc
11581 tctataccat ctacgagcag ccgcagaccg aagcatcggt gttggaacag ccgatattgg
11641 aaaacccggt gttggaaaat ccaacatcgg ataacccgac accggaaaat ccaacgcaaa
11701 taaataaaga tagatcaagt aaagagaaat caaatacaga tggacgaatt accgattcca
11761 ttcctatcct ttcccctccc tctcctttgg agaaagcagc ggctgcgccg ccggaacgga
11821 aaggaacggg agcgaagtca cagagcgccg tagcggttta tcgggaaatc atcaaggata
11881 acatcgagta tgaacacctt tgccagcaca ccaaggggat tgaccgggaa ctgctggacg
11941 agatcgtgga cttgctggtg gaaaccgtat gcagcgcccg aacgaccatc cgcattgccg
12001 gggacgatta ccccgctgag ctggtgaaat ccaagttgat gaagctgaac agctcccaca
12061 tggagtttgt ctttgactgt atcagcaaaa acaccacgga aatccgaaat atcaagaaat
12121 acctgctggc ggttctgttc aacgccccaa gcaccatcaa cggctattac acggcactgg
12181 tggcccacga tatgaacacc ggcaaaatct gaaaaaggag gacagcccta tgaaacaggg
12241 aactttgatt tttgacgagc aggcagaccg ctatgacatc cgttttgacc tagcggacta
12301 ctacggcggc ctgcactgcg gagaaacctt cgatgtactg gtaggcggca gatggaggcc
12361 cacccgcata gaaatggctg aaaactggta tctggtgggt atccgcaccg acgacctctc
12421 cggcctgcgg gtgcggatct aagccgtgcc gcctgcctgc tttgacttcc aaggagggat
12481 agcagttgca ggaagaaatc gaacagaaaa ccatcgcgtt agcggttaag acgggcaagc
12541 tcaccggcca ggtattgcag gcggccatga agaaatttct ggccgcccgg cagaaaggga
12601 caggaaaggc acatcacgga aaacagagct tgcggcagct caaaaaggac ggttccgccc
12661 tgtccaacat tgaaatcacc gatgccaaca tcggcctgtt tcggccctgc gccaagaaat
12721 acggaattga tttcaccctg cggaaagacc gcaccaccca tccgccccgg tatatcgtga
12781 ttttcaaatc caagcaggcg gacaatctgg aacaggcgtt taaggagttt acggccaaga

12841 agctcaaaca gcaggaacgc ccctccatcc gaaaggccct gtcggctatg aagcaaaaga
12901 cggcggccag aaaccgccag caggccaagg agaaaatcaa ggaaaggggg ctgtctctat
12961 gaggccggaa gtgaaaaagc tgataatcgc aaacctgccc tatctgctgt tcgtttacct
13021 gtttggcaag ctgggacaga cctaccggct ggcggctggg gcggatgtct cagaaaagct
13081 gctgcacctt gcggacggct tttcccttgc ctttgaaaac gccgccccca gcttccacct
13141 gttcgatctg gcggtgggtg tggccggtgc ggtggcattg cgtctgatgg tgtactgcaa
13201 gagcaagaac gccaagaaat accgccgagg cgtagaatac ggcagtgccc ggtggggcgg
13261 ccccaaagat atagccccgt acatcgaccc ggtgtttgat aacaacattc tcctgaccca
13321 gacggaacgc ctcaccatga acaaccgccc caaagacccc aagaccgccc ggaacaaaaa
13381 cgtgctggtc atcggcggtt ccggcagcgg caagacgcga ttttttgtga agccgaacct
13441 gatgcagtgc gtgtccaagg attatccggc ctctttcatt atcaccgatc ccaaaggggg
13501 cctgatcggg gaggtgggcc agctccttgt gcgaagcggc taccgggtca aagtgctgaa
13561 taccatcaac ttttccaaga gcatgcggta caacccattc cgctacatta attcggaaaa
13621 agacatctta aagctggtca acaccctgat ctgcaacacc aagggggagg gcgaaaaaag
13681 cgccgaggat ttttggatca agtcggaacg tctgttgtat tcggctctca ttggctacat
13741 ctggtacgaa gcgccggacg atgaaatgaa tttcacgacg ctgcttgaaa tgatcaatgc
13801 gtcggaggcc cgcgaggacg acccggaatt tcaatccccg gtggatgcca tgtttgaacg
13861 gctggaagaa aaagacccgg aacactttgc ggtgcgccag tacaaaaaat tccttctgtc
13921 ggcaggcaaa acccgcagtt ccattttgat tagctgcgga gccagattag cgccatttga
13981 catccgggaa gtgcgggagc tgatggagga cgacgagctg gagcttgaca ccatcgggga
14041 tgaaaaaacg gccctgttcc tcattatgag cgacacggac acgacgttta acttcattct
14101 taccatggtg cagagccagc ttatcaacct tttgtgcgac cgggccgatg acaaatacgg
14161 cgggcggctg cccgtccatg tgcggatgat tctggacgag tttgccaaca tcggtcagat
14221 tcccaacttt gacaagctga tcgccaccat ccgaagccgg gaaatctcgg cttctatcat
14281 tttgcagagc cagtcacagt taaaggccat ctacaaggat gcagcggaaa tcatttccga
14341 caactgcgac tgtacgcttt tcctgagtgg aagaggtaaa aacgccaagg aaatcgctga
14401 cgtgctgggc aaggaaacga ttgacagcta caaccaaagc gaaaaccggg gcgcgcagac
14461 ctcccacgga ctgaactacc aaaaactcgg aaaggatatg ccataattgt tcgtgaagag
14521 ttcagttgcc ttgaatttta catgaacagg gacacgatca actggattct gaaaaaacga
14581 atacaggagg tcggaaatgg agaaattacc gaaatacacc cacgcaaacg gcatcgacta
14641 tgttcttgcc ggggattatt acatccctat gctggcgctg ccggaggaaa gccgccctat
14701 tgggcgctgg ggacggatgc accaaaacta tctgaaagag ttccgtcccg gatattatag
14761 cagcttattg ctgtcgggta aactgtggac ttaccttgcc gatctgaatg aacaggcaca
14821 ggatcggctg gatacggtca cagcgcagat gatggctgcg gagggcgtga ccgaggaatt
14881 gaagcgcact tcccaaatgg agtgggttcg ggctgctaat tccatccgca gccgggcgga
14941 agaaatcatc ctgcacgaaa tgatttatgt ataaggagac tgccatgaga tatcgggttg
15001 aacggcggga gggccagcac tgcctgatta tgaacagccg ggcggagctg ctggaatacc
15061 tgaagcatac cccgcccgga acaatcgcag acatacggaa aatttataag aatgggatta
15121 cggattctgt catggaaacc tacttccctt atggcggata cagaagcaag ggataacggc
15181 catcaggctg ccttgtcagt gctccgccaa atccaaaatg atattcgttt gctgcacgag
15241 ccagtccgta aattcattcg ggctggcttt ttccgtcttg atcgcccgtt tccttttcag
15301 ggcttccttt ttgaacgctt caaaagccgc tagcatttct tccagccggt cagccgggaa
15361 gttgggattt ttctttgcct tattgatgat attgcgccag tgctggcact cgtttttata
15421 aacgaggtca tatttgttat ttctggcccg tttatcaaag ccccgcttgt tttcaagcgc

15481 ctgcgctttc cggcatttct cactgcacaa ttcgtagcgc tggctcctgg caagaaagat
15541 tttcccgcag gttttgcatt tccggatata caagccccag tcggaaagcc ggttcagata
15601 atagataatc aaaggatgga gggacacata agcggcaacg cattcagccc ctcttttatt
15661 gtcgggatac cagaggtcga gccgggtatc ttcggctggc accgatcccc atatccgatg
15721 ggagatattc agccgctggg gtctgccggt atccgggtca aattccagcg gcgggtcttc
15781 cataaatgtg gctgtaaagc cgttcatttt acggataaag cggtcacagg cggctggcct
15841 gtcccgtggt tccctggctc gcagatactc gttccagata cggaacgccg catacattct
15901 caccggatcg ccggtgagat atttttttga aagaaattct ccggctgcct gttccaacgc
15961 aggctgtttc cagcggttcg agtgcagggc ttcccgcagc ggggacagtc ccaaacagtt
16021 ccattccccg tccgggtcat gttcaaattc agaaagaaat gtccccagag gccggatgac
16081 gtcacagaga aaatccgcat ccatgtcccc ttgcaggcgg aaacggactt gcgcctcttc
16141 gccaatcaaa atacgctctt tcattttctg cctgtccagt gtcaggacga acaaaatccg
16201 gtcatagacc ggttcatgcc ggttaaatgc aaattccata ggtttgttcc tttcaatgct
16261 tatggtaatt ttataattga gttcaatccg ttacataaag agtaccgcag aagtattgat
16321 ttgtcaagtt taagccgata tgatgattgg cacatggtaa tttcattaca tgcagtacct
16381 tgacaagtga atgaccgtct gacagggata tgaccggcag gagaagtgac gcatgaagcg
16441 caccaatgca gggatacgcc gcaggaatgg ggcagccgca gggatgcggc tatgaggata
16501 acggctgtca ggcagaacgg cgaaaggaaa caaaaatttt atcaaattga ataacaggag
16561 gaacaacaaa atgagtgaaa gcaagagact gaaaaactac gacgatctcc cgctggtgct
16621 ggatgtggcg gacatccggc gtattatggg aatttccaga gtgagcgctt acgagctggt
16681 tcacacgccc ggttttccgg cgttcagaag cgggaggctg attaaagtca gcaaaaaggc
16741 gttctttgac tggatggcaa aggggactga aaccgtaccg gaaagcgaca aatgaaggaa
16801 aggtatcttc ccctgtctgg aatattaccg gcctttttca aggggcatac agaggggaaa
16861 atgcccaaaa atgacaccga aacatgacag taaaacggag ggaacagcaa atgccgagaa
16921 tgagtaaaaa acgtaaggaa gaactggctt tcttcctgaa tgaccgcaac cgtgtcattt
16981 ataatgaact gtgccggaaa tgccgccata gctgcaagca gagtttccgg gtgcagctga
17041 tcgaatgccc gcagtatctt tcaaagcggg cggcgatccc cgtcatagag ccggatgatc
17101 ttccaccggc ggcatgacgg gatttccaat ttcggcagga aattggaaac cggcagaatc
17161 ggaaataaga cagctccatg ctggcagttt ccgattttgc cgggcgggag tacagagggc
17221 gcagcccttt gtgccgggta cagggcggca gcgcctgtgc agagtgtaga ggcagcgagc
17281 ctttactccg gtgtccagag gtggaaaacc tcggcccagg gaaagttgat gtgtgcgtca
17341 acttatactg ggtattacct gacgcagact ttcgcaggat ttgcggcttc gccgccctcc
17401 cccggccccg gaacatcttt cgcaggaagg aggaaatctg tttgaaactg acacgacaca
17461 atgggcgctc cggcaagcat ggcacttaca atccccgtca caatgaccgc cggttcgatg
17521 tggagaacag cgaacacatt gacgctgacc gggccagaca aaatatctac tgggactgct
17581 accgaggctt taccacccat gagttccggg agaacccgga acagccggat ttcagctttg
17641 aggaaattga gcggatgttc tactacgagc attacggcgg ccatgtggag gcccagaacg
17701 cccgcaatga gaaaacacgc cacacggagc gcaaccgtac tgtggaggac ttgctgaaaa
17761 ataacaagac ctgcccggaa gaaacgctct accagatcgg aacaatggaa caatctgttc
17821 caccagaaac gctctttcgc atcgtcacgg aatttcagga ggaatttgac cgccgcttcg
17881 gttcccatgt ccatatcatt gattgggcgc ttcatctgga cgaagccacg ccccacattc
17941 atgagcggca cgtctttgac tgtgagaacc gatacgggga actgtgtccc cagcaggaga
18001 aggcgctgga agaacttggc gtccctctcc caaaaccgga caagcccaaa ggccggaaca
18061 ataaccgcaa gcagaccttt gacgccatct gccggacact gctttttgac atctgccgca

18121 ggcatggcct gcatctggat caggagccgt cctacggcgg gcgggactat ctggaaaagc
18181 aggattatat cctgatgaag cagaaagagc agcttgcggc gcaggagcag aaactggaag
18241 aactgacgct caaaattgag gacgtggaga cgctgttaga tgatgtttcc gatgtggcct
18301 atgacaaggc ggtggaggtc gtgaccgata ccgtccggca ggagacgcac aaagaagata
18361 ttcgtctgat tgaggatacg aaaaaatggg tactttcgcc ggagcgcaag gccccgaaaa
18421 aggaacgtga ttacgccgct gcccgtctgg atggggtcat taccaaaatc aaaaacgcta
18481 tgcagcatgc tctggcaaaa atccagcgga cgctgatgca gccggagatc aagcaggcca
18541 ggaaggagca gatcaagaag aaagccaaag aatccattat ggataagctg gcaaaaggaa
18601 aaatcagcgc tgatcggaag aaccgggagc gctgggaacg tgaggggcgc attgccccgg
18661 aaaaaaagaa cgatatggag ctatgaggca tctgattttc tatggaaacc gggtgccttt
18721 tgctctgtct ggaggtgaga aaatcgaatg tgtttgaagc agtaaagcag gctgtcccca
18781 caaggcaggc tgcggaatct tacggggtgc aggtcggcag gaacgggatg gcctgctgtc
18841 ccttgcacga tgacaagcat cccagcatga aggttgaccg ccgtttccac tgtttcggct
18901 gtcaggcgga cggggatgtg attgatttta ccgcccgtct gtttggtctg agcagtaagg
18961 aggcggtgtt gaaactggcg gatgattttt ctgtccgcta tgacgccaaa ggccatgacc
19021 cgccccgcag aaggccggtc aagcgaaaaa tcagcgagga actgcgctac cggcaggcgg
19081 agcagaaatg tttccgcgta ttgtgcgatt acctgcatct gctggaacgc tgggagcagg
19141 aatacgcccc gcagaacccg gatgaaacat ggattccgct gttcgtggag gccctgcaaa
19201 aaaaatcgta cacggagtat ctgctggata ttctcttgtc cggcagcata gaggaacgtg
19261 cttccgtggt cgctgaatac ggaaaagagg tgaggaagat tgagcagcga gtatcagagt
19321 ttaccgccag ctacccggca ggccgtcatg agcgcagccg aagccttagc accggagcag
19381 agcgttgacg aagtgcggga aagcctttcc gtcacggaaa aaggtcagcc cgccaacacc
19441 attggaaact gccagacggt gttctgccac gatccgctgc tgcgaggggc gattcgtcta
19501 aacattttga ctgaccgggt ggacattgtg cgggatttgg gctggcgcag gaataccagc
19561 gccctgacgg atacagacgt gaagtatctg cttctctatt ttgagcagac ctacggcctg
19621 accagcgaga agaaaatgac agcggccctg tccatcgtgg cgaatgaaaa ctgctaccat
19681 cccatacagg acgtgctgaa cagccttgtc tgggacggga cgccccgcat ccgctcctgt
19741 ctccatcatt tcttaggtgc agaagaaagc gactatgtgg aggaaatgct gaagcacttc
19801 ctgctggggg ctatccggcg ggtgttttct cccggctgca aatatgagga aatgctctgt
19861 ctggtgggcg gacagggggc tggaaagtcc agcttcttcc ggctgttggc aatccgggac
19921 gaatggtttt ccgatgacct gaaaaagctg gacgatgacc gggtgtttct aaagctgcaa
19981 ggccactgga tcatcgagat gtcggaaatg ctggctacca gcagcgccaa aagcattgag
20041 gaaatccgat ctttcatcag ccgccagaag gaaacctatc gcacacccta cgaggcccag
20101 cccaaagacc ggctgcggca gtgcgtgttc ggcggttcgt ctaacacgct ggactttttg
20161 ccccttgacc gggccgggaa ccggcggttc cttccgatca tgatttaccc agagaatgcg
20221 gaagttcaca ttttggagga cgaggacgct tccagagcct atctgttgca ggtctgggcg
20281 gaggccatga cgatttaccg cagcggccac tattccatga agttcagcaa gtctatccag
20341 cgccagctgg tggaggtaca gaaggatttc atgccggagg acaccgaggc cgggcagata
20401 cagggctttc tggaacacta caccggcagc atggtctgct ccaaacagct gttcaaggag
20461 gcgctgggcc acacctatga cgagccgaag cggtggcagc ttcataatat caacgagatc
20521 atgaacacgg tggtgacggg ctggaagccg ttctctaatc cgagaatgtt cgccggatac
20581 ggcaggcaga agggctggga acgggatgtt tccggcaacg aactgcccgg caacgaagat
20641 ggatttgtgg aactgagcga ggaagaatgc cgccagttgg agcttccaaa ggagtggatc
20701 gcctgaaaat agcggtctgt tgccgggatg gttgctgctt ggttgccggg ttcgttgccg

20761 ggtttttccc tgtctggata cgggaaaagc ctgtaaataa gggatatttt tatcattatc
20821 tttatctccg gcaacgaaag caacgagatt tttcaagaaa attcacaaag taagatttgc
20881 gggccagacg gtagtttaca ggttttcaga ggttcgttgc cggacttcgt tgccggaatt
20941 gccgtccggc ctgcctttct atggaggtag cctatggaaa aaagcaaaaa tcagaagaat
21001 aaagtccgtt ttgtggtggt gcgggagttc tccggggaga agtccatgcg ggaagccttt
21061 gagcagctga tcgagcgcca gacctgtgaa cacttcgaag aatggcgtca gcagcgcgaa
21121 atggcagaaa aagcggcgca aaaacggttg attcggggac aggggcatgg tactataatg
21181 gtatcgtgtc cctgttcata gaaggagaac gctatgagca ggaaaaaaca agtctatgaa
21241 aaaatcaccg ctctctactg ccgtatctct ctggatgatg gcggcgacaa tgagagcatg
21301 agcatcagca accagaaact tatgctccgg gactttgcag aaaagcatgg gatgttccag
21361 tatgagtatt atgtggatga cggctatacg ggccgcaatt tcaaccgccc tgccttccag
21421 cgcatgatcg ctgacattga ggcggggaaa atcggctgcg tcatcacgaa agacctatcc
21481 agacttggta ggaactatat cgaagctgga agctatatcg aaatcttttt cccaaaacac
21541 aatgttcgct atatcgccat cacggacggc gtggacagcc tgacccgtca ggaaatggac
21601 atcacgccct ttaagaatat cctgaacgat atgtatagcc gggatatatc caaaaaggtg
21661 ctggcagggc gcatgacccg ttccagacag gggaaatacg gcggcggtcc gacacctctc
21721 ggtctgatgc gtgacccgga ggacaaaggg catcttatcc ttgacccgga gacagcgccc
21781 atcatccgca agatttacga ctatgccctg gacgggctgg gctgcatgag gatttccaaa
21841 cggctgatgg aagaaaagat cccgattacc catgtgaagg caaacacaga atgcgacgcc
21901 aactattatg cctggggcgg ggcgagaatc agccatatcc tgcgaaatcc cttttataaa
21961 ggtgcacatc ttgtctgccg gacacatcag aaaggcatcc gctccaacac ctatgacatt
22021 atcccccgtg aggactggga agtgattgag ggctgccacg aagcgattgt gacgccggag
22081 gaatgggagc aggtgcagga aatcattgac cgcaggccca ccattatgaa aggcaacgcc
22141 tgcccctttt ataatctgtt ccacgggctg gtctactgtg ctacctgcgg gaagtccatg
22201 caggtgcggt atgaaaaagt gggccggact gggaagaacc gattcaccgg cgagatgcgg
22261 gagccgattg acaaggcgta ttatatctgc cagacctaca accggatggg ttgtaaggtc
22321 tgctccagcc acaagattga agcaagggat ttgtataatc tggtgctgaa ggatatacag
22381 gaattggcgg cgcaggccat gaaggatgct gatgcgttct atcagcggct cagcatccgg
22441 atggagcacc ggtatctggt ggacgcttcc cagacggaga aggaacgcaa aaggctggag
22501 gctcggaatc aggaaattga cgggatgttt ttgagcctat acacggataa ggctaaaggc
22561 atcctgaccg agcagcggtt tatgaagctg acggcggcgc tggaacagga gcaggaagcc
22621 aaccaaaagc gcctgcatga tctggcagtg atgcagagcc gggttgacgc ccaggaaagc
22681 gaagtccgca cctttattaa ggaaatccgg cgctatgccg ccattgagga actggacgag
22741 gcggtgttga accggcttat cagcaagatt ctgattggcg aagtcaagaa ggtggacggc
22801 cagaaggtgc aggaagttaa aatcgtttat aactttgtcg gggaaatccc aggaattaac
22861 tgacgcataa gagcagcaag gccctctcca tgccggggag ggccattttt agtgcgccag
22921 gcgggcgcac gttctatatt cttgattcaa cacctatggc tatcaccttg tgtggggaaa
22981 ccccgtaata tcttttgaag gtctctgaga aatgactgat attcgtatat ccgacagtaa
23041 aagcagcctg actcacatta tactgcccgt cttgaagtaa tacatatgct ttcttcattc
23101 gttcttctcg gatgtactga tagattgttt ttctgtaata ttgcttgaaa aggcgcttca
23161 tcttatagtc attcatccgg attaagtggg aaagctcctg aagggagggt ggacaatcta
23221 atcgacggat gagaatctcg cgggcattat ggaggcattc catatctgcg gcagagagag
23281 aaagagggcg tttttctttc caatcggggc atatgagccg ttcgatattt atggaaagca
23341 actccaacac ttgactttcc aacaaaaggt gattcaactg atccttcggc atttccattt

23401 cgttgtcaag cttgttcagc aatatttctt cctgtacatc ttgacaaaag gaataatacg
23461 ggcaattaaa attttgaaaa gcggagaaat tttgtccttt tttcccgcag actgcttcac
23521 agaaagcatc aaatatatgt ggttcaaccc aaatagagaa tgtcctaatt tcgtctccac
23581 attgatactc tgaataactt ctggtttccc ctaaaaaacc caaattagaa taaccgggtt
23641 tgacttcaac aggaacgttc cctacctcac tgtacaaaga atttttctga ctatatgcaa
23701 gttcaaaaat agaaccagta gtagctgtca cccactgaaa ttttcgatct atgatcccgg
23761 aagaaaacaa aagatgaaca ccaggtcgaa cttcaatctg cctgcgcaat ggaaatccag
23821 atgacgattt gaatttcaga gataagtcgg acatacggca cctcatagct aaaccaattt
23881 gcaaatttga acactcagtt tcctctaaac atagtatagt tagttgttgc taacttgtca
23941 agctgttaat atccctttag gggcaaaaaa aatccctctg ggggcagaaa gtggtatgac
24001 aatgtgttat tattcgtatt gattagttga tactaaccaa taaacatatc taaacaagga
24061 ggacaatcaa tgaaaaaaat gttgagtgga ttgctttgtt ttgcgatgct attcagcatg
24121 gcggcttgca caagcgccac aggaacaaat gacacttcag aatcgcagtc aggtgaagca
24181 gaaatatcac aaacggagca gaacggagac acgattacag taacggacat gaagggaact
24241 gttgaaatcc cagcaaatcc acagcgcatt gtagatactt ccggctctgc cgatgaactg
24301 atcattcttg aaatgccttt tgtcggatca gcaaatacta gcatgtttga ttccactacc
24361 gttcctgaat atttgcagga atactttgca gaaaacaacg tggaaactgt gggcaattat
24421 tcaggctccg ccggcgacct ggatctggaa aaaattgcag agttggagcc ggatttgatt
24481 attatgaata tccgccacga caatgtttac gatcagatgt gtgaaatcgc tcccaccgtc
24541 atgctggacg atgatatcag ctatgtaaac tggagaggtc gattccagca actgggggaa
24601 tggttcggga aagaagatgt ggttgaggaa tggcttgcga actttgatac acaggccgct
24661 gaatatgcgg aacagctcaa agcggtcaca ggaaatgaaa cttttgctgt cattgaagac
24721 aactccgtgc gttctggaac atattacgtc tacagtacag gaggccctgg agaactgatt
24781 tatgacgctt tggagcttcc ggcatcttcc ggagtgcccg aggatgtctg gggcgaagaa
24841 gtagatgcgg aatatttttc ccaaatagat gccgaccaca ttctttactt tagcgatgac
24901 ggaacagtag gcgatacggc agatcttccc acctggcaaa atttgaaagc tgtgcaggca
24961 gggaacgtgt actgcggtgt caacgaacag cagtatgatt tggcttatac gcctaacggc
25021 aagttgattt acatggagcg gatagtcaat gcaatcctca accaccaaaa cattgatgag
25081 taaaaagcgc tttgcgggtg gatccactgc ttccgggcag tggatccact ccaagaccgg
25141 atacatcctt tttttgctga ttttaactgt acttttaatt gcttctgcac ttatagcagt
25201 aggcgtaggt attgccgggg ggagcgtttc ggaattgatc catactcttc tacacttgga
25261 tctgtccaca tcgctttcac agatcatgct ggaaatgcgc ttgccaagag ttgctgccgc
25321 ctgtatcacc gggggggctt ttgcactgtc tggcactgtt atgcagggaa tcacctgcaa
25381 ccctctggct gacgccggcc ttttaggtat caatgcaggg gcttgctttg ccgttacact
25441 ttgcactgcg ttcttaccga cactatctta taatgggttg atgggtgcag cgtttctggg
25501 cgcaactgta gcagccctaa tagtttacgg attcggcgcg aagaagaaaa aagccgatcc
25561 gattcggcta attttggcag gttcggctgt ttcatctttt ttaacagctc tcagccaagg
25621 aatgtctctc gcatttggcc tttctaaaga cctttccttt tacacggcgg gtagcctttc
25681 cggaatactc tggcctcagg tcaaagcagt gactccctgg ctgctgatag ctgttataat
25741 cggacttctt cttgcaccta aactatccat cttggcgctg ggcgatgaaa gcgcttcagg
25801 cctcggagta aatgtcggcc ttgtccgtgc atttggcctc tttgccgtgt tgctattggc
25861 cggggcaagt gtatcgctgg ttggaggcat atcctttgtc gggattatta ttcctcatgt
25921 agcacgaaaa ctggtagggg cagattatcg acggctagcc cctgctgccg cactgatggg
25981 ggccgtgctc ttgatacttt ctgacgttgc ggcccgcatg ctccaagcac cttttgacac

26041 cccgacagga gcgctggttt ctgtgatcgg tgtgccgatt ttcctattgc tgacctatcg
26101 gaatgggggg actaagttat gagaagaaag cgcacatgcc tgatacaatt catcttagtc
26161 cttggcctca tcgccaccgt gctgctttca cttaacagcg gatactcaca gactggcctc
26221 gctgacctgt ggggggcaat taccggtacg gcagagaaaa caagcacact gattctgtta
26281 aagatgcggt tgccaagaat catattggcc gttttatgcg gtatgggact ggctctgtct
26341 ggctgcactt tgcagaccgt gacagataat tctctggcag atccagggat cattggtatt
26401 aacgctgggg caggcttttc cgtcatgatt tttatgtgga tgttcccgtc cctgcatatt
26461 gacaccaagt tctatcgccc cttgtttgcg atgatgggcg gtctgctgac agcatttttc
26521 ctttataact tcgccaaacg agacggcaga atacgcccgg cgcatttcct tcttggtggc
26581 attggcgcct cagctggttt cactgctatg atgaccatat tgggaacaaa tatcaacagc
26641 tccatttatc aaatggttca gcgctggctt gtgggcaata tttggggtac tgattggtat
26701 cagatcaaga ttctgttgcc ctgtctgctc gtattggttc cgcttctctt ctggcgggtt
26761 aatgttctcg atcttctttt gctgggagaa gattcagcgt ctgcattagg cgttagagtc
26821 ggacaagagc gcagggtgct actctttctt tctgccgcac tagccgcaag ttgcgttgcg
26881 gtcagtggag ggatcgggtt tatcggtctg gttgcgcctc acattgcccg aaggattacg
26941 gaggtgcggc accgcaattt gcttcttact tctatgctga ctggcggaat tctgctggtg
27001 ctggcagata cagtggggcg catcctgttt tctgggatag aagttccggc aggtatcgtc
27061 atctccatat tggcagcgcc atatttcctg tatctgctgc acaagcaaat ttaaggaggt
27121 gggtttgatg caacggggaa tcgtagcgga aaatttgtcg gcaggttatg ggacacacc
27181 tgtttttcaa aatcttaatt tgaaaatatt gcctggcaaa atcacaacgc tgatcggaac
27241 aaacgggtct ggtaaatcca caattctaaa gacgctgtcc agattgctga cacctgaaga
27301 gggaactgtc tgcttggatg gcgactcgat ctttcgtatg tcaacaagag cagtggcaca
27361 aaggctggca gtgcttccac aaggagcgca ggccccccagc ggcattacgg tgaaagactt
27421 ggtcgagtac ggaagatatc cttaccgtaa aagactatct ggcctgacca gtgaggatga
27481 agaaattatt aagtgggcta tggaaagtac aaatatactg aatatggctg gccgggaaat
27541 ggatcagctc tctggcggcc agaagcagcg agcatggatt gcaatggctt tggctcagaa
27601 aacaggaatc ctttttctgg atgaacctac gacctatctg gacatttctc atcagttgga
27661 aatcctgcag ctgcttaaac aattaaacag gaagcagggt gtgacggtgg ttatggtact
27721 gcatgatctg aatcatgcta tcatgttttc cgaccatttg gtgactatca agaacggaga
27781 aaaatatctg gaaggtccgc cggagcgggt tgttacccca catatgctgc gagaggtttt
27841 tgatgtggaa gcggtggtta ttcaacatcc tgtgttgaat gtgccagtat gcctgcctta
27901 tggtgtgcgt gatcaaatat ttggaaaact tgatagctga atatcgtata ggaaaagaac
27961 ttttcgagaa atggacataa acattggaca tagcgagtca gctttgaaag gaaaaatttt
28021 gaaagcccta taaattcagg gctgaagatt ccgagaattt ataaagagag gaggaatgtg
28081 atttgaaaaa gcaatcaaat ctgtcaagat tgctgacgct ggctggaggg cataagaagc
28141 tcacatttct gggctgcgct ctttccggga tcgcggcggt actcggcctg attccttatg
28201 tgtgtgtatg gctggcggca agagatgtgc tggctgccta cccggacgtc acgggagcct
28261 ccggcatcgt ccggtggggc tggatggcgg tctggtttgc cgtcgccaat atcctggttt
28321 actttgcggc gctgatgtgt acgcatatcg ccgctttccg cacggcgcgg aatatgcgcc
28381 atacaggagt tgctcatatt ctggagcttc ctctgggatt ctttacgggg aaccagtcgg
28441 gaaggctccg gaagatcatc gatgacaatg cggctctcac agaggacctg ctggcccata
28501 agctgccgga tctgacggcg gcggtggtga cgcccatagc ggctgtgatt ctgctctttg
28561 cctttgactg gaggatgggg cttctctgcc tgctgaccat gattctggct tttgtgtgca
28621 tggccagcat gatgggcggc aagaacgcgg gctttttcca ccgctaccag caggagatcg

28681 agaaaatgag cgcggaggct gtagaatatg tccggggaat cccggtggtg aaggtgttcc
28741 agcagaccat atattccttt aaggcctttt atgcggccat ccagtcctac agcgatctgg
28801 ccagccagta tgccatgagc tgccggaacg ggcagacctc atttctcacc tgcatcaacg
28861 gaggctttgc cctgctgata ccggcggccc tgcttctggc ttccggcgga aacgtctggg
28921 aagtgctgac agactttatc ttttacgccc tgtttgcccc ggcctgcggg gccatgatca
28981 accggattat gtatgcctcc gagtctgtca tggaagcaaa tgaggctgta atgaagctgg
29041 atcatattct ggagcagaag cccctggagg agacggacca tccgaagaag ccggggaacg
29101 agaccgtatc cttcgagcat gtgaccttcc ggtatcccgg cacagagcat ccggcgctga
29161 aggatgtgag ctttacagta aaagagggaa ctgtcaccgg actggtgggt ccttccggcg
29221 ggggaaagtc cacggcagcc tctctgatcc cgcggttctg ggacgtccag ggcggccatg
29281 tgcggattgg cggcgtggac gtgcgggata tggacagccg ggagctgatg aagcatgtgg
29341 ccttcgtctt ccaggacacg aagcttttcc aggtcagtct tctggaaaat atccgtatgt
29401 ccaggccttc cgccacagtg gaagaagtga aggcagctgc ccatgcggcc cagtgcgatg
29461 atattctgga aaagctgccg gcaggcctcg acactgtggt gggaacgaag gggtctatc
29521 tctccggcgg cgaggcccag cggattgcgc tggcgcgggc gattctgaag gacgcgccca
29581 tcgtggtgct cgatgaggcg accgcattcg ctgaccccga gaacgaggcc ctgattcaga
29641 aggcgtttag ggagctgacc agaggaaaaa ctgtgattat gattgcccac agactctcta
29701 cggtgcagaa cgcagacaag attctggtga ttgacaaggg ctccgtgcgg gagcagggaa
29761 gccatgaaga gcttctggcc gcaggcggac tgtatgcgga gatgtggtcg gattatcaga
29821 aggcggcggc ctggaaggta gaaaggggtg aggagatatg attcgcaggc ttcggcatgt
29881 atttgccctc agtgaaaggg gtgcgaagga tttcgtaaaa gcagtcatct ggtgcttttt
29941 ctgcaatatc agcctgatgc tgcccgtagg cgtggttatg gctgtgattc agtatctgtt
30001 ggggacgctg gaggcaggag gaaacccggc ggagggctta tggctctata caggagcggc
30061 agttctggtc cttctggtcc tgttcatcct ccactatttc cagtacgcgg ccctgtatct
30121 ggctacttac aaggaaagcg cggcaagaag agtgagcctt gcggaaaccc tccggcggtt
30181 gcctttaagt tttttcggaa accgggatct gagcgacctg acggcgacca tgattgcgga
30241 ctgctccagt ctcgatcaga tgttttccca ctatatcccg caactttttg cttccatctt
30301 ctctacgctg gtgattggaa tctgtatgtt tgccatcaac tggagaatgg ctctggctgt
30361 gctctgggtg cttccggtgg cggttcttct gacggctggc tccaaaaaac tccaggatat
30421 ctgcggcaca aagaatattc tggccaagag gacagtcgcg gacaatatcc aggagtgcct
30481 ggagaccatc cgggatatca aggcctgcag ccgccaggag gaatacctaa agggactgga
30541 cggaaaactg gaatacgcgg agcgctgcgc catccgctcc gagcttgcca caggcgtttt
30601 cgtgtgctcg gcccaggcgt tcttacggat cgggcttgcc accacggtgc tcacaggggc
30661 cgggcttctg atccggggag atctttcctt cctgtatttt ctgggctttc tgttcgcggc
30721 ggcaaggctg tacgatccgc tgggaatggt tctgcagaac atcgcggcca catttaacgc
30781 caagctgcag atcaaccgga tgcgggccat cctggagcag cctgtccaga ccggggcgga
30841 tacctgtgag ccggacaatt atgacattac ctttgaccgt gtgaagttcg cctaccggga
30901 gaatgaaggc gtgctggacg acgtatcctt tacggcgaag cagggagagg tgacggcgct
30961 tgtgggaccc tctggtggag gcaagtccac ggcctgcaag ctggccgccc gcttctggga
31021 tatccagggc ggcaggatta ccctgggcgg catcgacatc tcgaaggtag acccggaggt
31081 ccttctgaag cattattcct ttgtatttca ggatgtggtg ctgttccggg atacgattct
31141 ggaaaacatc cgcctgggac ggcggggagc tacggatgag gaggtctacg cggcggcaaa
31201 ggcggcatgt tgcgacgagt ttgtccggga tctgccggag ggctatcaga ccatggtggg
31261 agagaacggc tccacccttt ccggagggga acgacagcgg atttccattg cccgcgccct

31321 cctgaaagac gcgccggtga ttctgttgga cgaggccact gcctccctgg atgtggaaaa
31381 tgagacggct gttcaggagg ccatctcccg gttggtaaag aacaagacgg ttctcatcat
31441 cgcccaccgg atgcgtactg tgaccggggc agataagatc atcgtgctat ctgacggtca
31501 tgtggccgag cagggaagac cggaagagtt gtatgagaag aacggccttt acacccatat
31561 ggtaaggctt cagacccaga gccagagatg gactctggag acatgaaaag aacctgagga
31621 gtaaaaccgc cggcgcgcag agggctgtat agtcttgtat acagacgctg ccagaaaata
31681 cataaggaaa aacgactgcg gcaggatatc ttcaggtgtg accaggggat atatctgtcc
31741 ggtctgcttc ctgaagcaca gccttcaact gattcagata tttttaggcc actttgaaga
31801 gggcgtggtc agggcaaaat ttcagcaaat cggatacgaa agcatcctaa taggcagcat
31861 gttctccagg attactaggt gattaaaaat tggcgccagt cgattgaaat aaaatataat
31921 caatcggcct cgccgcgaat tgtgagcgat ttgtccagtg tttaaggaca aataagtggg
31981 tgcgtttttg agatagaggt ctgaaattct tataaaatca ggactaaaga ttccgaaaat
32041 ttatagttat tggcaggagg aaaacaagtt atgaaggcaa tgaaattttt atctatggca
32101 gcgatttgcg tctttggcag tgcgctgctt ttctccgggt gctctgaatc cggaaaatca
32161 agtggcgtat atcagcctgg aacttatacc gctagtgctg aagggtatgg aggggacgtg
32221 actgtcgaag taaaatttga cagcgaaagt attctgtccg tcacaatttt agatcaaaac
32281 gagacagaaa gggtatcaga ccgggcttta gaagaaatcc cagaacgaat tgtggaggca
32341 cagaccagcg aagttgatgc gattacaagt gcaactgtca caagtgatgc gattaagact
32401 gctgtaactg attgtatccg acaggccaca caggaataag gaaggggttc caacaattca
32461 acagaagtga agcatgatgt actcgtgggt atgccctgac atttacccta gtgtgctgaa
32521 cacgatgtct ttttcacggt ttctaaaaat cttattttca caaagaagag gctgtctatt
32581 tcccgacagc ctcttcttta gaattagtga atatgggatg ttgcctatca gctctttttc
32641 tactggatat gttggaagat gaaagatgcg agacaaacac acagaaattc tactctaacc
32701 cttctcttaa cggagagagg ggtgtttttt tctttttgaa ttcatcactt gacattgtgg
32761 caaagagctg atgagccagg acgaaattgc aaccatggac ggaggcaagt gtattttgca
32821 ggtgcgcggg gtgaggccgt ttttcagtga gaaatacgac attacccgcc acccccgcta
32881 tcaatacctc tcggatgccg acaaacagaa cacctttgat gtggacaggt acttgtcgtc
32941 cctgcgccgg aaaaaacggc gcgtggtatc ggaggatgaa acctttgacc tttacgacat
33001 tgacctgtcg gacgaagatt tagccgccca gtgaagcggc tttttcctg taatcaatca
33061 aatttggtcg ccggaaaccg gcagaaagtg aggcatatat ggaatttttc aatagtgcag
33121 tggatacgtt acagaccatc gttgtgggcc tgggcggggc gctctgtgta tggggcggcg
33181 tcaatctgtt ggagggatac ggggccgaca accccgccag caatgctcat gtgtggtaaa
33241 gtagcaacta agaatttgat agacagccgc tgactattcc gtttttaact tttagttaat
33301 tttctaaatc atcattttaa caacggctct attgttcctt aaagcctacg ctgctatcat
33361 aaaagcataa acacaaggag gtttgcttat gaaaatcaat caaattatcc gagaaaaacg
33421 gaaagaattg tcattaacac aagaacaagt tgccgagctt ttaggtgtat ccaccccggc
33481 tgtaaataag tgggaaaaag gtagcacata tccagatata accctttac cggctcttgc
33541 ccgtctacta aaaatggatt tgaataccct gttgtcattt aatgaggatt taactgatgt
33601 agagattgaa aactttgtca atgaacttga taaagttgtt caagagcagg gctatcaaac
33661 cgcttttcag atggcaatcg acaaaataca tgaatatccc acttgtgata gactggtata
33721 ttcagctatt ctttatttag atggcgcaag atggctttat agtgtaccag agccagaaca
33781 atatgaggaa acatttgaaa ctttttataa aaggctgtcc gtaaatgaaa ttccagaaat
33841 cagagatata gcaattagta tgctaatttc ctatgcaaga aatagggggcg aatattcaaa
33901 ggcagaggaa ctaatcaata cgctaccatt ctccacaata gacaaagaag aacagcttgc

33961 aattctttac cagcaacaga aaaaatacac agacgcagaa aaattctggg aacacagaat
34021 attgaacggt gttacggaga tacaaaccgc tttaataaat atgcttgaaa tagcgttatg
34081 cgaaaaccgc aatatggatg cagatttttt tgcggatatg tacgaaacaa ttacttgtac
34141 gttttctttc ccggaatgga tgcgttacaa tgctcatttg cagcttgcat tagataagaa
34201 agataaagac gaaagtttat ctattctttc aaaaatgttg cctgctatga gaaaagaatg
34261 gaaatcgcag aattgtaagt tgtaccgcaa tttggacgga acagaggtaa cagctctttc
34321 aagtaaagtg gcagattctc tttgcgagga attgaacagt aaagatgaat ttgcatttat
34381 gcgtgattgt acggagttta gagaactaat gacacaaatg aatagttaag catatctcta
34441 atttgtatag caaatccagc cgagccagtc aacggcaggt aaatgggctg cgcccaccgt
34501 tgacagcccc gactgtcctt gctggtgggt aatcaagggg cgacagcaaa aagtgctgcc
34561 gccctttccc ataatcgaag aaagggggat tttccatgac tgaatacgaa gcccatcaag
34621 aacatatccg ctatacccat gatacctact gccggattgt tattcgccat gcgtcctttg
34681 acgctgcccg tatgctggcg gcgaggtgga aacgggaaat ctcccttgaa tatttgaccg
34741 aagaaaagtt tgtcccacta agcaccacag acgagtattt tcaagtgcca gactatggcg
34801 aaacttatcc attctccgtc cgggggcaga cgatactttt agatagctgc tcccttgcgg
34861 cggctcttgc tggactgccg gaacagacgc aggaggaaat ctttttgtat tacttccagc
34921 acttgacgca gaaagaaatc ggggaacaaa gcggctggac acgcagcaca atcgggcggc
34981 atattcagct tgctttgaag cggctgaaag cggaaatgga ggtgttgtcc catgagtaac
35041 cgacttctcc cctatgacac tatcataaag gcacatgagg gcgaccccat agcgatacag
35101 gctgtccttg accgctatgc cggatatatc cgctacttct ctaagatgaa cggctattat
35161 aactctgata tggaggacta catcagaaca aagctgattg aaagcctgtt taagttccgg
35221 cttgaccgtt gagtttggat tgatgatatg ataaaattag tccaataaat gcgaacggga
35281 ggttagtatg aaaaaaatta tattgatagt ctgtacatta gtctttctta tatctttaac
35341 tgcgtgtggc acaaaatttg atggtagtcg tactggaaac gatagtgaat ttatcatgga
35401 atatagtgtg ttaaatacaa cggacgcaca ggatttaact attgaagctg gagatactat
35461 tagtgctaaa attgtagtag ataaaggtag cttatcaata aaaattcaaa aagacgggga
35521 aaaacctatc tatgaaagta acggaatttc cacatctaat gagtttgatg ttgaaattga
35581 tgaaagcgga acatacacag ttgaagtaac aggaaaaaaa gcaaaaggaa gtgttagttt
35641 catagtgcag agtaattgat gaacttccag tttactacac tgaaccacca taaaaactga
35701 atatcccgcc gccccgcagc ggcagccatg agcagtaaat ccgaaaaaga tttgctgctt
35761 tttttacgcc tatttttggc aaatttccaa atgttccgta ttagatagtg aaacccaaaa
35821 aaggctgcca ttttttcaga cagcgggtaa aacacagttt ccaaaacgcc ttattatcgg
35881 gaaaaaccat gccgcaggaa aagttcttgc cgaaaagtct gtcatttttg ctttttgcgg
35941 tgttgtaggg agtaagggga gaacgccagc ccgcagcctt ttataaaaaa gctggttata
36001 gatttcccga aaaagtggca gcagcaagtg aacggcagtc tgacagtttc ttagagcaag
36061 attctaccga ggtgccaaaa ttcgctttct gctcattttg cccctgcggg aatcttgttg
36121 gggagtgcct tccccaaacc ctgcttatac ggcttgcgcc gcttaaaaat ccctcaaaat
36181 attttttcgg atttttcaaa aacagttccg ctttacaccc tgtttttctc ctattagtga
36241 gaggacacca cgcacagaaa ggaggtctgc cgaccatgaa acgatacaac acgccgcacc
36301 gcagccgggt agtcaaaaca cgcatgaccg aggaagaata cgccgagttt gcggaaaggc
36361 tttctgcctg caacatgagc caagccgagt ttatccggca agccataacc ggggcagcca
36421 tacgccccat cataaccgtt tcccccgtca atgatgagct gcttgccgct gtcgggaagc
36481 tgaccgccga atacggcagg atcggcggca acttaaatca gatagcccgg acgctgaacg
36541 agtggcacag cccctacccg cagcttgcca gggaggtacg ggcggcggtt tccgaccttg

36601 ctgccctaaa gtttgaagtc ttgcagaaag tgggtgacgc tgttggcaac attcaaacat
36661 atcagctcta aaaatgccga ctatggcgca gcggaagcct acctcacatt tgagcatgac
36721 gagtttacca tgaagcccac ccttgatgaa aacgggcggc tgataccgag ggaggattac
36781 cgcatttctt ccctcaactg cgggggcggg gatttcgccg ttgcctgtat gcgggctaat
36841 ctccgctatg agaaaaacca aaaacgggaa gatgtgaaaa gccaccacta tatcatcagc
36901 tttgacccac gggacgggac agacaacggc ttgaccgtag accgggcgca ggagttgggc
36961 gagcagttct gtaaagagca tttccccgga caccaagccc ttgtatgcac ccacccggac
37021 gggcataacc acagcggcaa tatccatgtg catatcgtca tcaactccct gcggatttat
37081 gaagtcccgc ttctgcccta catggacaga ccagccgaca cacgggaggg ctgcaagcac
37141 cgctgtacca acgccgctac ggaatatttc aagagcgaag tcatggagat gtgccaccgg
37201 gaggggcttt accaaatcga cctcctaagc ggcagcaagg aacggataac ggaacgggaa
37261 tactgggcgg caaagaaagg gcagcttgcc cttgataaag aaaacgctgt cagagaagca
37321 gcaggacagc cgaccaagcc caccaagttt gaaacggaca aggcgaagct gcgccggacg
37381 atacggcagg cactttccca agctggtaac tttgacgagt tttcttccct tttgctgcgg
37441 gagggtgtga ccgtcaagga gagccggggg cggctttcct acctcacgcc ggacaggaca
37501 aagcctatca cagcccggaa gctgggggac gattttgaca aggctgctgt ccttgccctg
37561 ctcacgcaga acgcccacag agctgccgaa cagaccacag ccataccaaa atacccccac
37621 acccaaaagg gacgcttgca ggaagaaaaa gccgcaaaaa ccaccccggc agacaacacc
37681 ttacagcgca tggttgaccg ggaagcaaag cgagccgagg gcaagggcgt aggctatgac
37741 cgctgggcgg caaagcacaa cctaaagcaa atggcggcta ccgttacagc ctaccagcag
37801 tacggctttt cttccccgga ggaattggac gaagcctgtt ctgccgccta tgccgccatg
37861 caggaaagcc ttgcagagct gaagcaggtg gaaaagacgc tgaacgggaa aaaggagctg
37921 caacgacagg tgcttgccta ttccaagacc cgccctgtcc gggacgggct gaaacagcag
37981 aaaaacgcca aagcaaaatc agcctaccgt cagaagcacg aaagcgactt tatcatagca
38041 gacgcagccg cccgctattt cagggagaac ggcatttcca agctgccgag ctataaagcc
38101 ctgcaagcag agattgaaac ccttatcaaa gagaaaaaca gcggctacaa cgattaccgg
38161 gcaaaacggg aggaataccg ccgcttacag actgtcaagg gcaatatcga ccagatttta
38221 caccgggagc gcaagcctgt gaaaaggcag gaacaggaac gataaaaacc gccccaaaat
38281 gtacccgaac ccatacagaa caagggggct gccccatacc ctatccgcaa taccaaagga
38341 ttttttaacg gtcttacagg gcagaaaaac cccggaaatg ataccgagat gatacagaat
38401 taccgccgat accgtcacac cagcggaaac ggcagaaagg agcgcaccca tgccaagaat
38461 gagcaagaaa cggcggctgg aatggtcttt tttcctgcgg caagtgaaag tcgggaatgc
38521 cacctgcgac cgtatcacat acaacgacct ctgccggggc tgtacccata gctgcaagca
38581 gagcttccgg gcggttatca tactctgccc ccgctactac tccaaacgcc ggaaaaagga
38641 ggacagggac aatggcagat aaccgcaagt attactacct caagctgaaa gagaactttt
38701 ttgacagcga ctccattgtg ctgctggaag atatgaaaga cgggatttta tattccaata
38761 ttctcttgaa gctgtactta aaatcgctga aaaacggcgg gaagttgcag cttgacgagc
38821 atatccccta cacagcacag atgatagcga cactgacccg ccaccagata gggacggttg
38881 agagagcttt agagattttc cggcagttgg ggcttgtgga gcagcttgac agcggggctt
38941 tttatatgac cgatattgag ctgatgatag gacagtcctc taccgaagcc gagaggaaac
39001 gggctgcaag gctggaaaat aaggcacttt taccaccccg gacaaaagac ggacatttgt
39061 ccgacattcg tccaccagag atagagatag agttagagaa agagatagaa aaagagagag
39121 agggagaaac gggacacccc gcccccgccg cttatggcag atacaacaat gtgatactga
39181 ccgatacaga gctttccggg ctgaaaacag agctgcccga caagtgggag tattatattg

39241 accggctttc ctgccatatc gcttccaccg ggaagcagta ccacagccat gcagccacca
39301 tttacaagtg ggcgcaggag gacgctgcca aagacaaggc tgccccgaaa cagggcatac
39361 ccgattattc atgcaaggag ggcgagagct tatgaaaaac gggattgaag ctatgattac
39421 ggacattaca gccactaccg ccgaagcgga ggactacaca ggcgaggacg ggcttttata
39481 ctgcggcaag tgccatacgc ccaaagaagc ctatttttca aaagaaaccg cccaatggtt
39541 agggcatgac cgacacccga cagagtgcga ctgccagcgg gcagcccgtg aaaaacggga
39601 agccgctgaa agccgacaga agcaccttga aacagtagag gacttgaaac gccggggctt
39661 taccgacccc gctatgcgga actggacatt tgagcatgac aacggcagaa acccgcagac
39721 cgaaaccgcc cgtttttatg tggagagctg ggaaaccatg caggctgaaa atatcggcta
39781 cctgttttgg ggcggcgtgg ggacaggaaa aagctacctc gctgcctgta tcgccaacgc
39841 ccttatggag aaagaagttg ccgtctgcat gacaaacttt gcaacgatac tcaatgacct
39901 tgctgccagc tttgagggta ggaacgaata tatttccagc ctttgcagct atcctctgct
39961 gatacttgat gatttcggta tggagcgagg gacagagtac gggctggaac aggtttacag
40021 cgtgattgac agccgttacc gaagcggcaa gccgctgatc gccacgacca acctcacgct
40081 ggaggaattg cagcacccgc aggacacgcc acacgcccgt atctatgaca ggctgacttc
40141 catgtgcgcc cccgtccgct tcacgggcag caacttccga aaggaaaccg cacaggaaaa
40201 gctggaacga ttaaagcaac tgatgaagca gcgaaaggag agcctatgac agaaaccaaa
40261 cagacaagca ccaccaaaac agaccgccgc ccggactgtg taacggaaat ccgcatgggc
40321 aactccgtcc ttaccgtttc cggcttcttc aagcagggcg caaccgacac cgcagccgac
40381 aagatgatga aagtgctgga agcggaagct gctacacaaa aaacggcgat ttgaccgacc
40441 ataaagaagc agatttgacg cttttgccgc tatacagaca gccgccccat gtggtacaat
40501 caaggtacgg aatagtgggg ctggctgtcg gaaacggagg attttatgtt aagacagacc
40561 aaccaacaac caattaccgc cctttaccca agactttccc atgaggacga gctgcaaggc
40621 gagagcaatt ccatttccaa tcagaagcgt atccttgaaa cctatgcaaa gcagaacggc
40681 ttttccaatc tgcgctggta cacggacgac ggttattctg gtgcgaactt tgctggggtt
40741 agatagcgat actttttaca tctaccatgt tgcggcggtc atggttgaca acttgcgtcg
40801 gctcctgctc ggtatcaacc tgtccgacaa agctataatg gatttcgatt ttccgggtgt
40861 tggtttcccg gtctaattca tgtatcaaaa tacggtcgat aaactcatgg acgttttcat
40921 aggtcaattc ctgtatgtcg ctgtatttcc cgacaatctg aataaacctt gatatatccc
40981 tttttcgctc ggtaacggtt gcgatctgct gttgtaactc gtctatcctt gcggcaaggg
41041 atttcttttc gtcctcatag ccggaagtta gaaacacaaa ccgttcatct gtcagtctgc
41101 ctaatgcgtt gtcctcatac agcttgcgga aaagcgtgtc aagttcggtc atacgcgcct
41161 gtgctttgaa aagttccttt tgctgctgcg ctaatgcctt tctcgcttcc atgtcgccgt
41221 actcggtagc tttacagata aagtcctgtt cgtgttcctt aacataccgc aaaacccgcc
41281 gcatatcttc cagcactaaa tcaagaagta cgtttttgcg gatataatga gaggtacact
41341 cgctgcctgt ccttgccctg ttgcgccatg cgccgcaaga ataggaaaac ttgcgcggct
41401 cgatatttac gccctgttgg taatacatct tatgtctgca accagcgcaa aacagcaagc
41461 cggaaaaaat atcaatctcc gctgcttttg tcgggcggtg gcgggtagca atccgctttt
41521 gcgcaagttc aaaggtttcc tcgtcaacaa gcggctcatg ggtgtttggg aagaaatagc
41581 gttgttcttc ttcgttcttc cgggtctttt tcgacttgta ggataccttg tgagttttcg
41641 cggttatggt atgccctaaa tattcctgcc ttgctaatat gtcatagagc gttttgtccg
41701 gccaagtata gggggcgatc attgcccgct gataccgcgc ctgtcctgta cgctgatagc
41761 gcaacgctcc aatcgtcaag actttgtttt ccgcaagcca tttttggatt tcgcacatac
41821 gcacgccgct aacaaacata gcgaaaacct gtttgacaat cggcgcggtt tccgggtcgg

41881 gtataaggtg gtggcggttg ttcgggtctg ggatatagcc gtatggatat tccccgttga
41941 cgcgctcgcc tttttgcgcc tgtgcctgtt taactgcgcg gattttcttg cttgtgtcgc
42001 gggcgtaaaa ctcgttaaac cagttccgca agggggtaaa ctcgttatcc tcccgcgctg
42061 tgtcaactcc gtcgttgatc gcgatatagc gtactccgtt ttcgggaaat acaatctcta
42121 tgagttcgcc cgttttcaga tagtttctgc caagacggga aaggtctttg gtaatgaccg
42181 tcgccactcg tccggcttcc acttcctgca acatcgcttg cagcccctcg cgctcaaaac
42241 tcacgccgga aaatccgtcg tccacaaaaa actttgtgtt caaaaaatgg tgttcgtccg
42301 catagcgttg aagtatcatt ttttggtgct gtatgcttcc gctttctccg gcttgcatat
42361 cctcttgact taaacggcag taaagggcgg tgattttatt cgactgtaac attagtcctc
42421 ctttccgaca gccgaataaa caggtataat gtgttgctat cattatacca tatcccgccg
42481 cctaatgcac tactcggacg ctaaaaatcc cggatttccg ggctttttct gcaaatcctt
42541 tacaatgagt ttctcaatct ttttatggat tgtatcggtt gccttttcac tcggcaacgc
42601 ttgaacaaga taggtgattt tccctatttt gcgttcggtt gtaattgtct gtttttttatc
42661 cattagaaaa acctcctttt cctgtatgga taaactatat tcgtcaaaag gcaaaaacgg
42721 gcggttgtta gctgcaacct cacgggagtt tcaccccggc ccatgcttat gggtgcgccg
42781 cgcaatactt tgagggtgtt caacgcggga gtatcattgt gccgccttgt atgtcgtcgc
42841 ccacaagctg ctaaacctgt tttacggcgc ggtagttctc gctcggcttt tcccctatgg
42901 ggaaaagctg tcatggcgta cccgccgact ggctcggtac agacggaatg tacccgtttg
42961 cctgttatgc tgcgcaccaa aggccgcttt ctttgtcaaa gaacaatagg ctttgtcggc
43021 ctgcaaaagc acatcaacag cggatatgct tttgcggaac gacaaatagc ccataacggg
43081 aagcgtggaa aggggacacg ctccccgcat gggctaaaag attatcctac atgaaaagca
43141 agggtgcggg taataaggcg cacttgcagc ctgttacgca tttcctcgtc cacataggaa
43201 taggtattgc ccgcgtcgtc tttcagcgtg cgggtacaaa gtttcgctat gtaaccctcg
43261 tagtgcttca agatcgcgca catggcggtt gtgtcgccgt ttgccgctgc ggaaatgaca
43321 gggaacggca acagattttc agacttccta acggtattca tcatctgctt ttccctccaa
43381 atactgtttg attttcgcaa gcgcggattt cctgtgccgg aaaaccgtcg tgcgtacaac
43441 attcagcagt tcgccaattt ccgcgtcgct catatccaag aagtaggaca aaaggataat
43501 gtcgcgtttc ctttcgggca aagcgttaag ggcttcggca agcagttcat ttttgacgag
43561 tacatcaaag ccggacactt gaaaacggaa ataatcgctt tcgtattcgt ccgttgtgaa
43621 aagctgcgcg agttcgcttt cgctcaaatc cgaaaaagta acttcgcgtg ctgcgcgttt
43681 cgcaagagtg cggcggtggc ttttcgcttc gccgaccaaa gttttctttg ctaatgcgtc
43741 gtactgatgt tgtattcttt ccttgtcgga agaagatagc tccatagagt tcacctcctt
43801 cccgcgtgga gtagaggacg ggagttaagg gcttgtcctc tctgcccctt tcgctccgta
43861 cccgttcggg agatggctct tgagtgcgct tttcagaaaa aagttgaaaa aagcaaaatg
43921 cgcccgtccg caagacgcgg acgggcgcaa aggaaatgta agcagtagct aaatgtattg
43981 acagttcaca ttcatagccg gaatatcccg cttgcggagc atagcttttt ttgaccgcaa
44041 agcattaggc gggttacagt aaataaaaat ggacacggcg atacctcccc gataccgccg
44101 tatccttaaa aaagggcgac tttaatacac tacccgcaat ccattctata atagggaaca
44161 gcggcttttg cgcaatatta aataaaaaag ccgataacac ataggttttt tgacctaatg
44221 cgttatcggc tctgcgtcta tgcgtctggc acttttaatc atttttttt gaatttatta
44281 tatgtgttag ctaactgtcc acaagccgcg ttaatctctc tgccatgaga aactctcata
44341 gtaacttcaa gtcctgcttg ctctaattga tgtttgaatg caactatttc ccgtttctgt
44401 ggtgctttaa ttcttgaatt gcttgttggg ttgtattgta acacgttaat cataactttt
44461 ttgccccgaa accattttgc aagttgtctt acatctgagg gccggtcatt tatacccggt

44521 aaaagcaaat acgcaaagac aattttgcga ttatgccttt cagaatagga taaggcttgc
44581 ttaacaacat cttcaatagc atatatgcgc atgtgaggaa taatacgatt tcttgcagat
44641 tgtgttgctg cgtgtaaaga tattgtcaac tgaattttaa gatgttcctc gcgcaatttt
44701 tttaattgat cgaccggacc aactgttgat atggtaatgc cgtcggttgg aaagttgagc
44761 ccatatctat ctcggagaat atggattgct tttatcaagt tgtcataatt gaataaaggc
44821 tctcccatac ccataaaaac gatacggttt acttttcgac gcaacaatat aatctgttgt
44881 acgatttctg acgatgttag attacgaaca aagccatttc gcccggactc acaaaaaata
44941 caaccaacag gacaaccgac ttgtgtgctc acgcaaacag ttccaccatc tcgccgcttg
45001 ataaaaaccg tttcaatgta tttgttgtct ttcagttcgt aaacatactt ttcagtatca
45061 ctgcttttgc aaatattttt taccaacatt gatagatttt ttttgcgtgg taattgctta
45121 tataattctt catataaggc ttttgcttca ttctcgccaa taacttccga catttctttg
45181 taagtaaatc cgtatggatc attccgtact tccgcaggcg tatatttagg taaacgtttc
45241 atttttatat cctttcttct aaataataaa agtttgtttt tctgtatttt tgattacaat
45301 ctctaatttt tctacatcaa tgatatgcgt caatttgcat ttagggcaga aaagaggaaa
45361 atcttttaat acagtattat gaaatacttg aactctcgtc tttccgttac aaatcggaca
45421 ttttatccaa tattttttaa gcattatatt tcactcgtcc tttttacaca aaaaaatgca
45481 ggtcaatcct caacatgagc attgacctgc atttataatg cacagagcag tacaacaaaa
45541 ctaaggcata gcttgcacta tgtctatact atatctatac gtcgttagtt ttttgtatag
45601 catccgcaaa ataagcatga caaaaaagcc catgttgaaa atgggaaaat cctttttttc
45661 aatgcttatc taatacgcat gctatgcaac ataaacgccg cctcctttta cataaatttt
45721 attttatcag aaaatcagtc tactgtcaat acacaacagg aagtatttaa cctaatgata
45781 tgaattgtgt ggacatatcc aaaaagaaag gtgaactgta aaatgtaaca gggtgaatga
45841 aaatggcaaa aagacccgta ccattgtacg actttaaggc tttcggggca gctataaaag
45901 ccgcgagaaa tgaatacggc gagagccgca aaaaggtaag cgacgagtta tatatttccc
45961 cgcgctacct tgcgaatatc gagaacaagg gacaacagcc gagtttacag gtattctatg
46021 accttgtaac ccggtatcat atttcggtag atcaattttt cttcccgaac agcaatgcgg
46081 agaaatccac cgggcggcgg cagcttgacg cgctgctgga cggtatgagc gataaaggca
46141 tacggattgt aaccgcaaca gcaagggaga taacggaagt cgaaaaagca gaggattaac
46201 ctcacaatat gagaaatcgg gagattgcag cgcatggcgg ctgcaatctt tttttgcgtc
46261 caaaatcaaa ggaacgcgca acaaataccg cctttcggtg ggggtatgga gtagatagca
46321 agcacagcaa acgagtaccc gtttgcggaa aatgcttgtt gggataatcc caaaccctta
46381 taccgaaagg cggtgcggaa atggaaaacc gaaagagaaa tatacagatg aagttttacg
46441 ttacggaaga agaaaagcgg ctgatcgacg agaagatgaa gcagcttccc ataaagcagt
46501 atggggcata cttccgtaaa atggcgatag acgggtatat tcttgtcgtt gaccgaagcg
46561 acacaaaagc atatatccgg gaactgcaag cggtgagccg gaacatcaac caaattgcaa
46621 aacgcgccaa tgcgacgggg acggtttaca ggcaggatat agaggacatt aaaaaggcgg
46681 tggacgagat atggcggtta caaagacgca ccctattaaa tcaaccttaa aggctgcgat
46741 agactatatc ttaaatcccg aaaagacaga cgggaagctg cttgcgtcct ctttcggctg
46801 cgggctggaa accgccgata ttgagtttgc atggacgcgg gaagctgccg gagatcgcgg
46861 cacacattta gggcggcact tgatacaatc ctttgcggtg ggagaaacca caccggaaga
46921 agcgcacaaa atcggcatgg aacttgccgg ggcggtatta ggcggcaagt atgagtttgt
46981 tttgacaact cacgtcgata aagaccatct gcataatcac ttgattttca acgcggttag
47041 cttcgttgac tacaaaaagt accattccaa caagcaaagc tatcacttta tccggcgcac
47101 cagcgacagg atatgtaaag agcatgggct atccgtcgtc gtaccgggac aggacaaggg

47161 aaaaagctat gcagaataca ccgccgaaaa gcaagggaca agctacaaag caaagctgaa
47221 aacggcgata gatactctca ttccccaagt gaaagatttt gacgaactgc tgcgccgctt
47281 gcaggaaatg gggtatgaaa tcaaacaggg caaatacatt tcctttcgcg ctgccggaca
47341 ggaacggttt acccgcacaa agacgctcgg cgcggcctat acggaagaag cgataaagga
47401 gcgtatcaag ggcgtgtatg ttgccaaaac aaaaacgctg cgggaagata agaaaatccg
47461 gcttgtcgtc gatcttgaaa acagtatcaa agcccaacag tcggcgggct atgaacggtg
47521 ggcaaaaatc cataatctga aacaggctgc taaaagcatg aacttcctaa ccgaaaacaa
47581 gattgagtat tatagcgaac ttgaaagcaa gatagccgat attatgaccg ctcatgacgc
47641 ggcggcaaag gcggttaagg aagtggaaca gcgtatgtct gatttgtcgc tgcttatcaa
47701 gcacaccacc acataccgac agttaaaacc gatttacgat gaataccgaa aatcgccgga
47761 caaggaaaag tatctgcggg ggcatgaaag cgaaattatc ctgtttgaag ctgcggcaag
47821 ggcattaaag gaaatgcaga taaagaagct gcccgatctc gccgcgctgc gcaaggagta
47881 tagaagctta aacgacagga aaaccaaatt gtatgaagat tatcggcaag ccaagaagca
47941 aatgcaggaa tacggcgttg tcaaaaagaa cgtcgatagt attctttacc cgtcccaaag
48001 cagggcgcgg gagcaggagc gataaggcgc aaaacatggg gtggcaagtg gaatgttaag
48061 ggctgaaaac gcctgtgttt ctgcggcttc cagcgcatga aaacgacata gacgataaag
48121 tatattcaaa cccgcaaaaa cggctttcta attgtccacg caaggacaaa aaaagaacag
48181 gggcgcggtg ccggaaatcg gcaaccgcgc ccctgttctc tatgggctat tcctcgtcgg
48241 tcaagataaa ctccctttcg gaaaactcgc tgtccgtcat ggcttgcagc ttgttcaccg
48301 ctgcggcggc aagctggcgc atatcctcgt ccatgtaggg catggcggca aggatatttt
48361 caagcgttgt ttcccggctg tcctcggcgt agatcgcaac aatgttttct tcctcaatgg
48421 taaagcgggt attcatgcta tcaaacctcc aaatcctttt tatgctcttt ctgtttatct
48481 ttcggctgct gcgccgcctg tttcttgtat tcgtccagct tcttcaaaat ggacggtttt
48541 ttctgcggct gttcccgttt ctcggctttt accgcttttg caaggtcggt aacggaaatc
48601 gcttcccccg cctttgcccg ctgctcaaaa tctgcaacgg tgggcgtttg cggcgtattg
48661 ttgataagcc cgtcaaaatt gttgtcgttc tgctctatgg tgtcctcgac gtgctttaag
48721 ggattatccc gctgcggctg ctcggcggct atggcataag cctgtgtgcc attccccata
48781 atgagatact tcccgtcctc cgacgcatgg tgaaaaccaa atcccgccgc ttcgatctgt
48841 tcacggctca tatcggtaat atgaaaattg cccctcggcg ttctaacggt ttcacccgtc
48901 aaaaggctgt cgggggtcgc ttccggctgc tgcttttcaa gctgcccgtc cttgtaggaa
48961 tatccttgcg ccgcgattgc ttcaagggtc ttgcccgtaa cgtcgccata aatccgcttg
49021 tctgcgtcaa ccagcatttg caaagtgtct tgtacggtgt cggacaaggc ttgctgctgt
49081 tcgggcggca gcttgtcaaa tacgggggtc tgctgctcct gcaaaaactc cggcacttgc
49141 tgaaaaccga tactgtctac atagtgggcg gtgtcctcgt cgttctgatg aagcacaaca
49201 atgtcggaaa cggaaaggct atgcccctta aagtcggcgg ggtggtcgat attaaacctt
49261 tcccaaatgt cgcccagcgt catatccttt gtaagcggtg cagtatagac gagttcataa
49321 tttacccggt caatggtaag tccagccgct tgcagacggt cataaggctc aaagcgcaag
49381 tcccttgttt cgtccccgcg ctttaactga taaatggaaa aggtatcgcc ctgctcctgc
49441 tcggcttgcc gctgctgctg caactcggta aaatgcccct caatctcggt aatgagttcc
49501 tttgcggtgg tctgtattgt ttcaagggac gcttttaact cggtcatttc cttgccgctg
49561 ctccaaccag ccacatagcc gaaagagtaa tcggaagtat caaggccgaa atgctggcaa
49621 accgtataag cgacgctttc cgcttctacc tcgcaggtat ggcggtcaac gcggttttgc
49681 tgctccggcg gggcgtttaa gtctacgtcg tgcagcttcg catgggcgat ttcgtgaatg
49741 gcggtcttta ccgcctgtaa ttcactcacg ccctctttga ttgcaatacg cttgtcggtc

49801 aaactgaaat acccatttgc gccgtttgtg atctgctcaa atgcaatcgg gacgggggac
49861 gctttttcaa gggcggcgaa aaaatcctcg tactgctcca catcggcaag cagcggttta
49921 acggaaaggt cgggaaactc cttgccctcg gtctgcgaaa tatcaaagac ggttataacc
49981 ttaaaggcgg ggacggtgat ttccttttct tcctttacca cttttccctc gcggtcaagc
50041 atgggctttc tcgtttcggg gtcgatttta tcaaccagct ttttcacctt gtagggcgcg
50101 ggggcaaaga ttttaatgcc tttctcgcct ttctttacat ggcggtcaaa ctccttttcc
50161 cattgacgga agcccattac caaattgccg ccctgcatag caattaggat agtgttattc
50221 aaactataat catggaactt tgacatagcg gttagaaact ctttgtagcg gtcgctgtca
50281 aaaatcgcct gtacgccctg ttccaaacgc gcggtgatct cctttacctt ttcggcgggc
50341 ttttcggaag taaggataat ggggacgacg gggcggggct gctgcggctc tgcggcctgt
50401 gcgtctgctt gctgcgtttc ggctgcgtcc atataggctt tgtcctgttc gccgcgctcc
50461 ggctgcgggt aactcataat gcggtattct tccggcactt cgcgcccctc aaactgccgt
50521 tcccactcgt cgccgctttc cacgatatag ccgtactcgg taaacgcgcc tttttcctgc
50581 gcggcagcgt tcctgccaaa agtagaaagg tcaatgccgc ctttccattc ttcgggcatt
50641 tgtaccatgc cggattgatt taagtaatag tcgcccaaat cgtggtagtt atgtacttcg
50701 gggatatgga cgtaaaaatc cacgttgtag gtgaagtcaa ttacctgtcc aatatttgca
50761 agcccgtttt tctgctgcaa agctgcgttc agcttgccaa cctcggcggg gtcaagctgt
50821 tcgagccgcg ccgcaagaaa attgagttcg tccacgtcgg cggcgcatac catttcataa
50881 ggcaaggcaa tatgtttctc cgggtcgtcg gaatagccgc gaatggaaaa atcctgcggg
50941 ttgtcggcgg taatgccgac gttccgcata gcttcgtgca actgctcctt tgtggcgggc
51001 atatccaacc aatagccgcc ggggtcgccc gcgtcaaagc ggctgcggtt gcctatcata
51061 atagaaaact gttcgctcat gtgttcacgc tccttttcct gcccggtatt ttcggggata
51121 gctgccgcct gttgctgctg cgggggttct tcgccctcta tggtgtagtc cggcatgagc
51181 ataccgttta ctttgaaata gtcggcgtac tcctgcggga taggcgggga gatttccgta
51241 aacaagcggt catacgcttt gatacgccca ttcagatact tttccatagc tgccttgtcg
51301 gtgaaaacct ttctatcttg tccggcaatc ttcgcttggg ttctgcttgg gctgttggtg
51361 cgcacgctcc atgtaagcgt ccatgaatac gggatagatt tctgcgcttc tttgtcgtag
51421 cgggtattct cgtaaatgta gtaattcatc tgatataccc gattgctgcg aatatggcgg
51481 ttatgctcgg tggtctgcca ctcgttagcg gtgtgctgga cgtgcggcgt tctcacatat
51541 tcaagtgttt tatccagcaa aagcgttctt cccgcctgtt cttcccatgc ggcggcgcgt
51601 tctttaaggt cgtcaaaaat cgcctgttct gccgccgcgc tttcgctgcg catggaaagt
51661 agctgcgcaa tcggcaagga agcataaggg gcaatctccc tgtcctttgc ttcaaaatat
51721 atccggcgtt ctatccgcat ggtttcggcg gcttctaatt tgtcatggtc gtatgaggaa
51781 tacggcatat tctttcatca cgtcctttct cggtcatata agggtttggg actatcccaa
51841 caagcaaaat gcccgcgctc tctatgggaa gcgcgggcat ttctcgcaaa cgggtactcg
51901 tttgctgtgc ttgctatgta gttgttttta tccgtaaacg cctgtttctc tcgctttggg
51961 ctttctgtct gcggcgtatc tttgccgcgc aatcggggca gtatttcgcc cggttgctgc
52021 ccgccgcaaa cgtgctgccg cataccgcgc aacgcttcaa atcggtttcc ccgctgcggg
52081 tgatctgttc gcaaagctgc ctgtcggcgg gtaataccgc ttcgcggaaa tagcggcata
52141 gcagcgaata gctgatactc tgcacacaaa cgcaagtttc cccgtcgtcc agcagcagac
52201 aattcccgca atcgttgttt gcacaagcgg cgcgggtcat tttctttacg gcgcggtatt
52261 gcttttcatc aagccttatc attttcggtt ctgctcgacg cggaagttta catacttgcc
52321 gcctgtgtcg gcaagcagca caactccgtc aaaggtcttt cccgtcttgg tcgagtacat
52381 acctttgatt tttacctttc cgtctttcaa aagggcggcg gcaatcgcct ttgtgaaagc

52441 ctttttgcgc tgctcaaaaa agcggtcatt cttccacatc acaaagctgc aagccctgtc
52501 ggagcagtaa aagtttttct tgccctcgta aaccgccgcg ccgcaacggg ggcaagtgcc
52561 gattgcttcc cgctgcgggg taaacagctt cgctttatcc tcggaaatcg cggaataggt
52621 cttgacaagc cccgctgtca tttcctcaat cccgcgcatg aagtcggcgg ggctgtcgct
52681 gccctttgca atcccggtaa ggcgcgtttc ccattctgcc gtaagctgcg gggacgtaag
52741 gctgtcgggc agaatagcga caaggttata cccgtccttt gtggggatta gctttttgcc
52801 cttgcgctcg gcaaagcccg cgctaatgag tttttcaatg atacccgccc gcgtcgcggt
52861 ggtgcctaac cctttgcgct ccgcgtcctc cggcgtgtcc tcctttcccg cactctccat
52921 agccgaaagt aacgtatctt cggtaaatgc ctttgggggc tgcgtgtaac gttcggagat
52981 ctccgctgcc ggattgtcaa aggtctgccc ctcggcaagc gggggtaaaa ccgcttccgg
53041 ctctgcgtcc tcgtcctgtt ccccggacgc ggcgcgggat agtctttcaa tctcgcgcca
53101 accctcgcaa agcgtcgtcc tgccctttgc ggtaaaggta tagccgccgc aagtgaaaac
53161 cgccgtaacc gcttcatact catagggcgc ggcaacggcg caaagcagtt ttgcgcacac
53221 aagggtcatt agctttttct cactttcggc aagggaagaa aagcccgttt tcacaaactc
53281 cgctgtcggc aaaatggcgt gatggtcggt tactttgctg ttgtcgcaaa tcctcgcaag
53341 gtcggcttgc agcttcacgc cctgcataaa ggggagagca gcgcaaaggc cggtgataag
53401 ttcctttgtg ctgccctcca tatcggaagt gatatatttg ctgtcggtgc ggggataggt
53461 caaaagccgt ttttcataaa gggcttgcgc atagtcaagg gtctgtttcg ccgtaaaccc
53521 atacaaacgg tttgcttccc gctgcaaggc ggtgaggtca aacagtttcg gcggggctgc
53581 ggtctttttc tcttttttaa gggaagtaca aacggcctgc gatttctcgc aagccgcttt
53641 catctgccgc gccgccgctt cgtcgtcctg cctttcggaa agggcggcaa cgccgccaac
53701 gtccagccgg acgacatgat atttctcttt cttaaagctg ctgattgcat agtcgcggtt
53761 taccagcata gcaagggtgg gcgtttgcac tctgccgaca ttcaatgttt tatggtagag
53821 gatagagaaa aggcgggtcg cgttaatgcc aatgagccaa tcggcctttg ctctgcaaag
53881 cgcggattga tagaggttgt catagccgcg cccgtctttg agatttgaaa agccctcgcg
53941 gattgcactt gcttccatgc tgctaatcca aaggcgggaa aagggctttt tgcaattcgc
54001 ctgttcatag acaaagcgga aaatgagttc cccctcgcgc ccgctgtcgc aagcattgat
54061 aatctcggta acgtcgctgc ggtgcataag gtctttgaga atggaaaact gctgctcctt
54121 gccgctggca acaataaact gccactcctg cgggacaatg ggaagatcgt cgtacttcca
54181 tttacggaat tgttcattgt aggtcgccgc gtctgcaagg ctaatcaaat gcccgataca
54241 ccacgacaca aggtagccgc cgccctcaaa atatccgctg cgcttttcat tcgcgcccat
54301 aacggcggca atcgccgccc cgacgctcgg tttttctgca attacaagtt tcatagatta
54361 tcgtcctcct gctcggtctg ttcggtttcg actacttccg tttcctcgtc gtcctcgtca
54421 aaatcaaggt cgtcaaggct gctgctgcct ttcgccgcgt cctttttggg cttcaaaatc
54481 ttaaagtagt agaacgcgcc gccgcccgct gcggcaacaa gcagcacgac aagcagaata
54541 cccatgccgc cgcctttttc ttcttccggc ggctgctcgg tttcttccgg ctccggctcg
54601 gcttccacac ccgcgcactc gctcatgttg acgctgcaaa ccgggcaagc ggtgttgacg
54661 cttccggcaa cgcatttatc ggtgcagcta caaacggggg cttcggtctg cccgtcctcc
54721 ataagggcgg ttaaatccgc ttcgtccact tggttaagga aatggacggt attttccccg
54781 tcctcggcgc ggtcaatgat gatgtaaaaa tagttgccac ttttggtaac gacggtgata
54841 aactgcttgt cgccggacgc ttcacccgtt atatcgtcca caagggacat attcccgtcc
54901 ggggtaaggg gctgcggctc atagccgccc gtcgtttctt ccggctcggt ttccggcgcg
54961 ggggtttcct cggttacggg ggtaggcgtg tcgtcgcttt catcgccacc gcccgcataa
55021 gcggtaacag aaaagccgca catgataacc agtgcggcaa gcaatacggt aaactttcgg

55081 attatctttt tattcttcat tgtcgttgtc ctcctgttcg gcggggtaag tggcttccgg
55141 ctgcggcaag gtgatttcgc cgctggcgta agcggagagg aacgcggcaa gctgcgtccc
55201 gtcaaggtgt accgccttta ccatttgcac gatctgcgcg ttttcttcct cgactttctg
55261 cgcttcaagg tctttcagct tcttttgctg ttcaaggatt tttgccctcg ttttctcaat
55321 atccctttcg atacggtcaa gtttcgtgtt cgccatagtc atttactcct ttccaaactg
55381 ttctttccat gaatagctgc gccccaatcc cttaacggtt ttaagatagg gcatagcgtt
55441 ttcttccaaa gtcgccgccc ggttaaatag atcgggatag tgttctttga gatataaaat
55501 ctcttgtttc ttcatgctcg gacaaaagaa acaggacgat ttacccggct gcggcaatcc
55561 tgccgctttg atttcccgga tacaatcgtc cctgttccag ccccattctt caatgagggg
55621 gtagcggtta tgatattttt tgtcggcttc gttgtacttg cgggaatgtt cataccgttt
55681 cacttctccc gcgtcatagc cgatatagcg gttgactttc tcgccgcgct gccatacttc
55741 gcggcaaggc gcatagtggt tgcaaaactt ttcctgcggt gcgattttgt gtttctgcga
55801 acaacgctta tgcccgtagg caatcgacgg gagcgtatga gagcgcaagc actcggtttc
55861 aagagatagg cggttgccgt agcggtcaac atactgcaca acggtaatct gcggcatacc
55921 gtgttccgca agccattgat tgattatctc gataaattga taggtatgcg ggtgttccgc
55981 gcccgtatcg gcaaaggtga taaggtcaat cgggattttg tgcttgtata ggccaataag
56041 taaggcggca ctattcgtgc cgccgccaaa agatacaata ttcatttttc aactcctttt
56101 aattgtaatt taaccgcccg taggcgtaaa agtgggattg ccaatagctg gtgtccagct
56161 ttgaatactg gatagggtcg ccacaatgca gcatgacggg gcttccgtcc ccgtcctcgc
56221 ccacataaat ccccacatgg gaaacgccgg gggtgtcgta tgtacccgta aagaacacca
56281 aatccccccgg ctgcgggtta gaaacagggg ttgagatgtt gtaaagccct tgcgccccaa
56341 gtctgcccgt attgtatagg ccgctgttcg tcaacacata actcacaaag ccgctgcaat
56401 caaaagaggt ttccggggtg ctgccgcccc acacataggg aaaaccgacg tatttatctg
56461 cttcctccat gagccgcgca aacttttcat cggaacccag cagcgcggcg ggtacttcat
56521 aatcggcggg cgggttttca tagtatttag acacatagcc ggattgcggg aaaaggtcgg
56581 gtctgtttcc cagcgacgac atatacatag cgtacatgga tagctgttct tcgcccatga
56641 tgtaaacggg gacatgggag aggttgaagt tttccaaatc cacatgaaga atgtaatagt
56701 tataggggac ttcaaccgta taggggttgc cctcgctgtc cgtccttgtt tcggtgcggt
56761 agcgtacctg tacttcttcc tcgaccgtca agatatactg tttctcaaag agcatttcaa
56821 gggtgctttg cacttcatcg agtgtaaaca cgccctcatg ccacgcggaa aggatagaaa
56881 ttagcacata ggggtcatgc tcgatctcgt ccgcgtttac ccggtattcg tcatagccgg
56941 ggtattcgct ttcgatattg tccagcatat cttgtaactc ggcttccata gcggcatacg
57001 cgctttccgc gcccgttata tccgcgtcct cggagagata ggacgacgca atcaagccgc
57061 tgccgctgtt gccgccaaag agggagcaag aggacagccc gccgaaaagc agcagcacga
57121 taaaggcaat cccgcccaca atcaagcagc ctttccaatg ccggacaaca aagaacgtcg
57181 cccgtttggt ttcctgcgcg gttttcttcg ccgcctttgc ggtggcttca gcggtttttt
57241 tcgctttgcc gcccgctttc agcgtttttg catactgctt tttaatgcgc tgcttctgcc
57301 aaaaacggga aatggggttg gagtgcgcaa gcgcggggtt atcgtgcaac gccttttgat
57361 agagaaaatc tgcgtttgcc tttaccgctt gtttttccgc tgccgccgct gcccggtagg
57421 gtttcagctt atggcggcgt atgccgccct taactgcccg cgccgccatg ccgcccgctt
57481 tctctccggc aagttctacc cggtgcccgc tttccacacc gacgttttct ttctctacct
57541 cgtagatttt cccatgtacg gtactgttta actcctgtgc cggacgggat aaggggttat
57601 gccgcaactt gccgtttggg ggcttatccg tcttttcaaa ggaaaggcgc gtctttgcct
57661 tgcctttcgc ttcgtcaaaa acgcgctctt tcttgatttt ggtctgcttg gggatagccg

57721 cccgcgctgc gtctaaacag tctgccgcct tatccgattt cttgatcgct ttttgaagtt
57781 ccggcgttgc ccgttccgct tcggaaaact gcaagcggga agaatgggtt ttaggctgcg
57841 tcgcgtcaag ggctttttc tccgcgcctt ttgctgcgtg tcgctcaatc tctgcgccga
57901 tatgctgcac aaccttttcc gctgccaccc ctgcgggggc tgaatagttt tcttcctgcg
57961 gcctttcgct gatactggac gtttcgcccg tcgtgaggtt ttccgcaacc gcaccgtcgc
58021 gggtcatttt ttgcgttacc ttgtcggggg ctttcagttc cttgctcaat cttcatcacc
58081 tccaatccgc tgttttgcaa gttcgcaata gtcggggttt aactcaatgc cgatatagcg
58141 gcggttaagc cgttttgcta ccatgcccgt cgtgccgctg ccgaaaaagg ggtcaaggac
58201 aatcccgccg atgggacagc ccgccaaaat gcacgtttcc gcaagtttcg gcgggaacgc
58261 tgcgaaatgc ccgccatgat agccgacatt gttaatctgc caaacgttgc ggcggctgcg
58321 gacgggagaa atcaactcgt cggcgtatgc gcccttttcg cgggggcggt tgattttctg
58381 cggttggttt tgccccggaa cagtaacgtt atacttattt cctttgctaa ctccgctttt
58441 cagccgcccc gccgttgtgg gggctatcgg ctcggctact gcttgccaat cgtaataata
58501 cttctttgac ttggtaagca gaaacacata ttcatagcag cgggtcggcc tgtcccgcgt
58561 gctttccggc atggggtttg ttttgtgcca aatgatagaa ctgcgcaaat accaaccgtc
58621 gccccgcaag gcgagggcta caagccacgg aataccaatt aaatctttgg gcttgcagcc
58681 gggggcgcgg tggttaaacg ccacttgctg cccgttcctg cctttggggt atttggggtc
58741 ttggtggtcg gctttgctgc ctgtgccgca ataagtgtcc gcgatattca gccaacaagt
58801 accctccggc gtaagcaccc gctttacctc atggaatacc gccgtaatac gggaaacata
58861 ttcctccggc gttgcttccc gtccgatctg cccgtctgcg ccgtagtccc gcaaggcata
58921 gtagggtggg gacgttacgc aacaatcaac tgcattttcc ggcaaggttt gaagcacctg
58981 caagctgtcg ccgcaataga ttttgtccat ttctatcctc atgccgacac ctcgccgggt
59041 ttggtcgtca ttacacgata gagttccgta tcttgcggga aacggtcaat aaagggcaaa
59101 atcacgttgc catagaagat aagcccctcg cccgcgtcgg aatgggttac ataggtcatt
59161 tgctgctgcg aaatgttaag ctgctttgca aggatttccc tgtcgccctg ggcttggttg
59221 agcatataca caaaatcgct gttttcaaaa atgttctcca cctcgcggga cgataacaaa
59281 tctttcacgt tttgggtgat acctgtcgga ataccgcccc attttctaaa acgcttccaa
59341 atctcaacgc tgtacgctgc ggtctgttcc tctttgagca gcaaatgaaa ctcgtcgata
59401 tagtaccgcg tcgccttgcc ttcgctgcgg tttatggtaa ctctgttcca aacttgatct
59461 tggacaatga gcataccgag ttttcgtagc tgggtgccga gttctttaat atcaaagcag
59521 acaaggcggt tgtgaatgtc aacgtttgtt tggtggttaa acacgttcaa actgccattt
59581 acatagagtt ccaatgccga cgcgatacgc tgcgcttcct tttccggctg cttcttgatt
59641 tcattgtaga ggtcgcccaa aatcggcata ttctccggct tcgggtctgc aaggtagctg
59701 cggtacacgc tgataacggc gcggtcaatc aaagtctttt caaccgcttc cagcccgttt
59761 ttgccgccca taatcaactc gcaaaaggaa aggataaaat cgctctttaa ggctatgggc
59821 gtgtcgccct cgctgtaatt gaggtttata tccatcgggt taatgtactg cgtgctgttc
59881 tgcgaaacct tgataacctg ccctttcagc cgttccacaa gggggtaata ttccgcttcc
59941 gggtcgcaaa tgataatatc gtctgcggta atgagaaaag cgttggtgat ctcgcgcttt
60001 gccgaaaaac tcttgccgct gccgggtgtg ccgagtatca agccgttagg gttcttcagc
60061 cttttcctgt cgcacaaaat catgttattc gataatgcgt ttaagccgta atacaacgct
60121 tcgccgcttt ggaacagctc ttgtaccgta aaggggacga aaatggcggt gctggaagtc
60181 gtcaagcccc gctggatttg gattaggttt tcgcccaatg ggatagagga cataagcccc
60241 tgttcctgcc ggtagtcaag gcgggtgagc atacagttgt atttctgtgc gatacttgcg
60301 gcctgaaata cgtcgttgtc caatttctgc tttgtgtctg ccacattcaa aactaacagc

60361 gtaagcaaaa acattcgctc gttccggctc tgcaaatccc gcaaaaggtt tttcgcttcg
60421 ccgccgtagg tggcaaggtc ggacggaatt atatccatat cgtagccgct gcggattgcc
60481 tttttctgtt cctcgatttt catacggtca aggtcggtga ttttccgctt gatcgtcttg
60541 atcgcttcgt tgtggtcgat actctgaatg tgcagattga cgacaatccc gttttccgct
60601 tccatgaagt cggcaagcat acggtctgaa agttccggcg cggcaatctg caaaaaggaa
60661 accgcgccaa acttcccgcc catttgaaac attcgcccgt cgccaaaacg gaaagaggac
60721 ggggcaataa agtctttgac ggaaaggccg cttacgggta gccattccca cgcaaagttg
60781 aagcgttccc cgtccgggtg gaaaagccca tgcagcactt caagccgctg cttcccgtcc
60841 aatacccgcg ccgccgcgcc gatcacctta aaatggttga gggtgtccgt ttcgatacgg
60901 gcaagccgcg cccgcgccga tttaatatct tttgcttcaa tggtaaaggt gagaaacttt
60961 gtcttgataa gcccgttgtt gccgcgttca agctgtgttt gcagcatttc ggtgtattcg
61021 gctctgatac tgtcaaagtc gtcattctgt ggggcaatcg tgatactgct cttgaaatcc
61081 tccttgttcg catggcggct gattaaggaa agctgcacac tgatagaagc gtcaaaagag
61141 ttatacatat cgcaaagggc ttcaaaaatg gcggtcttgt cgtccggctg cgcaagctga
61201 tagttaatat cttcaaactc aatgcacttg gaataccgcc cgtttgcaag gcggcaaatc
61261 ccgtccttat acattcgttc aaacggtatg ctgtcctgcg ccgtatgggg cttcccgtcg
61321 ccctttgact gccggataag cgcgtcgatc tgtttctttt cctgccgcgt aagtttacgc
61381 tttttgtttt gggtctttcc cttttccttt gacaatcgcc cgtacctcct tatctaatcg
61441 ctcctgccgc gcaagggcgg tataaaaatt gttggtctga taaggtcgct ctttggggcg
61501 aagaaacttc acttccacaa attgccgtat aatcacttcc agcggttgcc cgttcttctc
61561 gtacatagcg agtaggaaca tgggcagcat ggcaagcacc atgcacaaag ccgccgttgt
61621 cgttcccgcg ctgtcctttg tcaaaaagaa tagcggtagc ccgatagcca ccgccgcgcc
61681 gaaacagagt aattgtcttt tggtaaggtt gaacattacc ttgctcttga ttttggttaa
61741 atcctttggg acggttacat acgccatttc aaacctcctt tccggcaaac cgtatgtcgc
61801 ttttgatttc atttccccat acgtcccaac caggggtatg ctgccttgca aacagttcaa
61861 tacgcggtag gtcgcccata agggcaagta tcttttcccg cgttatatcc ggctttttgc
61921 tgtgctgctc aacaggggaa ataataaatt gatgtacctt gtttgattgt cgcttcgggt
61981 gtcctttggt agcaagcaag cagatttccg catttcctcg cgtccaaaag cccaagccat
62041 aaaaccaagt gggcgatttc cgattttgct ttaaccatac aaaggcaacg gatttatagg
62101 taaatcccca agccttgatt aactgcaacg cttctttaag ctgcgggaac gtcgcccata
62161 ggaataaaat acagtcctta tccgcaattt ctttgacggg taaagcgcaa agttcctcaa
62221 tactcatagt gggatagtgc ttttccgctg cgccctgtac cttgcttctc gcgtatctcc
62281 aagggggatc ggcgtaaatg attttgtatt tcttcatctg cacccccttta atgcgcgctg
62341 aatacggatt ttgcaaggct tcccgttttg aaaagcgtaa aacacaagag gacggtatag
62401 cccatgcaag accagatcgc gccggatatg tcatcggttg ctgcgattgt ctgtatcaaa
62461 accgcataaa taccgacgca cacaagaatt agaaaagcct gaaaaccgag ggcaagcaag
62521 gattttaagt agttctgtcc catgccgccc cattcacgat ttaccattgt cgcaacgggg
62581 ataggggcga tactcgttac aaggtatatt tcaatcatgc gcccgtacac caccagcatg
62641 atacagatag acaacgcttt catggtaagc cccacaaaga gtgattgaaa ccatagcccc
62701 aaaagcccgc cgacgctcat agcgtccagc tttgcttcta tatcggtgat aacggttgca
62761 atatctatgc ttgtgtcgga tatgataacg cccgcgctgt cgttgacgac gctctgtgta
62821 atgtcaaata cgcccatgac gatgttccaa gtatttgtta ccagcagcac cgcaacaaag
62881 gttttgaaaa tccacttgaa gaaaatccaa gtgtcgagat cgtgcagatt gtttttctca
62941 attacaagct ggattagttc gtagcacatg acaaaggtaa tgataacgcc cgcaatcggc

63001 acaatgacgt tttccgatag gctgcgtatc atattgaata tgcccgcgtt ccacccggcg
63061 ggggttgtgc cgacttgcgt tgcgatttct ccaacctctg tatttacgct gtcgaacagc
63121 ccggagagat tagagagtat tccgtctgtc aaaatgccgc gaaaccaatc ttcaatcatc
63181 tgccatatca aaagctaatc cctcctttcc cgcgccctcc cgcgtgaagc gggagagcgc
63241 ggcgggtagt gttagtttgt ggcattatcc aaacaggccg gagagcagcg ggacaagggt
63301 aaggccgata agggcaacgc cgccgcccgc catcaactgt ttcatgccct gcgactttgc
63361 tgatgtgtga taaagaagta gtagaagtgt aaaaaatttt gccgttccta tccggcgttt
63421 gcgcattgcg ccggataggt cgggcggtca ttcgggcagg tctaccttgc caacgaaaga
63481 gtaataaata tcaatgtcct gtctgcgtgt gccgttctcg tcatagctgc actcatgcac
63541 aacgattttc tccacaaact cacgcaaaag ggtaggggta agttcttcaa agctggtgta
63601 cttgcggaca acattcataa acttttctgc gtttaccgtg gcttcctgcg ctttggaaag
63661 ctctgcccgg atagcggcgg ctctctcttt cagttctttc tgctcggctt catagtctgc
63721 cgacagctcc gtgaaacgct cgtctgaaat gcgcccggtc acgctgtcct catacagccg
63781 cttgaagata gcggataact cgctgatacg cttctcggcg gcttccagct cctttttcct
63841 tgcggcgttc ctgcgcttgc ccccgtcctc gttctgctcg atcagcagct tcataaaccg
63901 ggcttcgtgc ttcgctgcat agctggtgac tttccgcaga ttgtcggtca ctccggcggt
63961 caacaggtcg gtgcggataa agtgcgccgt acagtcacgg gtacgcttct tgtagctccc
64021 gcagatataa caatcctgct tgcgggtggc gttctggtaa cgctgctggt aaaggacgct
64081 gccgcagtcg gcacaaaaga gtatgccgga gaataagccc acttcatcat agcggttcgg
64141 gcgtttgcgc tgcttgcgta actcctgcac acgctcccac atctgggtgt caatgatagg
64201 ctcatggtgg ttctcgaaaa tcgcctgttt ctcaatgggg ttctctacgc tgtgcttggt
64261 cttgtaggac ggcttctccg tcttgaagtt taccagacag cccgtgtact ccctgttttc
64321 aaggatatgt accacggtat tggtcgccca cttgcactca tagccggggt ggtagcggcg
64381 ggtgcttccc gtccgacggt attccagcgt tcccggcgtg gggatctcct gctctgtgag
64441 catacgggct atcttggtcg gcccgttccc ggcaaggcac aagctgtaaa tctgccgcac
64501 aacaggggcg gcttccccgt caatgataaa attttcgtcc tcgtccataa agtagccgta
64561 tacgggtttg ctcgtgacgg gcttcccgct catacccta gaccgtttta ctgctttgat
64621 tttcttgctc gtatctctca ccagccattc gttaaaaatg ttccgcagag gggcaaaatc
64681 attgtcgccc tgtgcgctgt ccactccgtc attgatagcg ataaagcgga cgcctttctg
64741 tgggaaaatc atttccgtat acattcccac ttgcagatag tttcgcccta accgtgacat
64801 atccttgacg ataacggtgc cgacttttcc ggcttcaatg tccgcaagca tggcttgaaa
64861 accgggtctt tggaagttcg ccccagaata accgtcatct gtgtaccatt gcagattgga
64921 aaagccgttc tgcttcgcat aggtttcaag aatacgcttc tggtttgaaa tagaattgct
64981 ctcaccttgc agctcgtcct cgtgggataa tctcggataa agggcggtaa taaggtttcg
65041 ggtggtctgt cttaacatag tgtcctccat ttccgacagc cagccccact attccgtatg
65101 actatcatac cacacgccgg ggctggctgt acagtccttt ctgcttcttt acgtcccgtc
65161 aaattgccga tttttgtgta gcagcttccg cttccagcac tttcagcatt ttatcggcgg
65221 cggtggcggt ggtgtcctgc ttgaaatagc cggaaacgac aaggacagaa ttacctatgc
65281 ggatttccgt cacgcaatcc gggcggcggg tgctgcggtc attctttggg gtgttggtca
65341 taggcggctc tcctttctgt tcatcagctt tttcagttgt cccattttct cctgcgccgt
65401 ggcttttcgg aagttgctcc cggtaaagcg gacaggagaa cacatttcaa tcagacggtc
65461 ataaatgcgg gcgtgggcgg tgtcctccgg gtgctgcaag tcctccagcg tgagattagt
65521 cgtgacgatc agcggcctgc cgctgcggta acggctgtca atcacgttgt agacctgctc
65581 taagccgtat tccgttcccc gttccattcc aaaatcgtca aggataagca gcgggaagct

65641 gcaaagtcgg gagatatatt cattcctgcc ctcaaagctg gcggcaaggt cacccaatat
65701 caatgcaaag tttgtcatgc acacggggat ttcacgctcc ataagggcat ttgcgataca
65761 gcccgcaaaa tagctcttgc ctgtgccaac gccgccccat agcagatagc caatgttgcg
65821 ctctttcatg gtttcccagt tctccacata gaaatgggcg tgttccattt gcgggcactt
65881 gccgttgtcg ttctcaaacg tccagccctg catagccggg tctgtaaagc cccgccgctt
65941 caaccgctcg acagtttcaa ggtgactgcg ctgtttctct gcggcttccc gttcctcacg
66001 ctctgcccgc tggcagtcac attctgccgg gtggcggtca cgccccagcc atgcggcggt
66061 ttctttgggg aaatagccct ctttgggctt gtggcacttc ccgcagtata aaagcccgtc
66121 ctcgccggtg tagtcctccg gctccggcgt ggtgtcggtc atattcaaaa tcatttcatc
66181 aaatccattc gtcataagct ctcgccctcc ttacaggtat aatcgggtat gcccttttc
66241 ggggctttct tggctgcgtc ctcctgcgcc cacttgaaaa tcgtggctgc atggctcttg
66301 tatttcctcc cgctggaagc gatatggcag gataggcggt caatgtaata ctcccacttg
66361 ccgggaagct ctgttttcag cccgtcaagc tccgtatcgg aaagaatgac attgtggtat
66421 ctgccatagg cggcgggggc ggggtgtccc gtttctaact ctctctcttt ttctatttct
66481 atatctatct ctttctctat ctctatctct ggtggacaaa tgtccgctcg atatggtgga
66541 catttgtccg cccctgtctg cggcagggct ttttgttcct gcaaagccag ccgcgccctg
66601 cgcttgcgct ccccctcggt agaggactgg ccgattaaaa gctcaatgtt gctcatgtag
66661 agtgcgccgc ttggcaaagg ctccacaagc cccagcttca taaagatttt caaagctctc
66721 tctacggtac ctacctgctg gcgggtaatg gtcgcaatca tctgggcggt gtaggggata
66781 ttctcgtcaa gctgcaattt cccgccgttt ttcagcgatt tcaagtacag cttcaagaga
66841 atgttggaat agataacgcc gtcctgcata ctttccagca gaacgattgc atcatcgtca
66901 aaatagctct ctttcagctt gaggtagtaa tatttgcggt tatctgccat aggctgcgtc
66961 ctcctttttc cagcgtttgg aataatagcg agggcatagt atgataaccg cccggaagct
67021 ctgtttgcag tcacgggtgc agccccggca taggtcgttg taagtgatac ggttgcggtg
67081 gttgaggaag aaagaccatt ccagccgccg cttcttgctc attctcggca atggtaagct
67141 cctttctgcc gttcctatcg gtgtagcggg atagggggca ttttctgtat gttctcggta
67201 tcattttcgg gctttttctg ccctgtaagc cccgtttgga agtcggggag tattgcggta
67261 agggttcata tcatacctgt atttttgtag ggtttcggta tcatttcggg gcggtcttta
67321 acgctcctgt tcacgctttt tctgctggct cggtgcgccc cttaaaatct ggtcgatatt
67381 gcgtctgact gtctgaagct cctgcgcccg ctggcgtttc tcccggtagt cgttatatcc
67441 ggcgttcttc tggacggtaa gctgctcaat ctccctttgc agggctttgc tggctggcag
67501 cttggttatt ccatgctccc ggaaataacg ggcggctgtg tccgctatga taaaatcgct
67561 ttcgtgctgt tcccggtagg ctctccgggc tttctccgat ttctgcgctt tcagaccgtc
67621 acgggcgggc ttcgtgccga tatagcccaa aaggttgcgc tgcaattcct ttttctcccg
67681 gatagcggct tccagccctt tcagtccggc aaggctctcc tgcgtggcgg tgttggctgc
67741 ggaaatggca gcgtccagct cgtccgggga agaaaagcca tactgctggt agagcatgag
67801 ggtagccgac atctgtttga ggttgtgcat ggtcgcccac ttctcatagc cccgtccttt
67861 cccctcggct cgcttggcgg ctatgtctac catgcgctgt acagcgtcat tgttcggggc
67921 gtttttcgct tcttttttcg tccgtaagcg gtctttgata ctgccggggt attccgctat
67981 ggctgcgggt ttttcggcgg ctctggcggc gttctgctcc aaaacggaga ggacagcagc
68041 ccggtcaaaa tcgtccccca gcttccgggc ggtaattggc ttcgtcctgt ccggcgtgag
68101 gtaacttagc cgcccccggc tctccttgac ggtcacacca tgccggagaa gcagggcggc
68161 aaactcatca aagccggaag cagcggcaag ggcttcccgg atagtccggc gcagcttcgc
68221 cttatccgtt tcaaacttgg tctgccgggg cgcgataccg tccgcagcaa taggggcgtt

68281 ggctctgtca agggcagcct gtcctttctt ctgcgcccaa tactcacgct cggtaatgcg
68341 ttctttgctg ccgttcaaaa ggtctatctg gtaaagcccc tcgctgtggc acatctccat
68401 gacttcggct ttcagatagt tcatagctgc gtccgtacag cggtgcttgc agcctgcttt
68461 cctgtccgct ggcctgtcca tgtggggcag catgggaacc tccgcaatcc gcagactgtt
68521 aatgacgata tgcacatgga tattttcggt gtggctgtgt ccgtccgggt gggtgcagac
68581 aagggcctgg tgtccgggga aatgctcggc gcagaacttc tcgcccagct cctgcgcccg
68641 atcaacggtc aagccgttgt ccgggccgtc ccgtgggtca aagctgatga tgtagtggtg
68701 gctctttacg tcctcccgtt tctggttctt gccgtagcgg agattggagc gcatacaggc
68761 aacggcaaaa tcctccccgc cacaattcag agaggatatg cggtagtcta ccctcggtat
68821 gagccgcccg tctgcgtcaa gggtgggctt catggtaaac tcgtcatgct cgaatgtgag
68881 gtatttttca gcgtccccgt agttggcatt tttagagctg atatgtttga gtatcgccaa
68941 cggcttcacc tactttctgc aagacttcaa acttcaaggc ggcaaggtcg gcggcggctc
69001 cccgcacttc cttttccagc ccccggtagg ggcttccgta ttcgttcagg ctccgggcaa
69061 tctggtttaa gttgccgccg attttcccgt actctgccgt gagcttggag agggcggcaa
69121 gcagcccgtc attgaccggg gaaacggtga caatggggcg tatggtcgct ttccgtatgg
69181 cttgccggat aaactcggac tggctgattt cataagccgc cagccgcccg gtgaagtcgg
69241 cgtattcttc atcggtcatg cgggtcttta ccacatgacg gcggtggggc gtattgtatt
69301 tctttcgcat ggtctgtgac ctcctttctc gcccgacagg gcgcatagca gggtttgggg
69361 aaggcactcc ccaacaagat tcccgcaggg gcaaaaaataa gcggagagcg aattttggca
69421 cctcggtaga atcttgctct gaaaaactcc ggcgttcccc ggctcccgtc ctttccgcta
69481 acaaaacgtt tcaaaagggc tttgctaccc gctattccgg cttatcttga aaattttcct
69541 ttcactacct acaacaccac gcaaagcaaa atggacggag atttttagaa tttcccctct
69601 attccctaca acaccacgca agccggaata ggcggacaat ggcagtattt ttttctttga
69661 taccctctac ggataagtaa gggattttgc caactgcggc ggctattttc ccttttgttt
69721 tttcatactg gggattttca aaaatgccaa acaggaacga aaaaaagcag caaatctgtt
69781 gaatagattt actgctttct ggttgccggc gaagcggcgg ttattcagtt gtaggcggta
69841 gggctatctc ccaaagcgga acttgaaaat agctgtgaga agcgtctggg tgatgtagtc
69901 ctctgtatct tggtctaccc gtccattcac acgggcggca cattgtatgc ggcggcggta
69961 atactgcaat acagttccca ccgcttccgg ctccccgctg gcggcttgga cgattgtttc
70021 ataggggaga agcctattat ttctcatatg ccattccctc catttccgct tggagctgcc
70081 gtagggcttt tcggatatgg tagcccgctg tgctgcggct gcgcccgtac tgtctgccaa
70141 tttcgtgctg tggaatacgc ttgaaaaagg acaggtaaat catttcccgc tcccgttcgt
70201 ccagccgtga cagggcttcc gcaagtaaac cgctgctgaa aatgaccgta tcgccgcaaa
70261 gggtaagtat gtattcctcg tccggctccg gggcttggaa atattcatct gtcgtgccta
70321 aagggtagtg cttttcgtct gtgaggtatt caagggatat ttctcttttg tgcttcctgc
70381 tccgtgtcct cgctgcgttg atcgttgcat ttcggatatt tagtttgtca gcttgacaaa
70441 taaagggtta aggaatataa ttagattcag tattatacaa ggagttaata aatatgcggc
70501 aaggtattct taaataaact gtcaatttga tagtgggaac aaaaagtagc agtcccgttt
70561 cacttttaat atggggctta gttttttgta cccagtttaa gaatactttt atcatgtaat
70621 tttatatgcc cgaaaacata taagtgtttt ggggctattg gagttattta cccagtgata
70681 ggagtattta tcactgggta tttttatgcc cttttttggg tgttgatagg aggaaaatca
70741 catgaaaata attaacttag gcattctggc tcacgttgac gcaggaaaga caacattaac
70801 ggaaagttta ttgtatacca gtggtgcaat tgcagaacta gggagcgtag atgaaggcac
70861 aacaaggaca gatacaatga atttggagcg tcaaagggga atcactatcc agacagcagt

70921 gacatctttt cagtgggagg atgtaaaagt caacattata gatacgccag gccatatgga
70981 ttttttggcg gaagtatacc gttctttatc cgtattagac ggagcagtat tattagtttc
71041 tgcaaaggat ggcatacagg cacagacccg tatactgttt catgcactac agataatgaa
71101 gattccgaca atttttttca tcaataaaat tgaccaagag gggattgatt tgccaatggt
71161 atatcgggaa atgaaagcaa agctttcttc ggaaattata gtgaagcaaa aggttgggca
71221 gcatccccat ataaatgtaa cggacaatga cgatatggaa cagtgggatg cggtaattat
71281 gggaaacgat gaactattag agaaatatat gtcagggaaa ccgtttaaaa tgtcagaact
71341 ggaacaggaa gaaaacagga gattccaaaa cggaacgtta tttcccgttt atcacggaag
71401 cgctaaaaac aatctgggga ttcggcagct tatagaagta attgccagta aattttattc
71461 atcaacgcct gaaggtcaat ctgaactatg cgggcaggtt tttaagattg aatattcaga
71521 gaaaaggcgg cgttttgttt atgtgcgtat atatagcgga acattgcatt tgagggatgt
71581 tattagaata tctgaaaaag agaaaataaa aatcacagag atgtgtgttc cgacaaacgg
71641 tgaattatat tcatccgata cagcctgctc tggtgatatt gtaattttac caaatgatgt
71701 tttgcagcta aacagtattt tggggaacga aatactgttg ccgcagagaa aatttattga
71761 aaatcctctc cctatgctcc aaacaacgat tgcagtaaag aaatctgaac agcgggaaat
71821 attgcttggg gcacttacag aaatttcaga tggcgaccct cttttaaaat attatgtgga
71881 tactacaacg catgagatta tactttcttt tttggggaat gtgcagatgg aagtcatttg
71941 tgccatcctt gaggaaaaat atcatgtgga ggcagaaata aaagagccta ctgttatata
72001 tatggaaaga ccgcttagaa aagcagaata taccatccac atagaagtcc cgccaaatcc
72061 tttctgggct tctgtcgggt tgtccataga gccgctccct attggaagcg gagtgcagta
72121 tgaaagcaga gtttcacttg gatatttaaa tcaatcgttc caaaatgcgg ttatggaggg
72181 ggttctttat ggctgcgagc aggggctgta tggatggaaa gtgacagact gtaaaatctg
72241 ttttgaatat ggattgtatt atagtcctgt aagtaccccc gcagactttc ggctgctttc
72301 ccctatcgta ttggagcagg ctttaaaaaa agcagggaca gaactattag agccatatct
72361 ccactttgaa atttatgcac cgcaggaata tctctcacgg gcgtatcatg atgctccaag
72421 gtattgtgca gatattgtaa gtactcagat aaagaatgac gaggtcattc tgaaaggaga
72481 aatccctgct agatgtattc aagaatacag gaacgattta acttatttca caaatgggca
72541 gggagtctgc ttgacagagt taaaaggata ccagccagct attggtaaat ttatttgcca
72601 accccgccgc ccgaatagcc gtatagataa ggttcggcat atgttccaca agttagctta
72661 acagcttgca aaagtcatat aaaatgagat ttgaaaggat tagagactaa ttatgatgaa
72721 atgtaacttc atgtttgcta aaaattcaaa ggcatattct gtctacctgc tgttccgatt
72781 tgtctgttcc ctggcggttt ctatgtccac agtgctttcc atcgtgtacc acctggaggt
72841 ggtgcagctg gatgttttcc agcttgtcct ggtagggacg gttcaggaga cctcctgctt
72901 tctgttcgag atacccaccg gtgtggtggc ggatttgtat agccgtcggc gctcggtgct
72961 gattggaatg ttcctctacg gcctgggctt tctgatggag ggtgcgctac cgtggttcgc
73021 gccggttctg ctggcccagg ttgtctgggg ttgcggtgat accttcatca ccggcgctct
73081 ggaggcgtgg attgcctcgg aggaagagga caaacccata gacaaggtgt tcctgcgggg
73141 cagtcaaatg gggcaaatcg gcggcgttct gggcgtggtg ctgggcacac tgctgggaaa
73201 cataaacctg caaatgcctc tcatcttggg gggcagtttg tgcttgttgt tggggctggt
73261 gttgggtcgc atcatgccag aaaccaactt ctcccctgct attgaggaac ggcagggctt
73321 gcttaaagac tttgtctgcc tgttcaagct caacctgggc tttgtgaaag gcgcacctgt
73381 gttgctggcg ctcttagcaa tcacactatg cgggggactt gccagtgaag gctttgaccg
73441 gctctccacc gctcattttc tggatgacac ggtaataccc gttatcgggc cgctgaacag
73501 cgtcacttgg ttcggtgtta tcagtcttat cggcagcggc ttaggtattc tggcttctca

73561 gttgctcatc gcccgcatgg agaaaaaagg gactgtcagc cgaaccagtg tggtcatgtc
73621 caccagcgcc gggtatatcc tgtgcctggt tctcttcgcg gtggggcgga gcttttggtt
73681 catgttgttg gtgttcctgc tggcggggct tatgcgcacc atcaaggagc ctgtgctggc
73741 cgcctggatg aacgaccatg tggatgagaa aatgcgcgcc acagtctttt ccaccagcgg
73801 acagctggac tctttcgggc agatcatcgg cgggcctatt gtggggctgg tagcccagca
73861 ggtgtccata ccctgggggc tggtctgtac cgctttcctg ctgttgcccg cgctgttctt
73921 agtgccggtg gcgggaaaga agcgggattg atatggcata gttaagttta ctgacgagcg
73981 ggagattact tccgctcgtc ttcatttagt agcgcaaaat ttgaggactc tttatttcct
74041 tattcggtat agcggaggcg gctgtcaatt cgacataatt ttcttgtgat actttaccgt
74101 acatgagcat tcgcgccggg gttatcgttg ccgtacccct ccatgaggtt aatgccgccc
74161 caaatgccaa gtcctgcgcc aagcgcgatt accaaagtct gcaatacgtc aactgcgctg
74221 ttgaaaaaat ccatataaac ctcgctttct gccgctaaaa cggctcaaaa aatgttgttg
74281 gaaataaaaa tgccgccgtt tttattcggc ggctgcgtcc tcgtcggaca aatctatttc
74341 gtatatgtca aagggttcgt cgggctttac gattgccgga cgggtggaca tataccgttc
74401 aacgtcaaaa gcgtttttct tgtcgaagtc ggaaaggtat ttgtagttcg ggtgctgcgt
74461 tatatcgtac ttatcgctga aaaaggggcg cacgccgcgc aactgcaaaa tacattttcc
74521 tccgtccata actgcgattt cgtcctgtgt catcaactcc tttcccaatt tctgataatt
74581 gaggccatga gaaacctgcg tgccgcgatt ttcggaagtg ttgaaactgt cgattgtttc
74641 acgtcccaag ttctccgaaa tatcctttgc atttttcccg cgccctccca aaaacaaggt
74701 gctgtcggca ttgtcaagta tgatttctgc cgcgtccttg taaatggctt ttagctggct
74761 ctgcgattgc agaatgatag aagccgagat ttcccggctg cggattgtgg caatcaattt
74821 ttcaaactgc gggatttgtc caatgtttgc aaactcgtct aacaggcagc gaacatgaac
74881 cggcagcttg ccgccgtatt cgtcgtcggc cttgtcgcac aagagattaa ataactgcga
74941 ttgcagcata gcgataacaa aattaaacgt gctgtccgta tcgctcataa tcaaaaacaa
75001 agctgttttt ctgtcgccta acgtgtcaag ttctaattcg tcgtaggaca taagatcgcg
75061 caattccttt atatcaaaag gagcgagcct cgcaccgcaa gaaatgagga ttgattttgc
75121 ggttttgccc gaaacaagtt gtcaaggact ttttaatcaa attcacaaaa aagccacaaa
75181 aaagtgccag aagcactttc atgctcctga cactcaggtg ggtgaggtta cttgcgtctg
75241 cttaacagtt catcgcaaag atacgcatat tctggcaact ggtcgatata gggtttccaa
75301 cgccgtttga tttcggccaa ttccagcggg tcggcttctt ctatcgcgtg gtcgttgcct
75361 gtcatataca gaacatccat cgcaacatca tccaggcact cgtgacagag atcggcggca
75421 gactctatcc tccttccggt atcaacataa agcgggttta ggccaatcgc cagccagaca
75481 aagcccactt tatcagacca tagcagttca aaattagtgc tttgttgaag atgctcggca
75541 aacacctttc gtacccgttc aatctcagtt ttctcttttt ctgtgtacag cattttcatg
75601 tcctccttgg ctaaaagttt tggtgtgtac catcatcagt ttaacacaag gcggctggct
75661 ttatcagccg gtcacatctg ctgtatctcg tttttgagca tacgcttgat tttgtcgctc
75721 atggtttcgg tgctggtatc gctgaaatgg acgcgcacct tgtaggtggt cttgccgatt
75781 ttctttacca gcgcgggggc gtcctccggc gctctggccg gagtggtgtc tgcggtctgc
75841 atggtacggg tttcttcact catggcgctc ctttctccgg gcggggctgc gccgtccggc
75901 aaaaaattac tcatgcttac ggtttacaat gtctttcgcc gcctgcttgg tagctccggc
75961 gttttcctcc cggaagttct tcccggcaaa gaggatcggg gcgcaccgtt ccaggatacg
76021 gtcatagatg cgggcgtggg ccaggtcggg cgggttcttg agttcagcca gcttcaggtt
76081 ggtggtgatt atcatgggcc tgcggctgcg gtagcggctg tcaatgacga aaaacatctg
76141 ctccatcgca tactcggtac tgcgctccac acccaaatcg tcaatgatga gcaggtcgta

76201 ctcgtcaaag ctggcgataa agtcggccct gtcctcggag aacatccctg tcaggcggtt
76261 cagaatggtg ggaaagttcg tcatcagcac cggcacatcc cggtcaagca gggcgttagc
76321 gatacaaccg gcgaaaaagg acttgccggt tcccacatcc ccaaacagca gcaggccggt
76381 gttgttctta taagcctcct tccagttctc cacataggcg cgggccttgt ccatcagggg
76441 attttggccg ttgtcgttgg caaaggtgta gtcgtacaga tagcggtctt gcaagccctg
76501 ggccttccgg cgtttgatac gctccatgcg gcgctgccgt tcctcggcgg ccttgcgctg
76561 ttcctcggcc tcccgctggc actggcagat acagcggggc atgaaatagc cgggtttccc
76621 aaagcatggc acaacggtct gtctctggcc gccgcacttc ttacagtgaa tgaggccgtc
76681 tgccgggtct atgtactcgt ccccggccag ttccacaccg gcggctgcct tgtcgatcag
76741 cgcccggatt tctgcggtaa tctcaatcat acggtttcat cctcctttac gctgtaatcc
76801 cggttgcgag agggcggggc cgctttcctg gcgtcctcgc tggcccagcg ccggatggtg
76861 gcggcatggc tctgataccg cttgccggta gaagccatgt actcggacag cttttcgatg
76921 tactggcccc ataccgtggg aaagctggcc tgcaagtctg ccagttcctc gtccgtcaga
76981 aaaacattct ggtaacggcc ataagcgcgg gcgctctcac tctctctatt ctctatactc
77041 ttatctctaa tatctaatct cttatctcta atctctggtg gacaaatgtc cccccggcta
77101 tatggtggac aaatgtccgc cccgccggat gggagcagtt tttgacgctt caatcgcata
77161 cgctctttct ttttccgttc cccctcggta gaagactggc cgatcagcag ttggatgtcg
77221 gccatgtagt aggccccatc ggtcaaaatt tccacaaggc caaattcaag gaacaccttc
77281 aaggccctct ccaccgtccc cacctgatgg cgggtaaagg tggcaatggt ctggggcgtg
77341 tggggcaaat ggtcattgag catgaggata ccgccgcttt tgagcgacat gaggtacatt
77401 ttcagcagca gattggagta cagcaggcca tcctgcatac tctccaatat aaccatcgtt
77461 tcgctgttgt aaaagttttc tttcagcttg agatagtaat attttcgatt gtctgacatg
77521 ggttcctcct tggggttcgt tttgggcttc tgtacgcatt tagagggccg ttttgggtgt
77581 cagaggataa atctatcagg cccccagcga caccctcgaa aaagtccttg atttacaagg
77641 ggttttcggc ccctaaagcg tgacatttag ccctctgttt ccgcttgcgc tgggcggcct
77701 tgatacgctt catgcgtccg gcacattccg ggcagtattt ggcccggttg gagccggggg
77761 tgaacagcgc cccgcagacg gcgcaccgtt tggcgtccag gcggtggaac agggccgttt
77821 ccagttcctt gtccagcggc agcaccgccg cccggaacca ccggcacaac agggagtagg
77881 aaatggactg gacgcagaca cattcctccc catcatccag cgcgatacag ttgccgccgt
77941 catagttgca gcactcatgt accagccgcc gcgctctgcg gtattggcgg tagtccatga
78001 cggggacggg ttcaggcttg cttttcttca tgctggcact tcccacaccc gccgcagttg
78061 ccgcagccct ggcaggcatg ggcctccggt tccgggccgg tgggcgcgtc tgccggttcc
78121 ggctgcaaga gggcaagggg caggcggata ttcaggtgga cagtgacggg taaaataatc
78181 tgcttcatag ggcaatctcc ttccatgagt gaatgttgac agttctgtgc aaaggggtgt
78241 cgtgaaacgc gacgccctcg aaaggggtcg ccaaaacgcc gtacccttta ccgttccggc
78301 ccccggtcat gggacttgtc ctgttcctgg cgggccagtt ggtcggcggc cttgcggagc
78361 cgttcaacgg ctttcagttc ctcccgcatg gccttcatgt cgatggtgtc cacctgcttc
78421 tgggcagtca gcaggtcaat ctcgcgctgc catgccttgg gggtgatttc ctcgccgctg
78481 tctttcagtt ccttcagata ccgggcagcc gcgtcataga ggatcagttc ccggctgtgg
78541 cgctgctcga acagttcccg tttctccggc ttgaccttgg caaactgctt gcggatgggc
78601 ttgtacttgt tgtactgcgc ccacatctct ccgcgctcgg tgagtgtggc aatccggcgc
78661 tcggctttga cgatctctcc ccgcaggtca tagtagcggg tgttcatatt ggtaattttt
78721 ttatggagct gctgcatggt cttgattccg tttgcctgca aaaagctgaa cagggcggcg
78781 ctctcctgca aagcccggat tttgccggtg cgggtgtcgg gccgcttgag ctgctgctgg

78841 gcctcccaca agtcaatcat gctgggctgc tggccctccg gcttctccgc ctcggcctta
78901 gaccagttgt agaggcgggt aatgcgggcc ttgatttcct tgagcagctt gttgtcggcg
78961 gcgatctgcc ggttgacttc tcccttgtcg gtgcggatgc cgcgcttctc catctggctg
79021 gccgctggcc ccaggtggac agaggggatt ttatcaatgc cccggcgctt gtagctgcgg
79081 tggtctacca gggcgggctg accggcggcc tccaatgccc ggttggtata agccgcccat
79141 gccgcccgcc aaatctccac attccctttg tcgttccagt cggtggtgtc ctcccggtgg
79201 ttcttccagc cgcctttccc atccgggata cgctggccgt tctcgtccag gtcgtatgcc
79261 ttgcggcact tcgcgcccca tgcgccattt tctttcaggg gccggacggt cagcatgata
79321 tgaacatggg ggttgccggt tcccttgtca tggatggcaa aatcggcgca catccccttg
79381 tccacaaagt tgtccttcac ataagcccgg acaagggcaa gctgctgttc cctggacagt
79441 tcgcggggca aagccgcctc aatctcacgg gcaagctggc tgtccctggc tttctcgatc
79501 tgctcgacac tgttccagag ggtggagcgg tctgcaaact ctggcggggc gtgggcgggc
79561 agcatgattt ccgcatggac aacaccgccc ttccgggtgt aatcatgggt caggccgtcc
79621 cattcattcg tcagtttggt tccgctgcgg taggcggctg cggcaacagc ggaacggccc
79681 tcgctgcgct tgatgatact gacgggaata tgacaaaaat ctatgggtgg cacctccttt
79741 cggcggggat atttcaggct ccgtggagcc gaacagacgc gaagcgggtg tttggctgcg
79801 cggggcgcac aaggggtttg gggagtgtag ctccccatcg cggacgaagt acgcaacagc
79861 ccccggcagg gcgcacgaca ccggcaacgg tgtataagtg cgcccttgta aacaagggac
79921 ttatggcttg tccccggatt tcggcagttt cgcggccagt tccgccgccc ctggaagatg
79981 ggacagggca atcaggaacg ccttgacctc tacgccggga agggcggggg ccagagggaa
80041 aatgccctcc aaaatggccc cgcgctcaat caggcggcgg gtgcgggcct tgcgttcctc
80101 attgcgctgc ttgttctcca aaatcttctg gcggttctca agctgccgga tttccttcag
80161 gacttcttcg cgctcgtctt ttttctgttt cgcttgggtg gtgctgtctt tgtgcatatc
80221 gctttctcct ttggcggctt catccagccg cgcttgatat aatatttctg taacttgata
80281 atagcgttgc gaaccgtgac aatggctgct gatgtcgtga agccttcggc ctcggcaatc
80341 tccttaaact gcttgccctc ccaaaaccgg gcgtgcaggc agcttgcctg ccgtggcgtc
80401 agggtggcaa gggcctcccg gagccgttcc atcatcaggg cataggcggc ctcctgttcc
80461 cgttccgttt ccttctccat cagaatatct tccggggaga tgacctgttc cggaaaatgg
80521 tgttctatgc cgggggatgc gtccagagaa taacaatggt agtgctgctt ggcgtcattg
80581 gcgttctctg cgcggctttc ctccacaatg gcgtctgcaa cctcgtccgg cacttccaca
80641 aaagtgtctt tcttgcaaat cggatagtag tattttttca ggttgatggt tttcataggc
80701 cgcttgtcca tttccggcgt atgaagtacc ggcgcagccg ccggagcgcc gcatggatgg
80761 actttccagc ctgggatggg gtgataccct ccatcgcggc aatgtccttc actttcatcc
80821 ccagcatata ccgggcatga acgcggcggg cctgcgttgg ggtcagagag gaaatggcct
80881 catagagccg ccggagcagt tcggcatatt cggcctgcgc ctccttttcc atcaggcaat
80941 cctccggcga aggctgcgcc cagccgatgg cggcgttctc aatcccatca ttgcagtcaa
81001 gggagtaaaa cgccttgtag cggtacatct tgcgctccct ggcggcctcg gcccgcttgt
81061 ccagcaggaa cgcctcgaca atttcatccg acacctccac aaaggtgtcc tcggtgcaga
81121 acggataata ctgtttgagg ttgatggtct gcataggccc gccctcactc tgccggaaaa
81181 aggccgtcat agcagcggcg catattccgc aagccccgcc ggatggcaac gctgaccttg
81241 ctgctgtcta cgccctccct ccgggagatt tccggctggc tgataccggc gatgtagtag
81301 gccctgacgc ggcgggcctg tgtgggggtg gcatgggcca gggcctccgg caaggcagcc
81361 agcagccgca gatgggcggc ccgttcctcg cgctccaaaa ggatttcctc cggcgatctg
81421 ccgtgttcca gcgcatagtt ctccgtccag ctatatgcgt ccagagagta gtaggcgcgg

81481 tggtagacct tccggcgctc atagttgttc atctcgcgct gggcctgccg catgacctcc
81541 caaacctcat cgctgacctc cacaaattca tcgtgccggt agtgaggata aatccaccgc
81601 agattgattg ttttcatagg cttacctcct gaaaatggga gaggcgggca gcttgtccgc
81661 tgtccgcctc ccgctgggat aagggaaaca atccctttca cttggtagcc cgaaaagggg
81721 tcattcgtta agcctgtttt caaagaaatt tataaatttt tttctgactg cctccacact
81781 ctgatacacg gccactttgg agcagccgca caccgtcccg atctccgcaa gggtcagccc
81841 gtccagggca tagagcagga accggcggcg ctgggcctcg gtgcaggtgt ccaggacggc
81901 catcagctcg tccagtgttt ccatgcggca aaggtattcc tccggcgttt cgccgtagat
81961 acccacgccc tcggtggcaa ctatgtattc gtcaaactcc cggccatccc aatgtcgccg
82021 ctgctcatgg gacaggttct cggctttgtg gtcggcccgg tcaagaaatt catagacttc
82081 cagcgtgacc tccacggcca cagcttgtcc gttataattg atggtgactt ccatcggcct
82141 ttcctccgtt cagatttttg cgggaaaatc tgaacggggc ggcggggaac gacaccgggg
82201 acagggttcg ggccatgttg gcccgatgtt ccggccccac aggtatgaaa aagcgcccgg
82261 acagcagaaa gctattcgag cgcaaaaaat acggtattca gttacattgt gacaattttc
82321 gcactggctg ccagacgggg cctgttcggt ggccgggatt tttttgacga tagtgcaaat
82381 atcgtctgaa tttgttggcg gtcagtgtgc ttcatggcgg ctccctccta aagccgccgt
82441 tcttgttagt attagaacgt cattcctatg agagaataga aaccggcggc cttgaatgtc
82501 ttttaaaacc gccgccggat aaaaatgtaa attgagcgca caaaagcagc ctgcccggtt
82561 acggtttttt tctttccccg tttacggggc aggagcgttt gtaagttata tttattctat
82621 tgtttatttt ttctcgatat tgataatccg ctgagtccaa tcccggtaaa aggcctcctc
82681 gtgcgatacg agcaataccg ttcccgcaaa agttgaaagg gctgttttca aagcctcttt
82741 tgcctgtaca tctaaatggt ttgtaggttc atccataatt aagaaattgc agggagtcag
82801 agtaagcaga cacattttta cctttgcctg ctctccaccg cttaatgttc ctatggcctg
82861 catagcgtgt ttactggaaa tcccacatcg tgctaaatgt ttgcggacat cttttgtcac
82921 catattggga taagcattgg caacaatttg aatggcgtc aattcaggtt cttgccactc
82981 taaatcctgt tcaaaatatc cgatagtaac ctgatcggaa aatttgaaat ggccctgcat
83041 agatggaatt tccccaatca gggtttttaa cagtgtcgat tttccaatcc cattaaatcc
83101 agtaataaca actttttgtc ctcccttaat gctaaattca atgtttgata aaacggggta
83161 atggtagcca actgccagat gtttcaccaa caagtgttct gtattggtta atggaagtac
83221 gggaaaatgg aagtatggaa taatttcctt ttggtctaag gcttccatct tttccatccg
83281 gtcaagctgc ttttgtcgtc ccctcgccat ttttgctttt cttcccgcta tgttcttccg
83341 tataaattcc tccgtctttt taatttcctt ttgctgggcc gcatattgtc ttacataatc
83401 ttcccgcagc aaagttttct ttcgcaaaaa ttcggaatat gtgccatagt attttgtgat
83461 agtatcgtta tcaatatcac aaatgcggtt cgctatctta tccagaaagc cataatcatg
83521 agaaactact aaaaaagcat tttccaaagc agataaatat tctgccagcc atgtcacatg
83581 gtctttatca aggaaattgg tcggctcgtc cagcaacaaa acatccggtt tttccagcag
83641 caacttcgct aaaattactt tggctcgttg tccgccactc atttctgcaa tcgaacggtc
83701 aaggccaata gccagaagtc ctaacccatt ggcgacacgc tcaatggctg tatctatcga
83761 gtaaaaatcg tgggcttcaa gctgctcctg ataataggcg gccagttcaa ggcttttcat
83821 atcgccgtca gcagcttttt catagagctc cattgcctgt gtttccatat caaatagctt
83881 tgagaaagca gatttaagaa attccttcat cgttagggaa tggtcaattt ccgcatattg
83941 gtctaaataa ccaaccgtag tatttggttg ccagacgata cgcccgctgt ctggaatgat
84001 ctgctccgta cagattttga ttaaagtgct tttgccggtt ccgttctgtc cgacaatacc
84061 aatatgttcc cccttgttga gtgtaaagcc tgcatttttg tatagcacat tctctccaaa

84121 agagtgtgtc agtccgtcaa tttctaataa gctcatgttt atcaaccttt cctgtatatt
84181 ctgggcaaaa gaaaaaggca gaaagcaaag atacataagt gtatctgcct tctgcctctg
84241 gcatagtagc tcaaaaacaa aaggccatgg acaaaacatt gtccatagcc tcatacacat
84301 tataactgca tgggaagccg ctttctaaaa atgggtataa aatccccttt gatggcttcc
84361 cgatgaaatc taaatgcgtc gtttatacgc tatactctgc acaatgtttc atcggtatgg
84421 attgtacaga gttgattttc tttttacgct gatcggtcaa atgaaaattc atagtcatgc
84481 ctttatctcc ttttgagagg tacaactctc gattctaata atactatgag gggttcaatt
84541 tgtcaatcgt tgcctcaaga agtacagata ataagtaaaa aacttctgct gtttcatcta
84601 tgtccaaatt agttcgctaa aaataacttc ctccgcctgc gctttcagcg tgttcatcag
84661 ccccacccag cgcatggggt cacgggattt cagttcctcg gtgacgcccg ccgcctccat
84721 catgcggggc agcatggcgt ccatccgttc atgcgccgcc cagtcaatct ccaacaggtg
84781 cgggtacagc ttttcgctca aaatcaagct gctgtaaagg ccgggccggt gttccttcag
84841 gtagcgttgc cgcatacggc catacttgcc gatgggggcc tccggctgct cggaaagttt
84901 caggtcgggg atgtagtaat ctccgcagcg ggtgtaagtc aatttgctca tgttcattct
84961 cctttgcact gtgatggtta aactgcggca ctggccgata cggtggcctt gcgcggctcc
85021 tgcttaagta actggctgac ggtggtaaaa atgcccttct ccatgtcctc ggcccggatt
85081 gcggcctttg ccgcctccat ttttgccttg cgcttggcgt tgtagcgttc ttcccattcc
85141 ttctgcttgc cgtttgcctt gcggcgtaag tagttctggt gcaacctgtc cttgcgttcc
85201 tctttcttcc gcagttcttc ctgttcctcc ggggtcaggg gaaccggctg caaggcgggt
85261 gggatatacc ggcccacaaa gttgaagtag atttcaacct cctgcgtggt gtcctggctg
85321 cctttgcggg cgcgttcatg gacaaggatt ttctctacaa actcgttgag catggtattt
85381 gtcagcgtgt cgaaattttc atacttgtca atcagggcaa taaacttctc cgcagatttc
85441 cggctctgct catagccggt aacggccttt tccagttccg tgatttccgc attaagggcg
85501 tcttgctctt tggcgtattg tgcgtccagg gcctcatacc gggcgtccgg cagtttgccc
85561 agggcgttgt cctcgtagat tttgcaaatc agcttttcca gttccccggc ccgcttttgg
85621 gcggcggcca gccgtttccg cttcttggat atgtcggcgg tctgttgggc ggcctgcgtt
85681 tcctggacgg tgcggataaa ctcagcccgg tcattcttgg aatactccgc gatggcccgg
85741 agcatatcgg caatcagggt cagaacagct ttggccttga tacggtgctg tgtagggcag
85801 agggagccga tagggtactt actatactgg ccgcaagtgt actgcgggac gcgcttgccg
85861 ttgtatgtgc ggtgaacata catcttgccg ccgcaatcgg cgcagtacat caggccggtc
85921 agaggatggg cctccccaaa gccgtctggg tagcgtctgg cgtttccccg gatacgctgc
85981 acattgtcaa aggtttcctg gtcgataatg gcctcatggg tgttctcgaa aatcgtccac
86041 tcgctttcgt caacatagtg gcttttcttg tccttgaagt gcttgcgggt cttgaaattg
86101 atggtgtggc ccaggtattc ccggcgcttt agtatctgga caactgtgga cgatccccag
86161 ccgcaggggt cttttacagt cttggtttta ttcacgcctt ccccatgacg ggcaaggtgt
86221 accgatggaa gttcgatttt atcagctgag agttgctgtg caatctgata aggcccgtag
86281 ccttctaatg ccagcgcaaa aattcggcgg acaacatcag cggcttcttc atcgacaatc
86341 caattttccc gtttctcgtc ccacaagtag ccgtaggcca caaagccggt cagatgcttg
86401 ccgctcatgc ctttggcttt gaacacggag cggattttcc ggctggtgtc gcgggcataa
86461 aactcgttca taatgttgcg gaaaggggta aaatcatcat cgcccttcag actgtccaca
86521 ccgtcattga tggcgatcag ccggacaccg cgctgccgca aaatctccat gacctggccg
86581 actttcagat agtcgcgccc caggcggctc atatctttga taacaatggc ctccacccgt
86641 ccggcctcga cttcctccat catcgccaga aaaccaggac ggtcaaaacg ggttcccgat
86701 acgccgtcat cggtgaagtg ggtggggttc ggcagaccat tccggcgggc aaagtcctct

86761 aacatctttt tctggttcga tatggagtta ctctcgccct gtaactcatc atcgcggctc
86821 aaacgctcat acagaggggt gattttctcg tttctcatgg ctgtacctcc tgaaatgaaa
86881 atgctctaag tgactgcttc actaaccatg acaaaaacta tgattaagaa gtcacttaga
86941 gcatacatct cccgccgcca atttatattt tcggtactgc ttcaccgcga aatggtcggg
87001 gtcgcgttct tccaagtcgg caaacagcaa atcgacgggg ctttgatatt cctcgtcgtc
87061 ctcgcgggct tcgctggcgt tgataagttc caacagggta ataaagttca tttcctcggc
87121 gggggcttcg taccagatgt agcccaccaa cgcgcaatac aaaagccgtt ctgcctttac
87181 ccaaaaatcc tccgcgcttt tttctccctc acctttggta ttcgctatca acgtatttac
87241 cagcttcaaa atatctttct cgctgcggat atacacaaag gggttgtagt gcatagattt
87301 tttgaagtta atcgtattca gtaccttaat gcggtagccc gcccgttgca gcagcttccc
87361 gcactcaatc aaaaccgtac ctttcgggtc ggttaagacg taagaggaat gaagctgcat
87421 tagattaggt ttgacaaaaa atcttgtttt tccgctgccg ctgccgccga tcacgacgac
87481 gttcttattt ctcgcgtact tcggctgctt cgggcggctg ttcatggtaa gccgttccgt
87541 ctgcgtcaaa atgatgttgt tttggaaatc cgggtctatg tagggcttta tatcttcggc
87601 gtttccccat cgggcagagc cgtactccgc gcccttgcgg tatttcttcg cgtttttccc
87661 tttgacatag actatcaagc ggataatcac cgcaccgaca atgccgacaa gcaaatccaa
87721 cgggtaaacg ctcggcaagg cgttttcaaa ggctgcggaa aatccttgtg tgatattcag
87781 tatctttgca gaaatatccg cgccgggggc aagccgcacc gcctgtccga ctttatcaaa
87841 gagatagaca aagaacacat aggggatatt cggaagtatc agctttttgt agttgatctg
87901 cttcatatct cgcgcccccct gtccattttc tttaccttgt cgcgggcggc gttaagctgc
87961 ttcgccttgt cctttgcagc ggcaagggct ttgcggatag agggcttttt ctcccggttc
88021 agcttctttg cggaaaactc ttgaaacgct gcggtcataa cgtccgcgtc ccggcctttg
88081 aaaaacacaa ggtaacgggt ctgctcgccc ttaacctttt ttagcgaaaa atctatgttg
88141 tatttcttcg ccgtactctc aaaggcttta atattgctgt cggtgatctc gatgttggaa
88201 acacccgcgt tctgcttcat aagctgccgc atggtctgtt tcccatgcgc gggtttgctt
88261 ttctgctcca aaaacttttg gattgctttc tgtaacgcct gttcggtaag ccgcgccgcg
88321 ccttttccca ctttgacgta tagggcaatc gtttttttggg ttacttcctc ctgcaaggta
88381 tcaactcctt tcccggtaaa gtgcggctat gctcattcgc cgccgtataa gtcgtgattg
88441 acaagggcgg tatagtagct gtcaatggta ctcggcgcgt tgaacagcac cgcttttaga
88501 tactgcttga tgttgcgcac tttggttgtg ttctccctca tgcaatccaa aacaaactca
88561 atatggctgc tgttcagctt caaaaacttg gatttcacca attcggcggg gtagtcgtcc
88621 ccggcaatac ggattgtctt tcgggctgtg cagacggttt ccagcataag gtcaacaatc
88681 tcgtccaaac ggtctttgtc aattttgcag ttttgaatga gatggtcgta ttcgatattg
88741 tccttgataa tctccctgta aatctctact gcgctattgc ttttcgcttc cgttcttttc
88801 ctttccggcg gcgtagccgc ttcgccatgc gcaaagggca aggggtttag ggaatggaaa
88861 ggaatggaat cggtacttga taaatcagta attgattgtt ctttacttaa tatatcttta
88921 tttaattgcg ttggattttc caacgtaggt ttttcctgcg taggtttttc caacgttgga
88981 ttatccaatg ttggattttc caatgtaggt aaatccaata taggcggctg tccgccgctg
89041 gtagctgctt ccggctctcg cggctgcggc tgctcgtata tgacgtaatc tgcgccccgc
89101 aagcgtccct tttcgtcgcg ctcccggctg cgcacgatat atccggcttt ttcgagttcc
89161 tttaccgctt cgcggattgc gtcgatcttc tcccggttga tatgggataa gcccgcaagg
89221 gtataatccc aatcttcggg cagcgaaagc atttgcgaca acaagcccctt tgcctttaag
89281 gacaattcct tgtttcgcaa gtggtggttg ctcataacgg tatatcccgt atttcgctcc
89341 actctgaaaa ctgccatatt tcctcacttc ctctcttggt atgcgtggac aattagaaag

89401 ccgttttcgg ctgttttggt attgcaatat gccctttgcc gttttcgcgt ctgcaaagcc
89461 gcatggcaca agggcttttg cccgtctatt tgtccatgct ggcgtgcgtt ttccgcgtcc
89521 ttttgccggg gcaataggga cactctttga aaacgcaaaa ctcatatttc cacccggggc
89581 ggtagaaacg gcaagtgccg caatcttcat catctccggc acaaccggat tgatagcggt
89641 caaatcccgg cttctgctgc atgagcaatt caaacgcccg ctgcttgccc tttgtgaaat
89701 acattccggc tgcacctcct ttcctgcggg cataaaaaaa cggcgttact ttctccccca
89761 aaggggttaa ggtaacgccg tttgtgtgcg tattcagttt ttaacacatt ccctgtctat
89821 ccgctgtccg caaaaccatt gattttatta gctttttgcc gctatcgcta tcacacctca
89881 ttatttcaaa gacccggttt tcaagccatg cttgcggaca ttgaagccgg aaaagtcggg
89941 acagttatcg taaaggatat gtcgaggtta gggcgaaact acctgcaagt gggaatgtac
90001 acggaaatga ttttcccaca gaaaggcgtc cgcttcatcg ctatcaatga cggagtggac
90061 agcgcacagg gcgacaatga ctttgccccg ctgcggaata tctttaacga atggctggtg
90121 agagatacga gcaagaaaat acgggcggta aaacgctcaa aaggcatgag tggcaagccc
90181 atcacaagca agcctgtgta tggctacctc atggacgagg acgaaaattt cattattgac
90241 gaggaagctg cacccgtagt caagcaaata tacaacctct gtcttgccgg gaatggtccg
90301 accaagatag cccgtatgct cacagagcag caaatcccca cgccgggaac gcttgaatac
90361 cgcaggacgg gacacaccca ccgttaccac cccggctatg agtgcaagtg ggcgaccaat
90421 actgttgtgc atatccttga aaaccgggaa tacacgggct gtctggtaaa cttcaagact
90481 accacgcaat cctacaagtg cagcaaaatc atttacaatg acgaggataa gcaggcaatc
90541 tttgaaaacc accacgagcc tatcatagac actcagacat gggaacgggt gcaggagcta
90601 cgcaagcagc gtaaacgccc caaccgctat gatgaagtgg gcttgttctc cggtatactg
90661 ttctgtgcag attgcggcag cgtcatgtat cagcagcgat accagacgga caagcgcaag
90721 caggactgtt atatctgcgg caactacaag aaacgcaccc atgactgtac ggcgcacttt
90781 atccgcaccg acctcttgac cgctggcgta ctctccaatc tgcggaaagt gaccagctat
90841 gcggcaaagc atgaagcccg gtttatgaag ctcttgattg agcagaacga ggacgggggc
90901 aagcgcagga acgccgccaa gaaaaaggag ctggaagccg ccgagaaacg catagccgag
90961 ttatccgcta tcttcaagcg gctgtatgag gacagtgtga ccgggcgcat atcagacgag
91021 cgtttcacag agctgtcggc agactatgaa gcagaacaac gggagctgaa agaaagagcc
91081 gttgctatcc aagcggagct ttccaaagca caggaagcca ccgtgaacgc agaaaagttt
91141 atgaatgttg tccggcggca taccagcttt gaagaactta cccccactct gctgcgggag
91201 tttgtagaga aaatcgttgt gcatgagtgc agctatgacg agaacaagac ccgcagacag
91261 gacattgaga tttattattc ttttgttggc aaggtggact tgcccgaata accgcccgac
91321 ctatccgaca caagagctaa gtgccggata ggaacggcaa aattttttac acttctatta
91381 cttctttatc gcacatcagt aaagagccag gggatcaagc agcttgtcgc gggcggaggc
91441 gtggccctca ttggcatgac ccttgttccg ctgctgtccg ggctgctggg ataatcgttt
91501 ggtccatctg caagcaatac gccgcgccct cctgctgctg ggagggcgcg ggaaaggagg
91561 gatacacaca tgattgatga tttaatcggg gaatggatca aaggcatcct gattgacggg
91621 atcaagggaa acctgtccgg ccttttcagt accgtgaata ccaaggtcgg ggaaatcgca
91681 tcggatgtgg gcagcactcc gcaggactgg aacggcggca ttttttccat gctgcaaacc
91741 ctgtccgaaa cggtcattgt ccccattgcg gcggcgattc tcgccctggt gatgtgctat
91801 gagctgattg aaatgattgt ggaaaagaac aacatgcacg actttgatac gtccctgttc
91861 ttccgctgga tgtttaagag cgcatttgcc atcctcattg tgacgaacac atggaatatc
91921 gtcatgggcg tctttgacgc cacccaggcc gtcgtgaacc aaagcgccgg ggtcatcatc
91981 ggggaaacca gcattgactt tgataccctg attccggatc tggaaacccg gctggaagcc

92041 atggacatcg gccctctgtt gggcctgtgg tttcagaccc tggttgttgg gctgacgatg
92101 cacgcgttat ccatctgcat tttccttgtc acctacggga gaatgattga aatatacgct
92161 gtgactgcgc tggggccgat tcccttgct acgctgggaa actcggagtg gcgcggcatg
92221 ggccagaact acttaaaatc cctgctggca ctgggcttcc aagccttttt aatcatggtg
92281 gtggtgggga tttacgcggt gctgattcag gacatcggca cgatggagga tatttccggc
92341 gcgatctggg gctgtatggg ctatacggtg ctgctatgct atggcttgtt caaaaccggc
92401 agcttgtcca aagccgtatt cacagcccat tgacgcgaca ctatttgcac aggaggaaac
92461 aggcttatcg agaaatcaca gaagaaaacc tatcacatta tttacgctga cccgccgtgg
92521 agatatgacc agaaaacggt acagggagca gcggaaaacc actatcccac catgggaatc
92581 gacgcgctgt gcgctctgcc ggtggaaacc ctggccgcca aggactgcct gatgtttttg
92641 tgggccacgt tccctatgct gccggaagcc ctgcggctga tccaggcatg ggcttcacc
92701 ttcaagaccg tggcctttgt atggctgaaa cggaataaat aagtgtcttc gacacttcag
92761 ctcccctgcc cctcaatctt gaggggtggg cttgcttttt tcttctatct gattcataat
92821 tatctcatga acatcttaca agagattttt aaagacaatt acgaaatcat tcaatatacc
92881 cttcatccca gagatgttga aatggaaaat atctctaaga tgatttattg tggtgaccct
92941 tcctttggcg gctccatgta tgcctgccct cactgcggca acttgaagtt cgttcctttt
93001 cgctgcaaaa gccgtttctg tcctacctgc ggggtcaaat actcccagga acgctccact
93061 caaatggcct ttaaactaat acgctgtact caccggcatt gtgtctttac cattgatgaa
93121 gaattacggc actttttct ggatgaccgc tctcttcttg actgcctttt ccaggctgtc
93181 cgcagtgttg tcctgcgtat gttccataat ctcaacaaat ctaaaaactt tgttcccggc
93241 atcgtctgcg tccttcacac ttttgggcgt cccctggaat ggaatcctca tatccactgc
93301 ctcatttcgg aagggggatt ctcggatgac ggcttctggc gggtcgtcaa gcattttaat
93361 tataccctct taagaaagtc ctttcagaca gctcttctca atcttctcga agcccgcatt
93421 ggaccttctt ttaaaaaaac gaaagctctt atctaccata aggataaaaa tggtttttat
93481 gtttatgcca aaccaaatct ctgtgatacg gatgcggtca ttaaatatat cagccgttat
93541 ctcggccgcc ccgtcattgc cctgaaacgg attgattcct atgatggcac taatgtcaca
93601 tttcactata accgccacga agacaatgcc tttgtaaaaa ggactcttcc tgttatagat
93661 tttataaagc tcctgattca gcacatcccg gaaaaaaact ttaaaatgac acgttactat
93721 ggtctttatt ccaggcaccg tgataaagat aaatcacttt ggaaggctgt tccaaaatcc
93781 aggcaccgcc tgatgaaaac ttttaataaa tggcgtgaag acatccttct gtcttttggc
93841 tatgacccac tcaaatgtac acaatgcagc cacgaaatgc agttcgtaga gctttattac
93901 aatcatcgcc gtgtttctct ggaagaatta tatgaaaggg caatggcaaa agtaaaatgc
93961 cgttctgctt gatttttcca cctgtctgta ccatacttat cttattaaca gacaggaggc
94021 ttctcatgga ttcaaaattt ttagaggaac tacggcaaaa atatattcaa aacccgccgg
94081 aaggaatgac agccaaactg gtcagaacta tgtctgattc tgatctgctt gatatgcatg
94141 aatttcttac ggaggataat gacctggatg acgatgaatt tgaagaaggc ttttatattg
94201 accttttta accaccgtct gtttgctttg cccgcatttt gcgggctttt tttatcaatt
94261 ggttcgccat ttggcgaact ttcctattat acaagaaaag ccccacctgg ttttacgggc
94321 tggggcactg gacaaggggc aatgcggaaa tctgcctgct tgccaagcgg ggacacccca
94381 aacgctattc ccgcagcgtc catcagttta tcatttcccc cattgaggaa cacagcaaga
94441 aacccgacat cacccgcgag aaaatcatcg agcttgcggg tgacctgccc cgcgcagaac
94501 tgtttgccag acagaaaacc cctggctggg atgtttgggg caacgaagtg gacagcgatt
94561 tttccctgtc cgcaccagaa acgaggtgat tttatttggc ttatgtcact gtccccaagg
94621 attttaccaa ggtgaaatcc aaggtggtat tcgggctgac gaaacgccag ctcatttgtt

94681 ttggcggtgc gctgctggtc ggtgtcccgc tttttttgtt gatccggggc cgcatcccca
94741 ccagtgcggc ggcgctcctt atggtgtttg ccatgcttcc gggctttctg ctggcgctgt
94801 atgagcgcca cggccagccc ctggaagtgg tggtacacca aatcctggag tgttgctttt
94861 ttcagcccaa ggaacggccc taccagacca acaacgctta caccgccctt gtgcggcaat
94921 tccagatgga acaggaggta aatgccattg ttcaaaaagc aaagaaacga aacgagcgca
94981 aaagcgccgg tcaggctcac ccgtgccgag caaaaagaaa ttcaggccgt catccgcagg
95041 tataagggcg acggaaagcc ccactccgcc caggagagta tcccctacga ggccatgtat
95101 ccagatgggg tttgccgtct gacgccccgc aggttttcca agtgcatcga gttttcagac
95161 atcagctatc agcttgccca ggcggacacc aaggcggcca tctttgaaag cctgtgcgac
95221 ctttacaact atctggatgc ctccatccac atccagtttt cctttctcaa ccacaagatc
95281 gaccccaggc agtacgccaa gagccttgaa atccgggcgc agggcgatgc ctttgacgac
95341 atccgcacgg agtattccgc catcctaaaa gaccagctcg tcagcgggaa caacggactg
95401 gtaaagcgga agtttctcac ctacaccatt gaggctgaca gcctgaaact ggcccgtgcc
95461 agactgcacc gcattgaaac cgacctgttg ggctatttca agagcatggg ggccgtggcg
95521 tgggggctgg acgcaacggc ccggctggaa gtcatgcacc gcatgttcca cccggacggg
95581 gagccgtttt ccttcgattg gaagtggctg gcttcctccg gcctctccac caaggatttt
95641 atcgccccgt cgtcgttccg tttcgggaac gcccggatgt tcgggctggg aggcaagtac
95701 ggggctgtca gcttcttaaa catcctgtcc ccggagctgt cggatgaaat gctggccgac
95761 tttctcaata cggaaaacgg cattgtggtg aacctccatg tgcaggccat cgaccagagc
95821 aaggccatca agacggtgaa gcgcaagatt accgacctgg acgccatgaa gattcaggag
95881 caaaagaggg ctgtccgcag cggctacgat atggacatcc ttcccagcga ccttgccacc
95941 tacgggcagg acgccaagga gctgctaaag accctgcaaa gccggaatga acggatgttt
96001 cagctcacct ttcttgtcct caataccgcc gacacccggc aggcgctgga aaacgatgtg
96061 ttttgggccg caggcgtggc acagaaatac aactgttcca ttgtacggct ggactaccag
96121 caggagcagg gccttatgag cagcctccca ttaggggcca gccacatcca gattgagcgt
96181 tctctgacca cttccagcgt ggccgtgttc gtcccctttg taacgcagga gctgtttcag
96241 gatggagagg ccatgtatta cggcgtcaat gccaagaccg gcaacatgat tatgcttgac
96301 cggaagcggg cgcgctgtcc caacggttta aagctgggta cgccaggaag cggcaagtcc
96361 atgagctgca aatccgaaat tttgagtgtg ttcctctgta cgccggatga cgtgtatgta
96421 tgtgacccgg aggccgagta ttatcccctt gtaaaacggc tgcatggaca ggtggtgaag
96481 ctgtcgccca ccagcaaatc ctatgtgaac ccgctggaca tcaacttaaa ttacagcgag
96541 gatgaaagcc cgctggccct aaaatccgac tttgttttgt cgttttgtga actggtgatg
96601 ggcggcaaga acaggctgga cgccatcgaa aagacggtca ttgaccgggc ggtgcaggtg
96661 atctaccggc cctatctggc agacccacgc ccggagaaca tgcccatcct ctccgacctg
96721 cacaaagcct tgttagacca gcacatcccg gaggccgacc gggtggccca agcccttgac
96781 ctgtatgtga acggcagctt aaacgtgttt aaccaccgca ccaatgtgga cattgaaagc
96841 cgcattgtgg cctttgacat caaggagctg ggcaagcagc taaagaaaat cgggatgctg
96901 atcgtccaag accaaatttg gggccgggtg acgcaaaatc gcagccaggg caaggccacc
96961 tggttctttt gtgatgaatt tcatctgctg ttgcgggagg aacagacagc ggccttttcc
97021 tgcgagattt ggaagcgttt tcgtaagtgg ggcggcatcc ccacaggtgc cacgcagaac
97081 gtcaaggatc tgcttttatc gccggagatt gaaaatattt tggaaaattc ggacttcatc
97141 tgcctgctca accaggcgtc cggcgaccgg cacatccttg cggagcgatt gaacctatcc
97201 ccccagcaat tacggtatgt ggaaaattcc gaacccggcg aggggctttt gatttacgag
97261 aatgtagttt tgccctttaa gaatcctatc ccaaagcaca cccagcttta ccagatcatg

97321 accacccgct taggcgagag ggccacggta tgaggtgcgg ggccgcaaaa ggaggtggcg
97381 caccatgaag cccttaaaac cccgtgataa ggtgacgcag cgcatgaccc gcgctgggct
97441 tacactggac aaccagacca ccggcgagag catgaacatt tccagccggg aggccgaacc
97501 ggaatacacg gccaagccgg atggtacagc ggaaaaggcg ctggaacggg ctgtggacat
97561 ccgcgaccgg cataaggcaa agcaggcggc ccggcatgga gagcgtatgg caaaggaagc
97621 cagcggcccg gcctcccgac tgcaatttac cgccgaggaa cgggcctccc cggaactggc
97681 cccgtatatc aaacaggcgg agaaacgggc cgaccggctg gatacagcaa aaagcgccct
97741 cccgaacagg cgcgtgatta ccaaggaaac cgtctacaac gaagccaagg gaaaggccag
97801 aagcaaactt cactttgaga aagtggagaa acatccgcca aagctcaaac ccaatccggc
97861 cagccgcccg gtgcaggagg cggggctata cctccacgga aaaatccatg aggtggagca
97921 ggaaaatgtg ggtgtggaga gcggccataa ggcggaagaa ctggccgagc gccaagcggg
97981 caaggccctg agaaacgcca gacggcggaa taagctgaaa ccctaccggg ccgcagccaa
98041 ggcggaacga aaatccatgg ccgccaatgc cgagtttgtg tatcaaaaat ccctgcggga
98101 caacccggaa ctggcgcagg cggtcacgaa tcccatttcc cgcctctggc agaaacagca
98161 catcaaacgg gaatacgcca aggccgcacg ggcggcgggc cggggcgcgg caggcagtgc
98221 caagaccacc gcatcagccg ccaggaaagc cgcagagaaa ggtaaacagg ccgcgtccct
98281 tgtggcccgt cactggaaag gggcgctgct gattggcggc gtgggcctca tgctcctgtt
98341 tctgatgggg ggcctgcaat cctgcaccgc catgtttggc agtgccggga cgggacttgc
98401 ggccacctcc tatctctcgg aggacagcga tatgctggga gccgaggctg cctacgccgg
98461 gatggaagct gatttacagt acgaattgga ccactacgaa accctgcatc ccggctacga
98521 cgagtaccgt tttgagctgg acgagatcgg ccatgacccc tatgtactga cctctatcct
98581 ctccgcgctg cataacggcg tgttcacctt ggaggaagta tagggcgacc ttgccatgct
98641 ctttgagcag caatatatcc tgacccagac ggttgagacg gaaatccgct accgcaccga
98701 aacgagtacg gacagcgagg ggaacgagta cgaggaagaa gttccctaca cctattacat
98761 ctgcaacgtg acgctggaaa acagggattt atcccacctg cccgtctccc tcatggacga
98821 ggaagcatta agcctgtacg ccgcctatat gcagacgctg ggcaaccggc cagacctctt
98881 tccaagcggc agctatccca atgcctccac cattaaagag ccgacctact atgaaatccc
98941 accggaggcg ctaaaggatg aagcctttgc ggccatgatc gcggaggcgg aaaagtatgt
99001 tggctttccc tatgtctggg gcggcagctc ccccagcacc agctttgatt gttccggctt
99061 tattagctgg gtggtcaatc attccggctg gaatgtggga cggcagacgg cgcaggggct
99121 ttacagtctc tgcacccccg tttcaccgga acaggcaagg cccggcgacc tggtattttt
99181 cgtcggcacc tacgacaccg ccgggatgtc ccatgtgggg ctgtatgtgg gcaattcggt
99241 gatgctgcac tgcggcgatc caatcagtta cacgaaccta aattcaagct actggcaaca
99301 gcatttttat tgttacgggc gtttacctta aacgccagaa aggagtgttt gactatggca
99361 atgaacaaat tggaacgcat tgaaaaggac attgagaaaa ccaaggacaa gatcgcggcg
99421 ctgcaaaagc agcttcggga tctggaagcg gccaagacgg aacaggaaaa cttgcagatt
99481 gtccagcttg tgcgggacct gcacatgacg ccccaggaat ttgccgcctt tgtccgggat
99541 ggcgcgttgc aggcaccccc agccccgcag ccggattttg agcaggacga acaggaggaa
99601 accgccgatg aagaataaag ccattcgcat gttcaccgca acactggccg ctgtgctttg
99661 catggcggct gtttccgtca ccgcctacgc cggaggtgtt gaccctcacc cgcagccatt
99721 gccggaggaa acggaaaaac ccaccaccgg ggggatcgag atggagccgg aggacgtacc
99781 cgtcacaccc aagggcaatg cggcgctggt ggatgatttt ttcggggata agcagttgat
99841 caccgtcacc accaaggccg ggaattactt ctatatcttg atcgaccggg ccaacgagga
99901 caaggaaacg gcggtgcatt tcttaaacca agtggacgat gccgacctgc aagcactgtt

99961 gaaggacggc gagaccaagc cggaacactg tacctgcaca accaagtgtg aggccggggc
100021 tgtcaatacg aattgcccgg tgtgtaaaaa caacatgaac gcctgcaccg ggccggagcc
100081 ggaagaacac aagccggagg aagccaagcc gcaggaggaa aagccaaacg gcataggcgg
100141 ccttgtggtg ttcctggcgg tcgttctggc aggcggcggc gcggctcttt acttcttcaa
100201 gttccgcaag ccaaaagcgg acatcaaggg cggcgacgat ctggacgagt acgactttgg
100261 ggaggacgag gacgacgagg aagaagcccc ggagccagat gacacccaag agccggggga
100321 agattggttg atcgagaaag aagcacctga accaaacaag gaggatgagg aatgagtatt
100381 cagttagtga ttgcagagaa accgagcgtg gcccgcagca ttgcggcagt gattggagcg
100441 gagaaaaaac aggacggcta cttagaggga aacggctatc tggtgagctg gtgtatcggc
100501 cacctggttt cccttgccga cgccggggcc tatgacgagc ggttcaaaaa atggcgctat
100561 gacgacctgc cgattatccc gcagcagtgg cagtacatca tcccggacga gaagaaaaag
100621 cagtttgaaa tcttacgttc cctgatagag cgcaccgacg tggagagcct tgtgtgcgcc
100681 acagacgccg ggcgagaggg agaactcatt ttccgtttcg tctaccagat ggcgggctgc
100741 aaaaagccgt tcaaacgtct ttggatttcc agcatggagg acggggccat caaggacggc
100801 tttgcccatc taaaaccggg ggcagactac gacagcctgt atcagtcggc tctttgccgg
100861 gcgcaggcgg attggctggt ggggatcaac gccatacggc ttttctccat cctctaccac
100921 aagaccctga ccgtgggccg ggtgcagaca cccaccctca aaatgctggt tgaccgggaa
100981 agccaaatcg gcaatttcaa gaaagagaaa taccacgttg tccatatcgc ggcgggcggc
101041 atggaggcgg ccagcgaccg ctttacaagt cctgacgatg cggaggcggc taaggcggct
101101 tgcgctgggg cacaggctgt ttgtgtttcc gtcaagcggg agcaaaaaac ggagaaaccg
101161 ccccgcctct atgaccttac caccttgcag cgggaggcta accggctctt tggctttacc
101221 gccaagcaga ccctagacta tgcccagcag ctttatgaaa agaagcggct gacctatccg
101281 agaacagaca gccagcacct taccgacgat atgcagccca cagcggaaag ccttgtgtcc
101341 ggcctctggc cgttgctccc ctttgcaaag gggcttgacc tctccccgca gttgggccgg
101401 gtgctgaaca gcaaaaaggt gagcgaccac cacgccatca tccccacgat ggagtttgtg
101461 cagaaaggct ttgacgggct gaccgagggg gagaaaaaac ttctctccct tgtgtgctgc
101521 aagctcctgt gcgccgtggc cgcgcctcat gtgtttgagg ccgtcaccgc cactttcacc
101581 tgtgccggga aagagtttac cgccaagggc aaaaccatcc tgacgccggg ctggaaagag
101641 attgacctcc gtttccgtac ttgctttaag gccgatgcca acgagaacgc gccggagctg
101701 gcgcgggtac tgcctgagat caccgagggc cagacctttg acaaggcgga ggccagcatc
101761 accgagcatt acaccgcgcc cccgaagccc tacaccgagg acacgctact ttcggctatg
101821 gaacgggccg gggccaagga tatgccggag gacgccgaac gccaaggttt gggaaccccg
101881 gccacccgcg cctccattct ggaaaagctg gtgcagatgg gctttgtgga acgaaaaggc
101941 aagcagcttc ttcccaccag ggacggccat aaccttgcct gcgtccttcc ggatgtgctg
102001 acctctcccc agcttacgtc tgaatgggaa acgaagctga cggccattgc caaaggtcag
102061 gccgacccgg aggacttcat gcatgacatt gcagaaatga caaggagcct gattgtgggc
102121 tattcgcaga tcagcgagga cgcacagaag ttgtttcagg atgagcgtgt ggctatcggc
102181 aaatgccccc gttgtgggga gagcgtctac gagggcaaga aaaactattc ctgcggaaac
102241 cgggcctgcc agtttgtcat gtggaagaat gaccgatttt ttgaggaacg gggcaaagcg
102301 ttcacctcga aaatcgccgc cgctctgctg ggagacggaa aggcaaaggt aaagggactg
102361 ctctctctga aaaccggcaa gacctacgac ggcacggtgc ttttggccga tatcggcggc
102421 aagtatgtga actaccgcgt ggaacagagg ggcaaaaact aaaaaacagc aagcataggg
102481 agtgtagcca cactcccggt aaactccctg tcccggccat tctgccgggc gggaattttc
102541 actttgggaa ggcttcaatc cccctgtctc atgaaccccc gtttgtgcgc aatggattca

102601 gatttgcttt tcaacatcgg gtatccctgc tataatatgc gcaaggagta tgtcatgcaa
102661 aagaggaacc gatgaaacaa ccggataaaa aatcacgttt gaatggtatg gctgcgattg
102721 ggatggggct tggcctgatt tttggaattt tgttgaagga cctcccgctg ggccttttga
102781 tgggagcggc actggctggc tttggaaata tgacaacgaa aaggtaggcg gatggttatg
102841 aaaaaatgga tacttatttt tagcgcaatc gcattgcttt tgttttcact gacctttgtt
102901 ctatgtagcc gggaaaataa tctggcagca ctgtccagac aaagcgggat tgacttaacg
102961 atagggaaag ttgtttccca caaggatacg catggtggat ttcatggaga tggtgtttcc
103021 tacacagtgg tacagtatcc cgatgattcc atcggtgaac aaatggaaga aagtgaaacc
103081 tggcatacac ttccacttcc cgaaaactta gatactttcc tataccagcc atacgatgat
103141 gaagtatcta taccggaaat acaggacgga tactattatt tttatgaccg ccattcggaa
103201 agtaggaatc catatgatga ttccgagttg ttccaacgtt tttcctttaa ttttaccttt
103261 gcaatttatg atcaaagctc caatcaaata tatctcattg aatatgacac ataaaggtag
103321 tcgatctgtt tgattttaag aacggcttag accaggagga acaatgtata cgaacagccg
103381 ggagaacgaa gtccgtttgc tgaaacggtt atgggcagag cttccaacca tatgccccaa
103441 atgccaccaa gcggaattag agcatcttca taagaaagcg aaaaagagta atctggactg
103501 gaagtgccct gcctgcggag aaatttaccg caccataaac atgctgagtg agctgccgga
103561 ttcttaaatt tctgtatggc ctggctcttg caacctacgg ttgcgtttta gttggtagcc
103621 tgcaaccgga atgttgatgt atgatgaaat ctgaaaggag ggcgtgaaca tggcactgag
103681 cgaaaacatt aaagccaggc gtacacaact aaaaatgtcc caagagtatg ttgccgatca
103741 gttgggcatc agccgtcaag ctgttgcaaa gtgggaagca ggaaccagtg aaccgacatc
103801 taaaaactta tcggaactgg cttctctctt tgaaatgagc atttctgaat tggttgatcc
103861 tcaaacctat gccgaggaac aagagacgca ggaacaaaaa ttaagtacta agcagcgaaa
103921 tgccaagatg ctttttggca ggtgggggc tgtcatactg atgaatgctg gctgggacgg
103981 ctattcttcc ggtctgtatg gtacagactt cccgtactat tggttagcca ttttggcggt
104041 tggtcttgta ttgctgttta tcacatcaaa ggatatgggg aaaaagcata ggcttgaagc
104101 attgcagatt atacttggtc tttccatgat tttctctgtt ttctttttgc cgcgcatcat
104161 tcctttggag caggttggag tgacgtatct tctggcagat gtgattactg caatctgcgc
104221 gattctatta agtttgaaat attggcggca tatttggaag gttaaataaa acaaaggccg
104281 tgtaggcccc tcttattcac aactaaatac cagccgccca ccggcggcac aacccagcag
104341 gaaatctttg aaacggtttc ctgcttttct tatgccattt tttaggaagg atgtgacacc
104401 aaatgtctta tagctcttat gaccatgacg atttagaacc ggccagcacg atgcggatcg
104461 agcgccggat ttactttgaa agcggcaagg ccgatctctc cgaagcggta aagttgcccc
104521 ttgcggagct tctttctctg cgggcggaaa gcgccgccgc agaacaggaa gtgtttgacc
104581 gtttgaaagc gcaggccgcc gcatgggaga aacaggccgg aagaaccctg tttctggaca
104641 aagtcctgga atatgcccgg acgcttcccg tcacgcacac ttccaaccag tgggagcagc
104701 cggacaacta ccggcatatc cgcagcaaca tggtttacca gatgtattac tccatttcag
104761 agaacacccg gtacgacagc gccgcccaaa cctccgtccc ttactcctgg acattaagct
104821 ggagcctgcg caccaacgcg ccgggcagct accggcaggc caagattgcc ggacaggatc
104881 ggaaagtctt tgccagccgg gaagcactgg acaagtacct gaatggccgt atcaaggcac
104941 acgcccatta ctttacggaa atttccccgg ctatccctaa ggaatacgcc gactatttca
105001 aggtcaacgg ctgtctgctc cccggctaca ccatcgaagg ggaggaaccg gcaaaagccg
105061 cagagcttcc cacacaggag gaacccatgc agccgccaac gttcaacacc accccggaac
105121 ggagagcctc cgtgaacgaa gtatgttcca tttttcttga taaccacgcc gaagcacaat
105181 ccgacgaacc acatggctgc tggctttccc tgccaacgac cgccgaacag gtgcaggccg

105241 ccttaaacga aatccatatc accgccgaca accagcagga ttgtttcatt gccggtgttt
105301 ctgccccgga ggggcatccg ctggaactgc cggaggactt gatacaatcc gccagtgtgg
105361 acgaactgaa cttccttgcc gtgcagcttc aaaagttgga tgctgtggaa cggtcgcagt
105421 taaacgccgt catgcagtcc ctggagaaat ttcagaccat cgggcaggtc atcgactatt
105481 acaaaatacc gattgcttta ttctcattga cgccaaagac taccgcacct taggggatta
105541 ctacctgaat cattcgggcc tgatggtgat tcccgatgaa tggaaaccgg ccattgacac
105601 ggaacggctg ggccagttta ttgcccaggg ggaacaaggc acattcaccg agtacgggta
105661 tcttctgcga accggggaca aatggcagcg cgtccatgag ggccagccag tgccggaaga
105721 ataccgggtg atggcctatc ctgcgccgga aattatgcgg gaggaaagca aggtgcagcc
105781 ggagacggca gctcccgcaa aggccccgca gcctgtcacc cccattctat taaacagtca
105841 gaacagtacc gaccgtatga aagaaatcac cgaccggctg gaaatcggca tacaggagct
105901 gtttgagagt gaacgctaca aagcctatct cacctccata gcgaagttcc acagttacag
105961 cttcaataac accctcttga ttgctatgca ggggggccag ctggtagcgg gctacaataa
106021 gtggcgggac gagtttcacc gcaatgtgaa gaaaggcgaa aaagccatca agatcctggc
106081 cccggccccg ttcaaagcaa agaaggaagt gccgaagctg gacgcccagg gcaaggtggt
106141 gataggccag gatggaaagc cagtcaccga ggtacaggaa atccaagttc cggccttcaa
106201 aatcgtgtcc gtctttgatg taagccagac cgagggcggg ccgctgccct ctatcggcat
106261 ggaagaactg accggcagcg tggaacggta cgaggaattt ttcaaggcgc tggaacagac
106321 ctcccccgtc ccgatgtcct ttgaggacat ccccggcggc tcccacggct actaccactt
106381 ggcagaacag cggatcgcca ttcaggaggg catgagcgaa ttacagaccc taaagacggc
106441 catccatgag atcgcccatt ccaagctcca cgccatcgac ccggaagccc tggccataga
106501 gcaggccgac cgccccgaca gccgcacccg cgaagtgcag gccgaaagcg tggcttatgc
106561 cgtttgccag tactacgggc tggatacctc cgattattcc tttggctatg tggccggttg
106621 gagttccggc aaggacttga aagagctgaa agcctccctt gaaaccatcc gggccaccgc
106681 ccatgaactg attaccacca ttgacggcca ccttgcccag cttcagaaag aacggcaggc
106741 ccagcaggaa cagccgcaaa ccgcaccgct ggaacaggcc gctgagcagc cagacccagg
106801 cagtgcattt tccaagttgc cccccaagca acagcaggaa atgaccgaca gcgtaaaagc
106861 catgctgcaa accctcattg aagcggattt aaaatccacc ggcgaggtat cacagggaac
106921 caaggaggcg gcgcaggaac agggctttac cattgccggg gacggttcac tggaaaaggc
106981 cgaggccccg caggaagccg cctaccgtct ggaaagcggc gattacctgt atatccagac
107041 atcggaaaac ggttatgact acacgctata tggccctgac tacaaggagc tggacggagg
107101 ccagcttgac aatcccgacc tgtcccttat ggaagctgga aaggaaatcc ttgctgccca
107161 tgagctaccg gctgggacga tggaacccct gaccggcgac gcgctggacg gttttctgga
107221 agccgccaag caggccaacg ccattcccca gccgcaggca tggaacggaa tcgacggtat
107281 tttgaacggc aatcccttga tgccggatgt atccccggct gaccgggcgg ccgccctgat
107341 cgcgctggcc gaacaaagcg ccccgctttt gggaagggaa gaacggaagt tgattatgga
107401 ctatgccgac gccgtgaacg acccgcaaaa aatcataggg ctgatcaacc acctatgtga
107461 acaggggtat gaacttcaac acggcaagat ggatgacgtt gtaaaaagcc agattgtaag
107521 tgagatggcc gttgccaggg cggaacagac catcgcctgt gacccggaag ccgagccgat
107581 tgtcacgatt ctttggagtg aaagccccca cttaaaagat ggccagcaaa tgccattaca
107641 tgaggccgac gctgtttttc agtcactgga cgaggcccaa cgccgcggcc gagaacagcc
107701 gggctatacg ggaagctggt acgacaagac caggttccgc attgacttta ccatgcaggg
107761 ccagccggac aactacgagg ggcggcagga ctttggcgat ggggacggtt ccctgattca
107821 gcatatccgg ggctgccacg aatactacga acaggatgaa agctggaaaa accatgtgct

107881 gcaccatgag gggccggagg catgggaagc ggacaaggcc cagcgggata tgctgctgca
107941 tacctttgtc ccatatctca aacagcactg caacctttcc cgtatggaac aggaggccca
108001 acacctttta cgatccggcg atcccctgtt gccagagcag accgcctatt tggacgcctt
108061 ggtggagtat gtaagagagt gccgtccgct tctcaatcag ggagaatcgt taccggagcc
108121 gccgcagctt ttcgactttg ataagagctt gcaggactat aaggcgcagg tcaagaccga
108181 gctggaacag gaagccgcca ctgccgggct gacggtggag gaatatattg actctggttc
108241 ctatgccccg gcgcagccta acttttccat ctaccaagtg ccgcccggcc cggaggggcg
108301 cgacttccgc taccgttcct atgaggactt gcaggcggac gggcttttgg tggacaggaa
108361 gaattaccag cttgtctaca ccgcgccgct ggataaggac accacactgg acgagattta
108421 ccgccgcttc aatatggaac atcccgccga ttacaaggga cgttccctct ctatggggga
108481 tattgtggtg ttccggcagg acggggaaca gacggcctac tatgtggacg agggggccga
108541 ctaccggcag gtgccggagt tcttctccca gccggaaaag cagctcacgc cggacgagtg
108601 catgaccggc gaacaaatcc agacgccgag gggacgtttc tatctgaccg acaggagccg
108661 ggaacagatg gaggctgccg ggtatggatt ccaccaccag tcggaggacg gcaagtatct
108721 tattatggca aacggcacaa gggcctttgc catcccggcc cagcaggaca gccacatcaa
108781 aaccgctgaa atgtccaccg agcagaacta caacatgatc gacggcatga tgaacaacgc
108841 cccgtctatg gaggaactgg aagcaaaggc aaaagcaggg gaacagattt cccttttgga
108901 tgtggccgag gcagccaagg cggaggccaa aaagcccaag caaacgcgca ggacaacaca
108961 aaaaccaaag aaaccgtcca tccgggcgca gcttgccgcc gccaaggagg aacagaaaaa
109021 gaagtccccg gcgtgggaaa aatcaaaaga aatggaggta tgaggaatgt atttcaccgt
109081 ggaagaagaa aacctgctct gtctgtatca taatgccgaa cgccgcagga cagcagcaaa
109141 tctccggaca gttctgccgg atatggacaa ggaaatggct acgctggcct gccagaccgc
109201 cgataagctg gacgccatga gcgacgccga ctttgcggcg cagcggttcc acttcacaga
109261 cgagtaagca aaacgccggg cgcgagggct atatggccct tgcgtccggc attttgtttg
109321 cccagcatgg gcgttttcta atgggtgaaa gtcccaagtg cgcgtaggca acaacgaagc
109381 acatagccga acagcaaggg tgtccgccgt gagacggaat ctgaaagaag ctgtaagcaa
109441 acctctgacc tgacggacag gaaccgcata taaggctcgg aaatacggat aaggtggcaa
109501 aaggcactga agtccaaaag gttgccggaa gtacgagtaa atgcggcagg tacatggagg
109561 aaaagaacac gcaccttaac tggggaggtc tcacaggcgg tctcattagc cgtagtaaca
109621 acgaattgtg agaagtcagc agaagccata gtagtgagga agttcctgta atggggacgg
109681 agcgaagggc tgaacaatca atcagttgaa gtacgttcca cttcgtagcc ggagcatacg
109741 cctgtcgatg acttcaaagg cggcaaaggc aaaaggggca gaaaggaaac aacgcatgga
109801 cacaagcagt ctcatggagc agatattaag cagggataat ctaaatgcgg cgtatctgca
109861 agtcgtaagg aataaaggag cggcaggcgt ggacgggatg accgttgaag aacttggcgc
109921 atatctttcg gaaaacggcg aaaacattaa ggaacagttg cggacgagga agtataagcc
109981 gaagccagtc cgcagggtgg agatacccaa acccgatggt ggtacaagaa atcttggagt
110041 gccaacagca gtagaccgct ttgtacagca ggcggtggca caggtgctta ccccgatatt
110101 tgaggagcag tttcacgacc acagctatgg attcagaccc aagcggtgtg cacagcaggc
110161 agtccttaaa gcattggaaa tgatgaatga cggacacaac tggatagtgg atatcgacct
110221 agcgaaattc tttgacacag tagaccatga caagctgatg acgattttcg gacggacaat
110281 aaaggacgga gatgtcatat cggtggtaag aaagattctg gtcagcggcg taatgattga
110341 tgatgagtat gaagatacgg tagtcggcac accgcagggt ggaaatatct cgccgctgtt
110401 agcaaatatc atgttaaatg agctggacaa agaactggaa gcaaggaggc tggatttcgt
110461 ccggtatgca gatgacctta ttataatggt cgggagcaga caggcggcag agcgggtaat

110521 gaagagcgtg gctcggttta tagaggaaaa gcttggactg aaagtgaacg cggaaaagag
110581 cagggttgat aaaccaaagg gcattaagta tctggggttt ggattttact atgactcatt
110641 tgccaaaggg tacaaagcca gaccacaccc gaaagcggca gcaaagttca aggcgcagat
110701 gaagaaatat acaagcagga gctggggagt gggcaatggt tataaaattg ggaaactcaa
110761 ccggcttatc cgagggtgga taaattattt caaaatcgga agcatgaaaa ggctgtgcgc
110821 aaaaatggac ggacagattc ggtatcgact gcgcatgtgc atatggaaac actggaaaac
110881 gccaaagaac agggaaaaga atcttatcaa acttggtctg ccgccaaatg cggcacatgg
110941 catttcatat gccaaaggat atgccagagt gtgcagaagc tggaatctcc acatttgtat
111001 cagtaaagag agactagcta agtttggtct tgtatccatg gaagactact acgccgaaaa
111061 ggctgttaca tgttaagttg attgaaccgc cgtataccga acggtacgta cggtggtgtg
111121 agaggtcgga aggcgaaata atcgccttcc tcctactcga ttatatggtc ttaaacgctg
111181 aaatcatatc gcgaatattt tgaaatcttt tggttttatc cgggttcaat cctctctcaa
111241 caaaatttcg taaattctga ttcgtaattt tttcagtatt agttttacct gtcattacaa
111301 aataaataac tctggtgaga gcataagttt cgtgaactat gccataagtg ttaaacccct
111361 cgacaaccag tgcaggatca ttaaaataac ccttgaactc cgtattaacc gttgttagcg
111421 tactatcagg aatttttaca agtccaaaat ctgaaagttt aactacaagg gtatcgtcgt
111481 actcttttat caaaatattt ttagggctta tatcacgatg cagatgtcct tttgaatgaa
111541 gataatcaaa agcgcgtaat atttgttggg caattccttt tcgctgtata attgttaagg
111601 tggagttatg agcagcgata tatccgtcca gtgtataatc catatattcc ataatatact
111661 catttttatc tggattgtag cagtaaactt caagtatgta tggagaagat agatcattca
111721 ttacatcaaa ttctcttttg aaacgggcca tttctttatc tgtaagttcc ttttttgcac
111781 gctttaaaat aaaaggccga ttataaaaag tgtccttata tttataaaca ttagcataag
111841 aaccgcttcc gatcagcttc aaatcaaatg taaaatcttg ttgcttatgt gaaatggtta
111901 tgctactaag aggtaaaaaa ataggaagtg tataatataa ttcgaccttt gccatattgg
111961 gcggtaatga actgcctcca cttgaactaa ggaaatctcg acagcgtgta agtaactcta
112021 aatagtatgg gtctatttga aaggctaaag gcgttgcttt tagtgaactt actaagctga
112081 atataatttc gattctctta attaaatctc tgctcggttc tgcccaaaag tgtgcttcat
112141 gttccccagt tggtagcctt tcgttcattg ttctaaaaag tccaaccaga tcatgatgta
112201 aggtcataag tatttcccga agtttttgat ttctaaatga tttatataaa tcggagtatt
112261 ccacatttat ttgactacat gataataatt ctctgtactg tgattcaata tagttttcaa
112321 tattcataca agttccaccg tttcccgctt tgttttactg tattatatca tgcaattaca
112381 taatcggcaa ttatccaaaa cgccgggcgc ggggctgtat tgcctctgcg tccggcattt
112441 cttggtgcgc ccggcggcgc acgtttctaa ccggtgcaag tccggaatcc gcccggtggt
112501 gggaaggata tagccgaagg caagggtgtc catcgtgagg tggaatctga aggaagctgg
112561 atatggggaa catactaacc catgggcaaa tctctggtct gacgaacaga aatcacatca
112621 ggccatgcgt gagggtaagg ctgccaaaca agtcgaagcc caataaccac acggaatcat
112681 gtatggtaaa tgtggcagat agatggagag aaagaaacgt gtggtacccg gggaggtctg
112741 tgcggaatgt cctgaaaggg ataaccacta tcgtgaggta gtgctgaacg tacagaagtc
112801 agcagaggtc gtagtagtcg agagacgaag gaccgaatca acaggagtct ttagtacaac
112861 cgggaaagga ggaaagaaca ttgggtacag aaaacagaga tagctgctca caaagagaca
112921 gcgcggaacg tgaagggtat gtgagagcgc accgttcttt ccgccggata tggaaggaaa
112981 gagacagtgc acagccggaa ctcttagaga agatactgaa taaggacaac ttaaacagag
113041 cttttaagag agtaaaggcg aataagggag cgccgggaat cgacggaatg accgtagagg
113101 aaacaggtga ctgcctaagg gaaaatcaaa aggaactgat agagagaatc aaaaggggaa

113161 aatacacccc cgacccagta agacgagtag agattcccaa gccagaggga ggaatgcgaa
113221 agctgggcat tccaaccgtg aaagaccgta tcttccagca agccatagga cagcagttaa
113281 tgccagtcta tgaggcgctg ttctcggaga acagctatgg gtatcggccg ggaagaagcg
113341 ccagggacgc cattttaaag gtaaaagagt atgcagaaca gggatacacc catgccgttg
113401 cattggacct gtcgaaatac tttgatactc tgaaccaaga aatgctatta aatatcctgc
113461 gcagagatgt aaaggatgaa agggttatcc aatggataaa gagataccta aagagtggtg
113521 tcatggaaaa cggtgtagtc atggagacag aggaaggatc gccgcaagga ggaaatctat
113581 ccccgctgct ggcaaatgtc tatctgaacg agttcgacca ggagtacgag aaacgaggag
113641 tggcttttgt acgttacgca gatgatatag tattgttagc taaaagtgag agagccgcaa
113701 agagacttct ggaaacaagc acgaaatatc tggaagggac gctgaaattg caggtgaacc
113761 agaagaaaag ccgtgtagta agtgtatttg caatccgaaa ttttaagttc cttggcttca
113821 cattgggaaa gaacggaaag ggcatttatg tccgagttca tgggaagtca tggaagaaaa
113881 tgaagtcaaa actgaaagac cttagttccc gaaggtctgt tcagagcata cgcccatcgt
113941 tagaaaagat aaaggtgtat atgcggggct ggctgaatta ttatggagta gcagatatga
114001 agaaccgaat agagggactg aaccaatggc tatatcacag aatccgaatg tgtatatgga
114061 aacaatggaa gaaaccgaaa acgaaggtaa agaacctgct gaaaatggga gtgccgaaag
114121 aattggcatg gaaggcaggg aacagccgaa gaggctattg gtttaccaca caaacggtag
114181 cggtaaatat ggcaatgaca aaagaaagac tgataaacag aggctattat gatttagcca
114241 cagcctatca gtccgtgcac gtcaaccgtt gaaaccgccg tgtaccgaac ggtatgcacg
114301 gtggtgtgag aggacgggag ttaatcactc cctcctactc gattttgtcc attttcccgc
114361 atttagaacc ccgttttgcg gggaacagga taactttatc catctgcctg ctgtccatgc
114421 ggaaaagccc ttgatttccg tgtattttta ccgtcctaaa tgcgtagaca ggaggtatct
114481 tttccgctgg cgttccgcct ctttctttcg gcgtacctgc accgcacaat ccgggcagta
114541 tttggcccgg ttggagttgg gcgtaaaggc cgccccgcag acggtgcagc gtttgatatc
114601 atcccggcct ttggtgattt cggcgcacag ggccgcgtcc agcggcagga cggccacctt
114661 gaaccaccgg cacagcagcg aataggaaat actctgcaca cagacacatt cctccccatc
114721 gtccagcagg aggcagtttc cgttacagta gttgcagcac tcatgggtga gcctgcgggc
114781 cttgcggtat tgggggtaac tcattctcgg tatattcatc gttcctgcgt aagctccttt
114841 cccggcgctg tccgcaaaat gctgtccaca ttctgtttga caatgccgta ttctttcaat
114901 ttttgtttcg cttcgctgta ttgtgcggac agccgttccc gttctgcgtc cagcttttta
114961 tactcggttt tgagggcgta caggttgggg agctttgtga tcccagcctc tttgagcgcc
115021 tttgccgccg cctcatacaa tatcagttgg ctttcatgct gccgctggaa cgcggccttg
115081 tctttgacat ttcgcagttc cagcaccaca ggccggagct gcttgtaggc ggagaggttt
115141 tttatcagca gcgccatatc cccaaggcgg cgttctgcgt ctttcagggc ggaggccgcc
115201 gcgtcgccca ccgcgctgat ctcggccagc ctgctttcca agccctcgta gctgtcaatc
115261 tggtgttccg tgatgaagtt taaggagttg gcggcctgtt tcaagttgtg cagcttcgcc
115321 caccgcgcat agcccgccga ctgctgggcc ttgatactgt cctccaaggc gatccgcagg
115381 gagatttccc cggcgcttct tctgcgcggc cctcggtcga tggcaatgcc cttgattcgc
115441 tgggtgatcc gttcctcggt gtaatcctcc cccaaggtct tgcagcgggt gaagcgttcc
115501 tgtccggggg cgcggaatga aataaacttg ccctgtttga cttcatagcc ctgcgcctcc
115561 atgagctgca aaaagctgtc aaaatctttg gactgcggga tggccgcgtc gatagcaatt
115621 tttagttttg ctttccagct cttgcccttt ttctgtgcgt cccactcggc ataggtcttg
115681 cccttgtcct tgccgggctt taccacagac aggccgtgtt ccttgcacag ccggtcgctg
115741 gtgcgccgga taaaggcgta gctctgacgg ttggaaatgt acttgcgctg gttcgccata

115801 tccacggcgc agataatgat gtggttatgc aggtggccct tgtcaatgtg ggtggtaatc
115861 acataggggt atttgccttg cagcacttca tcggcaagct gccgcccgat ctcgtgggcc
115921 tgctccgccg tggtttcccc cggttcaaag gcttgtatca ggtgatgtgc cagattgttt
115981 cccttttggt acgcctgatc cagcagcatt tgaaattcta tatccgccgt ttcataggaa
116041 cagccatagg aagaaacgaa cagcttttca tcggtcttgt ccgggttctc gatgtagtcc
116101 agcgcctttt tcagcgttga cttgatagga tggatctttg taactgccat atctgtgcca
116161 gcgcccccctt gatgtcctcc aagtcctgcg cgtacacagt cccggtactg ttgatacgcc
116221 ttgcaatctg gttgatgttg acgccgattt tttgcagctc tgcggtctgc gcccggacgg
116281 ggccggtgtc gatatggacg atatagccgt cgatcaacat tttccgggcg taggcagaaa
116341 agttctttgt gccaagctgg gccatcttct gttggatcag cgcccgttcc tccggcgtca
116401 ctgggacgcg cagcacaaat ttccgttttc ggtttgccat tggttcacct ccgttcattc
116461 aggggtttgg ggacttcccc aagatacaaa cgggaagccg tttgccggga gtgtggctac
116521 actccctatg cttgctgttt ttcttttcct ccactcctaa tactctgaga agccatcaaa
116581 ctacccgcaa aatagcgctt tggaaagaaa aatttgcaaa aaaataagag gcccttggcc
116641 tccatacaac aaatgataaa caaatgccga atccgtttga cgcgttcgct tgacggtccg
116701 tgtctacagc acatcgttaa acttacccgt tcttttgttt tacctttgca gagaatacgg
116761 tataaaagat tctgtcatac tactttcata ctatcaattt agaagaggta aaacacgatg
116821 aaacaaaatt acaatgaaat cttgcgggag taccgcatct acttgaccga acatgaaaaa
116881 tctcacgcaa ccatacaaaa gtatgttcgt gagttggttt ggtttctttc atttttacag
116941 ggggaagaac ctacaaaagc aaaggtgctg gaatatcgag agcaactgca acagagccac
117001 catgcccgca ccgtaaatgc caagctgtcc gccattcatt cctatctgga ctacctgggg
117061 ctggcggcct gcaaggtgag gtttttgaaa atccagcata cggtatttgt ggatgacagc
117121 cgagatttga ccgaagcgga ataccaccgc ttgttagacg ctgccaaaag gaaaaaggac
117181 agccgcctgt atcatgtgat gcttgccatt tgcaccaccg gaatccgggt cagtgagctt
117241 tcctttctga ctgtggaggc attgcacaaa ggaaaagcgg aaatccgcat gaaaggcaaa
117301 atccgcacca ttctcttgac aaaagagctg tgccggaaat tgaacgccta tgccaaagaa
117361 aaaggcatcc gaacggggta tctgttctgc acccgtaccg gaaagccctt agatcgaagc
117421 aatatctgtc atgatatgaa aaagctgtgc cgggcggcgc gggtaaatcc tgaaaaggtc
117481 tttccccaca atttgcggca cttgtttgca aaatgctatt atgccatcaa aaagaatctg
117541 gcatatcttg cggatatttt aggacacgcc tccgtggaca ccacgcgtat ctatgtagct
117601 atgggcacac gggaacatga gcgaacctta cagcggatgc atctgatctc atagcaccac
117661 ggaataatga ttctgtggtt ccatggcccc cttgctcagt tacaagtaga actgattagc
117721 tctattataa gtgtatcaga gcagattttc aaccatttat tacatataga acagcaaaat
117781 aaatacgaaa aatagacctg caccgcgaaa tatgtgcgca ggtctgcttt ttatgcctgt
117841 cagaacgccc ttagttgatt aaaataataa ttttcttcct agaagcagta acgattaccc
117901 cagaatcatt attctgcggt aattgaatcg ttcagaaaag gaaagtttgt ggttaggata
117961 ttccctttga aatggcagga ggtgaataaa gtggaagaaa gatggtcggt agaattgttg
118021 gagtgcttgg cccagaaggc cggatgtggg tatctgtcgg atttgcggta tctcagccca
118081 tgggagcaat tccatctgtc cagggagctt ttacatgttc cttcggaagc cttttcgcta
118141 aaaacgtgga atgacgcctt ggcttatctg acgggtcttt cccagcaagc ggatgcatcc
118201 gccgcgaaag aaacgttgat gaagcatcta tcggaacact gcagcggttt gcaacgtcat
118261 ccgagccgag aatgaatttg aatagaaagg tggttagaat gctatgcaaa ttactgttgt
118321 aggcgcaggt tatgtggggt tatctcttgc cactctgtta ggccaaaagc atgaagtaat
118381 agtgcttgat attgatgaag aaaaggttgc gaaagttaat tcccgtattt ctcccgttca

118441 ggacgtatat ctcgaaaagt tttttgccga aagaaaacta aatttgacag ccactgtaga
118501 tacggcaata gcatataaag acgccgagta tgtaattatt acaacaccta ctaattatga
118561 ggatgaaacc aatagctttg atactcatgc ggttgattcc accatagaaa tttgtatggc
118621 aaataatgat cattgcatta tgatcatcaa aagtacggtt cctgttgggt atacaaggag
118681 tgtgcgaaaa aaatataaca ccagccacat tctattttcc cctgaatttt taagagaaac
118741 gaaagcccta tacgacaatt tgtacccttc cagaattgtt gttggtacag atattgcaga
118801 ccctgctatg gttatccatg cacaggtttt ttcaaccatc ttgcaagagt gcgcaaataa
118861 agaagatatt cccgtcttga ttatagggct tgatgaagca gaagccgcta aacttttcgc
118921 taatacttat ttagccatgc gtgttgcgtt ctttaacgaa ttggatacct ttgcagaaat
118981 gaatgggtta agtacaaaga acatcataga tgcaatttgt catgatcctc gaattggtaa
119041 gcattataac aatccatctt ttggctatgg tggatattgc ctcccgaaag atacaaagca
119101 gttgaaatca agttttcatg atattcctga aaatttgatt accgctgtgt gtcaggcaaa
119161 ccacaccaaa aaaggccatg tcataaaggg aattctgaat aaacatcctg gcacagttgg
119221 tatttaccgc ttaacggcaa agtctaattc tgacaatttc cgttcaagtg ctgtttgggg
119281 tgttatggag gggctttcca aagaaaaaca agaaattgta atatatgaac ccttattagg
119341 cgatgccgca gagtttatgg gttataaggt tgtacattct tttgcggagt ttaagcgttc
119401 atgtgatgtt attgttgcta atagagtttc cagtgaactc tcggaagtta tgtataaggt
119461 ttatacaaga gacatctttg gaagagattg acccggcgag gcggtgattg taatgtacac
119521 atttttcttg atggtcgagg attatatcaa aactcataac cttcacctct ttatgttttt
119581 cttcgcattc attttcattc gatggggaat agtatttttc catgctattc gatataagcc
119641 atatgattat gaagataaag aaataaatta ttttacatcg gtgctgctcc cagtcgttga
119701 cgaaccatta gatctatttt atagtgttct gatgaaaata gcacgccaga atccgtctga
119761 gattatagtt gtcattaatg ggccaaaaaa cgaggggctt gaaaacctgt gtgtagattt
119821 caacaggaat ttaccaattt gctttactcc tgttcagcat tattatacac ctgttgccgg
119881 taagagaaat gggatccgtg tggcaatgga gcacattaat ccgaattccg atataacagt
119941 gcttgtggat agtgatacgg tatggacaga ggatactttg tccgagttgc tgaagccttt
120001 tgcttgtgat caaaaaattg gcggtgttac aaccagacag aaaatacttg atcctgaccg
120061 caaactcgta accatgtttg caaatttgct tgaagagatc agagctgaag gaacgatgaa
120121 agcaatgagc gttacaggca aagtgggctg ccttcccggc aggacaattg ctttccgtac
120181 tcagattttg atagatgtga tgtatgactt tatgaatgag acattcatgg gatttcacaa
120241 agaagtatcc gatgacagaa gcctcacaaa cctgacgctc cgaaaggggt acaaaacagt
120301 tatgcaagac acctcagtaa tttatacaga tgcgcccacc gaatggaaaa aatttatcag
120361 gcagcagtta agatggtcgg aaggttcaca gtataacaat ctgagaatga ctccttggat
120421 gttaaaaaac gccaagctga tgtgctttat ttactggtca gatatgatta gtccgatgat
120481 gctggttagc gtttatgcca atactataat ttgcaaggtg ttaaatatac ttggctgcgc
120541 gattccaaca ttggcatata ccgctccatg gtggcagatc attctcttta ttcttcttgg
120601 ttgtatcatt agttttggtt cacgaaatat taaggttatg agaagtgtga agtggtatta
120661 tacattactg cttccagtat tcatccttgt tttaactgtt gtcatggttc ctatccgatt
120721 gctaggcctt atgctctgct ctgacgatat ggaatggggc acccggaaat tggaggagga
120781 caatgataaa gtggaagttc cttaattgca tttgtatatt cgcggctgct atatttatat
120841 tatgggttgg ttgcaacgaa cagcctgtcg ataccgagaa ggccgatggc tctgacagca
120901 ttgcgtctat tgaggctccg agtaatagtc aagctacact ggaagaacca actacggaat
120961 ctattgtaac agaatcctca gttgtcgagt ctgaatatta tcctattaca gagggacaaa
121021 gatatttggg cgtttatgtt caaggccccg aagagactga tgttttagct gacagcatca

121081 atactttagc ttggtttgat cgttttgatc agacaagtga ttacaagatt tctttatgtt
121141 tagacgacaa taaatatata gcgtttatca ctttgcagcc taccgactgg gacttaaaat
121201 tggtttctga cgggtattat gatgatttaa ttattgaata ttttaagaaa ctctcttcgg
121261 ataacagagc caatacagaa ctttttgttc gattagcgca tgaaatggaa atgaggccct
121321 cttacaaatc tggttggtac agttggcaaa cagatgatgc tcacgcatat gtaaacgctt
121381 gggttcacat tgtgaacttg ggtagggaat atgctcccaa tgttaagtgg gtatggtctc
121441 ccaacagggc agacgaatac acaacgaaat attatccagg tgatgagtat gttgattatg
121501 tcggattgac attgaataac acattggatt cccgtgagag ttttcagcaa ttctatgaaa
121561 atgaaggtca aagagactat ttggaagcat ataataagcc aattattttt ggagagatcg
121621 cggagcatag caccagtgat gaagttcgca atgaatacat acaatctgtt tttgattatc
121681 ttggaaccta cgataaatgc atcggtttta ttttcttaaa tcaagatata gaaagtgcaa
121741 gacaatacaa gtttacagat tgcgaattga tattagatac atttattgag aatgcgaggg
121801 attacatttg tgcgaaataa gaagtttaca aagtatcaga tcatgacgag agttggaacg
121861 attatcagca ttttactgtt tgttgtcctt ggattttgta tcgaagttat tttctaaaaa
121921 aatggagaaa gtgaaagatg gaacatcaaa ccaaaggaac gttgcgcaat caggccgtca
121981 ttacatcagc cggatcgccc cga
//
DEFINITION Integrative and mobilisable element IME_2
fasta_record 1..9797
/ID=B77|SC|contig000001
/colour=5
CDS 969..1193
/color=7
/locus_tag="B77_00649"
/note="transcriptional repressor DicA
Helix-turn-helix, protein motif:CLUSTERS:PRK09706 protein
motif:TIGRFAMs:TIGR03070 protein motif:Pfam:PF01381.16"
/product="transcriptional repressor DicA
Helix-turn-helix [transcriptional regulator]"
/"ID=B77_00649
/protein_id="gnl|SC|B77_00649"""
/gene="dicA"
/codon_start=1
CDS 1433..1858
/ID=B77_00650
/color=1
/locus_tag="B77_00650"
/note="hypothetical protein"
/product="hypothetical protein"
/protein_id="gnl|SC|B77_00650"
/codon_start=1
CDS complement(1807..2673)
/color=2
/locus_tag="B77_00651"

/eC_number=2.7.7.-
/note="Aminoglycoside 6-adenylyltransferase Streptomycin adenylyltransferase, similar to AA sequence:UniProtKB:P17585 protein motif:Pfam:PF04439.6"
/product="Aminoglycoside 6-adenylyltransferase Streptomycin adenylyltransferase [antibiotic resistance]"
/"ID=B77_00651
/gene="aadK"""
/protein_id="gnl|SC|B77_00651"
/codon_start=1
CDS complement(2847..3362)
/ID=B77_00652
/color=1
/locus_tag="B77_00652"
/note="hypothetical protein"
/product="hypothetical protein"
/protein_id="gnl|SC|B77_00652"
/codon_start=1
CDS complement(3399..4199)
/color=7
/locus_tag="B77_00653"
/note="HTH-type transcriptional regulator immR, similar to AA sequence:UniProtKB:P96631 protein motif:Cdd:COG1426 protein motif:Pfam:PF12844.1"
/product="HTH-type transcriptional regulator immR [transcriptional regulator]"
/"ID=B77_00653
/gene="immR"""
/protein_id="gnl|SC|B77_00653"
/codon_start=1
CDS complement(4266..4742)
/color=8
/locus_tag="B77_00654"
/note="Pyridoxamine 5'-phosphate oxidase, protein motif:Cdd:COG3467 protein motif:Pfam:PF12900.1"
/product="Pyridoxamine 5'-phosphate oxidase [oxidase]"
/"ID=B77_00654
/protein_id="gnl|SC|B77_00654"""
/codon_start=1
CDS 5120..5533
/ID=B77_00655
/color=4
/locus_tag="B77_00655"

/note="Conjugal transfer protein Cysteine-rich VLP
containing protein"
/product="Conjugal transfer protein Cysteine-rich VLP
containing protein"
/protein_id="gnl|SC|B77_00655"
/codon_start=1
CDS 6223..7167
/color=9
/locus_tag="B77_00656"
/note="Plasmid recombination enzyme Mob_pre, protein
motif:Pfam:PF01076.13"
/product="Plasmid recombination enzyme Mob_pre"
/"ID=B77_00656
/protein_id="gnl|SC|B77_00656"""
/codon_start=1
CDS 7240..7434
/ID=B77_00657
/color=1
/locus_tag="B77_00657"
/note="hypothetical protein"
/product="hypothetical protein"
/protein_id="gnl|SC|B77_00657"
/codon_start=1
CDS 7535..9157
/color=9
/locus_tag="B77_00658"
/note="Serine recombinase, protein motif:Pfam:PF07508.7"
/product="Serine recombinase [recombinase]"
/"ID=B77_00658
/protein_id="gnl|SC|B77_00658"""
/codon_start=1
CDS 9273..9797
/ID=B77_00659
/color=7
/locus_tag="B77_00659"
/note="HTH-type transcriptional regulator immR"
/product="HTH-type transcriptional regulator immR
[transcriptional regulator]"
/protein_id="gnl|SC|B77_00659"
/gene="immR"
/codon_start=1
BASE COUNT 2852 a 2117 c 2311 g 2517 t
ORIGIN
1 caaccgtcag ttaaaggcaa aacggccata caaagatacg atttttcgaa tgttgtatca

61 tgataaagaa aatctgctat cattgtataa tgcagaagat ttgcagattg tcacattaga
121 aaatgcaatt tatatgggaa tgaaaaatga tctggcgttt attatggata tgaatttata
181 tttgtatgag caccagtcga cgtataaccc gaatattccg cttagaaatt tattctatat
241 tgcggatgaa tatcaaagat tagtagttcg gaaatcttta tattccacgg taatacaaaa
301 gataccgaca ccaagattta ttgtatttta taatggaacg aaaaaggtag acgattataa
361 tgaattccga ttatcttcag cttatgagaa tccgatggat gatccggatc tggaattaaa
421 ggtgactatg ttgaatgtaa atgatggtca taacctagaa ctgatggaac actgcagaac
481 attaaaggaa tatgccaaat atgttgcgag ggtaagaaaa tatgtgacgc agaatattcc
541 tctggaagag gcagttacgc gagcggtaga tgaatgcata gaggaaggaa ttctggcaga
601 atttttggtg aagaataagg cagaggtgat caaagtgagc atctatgagt atgataaaga
661 atttgaagag aaaaaattac ggaaaacgga gtatgaagct ggtgtggagg ctggagtaga
721 agctggaatt gaacttggag aacggagtct tctgaaaaat cagattagaa agaagttgaa
781 aaaaggaaag tctatagaac aaattgcgga tgaattggaa gaagaagtaa ctacaattca
841 acgaattata gaagaaataa aaaaagaaaa ataagtatat gatcaggaag gctgttgcgt
901 ctgcaatagt cttcttttac gattcagagc tttttacctt aaaattcaaa aagtaaagga
961 ttcgtcgatt gacgtattta acaaaaaatg tatatgatgt aattaaacaa agaagttcat
1021 tcggcttcaa agagtggaaa aaggaagtga attacatgga tgcaaaagta tttggaaaat
1081 ttatcatgga gcggagaaag cagctgggta tgacacaggc ggaattagca ggaaagataa
1141 atgtaacgga taaggcggtc agcaggtggg aaagaggggg ttcacttaga taaagatacc
1201 acaccaatcc cacgctgact tttgctcaac caatacagcc ggagggctgg ataacaagaa
1261 aaggcggtat gcgccccgtg gaggggtagc atatcgcctt ttcttgtggg ggtttccaaa
1321 gggggcaacg cccccatttg gcacacgact ttgcgaagca aagtgtagtg tgttatacgc
1381 tctgtcggcg ttgtcgtgaa aatgccgttg ccgccgggga gggcaagggc atttgaagcg
1441 gcaagatttg aagcggcaag aaaacagggc tgtgcttgcg tggcgtggag ccgcaagcgc
1501 aggtctggtt tcatcagcag acagggcgta tatgaaagcc cccgtccctc gtcctacgct
1561 ccgacttgtg gacaaagtgt cccgaagttg tggccacact cccggaaaag tagcaggaca
1621 gcgggcaacc gtcaaggctg aaatgaacgg ggttacaccc ggccttgacc gccgcccgcc
1681 gccccgctgg atgggtggac aaggcggcca atccctcaaa gtgcttggcc gcgctctttc
1741 tgattttgag gtattcagtt tttcaaaatt gaataatcaa aataaatttt ttcgattgaa
1801 aaaattttat ttgttatcat cttcaatgcc atattttttc ttttgccgaa tcacataatt
1861 actgattttt tcatcatata gtggatactg gtaatccaac tggcatgcca cttctgacga
1921 aacctcccgg aacaatgcca tacattgttc aaaggattcc cacatatggg gataggaatc
1981 catattataa gtggacataa gtcgttccca caattcctct gggacatatt gcttcataaa
2041 tttatagttt tttcccaaac tgaaatgaaa tccctgtttg ataccaatca gccaggaaat
2101 catgcgaagc aattcttttc gtactatatc attcatatgg tcaatagcaa aaagaatttc
2161 tttgcggcat aagcccttta ctacatatgt tgtagtattc caaaattcat tacagcaatc
2221 gtcaaacatt ctttgagtag gcttctgcaa gtggtagtct atatccgttg gtattggcgg
2281 ctttacgata cggttgtctt tatccaacag taactttacc agtttatccc atgtaaaata
2341 ctcgtctatc agttccagcg gcagcaaagt taaatctatc ttaacatcat cagtaaacag
2401 cattaaatat gaaaatccct tttctacagc tggaaataat tccatatctt ccggcttttg
2461 cagaatcaac ctttcaccaa atatatctag ccatttatca tcacttgtga agctgtccat
2521 atccgtaaca aaaaaagtaa tatcaaaatc ctgaaaatca tcaggcggaa tatttgtatt
2581 tgttctagat ccttccaaag taaccatgcg aatgcgtttg tctgtttttg caaaattcaa
2641 aacaatatca taaacttcct tttctgatct catttttatc tcctccaatt attgaaatag

2701 gtgtgaagcc attttagggc tttcccgcct tgattaccct ttagcaaaga cgggcgggac
2761 tgtcaacggc ggcgcatgaa atgcgccgtt catcttgacc gttgactggc tcggctggct
2821 ttgctatttt ctcgatgaaa aacaatttat tcgtatttat ttccatcatt gtcatagtaa
2881 gccacatgaa tggaatactc tttttctggg tttttaattg agataatatc catatcatct
2941 gtgttgtatc tcaatgtatc ttgcacttgt ccatttattg tataagtgat ttcagcatac
3001 tctgttttag caccattaaa ccaaattaaa tcataccatt ctccgtttat tgcaacgcca
3061 ccgacaataa tctcatcact atttcggttt gttgatgtac taaatttgta gtctccattt
3121 cctgttggtt caaagatagc aagtgtagat ttattatctg tactgtatgc tccactcacg
3181 atataaccgc caagttccag ttcttttgcg atagtccatt cattcccttt tgaaattgcg
3241 gtgttcaata tgctttctct gctggtaatg gtatctcttt tgggagttaa actaaagaaa
3301 tataaaaaag cggcagcaac gcaaataaca gtcgctgaaa ttatcagtaa ccactttttc
3361 ataaccaata cctccatata atctgattta ttattctctt acttgcgtcc ccgctgcggc
3421 tttggatttt tcagcaaatc cgatttctgt gcaatctttt tcaaccactt cttttcttcc
3481 tctgacagtt tcccatagtt cagcttgagc cgcttacaga taaatgccat agccgcctgc
3541 tccgggtcgc tgtcctctct ggtggtttcc tctgcggtct gcaaaaaatc ttccagagga
3601 ttgtcctccg ggacgctgaa aaaatcgtcc ttgtgggctt cccgcaggtc aagggctatc
3661 ttgtttatat cctgctgtaa cacatagcgg caaaagctgt catcgtcaat gtgggcggcg
3721 atcagctgcc ttaactgcgg gtcggtcaga ccgggattgt gctgcttcat aatggttgcg
3781 ctcacagcgt ccacaatggc gtttgccccc tgcacctgtt tgccagcgat accgtttaca
3841 tagatttcaa ggtcagccat caaccgaggg aaatccgggt gtgccgccag ttcacacaag
3901 agggaattat ccaccagccc actttttcaat aattcaatca tttcatcact caaacgcagg
3961 tcggcaagat cggcgtttgg gtgatttttt gtgtgagagc ggcacagaag ataatcaacg
4021 gacacttcat aaaacttcgc cagctggata agggcatagt ggctgatgtc cttgaaattg
4081 tccccctcat aacttcccag cgcagactta gaaaggtggg tctgttccgc aagctgttcc
4141 agcgtcaagc cacgctctac acgcaggtct tttaaccgtt ctggtatgga taattccatg
4201 cttccctctc cctccttgtc tgcccgataa atgggaagtt gtcgctctaa atacaattct
4261 tattgttatt tttcgtgtac tttttttccg gaaatatgtt ctatttctat ttctatcaac
4321 tgtacgcctt taattccctt tctgatttct tcttcaacct cctccgctga cggataatat
4381 ttaagtgcaa attttctcac tttttcttcc gtagtcttac agtcttcaac tagccttgct
4441 ctgccaaata ccacacagct tgaaacatga taagcccaat ctccatcatc tacatatcca
4501 tcattccatg tcgtgaagca aactttgtcg ttatttataa tggaatcaat cttatgtcct
4561 ttttttgcgc tatgaaaata aattttattt tcattttcat catagtaaaa atttattggg
4621 attgtataag gatatccgcc atctccgttt acggagaaag ccgctctctt attatttctc
4681 aacaaatctt ttgcttcttc aatcgctatt tcgttcttaa tctttcttac ttttctaaac
4741 ataacaactc ctcctgcaaa taccgattta ccgttctgtt tcttccattc taccacaatt
4801 ccgagagcgt ggaaatagcc tgtttttgaa gccaatttcc cacctttcgg atatacggga
4861 cgggcggtca tttaggtgta gacttagatt agttcatcga tggacagcac cttgaaaacc
4921 aaatgaccgt ccgaaaagga ataccctggc tggggaagca ccgcaaggag ccaacttctg
4981 gacactttgt ccagaggtca cttcgggaca agttgtcccg aagttatggg gagcgtgcca
5041 atgccggaac acgccgcagg aatggggcag gaagcctttt cggggagaac ggcaacggaa
5101 agcaaactgg agggaacgca tgagagataa cccctataaa gacttgccgc cgctggaacg
5161 caagccggac ggttcccttt accgcatgac accagcgcag cggaaacggg caagcccccct
5221 gatacgccgg gcgtgctgta actgcgaagg cggcaactgc attgtccttg acgatgggga
5281 cacctgcacc tgcccgcaga cggtttcttt ctcggtctgc tgtaagtggt tccgctgggc

5341 ggtcttgccg ctggacggaa cgctggaagc ggagattttc cgggataagg acttgaaacg
5401 ctgtgtggtc tgtggcggta tgttcgtccc caaatccaac cgggcaaaat actgccccgg
5461 ctgcgccgcc agagttcaca ggcggcagaa aacagaaagt gaacggaaaa ggaggtctgc
5521 tgtggacagt taggagcgga aaaagccttg atttatcagg cttcgcaagc cccaaaccgg
5581 ggcaggtggt ataaagtatc gcctaccccg gaaaacaggc ttctaaccgt ccacaaaacg
5641 cactatgaca aatacaatct atatccatca gccggaaaag gcgttcagct tcacccggct
5701 cccgaatttc ctctttgaag cccccacatt caagcccctg tccaacgagg caaaggttct
5761 gtacgccttt atcctgcgcc ggacagagtt atcccgcaag aatgggtggg cggatgactg
5821 cggacggatt ttcctgtatt accccatctg cgaggtggtt gacctgcttc actgtgggcg
5881 gcagaaagcg gtgaacaccc tgcgggaact gcaatatgcc ggactggtgg agatccagaa
5941 gcagggctgt ggaaagccca accgcattta cccaaaatcc tatgaagcgg tttcaaacac
6001 cgacttcaag aaatccggtt atggtacgcc ggagggctga aaaccgtacc catgaagtac
6061 ggaaatcacc cgtgagaagt acggaaaccg gacggtatat agaaatacaa agattaaaag
6121 attgatttag attctatcca ttccaatcct atccgagcta tttccggcgg gattttccct
6181 gtggaaaacc ccggatggga aaggaatggg gaaaggagtt tcatggcaca acacgcaatt
6241 ttgcgtttcg agaaacacaa gggcaacccg gcaaggccgc tggaagccca tcacgaacgg
6301 caaaaggaga agtatgccag caatcccgac attgacacca gccggagtaa atacaacttc
6361 catatcgtca aaccggaggg cagatactac catttcattc aaaaccgtat tgaacaggcg
6421 ggttgccgca cccgcaagga cagcacccgg tttgtggaca cgctgataac cgccagcccg
6481 gagtttttca aggggaaatc ccccaaaggag atacaggcgt ttttccagcg ggcggcggac
6541 tttctcatcg ggcgggtagg gcgtgaaaat atcgtgtcgg cggtggtaca catggacgag
6601 aaaacgcccc acctgcattt gacctttgtt ccgctgacaa aggacaaccg cctgtgcgct
6661 aaagagatta tcggcaaccg ggcaaacctg acgaagtggc aggacgattt tcacgcctat
6721 atggtggtga aatatcccga cttggagcgt ggggagagtg ccagcaagac aggccggaag
6781 catatcccca cccggctttt caaacaggcg gtttccctct ccaaacaggc cagagccatt
6841 gaagccgccc ttgacggcat taacccgctg aacgccggaa agaaaaaaga ggaagccctc
6901 tccatgctaa aaaagtggtt cccgcagatg gagaacttct ccgggcagtt gaaaaagtac
6961 aaggtcacaa tcaatgacct gttggcggag aatgagaagt tggaagcaag ggcaaaggcc
7021 agcgaaaaag gaaagatgaa agatacgatg gaacgggcaa agctgaaaag cgaactggac
7081 aatttacagc ggctggttga ccgtatcccg ccggatatac tggcggaact gaaacgtcag
7141 cagcggcaca cggtaaagga aaggtgatag atataagcag aaattttcgc cgcctttgtg
7201 cggcattgta ccttgaaaac tgaatatgga ggacactgga tggcaaaaag cgaaagcgat
7261 atttttacgc cccggacagg gcaggtcata caaacagaga acggcacgca gtattttgta
7321 tgtgggaaca accgtataaa aatctccgaa cacttcgccg caggcgggaa gcccctcggt
7381 gatctgattg tagatgtggt gcggcatacc gcagaaaaag ccgcttcaac ctgatagccc
7441 atcattgata acacgcccac gcttatgata taattgccat agagcaaaag tattgtaagc
7501 gtgggttgtt tctttagaag gaggattttt aacggtgaaa caaccttaca atactacgat
7561 ttacaacacg gcgctttata tgagattgag ccgggacgat gaaaactatg gtgacagcgt
7621 aagcattgaa acacagcgga caatcctgca acagtacgca aaggaacagg ggcttcatgt
7681 agtcggtgag tatgtggacg atggctggtc ggggacgaac tttgaacgcc cggattttca
7741 gcgcatgatg gacgatgtgg aagctggaaa agtaaactgc attgtcacga aagacctttc
7801 ccgtttcggg cgggaacatg tgatgatgga ctactatctg gaatttctgt tcccggaaaa
7861 acgggtgcgc tacatcgccg ttgcggagaa tgaggacaca gagaaagggc tttccgattt
7921 tgtcccgttc aaaaacctgt tcaatgaatg gtttgcgaaa gatacaagcc gcaaggtaaa

7981 ggccgctttc aaagcaaagt ttgccacagg acagcgtatc ggcgcctatg ctcccatcgg
8041 gtacaggaaa cacccggaaa tcaaaaacaa gctgataatc gacgaggaaa cccgctggat
8101 agtggagaag atttttgacc ttgccattca tggcagaggg gcggccagta tcacaagaat
8161 attgattgcg gaaaaggttc ctacgccggg atttatcaac ttccagcgtg acgggacttt
8221 cgcaaacatc tatgcgggtg cgccagagga aaaaagctac gcatggacga tagcgcaggt
8281 caagagcatt atgaaagatg aaacctatat cggccatacc atccactacc gggaaacaaa
8341 catttccttt aagaacaagc gaaggatacg caaaccccaa agtgaatggg tacgggtgga
8401 gaatacgcag gagccgatta tcagcgaaca ggttttccgg caggtacagg agcagatagc
8461 aaaccgccgc agaaagtgca aggatggtac gacgcagatt ttttccgggc tggtaaaatg
8521 tgcggattgc ggctggtcgc tgtcctatgg ggagaacagg cagaacagca agccttacgg
8581 ccattatcat tgtagcaagt acggacaggg cacacgccaa tgctccatgc attatatccg
8641 ctatgatgtg ctttacgcct atgtcctctc ccgtctgcaa tactggtcgg ggctggtaca
8701 gcatgatgaa gaacggcttt tgaagcggct gttaaatgcc aatgacaagg gacaggccgc
8761 cgcaggaaag aagcaagccg cagagttgaa caaagcggag aagcggaaag ccgaagtcga
8821 taccctgttt gcccggatgt atgaggactg ggcggcgggg cgtatcacgg aatacaactt
8881 ctctatgctg tccgggaagt accagagcga acaggctgaa ctggacacaa agattgaaca
8941 gcttcaatcc gctatcgcca ctgaaagcca gaacgccgca gacgcagaaa aatggattgc
9001 cttgatgaaa gagtgcgtca atccaacaga actgaccgcc gaacttttga atacgctgat
9061 tgaaaaaatc gttgtccatg aagcggtcaa aggtgaggac ggaagccggg aacaggaggt
9121 agaaatcttc taccgcttta tcggcaaaat cgaatgacac atcttgagat acccaacaat
9181 atctttaact aagggaaact ggactgggct tccctgatat taatacactg gaacctctgt
9241 cagatgcact cgagatagga gtggtggaac ttatgagatc ggaaatcaac acagaaaaaa
9301 gagaagataa aataaatcaa gaatgtgtga ccgagattat gaaatcaaca gcagagatag
9361 cggaatatca taaaatgaag aagatgaaat ggttgatcgg cggaatttgc tttgtagtag
9421 cagtgctggg agtgatggtt acacagtcaa caaaagtgtt ttcttcgatt gtattttatg
9481 cttgcatcat aggatgtact acatgtgtgg ttcttttacg gcagcattgg aaaattcgtg
9541 aaaaaagaga attttatttt tcgcttatct gctgtctggg agcagccagt ctgatatgga
9601 tgttccaggt gttttcgata aaagcaataa acagtatcat ggatgtgggg agcattctta
9661 tttgtattgt gattgatttt acatttttaa aagattttat tgaaagtgta attaagaaag
9721 aatatctgca aatggcagtg tgtggaatat ttcttctttc aacgacagta gaatattttc
9781 ggattgtatt aaagtaa
//
DEFINITION Integrative and mobilisable element IME_3
fasta_record 1..30313
/ID=B77|SC|contig000001
/colour=5
CDS complement(163..783)
/color=8
/locus_tag="B77_03249"
/note="NUDIX hydrolase Hydrolase of X-linked nucleoside
diphosphate N terminal, protein motif:Pfam:PF12535.2"
/product="NUDIX hydrolase Hydrolase of X-linked
nucleoside diphosphate N terminal [hydrolase]"
/"ID=B77_03249

/protein_id="gnl|SC|B77_03249"""
/codon_start=1
CDS complement(867..2033)
/color=8
/locus_tag="B77_03250"
/note="ABC transporter permease subunit, protein motif:Cdd:COG1277 protein motif:Pfam:PF12730.1"
/product="ABC transporter permease subunit [transporter]"
/"ID=B77_03250
/protein_id="gnl|SC|B77_03250"""
/codon_start=1
CDS complement(2038..2904)
/color=8
/locus_tag="B77_03251"
/eC_number=3.6.3.-
/note="ABC transporter ATP-binding subunit, similar to AA sequence:UniProtKB:P32010 protein motif:CLUSTERS:PRK13537 protein motif:Cdd:COG4555 protein motif:TIGRFAMs:TIGR03522 protein motif:Pfam:PF00005.21"
/product="ABC transporter ATP-binding subunit [transporter]"
/"ID=B77_03251
/protein_id="gnl|SC|B77_03251"""
/codon_start=1
CDS complement(2895..3650)
/ID=B77_03252
/color=1
/locus_tag="B77_03252"
/note="hypothetical protein"
/product="hypothetical protein"
/protein_id="gnl|SC|B77_03252"
/codon_start=1
CDS complement(3640..4194)
/color=7
/locus_tag="B77_03253"
/note="RNA polymerase sigma factor Sigma-70, similar to AA sequence:UniProtKB:P0AGB6 protein motif:CLUSTERS:PRK12513 protein motif:TIGRFAMs:TIGR02937 protein motif:Pfam:PF08281.6"
/product="RNA polymerase sigma factor Sigma-70 [transcriptional regulator]"
/"ID=B77_03253
/protein_id="gnl|SC|B77_03253"""
/codon_start=1

CDS 4585..4893
/color=8
/locus_tag="B77_03254"
/note="Type II toxin-antitoxin system RelB/DinJ famil
antitoxin RelB , protein motif:TIGRFAMs:TIGR02384 protein
motif:Pfam:PF04221.6"
/product="Type II toxin-antitoxin system RelB/DinJ famil
antitoxin RelB [toxin-antitoxin]"
/"ID=B77_03254
/protein_id="gnl|SC|B77_03254"""
/gene="relB"
/codon_start=1
CDS 4890..5219
/color=8
/locus_tag="B77_03255"
/note="=Type II toxin-antitoxin system RelE/StbE family
toxin RelE, protein motif:TIGRFAMs:TIGR02385 protein
motif:Pfam:PF05016.8"
/product="Type II toxin-antitoxin system RelE/StbE family
toxin RelE [toxin-antitoxin]"
/"ID=B77_03255
/protein_id="gnl|SC|B77_03255"""
/gene="relE"
/codon_start=1
CDS 5401..6369
/ID=B77_03256
/color=1
/locus_tag="B77_03256"
/note="hypothetical protein"
/product="hypothetical protein"
/protein_id="gnl|SC|B77_03256"
/codon_start=1
CDS complement(6589..7320)
/ID=B77_03257
/color=1
/locus_tag="B77_03257"
/note="hypothetical protein"
/product="hypothetical protein"
/protein_id="gnl|SC|B77_03257"
/codon_start=1
CDS complement(7418..7699)
/ID=B77_03258
/color=1
/locus_tag="B77_03258"

/note="hypothetical protein"
/product="hypothetical protein"
/protein_id="gnl|SC|B77_03258"
/codon_start=1
CDS complement(7719..7916)
/ID=B77_03259
/color=1
/locus_tag="B77_03259"
/note="hypothetical protein"
/product="hypothetical protein"
/protein_id="gnl|SC|B77_03259"
/codon_start=1
CDS complement(7950..9152)
/ID=B77_03260
/color=1
/locus_tag="B77_03260"
/note="hypothetical protein"
/product="hypothetical protein"
/protein_id="gnl|SC|B77_03260"
/codon_start=1
CDS complement(9184..9342)
/ID=B77_03261
/color=1
/locus_tag="B77_03261"
/note="hypothetical protein"
/product="hypothetical protein"
/protein_id="gnl|SC|B77_03261"
/codon_start=1
CDS complement(9548..9961)
/ID=B77_03262
/color=1
/locus_tag="B77_03262"
/note="hypothetical protein"
/product="hypothetical protein"
/protein_id="gnl|SC|B77_03262"
/codon_start=1
CDS complement(10022..10246)
/ID=B77_03263
/color=1
/locus_tag="B77_03263"
/note="hypothetical protein"
/product="hypothetical protein"
/protein_id="gnl|SC|B77_03263"
/codon_start=1

CDS complement(10604..11359)
/ID=B77_03264
/color=1
/locus_tag="B77_03264"
/note="hypothetical protein"
/product="hypothetical protein"
/protein_id="gnl|SC|B77_03264"
/codon_start=1
CDS 12774..13160
/color=4
/locus_tag="B77_03265"
/note="plasmid mobilisation relaxosome protein MobC,
protein motif:Pfam:PF05713.5"
/product="Plasmid mobilisation relaxosome protein MobC
[relaxosome]"
/"ID=B77_03265
/protein_id="gnl|SC|B77_03265""""
/gene="mobC"
/codon_start=1
CDS 13138..14691
/color=4
/locus_tag="B77_03266"
/note="Relaxase/Mobilisation nuclease domain, protein
motif:Pfam:PF03432.8"
/product="Relaxase/Mobilisation nuclease domain
[relaxase]"
/"ID=B77_03266
/protein_id="gnl|SC|B77_03266""""
/codon_start=1
CDS 14721..15047
/ID=B77_03267
/color=1
/locus_tag="B77_03267"
/note="hypothetical protein"
/product="hypothetical protein"
/protein_id="gnl|SC|B77_03267"
/codon_start=1
CDS 15034..15813
/color=8
/locus_tag="B77_03268"
/note="FAD-dependent thymidylate synthase, protein
motif:CLUSTERS:PRK00847 protein motif:TIGRFAMs:TIGR02170
protein motif:Pfam:PF02511.9"
/product="FAD-dependent thymidylate synthase

[methyltransferase]"
/"ID=B77_03268
/protein_id="gnl|SC|B77_03268"""
/codon_start=1
CDS 15890..16216
/color=8
/locus_tag="B77_03269"
/eC_number=3.6.1.23
/note="Deoxyuridine 5'-triphosphate nucleotidohydrolase,
similar to AA sequence:UniProtKB:Q2YRG4 protein
motif:CLUSTERS:PRK00601 protein motif:TIGRFAMs:TIGR00576
protein motif:Pfam:PF00692.13"
/product="Deoxyuridine 5'-triphosphate nucleotidohydrolase
[hydrolase]"
/"ID=B77_03269
/protein_id="gnl|SC|B77_03269"""
/gene="dut"
/codon_start=1
CDS 16367..16492
/color=7
/locus_tag="B77_03270"
/note="helix-turn-helix transcriptional regulator, ab
initio prediction:Prodigal:2.60"
/product="helix-turn-helix transcriptional regulator
[transcriptional regulator]"
/"ID=B77_03270
/protein_id="gnl|SC|B77_03270"""
/codon_start=1
CDS complement(16622..17050)
/ID=B77_03271
/color=1
/locus_tag="B77_03271"
/note="hypothetical protein"
/product="hypothetical protein"
/protein_id="gnl|SC|B77_03271"
/codon_start=1
CDS 17374..17748
/ID=B77_03272
/color=9
/locus_tag="B77_03272"
/note="transposon-encoded protein TnpV"
/product="Transposon-encoded protein TnpV"
/protein_id="gnl|SC|B77_03272"
/gene="tnpV"

/codon_start=1
CDS 18115..20034
/color=2
/locus_tag="B77_03273"
/note="tetracycline resistance protein tetM from
transposon Tn916, similar to AA sequence:UniProtKB:P21598
protein motif:CLUSTERS:PRK13351 protein motif:Cdd:COG0050
protein motif:TIGRFAMs:TIGR00484 protein
motif:Pfam:PF00009.21"
/product="Tetracycline resistance protein tetM from
transposon Tn916 [antibiotic resistance]"
/"ID=B77_03273
/protein_id="gnl|SC|B77_03273"""
/gene="tetM"
/codon_start=1
CDS 20752..21177
/color=7
/locus_tag="B77_03274"
/note="RNA polymerase sigma factor Sigma-70 , protein
motif:TIGRFAMs:TIGR02937"
/product="RNA polymerase sigma factor Sigma-70
[transcriptional regulator]"
/"ID=B77_03274
/protein_id="gnl|SC|B77_03274"""
/codon_start=1
CDS 21770..22114
/color=7
/locus_tag="B77_03275"
/note="ribbon-helix-helix protein CopG family, ab initio
prediction:Prodigal:2.60"
/product="Ribbon-helix-helix protein CopG family
[transcriptional regulator]"
/"ID=B77_03275
/protein_id="gnl|SC|B77_03275"""
/codon_start=1
CDS 22116..23471
/ID=B77_03276
/color=4
/locus_tag="B77_03276"
/note="Relaxase/Mobilisation nuclease domain"
/product="Relaxase/Mobilisation nuclease domain
[relaxase]"
/protein_id="gnl|SC|B77_03276"
/codon_start=1

CDS 23468..23758
/ID=B77_03277
/color=9
/locus_tag="B77_03277"
/note="transposon-encoded protein TnpW"
/product="Transposon-encoded protein TnpW"
/protein_id="gnl|SC|B77_03277"
/gene="tnpW"
/codon_start=1
CDS complement(24054..25727)
/color=9
/locus_tag="B77_03278"
/note="serine recombinas, protein motif:Pfam:PF00239.15"
/product="Serine recombinase [recombinase]"
/"ID=B77_03278
/protein_id="gnl|SC|B77_03278"""
/codon_start=1
CDS complement(25805..25978)
/ID=B77_03279
/color=9
/locus_tag="B77_03279"
/note="transposon-encoded protein TnpW"
/product="Transposon-encoded protein TnpW"
/protein_id="gnl|SC|B77_03279"
/gene="tnpW"
/codon_start=1
CDS complement(26437..26679)
/color=7
/locus_tag="B77_03280"
/note="helix-turn-helix domain-containing protein, protein motif:Pfam:PF12645.1"
/product="helix-turn-helix domain-containing protein [transcriptional regulator]"
/"ID=B77_03280
/protein_id="gnl|SC|B77_03280"""
/codon_start=1
CDS complement(26663..27100)
/color=7
/locus_tag="B77_03281"
/note="RNA polymerase sigma factor Sigma-70, protein motif:CLUSTERS:PRK11924 protein motif:TIGRFAMs:TIGR02937 protein motif:Pfam:PF08281.6"
/product="RNA polymerase sigma factor Sigma-70 [transcriptional regulator]"

/"ID=B77_03281
/protein_id="gnl|SC|B77_03281"""
/codon_start=1
CDS complement(27574..28557)
/color=8
/locus_tag="B77_03282"
/eC_number=2.1.1.192
/note="R23S rRNA adenine 2503 methyltransferase RlmN,
similar to AA sequence:UniProtKB:Q7A600 protein
motif:CLUSTERS:PRK14469 protein motif:TIGRFAMs:TIGR00048
protein motif:Pfam:PF04055.15"
/product="R23S rRNA
%28adenine%282503%29-C%282%29%29-methyltransferase RlmN
[methyltransferase]"
/"ID=B77_03282
/protein_id="gnl|SC|B77_03282"""
/gene="rlmN"
/codon_start=1
CDS complement(28574..28759)
/ID=B77_03283
/color=4
/locus_tag="B77_03283"
/note="conjugal transfer protein Cysteine-rich KTR"
/product="Conjugal transfer protein Cysteine-rich KTR"
/protein_id="gnl|SC|B77_03283"
/codon_start=1
CDS 29158..29514
/color=7
/locus_tag="B77_03284"
/note="anaerobic benzoate catabolism transcriptional
regulator helix-turn-helix transcriptional regulator,
protein motif:CLUSTERS:PRK08154 protein motif:Cdd:COG3800
protein motif:Pfam:PF12844.1"
/product="Anaerobic benzoate catabolism transcriptional
regulator helix-turn-helix transcriptional regulator
[transcriptional regulator]"
/"ID=B77_03284
/protein_id="gnl|SC|B77_03284"""
/codon_start=1
CDS complement(29517..29708)
/ID=B77_03285
/color=1
/locus_tag="B77_03285"
/note="hypothetical protein"

/product="hypothetical protein"
/protein_id="gnl|SC|B77_03285"
/codon_start=1
CDS complement(29737..29934)
/ID=B77_03286
/color=7
/locus_tag="B77_03286"
/note="Ribbon-helix-helix protein CopG family"
/product="Ribbon-helix-helix protein CopG family
[transcriptional regulator]"
/protein_id="gnl|SC|B77_03286"
/codon_start=1
CDS complement(30194..30313)
/ID=B77_03287
/color=8
/locus_tag="B77_03287"
/note="molecular chaperone"
/product="Molecular chaperone"
/protein_id="gnl|SC|B77_03287"
/codon_start=1
BASE COUNT 9471 a 5978 c 5527 g 9337 t
ORIGIN
1 ttttctttaa tgcaactcgt aacattagat ttttacattt ttcatgaaat taagtaaaga
61 aaaagagcat atggaattga atccatacac tctatgatta ctggaacttt atttcatttt
121 taattcaaca aactagaagt tgtcagttta gaaaaccgga agtcaatcaa ataaagcttc
181 ccatttttct gattgcattg cattaaaaca catcgcaatt tgttctttat tattcttttc
241 ttctgcaagt tctggaagct catcaattcc aaaataatca aaacctgttg tttcaatatt
301 cgttttaaat tgaccgcccg tcacttcaca ttgcataaaa atcttacata ctttatacgc
361 atatattgga agattatgtt tctccctatc ttgaacagca atcacacgtt ttacagtaac
421 atcaagtcct gattcctctt tgacctcctt tattgtattc tcctctacgg aaatatcgac
481 atcacaccaa cctcccggta atgaccaagt accatttgct tcatgaacca ataatatttt
541 cccgtcttta aaaatcgcag cccttgtatc tattttagga gtctgatatc ctacatcact
601 gcaaaataaa ttttgaactt tttctgtaga tatttcagtt ttatagctta tcatttcagc
661 agaaatttct cggattcttt catatcgttc tatatcaaat ttgtcctttc cataaaacaa
721 accaccctga gcaatagctt gtaattctac tgcccatttt agccattgtt cgtttttatt
781 catagattat cgctccttta aattccgatt tgttactcac atgatatcat attttttctt
841 caattgcatt ttcattgaaa aaatcattaa aacttccaat gtttaatata aataaagcaa
901 tatagttgca tccctactcg tattaagagc tgaaaaataa atataaatcc tatattaatg
961 aagaaggaat ggaagaaaaa agcatctttt gaatatagaa tacctaatat attttgtaca
1021 ttcaaattat ttattggaaa caaatacaaa attttgttac ctacccctgt ttttatcatt
1081 tgcaccagca ataatggtaa tatatatacc cctaatgaaa cagctaatac tgacatagat
1141 gaatgcataa gagctgatat taacaatgtt acactctcaa caaatccaat accaactaat
1201 tgcatgacaa gaattaaaaa gtatatttga atatggttat attgcaaagg gaagtcaaat
1261 aaccccaacg acaaatttgt ttgtatgtct gaattccatc cgctcatatc atacattgta

1321 gagaaaacac ctatgttgac caattgaaaa ataataaata aactcactgt aataatactt
1381 gtcgcaatca atttagcaaa tatcattttg tttcttccat attttgtaac taacagaatc
1441 tgattgatat ttttagaatt atcatctgaa aacactatgg aagataccaa tattgtaatt
1501 atggacaaaa gagcatatat atatccaagt actcgatata aatctgtcca tggaacataa
1561 tttcctaatg tcaaaggagt atcaaatttt ttaaatccag cctcgtccaa tgaatataaa
1621 tccacgtctt ctattgttaa tctttggcct ttattaacct gttcaatcat ttctgaataa
1681 acattttcgg catcttttga aaaaaaagta tctcccattt cccaataaaa tggataaaaa
1741 ggcttattaa caactttttg ctcttcattc tcttggaaat tattcataga atcagacaga
1801 atcattttaa ctttttcatc tgtaaactct ccagaatatt tttcatctat tattacatcg
1861 aggctctttt caaacccctt tttcttatta ctcgcttcac tcattgatag ctgtgaatag
1921 tgatacccaa caaaaaaaat ccccagaagg actaagcatc ctacaacaat cccaccaaat
1981 acagttcggt ttctcaatat tttcccaagt tcaaacttaa ttagcgatgt catactctca
2041 ttccccctct ctaaaataaa acaaatacaa atcttctaaa accggatcaa cttgtttcgc
2101 atttgcaaaa ggtttttccc ttgaaataag tcgaagaaca actccgtctt ctccatattt
2161 acgattactt actgtatata cttgctcaat tcgatttact tctttttctg gaaccaggca
2221 ttcccataca cacttgttaa tatctttcaa caaatatgtt attgtgcctg tattttcaat
2281 tctacctttt tttagaataa tgatttcatc tgctatatat tctatatctg atacaatatg
2341 tgtagacaat aatactattt tttctgatgc aaacgaactt atcagttttc taaatttaac
2401 tcgctctttg ggatcaagcc ctaccgtagg ttcatctaat attaaaattt ccggattatt
2461 gatcattgct tgagcgatac ctagtctctg tttcattccg ccagaatact ttctgatttt
2521 tttatttttc acatcatcta agccaacctg aatcaataaa tctaaacatc tcttttttgc
2581 tattttccca tttaagcctt tcaatgaagc gatatacatc ataaaattta atcctgtaaa
2641 atcaggatag tatcgaaaat cttgtggtaa gaatccaaga atcgagcgat aatattctgc
2701 ttgcagagca atatctttat ggtcatattg tatatttcca caatttgctc tcatcagtcc
2761 acaaataata cgaaataaag ttgttttacc cgccccatta gcacctaata ttccgtaaat
2821 tccaggagta aatcgatacg acacatcatc aagcacagtt acattcccat attgttttgt
2881 tatattatta atagttaatt ccatactgaa gctacctccc atttgattat tttagtttga
2941 gaatataaga taatccctaa tattgtcgtt gtaatgaaat atgctattcc ccaataaagc
3001 attcgaattt catatatttt aaagatattt gcaattaaaa taactgcgac acaaatactt
3061 cccgatatac accataggaa ataattatgt aattgctttc tccatatagt tataagcgaa
3121 aataaaataa tgcaaacaat attgtatgga acaagtacat agactaatat ctgatacaaa
3181 ggcaaagata ctatagattt agtcataaga gaaagcaaaa taacaattac catatcaata
3241 atgccaaaaa tcaataattt catcataata tgttgtcgca aatcatattt gaatgtctgt
3301 tccaattccc acattcctgt agtaaaactt ttgaataatt catctataaa aaataaagcc
3361 gaaatgaata ttacacctgt gatgcataac attatttcac ttatcggctg taaattttta
3421 aaatttaatg ccatagttat ccctgataga aaaatgaata attgaattat ccaagtatga
3481 acatgtatga agcgaaaagt tgacaggaat agctcaaagt aagaatgtcg actcattcta
3541 agtctaatca ttatttcata tcctaattcg atgatttttt cacgtccttc tatatcacct
3601 tctaatttat aaccagccaa tttcgatata atatcttttt tatccatcat aaaattcctc
3661 tctttctaaa actttttcta attttctaag gccttgataa atacgtgact gtacagtaga
3721 tacccccacg ccagtaattg tcgcaatatc ttttaccttt aaatcgtgat aataccgtag
3781 tatcacaacc tctcgttgta tttcaggaag taaatcaagt gctttttgaa ccgtttcttt
3841 tctctcatca tgtaatattt catcaatcaa tgaagtgtct aaatcagttg taaaattatc
3901 tgtgagttct atttgctcaa tatgtctttt tttcaaataa ttattgcaag tattgacagc

3961 catagtgtat agataatttt gaaattttcc tatgaatcgg tatctaggta aactttctac
4021 caattttaaa aaaatatcct gtgttaaatc agaagctata tcgcagtctc cataacatct
4081 gcgaacacaa taaaaataca cactatcata atatttatgt actaaaatat cccatgcatc
4141 aggatctccc atcttaattt tgtttattat catccagtct tgtatcatat ccatcacttt
4201 tctcttaatt attcttctct atatataaac aactaacctt aacaaatcat ttaaaaacac
4261 aaaaactttt tatattattt ttatatttat ttacaatatg ataaactatt cttcccctca
4321 tgggcaaggg attaggagtt ccctgtggcg gttagtatgc agtagttcga ttatcctgta
4381 agcgtgagtt ttccaaactt atcggattct tggtagttgc cttagtagcc gttggtcttg
4441 tattcaatgc gtccggcgta aaagatgtat tgttacagtt attcaataag attattggag
4501 cataaaaata gtattgctat tgtgcggaca tttgcatata ataaagatag atggaaatac
4561 tactcacgaa aggagatgtt ttatatggct acaaaatcag caaatttata tgcaaggatt
4621 gaaccagatg tcaaagaaaa agcagaaagt atcctgtcta cacttggtat ccctgcttcc
4681 agtgctatca atatgtttta taaacagatt atcttacaga gaggacttcc gtttgaagta
4741 aaaataccat ctgccagacc ggttgatatc agcacattat cagaagcaga gttcaatgaa
4801 gaattggaga aaggatatgc ggatatgcag gctggacgga caaaaaatgc gaagaaagca
4861 tttgctgaca ttcgcaagga ttatggctta tgatatatga agtagaatta tccgaacagg
4921 ctgacagtga tttgagaggg atttttgaat atatttcttt tgaattacaa tcgccggaaa
4981 atgcaagcgg acagcttgac cgtctggaag aacagatatt aagtctggat acgatgccgg
5041 agcgttatcg aaaatatgaa aaagaaccgt ggaaaagtcg tggacttcgt gtattaccag
5101 tagacaatta tgtggtactt tatattccag acagcgataa aaaagttgta acaatactta
5161 gagtaatgta tgccggacgg gacatagaca atcagttaaa tctccacaca aagcaataaa
5221 ggatatgatt taaagcagtc tttcatttat gatcgactgc tttttcata ttcaaaatta
5281 aaaagaaagg gctgaattac gatggaagaa tgtagcgtat tgattgaaac gaccaaatca
5341 gcagaggata acaacgaatt gctgaactgg aacagcagat agagaaagga cgtgatatag
5401 atgaacgaat ccagagaatt aaagaacgcc gaacagttgg aagagcttct gctcctgacc
5461 gaggagatac aagaagaact ggaacagaaa gaccagctta tcgtggaact gaagacgcag
5521 ctcaacgaat ctctgatctt gaacgagaaa ttaaacaaag agaacagagc cgggaatatt
5581 caagcattaa agaacgactt gaagctggca gacagagcat tgcggaacga gaaagagaag
5641 ctgcgaagcg caaacgtcat gatcggggaa tatcaaaata aaatacgctg tttaacacaa
5701 caacgggatt atgcccggac acatcagaaa atcgtagaga taccggtaga aaagccggta
5761 ctctatgaaa aatgtgaatc ctgtgaccgg acagcctatc agaatgaaaa agcgaaatac
5821 ggaacacaaa aagagcgact tgccggacag tataaagcaa aaacggtaat gttccaaaca
5881 accttgttcc ttcttgcatg gtattcgctg acaaccactc tttttcaggc agtacagtcc
5941 gatgtgttcc tttccgattg taaatcattc ttccatgatg cagcgtcatt cctacaaact
6001 tttatcggtt ggacaattga tgccgggcaa tctgtggcac agattagcac aaaaattcca
6061 aatcctttta tagccggaat ggtatattgg ttatttctga tattggttgt cggaatatgt
6121 gtggcaggga cagggatact ggcgatactg atagaaataa aggttataga attatataag
6181 aaaaactgtt gggatgttat tactctactg atgatactga caagtactgc tgtcatcatt
6241 tattttggag aagcaattaa aaaagcactt tccgtaaatc ttctattgct tttcgtactt
6301 tctcagggag tatatcttgg aatcagatgt tatttgaaaa atcggttagg taccaagcaa
6361 tattggtgaa aacatgctaa aaaacgtata atagcagggg ataaaacatc gtcccctact
6421 attatataaa cagcgaagtt ttacccatct ctctcttttg aaagattcgc tacttgtgga
6481 accgagatat aaggtttcgc ctgtgttctt tggtgtatca ctaccacatt tctctgtacc
6541 gatgtaatct ttctgcatac ggaagcaaga gatacaatat tccttatttt aaaacataat

6601 tacaattcct gtttgacaag aacgatcatc atagaaggat tgtatttgtc cccactcata
6661 ataatattcg acgcctgtca tactcttttt actgggagta cctaagcatt tccttaaaaa
6721 cctattattt tcttcttgca actttctaa gactgtacca ttcattgtct catatttcgt
6781 agaatacttt ttgtctgctc gaaccaattc aatctttttt acttttccat gtcttccata
6841 gattgtaaca taaaacttct tataaagcca ttttaaaggt ttcttagaaa tcaatattgt
6901 ataagaatac tctgtttgag caatcttaaa atcaatttct ttaaataggg aaatacttcc
6961 tttgggagta attttatatc cccctatttt taacatatct tcctttagct catcctttat
7021 ttggataatc tccatactta ctataaaaac aacatagcct tcacacacta caacggtaac
7081 tatttttgcc catagcacat ctggaacaaa catgttccaa gctataacag gaagcaaagc
7141 tactcctacc gcagtcacca gtagaagcac cctccaaata taatttttct ttctttttcc
7201 ttcattagtt aaattttcat cataataaag atgacctggg gcttcgaatg agtttaatag
7261 aaattccttt aaaaatgaca ccagaaccgc tccccaagaa gatttctttt gttccttcat
7321 attacccaat ctcctctgct atatgattat ttttcattct attttttact taacttatac
7381 attagcaaga tttttagtta ttcttctaac aatattttta tccccgcttt ttctgcttta
7441 tgcgaaaagt caacagcaaa tccccatccc atctctgcaa attgccattt cttcgttttt
7501 ttgtaatttt atagctttca actgtagaca ctttatttaa ttcaggatat aacttcttaa
7561 gtgtaagtat ttgtttattt ctgcttattt tttcattcca tataattgtt aatttaacct
7621 gttttctttc aaaaaaaccc accttacata aatctgcata gcaggcttct ggtattgata
7681 tatcattgtt ttttaacatt tttctccttt atttaatttt ataaaaaact tctaagccct
7741 attaaaagaa atatgaagcc tattagagga acaggtatca tacaaatgag aaacacatta
7801 aattttttcc ttgcttcatc agtattttct tcttttgtta cttttgcttt tttcataact
7861 ataaaaggga atatcagtaa taataatcct ccaactactg ctaacaatat tttcatctta
7921 tatattccct ccttatcttt aaataaatat taatcatgta tcgtactata ttccaatcct
7981 aatgcacttt ttcttctaca aattctttta ctatcgccta ttgatgtatt ttcatattca
8041 ctgtttgaaa cagtatatat ttctctttta ccattacaac ttacatataa ctctctatct
8101 ctagttttac cactgctaat ctttttgtct gttacaattg caatttcatg agttgcccca
8161 tcaaatgtaa ataaaaaatt tattgggaag ctaattgtaa atgcgattat aaaagcagca
8221 aatacaactg aaagtttacg tccaaatttc tgtggtacat ctgtttttaa agatttaata
8281 ataaatggca gtgcaagtaa agcagtaata attgccgtaa tcttcatata gtttccaaat
8341 tcataattaa agagtttact cgtattcagg gataaaagca ttgctattgc tgcacccata
8401 aatggtaatt gatacgccag ttcttgtcct ttttttgtgg tgacttctat aaatatatat
8461 ggataatatt ttatatacac tgcatatgta atcatcaaaa ccattataaa tataactaac
8521 attgttttgc caccaataaa aaatgctacc gcatcagcaa ttattagaat ccaacctaaa
8581 gcttttaaaa attgctttgt ttttttacgc ttttcttttt ctattgtttt tgacatattt
8641 ttgatagttg tattctcatt acatatagat ttatagaaat tcttttctat ccagtttaaa
8701 gcaggtacat actttgatac atctttgttc tgttcaacta actcaccaaa atccacaaga
8761 tctatttcat gttttataag tgtatcatat aaaatatcat aatttacata cacttgctcc
8821 agagtaacaa gtctattgta catcttatca attaatatga catctgttct accatttaat
8881 ggaatcatta aaatacaacg aatatcatta tattttatat gctgagtttt cttaaccaga
8941 tagctaaata caatttcatc ttcatttact acaattattt gtttcttgta tgccaaaatt
9001 aaagcaatac ctaaaaatac aaaaggcata aaacatatcg caaccccat atttcctgta
9061 acaaatgtta ccagtacgat tattaaaaaa agaactgttg ttactattcc taaatttttt
9121 acaaatttac ttaatctaat gacattatcc ataaactcaa tctccctata tgattcacat
9181 ttttcatttt ttacgttcta atttcttctt atccttttt atttctgcaa tctttacttg

9241 cagccagtcc acatttctat ttcgggcaga aattttagcg attttttttct gatttttata
9301 taaaataaga gcttctccaa ataaattctt ttttcttttc atctgggtaa tatccgttaa
9361 tcttatttta cagttcttcc ttaaaaatcc atttctatat agccatccat ccttgacaat
9421 tgttttgtcc attaaaacat aaaataaatt tctaattatc ttttcataag ttcttataac
9481 accgatatat ccctatttta caggctttat cggtgttttc tttttggcag atacttctta
9541 taaccactca taaatctgca ttcttactaa gtccgttttc ttttggaaat attttttcaa
9601 tgtcacgaaa gcggtttgca cagaatatat gtggtaaaat attccaagcg tgaatggaat
9661 ggtcgcaacc gctgtttgtt attgtgagtt gtccccatgt tactgccgaa ttgaacgcat
9721 tatttaaata gctcatatcc ttttcttgta atgctaagta acttcctaaa taaacaatct
9781 tttttccgaa attttcatgc aataatttca gccatataat aatctgttct tctggctttt
9841 tactttgaca gtatttcgta tacttcttta atcgctcatc aatatatgcg ttcatgttgg
9901 catatccaaa cattttatcc acattttgta aatatagttt gataagttcc tttttttcca
9961 tagagtacct cgtttcactt ataaaatttt gctatttgct acataaatgc tttcaatccc
10021 attagaataa gtataatagc aatatataca ccccacttat cccaattata agtgtcataa
10081 taaaattgaa aacaatacaa actttcttct gatttaaaga aaaatgttca aatttttcaa
10141 ggttgtattg acgaatataa cttttagtaa ttatggaata aattgaataa ccaatagcaa
10201 tgatacttga aataatagca ataatatcag tcattttaca atccatatct tccctcctaa
10261 aaagtttgca tatcattacc acaaaaatta aaatattata atgtttaaat ttattgtttt
10321 tcttttgtga aatagatact ttttatatca catcttataa aatgcggttt tgccttgtaa
10381 atacagcgtt ttccgcctgt tagtgtatag tgtcttttgt tttttgtgcc ccccttgtta
10441 gataacgggg gcttaatctt cgctcgccac aggctcgctc acagacgaag taaaccagcc
10501 acttgcaaga catggcagaa aaatctgaaa gattttcctg ccaccgtgtc tggatcactc
10561 cgtctgccta actaataata tagtattatg taaaaatgtc attttattta ggtacatcta
10621 ccgaatagct taaacagtca cctcttttgt tactttgtaa gtcattgtaa taccatagat
10681 acatgactct gtcatataca ttgatttcaa ttacaccgtc cggtttttgt ccttctactt
10741 ccaatgaata gatttcttta tcatgtgcat agccgtaaac aaaacgcttg cttggataat
10801 cggggtttaa gtcttgttgg aaatcactta atttcaaaga agagtgaata agctctgtaa
10861 tatcatctaa caaaattcta tctcgtttcg ttgtaaccgg tgtagcattt ataaatgcgt
10921 attgtgtatg cccattctgt atctgctttt tacaacgagc cattactaag ccttgaatat
10981 catcgttatc atttccagcc cgatagaaaa tcgtatcata ttttttgcct tgccatatat
11041 gcaaggtttc atcaacatga ttaagaaaat cagcaggaac cgaaagttcc tcattttcag
11101 cattcaaagt ggaaagttga atagcttctt caatcgtgtc ataatataca agcttggcat
11161 tatttttatc ataatcattt atcgtaaatt ccacgtattc attatcttta ttttcttctt
11221 ccggaacata agtataacga ccattttcaa tatggtcatc atattcaaca tcaacaaata
11281 acccaatccc tatcagagct acaaaaatca aagaaatcgc tccaattatg attaacatta
11341 ttttcaaaaa ttttttcatg gcatacactt ccttaatcac ttcaagttta tctattatta
11401 tttatcatat tatatcatta gaaaattgag tagtcatcac ttttctagta tctgtcagaa
11461 tattttttat tccataaatt atcaagagtg taaacattgc tacgcaacaa gctcttgaca
11521 attcatgtcc taaaaaatga ttaataatca agcagaaaaa gaatagttat actgtattac
11581 agcaacaaaa aagacatctc atattctgta tgaaatgcca attttgcctt tatttatgct
11641 gttttctctg tcgtatcaag gttctttcat tcgtagtaaa ttagtggtaa aattctttta
11701 ttttttgttt catattgctt aaaatccctg tgtttatgcg gataattcga tttttactct
11761 ttttgggata aaaactgcat cacaatatcg aaccaaaata taaaaatcaa ttttatctct
11821 ttatcataat tatatacatc aactaaccac attgtaaagc aaaaagcaag ccattttcag

11881 acttgctttc caccagatta ctgcacctta ttgccaaatc atctccttat tcaaaattcc
11941 cttttcacac tcttgtatag tatttatttg ctacccattc catctgttta tctgccttca
12001 actcttcctt aatcccctgt gtctatttca tttgctctac aattgactat tattccagcg
12061 attatataaa aaccataacc aatacaaaat atggaaacgg tgcttctaac tataccgcta
12121 aagcctttcg ccattatccc gaaaatagta ctccagcaga gaaatagcgc cagcagacat
12181 aaaaaactcg aaccccttgt aaaatgggca tcagaagttt tctacccgtt attcttgttc
12241 tataattcaa gtgaatcttc tgaatctatc cacatctgta aattgccatc gagatataat
12301 cttgcatatt ctaaaggctc atccacagcc aatgaagtaa gcgcacattc tgatccatat
12361 gtggtttta aatctttttc agctccccag caatcaatgc gaagcatata tccaacactt
12421 atgcttttct cttttttca ttctatcaca agctcaccat taatcatcat tttttgata
12481 ttttttcttt tcccccactg taaaacaggc tctcagcgtt tcgttatata gaggggtaca
12541 catacagagc aagatccacc catgtttaac agtattgccg tttggcatcc tgttctccta
12601 tgggatttat cttgatgggg atctccccat accctcttaa taacagacgt ttcacttgtc
12661 tgttggttcc acctgaaggt gttcacggca ctatttctaa tatcggaagg gacagccgat
12721 attgaaatac gtggactccc ggcgggagga atagtctcag cggaaggagg agtatgagaa
12781 caaaaaacaa agataaagaa ctgaatcaca gacattttat aggattacgt cttacagatg
12841 tcgaactgga tttattagac caaggcgctg cttgtttatc tattagccga tcagaatatc
12901 tgagaaaact tctgcttgag aatcaaatca atcatcagat tgaagttgtt gcagacatga
12961 atgacctcag aaaactggtc agtgaatatg gaaagattgg atcgaatctc aatcagatcg
13021 ccaagcattt taacagtggt ggcagtcagt cacgtgctat agaaaacgaa atccatcagt
13081 gtatcacgga cttattccta ttgagaaaag aggtactgaa actggcaggt ggtatgaatg
13141 gcaatcgtaa aacacattaa aagcagaaat gcaaattact cagctgctat caattatctt
13201 ctgtttgaac atgatgaaaa aaccgggaaa aagattgtag atgaatccgg acgaagcatc
13261 ttgcgaaaag agttttatat ggatgggctg aactgtgatc cgatgtcttt tgacaaggaa
13321 tgtgaactga caaatgctca ttttcacaaa aataaaaaac gtgaagatat caaaagtcac
13381 cactacatta tcagctacga tccgagcgat gttacagaaa acggacttac cggaccaaga
13441 gcacacgcaa tcagtttgga acttgcaaag cagatgtttc ccggatatca ggcactggtg
13501 gttactcata ccgatgggca caatgagtcc ggaaatatcc atacccacat cgtcatcaac
13561 agtgtgcgga agactgcagt agaaaggcag ccttatatgg ataaaccaca tgaggaagcc
13621 gccggttaca agcatcgttc cactgataaa ttcatgaaca cttttaagaa aacagtcatg
13681 gatcggtgcc aacaggaagg acttcaccag atagatcttc ttgctccggc tgaaagaaaa
13741 atcactcaga aagaatatat ggcacagaaa catggccagc aaaaactgga tgagatcaat
13801 caaaaaatca tcgaagatgg tctcaaacca acatccaccg tgttccttac acagaaagaa
13861 tatttgcgaa atgcgattga tgaatgcgct gccacatcca atagttttga tgaattccaa
13921 tctaaacttc tggaactatt tcagatctct gtgattgagc atcgtggcag atacagctat
13981 cttcatcctg atcgacagaa gcggatttct gaacgtgcac tgggaacacg atatggaaaa
14041 gaacacctgg aacagacctt cttacgcaag gatccattgg ccattctcta cgtcagatct
14101 cacctgtgcc tggttgtaaa tctccagaca aatgtgaaag caatgcagag cccggcttat
14161 gctcatcggg taaaactttc taatctgcag caaatggcaa ataccattat ctatgtgcag
14221 gaacatggat ttgatacgca gtcagatttg aaaaaaacac tgcttgcatc aaagcaggaa
14281 ttgaaagaaa tgcagacaca gtttgctcag catcggtcca atctaaggac tctcaatgat
14341 cagatccgct acaccggaca gtactatgca aacaaggaag tttattccca attttcaaat
14401 gcaaagtaca aaggaaagta ccggaaagaa catgcgaagg aaattcagaa atatgaagaa
14461 gccagagatt ggctgagatc cttttatcag gatggaaaga tgacaagcct aaaaaccctg

14521 actttacaaa aagaaaaact gcaacagcag atagcttccg aagaagaagc aataagcagc
14581 ttgaaagaaa aattaaaaga tctggacact gccgatcaga atgttgatgc cattcttcag
14641 atgcagatac cagaaccagt aagatcaaaa aacaaagatc tcgaacgata actatcacaa
14701 aacaaaggag gaactatcac atgacacaag aagaatttaa tattgtattt gaacttcaga
14761 tgagaaagtg tgccgatatt ctggcacata aaaagaaaga atataccggt gacaatattg
14821 accgcctcag tgcattcaag attgctgccg ccttacaaaa ctgcgatcca aaagcagcac
14881 tggccggtat gatgtcgaaa catgtcgtat ccctttacga tatgtgttat tccactctgt
14941 tacattttga catggaacag tgggatgaaa aaatcacaga ctgcatcaat tatctgatcc
15001 ttttaaaagc tctggtaaag gaggaacagg cctatggatc acattgaaac agctatctta
15061 aactgctatg gaatccggga agcagaacag aatatggtct ttgctgccag actcacacaa
15121 cacggtcata cgatccagaa catggcagat ctgatggatc tttaccgtca atcctacagt
15181 cagaagactg tagaaaacat tgcagggctt ccgcatccaa ccgtgcagaa attcatggtc
15241 attaccgtag ctgttgttgg tgcaagcaga cgcttcctgg cacagatcac aagacatcag
15301 aatgaagtaa agtacatgag tgcatctctg cagtacagca attattccgg tcatgctgca
15361 tttgctattc catatggaat catgaaagca gataaagaaa tccaggatat ttacaagaaa
15421 agctgtcagt cagatctgga tcattatgca gagctttgtg ccttaggcat tgatcatgat
15481 tcagccggat atgccacacc tcagggactt cgaaacgtac tgatcatcag cgcgacacct
15541 taccagtgga agcatatgat tggtcagaga acctgcagaa ggaatacgga tgaaaccagg
15601 attgtgatgc tctatatctg gcaaagattg tacaaattaa gcccgattct ttttgcacca
15661 gatctgaccg gtccattctg tcagagagga agctgcatgg aaaaacagat gtcctgtcag
15721 aatcccattt caaaactttg gatgcctgca gatatcttaa aaacagatta tccattgctg
15781 atccggaagg agatgagcag ccataaagat taaactgatt gactttggtc tgaaagcaga
15841 tcatcatcca ttccggccac atgaaaatga tgccggtgct gatgtattca tgccttatga
15901 ttgcgtatta aaaccaggcg aggttgcaag aattccgctt ggattcggtc tcatcatccc
15961 agatggattt gccggatatg tctttccacg aagcagcatg gcagcaaaag gactggtatg
16021 tgaacttcca ccgattgact ccggttatcg tggtgaaatc catgccatta taagcaacgt
16081 gagcacatct tctcagacca ttcataaaga cacacgcgtc ggtcagctgg tgattacacc
16141 ggtcgtgatc gcagattttg tcctggacct tggtgaagaa cgaggaactg gggcattcgg
16201 aagcacagga gaataatact atgattacat ttggaagaaa attaaaacac ttaagacaaa
16261 aaaatcatct cacccaaaaa gaacttggca tggctgttgg atttcccgac agctgtgctg
16321 atgtccggat tgctcagtat gagagtgatg tccgaaagag gatcttatga agctcttcgc
16381 atcaacactc ggtgttccgg ttgaactctt taccgtaccg gtactctcag aacccaggga
16441 gtatgaggcg gcggagtatt ggaggtatga gttaggggcg gaactagatt aagtgtcgta
16501 tactgtaatg aggcaacgga ttatccgctg cctctctttt atcattttag gggtaataat
16561 tgaaaataac ctgtattgag caatcctctt tttctgcgct ttatctacat atctatatat
16621 ttcactttat ttctgttcca tcgataagaa gcacatattt atccggatta tgaacaagat
16681 cataatcttt tccatattca atcggtctaa attctatttg cataatggat tcgccatttt
16741 cgatatcttc tttccaagaa tatacggtca tatcaaggct ggttgcatat ccatgatctg
16801 tagaaaattt tattgaatga aatgaattct ctttgtacat ttttaaaagc agtcttgcaa
16861 atgcttcttt atcagaaata ttttttctgt tagcaactac tgtaaaattt tcatctctat
16921 tcatagaaaa actgccaact acatctggct caccttcttt atactttaat aagctttttc
16981 caattatcat tccaacaata atcgaaaaaa cgatagcaac aaatacaagt gttatattat
17041 ttttcttcat tttattaagt tccttagaat tatatagttg aattaaatct gtatttcaaa
17101 taaaacgcct acaaagtggg cgttttatta tcttaatact atttaatttt tttcatttct

17161 tcaacaaact gttcttgtgt aattctcgca tttaatacat tttttcctaa aaacaattca
17221 tctttatcac atgtagcaaa acatgatttt acccaattaa ctttttatt taattttaaa
17281 taatgtatga aaccagataa ggagaaaacg taaattgtgt taatgatagc agacatgagt
17341 atggagtgaa tatcttattt ggaggaaaaa caaatggtac aatcaatctt tgaggaaatg
17401 ggcggcagat acgaacggca aggcgaatac attttaccct gccttaccat accacccgaa
17461 aaggaacagt cgatagactt attcggacgg cggcacttgg actatctgcg ggagtaccgc
17521 aaaattacat acaccaatct tctcacaagc ggcaggctca atgcttacct tgccgacatt
17581 gacagacagg cacaggaaca ctttgaacgg ctcatagagg gcatgaagca agcacagggc
17641 ataacggaat gcctaaagga agaaaacgcc ttagaatgga ctggaagaac gaataacata
17701 agggcttgtg cgagggagat tgtggaaaag gaaattattt ttgcataaac agatgattag
17761 tggcaggggg aaatcctgcc gctttttctg ctttagtttg tcagcttgac aaataaaggg
17821 ttaaggaata taattagatt cagtattata caaggagtta ataaatatgc ggcaaggtat
17881 tcttaaataa actgtcaatt tgatagtggg aacaaaaagt agcagtcccg tttcactttt
17941 aatatggggc ttagtttttt gtacccagtt taagaatact tttatcatgt aattttatat
18001 gcccgaaaac atataagtgt tttggggcta ttggagttat ttacccagtg ataggagtat
18061 ttatcactgg gtatttttat gcccttttttt gggtgttgat aggaggaaaa tcacatgaaa
18121 ataattaact taggcattct ggctcacgtt gacgcaggaa agacaacatt aacggagagt
18181 ttattgtata ccagtggtgc aattgcagaa ccagggagcg tagataaagg cacaacaagg
18241 acagatacaa tgaatttgga gcgtcaaagg ggaatcacta tccagacagc agtgacatct
18301 tttcagtggg aggatgtaaa agtcaacatt atagatacgc caggccatat ggattttttg
18361 gcggaagtat accgttcttt atccgtatta gacggagcag tattattagt ttctgcaaag
18421 gatggcatac aggcacagac ccgtatactg tttcatgcac tacagacaat gaagattccg
18481 acaatttttt tcatcaataa aattgaccaa gaggggattg atttgccaat ggtatatcaa
18541 gaaatgaaag caaagctttc ttcggaaatt atagtgaagc aaaaggttgg gcagcatcct
18601 catataaatg taacggacaa tgacgatatg gaacagtggg atgcggtaat tatgggaaac
18661 gatgaactat tagagaaata tatgtcaggg aaaccgttta aaatgtcaga actggaacag
18721 gaagaaaaca ggagattcca aaacggaacg ttatttcccg tttatcacgg aagcgctaaa
18781 aacaatctgg ggattcggca gcttatagaa gtgattgcca gtaagtttta ttcatcaacg
18841 cctgaaggtc aatctgaact atgcgggcag gtttttaaga ttgaatattc agagaaaagg
18901 cggcgttttg tttatgtgcg tatatatagc ggaacattgc atttgaggga tgttattaaa
18961 atatctgaaa aagagaaaat aaaaatcaca gagatgtgtg ttccgacaaa cggtgaatta
19021 tgttcatccg atacagcctg ctctggtgat attgtaattt taccaaatga tgttttgcag
19081 ctaaacagta ttttggggaa cgaaatgctg ttgccgcaga gaacatttat tgaaaatcct
19141 ctccctatgc tccaaacaac gattgcagca aagaaatctg aacagcggga aatattgctt
19201 ggggcactta cagaaatttc agatggcgac cctcttttaa aatattatgt ggatactaca
19261 acgcatgaga ttatactttc ttttttgggg aaagtgcaga tggaagtcat ttgtgccatc
19321 cttgaggaaa aatatcatgt ggaggcagaa ataaaagagc ctactgttat atatatggaa
19381 agaccgctta gaaaagcaga atataccatc cacatagaag tcccgccaaa tcctttctgg
19441 gcttctatcg ggttgtccat agagccgctc cctattggaa gtggagtgca gtatgaaagc
19501 agagtttcac ttggatattt aaaccaatcg ttccaaaatg cggttatgga gggggttctt
19561 tatggctgcg agcaggggct gtatggatgg aaagtgacag actgtaaaat ctgttttgaa
19621 tatggattgt attatagtcc tgtaagtacc cccgcagact ttcggctgct ttcccctatc
19681 gtattggagc aggctttaaa aaaagcaggg acagaactat tagagccata tctccacttt
19741 gaaatttatg caccgcagga atatctctca cgggcgtatc atgatgctcc aaggtattgt

19801 gcagatattg taagtactca gataaagaat gacgaggtca ttctgaaagg agaaatccct
19861 gctagatgta ttcaagaata caggaacgat ttaacttatt tcacaaatgg gcagggagtc
19921 tgcttgacag agttaaaagg ataccagcca gctactggta aacttatttg ccaaccccgc
19981 cgcccgaata gccgtataga taaggttcgg catatgttcc acaagttagc ctaacagctt
20041 gcataagtcg tataaaatga gatttgaaag gattagaagc taattatgat gaaatgcgaa
20101 tggatattat gtcctgtttg tgggagcaaa acccgtaata aaattaggaa ggacactgtt
20161 ttgaagaatt atccccttta ttgtccaaaa tgcagacaag aaagattgat taaagttgac
20221 aacttgaaga taactgtcat caaagagcca gacgcttaag acgcagagcc gatgaaattg
20281 tggaacaatt cacgaatcat cggctctttt tgtttcgtat ttgaaagaac aaactcacca
20341 aataaaaaaa acgatattcg ggtgggttat tttgttatac cctaaattac cctctgaatt
20401 tgtttttaaa tttggagggt ttttatgttc ctttttcgg gcggttgccc atctgcttat
20461 ataaagcaaa gtttatttat acatagcaca tcagggaatg gcgggaagcc agttaaaaaa
20521 atccctgcca gtgcaacggt acttcccacc atgaaggtag aagtacaatc tccatacaac
20581 agaattagta tttcccataa ccgtaaaggt cacaaagcct ttgcggtttt tcttttgtcg
20641 tttttggttg gaaagtgccg caggactatt attggtgtta agtgtcgttc gccgcctttc
20701 aatctggatt tttatgtttt caaagattca gattgattgg aggaaaaaag aatggcatat
20761 aacaacggac gtgaggacag gaagtggcgt atctggaaag aagccgagga aaaagtgttg
20821 cgggaacatg gagttgatga agctaccatc gaacaaatcc gtattgacga cagggcagac
20881 tttaattcca acaggcggtt ttaccattgg acaagcgatt tcggggaata ccttgaggga
20941 atggcagaca gggaacagtc agccgaagta aagaccgttg aagatttact ggatgaaatc
21001 gaagatgaaa atctgtatca ggcattactt acggtggaca ggcgtacact gcaaatcgtt
21061 ctgctgaaaa tgaaagggta ttccacaaaa gagattgccc cgcttgtgca tttaacgacg
21121 ggtgctatct atgcacggtt agaccatctt agaaaaaagt tgcggaagat tttatagtca
21181 tctaatacag catgctttct tatgggctat tgggtggggg ataaaattct cccacccaat
21241 tttagaataa gtgaataaaa tccctgtcca tagagaatac taaaaagttt gtgcaaaatc
21301 ttgacagcca tattgccgct atggtattct gaaataccaa tggggaaatt ggaagattga
21361 aaattaaata agcacataga tataagattg aaatgtcagc caatcggcaa ggctgacatg
21421 ggttttgaat ttcaaagccg ctacatagca ttgcgaatca gagcgggagg gaatcctctt
21481 gtcgttcgtg atgcagtgtg gcggcttttt catttatcca aaagtcaaag gcaataaaat
21541 ccagaacgga ggtggaaagg acataggatt tccttttcgg agggtaaagg caggcattaa
21601 gaaagccgtt acacagaaaa atgctctgcg gagttgtcca cagagatagc gagcatcgga
21661 ttgtgtcagc cacaatccaa tccacaccct aaaggcgtgt ggactaactt aggggacaac
21721 ccctagctac cccgataaaa ggaaaaataa aactggagga ttaagaaaaa tgagtagaga
21781 aaaatcagaa aaagtatgca ttacttttag attgagcgaa gaagaacatg aaaaattaaa
21841 gcagtattca tccgcctgcg gtctgtccac cgcagaattt atgcgtcagt tatgcagggg
21901 aaacgcaccg cagccacagc cagagaaaga attttgggaa ctattaggca cgctttacga
21961 agtacacgta gcctttaaaa agtgtatccc ctacgcccct tccgctgatg aaatatgcag
22021 ggagattgag gattttatct tggaactgca acggaattat accctcccac agcagtttga
22081 catggaaaaa ttgacggaac agggggcagt ctgatatggc ggcaacgagc ctctggcaca
22141 tcgaaggacg gttgaaagac ttaatttctt atgtggaaaa cccagaaaag accgtgccga
22201 ataaggagat gcaggacttc tttgacgtgt tctcctatgt gaagaatcca cagaaaacgg
22261 acaatggaga gtttgtttcc gccatcaact gtttaaaaga aacagcgtta cagcagatga
22321 ttctaacgaa aaagcagtac cagaagacag gcgggtatat cgcttggcac ggctaccaga
22381 gcttcaagcc agacgaagta acgcccgaac aatgccacga aatcggctta aaacttgcga

22441 gggaaatgtg gggcgacaag taccagataa tcgttgtcac ccaccttgac agggggcatc
22501 tccacaatca cttctgcttc aactctgttt cctatctgga tgggagcaaa tataattact
22561 caaaatcgga gcagaaacgg ctaagggaaa cgtcagaccg cttgtgcaga gagtacggat
22621 tatcggtgat tgagaacccg aagaaagcac cgtcaagacc gctgtatctg gatgaaaaga
22681 gcggcaagcc gacacgctac aacgtctatc tggaagattt agcggaggca atcagcggca
22741 gtcgggatat tccgcacatg atgaagtatc taatcgacaa aggctacgag gttgattttt
22801 cgggcgggca ttggaaaatg aagctgccgc agtataagca tttcacaaga ttagatacgc
22861 ttgatgaacg cttaacgccc gattttatcc gttcacattt aggcacaagg gcaagatatg
22921 ggaactacaa agcaaagatt tcctactctc cccatatgcc cgaagaatat aaaaaggctt
22981 ggaaaccgca ccagaaaaca aggggaatct tagcgttgta ttactattgg tgctatcagt
23041 tggggatatt tcccaaaggc acggactaca agccgacaag cccgcttatg aaagaggagc
23101 ttcggaaact ggatgagatt accacacagg tacgctatat gtcggagcat aacatctcca
23161 ccctctccga tttgcacgcc gacagggaga agaaccagac ggaaatggac aggcttacca
23221 actatcggcg gcatttgcag aataagatac ggcgtgcttc acccgccgag aaagaaacgc
23281 tccgagaaga aaaaaacggg gtgacggaac ggataacgga gctgcgaaag cggctgaaat
23341 atgcggacgg gattgaaaaa cgctctgccc acattgatga ctgcttgaac caaatccacg
23401 atacggttga aaatcaaagg tcaaaccaga atggcaggac agaccgcaag agggaggacg
23461 cattgcgatg agcagactga cagaacagga acttacggaa attttagacg aaaataaggg
23521 tatctcaatg gtaaacccga atagggtgta cccgcccctg ccgcctgctg aaaacattat
23581 gccccttatg cgtatccctg aacaggcaac gctctgcgag aaaatcggcg gcacggaata
23641 tacagtgacc gcacatttcc gagaggacgg gagggacttg cttgggcttc tcagcaggat
23701 attcatgcgg gacacagcag attacagttt tgattccgag ggttatgatg aggattaaga
23761 aatatcggaa agattatcgc tttaaagcgt tgaagtttga ccgcaggcgt ggtatactgt
23821 atctacactt catactatcg tgtctgctgt cggaaaggag aaagcattgc aaagacagca
23881 ggaagaaatc accgccttgt actgtcgtct gagtcaggac gatggacgtg aaggcgagag
23941 caacagtatc gtaaatcaaa aagaatatct attcaagtac gctaaggaac acgggtttcc
24001 aaatcccacc tactatgtgg acgatggcta tacgggtacg aattttgctg ggttagata
24061 gcgatacttt ttacatctac catgttgcgg cggtcatggt tgacaacttg cgtcggctcc
24121 tgctcggtat caacctgtcc gacaaagcta taatggattt cgattttccg ggtgttggtt
24181 tcccggtcta attcatgtat caaaatacgg tcgataaact catggacgtt ttcataggtc
24241 aattcctgta tgtcgctgta tttcccgaca atctgaataa accttgatat atcccttttt
24301 cgctcggtaa cggttgcgat ctgctgttgt aactcgtcta tccttgcggc aagggatttc
24361 ttttcgtcct catagccgga agttagaaac acaaaccgtt catctgtcag tctgcctaat
24421 gcgttgtcct catacagctt gcggaaaagc gtgtcaagtt cggtcatacg cgcctgtgct
24481 ttgaaaagtt ccttttgctg ctgcgctaat gcctttctcg cttccatgtc gccgtactcg
24541 gtagctttac agataaagtc ctgttcgtgt tccttaacat accgcaaaac ccgccgcata
24601 tcttccagca ctaaatcaag aagtacgttt ttgcggatat aatgagaggt acactcgctg
24661 cctgtccttg ccctgttgcg ccatgcgccg caagaatagg aaaacttgcg cggctcgata
24721 tttacgccct gttggtaata catcttatgt ctgcaaccag cgcaaaacag caagccggaa
24781 aaaatatcaa tctccgctgc ttttgtcggg cggtggcggg tagcaatccg cttttgcgca
24841 agttcaaagg tttcctcgtc aacaagcggc tcatgggtgt ttgggaagaa atagcgttgt
24901 tcttcttcgt tcttccgggc ctttttcgac ttgtaggata ccttgtgagt tttcgcggtt
24961 atggtatgcc ctaaatattc ctgccttgct aatatgtcat agagcgtttt gtccggccaa
25021 gtataggggg cgatcattgc ccgctgatac cgcgcctgtc ctgtacgctg atagcgcaac

25081 gctccaatcg tcaagacttt gttttccgca agccattttt ggatttcgca catacgcacg
25141 ccgctaacaa acatagcgaa aacctgtttg acaatcggcg cggtttccgg gtcgggtata
25201 aggtggtggc ggttgttcgg gtctgggata tagccgtatg gatattcccc gttgacgcgc
25261 tcgccttttt gcgcctgtgc ctgtttaact gcgcggattt tcttgcttgt gtcgcgggcg
25321 taaaactcgt taaaccagtt ccgcaagggg gtaaactcgt tatcctcccg cgctgtgtca
25381 actccgtcgt tgatcgcgat atagcgtact ccgttttcgg gaaatacaat ctctatgagt
25441 tcgcccgttt tcagatagtt tctgccaaga cgggaaaggt ctttggtaat gaccgtcgcc
25501 actcgtccgg cttccacttc ctgcaacatc gcttgcagcc cctcgcgctc aaaactcacg
25561 ccggaaaatc cgtcgtccac aaaaaacttt gtgttcaaaa aatggtgttc gtccgcatag
25621 cgttgaagta tcattttttg gtgctgtatg cttccgcttt ctccggcttg catatcctct
25681 tgacttaaac ggcagtaaag ggcggtgatt ttattcgact gtaacattag tcctcctttc
25741 cgacagccga ataaacaggt ataatgtgtt gctatcatta taccatatcc cgccgcctaa
25801 tgcactactc ggacgctaaa aatcccggat ttccgggctt tttctgcaaa tcctttacaa
25861 tgagtttctc aatcttttta tggattgtat cggttgcctt ttcactcggc aacgcttgaa
25921 caagataggt gattttccct attttgcgtt cggttgtaat tgtctgtttt ttatccatta
25981 gaaaaacctc cttttcctgt atggataaac tatattcgtc aaaaggcaaa aacgggcggt
26041 tgttagctgc aacctcacgg gagtttcacc ccggcccatg cttatgggtg cgccgcgcaa
26101 tactttgagg gtgttcaacg cgggagtatc attgtgccgc cttgtatgtc gtcgcccaca
26161 agctgctaaa cctgttttac ggcgcggtag ttctcgctcg gcttttcccc tatggggaaa
26221 agctgtcatg gcgtacccgc cgactggctc ggtacagacg gaatgtaccc gtttgcctgt
26281 tatgctgcgc accaaaggcc gctttctttg tcaaagaaca ataggctttg tcggcctgca
26341 aaagcacatc aacagcggat atgcttttgc ggaacgacaa atagcccata acgggaagcg
26401 tggaaagggg acacgctccc cgcatgggct aaaagattat cctacatgaa aagcaagggt
26461 gcgggtaata aggcgcactt gcagcctgtt acgcatttcc tcgtccacat aggaataggt
26521 attgcccgcg tcgtctttca gcgtgcgggt acaaagtttc gctatgtaac cctcgtagtg
26581 cttcaagatc gcgcacatgg cggttgtgtc gccgtttgcc gctgcggaaa tgacagggaa
26641 cggcaacaga ttttcagact tcctaacggt attcatcatc tgcttttccc tccaaatact
26701 gtttgatttt cgcaagcgcg gatttcctgt gccggaaaac cgtcgtgcgt acaacattca
26761 gcagttcgcc aatttccgcg tcgctcatat ccaagaagta ggacaaaagg ataatgtcgc
26821 gtttcctttc gggcaaagcg ttaagggctt cggcaagcag ttcatttttg acgagtacat
26881 caaagccgga cacttgaaaa cggaaataat cgctttcgta ttcgtccgtt gtgaaaagct
26941 gcgcgagttc gctttcgctc aaatccgaaa aagtaacttc gcgtgctgcg cgtttcgcaa
27001 gagtgcggcg gtggcttttc gcttcgccga ccaaagtttt ctttgctaat gcgtcgtact
27061 gatgttgtat tctttccttg tcggaagaag atagctccat agagttcacc tccttcccgc
27121 gtggagtaga ggacgggagt taagggcttg tcctctctgc ccctttcgct ccgtacccgt
27181 tcgggagatg gctcttgagt gcgcttttca gaaaaaagtt gaaaaaagca aaatgcgccc
27241 gtccgcaaga cgcggacggg cgcaaaggaa atgtaagcag tagctaaatg tattgacagt
27301 tcacattcat agccggaata tcccgcttgc ggagcatagc tttttttgac cgcaaagcat
27361 taggcgggtt acagtaaata aaaatggaca cggcgatacc tccccgatac cgccgtatcc
27421 ttaaaaaagg gcgactttaa tacactaccc gcaatccatt ctataatagg gaacagcggc
27481 ttttgcgcaa tattaaataa aaaagccgat aacacatagg ttttttgacc taatgcgtta
27541 tcggctctgc gtctatgcgt ctggcacttt taatcatttt tttttgaatt tattatatgt
27601 gttagctaac tgtccacaag ccgcgttaat ctctctgcca tgagaaactc tcatagtaac
27661 ttcaagtcct gcttgctcta attgatgttt gaatgcaact atttcccgtt tctgtggtgc

27721 tttaattctt gaattgcttg ttgggttgta ttgtaacacg ttaatcataa cttttttgcc
27781 ccgaaaccat tttgcaagtt gtcttacatc tgagggccgg tcatttatac ccggtaaaag
27841 caaatacgca aagacaattt tgcgattatg cctttcagaa taggataagg cttgcttaac
27901 aacatcttca atagcatata tgcgcatgtg aggaataata cgatttcttg cagattgtgt
27961 tgctgcgtgt aaagatattg tcaactgaat tttaagatgt tcctcgcgca atttttttaa
28021 ttgatcgacc ggaccaactg ttgatatggt aatgccgtcg gttggaaagt tgagcccata
28081 tctatctcgg agaatatgga ttgcttttat caagttgtca taattgaata aaggctctcc
28141 catacccata aaaacgatac ggtttacttt tcgacgcaac aatataatct gttgtacgat
28201 ttctgacgat gttagattac gaacaaagcc atttcgcccg gactcacaaa aaatacaacc
28261 aacaggacaa ccgacttgtg tgctcacgca aacagttcca ccatctcgcc gcttgataaa
28321 aaccgtttca atgtatttgt tgtctttcag ttcgtaaaca tacttttcag tatcactgct
28381 tttgcaaata ttttttacca acattgatag atttttttg cgtggtaatt gcttatataa
28441 ttcttcatat aaggcttttg cttcattctc gccaataact tccgacattt ctttgtaagt
28501 aaatccgtat ggatcattcc gtacttccgc aggcgtatat ttaggtaaac gtttcatttt
28561 tatatccttt cttctaaata ataaaagttt gtttttctgt atttttgatt acaatctcta
28621 atttttctac atcaatgata tgcgtcaatt tgcatttagg gcagaaaaga ggaaaatctt
28681 ttaatacagt attatgaaat acttgaactc tcgtctttcc gttacaaatc ggacatttta
28741 tccaatattt tttaagcatt atatttcact cgtccttttt acacaaaaaa atgcaggtca
28801 atcctcaaca tgagcattga cctgcattta taatgcacag agcagtacaa caaaactaag
28861 gcatagcttg cactatgtct atactatatc tatacgtcgt tagtttttg tatagcatcc
28921 gcaaaataag catgacaaaa aagcccatgt tgaaaatggg aaaatccttt ttttcaatgc
28981 ttatctaata cgcatgctat gcaacataaa cgccgcctcc ttttacataa attttatttt
29041 atcagaaaat cagtctactg tcaatacaca acaggaagta tttaacctaa tgatatgaat
29101 tgtgtggaca tatccaaaaa gaaaggtgaa ctgtaaaatg taacagggtg aatgaaaatg
29161 gcaaaaagac ccgtaccatt gtacgacttt aaggctttcg gggcagctat aaaagccgcg
29221 agaaatgaat acggcgagag ccgcaaaaag gtaagcgacg agttatatat ttccccgcgc
29281 taccttgcga atatcgagaa caagggacaa cagccgagtt tacaggtatt ctatgacctt
29341 gtaacccggt atcatatttc ggtagatcaa tttttcttcc cgaacagcaa tgcggagaaa
29401 tccaccgggc ggcggcagct tgacgcgctg ctggacggta tgagcgataa aggcatacgg
29461 attgtaaccg caacagcaag ggagataacg gaagtcgaaa aagcagagga ttaacctcac
29521 aatatgagaa atcgggagat tgcagcgcat ggcggctgca atcttttttt gcgtccaaaa
29581 tcaaaggaac gcgcaacaaa tgtagtatac cattcatccg tgttatctgc agaatatttt
29641 gcgttattta atattgaatt tcctgtttta tttatagtta taatttcttg atctttgccg
29701 ttgatcattt gcttgattct cccctgcatg aatttctcat ttgtcttgat tagattctaa
29761 ttcttttaat ttctttttca tttcttcata ttcatcgaaa taagaaatta atgctctttc
29821 aacagcaacc gtttttgtat gtcctgcatc tttacaaaaa ttttctaatc gctcataaat
29881 aggagtttca atacatacat taaggtatgt tccattcttt tttggtcttg ccattttttt
29941 ctccctgcat ttaatactat tacatgaatt ttcctcaaat aattgtacca ctgaaaacta
30001 cttaaatcaa ctattcacta cttgttatca aatatataaa agataaaatc aggttttgta
30061 ctactaaatg attttaaaca aaataaggcc caaaagggaa ctttcccttt tgagcctata
30121 cactttatca gctaacatat tttgctttca tttttcctca tggtcatgag gcttttttct
30181 cttaaatcgc attctatgaa tcctttgatt gttccttctg ttcttcaata attcgttctg
30241 ttgctttatg aattctgtcc ttagaatagg gtggtcttaa attcaatgac aaccggccct
30301 ttttgattat ata

```
//
DEFINITION     Integrative and mobilisable element IME_4
   CDS           complement(1..1161)
                 /color=9
                 /locus_tag="C83_01573"
                 /note="Tyrosine recombinase, similar to AA
                 sequence:UniProtKB:P0A8P6 protein motif:CLUSTERS:PRK00236
                 protein motif:Cdd:COG4974 protein motif:TIGRFAMs:TIGR02224
                 protein motif:Pfam:PF00589.16"
                 /product="Tyrosine recombinase [recombinase]"
                 /"ID=C83_01573
                 /protein_id="gnl|SC|C83_01573"""
                 /codon_start=1
   fasta_record    1..51034
                 /ID=C83|SC|contig000001
                 /label=C83|SC|contig000001
                 /note="C83|SC|contig000001"
                 /colour=5
   CDS           complement(1286..1525)
                 /color=9
                 /locus_tag="C83_01574"
                 /note="DNA-binding domain  excisionase family
                 helix-turn-helix domain-containing  protein, protein
                 motif:TIGRFAMs:TIGR01764 protein motif:Pfam:PF12728.1"
                 /product="DNA-binding domain  excisionase family
                 helix-turn-helix domain-containing  protein [excisionase]"
                 /"ID=C83_01574
                 /protein_id="gnl|SC|C83_01574"""
                 /codon_start=1
   CDS           complement(1970..2575)
                 /ID=C83_01575
                 /color=7
                 /locus_tag="C83_01575"
                 /note="RNA polymerase sigma factor  Sigma-70"
                 /product="RNA polymerase sigma factor  Sigma-70
                 [transcriptional regulator]"
                 /protein_id="gnl|SC|C83_01575"
                 /codon_start=1
   CDS           complement(2645..2893)
                 /ID=C83_01576
                 /color=1
                 /locus_tag="C83_01576"
                 /note="hypothetical protein"
                 /product="hypothetical protein"
```

/protein_id="gnl|SC|C83_01576"
/codon_start=1
CDS complement(2890..3624)
/color=8
/locus_tag="C83_01577"
/note="ImmA/IrrE family metallo-endopeptidase, protein motif:Pfam:PF06114.7"
/product="ImmA/IrrE family metallo-endopeptidase [pepidase]"
/"ID=C83_01577
/protein_id="gnl|SC|C83_01577"""
/gene="ImmA"
/codon_start=1
CDS complement(3621..3881)
/ID=C83_01578
/color=1
/locus_tag="C83_01578"
/note="hypothetical protein"
/product="hypothetical protein"
/protein_id="gnl|SC|C83_01578"
/codon_start=1
CDS 4086..4508
/color=10
/locus_tag="C83_01579"
/note="Replication initiation protein RepA, similar to AA sequence:UniProtKB:P96631 protein motif:TIGRFAMs:TIGR03070 protein motif:Pfam:PF01381.16"
/product="Replication initiation protein RepA [replication]"
/"ID=C83_01579
/gene="repA"""
/protein_id="gnl|SC|C83_01579"
/codon_start=1
CDS 4569..5078
/color=8
/locus_tag="C83_01580"
/note="ImmA/IrrE family metallo-endopeptidase, protein motif:Pfam:PF06114.7"
/product="ImmA/IrrE family metallo-endopeptidase [pepidase]"
/"ID=C83_01580
/protein_id="gnl|SC|C83_01580"""
/gene="ImmA"
/codon_start=1

CDS 5089..5415
/color=1
/locus_tag="C83_01581"
/note="hypothetical protein, ab initio
prediction:Prodigal:2.60"
/product="hypothetical protein"
/"ID=C83_01581
/protein_id="gnl|SC|C83_01581"""
/codon_start=1
CDS 5431..5910
/color=8
/locus_tag="C83_01582"
/note="Flavodoxin, protein motif:CLUSTERS:PRK07116 protein
motif:Pfam:PF12682.1"
/product="Flavodoxin [electron transfer]"
/"ID=C83_01582
/protein_id="gnl|SC|C83_01582"""
/codon_start=1
CDS 5926..6633
/color=8
/locus_tag="C83_01583"
/eC_number=1.1.1.100
/note="3-oxoacyl-[acyl-carrier-protein] reductase FabG,
similar to AA sequence:UniProtKB:Q9KQH7 protein
motif:CLUSTERS:PRK05565 protein motif:Cdd:COG3347 protein
motif:TIGRFAMs:TIGR01830 protein motif:Pfam:PF00106.19"
/product="3-oxoacyl-[acyl-carrier-protein] reductase FabG
[oxidoreductase]"
/"ID=C83_01583
/gene="fabG"""
/protein_id="gnl|SC|C83_01583"
/codon_start=1
CDS 6646..7689
/color=8
/locus_tag="C83_01584"
/eC_number=3.5.1.-
/note="O-acetyl-ADP-ribose deacetylase RNase III inhibitor
Macro domain-containing protein, similar to AA
sequence:UniProtKB:P0A8D6 protein motif:CLUSTERS:PRK00431
protein motif:Pfam:PF01661.15"
/product="O-acetyl-ADP-ribose deacetylase RNase III
inhibitor Macro domain-containing protein [hydrolase]"
/"ID=C83_01584
/gene="ymdB"""

/protein_id="gnl|SC|C83_01584"
/codon_start=1
CDS 7726..7959
/ID=C83_01585
/color=1
/locus_tag="C83_01585"
/note="hypothetical protein"
/product="hypothetical protein"
/protein_id="gnl|SC|C83_01585"
/codon_start=1
CDS 8035..8817
/color=4
/locus_tag="C83_01586"
/note="Conjugal transfer protein TraX, protein
motif:CLUSTERS:PRK13882 protein motif:Pfam:PF05857.5"
/product="Conjugal transfer protein TraX [mating pair
formation]"
/"ID=C83_01586
/protein_id="gnl|SC|C83_01586""""
/gene="traX"
/codon_start=1
CDS 8930..10438
/color=8
/locus_tag="C83_01587"
/eC_number=2.7.7.7
/note="Nucleotidyltransferase/DNA polymerase involved in
DNA repair, similar to AA sequence:UniProtKB:Q47155
protein motif:CLUSTERS:PRK03103 protein motif:Cdd:COG0389
protein motif:Pfam:PF00817.14"
/product="DNA methylase Nucleotidyltransferase/DNA
polymerase involved in DNA repair impB/mucB/samB family
protein [methyltransferase]"
/"ID=C83_01587
/protein_id="gnl|SC|C83_01587""""
/codon_start=1
CDS 10435..10842
/ID=C83_01588
/color=1
/locus_tag="C83_01588"
/note="hypothetical protein"
/product="hypothetical protein"
/protein_id="gnl|SC|C83_01588"
/codon_start=1
CDS 10969..11628

/color=1
/locus_tag="C83_01589"
/note="Uncharacterized protein encoded in toxicity protection region of plasmid R478 contains vWF, protein motif:Cdd:COG4245"
/product="Uncharacterized protein encoded in toxicity protection region of plasmid R478 contains vWF domain"
/"ID=C83_01589
/protein_id="gnl|SC|C83_01589""""
/codon_start=1
CDS 11625..12287
/color=1
/locus_tag="C83_01590"
/note="Protein of unknown function DUF541-containing protein, similar to AA sequence:UniProtKB:Q44642 protein motif:CLUSTERS:PRK11087 protein motif:Pfam:PF04402.8"
/product="Protein of unknown function DUF541-containing protein"
/"ID=C83_01590
/protein_id="gnl|SC|C83_01590""""
/codon_start=1
CDS 12553..12882
/ID=C83_01591
/color=4
/locus_tag="C83_01591"
/note="Plasmid mobilization relaxosome protein MobC"
/product="Plasmid mobilization relaxosome protein MobC [relaxosome]"
/protein_id="gnl|SC|C83_01591"
/gene="mobC"
/codon_start=1
CDS 12843..14207
/color=4
/locus_tag="C83_01592"
/note="Relaxase/Mobilisation nuclease domain, protein motif:Pfam:PF03432.8"
/product="Relaxase/Mobilisation nuclease domain [relaxase]"
/"ID=C83_01592
/protein_id="gnl|SC|C83_01592""""
/codon_start=1
CDS 14402..14611
/color=7
/locus_tag="C83_01593"

/note="XRE family transcriptional regulator
helix-turn-helix domain, protein motif:Cdd:COG3655"
/product="XRE family transcriptional regulator
helix-turn-helix domain [transcriptional regulator]"
/"ID=C83_01593
/protein_id="gnl|SC|C83_01593"""
/codon_start=1
CDS 14684..16585
/color=10
/locus_tag="C83_01594"
/note="DEAD/DEAH box helicase family protein Superfamily
II DNA/RNA helicase SNF2 family, protein
motif:Pfam:PF00176.17"
/product="DEAD/DEAH box helicase family protein
Superfamily II DNA/RNA helicase SNF2 family [helicase]"
/"ID=C83_01594
/protein_id="gnl|SC|C83_01594"""
/codon_start=1
CDS 16618..17526
/color=2
/locus_tag="C83_01595"
/eC_number=2.7.7.-
/note="Aminoglycoside 6-adenylyltransferase Streptomycin
adenylyltransferase, similar to AA
sequence:UniProtKB:P17585 protein motif:Pfam:PF04439.6"
/product="Aminoglycoside 6-adenylyltransferase
Streptomycin adenylyltransferase [antibiotic resistance]"
/"ID=C83_01595
/gene="aadK"""
/protein_id="gnl|SC|C83_01595"
/codon_start=1
CDS 17523..18065
/color=2
/locus_tag="C83_01596"
/note="streptothricin N-acetyltransferase Sat4 , protein
motif:CLUSTERS:PRK10975 protein motif:TIGRFAMs:TIGR01575
protein motif:Pfam:PF00583.18"
/product="streptothricin N-acetyltransferase Sat4
[antibiotic resistance]"
/"ID=C83_01596
/protein_id="gnl|SC|C83_01596"""
/gene="sat4"
/codon_start=1
CDS 18158..18952

/color=2
/locus_tag="C83_01597"
/eC_number=2.7.1.95
/note="Aminoglycoside 3'-phosphotransferase, similar to AA sequence:UniProtKB:P0A3Y5 protein motif:Cdd:COG3231 protein motif:Pfam:PF01636.17"
/product="Aminoglycoside 3'-phosphotransferase [antibiotic resistance]"
/"ID=C83_01597
/gene="aphA"""
/protein_id="gnl|SC|C83_01597"
/codon_start=1
CDS 19228..19800
/color=7
/locus_tag="C83_01598"
/note="deoR-like helix-turn-helix domain protein, protein motif:CLUSTERS:PRK11886 protein motif:Pfam:PF08279.6"
/product="deoR-like helix-turn-helix domain protein [transcriptional regulator]"
/"ID=C83_01598
/protein_id="gnl|SC|C83_01598"""
/codon_start=1
CDS 19824..20375
/ID=C83_01599
/color=10
/locus_tag="C83_01599"
/note="DEAD/DEAH box helicase family protein  Superfamily II DNA/RNA helicase  SNF2 family"
/product="DEAD/DEAH box helicase family protein Superfamily II DNA/RNA helicase  SNF2 family [helicase]"
/protein_id="gnl|SC|C83_01599"
/codon_start=1
CDS 20506..22044
/color=9
/locus_tag="C83_01600"
/note="IS21 family transposase Integrase core domain, protein motif:Cdd:COG4584 protein motif:Pfam:PF00665.20"
/product="IS21 family transposase Integrase core domain [DDE recombinase]"
/"ID=C83_01600
/protein_id="gnl|SC|C83_01600"""
/codon_start=1
CDS 22038..22784
/color=9

/locus_tag="C83_01601"
/note="IstB-like ATP binding protein  putative transposase
subunit, protein motif:CLUSTERS:PRK09183 protein
motif:Cdd:COG1484 protein motif:TIGRFAMs:TIGR00362 protein
motif:Pfam:PF01695.11"
/product="IstB-like ATP binding protein  putative
transposase subunit [ATPase]"
/"ID=C83_01601
/protein_id="gnl|SC|C83_01601"""
/codon_start=1
CDS 23026..24258
/color=10
/locus_tag="C83_01602"
/note="DEAD/DEAH box helicase family protein  Superfamily
II DNA/RNA helicase  SNF2, protein motif:CLUSTERS:PRK04914
protein motif:Pfam:PF00271.25"
/product="DEAD/DEAH box helicase family protein
Superfamily II DNA/RNA helicase  SNF2 family [helicase]"
/"ID=C83_01602
/protein_id="gnl|SC|C83_01602"""
/codon_start=1
CDS 24271..24927
/ID=C83_01603
/color=1
/locus_tag="C83_01603"
/note="hypothetical protein  DUF4391-containing protein"
/product="hypothetical protein  DUF4391-containing
protein"
/protein_id="gnl|SC|C83_01603"
/codon_start=1
CDS 24924..25745
/ID=C83_01604
/color=1
/locus_tag="C83_01604"
/note="hypothetical protein"
/product="hypothetical protein"
/protein_id="gnl|SC|C83_01604"
/codon_start=1
CDS 25763..26374
/color=8
/locus_tag="C83_01605"
/note="Molecular chaperone Tir, protein
motif:Pfam:PF08937.5"
/product="Molecular chaperone Tir [chaperone]"

/"ID=C83_01605
/protein_id="gnl|SC|C83_01605""""
/codon_start=1
CDS 26427..27401
/ID=C83_01606
/color=1
/locus_tag="C83_01606"
/note="hypothetical protein"
/product="hypothetical protein"
/protein_id="gnl|SC|C83_01606"
/codon_start=1
CDS complement(28215..29984)
/color=4
/locus_tag="C83_01607"
/note="Antirestriction protein ArdA, protein motif:Cdd:COG4734 protein motif:Pfam:PF07275.5"
/product="Antirestriction protein ArdA [anti-restriction modification]"
/"ID=C83_01607
/protein_id="gnl|SC|C83_01607""""
/gene="ardA"
/codon_start=1
CDS complement(29998..30213)
/ID=C83_01608
/color=4
/locus_tag="C83_01608"
/note="Putative tranposon-transfer assisting protein TTRAP"
/product="Putative tranposon-transfer assisting protein %28TTRAP%29"
/protein_id="gnl|SC|C83_01608"
/codon_start=1
CDS complement(30215..31408)
/color=10
/locus_tag="C83_01609"
/note="DNA mismatch repair protein MutS, protein motif:CLUSTERS:PRK05399 protein motif:TIGRFAMs:TIGR01070 protein motif:Pfam:PF01624.14"
/product="DNA mismatch repair protein MutS [replication]"
/"ID=C83_01609
/protein_id="gnl|SC|C83_01609""""
/codon_start=1
CDS complement(31421..33481)
/color=4

/locus_tag="C83_01610"
/eC_number=2.7.7.-
/note="DUF3849-containing protein  antirestriction
protein, similar to AA sequence:UniProtKB:P27189 protein
motif:Cdd:COG4227 protein motif:Pfam:PF12960.1"
/product="DUF3849-containing protein  antirestriction
protein [anti-restriction modification]"
/"ID=C83_01610
/protein_id="gnl|SC|C83_01610"""
/codon_start=1
CDS complement(33493..35562)
/color=10
/locus_tag="C83_01611"
/eC_number=5.99.1.2
/note="Type III DNA topoisomerase, similar to AA
sequence:UniProtKB:P14294 protein motif:CLUSTERS:PRK07726
protein motif:Cdd:COG0550 protein motif:TIGRFAMs:TIGR01056
protein motif:Pfam:PF01131.14"
/product="Type III DNA topoisomerase [topoisomerase]"
/"ID=C83_01611
/protein_id="gnl|SC|C83_01611"""
/codon_start=1
CDS complement(35628..36413)
/ID=C83_01612
/color=8
/locus_tag="C83_01612"
/note="DUF4366-containing protein  bacteriocin"
/product="DUF4366-containing protein  bacteriocin"
/protein_id="gnl|SC|C83_01612"
/codon_start=1
CDS complement(36406..36651)
/ID=C83_01613
/color=1
/locus_tag="C83_01613"
/note="DUF4315 family protein"
/product="DUF4315 family protein"
/protein_id="gnl|SC|C83_01613"
/codon_start=1
CDS complement(36668..38905)
/color=8
/locus_tag="C83_01614"
/eC_number=3.4.-.-
/note="Pepidase M23  Probable endopepidase p60 precursor,
similar to AA sequence:UniProtKB:P21171 protein

motif:CLUSTERS:PRK13914 protein motif:Cdd:COG3584 protein
motif:Pfam:PF00877.13"
/product="Pepidase M23  Probable endopepidase p60
precursor [pepidase]"
/"ID=C83_01614
/protein_id="gnl|SC|C83_01614"""
/codon_start=1
CDS complement(38905..39861)
/color=8
/locus_tag="C83_01615"
/eC_number=2.1.1.113
/note="Site-specific adenine/cytosine
DNA-methyltransferase, similar to AA
sequence:UniProtKB:P11409 protein motif:CLUSTERS:PRK11524
protein motif:Cdd:COG2189 protein motif:Pfam:PF01555.12"
/product="Site-specific %28adenine/cytosine%29
DNA-methyltransferase [methyltransferase]"
/"ID=C83_01615
/protein_id="gnl|SC|C83_01615"""
/codon_start=1
CDS complement(39886..40506)
/ID=C83_01616
/color=8
/locus_tag="C83_01616"
/note="Nitrogen fixation protein"
/product="Nitrogen fixation protein"
/protein_id="gnl|SC|C83_01616"
/codon_start=1
CDS complement(40527..42956)
/color=4
/locus_tag="C83_01617"
/note="Type IV secretory system protein TraC  VirB4,
protein motif:Cdd:COG3451 protein motif:TIGRFAMs:TIGR02746
protein motif:Pfam:PF12846.1"
/product="Type IV secretory system protein TraC  VirB4
components  AAA domain [ATPase  mating pair formation]"
/"ID=C83_01617
/protein_id="gnl|SC|C83_01617"""
/gene="traC"
/codon_start=1
CDS complement(42922..43305)
/color=4
/locus_tag="C83_01618"
/note="PrgI family protein  Type IV secretion system,

protein motif:Pfam:PF12666.1"
/product="PrgI family protein  Type IV secretion system"
/"ID=C83_01618
/protein_id="gnl|SC|C83_01618"""
/codon_start=1
CDS complement(43274..43864)
/color=8
/locus_tag="C83_01619"
/note="Adenine-specific DNA methyltransferase , protein motif:Cdd:COG4725 protein motif:Pfam:PF05063.8"
/product="Adenine-specific DNA methyltransferase [methyltransferase]"
/"ID=C83_01619
/protein_id="gnl|SC|C83_01619"""
/codon_start=1
CDS complement(43878..44747)
/ID=C83_01620
/color=1
/locus_tag="C83_01620"
/note="hypothetical protein"
/product="hypothetical protein"
/protein_id="gnl|SC|C83_01620"
/codon_start=1
CDS complement(44785..44997)
/color=4
/locus_tag="C83_01621"
/note="Conjugal transfer protein, protein motif:Pfam:PF12750.1"
/product="Conjugal transfer protein"
/"ID=C83_01621
/protein_id="gnl|SC|C83_01621"""
/codon_start=1
CDS complement(45085..45297)
/color=4
/locus_tag="C83_01622"
/note="Conjugal transfer protein, protein motif:Pfam:PF12750.1"
/product="Conjugal transfer protein"
/"ID=C83_01622
/protein_id="gnl|SC|C83_01622"""
/codon_start=1
CDS complement(45420..47207)
/color=4
/locus_tag="C83_01623"

/note="type IV secretion system protein TraG  VirD4 components, protein motif:CLUSTERS:PRK13876 protein motif:Cdd:COG3505 protein motif:TIGRFAMs:TIGR02759 protein motif:Pfam:PF02534.8"
/product="type IV secretion system protein TraG  VirD4 components [coupling protein]"
/"ID=C83_01623
/protein_id="gnl|SC|C83_01623"""
/gene="traG"
/codon_start=1
CDS complement(47204..47695)
/color=4
/locus_tag="C83_01624"
/note="PcfB family protein  conjugal transfer protein DUF3801 domain-containing protein, protein motif:Pfam:PF12687.1"
/product="PcfB family protein  conjugal transfer protein DUF3801 domain-containing protein"
/"ID=C83_01624
/protein_id="gnl|SC|C83_01624"""
/codon_start=1
CDS complement(47796..48632)
/ID=C83_01625
/color=7
/locus_tag="C83_01625"
/note="helix-turn-helix domain-containing protein"
/product="helix-turn-helix domain-containing protein [transcriptional regulator]"
/protein_id="gnl|SC|C83_01625"
/codon_start=1
CDS complement(48637..49413)
/color=1
/locus_tag="C83_01626"
/note="Toxin-antitoxin system BRO family toxin Bro  phage antirepressor KilAC domain-containing protein, protein motif:Cdd:COG3645 protein motif:Pfam:PF02498.11"
/product="Toxin-antitoxin system BRO family toxin Bro phage antirepressor KilAC domain-containing protein"
/"ID=C83_01626
/protein_id="gnl|SC|C83_01626"""
/codon_start=1
CDS complement(49397..49645)
/ID=C83_01627
/color=1

/locus_tag="C83_01627"
/note="hypothetical protein"
/product="hypothetical protein"
/protein_id="gnl|SC|C83_01627"
/codon_start=1
CDS complement(49860..50480)
/color=7
/locus_tag="C83_01628"
/note="helix-turn-helix domain-containing protein, protein motif:Pfam:PF12728.1"
/product="helix-turn-helix domain-containing protein [transcriptional regulator]"
/"ID=C83_01628
/protein_id="gnl|SC|C83_01628""""
/codon_start=1
CDS complement(50882..51034)
/ID=C83_01629
/color=1
/locus_tag="C83_01629"
/note="hypothetical protein:Prodigal:2.60"
/product="hypothetical protein"
/protein_id="gnl|SC|C83_01629"
/codon_start=1
BASE COUNT 12860 a 12295 c 12769 g 13110 t
ORIGIN
1 ttagttcgga gcaagtccca aaccattgag catcgcgtca gccgaagtca gcttcgagct
61 ataatccaga tgtgcgtaaa tgtttgccgt ggttgagaag tcgctgtgac caagccattc
121 ctgaatctgc ttcatgggaa caccgtttgc cagaagtaac gaggcacagc tatgtctcaa
181 gtcgtgatac ctgatgtgtc tcagcccatt tgccttgagc agcttcggaa aactctcagt
241 aacatagtgg ggcttgatca gatcaccaat ctcgtttacg caaacataat cgatgtagtc
301 cttgatgtag ctgcgtccgc acaagcgacg gttttcctgc tgttcttctt ttagggcgag
361 aagtctttct ctcatgaaag gaacaagcgg gagcgttctc atactcgatt tagtcttggt
421 ggtgtcggat gcgaccagaa cgcgcttgcc gtccaggtca caagaagtaa ctgtgtgacg
481 gatggtgatc gtattctgct caaagtcaat agcatcccat ttcagaccga ttgcttcact
541 ccggcgcaga ccgtagaatg cgccgaacag aatcggaagc tcaagtttcg ttcctttagc
601 tgcttcgaag agagcgttga cctcatcggc atcataaaag ctgccgacat agcgatcctt
661 cttcggacgt tcaaccttat ctgccgggtt tacatcaatc aggtcaatct tcacggcata
721 cttgagagcc ttatgaatgt tggcatgata atggatgacc gaggaagcac tgacgcgctg
781 gagctccttc atatagaact cctgaatatc ctttgcggtc agattcttga gagtgatctc
841 cttttcacgg aagtacggtg cgacgaccct tttgaccgtg gtcgagtaag aagcaaatgt
901 cgggactgca accgagcttt tgataatgtc gagccattgt tccatgtagt ccgcgaacag
961 gatgtccccg ttcatgacct tcggctcttc gggattgaaa ttgcgtcgcg catccatcag
1021 gattgcttcg gcgcgttttt tgtttccctt aacagtaagt ccggtagaaa tccacttggt
1081 ctttcttttt ccgagagaat cggtgtagtt caacacggcg taataatatc cttttttttc

1141 gcggagatgt cctgctacca tatattatcc tccttcttgc gtagcagtaa cacctacctt
1201 ataccgggat cctattatac cagaaccggg ttcgaattgg aggacctact gacgcactat
1261 tttgtaaact ttttgagtcc aaatcttacg cgttcactgt ttctccgcag ccgatgcaga
1321 gataagtgaa aagatgagcc ttcggaatac gataggctct gccgaccttg aggctaagga
1381 ttttgccttc gcgcaaaata cgatagccgg tctttgtgct gatgctgagt atttcacaca
1441 tctgctcaat gctcatcaca tcgggatagt ctttcagcat gagcttatag gcatcgcgct
1501 gagaaatggt tgatgcggta ttcacctgtc tcacctccgt ttatttgttt ttgctttatc
1561 agcaaaagga taggcagcag tcggttgaca actgctgccc atagtttttg atgataaatt
1621 gtgttccagc atatttgcag ctcgcacccc tgctgaatag ggaatgctgc ggactaccac
1681 cggtaaagca gtatcatggg actctcaccc ctccgaggat cgctccgagc cgccccttca
1741 aagaagaaac ggaagtatca ttgtacccaa cttctccatc atcgcaagca gccgcaccac
1801 acggctgttt agtctctctg ttgatcgctc actcctgtat gggaggtcat ggcggcagaa
1861 gacaagtcgc ttcgagaaaa gaggaaagat ccgcagcact gactattcag ttttcaagga
1921 accgtgcagc ggcgattaag cctcttcaaa atacaacgga catttttcat cagtttttgg
1981 gggtctcctt aaaaaatatt tttgcttttt tctgaacatc tctcttggct tcaaaaacag
2041 tagagacatc aattccacga gccttgctgt aatccgtcat gctcatgccg ccgagcaagc
2101 agtagaggta aacttcgagc tggcgttcgc tcaagagcaa gcggaactcc tgctccataa
2161 cggaagtgag aactgtttct tccaaatcgt tcgggtcttc cagccaagca ggactcatct
2221 cttctgcatc ctgcacactg acaaaatcaa gcgaaagaag agttgccgtc tcatcctcat
2281 cctcgcaacc ggcaataggt actccggtct tagagcgtct gagcttcttt tcctcgttgc
2341 ggagaactct catggtttct ctgctgacct cgcatacttc gccggtgagc ttcacgcgca
2401 ccatgcactt gccttcctcc gtagtccaga gatcgtagtc aaattcgata ggggttttca
2461 ttgtcttgtc ctttccgctg gcgcggagca gcggaaggac gagacaaaat aagagccgca
2521 tgacggtgag tttgaatccc atgccgataa aacagagtcc gaaaactctg ttcatgcggc
2581 attaggaaga ctcaccaatc agcggctcca cagcacagca atcatattgt tttgttgtat
2641 gttattatct ccggcctttc agacggagtt ttaaccttct catgctgagg acatcaaaga
2701 ctgtctcacg cccacaagcc ttgcattttg cttggacatg gccttttgtg tcctcaaata
2761 cggcgatgga attatgcccg caataggggc atttcatcat gcgcatcttc tgatttgtta
2821 tggcgcgtcg cgccctttgt atctttacct gcatttccgg cgagggctcg gaaacccgga
2881 tgcttctgct catggccaca cctccaacgg atcgtcatac tccgcaaagg ggcggtcctc
2941 gatgtagccg agttgcctca gccgaataat ggctgcggac ttggaaactc cgagagactg
3001 gcacagcaga gttagggata aacaatcaat ataattaaac cttccctcgt agttcctgag
3061 gacacggcca cgggcaaatt gcctcatggc aatgtcgatt tccttttggg gcatcagaat
3121 ggcagcgcct aacacattag cttgccattc attccaatcc tcgcgggtct tgagatcacg
3181 gagcgaatat gccgctctgg gggcatattt gctttcgtgc gttctcttat cattatcgct
3241 ttccatcccg aacaggattt gatgagcgca ttcatgggca agggtaaatc tgcgcttccc
3301 gcacaaagca tgaacctgtc caggctgaat aaagctcaaa tccaaaagaa cctgatttgc
3361 tttcagcggg attgtccgct taatgccgtg ttcctctatg gtgtactctg tatcggcata
3421 agcagtcagc ccacataggc ttccgtcaga ggatagatgc gcaaatgaaa cgctcaagcc
3481 gagataatcc ttggcaaact ggtcaatcgg tgtcgcctga gcagttgacc tctcattgtc
3541 atcgtctttt ccgaaaaaga actcattgaa gtccttggtc acggctgccg ctatttcttc
3601 gatctgctta tgggatagaa tcacgagcag ccctcctttg cttctacgaa ccatttgtca
3661 ccttcatgaa acagaaacga ctcctttcct ctgatcaata cggtgtagcg aataccgccg
3721 cctcccactt ttttagaggt tgcgcggcac ttatacaaga ctttgtcgat ctggaagatg

3781 agtccgtcct tccagcggat gaagcatgga tgaatagtgc cttcctcatc cacatcaaca
3841 ttaaccgaaa catacgcttt tctgcactga gcgttattca taaccagcct ccgttcataa
3901 aaaagttact taaaagcacg aacattattt catgctcctg ttgacacaca ccctttcgtg
3961 tggtatcata tagacgaaca tagtttcgtg ctgtgtgatc attataccgc gaacagtgtt
4021 tcgcgtcaat aggttttaac gaaaaagtgt tcgtgttcat gaaaaattaa agaaagaggt
4081 gccttatggg tttcaaagag cgactgaaag aaaaaaggat tgaagccggt ctgactcagg
4141 tagaactcgc taagaaagcc ggagttacag cgcgtacaat tcagaattat gaacttggaa
4201 gccgcaagcc cgtccacatg gaggttgttc agaaaattgc agatgctctc ggaactacca
4261 cagaatactt gctgggcagc agcggagctt atgtcgctga tgcttatgaa aagggcggtg
4321 cgaaagcagc cagagatatt gacgagttgg taggtgaagt gaccggcctt ttcgccggtg
4381 gcagtttgag tgaagaggct ttggaaggtg caatgaaagc actcaatgat gcctattgga
4441 tcgccaaagc gaaaaataag aagtacgctc ccaagaagta tcggaagagc gatgagccag
4501 cggaataagt ccgttctgtt ggacacctct tctgctatac ttttatttcc acgaatggag
4561 gtggaacgat gactgctgca tacctctcga aagtaggcaa cgaccttgta aatcgttgcg
4621 gaacacgaga tccttttcgc attgcacatg agctgggcat tgaagtcctt ttctgcgaag
4681 actttggtcc cctcaaggga atgtatagag taatcaaacg caatcgcttt attttcatca
4741 ataaggattt gagtcctcgt atgcagacga ttgtctgcgc acacgaaatc gggcatgatc
4801 agttgcaccg gagcatggca aagaatgacg ctattcaaga gttcatgctg tacgacatgg
4861 caacaaagcc ggagtatgag gcgaacatcg tagccgctga aatcttgctt gatactgata
4921 ccatcctcga atacatctat gagtacggct atacctcgga gcagattgcg cgagctatgt
4981 atacagacat aaacttagtc gctttgaagg tggcgcatct ggctgagacc ggatatgacc
5041 ttcgccggat tgaccatcgg agcgacttcc tgaaataagg aggatgctat gaccaaaacc
5101 gaagtcatga aggactttat gaaaaaaaca gtgctgccgg ttgtcgtggc actgctgctg
5161 ttctcaattt tcagccggat tttcattgag aatggcacac cgaattattt tctaatctgg
5221 cttgcttgcg gagtgccgtt tggcatcggt aagatgttca cactgattcc ggttggtttt
5281 ggcatctccg gtacagttgg cgttgttgcg ctcaacatcg ttcttggagg attgatcggc
5341 ggagtgatcc tgatttacag acttgcagtg gcagtttggt atctgcccct gacgatatac
5401 aggcttattg cctgagaaac gaggaaaaca atgagcagaa aacttgtggc atacttttcc
5461 gcatccggca cgacagccaa agtagccgaa acattggctg aggccatcgg tgcggatata
5521 tacgaaatcg aaccggcggc accttacagc aaggctgatc ttgactggac aaatcagaaa
5581 tcccgaagca gcattgagat gagcgatcca gcataccgtc ctgctatcgc cggtaagcgt
5641 gacaacatgg atgactatga caccgtattt gtcgggtttc cgatttggtg gtatgtggct
5701 ccccatatca tcaacacctt ccttgaaagc tatgacctga ccggcaaagc cattattccc
5761 tttgctacct ccggcggaag cggcatggga gaaaccaatg aaaaactcca accgagctgc
5821 ccaaactcca aactgataga gggaaaggtg ttcaaggcgc actcaagtaa agctgagatt
5881 accgcatggg cagacgctct tgagtactga tttgaggtaa ggcgaatgga ttttgaaaac
5941 aaagtcgtag ttataacagg cggcgctcat ggtatcggca aagcgattgc cgagtcgttc
6001 cgcagtgggg gcgcatcagt tgaaattatt gacatcgcac cgggagatca ctacatcgga
6061 gacatcagca gaaaagaaac attagaggcg tttgcttcct atgttctgga taagcatggt
6121 catgtggact acctgatcaa caatgctctt ccgctaatga agggaatttc cgaatgctcc
6181 tatgaagagt ttcagtatgc cttgtctgtc ggtgtcacag ctcccttcta tctcagcaag
6241 ctcttttctg agcattttgg caagggcggc gttatcatca atatttcttc atccagagat
6301 cggatgagtc agccgcagac cgaaagctat acagcggcca agggtggtat tgcggcgctg
6361 acccatgctc ttgcagtcag cttctctgga aaacttcgtg tcaattccat ttctcccggc

6421 tggattgata cggaataccg cgaatacgag ggagcggatg ccgttcagca accgtcagga
6481 cgggttggaa acccgatgga tattgcaaat atggtcttat tcctctgctc tgaaaaggcc
6541 gggtttatca ccggagagaa tatctgcatt gacggtggta tgacgaagca gatgatctat
6601 catggagatc atggttggaa gctggaaata taggaggaac aaagtatgcc tcttcatatc
6661 gtcagaaatg acatcacaaa aatgaaagtg gacgcaatcg tcaacgctgc caactcatcg
6721 cttctgggcg gtggtggtgt tgacggctgc atccatcgtg ctgcaggacc ggagctgttg
6781 gctgaatgta aaatgctggg cggatgtgaa accggcagcg ccaagatcac gaagggctat
6841 aagctgccct gtaagtatgt gatccatgct gtaggtcccc gctggcggga tggcaagcgt
6901 ggtgagcgag ataagctcat ctcctgttat cagacttctc ttgctctggc aagggaaaac
6961 aaatgcgaga cagttgcatt cccgttgatc tcttccggta tttttggcta tccaaaagac
7021 caggcactca gaattgccat tgatacgatc agcgagttcc tgcttgaaaa cgatatgacc
7081 gtctacatcg tcatctttga caggaaggca tatcagatca gcgggaagct ctatacggat
7141 attgccgagt acatagacga taactatgta gacgagcatt ctgacaacta ttctgcgcgt
7201 ctgcgccggt tgaatgcact tcgggatatg gagccggttt gcgaagcgtc ggtctgtgaa
7261 gaagccgacg aagcgattga gcctatcctg tctatgccaa tggcagcgcc taagaagaca
7321 aaatctctgg atgaggcgct gggtcagatt gatgagagct tttctgagat gctgctgcgc
7381 aagatcgacg agaaaggtat gacggacgca cagtgctata agaaagccaa cattgaccgt
7441 aagctcttct cgaagatccg gtcggacaag ctctataagc cgagcaagcc tacggctatt
7501 gcctttgcta ttgcgcttga gctgtctttg gatgagacga aggatatgct tatgaaggcg
7561 ggctttgcgc tgtctcattc caataagttt gacatcatta ttgagtattt catagagaat
7621 gaaaactaca atgtctttga aataaacgag gctctgttcg cgttcgatca gagtttgctc
7681 ggagcataag agaaagacaa atcggaattt gtagaggtga attttttgga cagagcaata
7741 ttaaaagaac ttggcttcag aagcaggaat gaggctctgg cttttctgaa aaaaatatgg
7801 agacaggaga aggcggaatg tccaatatgc ggcagtgaac tggaactgct gcataaaaag
7861 gcgaaaaaga ataattgtga ttggcagtgt aagaattgta ataaaactta taaaaccatt
7921 cacttactgg atgaaattaa tgagcagatg ccagattaac aacttccagt ttggagaggc
7981 aacttttcac cgaaagagga tgaatacaaa aagttcaagg aggaaatctc tgtcatgaag
8041 ttaattcgta tatttcacca aagccaatca tctcaacaaa aaaatggaac ttttcagggt
8101 atcagttcct ttacactaca tatcatcgct atgcttctta tgctgtgcga ccatctttgg
8161 gcgactgttt tttcccgata cgcttggctt acctgggtgg gcaggcttgc ttaccccatc
8221 tttgccttta tgctggtgga gggctatttc cacacccgca atttgaaaaa gtatatggga
8281 cggctgctct tttgggcaat cctctccgaa ctgcccttca atctgatgta cggaggcagc
8341 ctgatctatc ccttccatca aaatgtactt tggacattcc tgttgggact tttgtttatt
8401 accggcattg aaaaaacgcg ccataaagga aagcctctgc tgacgatgat cgtctttttg
8461 ctttcagcag ccgcaggcta tgtgctgggt ttccttgcta tggtagatta ttttggatat
8521 ggaatatgga tggtactgct tttctacctg ttccgtggta gaaaatggtg gtgccttctg
8581 ggacaatttg ttggactgtg ctggattaac gcggtgttaa tcggaggtct ctccgttccg
8641 gtacaaatct ttgggcacga agtatttatt gtgcaacagt caatggcatg tctggccctc
8701 atccccatct ggctgtacaa ggggaaacag ggaccgcata acaaaatcat ccaaaccgct
8761 ttttacgcct tttatccagt gcatatgctg attctttcac tggcagcact tctgtagcgc
8821 cagcttgctc cggaccactt ctcaaaaaac tcttaccgct catcaaatta cttataaaaa
8881 ttcctgtttg tcaaactgat atggtgatat gaaaggaggc caccggagca tgaaagagcg
8941 cacatacatc gcaatcgacc tcaagagctt ttatgcttcg gtggaatgcc gggagcgcgg
9001 cttggatccg ctggacacaa accttgttgt cgcagacgaa agccgtaccg acaagaccat

9061 ctgcctcgca gttacaccct ctctcaagtc ctacggcatt tcaggccgtg gccgtttgtt
9121 tgaggttaag cagcgagtta aagaggccaa cgccggacgg cagcatgacg ctccgggaca
9181 caagctggaa ggttcctcct gcttccattc tgaacttctt aaaaacccaa acctcgcgat
9241 tgacttcgtt atcgccccgc cacgcatggc ctattacatg gaatacagca cccggattta
9301 caatgtctac ttgaagtatg ttgcatcgga ggacatcatc gtatattcca tcgacgaagt
9361 gttcatggat gtcactgact atctgaacac ctataagctg tctccccacg acctcgcaat
9421 gaagattatt ctggatgttc ttgagacaac cggcattaca gcaaccgcag ggatcggcac
9481 aaacctcttt ctctgtaaag tggctatgga tattgtggca aaacacatcc cggcagataa
9541 gaacggggta cgcatcgcag agctggacga aatgggattc cggcgggagc tttggtcaca
9601 tcagcccctc actgacttct ggcgggtggg gcgcggctat gctaagaagc tggaacagaa
9661 cggaatgttc acgatgggcg atgtggcacg atgctctgtt caaaatgaag acattctgta
9721 ccagctcttt ggaaagaatg cggagctgct gattgaccat gcgtggggct gggagccttg
9781 tacggttgaa gcaattaagg catacaggcc gagtacaaac agcctcggct caggtcaagt
9841 gctgcatcag ccatatactg ctgagaaagc caagctcgtg cttcgggaaa tggccgattc
9901 gcttgtgatg gacttggtgg aaaaaggctt ggtcacagac cagctcgttg tcaccatcgg
9961 ctacgacatc gaaaacctta ccgatcccga gcgccgaaag aaatatcgcg gcgatattgt
10021 gaaggatcac tacgggcggc agataccgaa acacgctcac ggcactgcca accttgacgg
10081 tcacacatca tctacgaaga aaataatgtg tgctgcgtca gaactctttg accggatcgt
10141 cgataagaac cttctcgtgc gacggttgaa catcgtcgca aaccatgtgc tacccgaagc
10201 agatgcaccg aggaaaaacg ccggttttga acaacttgac ctcttcacgg actatgcagc
10261 tcttgaggta aagcaggagc aggaacgcgc cgagcttgag cgtgaaaaga aaatgcagca
10321 agccatgttg accatcaaga aaaagttcgg aaagaacgct atcctcaagg gcatgaactt
10381 ggaggaaggc gcgacggcta aggatcgcaa cgcgcagatc ggaggacaca aagcatgagt
10441 gaaaactacg atgatataat cgatcttcct catcatgttt ctgcaacccg gccgcagatg
10501 tccatgatgg atcgtgcagc acagttttct ccattcgccg ctctcaccgg ttatgatgcc
10561 gccatcaagg aaaccgcgag gctgaccgat cagaaaattg aactcgacga ctacgagaag
10621 gaagaaatca acggcaaaat ccagctcatt gcagagcatc ttggcgagga ttttgaagtt
10681 gtgattacct actttcagct ggacggcaag aaggccggag gcgcgtatgt ggatgcgctg
10741 ggtgtcgtaa agaaaatcga cgaatacgag aggatgatcg ttttccggga aggcagaaaa
10801 gtctcgattg atgacattct tgaaatcaag ctggctgagt aattgtcgct tgccatgcga
10861 ccaaaccgaa tatgattttt cttataatat gactgtaagg taaacaaccg agcagtcata
10921 ttttgttttt ggctgaaagc caaaaacgat atacaggagg tactgaacat gagaaagaac
10981 ctaactgaga tcgtattcat tcttgaccga agcggctcta tgagtgggct tgaggcagat
11041 actatcgggg gcttcaactc catgattgag aagcagaaaa aggctgaggg cgaagctctc
11101 atctccaccg ttctcttcga caacgcaagc gaggttattc atgaccgtgt gggtgttcag
11161 aacatcaagc cgatgaccga tgaggactat acagtacgag gctgcaccgc ccttctggat
11221 gctgtcggcg gagcaatcca ccacatcgga aatgtccata agtacgcacg agctgaggat
11281 gtaccggagc atacgatgtt cgtcatcacg actgacggaa tggagaatgc aagccgccgg
11341 tatgacagcg ataaggtcaa gaaaatgatc gagcgtcaga aggaaaagta cggctgggag
11401 tttctgttcc tcggcgccaa cattgacgca gttgagacgg cacgacactt cggtatcagt
11461 gaggaccgcg ctgtgaacta tcattctgac agcgaaggca cacagctcaa ctacgaggtt
11521 gtcagcgagg ctatctgtgc tgtccggtgc aacgcacctc tcggtgccga ttggaaaaag
11581 cgcatcgacg aagactacaa cgcaagaaag gatgggcgaa gaaaatgatg aggacaatta
11641 ctgttcgagg cgttggaaat gtatcagcaa gaccggacta cataaccatc tcaatgacca

11701 tagagtctat ccgaaaagac tatgacgggg caatgcgaga tgcctctgag cgtatatcga
11761 aattgcagga tgtggctgtt cgttcggggt atgcaaagga ggacttgaag accaccagct
11821 tcaatgtaga tacccgatat gagagcgtca aggatcggca aggcaactac aagagagagt
11881 tcgccggata tgcctgtgta tatcgtctca agctgtcttt cgactttgac agcaagcagc
11941 ttgcgaaagt gatttccgct atcgccaaca gtggagcaaa tcccgaattg agcatcgcgt
12001 tcacggtgag aaatccagca aaggtcagtg cagaacttct ggaaagtgct gcgctcaatg
12061 cacgagcaaa ggctgagatt ctatgcagag cgtcgggcag cgagatggga caacttctct
12121 ctatcgacta taactggggt gaactgaaca tcgtctcgcg cacaagctac gaaatggaag
12181 actgcctcat gcctcttgca gcaatggaaa aatgctgttc acctgagatt gagcctgatg
12241 acattgatgt atcagacacg gttgccttca catgggagat taagtaatta ccatatcgtc
12301 ggagaaagac gctttcgggc gtcttttttc gtgtccaacc cccaaaaacc attggctttt
12361 gtccgttgta ttacgaaggg gtattttacg gggagtccct acgggggaat acgattttag
12421 agaatagcaa gcaaatccgg tgagtaccca ccggagaaaa aagcgttaag ggaaccctcc
12481 ccttaaccgc tgatttttgc ttgttgggat agtcccaaac ccttataaat tacgaagaac
12541 ggagacagaa cgatggcaaa cagaacgcgg aaagtgcagc tcaagttctg ggtcaccgag
12601 gaagaacggg cgatcataga gcggaaaatg gagatgctga aaacgaagaa tatgggagcc
12661 tatctgcgca agatggcaat agatggctat atcattcagg tagactatac cgagcagaaa
12721 aaactcgctg cggcagtcaa taagatcgga gcaaacatta acaatatctg ccgaaggatg
12781 aactcaacgg gacgattcta tcaggacgat gttgttgagc tgaaaggaaa gctggatgaa
12841 gtatggcagt tactaagatc aaagcaatcc gaggaacttt gaacaaggct cttgcgtaca
12901 tcatgaatcc tgagaagacc gatgataaaa tgcttgtctc ttcctatgga tgtgccgctg
12961 agacagcggc tcaggaattt gaatggacgc gaaaaattgc cgagcagcaa gggatgaatc
13021 ccgtcaaaat tattgcccgc catgtcatcc agtcctttga tgttggcgag gtctcgccgg
13081 agcttgccca tgagatcggc aagcagtttg cggatgaaat cctcgccggg aagtatgagt
13141 atgtgctgac cacccatatt gataaagacc atgttcataa tcacctgatt ttcaatgccg
13201 tggactttgt ggatcatcac gcatacaaga gttacaagaa aatctactat gatatgcgcg
13261 aggcaagtga ccggctctgc aaagaaaacg gtctctccgt aattcctccc tctcagggca
13321 agggcatgag ctataaagag tacaccgagg ccaagcgagg aacaagctgg aaacaaaagc
13381 tcaagcagac cattgaccga aagatcatca cggcaaagga ctacgatgag ttcctgaaac
13441 tgatgcagga tgccggatat gaaattaaaa ccggcaaata catctccttc cgagccgaag
13501 gccaggagcg tttcacacga gcaaaaacca tcggagagaa ctacaccgaa gaacgcatca
13561 aggaacgcat tcagggacac aatccccgaa agcgtcagat gcagactacc cgcaaaggtg
13621 tctcgcttat cattgatatt cagaacagca tcaaggctca ggaaagcaag ggatatgaac
13681 actgggcgaa gatcaacaat ctcaaagaag ctgcgaaaac gctcaactat cttactgaaa
13741 acaatctcct gcaatatgcc gatcttgaga aaaaggttga ggacattcac agctcctatg
13801 cccgtaccgg tgaagagctg aaaagggttg aagccaggct tcgggaggta cagcctctca
13861 tcaaaaacat cgcaaattat cgcaagctca agcctgtcta tgatgcctat atgaaagcaa
13921 aagataagtc tgctttccga gctaaacatg aagctgaact tgtaattttc gaggctgcga
13981 agactacgct acttgcaatg caaggcgacc aaaaactgcc aagcctcaaa gctttccaag
14041 ccgaacagca aaggcttact gatgagcaac aaaggctata tgatgagcgg gcaaggttga
14101 aaagagaaat gaagcagata gataccatta aggccaatgt ggatatgctg ctccgaattg
14161 atgaagatca ggagcaaaga caggagcgtt ctcctcagcg cgagtaaccc aagcgggtct
14221 gactgttttg atgtcaggcc cgcttttata tgcgtagatt gtggataaaa ggggcagtcc
14281 aagaagttgg aactttttg ttaatcctta gagcatccga ttgaaaagat cgactatttc

14341 tgctataata taaactgatt ccttatatca gcttacagat gtaaatgcaa gaggtgctta
14401 tatgaagatc agttacaaaa aactttggat gcttctcatc cagcgagaca ttacaaaggt
14461 caggatgaaa agagatttag gtcttgctgc cggaacaatg agtaagctca acaagggcga
14521 ggatgttgcc ctctctgtcc tgctgcgcat ttgtgagtat ctggattgtg acatcgggga
14581 tatttgcgaa gctgtgcgag cgggaaactg agtcccgttt ttcgtacagc ctttatgcta
14641 taatttggga ttggagacgg ttgtgtctca atgaggtgat gaaatgcaga ttattgataa
14701 cataaacaag accgtcaagg acgatctgca agtgggtatt cataaaggca gtaagatgtc
14761 cgttgccgcc gcttgctttt caatctatgc gtatcaggaa ctcaaaaagc agcttgaggg
14821 cattgacgag ctacgtttca ttttcacctc gccgacattt gtaaccgaga aaacatcaaa
14881 ggcgcagcgg gagttctata ttcctcgcat ttcaagagaa agcagtctgt atggcactga
14941 gtttgaagta aaactccgca atgagctgac ccaaaaagcg atcgccaagg agtgtgcgga
15001 ctggatcaga cgtaaggcgg ttttcaaatc gaacgtcaca caggagcaaa tgatgggctt
15061 catgacggtc gatgaaagca cctatatgcc gatcaatgga ttcaccactg tcgatcttgg
15121 ctgtgagcgc ggcaacaacg cctattatcc agtgcagaag actgagagct ttgagaacgc
15181 caactacttt ctgggacttt ttgaacaact ctggaatgac aagaaaaagc tgcaagaagt
15241 cacggatatt gtgattgaga atatctccgc cgcgtatcgt gagaacgcac cggagtatat
15301 ctattttgtc gcgctctata acatcttcaa agagttcctt gatgatgttt cggaggacga
15361 gctgcccaat gaagcaacgg gctttaagga aagcaaaatc tgggggatgc tttacaactt
15421 ccagcgtgac gctgttctcg ccatcatcag taagcttgag aaattcaacg gctgtattct
15481 tgctgatagc gtcggtcttg gcaaaacttt tactgcgctc gcggtcatca agtattatga
15541 gaaccgcaac aaatccgttc ttgttctctg cccgaaaaag ttgtctgaga actggaatac
15601 atacaaaggc aactatatca acaatcccat tgcggcggac cggctgcgtt atgacgttct
15661 ctatcacacc gacctttccc gcgagcatgg gctgtcaaac ggaattgacc ttgaccgcct
15721 gaattggggc aactacgacc ttgttgtcat tgatgaaagt cacaacttcc gtaacggcgg
15781 tgaggtgtcc ggcgaggatg caaaagaaaa ccgctatctg aaactgctca acaaagtcgt
15841 tcgcgccggt gtaaaaacca aggttctgat gctttccgca acgccggtca ataacaagtt
15901 tattgacctt aaaaatcagc ttgcacttgc ctatgagggc gatgcatcgc agatgaacaa
15961 aaagctcgat acaacgaaga gcattgatga aattttccgt caggcacaga aggctttcaa
16021 cgcatggagt aagctccctg ccgatgaacg cacaacggac gagcttctgc ggactctgga
16081 ttttgatttc tttgaacttc tggacagtgt gacgattgcc cgttcgagaa aacacatcga
16141 aaagtattac aacacctccg acattgggaa atttccggag cgtctgccgc cgatttcact
16201 gcgtccctgc ctgaccgact tgaacgatgc gatcaactat aacgacattt acagtctgct
16261 catgtcactg agcttgacaa tctatacgcc ttcgaattac atcatgccgt cgaagctggc
16321 aaagtacatc gacatgacac accaccaagg aaacagcttg acccagcaag gacgtgagga
16381 aggtattcgc cgccttatga gcattaactt tacaaaatat catagaacac tgaccacata
16441 tctaaataca ctgctttcaa atagttttat aataaatcag attgtggagc cacagccgcc
16501 agagaacatg atggatattc cggggatggc ggatgaaatg cgacgcccaa tgatgctgat
16561 tgtatcggca aaaaagaaga tgtaataata tagaaaaaat aaacgaggag tatgtaaatg
16621 agatcagaaa aagaaatgat ggatttagta ctttctttag cagaacagga tgaacgtatt
16681 cgaattgtga cccttgaggg gtcacgcgca aatattaata tacctaaaga tgaatttcag
16741 gattatgata ttacatattt tgtaagtgat atagaaccgt ttatatctaa tgatgactgg
16801 cttaatcaat ttgggaatat aataatgatg caaaagccgg aggatatgga attattccca
16861 cctgaagaaa agggattttc ctatcttatg ctatttgatg attacaataa aattgatctt
16921 accttattgc ccttggaaga gttagataat tacctaaagg gcgataaatt aataaaggtt

16981 ctaattgata aagattgtag aattaaaagg gacatagttc cgactgatat agattatcat
17041 gtaagaaagc caagcgcaag ggagtatgat gattgctgca atgaattttg gaatgtaaca
17101 ccttatgtta ttaaaggatt gtgccgtaag gaaattttat ttgctattga tcattttaat
17161 cagattgttc gccatgagct gctgagaatg atatcatgga aggtcggcat cgaaacaggc
17221 tttaaattaa gtgtaggcaa gaactataag tttattgaaa ggtatatatc cgaggatttg
17281 tgggagaaac ttttgtccac ctaccggatg gattcctatg aaaacatatg ggaagcatta
17341 tttctatgcc atcaattgtt cagggcggta tccggtgagg tggcggaaag gcttcattat
17401 gcctatccgg agtatgatag gaatataaca aaatatacca gggacatgta taaaaaatac
17461 actggtaaaa ccggctgcct ggatagcaca tatgccgctg atatagaaga gaggcgggaa
17521 cagtgattac agaaatgaaa gcagggcacc tgaaagatat cgataaaccc agcgaaccat
17581 ttgaggtgat aggtaagatt ataccgaggt atgaaaacga gaattggacc tttacagaat
17641 tactctatga agcgccatat ttaaaaagct accaagacga agaggatgaa gaggatgagg
17701 aggcagattg ccttgaatat attgacaata ctgataagat aatatatctt tactaccaag
17761 acgataaatg cgtcggaaaa gttaaactgc gaaaaaattg gaaccggtac gcttatatag
17821 aagatatcgc cgtatgtaag gatttcaggg ggcaaggcat aggcagcgcg cttatcaata
17881 tatctataga atgggcaaag cataaaaact tgcatggact aatgcttgaa acccaggaca
17941 ataaccttat agcttgtaaa ttctatcata attgtggttt caaaatcggc tccgtcgata
18001 ctatgttata cgccaacttt gaaaacaact ttgaaaaagc tgttttctgg tatttaaggt
18061 tttagaatgc aaggaacagt gaattggagt tcgtcttgtt ataattagct tcttggggta
18121 tctttaaata ctgtagaaaa gaggaaggaa ataataaatg gctaaaatga gaatatcacc
18181 ggaattgaaa aaactgatcg aaaaataccg ctgcgtaaaa gatacggaag gaatgtctcc
18241 tgctaaggta tataagctgg tgggagaaaa tgaaaaccta tatttaaaaa tgacggacag
18301 ccggtataaa gggaccacct atgatgtgga acgggaaaag gacatgatgc tatggctgga
18361 aggaaagctg cctgttccaa aggtcctgca ctttgaacgg catgatggct ggagcaatct
18421 gctcatgagt gaggccgatg gcgtcctttg ctcggaagag tatgaagatg aacaaagccc
18481 tgaaaagatt atcgagctgt atgcggagtg catcaggctc tttcactcca tcgacatatc
18541 ggattgtccc tatacgaata gcttagacag ccgcttagcc gaattggatt acttactgaa
18601 taacgatctg gccgatgtgg attgcgaaaa ctgggaagaa gacactccat ttaaagatcc
18661 gcgcgagctg tatgattttt taaagacgga aaagcccgaa gaggaacttg tcttttccca
18721 cggcgacctg ggagacagca acatctttgt gaaagatggc aaagtaagtg gctttattga
18781 tcttgggaga agcggcaggg cggacaagtg gtatgacatt gccttctgcg tccggtcgat
18841 cagggaggat atcggggaag aacagtatgt cgagctattt tttgacttac tggggatcaa
18901 gcctgattgg gagaaaataa aatattatat tttactggat gaattgtttt agtacctaga
18961 tttagatgtc taaaaagctt taactacaag ctttttagac atctaatctt ttctgaagta
19021 catccgcaac tgtccatact ctgatgtttt atatcttttc taaaagttcg ctagatagag
19081 ttctatatta gagatgtaaa gagttttggt gaagaggttt gtgagtatgg cttttataac
19141 cagggtgcaa tgagaagcac taacagcagc tagcttaaga acgcatagga tatacttttt
19201 atggaagtca cagaggagga ataaaatttg aacaggatta acagagtaac ctctattctt
19261 attcagctgc aatcaaaaaa aataattcct gccaaagaaa ttgcacagcg atttaatata
19321 agcttaagga cagtttacag agatatccgg acacttgaag aagccggaat accaattgga
19381 tctgaggccg gaaaaggata ttttcttgta gagggcttcc tacttccgcc ggtaatgttt
19441 acagcggcgg aagtgggtgc attgattaca gcagggaaat ttttaaattg ccatggagat
19501 gaatcattta taaaggattt tgattcagct atgtataaaa tcaaatctat tttgaagcat
19561 ggtgaaaaaa actatgcaca ggagttggag aacagtatta acgtgtacag cacatctgga

19621 cagaaaaata cattggcgga taacgttatt gcagcaattc agactgcaat ctgcaataaa
19681 agggtaatat caattcaata tcctgcatca ggagggcaag aaccggaaag cagaatgata
19741 gagcctatat cactggggtt ttatgaacag aactggtact taatcggttt tgcaggctaa
19801 gaaatgaata taggaatttt cgtgtggatc ggattaaaag catttgcata gaagaagagc
19861 tctgtcaaag ccatgatggc tctttggaac aaatattaaa acaaatgctt tcatataaaa
19921 aactatataa tgtaattctt agagctgaaa aaggggagac atacaattct atcaagaaca
19981 ggttgctcaa gcgtctggaa agctcggttg cctctttccg tctgacgata gaccgcatta
20041 aaaagttgat caacgatacg attgagacaa tcaaaaatta tcagaccgga gactgtgttc
20101 tgaatctgac cgagatttcc ggcacagaag attttgatta cgacgatcag aacactgatt
20161 tcttctccgt cggcaagaag gttaaaattg acctcactga tatggactat gtttcatgga
20221 tgcgcgagct ggaaaaggat gcggagaatc tggaactcct gtccctcatg attgccgaca
20281 tcacgccgga gcatgacacg aaactgcaaa cattgtttga tttaatccgg aagaaaatcg
20341 agcatcccat caatgagtat tccgcccgag attgagcacc tcgaccgggc aaattgagca
20401 ccttcaccgg cggaattgag cacggacgcc gcagccgttg agcagcacgg taaaatgtag
20461 atatgtaccc ttcagggtgc aaattctaca aggaggtcaa ccaatatgac caatcaccgt
20521 gagattctca ggttgcacag cctgggactc aacaaaaccg agattgcggc cagtttacac
20581 tgcgctcgca acaccgtggc cgcaacgctg cagcgtgcag acaactgcgg cttgcagtgg
20641 ccgctgccgg atgaaatgtc cgacaggcag ctggcagaaa ggctttttcc cagcgctccg
20701 ggcaaacctg cgtacaagat gccggattac gcctatgtgt acaaggaaat gcagaaaagc
20761 ggtgtgacgc tgaatctgct gtggctggaa tactgcgacc agtgcaaagc gtccggcgaa
20821 attccgtacc agtctaccca gttcaacaag tattatagcg actatctcaa caagaccaat
20881 gccacgatgc acctgaacca caagcccgga gagatcatgc aggtggactg ggcgggtgat
20941 accgcttccg tcatcgacac ggataccggc gaaatcatcc ctgcctatgt gttcgttgcg
21001 accctgcctt acagcggcta cagctatgtg gaggccttct tctccatgaa ccaggagtgc
21061 tggactgccg cccatgtcaa cgcctaccgg tatttcggcg gcgtcacgcg gatgattcaa
21121 tgtgacaacc tgaaaaccgg cgtggaaaag cacggaaaag atgaggtggt tctgaacaaa
21181 tcctaccagg agctggcgga gcattatgcc actgccattg ttcctgccag ggtcagagcg
21241 ccaaaggata aagctgctgt ggaaggaacc gtagggatca tctccacctt tattctggcg
21301 gcgcttcgta acagacaatt cctgtctctg ccggagttga atgaagcaat ttgggagcgt
21361 cttgaagcgt tcaacaacaa acccttccag aagaaaaacg gcagcagagc cagtctcttt
21421 gcggaagaaa agccgtttct gcgacccttg ccgccctatc cattcgagct tgccgcctgg
21481 aaaaaggcaa ccgttggccc caactaccat atctccgtgg agcggatgaa ctactcggtt
21541 ccctttgagt atatccgcca gaaggtggat gtgcgtttga ccagagccac cgtggaggta
21601 ttctacaacg gaagccggat ctgttcccat cccaggctgt acggcaaatt caaccagtac
21661 agcaccattc aggagcacat gccgcctgac caccagaaat atgtgcagtg gaacggggaa
21721 aggtttatcc gctgggccgg taagattggc agcagcacag aatgtgctgt ccgcgctatc
21781 ctcagcagct acaaggtgga gcagcagggg tataaatcct gtttgggact gctgaagctg
21841 gctgacaaat attcccccgc gcgcctggaa ggtgcctgca agcgggcatt ggagttcacc
21901 cccagacctt ccctgaaaaa tattcaggcc attcttgcct ccgggcagga caaactcccc
21961 acggagccgg agcccacgtc ttcctcccaa tacggtttca cccgtggcgc tgactattat
22021 gacaggggga agaaggaatg ctgacgcaga caacgattac aaagctgcgg gaaatgcgcc
22081 tgagcgtcat ggcgagtgcc ttgaaagacc agatggcaga cccacaattc cagaacatgg
22141 cctttgagga caggctgggg ctgttggtgg atgcggaatg gaatgcccgc aaaaataacc
22201 atctgcaaaa actgatccgc caggccgcct tctccgaccc cggcgcctgc ctggaaaatg

22261 tggaatacct tccagacaga aatctgaagc gggaggaatt gctgcggttt ggcacctgca
22321 attacattca ggagcaccac aacatcatcc tgctgggagc gacagggagc ggcaagacct
22381 atctggcatg tgcgctggga atggcggcag tcagacagtt cctgacagtc aagtacatcc
22441 gtctgccgga tctgctggtg gagtttcaca ttgccagagg cgacggaagc atccgaaagc
22501 tcctgaccca atacaagaag tactcgctct tgattattga cgaatggctt ctctacccgc
22561 tgaaggagac ggaagccaga gatcttctgg agattatgga ggccaggtac aagaggtctt
22621 ctacgatcct gtgttcccag tttgacattc cgggctgggc agagaagctc tccgatccga
22681 ttctggcaga tgcgatttgt gaccgcattg tccatgacgc ctacaccatt gtcattggcg
22741 gcaaggagtc catgcggaaa cggaaaggct tgcaggaaac ctgatgtaag agcctgcggc
22801 tcactgggct ctgcccggac ccgaggtttt gcgcattccg gttttccagg agagtgattg
22861 gtatggtgca acaaaaaaga gagcgatgct gacgacatca ctctcctgcc aaaccccata
22921 tacggcgctc atgtcgctcc tcagcgttgc actatcctcc gatatggcta aaagcaaatt
22981 tatgatacca agcacagcat agtctgtcaa tattcaatcc gagcggtggc accgctcaat
23041 ccgggcggtg ccattgctca atttcagcgg aacgactgct caatttcgcc ggtcgaggtg
23101 atctggcgag gcggcgtacg caccatcaat cctggcaata agaagatcat tattttcacg
23161 gcattctctg ataccgcaga ctatctcttt gcaaacgtca gcagatttgc gaaagagaat
23221 tttgggctgg aaacggcaga gatcaccggc acagttgacg gtaagactac cattccaaag
23281 ctccgggcag acctcaacac ggttctaacc tgtttctcac ccgtttccaa aggaaagtcg
23341 gtactcctgc ccggaagtac cgatgagctt gacattctga ttgcgaccga ctgtatttcc
23401 gaaggtcaga acttgcagga ctgcgactac ctcatcaact atgacattca ttggaaccct
23461 gtgcgcatta tccagcggtt cggacgaatt gaccgtatcg gaagcagaaa cgaccatatc
23521 cagctcgtaa acttctggcc ggatatggac ctcgacgagt atatcaacct gaaaagccgt
23581 gtggaaaccc gcatgaagat ttcggtcatg acctctaccg gcgacgatga cctgatcaac
23641 gccgaggaaa aaggtgatct tgaataccgc cgcgcacagc tcaagcggct gcaagaggaa
23701 gtcgtggaca ttgaggacat gacaagcggt atttccatca tggatttggg acttaacgag
23761 ttccggcttg atctgctcga atacatgaag cagcacgagg acattgaaac cgcgccgaaa
23821 ggactgcaca ccattgttgc acagacagag gattctccta ccggtgtgat tttcgttctc
23881 agaaatatca atgacagcgt gaacattgac aatcggaacc gcattcaccc cttctacatg
23941 gtttacatca gcgaagaggg cgaaattgtc tgcgattacc tcaatccgaa aaaactgctc
24001 gacaccatgc gtttgctctg tcgtggcaag agtgagccgt gcatggagct ttgcaagcgc
24061 ttcaatgacg agacggatga tggacgtaaa atgggtgaag tctctgagct gctgagtgag
24121 gcgatcaatt ctattatcga cgtgaaagaa gaaaacgata ttgacagtct gttcaagagc
24181 ggcggcacat ctgcgctgct cagtgcggtt tcaggactgg atgattttga gctgatttgc
24241 ttcttagtcg tgaaataagg agggcttgcg atgcttgggt ttcctgcatc tacggagttt
24301 ggcaaaagaa taccaaagca gaaattctat gagaatcttg aggtttcccc tgctttacgc
24361 cgtgtgtttg tcgatcagat ccggctggtg tactggcgaa ataagctggc agcgtctaca
24421 ctgaatattg ctgcgggcga aatagtgacc gagattgaag tgttcgaggt gcgtctgaat
24481 gatccgcagc ttgatgaagc ggtgctgaaa caaatcgaca aggaaatccc gtaccatatc
24541 ctctttattt tgacctgtga cggcaaggct caggcgtgga tcggctacaa ggaggctgcc
24601 gcatccggca gcaacgcctt taaggtgagc cgctattatc ataccgactg gatgccggaa
24661 gacgaactgc atctctgcct cgacggcttg aatatggatg ccatctatga aagcctcgtt
24721 cgccaaatcg ccggggagaa gctgcaaacg gattccggcg agagtttgaa agagtctgtc
24781 gagcgggatg aaaagaagaa gcagctcgaa aagcaaattt ctacgctgga aaacaagatg
24841 aaaaaagaga agcagcttaa ccggcggatg gaaatgaatg cgaagttaaa gcagctaagg

24901 aagctgttta gggagattgc aaaatgaggg ttaaactatc gaatcgaaag ctttggaaag
24961 agtattatgc ggtcctttct gcgataagtg taattgcatc gttcgttttc ctgtttctga
25021 aagtaccgga tggaaaagag cttttatgct gcattggctt tttgatctgt ctcataattt
25081 cattcctaat agtgtggtat cgagccaacc acatccaaaa cgctgtactt aacatcaata
25141 attcaacgct ggaaattgga cttggagata tttttgagga agagggcttt aagattatcg
25201 cattcaatga gttttttgat acccaagtag atgatgtggt tatatcgaga aaaacgctta
25261 atggaaagta tataacccat ttttatccac aggtatcgga attggatggg attatttctt
25321 ctgataccca ttcatcagag caaattgttg gaaaactcga aagagaaacc ggaaaatcat
25381 caaaatatag gctgggaacg attgtgaaaa acggggagta ttttctcctt gcctttaccc
25441 attttgacaa gaataatagg gctttcttag aaatcaatga ctatattgac tgcctcatga
25501 atatgtggaa cgagtgtgat attcattacg gaggaaacac cgttgttttg ccgttgcttg
25561 gatctggaat tactcggttt cggggttatg aaaacataac ggatcaagaa ttgcttgaaa
25621 tcattatttg gactttcaag atgagtcgta ttcgattcca atatccagca aaagcaaaga
25681 ttgttttaac aaaggactgt ctgagtaaag ttaatcttta tgatttaaaa ttacaataca
25741 aatgattgga gggggagcaa acatggcaca taaatgtttt atttcattca aaaaagagga
25801 tcagtggtac cgtgataaaa tagacaatct cttttctgcc tcggatataa ttaacaaagg
25861 gcttgatcga gtgattgata gtaatgacgg agactatatt atgaaaacga tccgccaaga
25921 ctatcttaaa gactcaactg ttacggtatt tctcattggg acacattcgg ctgaaaatga
25981 gggatgtgat tattggggac ggcgtaaaaa ctatttcatc gaacgtgaat tgcaagcctc
26041 cttattcaat ggtgaaggta acacgcgaaa cggtattttg ggtgttgttc tccctgagat
26101 gtgtgatcaa atctacaagg gaacttacca atgccaaaaa tgtaatggaa cgcatagtta
26161 tgttgctatt gatgactcga cagtgatccg tgaattcagt gtcaactatt acattgagcc
26221 tcactctggc tgtgcgtggt cagatgatga tcgctattgt attcttgtaa aatgggatga
26281 ctttgtttcc aaccccgaaa aatatgtaga catggctttt gaaaagagaa ccagtccggt
26341 tgcggaaaga gttaaaattc gtaatcttcg atgatagttt tgaagatcaa caggcattcc
26401 ttatgtttta attatgggag gtcaaaatga aattttcatt tactgcgaaa aacgagccag
26461 aatgtaagtc tttcctcgaa atcaggatgc ctgatggtga gtatcagccg atagacgagt
26521 tcattaaagc acagcctgct atttctgagg cagatcttga gacatgcttt ataaagtatc
26581 gcggggacga gaaaacttat aagcaggatt ctgatgatta cgttgccttt tcggaaacca
26641 tgaagtttgt aaattttatc ccgccaatgc taattgtaga tcaatccatt aaaccagcaa
26701 tttatttcgc gcagaaagcg attgaatgct tacaatttgc tcggttcttc acaatgaaaa
26761 gcaatttgat tcttgacacg gactacaata ctcattgggc gcagggatat gttccacagt
26821 atctcttccg ctgtatctac tttgggacgg cgacaacatg gtattctaat gcctttgacc
26881 atgtactgca aactgtatat tgggggaaga aactatacac ctctgctgtt gacagagatg
26941 ggatccaata cagtgacaca tgggatgcaa agaaaatcat gacatattgt acatacgagt
27001 ttgttgtggg cgaattgaag aaatgcggag atactgcact cagaaagtgc ttgacctcat
27061 gctcaggcgc aatcgagcaa gtacggagtt gggcaaacta tatcaagcat aaaggcggca
27121 tcgactataa gtacctcgaa ccggaaaacc cgtttaagct atactgccat tcccatcagc
27181 caaagggcga aacagaagaa atcatctatg cctttacata ctatgagtcg ccaatcggac
27241 gtccaaactc tgcccctccg ttgacaccac cgagtacatc gggaggaagc aattccggaa
27301 gcatccggga tgactatgac aatcccgaag acctctggga agataaccaa gattggtatg
27361 aggacgagga tgaagcgtgg gatgagtggt acgacggtta agctccaaat gagagatttc
27421 ctgtttccag cgattttaat tccctgagtc gatgctctat tcataggaga cgctattcga
27481 ctttttagaa ccgcgtgact tttcactatg acgtgcaaag tcacgcgaaa caatgagggc

27541 agttcattat ccagataatt accggatagc aaataggttt ctcggaagcg aaaagacttt
27601 cacgattccc accgccgttg atcgaaacgt ccgattttag ggtgttttac agccgctcaa
27661 ccgcaagcgt cactgggtca atcacaatga tatgctttcc gttcctgccc tgtgcatagg
27721 tcacggtctt tcccgtccca ctgggtttgc catcccatac cgtaataatg aagtcggcgt
27781 gatcgaccat ataacggttg cgcttgtcca tgcagtcggg cgagtagtgg ctttggggta
27841 cggaaaagat acaaaaaaga gcctctgccg atacccaata cgggcatcga cagaggctct
27901 tggttcgttt tccgggcatt caaacaagtg ccgaaatcga ctcttatttc cgcgctaaag
27961 aagaaaccgc gcagggaata caatcactta ggttggtcaa tttgcgccgc tctgaggctt
28021 gcaatcacag gcttttccga cccctaactg tctacacctc gatgccttgc gggagaagtg
28081 cggacaggcg atgagaacgg cacggaagct ctgcttgcag tcatggacgc agcggacaca
28141 gaagcggtta tacttccgcc gtccgctgtc agctatgaag aacgaccact cctgcctcca
28201 tttctgactt ctgctcacaa ctcaatctct cctttgtgct ttttgggagg gcagcattcc
28261 ttctgctttt cagcctttgc agacttgagc ctgtccatga tggaggactt ctcgcccctt
28321 tcgggggctt tgacctctgc ggtcttgtcc tccttcggac cgttgttgat gatcccgtca
28381 atcatgccgt agtcatcctc ggtagacatc tcggcggctt tcagatagtt ttcccgttcc
28441 agcgcggcct tccactcgac cttctccaca ccgaagatac cacccttgc agcatgatcg
28501 tggatttcct ttgcatccag aaccagcccc tcggtattat caccgtacag ccggtagaca
28561 gaacaggttt tcatgacctc ggcggcagct tcctcccgca tggggagcat accgccccat
28621 gcgtagccgt agtccttcat ttcgtcagtc gtgatagaag gatccggcat ccggtcaacg
28681 gtcatgacct cctgacggag aagatcaacc gtatgttcat cgaagccgtc atacacatcg
28741 ctcagggaga gcttgccgtg ttcatccacg atgagacaag cagcttcatc gccgtagcta
28801 tcatgctcca tcaggaaaaa ggtatgaccg tcaacctcca tgtgatcaat ggtgtgccat
28861 gttcccacat ggccggaaac agccagaccg gaggtgtcca tgttgacctc aaaatatttt
28921 tgttccttga cttcctgctt ctgctgctct tcggagatca tgggaatcgc gacaattctg
28981 tctccgagct tggcgaactg ctccggctgt tcaaagagct ttgcatactt gagtgcaagc
29041 tcactatcca gagaaccgaa gctgtcatcg gtcagaccga caatgacaaa ggttccggca
29101 acaatatcat agatttctcc gtcttcatcc cgcagagcgc ggttgagagg aagaccttcg
29161 agctttgctt cctcgttgca aacaataccc accgggtcct catagggata aactgcctga
29221 atgtagccgc ctaccagagc ttgcagccct tcaaggcttg aatcgatctc tttgatttcc
29281 ggcttttttc ccggctcaac cacaacagca agaatattgc cgtccttcat aggctgttcg
29341 ccgcctctgg tgagacggta ttcgtcggga atgtcatcaa gactgccgtc aaacatccgc
29401 gtccaaccgt ctccggtatt gcgtatatag ccatcatcgg tgaatctgcc gccttcatcc
29461 attgccacat cgcgtccaaa acgctcaaag tcaatgtagt tagcgagatt acccattgct
29521 tttgtgtcgt agatacctgc ttcttcgatg tagtagcgcc ccagatcctc ttcatccgtg
29581 acatcgggaa ggtagtcgta gcagtcgaga ttgtaggtca ggttgatgag atcgtcgatg
29641 tcgctgactt catcgcagcc agcctccata atagcaacaa acttttcctg atcccatttg
29701 tccatctcat caatgcgaga ggcaaggtag ttcagcttat cgaggctttc atactcgccg
29761 agcagctttt gaaccccata gatgggacac tcgtagtcgg tgataaacca ttcctcgtac
29821 acatagccga actcatcctt cgagccgata ccgatgcgct caaaaacttt ctgcatttcc
29881 tcttcggttg tggggaactt cacccattca ccgacgagcg caccttcatt gtacttgccg
29941 agattggtga caaacgcttg gaaactgtca tccataacag gcataagcat cctcctttca
30001 gtcatccatc atgctgtccg gcgcaatata gagcgcgtcg aactcagcgt cagacattgc
30061 ttcgagcttc tggacggtct tgtgtatcag ctccgcgatc tcgctgtcct tgttgttcag
30121 agtgacgctc tctatgtcgt caatcagccg cttccggctg gatgtgttga agcagcacat

30181 gagattgatt tcttccgcgg taaagtattt catatcagag taccatatcc tttcctttgg
30241 ttttctgcgg tttcggaatg tgaggtgatt gagctttttg cgcggatttc agcttatcgc
30301 ggatagattc acgagcagca ggcttctttg gaatggtgtc atagatggca gtcccggcat
30361 cctccacata cgaacggaca tactggatgc tctcagaaaa gcggattggc ctgttcttgt
30421 cctcaatgtt ggtcagccgg acttcatcgg cagcaaaatc aaccgagtca attctgcacc
30481 tgatcccgtc cacattcagc gtcgcaccga caggaatgta atctgcggct ttgagttcct
30541 tgaaggtctt gccgtccttg ctgagaaaga catctgtgcc ggatttccag agcttgtgag
30601 aaccggctac ccacgcggtt tcacggaagc ccgtgaccgg catggagggc ttgcctcgca
30661 ccttcttctc aaggagcttt gtgccgagaa gaggggcggc tttttcggca tccttgccgt
30721 acagctcaaa atagccgttc tgggcaaagc aaaccagcgc gtccgggtga gcctgtttga
30781 ttgcctcata ctttctgagt tctgcaatag gcagaggtgc aagcacttct ttcgtttctg
30841 cctccggcag cttgtcacgg cggcttttct gcgtggacaa tacttctgcc aaagcatccg
30901 cgtcttttgg gagcgtctga tgggttacaa caccgccatg ctcaccgacc agagccattg
30961 cttcatcgaa gctgctgcgg cagggagcgt ccagcacata accgtccgca aaggtcagcc
31021 ggtcatgatc cggctcaact gagttccgcg ctacatttgc cgcctgttcc cggtaattga
31081 tctcaaagac atctccgaat accttgccac gcttaacatc tttcgggaca atcacgcggg
31141 caatgcactc gtcgtaggtc tgttccgcat aaaagcggaa ggtgttatga tcccgcgtcc
31201 cctgaataaa gacctgattt tcattgaggc agtgtgtccc atgcggacgg cagaaccaca
31261 taagcgtctt gtcttcggga ttctggcttt cggcagcgcg gcggatgatg cgcttgtcga
31321 tgtcgaagtc ttccttgaag ctgtctacat gactatccac gagcttctga agctccgcga
31381 tgatgtcaac gtctgtgtat ttcttcatgc gtctcctttc tcacaactcc atgtcgtgcg
31441 atttagcctt ggccggagct ttcttttccg gctgagattt ggcagcagcc ttgagttggt
31501 cacagatcga gggcttttcc tgctttttct cctgtcggat atactcaagc gtaggtgtca
31561 gctcccgata gcagccctgc gttttcttgg ccgaatgacg gcggtgaatg gcaaatagcg
31621 gctctccgct ctgcgtgacc gtattgccct cgatcctgac tccgttccga gccatgacac
31681 ggaaggactg ttccgatgac cgataagtgg aaagcatccc gtcgttattg tgagcaatct
31741 gcgccttgag ggtgtccata acagcgtccc tgatctgttt ttcctccggc gtgaggcgat
31801 gcttcgggac tgcggcttcc tgcggcttct ctgcggttgg tgccttgact tctacaacgg
31861 ttttctcgat ggagacaacc tcgtcgcgtt cccatgcagc aacctccggg ttctgcgtcc
31921 gggcgttggc ttcctgttcg cgctccttca catccatctg ccgcataaac tcggtgatga
31981 agagatcggt caaaccgggg tgtgccttat ccacaatgaa ccgggtagaa agctccgggc
32041 aaacaggtat gtcctttgcc cacgcctttg ctttctgcga gatccttccg tcccaatcct
32101 tgtgctggac ggtgtttgcc agaacatatt gcactcgctc cggcgagaac ttttcaagaa
32161 cgctcgtaac cgcagattca gcagcaaggc ggttattgcc gtaatactcg ctgatcgtcc
32221 gctctatggt atctctgcaa gcaatattcg cttgggcgga gagcctataa gcatccgttt
32281 ctccggctgc ataagcgtga tccgccgttt cggtgtaaac gggaataccg gcatcacgca
32341 gagaaggagc tacttccggc tctttggcgg gagtatctgc ttccagcgcc gcagtctggt
32401 ttttctctgc cagcagcacc ttgagcttgg cgtcaatgct gtcgatcatc tcagcggcgg
32461 tcttgcggat ggtttccaaa gagcctttca gctccttggt ctccttatct gaggaccaac
32521 cggcgatata gccgaaggaa taatccgaag tctcaatgcc gtaccgctgg cagacggtgt
32581 aggccacgct ctcggcttca acctctttgg tatggcgatc tttccgcatc tcctgaacag
32641 cggaggtatc ttcctcaacc tcaagccgcc actcaaaacg gttttcatca acgaaacacc
32701 atccctcgtg ttcagcagcc gcatttgctt ctgcctcaga cgcgaaacct gagagaaagt
32761 cccgcttggt gccgccatcg ctgatcatga cgattttcca tgtgggttcc ggctcaggcc

32821 ggtcaataga gtggagcttc gcatgggcaa tttcatgaat cgctgttttg atggtctgga
32881 tttcgctcat gccctcctga atggcgatcc ggttctctat cggagagaaa tagccctttg
32941 cgccgccggg aatgtcttca aaggaaatcg ggacgggaga gtcctgtttc agagcttcaa
33001 aaaaggcagc gtagttctcc acgctgccgg aaagctcatc cacaatgatg tcaggaagct
33061 ccttgccgtc ggtctgggag atgtcgaaga cactcaccac cttgaatgcg ggacgcataa
33121 cctccaccgt ctctgtgatg ggttttccgt ctgcatccag caccggcttc tgcgtggaag
33181 gatcgatctt ttcacgctct tcctttgcct tgtagggcgc aggagcgagt atcttgatgc
33241 ctttctctcc cttcatgacc tgacggtcaa aattgcgctg ccatgaggta tatccggcaa
33301 cataggtggc ctccggtttc tgcatggcga tgagaagggt gttgctgaat gagtagtgat
33361 aaaacttaga catcgtgcgg agatattcct tgaaccgctc ggattcaaac agctccttga
33421 tgccctgttc cagcttgtcc gtgatctcgt ggacttgctg tgcgcttttg ttttctgcca
33481 tgtgacacct ccttattttt caaactccat cttgaagctg acatattttc cgccggtatc
33541 atccagccgg atgatcgcat catagagctg gggccgcttc ggggtataaa gcccggtcac
33601 ccggcaccag cccttgtcca gcagctcctt ggcaatcttc ttggtcagct tctttttctt
33661 gctggaaaag aacttattgt cctcccacag acagaaggga cattccttgc tggaacagta
33721 ataattgccc ttccccacat acaccgggga gccgcagcgg gggcatttgc ctatgacctc
33781 cttgccggag tcgaagcgat ttgcctcagc atcggagagg aaggggtacg ccttgaccag
33841 cgaggcagtc atatccacga tcccggcgag gaacttgtcc ccatcggcct gaccgcgctc
33901 gatctgcatc agggtatttt cccattccgc agtcatggca ggtgaagtaa tctgttccgg
33961 cagaacacat acgaggttca ggccatcctt ggtaggcacg atggattttc ccttgcgctc
34021 ggcatagccg cttttgacca acttctcaat gacaccggcg cgggtggcgg gagtgccgag
34081 acccttttc tcggtatcat catcaaacgc ttcatttccg gcagtctcca ttgcagaaag
34141 cagcgtatct tcggtgtact gcttcggagg ggttgtataa tgctcggtta cactggctga
34201 tacggcatcg aggacatccc cctcgctgac ggcaggaagc atttttttccg gctcgtcatt
34261 ctccttgctc ttgagcgcgg ccttgaatcg ctgctcaacg gctttccagc cctcctgcac
34321 aacggttttg ccctttgtgc tgaatacata gccttcacag gagatcgtca aggatgtctc
34381 ttcgtaagtg tgcttctcac cggtggcaga cagaaggcgc atggcaacca gccggaggat
34441 tttctgctcc gactccggca atacggaaat gtcctgcttc tcaagctgga cggtagggat
34501 aatggcatgg tgatccgtga ccttcgcatt gtttgtgata cgccggatgt tgggagcata
34561 gctcaggccg tcgaaagccg acaccttccg gcacacaatt ccgatgacct gacggacggt
34621 gctttccatg tcatccgtga tgaactgact gtcggtgcgg ggataggtca gcagcttctt
34681 ttcgtagagg gactgcacca gatcgagggt ctgctgtgcc gtgaagccgt aataccggtt
34741 ggcttcccgc tgcaaggtgg tcaaatcgta gagcttcggg ggattgaccg tcttcgtctc
34801 gcgcttgaca gaagaaacga cggcctgttt cttatcacaa gccaccgcaa tcgtctttgc
34861 ctcatcttcg gttttgacct tctccatatc agcggtgaga ttgctgccgc tcacatggac
34921 attgaagtat ttctcttcct tgaaggtgct gatcttcccg tcgcgctctg ccagcattgc
34981 cagcgtgggg gtctggacac ggccaaccac cagcttcttg tgatagagcg tggtaaacag
35041 ccgggtgtca ttgatgccca caatccaatc tgcctttgac cggcagaggg cggcttcata
35101 gagcctgtcg aattcactgc tatcccgcag atgagcgaat ccctcccgga tggcagagtc
35161 ttcaagggag ctgatccaga ggcgcttgaa gggcttggtg caaccggcct tgttgtagac
35221 cagccggaag atcagctcac cctcgcgccc agcgtcggtg gcgcatacca gctcggtgac
35281 gcgcttatcc ttcatcagat cgcgcagcac cttgaactgc tgtgctttgt ccttcggaac
35341 ctcatacagc catttctccg gcacaatgag cagatcctcg tacctccact tggcatatcg
35401 ctcatcatag gaactcgcat ccgcaagctc cacaagatgc ccgacgcacc atgatacgat

35461 gtagttgttt ccttcgagat agccgtccct gcgggatgtt gcgcccagca ccttggcaat
35521 ggactgcgct acactcggtt tctcagcgat gactaaaatc aatccgtgaa cctccttctg
35581 aataagatga agcccctcgc atttctgcaa ggggccgcga tcaatcctta ttctgcgtca
35641 tcctccggct cgtcatcagc ggcaatctga ggctcttcat cctcgttgat atacggctct
35701 tcctcgtagc cttcatcgtc aaagaagtcc atatcctcgt ccttctgctt cttgccgcga
35761 acgaacttga tgtagtaata gcccgcgccg ccaataccgg ccacggcaat gatgacgatg
35821 atcatgccca tattgctgtt cttctggggc ttctcatcca ccacaggctc cggttcctca
35881 gcaggagctt ccggctcagg agccgcgccg gtgcattcac tcatgttggt tttgcagaca
35941 gggcaggaag tgttgacatg accggcctca cacttttcgg tacaggtaca ggtggtcagt
36001 gcagccacgg agtcctcatc catcagggca agcagatcac tttcgtccac catgttcagg
36061 aagtaggtct ggtactgttc ctcttcctcg ttgatgggcg catcgtagtc gatcacgata
36121 tagaaggtgt tgccgttttt cgtctgaatg gtgatgaact gcttgttggt ggctttgtca
36181 tacagcagat cgcgggtgga ggcgttgccc tcatcgtcaa tcggctcccc tgcgggaagc
36241 gtctctgcgg gcggctccgt ttccggcatt tcctccggct gcgtggcctc ggtcacggga
36301 agattctgct cggtgtcatc cgcgtaggca aaggcggtca gggagaagct gcaaaccatc
36361 acagcacaga gcgccagcac ggaaagcaca cggattcctt ttctcttact cattctcaga
36421 tacctccgtc tcagtgattt cagggatggc cgcgctgccg gttttcatcg cagacaggaa
36481 ggacatgatc tgatccttgt ccataaccat agcgcggaca gtgttgatga tttcgaggtt
36541 ttccatctcg gttttccgat catacagctc cttgagctgc gcctgaagat cctcgatttt
36601 cttttcggtc ttcgcaatct cagcgcagac cttctgatat ttgggattca tagaaaactc
36661 ctttcgctta atagggcggt cttccaaagg caaagaagtg agactgccag taggaagagc
36721 tgatggatgt gtactggatg ggatcgccgc agtgcagcat cacattgtca ccgacataaa
36781 tgcccacatg ggacacgccg ggggtgtcgt aggttccttt gaagaagatg agatcaccgg
36841 gctttgcgtc ggcctttgaa accggtgtgc agatgttgta tagcccctgc gcacccagcc
36901 gcccggtatt tacaagcccg ctgttggtca ggacatagga gacaaaaccg gagcagtcaa
36961 aggaggtttc ggggttggag ccgccccaca catagggata gcccaggtat ttctccgcct
37021 ccgcaagcag tgtcgtaaac cgctcatcac tcagatactc cgcaggaacc tcataatcgc
37081 tgggagggtt ggtgacatat ttgtccacat agccggatga cgggaacagg tcctcccggt
37141 tgcccagcac cgacatatag gtggcgtaca tggatagctc gtcctgtccc ataatgtaaa
37201 cggggacatg gctgagattg aagttttcca gcgtcacatt gcagatgtaa taagaatacg
37261 ggacctgaac ggtgtactca tattcctctt catagctttc accggtgtcc gggtctgtga
37321 ccgtcctcgt gccggtccgg gtttccgtcc ggtatcgggt ttcgacgatg accttttcgg
37381 tgaggatgta ctgcttctca aagagcatat cgagcgtccc ctgaacctcg gagagcgtcc
37441 attcgccttc atgaagggca gaaagaatgg agatcagcac atаggggtca tgctcgatgt
37501 catccagatc gaagtgatac tcgtcataat cgtgggtggc ctcgtagtta tccagcttat
37561 cttgcagctc tgcctccata cggcaatact gttcctcagc ccccagcatg gcatcatcct
37621 cgctgagata ggaggaagcg atcaccgcct gaccggtaga ggaaaacatc gcagtgcatg
37681 agctgacacc ggcagcaagc agaacgagaa tgcccagccc cacaccgaac cagataaagt
37741 atttcttgtt cttggagaag aactctttca ccttatcggc agctttctca ccggccttct
37801 tggccgtgga ggtagtcgcg gttccggcag tctgcgttcc ggcctttcga gccgccgcgt
37861 actcctgctt gatctgctgc ttctgatagt gtctgttcat gttccgctgc tttttcatct
37921 caggatgatc ctgctgggtc ttttcatagt gcagcttcgt atcggcaatt tcagcctcat
37981 gttccagctt ggacaccttc tcatagggct tgtttgcctg attctgcctg tgatgactgt
38041 aatgacgggc agcggattcc gcaaggattt cggtcttgtg ggcggcttcc acggcagagt

38101 tgtcctgctc gacctcatgc agctttccat gaacgacggt ggcaagggag tctccgacct
38161 tgttttccgt ctttttcgtc tcgaattgga gcttactcgg ctgcttggga ggcttgatct
38221 catcctcaaa gtggagacgg gttttgccct tgccggtttc ttcatcaacg acgcgctcct
38281 tcttgagaac cttcttggtg ggaagacttt cacgggcagc gtccagccgc tcatgcgcct
38341 gttccgattt tctttcgagc ttctgaatcc gcttcgtggg cgggacagca tcgtccatga
38401 ccggctttgc ggaagcagtt tcagccgctt gagaaagtac agcatccgcg tcaactgcct
38461 caaccgcaga ggtcttgcgc agcttgtgcg tgacaatcgc ttctgcaata accgtaccgg
38521 tatgagagga aacaccgtgc ggctccaaaa ccggaggttt gaacggtgct tctgacggag
38581 gcagcgtttc gggtacattc tggggtagtt tttccggctt tttctcttct gtgacagacg
38641 tttcggagga cggctgagta ttgctatcct cggcgtgttc cgcctgaaac tgacgctgct
38701 gactgcgcat ttgcaccttt ttttgttcct ccgcagaaat tgcttcggaa ggcgcatctt
38761 cctcaaccgg cttgaccagc tctgcatctt ccggcctccc tgaaactcgc ttctctgtgc
38821 cagccgtcag attttcctca accgcaccgt cctgcgtcat tctcacgaca accttgtcct
38881 tgggcttcaa ctgttgcttt accatcagat ttcacctccg atccgcgcct ttgcgagttc
38941 tgcgtactca ggattcagtt caatgccgat ataatggcgg tttagctgct ttgccaccat
39001 accggttgtg ccgctcccga aaaagggatc gagaacaatg ccgccttcgg gacaaccggc
39061 cagcagacag ggttcaacca gcttcggggg ataggcagca tagtgaccgc ctttgaaggg
39121 aaccgtattg accacccaca catcccgctt gttgcgcatg ggattgatca tgctgtcggt
39181 gatctcgccg tgcgaacgag gccggttgat ggtctgggtc tgaggctggc cgggaacggg
39241 cttgccgtat tttttgccgc cattcatacc gcgcttgagc ctttctgccg ttgccggagc
39301 tatcggttcc gaaattgcct tggcgtcaaa gaaatacttc cggcttttgg acagcaggaa
39361 aatatgctcg tagcagcggg aacagcgatc cttgcagctc tccggcattg ggttttcctt
39421 catccagatg atgtcgttac gcagatacca accgctgtcc cggagggcaa aggccagcat
39481 ccacggaatg ccgatcatat ccttcggctt gcagccctca accttgtagt tgagagccac
39541 agtctgaccg ttcctgccct tgggattctt ggggtcaacg aaatcaccct gattgccttt
39601 cccggcatag gtgtcggcaa tgttgagcca gagtgttccg tcggaacgca gaacgcgcct
39661 gacttcggta aacacttcgg tcaggcgcga gatatattcc tttggtgttg cctctctgcc
39721 gatctgttcc tccatgccgt agtcgcgcag cgcgtagtac ggaggggatg tgacgcagca
39781 atggacgctc tcgtcgggca gggctttcag aacttcgaga cagtcaccaa tgtagattgt
39841 gtcaagctgt atggggatca ctcctttcgc tgagattgaa gaggctcagt cggagaaata
39901 ctcgtccggc tcatcgtcct cgtcgatttc atcctcccag cggttcagaa tgtcctcggc
39961 ttcctgcgcg gcctcgcggt acaggctcag gcggttcttt tcaaacgcgg caaaggcggg
40021 aacatacttt ccgaactcct gaatggcgcg gacatccttg gtgcgcatat ccgcaaccat
40081 gcccagcagc tctgccatcg tcagcagatc ctcaatcatg agatcgtact tatccttggg
40141 aatggtgacg gcttcgccct cgtcctcagc ctcggcaaag gggattgtgc cctcagcgga
40201 cacatggtag aaatgctcct tcgcgttcag ctcatccacg ataccggcga tgaactccat
40261 cacggcatca gcacggtttt cgatctcggc cctgacatgg aggtacgccc gatggtgctt
40321 tttcagctct tcctccttgt ggatgagatt gaccagctcg atggtcatac ccatcaggcc
40381 aaagaagctg tccatgccct tcacaacctc gcggaaggtc atctcgacgg cagcatcacc
40441 gaaagcctcg gtaaactctg ccatcatgcg tttgcagtcc atattggtgt tgtagttctt
40501 atccatattc aaaatcctcc ttgtaatcaa gccgtttctt cgagcttggt ggtcatgatg
40561 cggtaaagct gggtgtcctt cgggaagtca tccttgaagg gaacgatggt ggagccgtag
40621 aagatcagcc cctcaccggc gttggagttg gtgatgtagt tctgctggct gggagagata
40681 ttcagagctt cactgaggat ttttctgtct cccgaagcct gattcagaag atagacaaaa

40741 tccgagttct cgaagatgtt ctcaacctca cgggaagcca gaagatcctt cacattctgg
40801 gtgatcccgg tggggatgcc gccccattta cgaaaccgct tccagatttc cacgctgtat
40861 gcggcggtct gttcctcttt cagcagcaga tggaactcgt ccatgtagta ccgagtggcc
40921 ttatgctggg cgcggttgat tgtgacgcgg ttccacacct gatcctgaac aaccagcata
40981 cccagcttct tgagctgttt gcccagctcc ttgatgtcgt agcagacgaa gcggttgttg
41041 acatccacat tcgttctgtg attgaagaca ttcagagagc cgtgaacata gatttccagc
41101 gcagttgcga tgcgctgtgc ctccggctct ttttggttgc gcagaatgtt gtagaggtct
41161 tccagaatgg gcattttctc cggtctggga tcggcaaggt agtcctgata gaccatgcgg
41221 acgcttcggt caatgatggt cttctcaacc ggctgcaagc cgtccttgcc gccaacaatc
41281 agctcacaca tggagaggat gaagtcggac ttcaaggtca gcgggttttc ctcttcggag
41341 tagttgacat tgatgtccag cggattgatg tagtcggtac tgatgggcga aatgcggata
41401 acctgaccgc cgagcttctg cacgagggga taatactcag cctcggggtc gcacacgatg
41461 atgtcatcct cggtgatcag gaaggcgttg gtcatctcac gctttgccga gaaggacttg
41521 ccgctgccgg gagtaccgag gataagccca ttggggttct tcaatttctt cctgtccacc
41581 atgatcatgt tgttggacag cgcattcagt ccgtagtaca gagcctcgcc gccctgaaaa
41641 agctcctgcg tggtaaacgg gacgaaaacc gctgtactgg aagtggtcaa cccgcgctca
41701 atctcgatct ggttcaggcc gatggggagc gaggacatca gcccctcttc ctgctggaag
41761 tccagacgct tgagcgcaca gttgtatttc tgtgcgatgg atgccgtctg gaagacagcg
41821 ttttcgagct tctgacggtt tggcgctgtg ttcatgatga ggatggtcac gaggaacatt
41881 ctctcgtttc gcgtctggag atcctgcaac agacgcttgg cctctccgcc gtaggtggca
41941 aggtcggacg ggatgatgtc catatcgtag ccggaacgga cggccttctt ctgctcttca
42001 attttcatct tgtccagatc ggtaatcttg gacttgatgc gcttgatggc ctgagcctga
42061 tcaattgtgc ggatatgaag attgaccgta atgttagagt ccatatcgag gaagtctgcc
42121 agcatacggt cattcagctc cggcgcgaga atctgcaaaa agctcaccgc gccgatggtt
42181 ctgcccatct tgaagcactt gccttcccga aagtcgaatg aggtaggcgc gatgaagtct
42241 ttgctgctga gaccggtacg ggcaaccata tcgaaggaga agcggaatgg ttcgccggag
42301 tccatgttga acacatcatg caggactttc agacgttcat agccggacag cggctcggtt
42361 ttcacaccga gcgtcttgaa gttgttgagg atgtcagctt ccacccggtc gagctttgct
42421 ttggcggtac gcagagagtc tgcctcaatg ccgaaggtaa tgtacttgcg cttgacaagg
42481 ccgttgttgc ctttggtgag ctgggtttgc agcatatccg aatactcagc gcggatgtca
42541 ttgaaagcat cgtcctgttc ggggatgcgg attgccttct tgaactcatt gatgttcact
42601 ctctgattga tgaaggagag ctgaacaaaa atggagctgt cgaagtagtt caggaagtca
42661 caccagttct caaagatggc ggtcttgtcc tcgttctggg caagctggta attgatgtcg
42721 ttatagcgga tggtcttggt gtagaagcgg ctgttcacct tgcagatacc gtcgcggcac
42781 atctccacat aagggatggt ctgctgggcg gacttgggga ttttctttgc cttcttgtcc
42841 tttttcttct gcatgacgat ctgcttcttt tccgcagcgg aaagagcatc gccgtacacc
42901 ttgccatttt tcacggtctt cttaggcgtg tctgcctttt tggtgctggt tgtcaatctg
42961 caattcctcc tgttccttga tttctcgctg aatggcagcg taaagattgt ttgtgcggta
43021 gggacgcacc ttgtccctca gaaacatgga cttgatcaca tgaccgagga ttttttcagc
43081 cggttgcccg tccttctcat agagggcgaa gaagataaac gggagcatca gtacgaccat
43141 cagcattgct gccgtggaga tgccgaggct gggcttggtg agaaagtaca tgggaacacc
43201 taccgcagca gccagcgcga agcagatgat ctgccgcttg gtcaggttga acatgacttt
43261 ggttttcaca cggttaaggt ccttcgggac cggtacgaat gccattttct gaaacctcct
43321 ttccgagaga tacatcgctt tccacttcgt tgccccacac gtcccatccc tccggggact

43381 gtctggcaaa aagctcaatc cgtggaagat cgcccatgag ctgaacaatg cgttcacggg
43441 cttcatccgg cttcttgctg tgttcctcca catggctgag gatgacctga cggacaccgg
43501 cgttcacacg cctcggatgc ccctttgtgg caagaatgca gatttcggca tttgcacggg
43561 tccagtagcc catgccggtg aagagatcat cgttcttccg gttctgcttc acccacacga
43621 aagcgacggt tttatatgaa aatccccatg cctcaagcac ttcgagcgct tcatagagac
43681 atgggaatgt gatccacaga aacagagcgc agtctttggc ggcaagctca ccaacaggta
43741 aagccttaat gtcctcaatg ctcatcgttg ggtagtggct ttccgctgac cgtccctgtc
43801 ctttctttga ataggtccga tacgcccaag gcggatcggc ataaatgatg ctgtactttt
43861 tcatcccgcc gacctcctta atgcgcgttg aacaacgact tcgagaggct gccggtcttg
43921 aacagcgaga agcacaggat gaccgtgtat gctgccaccg agaaaagagc cgagtgaatg
43981 ttggctgcta tgatcatgtt gttgatgagt acggcgtaga tgccgacgca gatcatgatg
44041 aggaagcctt ggaatgccag agcgaaaaga cccttgaggt aatttgtccc gatgctgccc
44101 cattctcggt tgctcattgt agcaataggg atcggcgcaa tgctgacggt gcagtaaatt
44161 tcgatcattc gtccgtacag gatgactgtg atcaggatag acatgatctt gaggcatagg
44221 ctgatgacca gcgtttcaat ggcaagaccg agcagttcac caacacccat tgcttccata
44281 ctcgttcgca tggtgtccag tgtcgaggcg atatcaatat ttgtggtgcc gtgaatgact
44341 ccggctgcgc ccgcaaccac ggactgtccg atgtcgaata ctgccatcac gatgtcaaag
44401 gtattggtca cgaggtagat tgcaacagcc gctttgaaga accacttgaa gaacatccaa
44461 gtgtccatgt cgtgcatatt gtttttctcc gtgatcatgg agatcagctc gtaacacaaa
44521 acgaaggtga taatgatacc ggcaatgggg actatcacgc tttcggatag tccccggatc
44581 atcgtataga tgctgccgtt ccatgtagaa ggggtcctgc cgacctctcc ggcaatcgtg
44641 cctaccttct cattgacatc ggtgaacata ttggtcaggt tgctttcgat catgccgatg
44701 agcatttcct tgatggcttc tgtgattttt tctaagatac tgcccaatat ttcaccacct
44761 ttcggttaag ctgggcttgc tggattagaa caaaccggac agcagaggga tcagcgtagt
44821 tccgataacc accacaccgc cgccagccat gagctgcttg atgccctgac tcttagcgcc
44881 gggattgtca ttgccgtagc cttccatcaa gttgaccacg ccccacacgg cgaggccagc
44941 gccgagggcg atgacgagag tctgtaaaac cgtgactgcc tgagtgataa atgccatagg
45001 ttatttcctc cttattgttt tgaaaaatga agggagggcg cgtttcctgc gtcctccctt
45061 gaggggtgaa agggactgtc ggtatcagaa cagaccggac agcaagggaa tgagggtggt
45121 gccgatgaga accacgccgc cgccagccat gagctgcttg atgccctggc tcttggcgcc
45181 gggattgtca ttgccgtagc cctccatcag gttgaccacg ccccacacgg cgagaccagc
45241 accgagcgca atgacgagag tctgcaggac cgtgactgcc tgagtgataa atgccataga
45301 tttttcctcc tgatttcatg ttgtttttttg aaaatgggca taaaaaagaa gccccgtcgc
45361 tgttctgctt tgaaccttgc cgtcaggctc ggctcactgc gcagtgcagc gacggggctt
45421 tagtcttctt tgatctctcc catatcatag agatcaaagg tgtcggtggg cttgaccacc
45481 agcttgcggg agcgatattt ctcgatgtca aaggcgtttc gcttatcgta gtcagccagc
45541 agtttgtatc ttggatgctt cgttatgtcg tacttatcac tgagaaaagg tctcacgcct
45601 ctaagctgca agatacattt gctgccgtcc atgacggcga tttcgtcctg agacatcaac
45661 tccttaccaa ccttctgata gttcagcccg taggaattgt tggtggaacg ggtttccgaa
45721 gtgttgtata ggtcaatcgt ttccttgccc agcacttcgc tgagttcttt cagggtggat
45781 ttttcctttc cgccgaggaa gagggtgcag tcgcagttgc ccgcaatcgt gtcagccgca
45841 tccttgtaga tggtcttgag ctgggactgg gactgcaaga tgattgccgc cgagatttcc
45901 cggctgcgga tggtggcgat gagcttatcg aacttcggta tctggccgat attcgcaaac
45961 tcgtccagaa gacagcggac atgaacaggc agtctgccgc cgtactcgtc atccgcctta

46021 tcacagagca gattgaacat ctgagaatac atgatggcac agacgaaatt gaaggtgtca
46081 tccgtatcgc tgatgatcac gaacagggcc gttttcctgt ctccgagggt gtccagctcc
46141 aattcgtcgt agctcatcag ctcccgcagc tctgcaatgt caaagggagc aagtctggca
46201 ccgcagctaa tcaggatcga cttcgcagtc ttgccagccg ccaacttgta tttgcggtac
46261 tgcttcacag caaagtgatc cgggtccctc gcctccaact cgtcaaacat gatgtccacc
46321 gggtttttga agttttcgtc atcctctctg gcttccgaag cattgatcat ttcgagcaag
46381 gtagtgaagt ttttctcgtg ttccggtgcc tcgtaccaga tgtacccgat gagggccgta
46441 tagtagagct tttcagcctt cacccaaaaa tcttcgccgg acttatcgcc gtctcccttc
46501 gtgttgacga taattgtatt gaccagcttg aggatgtcct tctcgctgcg gatgtacgag
46561 aacgggttat agtgcatgga ttttttgaag ttgatcgtgt tcagcgactt gatcacatac
46621 ccgtttttct caagcatttt gcccgtttcc accaagaggg tgcctttcgg atcggtgact
46681 atgaagctga ccggataatc cttcgagacg cactgcatga ggttgggctt cacaaaaaaa
46741 cgggttttgc cggaacctga accgccaatg accatgatgt tcttattccg ggcgtacttt
46801 ggctgcttcg ggcggctgtt catggtcagc gcctcggtct gggtaaggat gacattgttc
46861 tcaaaattat catccatata cggcctgata tcctctggct tgccccatcg ggcagatccg
46921 tattcttcgc cctgacggaa cttcttgcgg tttttgcctt tcacataaac cgccagcttg
46981 agcaaagcgc ctgccgctac gccgatgagc aagtcaaccg gatgaaagct gggcagagga
47041 tttgcaaatg ctgtcccgaa gttggcaaag cccccggtga gcttgtcgat cagctcagtt
47101 ccgggagcct gtctgaacac cgcagcaatt ttgtccacga aatagaacgc gaacacatac
47161 gggaagttga gaagaataag gcgtttggtg tcgagcttct ttttcacagc tccacgcccc
47221 gatccttggt cttgaccttg accttctcct tgtgcttggc cttggactgc tcctgttcat
47281 gggagagctt gcgccggaga gaaggcttgt ccttctgctt gactgtcctt gcggaaaact
47341 ccttgaaagc ctgtgtcatc acatccacat cacgcccttt gaagaatacg atgtagcgtg
47401 gcggctcttc cgaagtgtcc ttcttgagcg catagtcaat cccgtatttg ttggcggtgc
47461 gctggaagga cttgatattt ccgtcggtca cttcaatgtt gctgatggcg gcattctgag
47521 ccacgagatg ctttatggac tgcttgcccc ggtaggtctt aggctgagag gcggttttct
47581 gagccttctc aatttcctcc acaaacttct tgagagcttt gtccagcacc tcagcagtga
47641 ttttgccgcc cttaatcgca atagcaacaa ctttggtatt tacttcatcc tgcatgggaa
47701 cctcctttct gagaatcgcg agatcagttc aacactaagg tggtactctc cgataaaaaa
47761 tgattgtcac gtttgatacc tccttcctgg gtggattact tctgaccgta gagatcatga
47821 tttacaagag ccgaatagta gttgtccata gtcagggatg cgttatacag tgttgtgagc
47881 atatatgctc tgatgttccg aatgtcggat gggcattcat ccattgcatg aaggacatac
47941 tctatatggc tgctgttgag cttgaggaaa cgggatttta ctacttgctg cggcatatcc
48001 tcgccgttga tacggatagt gggacgagta gaacaaatcg cgtccagcat aatctcgatc
48061 atttcattta cacggtcagt atcaaaccgg tgatcctgag agagtatttc aagctcaaga
48121 ttatctcgga taagctcttc atatctcgct ctttcgtcaa tcccatccat cccatcaaga
48181 ttgatagatt gatacttact caaagagtta tttctttctt gagtaattac ttgattagta
48241 tttatttgtg ccggatttcc ctcgtgtggt ttatccatgt ctggattatc cacatcggga
48301 ttttccatgt gtggactgcc cgtatctggt gataccggtt gtggctcttc aaatatgtcg
48361 aagattgtgc tgctcaatct gcctttgctg tcccgttgtc gatgccggac gagatagccg
48421 tgtttctcaa gctctttaag ggctgagagg atggcatcgg gaccttcctt gcagatcgca
48481 gacaagccgc gagttgaaaa cttccagcca tcgttaaacg acagcatttt tgagagaaga
48541 ccgactgcct tgagggacag gtttttgtct cttagatgga agttcgacat aaccgtgtag
48601 ttacgacttt tgtttacgcg gtagactgcc atgcgatcaa ctctcctttg gtaagagagc

48661 tgcaagttca ttctgctttt tcatgtactt gagccgaatg atttctttgc cctttggcgt
48721 gaaatatgtc tgtgatcctg tgaacgagaa ggtcacgaaa tcacgaacga tgaataagcc
48781 ctcgttgctg tgcttgttgt acggaagaag ctgaccggaa ggagagcgaa acaggaattt
48841 ctccttgaga agaaactgga cgaatttgcg ttccgggatt ttaagctcct tcgccgtcgt
48901 tcgtatattg gtgcagtcat ccggattgat gaaagcatca aaatagtctg ctttgggctt
48961 cgctttggat aactcagctt ccaatgccgt agcgcgtctt ctttctgcaa gaagcgcgtc
49021 tgcaaactca aagatgacag cgggatcatt ttgaatgcac tgcaataacg gctcggacaa
49081 ataacctccc gtctttcgga tggcggggat gacctcatga gtgacccagc gtttgaaggc
49141 tttggcttcg ggttttcgag aactgagaat aagcgagtaa aggccagctt cgtttacaaa
49201 agagagtttg gatttgccgc ttgctgcact atcatttgaa atgatagtgg taaatctctc
49261 gtcttcctcc agtcggctaa gagcttgggg agcgttcttg atctccaatg ctttacacac
49321 atcagaagcg gcaaaccaag gttcgccatc aatcagcgtt atgcggactg cgccaaactg
49381 ttcattctga aatacttcaa ttttgttctc catttgattt ttcctttctg gcagcagtgc
49441 tgtatgtgtt ttcgtttcgt gttgatacgg ggacgaggta ataagggcat tcctcaaaaa
49501 cgcaggactg atatttccaa taagggcgat ggaatctgca tgaccggcat tccgggcaga
49561 ttccatcgca gccgctatcg tagtggttat acggcgtttc tttcatcaat gcctcaaacg
49621 ccaatgagca tcctttcggg tacattttgc ttacctccat ttttgtgata gaaatagaaa
49681 aagccccgtc attgttctgc tacgcgcagt gcaataacgg ggcttgccgt gtattaagtt
49741 cttgcaggaa ttacagaaaa cttacggttt actgtttgga caaaatcagc gaaacaccat
49801 ttgaagggat attatatcaa gcaactgtga atgctgcgtg acatcattca tctctctggt
49861 caaagaagct gcttatccaa tttttttaat tctttcaaga aagtcgaaac gaacagtaga
49921 tgtcgccaag attttctttg tatgccgcat gagtatggcg atataagaaa ctccgcaaga
49981 ccgatttttg gaataataaa ctttcctgaa attttgaagc caacgagctt cttggcatta
50041 caccaacgat aaattgcagt gctgctgtaa ccggttattt ccattgtttc ctcaacagac
50101 aacaaatcac cataagaggc aagttctgtt cgtgtgtatt cttcaaagag ttttctttca
50161 atctccgata gactactcaa ctcatatctg aatgtatttc gtgatctccg ataacctgaa
50221 cgatctcgat accaatgatc cggtgcgcca tatttctcag gatgtatttc acggtcaatc
50281 aaataagtta ttacatcgtc ggtacggatt gtgtaccggc gtgttttttt gccagtgtcc
50341 ttacaaggga ctaatccgtt ttgcagaaga tgaagggctg ttgctttgct gatgtgagct
50401 atccggtaga actggtcttt ggagatcgtt tcaggatagg ctctgcggat agcatcatat
50461 tttttctctg tttcagtcat gctgataacc accttttag gtattcggta caaggtaggt
50521 gttatctgca tttatagggg tttgttctcc ccttgttctc cccaaaatcc ctttcaaaac
50581 tcgttttctc ccctgaatcg agcaaaagag gggattcatt cgaactcctt ttatatcaag
50641 gtttttgggc gtttgacacc agaaaagcct tgatattgct cactttttcg aagtccttca
50701 ttgtccgcta agttcccta tttcctctaa agttccaaag cctgactatg actcgaaatg
50761 cggtaggtcg ggtaaccggc gcgagggttc aaatccctcc atctccgtct aaggaaaaag
50821 ctgcattccc gcctgtgatt caaggcggga atgcagcttt tttaaatata ccacagtggt
50881 tttaattgtc tccctccgcc acgatttctt ccaatacccg gattaaatcg aaaactgttt
50941 ccaggcggac gttgcgttcc aggatacagt tttttaaatc gatagacaga gactcaaatt
51001 tgtcccggac tgccggggat acataggatg ccat

//

DEFINITION Integrative and mobilisable element IME_5

CDS 1..1548

/color=8

/locus_tag="C83_00259"
/note="Methyl-accepting chemotaxis protein  HAMP
domain-containing protein, similar to AA
sequence:UniProtKB:P05704 protein motif:CLUSTERS:PRK15048
protein motif:Pfam:PF00015.15"
/product="Methyl-accepting chemotaxis protein  HAMP
domain-containing protein [two-component system]"
/"ID=C83_00259
/protein_id="gnl|SC|C83_00259"""
/codon_start=1
fasta_record    1..129008
/ID=C83|SC|contig000001
/colour=5
CDS    complement(2254..3657)
/color=9
/locus_tag="C83_00260"
/note="Tyrosine recombinase, protein
motif:CLUSTERS:PRK00236 protein motif:Cdd:COG4974 protein
motif:TIGRFAMs:TIGR02224 protein motif:Pfam:PF00589.16"
/product="Tyrosine recombinase [recombinase]"
/"ID=C83_00260
/protein_id="gnl|SC|C83_00260"""
/codon_start=1
CDS    complement(3675..4544)
/color=9
/locus_tag="C83_00261"
/note="DNA binding domain  excisionase family
Helix-turn-helix domain, protein motif:TIGRFAMs:TIGR01764
protein motif:Pfam:PF12728.1"
/product="DNA binding domain  excisionase family
Helix-turn-helix domain [excisionase]"
/"ID=C83_00261
/protein_id="gnl|SC|C83_00261"""
/codon_start=1
CDS    complement(4656..4874)
/color=9
/locus_tag="C83_00262"
/note="DNA binding domain  excisionase family
Helix-turn-helix domain , protein motif:TIGRFAMs:TIGR01764
protein motif:Pfam:PF12728.1"
/product="DNA binding domain  excisionase family
Helix-turn-helix domain [excisionase]"
/"ID=C83_00262
/protein_id="gnl|SC|C83_00262""""

/codon_start=1
CDS complement(4921..5319)
/color=8
/locus_tag="C83_00263"
/note="Type II toxin-antitoxin system PemK/MazF family toxin MazF, similar to AA sequence:UniProtKB:Q7A4G9 protein motif:Pfam:PF02452.11"
/product="Type II toxin-antitoxin system PemK/MazF family toxin MazF [toxin-antitoxin]"
/"ID=C83_00263
/protein_id="gnl|SC|C83_00263"""
/gene="mazF"
/codon_start=1
CDS complement(5772..6440)
/ID=C83_00264
/color=4
/locus_tag="C83_00264"
/note="Membrane protein trbL/VirB6 plasmid conjugal transfer family protein"
/product="Membrane protein trbL/VirB6 plasmid conjugal transfer family protein"
/protein_id="gnl|SC|C83_00264"
/codon_start=1
CDS complement(6641..8257)
/ID=C83_00265
/color=9
/locus_tag="C83_00265"
/note="Serine recombinase"
/product="Serine recombinase [recombinase]"
/protein_id="gnl|SC|C83_00265"
/codon_start=1
CDS 8483..8683
/ID=C83_00266
/color=8
/locus_tag="C83_00266"
/note="GNAT family n-acetyltransferase"
/product="GNAT family n-acetyltransferase"
/protein_id="gnl|SC|C83_00266"
/codon_start=1
CDS complement(8748..8942)
/ID=C83_00267
/color=9
/locus_tag="C83_00267"
/note="Transposon-encoded protein TnpW"

/product="Transposon-encoded protein TnpW"
/gene="tnpW"
/protein_id="gnl|SC|C83_00267"
/codon_start=1
CDS complement(8957..9799)
/color=10
/locus_tag="C83_00268"
/note="Primosomal protein DnaI ATP-binding protein,
similar to AA sequence:UniProtKB:P06567 protein
motif:CLUSTERS:PRK08116 protein motif:Cdd:COG1484 protein
motif:TIGRFAMs:TIGR00362 protein motif:Pfam:PF01695.11"
/product="Primosomal protein DnaI ATP-binding protein
[replication]"
/"ID=C83_00268
/"
/protein_id="gnl|SC|C83_00268"
/gene="dnaI"
/codon_start=1
CDS complement(9796..10647)
/ID=C83_00269
/color=10
/locus_tag="C83_00269"
/note="Replication initiation protein RepA [replication]"
/product="Replication initiation protein RepA
[replication]"
/gene="repA"
/protein_id="gnl|SC|C83_00269"
/codon_start=1
CDS complement(10663..11049)
/ID=C83_00270
/color=1
/locus_tag="C83_00270"
/note="Cysteine-rich VLP domain-containing protein"
/product="Cysteine-rich VLP domain-containing protein"
/protein_id="gnl|SC|C83_00270"
/codon_start=1
CDS complement(11039..12664)
/ID=C83_00271
/color=4
/locus_tag="C83_00271"
/note="Relaxase/Mobilisation nuclease domain [relaxase]"
/product="Relaxase/Mobilisation nuclease domain
[relaxase]"
/protein_id="gnl|SC|C83_00271"

/codon_start=1
CDS complement(12636..13034)
/ID=C83_00272
/color=4
/locus_tag="C83_00272"
/note="Plasmid mobilisation relaxosome protein MobC [relaxosome]"
/product="Plasmid mobilisation relaxosome protein MobC [relaxosome]"
/gene="mobC"
/protein_id="gnl|SC|C83_00272"
/codon_start=1
CDS complement(13532..13744)
/ID=C83_00273
/color=7
/locus_tag="C83_00273"
/note="Helix-turn-helix domain-containing protein [transcriptional regulator]"
/product="Helix-turn-helix domain-containing protein [transcriptional regulator]"
/protein_id="gnl|SC|C83_00273"
/codon_start=1
CDS complement(13737..14159)
/ID=C83_00274
/color=7
/locus_tag="C83_00274"
/note="RNA polymerase sigma factor Sigma-70 region 4 [transcriptional regulator]"
/product="RNA polymerase sigma factor Sigma-70 region 4 [transcriptional regulator]"
/protein_id="gnl|SC|C83_00274"
/codon_start=1
CDS complement(14331..15098)
/color=8
/locus_tag="C83_00275"
/eC_number=2.1.1.197
/note="Malonyl-CoA O-methyltransferase BioC, similar to AA sequence:UniProtKB:P36571 protein motif:CLUSTERS:PRK08317 protein motif:Cdd:COG4106 protein motif:TIGRFAMs:TIGR02072 protein motif:Pfam:PF12847.1"
/product="Malonyl-CoA O-methyltransferase BioC [methyltransferase]"
/"ID=C83_00275
/protein_id="gnl|SC|C83_00275""""

/gene="bioC"
/codon_start=1
CDS complement(15149..17068)
/color=2
/locus_tag="C83_00276"
/note="Tetracycline resistance protein tetM from transposon Tn916, similar to AA sequence:UniProtKB:P21598 protein motif:CLUSTERS:PRK13351 protein motif:Cdd:COG0050 protein motif:TIGRFAMs:TIGR00484 protein motif:Pfam:PF00009.21"
/product="Tetracycline resistance protein tetM from transposon Tn916 [antibiotic resistance]"
/"ID=C83_00276
/"
/protein_id="gnl|SC|C83_00276"
/gene="tetM"
/codon_start=1
CDS complement(17709..17849)
/ID=C83_00277
/color=4
/locus_tag="C83_00277"
/note="Maff2 family  conjugal transfer protein"
/product="Maff2 family  conjugal transfer protein"
/protein_id="gnl|SC|C83_00277"
/codon_start=1
CDS complement(18048..19103)
/ID=C83_00278
/color=8
/locus_tag="C83_00278"
/note="PHP plymerase and histidinol phosphatase domian-containing protein"
/product="PHP %28plymerase and histidinol phosphatase%29 domian-containing protein"
/protein_id="gnl|SC|C83_00278"
/codon_start=1
CDS complement(19133..20179)
/color=8
/locus_tag="C83_00279"
/note="Fic Filamentation induced by cAMP family protein, similar to AA sequence:UniProtKB:Q9K0V1 protein motif:Cdd:COG2184 protein motif:TIGRFAMs:TIGR02613 protein motif:Pfam:PF02661.12"
/product="Fic %28Filamentation induced by cAMP%29 family protein [post-translational modification]"

/"ID=C83_00279
/"
/protein_id="gnl|SC|C83_00279"
/codon_start=1
CDS complement(20427..20915)
/color=8
/locus_tag="C83_00280"
/note="Cysteine hydrolase family protein
Nicotinamidase-related amidase, protein
motif:CLUSTERS:PRK11609 protein motif:Pfam:PF00857.14"
/product="Cysteine hydrolase family protein
Nicotinamidase-related amidase [hydrolase]"
/"ID=C83_00280
/"
/protein_id="gnl|SC|C83_00280"
/codon_start=1
CDS complement(20912..21571)
/color=7
/locus_tag="C83_00281"
/note="Transcriptional regulator Uncharacterized conserved
protein Uncharacterized conserved protein DUF2174, protein
motif:Cdd:COG4832 protein motif:Pfam:PF09942.3"
/product="Transcriptional regulator Uncharacterized
conserved protein Uncharacterized conserved protein
%28DUF2174%29"
/"ID=C83_00281
/"
/protein_id="gnl|SC|C83_00281"
/codon_start=1
CDS complement(21574..22491)
/color=7
/locus_tag="C83_00282"
/note="YafY family transcriptional regulator HTH and WYL
domain-containing, protein motif:Pfam:PF08279.6"
/product="YafY family transcriptional regulator HTH and
WYL domain-containing [transcriptional regulator]"
/"ID=C83_00282
/"
/protein_id="gnl|SC|C83_00282"
/codon_start=1
CDS complement(22836..24452)
/color=9
/locus_tag="C83_00283"
/note="Serine recombinase, protein motif:Pfam:PF07508.7"

/product="Serine recombinase [recombinase]"
/"ID=C83_00283
/"
/protein_id="gnl|SC|C83_00283"
/codon_start=1
CDS 24678..24878
/ID=C83_00284
/color=8
/locus_tag="C83_00284"
/note="GNAT family n-acetyltransferase"
/product="GNAT family n-acetyltransferase"
/protein_id="gnl|SC|C83_00284"
/codon_start=1
CDS complement(24943..25137)
/ID=C83_00285
/color=9
/locus_tag="C83_00285"
/note="Transposon-encoded protein TnpW"
/product="Transposon-encoded protein TnpW"
/protein_id="gnl|SC|C83_00285"
/gene="tnpW"
/codon_start=1
CDS complement(25152..25994)
/color=10
/locus_tag="C83_00286"
/note="rimosomal protein DnaI ATP-binding protein,
similar to AA sequence:UniProtKB:P06567 protein
motif:CLUSTERS:PRK08116 protein motif:Cdd:COG1484 protein
motif:TIGRFAMs:TIGR00362 protein motif:Pfam:PF01695.11"
/product="rimosomal protein DnaI ATP-binding protein
[replication]"
/"ID=C83_00286
/"
/protein_id="gnl|SC|C83_00286"
/gene="dnaI"
/codon_start=1
CDS complement(25991..26842)
/color=10
/locus_tag="C83_00287"
/note="Replication initiation protein RepA, protein
motif:Pfam:PF06970.5"
/product="Replication initiation protein RepA
[replication]"
/"ID=C83_00287

/"
/protein_id="gnl|SC|C83_00287"
/gene="repA"
/codon_start=1
CDS complement(26858..27244)
/ID=C83_00288
/color=1
/locus_tag="C83_00288"
/note="Cysteine-rich VLP domain-containing protein"
/product="Cysteine-rich VLP domain-containing protein"
/protein_id="gnl|SC|C83_00288"
/codon_start=1
CDS complement(27234..28859)
/color=4
/locus_tag="C83_00289"
/note="Relaxase/Mobilisation nuclease domain [relaxase], protein motif:Pfam:PF03432.8"
/product="Relaxase/Mobilisation nuclease domain [relaxase]"
/"ID=C83_00289
/"
/protein_id="gnl|SC|C83_00289"
/codon_start=1
CDS complement(28831..29229)
/color=4
/locus_tag="C83_00290"
/note="Plasmid mobilisation relaxosome protein MobC [relaxosome], protein motif:Pfam:PF05713.5"
/product="Plasmid mobilisation relaxosome protein MobC [relaxosome]"
/"ID=C83_00290
/"
/protein_id="gnl|SC|C83_00290"
/gene="mobC"
/codon_start=1
CDS complement(29727..29939)
/color=7
/locus_tag="C83_00291"
/note="helix-turn-helix domain-containing protein [transcriptional regulator], protein motif:Pfam:PF12645.1"
/product="helix-turn-helix domain-containing protein [transcriptional regulator]"
/"ID=C83_00291
/"

/protein_id="gnl|SC|C83_00291"
/codon_start=1
CDS complement(29932..30354)
/color=7
/locus_tag="C83_00292"
/note="RNA polymerase sigma factor Sigma-70 region 4,
protein motif:TIGRFAMs:TIGR03001"
/product="RNA polymerase sigma factor Sigma-70 region 4"
/"ID=C83_00292
/"
/protein_id="gnl|SC|C83_00292"
/codon_start=1
CDS complement(30526..31293)
/color=8
/locus_tag="C83_00293"
/eC_number=2.1.1.197
/note="Malonyl-CoA O-methyltransferase BioC, similar to AA
sequence:UniProtKB:P36571 protein motif:CLUSTERS:PRK08317
protein motif:Cdd:COG4106 protein motif:TIGRFAMs:TIGR02072
protein motif:Pfam:PF12847.1"
/product="Malonyl-CoA O-methyltransferase BioC
[methyltransferase]"
/"ID=C83_00293
/protein_id="gnl|SC|C83_00293""""
/gene="bioC"
/codon_start=1
CDS complement(31344..33263)
/color=2
/locus_tag="C83_00294"
/note="Tetracycline resistance protein tetM from
transposon Tn916, similar to AA sequence:UniProtKB:P21598
protein motif:CLUSTERS:PRK13351 protein motif:Cdd:COG0050
protein motif:TIGRFAMs:TIGR00484 protein
motif:Pfam:PF00009.21"
/product="Tetracycline resistance protein tetM from
transposon Tn916 [antibiotic resistance]"
/"ID=C83_00294
/"
/protein_id="gnl|SC|C83_00294"
/gene="tetM_2"
/codon_start=1
CDS complement(33904..34044)
/color=4
/locus_tag="C83_00295"

/note="Maff2 family  conjugal transfer protein, protein
motif:Pfam:PF12750.1"
/product="Maff2 family  conjugal transfer protein"
/"ID=C83_00295
/"
/protein_id="gnl|SC|C83_00295"
/codon_start=1
CDS complement(34223..34411)
/color=8
/locus_tag="C83_00296"
/note="PHP plymerase and histidinol phosphatase
domian-containing protein, ab initio
prediction:Prodigal:2.60"
/product="PHP %28plymerase and histidinol phosphatase%29
domian-containing protein"
/"ID=C83_00296
/"
/protein_id="gnl|SC|C83_00296"
/codon_start=1
CDS complement(34420..34947)
/ID=C83_00297
/color=8
/locus_tag="C83_00297"
/note="Fic Filamentation induced by cAMP family protein
[post-translational modification]"
/product="Fic %28Filamentation induced by cAMP%29 family
protein [post-translational modification]"
/protein_id="gnl|SC|C83_00297"
/codon_start=1
CDS complement(34962..35186)
/ID=C83_00298
/color=4
/locus_tag="C83_00298"
/note="Type IV secretory pathway VirD4 components
[coupling protein]"
/product="Type IV secretory pathway VirD4 components
[coupling protein]"
/protein_id="gnl|SC|C83_00298"
/codon_start=1
CDS complement(35230..36708)
/color=4
/locus_tag="C83_00299"
/note="Type IV secretion system protein TraG  VirD4
component [coupling protein], protein

motif:CLUSTERS:PRK13876 protein motif:Cdd:COG3505 protein
motif:TIGRFAMs:TIGR02759 protein motif:Pfam:PF02534.8"
/product="Type IV secretion system protein TraG  VirD4
component [coupling protein]"
/"ID=C83_00299
/"
/protein_id="gnl|SC|C83_00299"
/gene="traG"
/codon_start=1
CDS complement(36705..37196)
/color=1
/locus_tag="C83_00300"
/note="PcfB family protein, protein motif:Pfam:PF12687.1"
/product="PcfB family protein"
/"ID=C83_00300
/"
/protein_id="gnl|SC|C83_00300"
/codon_start=1
CDS complement(37231..37578)
/ID=C83_00301
/color=1
/locus_tag="C83_00301"
/note="hypothetical protein"
/product="hypothetical protein"
/protein_id="gnl|SC|C83_00301"
/codon_start=1
CDS complement(37603..39024)
/color=7
/locus_tag="C83_00302"
/note="Transcriptional regulator  AAA family ATPase,
protein motif:Pfam:PF12957.1"
/product="Transcriptional regulator  AAA family ATPase
[transcriptional regulator]"
/"ID=C83_00302
/"
/protein_id="gnl|SC|C83_00302"
/codon_start=1
CDS complement(39028..39627)
/ID=C83_00303
/color=1
/locus_tag="C83_00303"
/note="hypothetical protein"
/product="hypothetical protein"
/protein_id="gnl|SC|C83_00303"

/codon_start=1
CDS complement(39716..40642)
/color=10
/locus_tag="C83_00304"
/note="ParB/RepB/Spo0J family partition protein  ParB-like
nuclease domain [partitioning system], similar to AA
sequence:UniProtKB:Q83AH2 protein motif:CLUSTERS:PRK13832
protein motif:TIGRFAMs:TIGR00180 protein
motif:Pfam:PF02195.12"
/product="ParB/RepB/Spo0J family partition protein
ParB-like nuclease domain [partitioning system]"
/"ID=C83_00304
/"
/protein_id="gnl|SC|C83_00304"
/gene="parB"
/codon_start=1
CDS complement(40599..41423)
/color=10
/locus_tag="C83_00305"
/note="plasmid-partitioning protein A [partitioning
system], similar to AA sequence:UniProtKB:P37522 protein
motif:CLUSTERS:PRK13869 protein motif:Cdd:COG2894 protein
motif:TIGRFAMs:TIGR03453 protein motif:Pfam:PF01656.17"
/product="plasmid-partitioning protein A [partitioning
system]"
/"ID=C83_00305
/"
/protein_id="gnl|SC|C83_00305"
/gene="parA"
/codon_start=1
CDS complement(41436..42329)
/color=10
/locus_tag="C83_00306"
/note="Replication initiation protein RepA [replication],
protein motif:Pfam:PF06970.5"
/product="Replication initiation protein RepA
[replication]"
/"ID=C83_00306
/"
/gene="repA"
/protein_id="gnl|SC|C83_00306"
/codon_start=1
CDS complement(42349..42573)
/ID=C83_00307

/color=1
/locus_tag="C83_00307"
/note="hypothetical protein"
/product="hypothetical protein"
/protein_id="gnl|SC|C83_00307"
/codon_start=1
CDS complement(42566..42868)
/ID=C83_00308
/color=1
/locus_tag="C83_00308"
/note="hypothetical protein"
/product="hypothetical protein"
/protein_id="gnl|SC|C83_00308"
/codon_start=1
CDS complement(42868..43038)
/ID=C83_00309
/color=1
/locus_tag="C83_00309"
/note="hypothetical protein"
/product="hypothetical protein"
/protein_id="gnl|SC|C83_00309"
/codon_start=1
CDS 43106..44287
/color=9
/locus_tag="C83_00310"
/note="ISL3 family transposase [DDE recombinase], protein motif:Cdd:COG3464 protein motif:Pfam:PF01610.11"
/product="ISL3 family transposase [DDE recombinase]"
/"ID=C83_00310
/"
/protein_id="gnl|SC|C83_00310"
/codon_start=1
CDS complement(44470..45213)
/color=1
/locus_tag="C83_00311"
/note="DUF5131 family protein, protein motif:Cdd:COG4422 protein motif:Pfam:PF07505.5"
/product="DUF5131 family protein"
/"ID=C83_00311
/"
/protein_id="gnl|SC|C83_00311"
/codon_start=1
CDS complement(46022..46327)
/ID=C83_00312

/color=1
/locus_tag="C83_00312"
/note="hypothetical protein"
/product="hypothetical protein"
/protein_id="gnl|SC|C83_00312"
/codon_start=1
CDS complement(46413..47786)
/color=9
/locus_tag="C83_00313"
/note="Integrase  Tyrosine recombinase [recombinase],
similar to AA sequence:UniProtKB:P22886 protein
motif:CLUSTERS:PRK00236 protein motif:Cdd:COG4974 protein
motif:TIGRFAMs:TIGR02224 protein motif:Pfam:PF00589.16"
/product="Integrase  Tyrosine recombinase [recombinase]"
/"ID=C83_00313
/"
/protein_id="gnl|SC|C83_00313"
/gene="Int-Tn"
/codon_start=1
CDS complement(47791..48414)
/color=9
/locus_tag="C83_00314"
/note="DNA binding domain  excisionase family
Helix-turn-helix domain [excisionase], protein
motif:Cdd:COG3311 protein motif:TIGRFAMs:TIGR01764 protein
motif:Pfam:PF12728.1"
/product="DNA binding domain  excisionase family
Helix-turn-helix domain [excisionase]"
/"ID=C83_00314
/"
/protein_id="gnl|SC|C83_00314"
/codon_start=1
CDS complement(48532..48768)
/color=9
/locus_tag="C83_00315"
/note="DNA binding domain  excisionase family
Helix-turn-helix domain [excisionase], protein
motif:TIGRFAMs:TIGR01764 protein motif:Pfam:PF12728.1"
/product="DNA binding domain  excisionase family
Helix-turn-helix domain [excisionase]"
/"ID=C83_00315
/"
/protein_id="gnl|SC|C83_00315"
/codon_start=1

CDS complement(48775..49182)
/color=8
/locus_tag="C83_00316"
/eC_number=3.1.-.-
/note="Type II toxin-antitoxin system PemK/MazF family
toxin MazF, similar to AA sequence:UniProtKB:P96622
protein motif:CLUSTERS:PRK09907 protein
motif:Pfam:PF02452.11"
/product="Type II toxin-antitoxin system PemK/MazF family
toxin MazF [toxin-antitoxin]"
/"ID=C83_00316
/protein_id="gnl|SC|C83_00316""""
/gene="mazF"
/codon_start=1
CDS complement(49697..49912)
/color=8
/locus_tag="C83_00317"
/note="Antitoxin PezA XRE family HTH domain
[toxin-antitoxin], similar to AA sequence:UniProtKB:Q97QZ2
protein motif:CLUSTERS:PRK08154 protein motif:Cdd:COG3800
protein motif:TIGRFAMs:TIGR03070 protein
motif:Pfam:PF12844.1"
/product="Antitoxin PezA XRE family HTH domain
[toxin-antitoxin]"
/"ID=C83_00317
/"
/protein_id="gnl|SC|C83_00317"
/gene="pezA"
/codon_start=1
CDS complement(50264..50827)
/ID=C83_00318
/color=7
/locus_tag="C83_00318"
/note="Phage transcriptional regulator ArpU family
[transcriptional regulator]"
/product="Phage transcriptional regulator ArpU family
[transcriptional regulator]"
/protein_id="gnl|SC|C83_00318"
/codon_start=1
CDS 51114..51563
/color=7
/locus_tag="C83_00319"
/note="helix-turn-helix HTH transcriptional regulator
containing a XRE family HTH domain [transcriptional

regulator], protein motif:CLUSTERS:PRK09706 protein
motif:Pfam:PF12844.1"
/product="helix-turn-helix %28HTH%29 transcriptional
regulator containing a XRE family HTH domain
[transcriptional regulator]"
/"ID=C83_00319
/"
/protein_id="gnl|SC|C83_00319"
/codon_start=1
CDS 51615..52169
/color=8
/locus_tag="C83_00320"
/note="ImmA/IrrE family metallo-endopepidase, similar to
AA sequence:UniProtKB:P96630 protein motif:Pfam:PF06114.7"
/product="ImmA/IrrE family metallo-endopepidase
[pepidase]"
/"ID=C83_00320
/protein_id="gnl|SC|C83_00320""""
/gene="immA"
/codon_start=1
CDS complement(52314..53285)
/color=8
/locus_tag="C83_00321"
/note="Alpha/beta hydrolase [hydrolase], similar to AA
sequence:UniProtKB:Q01609 protein motif:Cdd:COG4925
protein motif:Pfam:PF12695.1"
/product="Alpha/beta hydrolase [hydrolase]"
/"ID=C83_00321
/"
/protein_id="gnl|SC|C83_00321"
/codon_start=1
CDS complement(53288..53899)
/color=8
/locus_tag="C83_00322"
/note="4-carboxymuconolactone decarboxylase
[carboxy-lyase], protein motif:TIGRFAMs:TIGR02425 protein
motif:Pfam:PF02627.1"
/product="4-carboxymuconolactone decarboxylase
[carboxy-lyase]"
/"ID=C83_00322
/"
/protein_id="gnl|SC|C83_00322"
/gene="pcaC"
/codon_start=1

CDS complement(53951..54052)
/ID=C83_00323
/color=8
/locus_tag="C83_00323"
/note="carboxymuconolactone decarboxylase family protein [carboxyl-lyase]"
/product="carboxymuconolactone decarboxylase family protein [carboxyl-lyase]"
/protein_id="gnl|SC|C83_00323"
/codon_start=1
CDS complement(54056..54595)
/color=8
/locus_tag="C83_00324"
/note="Flavodoxin family protein [electron-transfer], similar to AA sequence:UniProtKB:Q188R7 protein motif:CLUSTERS:PRK03767 protein motif:Cdd:COG0431 protein motif:TIGRFAMs:TIGR03567 protein motif:Pfam:PF03358.9"
/product="Flavodoxin family protein [electron-transfer]"
/"ID=C83_00324
/"
/protein_id="gnl|SC|C83_00324"
/codon_start=1
CDS 54702..55139
/color=7
/locus_tag="C83_00325"
/note="HTH-type transcriptional regulator AdhR  response to aldehyde stress [transcriptional regulator], similar to AA sequence:UniProtKB:O06008 protein motif:CLUSTERS:PRK09514 protein motif:Cdd:COG4978 protein motif:TIGRFAMs:TIGR02043 protein motif:Pfam:PF00376.17"
/product="HTH-type transcriptional regulator AdhR response to aldehyde stress [transcriptional regulator]"
/"ID=C83_00325
/"
/protein_id="gnl|SC|C83_00325"
/gene="adhR"
/codon_start=1
CDS complement(55272..56690)
/ID=C83_00326
/color=1
/locus_tag="C83_00326"
/note="hypothetical protein"
/product="hypothetical protein"
/protein_id="gnl|SC|C83_00326"

/codon_start=1
CDS complement(56842..57870)
/ID=C83_00327
/color=1
/locus_tag="C83_00327"
/note="hypothetical protein"
/product="hypothetical protein"
/protein_id="gnl|SC|C83_00327"
/codon_start=1
CDS complement(57909..58256)
/color=1
/locus_tag="C83_00328"
/note="MmcQ/YjbR family DNA-binding protein, protein
motif:CLUSTERS:PRK10250 protein motif:Cdd:COG2315 protein
motif:Pfam:PF04237.7"
/product="MmcQ/YjbR family DNA-binding protein"
/"ID=C83_00328
/"
/protein_id="gnl|SC|C83_00328"
/codon_start=1
CDS complement(58295..61357)
/color=8
/locus_tag="C83_00329"
/eC_number=3.1.21.3
/note="Type 1 restriction endonuclease subunit R, similar
to AA sequence:UniProtKB:Q7A801 protein
motif:TIGRFAMs:TIGR00348 protein motif:Pfam:PF11867.2"
/product="Type 1 restriction endonuclease subunit R
[restriction modification]"
/"ID=C83_00329
/protein_id="gnl|SC|C83_00329""""
/gene="hsdR"
/codon_start=1
CDS complement(61471..62652)
/color=8
/locus_tag="C83_00330"
/note="Type I restriction enzyme EcoKI subunit S
[restriction modification], protein
motif:CLUSTERS:PRK09737 protein motif:Cdd:COG1708 protein
motif:Pfam:PF01420.13"
/product="Type I restriction enzyme EcoKI subunit S
[restriction modification]"
/"ID=C83_00330
/"

/gene="hsdS"
/protein_id="gnl|SC|C83_00330"
/codon_start=1
CDS complement(62642..64147)
/color=8
/locus_tag="C83_00331"
/eC_number=2.1.1.72
/note="Type I restriction enzyme EcoKI subunit M, similar to AA sequence:UniProtKB:Q89Z59 protein motif:Cdd:COG0286 protein motif:TIGRFAMs:TIGR00497 protein motif:Pfam:PF02384.10"
/product="Type I restriction enzyme EcoKI subunit M [restriction modification]"
/"ID=C83_00331
/protein_id="gnl|SC|C83_00331"""
/gene="hsdM"
/codon_start=1
CDS complement(64236..64448)
/color=7
/locus_tag="C83_00332"
/note="helix-turn-helix transcriptional regulator containing a XRE family HTH domain [transcriptional regulator], protein motif:Cdd:COG3655 protein motif:Pfam:PF01381.16"
/product="helix-turn-helix transcriptional regulator containing a XRE family HTH domain [transcriptional regulator]"
/"ID=C83_00332
/"
/protein_id="gnl|SC|C83_00332"
/codon_start=1
CDS complement(64829..66502)
/color=9
/locus_tag="C83_00333"
/note="Serin recombinase [recombinase], protein motif:Pfam:PF00239.15"
/product="Serin recombinase [recombinase]"
/"ID=C83_00333
/"
/protein_id="gnl|SC|C83_00333"
/codon_start=1
CDS complement(66580..66753)
/ID=C83_00334
/color=9

/locus_tag="C83_00334"
/note="Transposon-encoded protein TnpW"
/product="Transposon-encoded protein TnpW"
/gene="tnpW"
/protein_id="gnl|SC|C83_00334"
/codon_start=1
CDS complement(67212..67454)
/color=7
/locus_tag="C83_00335"
/note="helix-turn-helix domain-containing protein
[transcriptional regulator], protein motif:Pfam:PF12645.1"
/product="helix-turn-helix domain-containing protein
[transcriptional regulator]"
/"ID=C83_00335
/"
/protein_id="gnl|SC|C83_00335"
/codon_start=1
CDS complement(67438..67875)
/color=7
/locus_tag="C83_00336"
/note="RNA polymerase sigma factor Sigma-70 region 4,
protein motif:CLUSTERS:PRK11924 protein
motif:TIGRFAMs:TIGR02937 protein motif:Pfam:PF08281.6"
/product="RNA polymerase sigma factor Sigma-70 region 4"
/"ID=C83_00336
/"
/protein_id="gnl|SC|C83_00336"
/codon_start=1
CDS complement(68345..69331)
/color=8
/locus_tag="C83_00337"
/eC_number=2.1.1.192
/note="23S rRNA adenine-C-methyltransferase RlmN, similar
to AA sequence:UniProtKB:Q7A600 protein
motif:CLUSTERS:PRK14469 protein motif:TIGRFAMs:TIGR00048
protein motif:Pfam:PF04055.15"
/product="23S rRNA
%28adenine%282503%29-C%282%29%29-methyltransferase RlmN
[methyltransferase]"
/"ID=C83_00337
/protein_id="gnl|SC|C83_00337""""
/gene="rlmN"
/codon_start=1
CDS 69583..70830

/color=9
/locus_tag="C83_00338"
/note="ISL3 family transposase, protein motif:Cdd:COG3464
protein motif:Pfam:PF01610.11"
/product="ISL3 family transposase"
/"ID=C83_00338
/"
/protein_id="gnl|SC|C83_00338"
/codon_start=1
CDS complement(70861..72081)
/color=2
/locus_tag="C83_00339"
/note="Macrolide efflux protein A Major Facilitator
Superfamily [antibiotic resistance], similar to AA
sequence:UniProtKB:P64783 protein motif:CLUSTERS:PRK10489
protein motif:Cdd:COG2814 protein motif:TIGRFAMs:TIGR00900
protein motif:Pfam:PF07690.10"
/product="Macrolide efflux protein A Major Facilitator
Superfamily [antibiotic resistance]"
/"ID=C83_00339
/"
/gene="mefA"
/protein_id="gnl|SC|C83_00339"
/codon_start=1
CDS complement(73086..73352)
/ID=C83_00340
/color=4
/locus_tag="C83_00340"
/note="Conjugatice transposon membrane protein"
/product="Conjugatice transposon membrane protein"
/protein_id="gnl|SC|C83_00340"
/codon_start=1
CDS complement(73426..73641)
/color=4
/locus_tag="C83_00341"
/note="Conjugal transfer protein Maff2 family, protein
motif:Pfam:PF12750.1"
/product="Conjugal transfer protein Maff2 family"
/"ID=C83_00341
/"
/protein_id="gnl|SC|C83_00341"
/codon_start=1
CDS complement(73709..75109)
/color=4

/locus_tag="C83_00342"
/note="Type IV secretion system protein TraG  VirD4 component [coupling protein], protein motif:CLUSTERS:PRK13850 protein motif:Cdd:COG3505 protein motif:TIGRFAMs:TIGR02759 protein motif:Pfam:PF02534.8"
/product="Type IV secretion system protein TraG  VirD4 component [coupling protein]"
/"ID=C83_00342
/"
/gene="traG"
/protein_id="gnl|SC|C83_00342"
/codon_start=1
CDS complement(75288..77189)
/color=10
/locus_tag="C83_00343"
/note="Replication initiation protein RepA [replication], protein motif:CLUSTERS:PRK11823 protein motif:Cdd:COG0358 protein motif:TIGRFAMs:TIGR02237"
/product="Replication initiation protein RepA [replication]"
/"ID=C83_00343
/"
/gene="repA"
/protein_id="gnl|SC|C83_00343"
/codon_start=1
CDS complement(77205..78593)
/color=4
/locus_tag="C83_00344"
/note="MobA/MobL family protein  DNA strand transferase MobA [relaxase], similar to AA sequence:UniProtKB:P07112 protein motif:CLUSTERS:PRK13889 protein motif:TIGRFAMs:TIGR02768 protein motif:Pfam:PF03389.9"
/product="MobA/MobL family protein  DNA strand transferase MobA [relaxase]"
/"ID=C83_00344
/"
/protein_id="gnl|SC|C83_00344"
/gene="mobA"
/codon_start=1
CDS complement(78945..79250)
/color=8
/locus_tag="C83_00345"
/note="DUF3847 domain-containing protein  AAA family ATPase, protein motif:Pfam:PF12958.1"

/product="DUF3847 domain-containing protein  AAA family
ATPase"
/"ID=C83_00345
/"
/protein_id="gnl|SC|C83_00345"
/codon_start=1
CDS complement(79372..80205)
/color=8
/locus_tag="C83_00346"
/eC_number=2.1.1.187
/note="23S rRNA methyltransferase
guanine-N-methyltransferase , similar to AA
sequence:UniProtKB:P36999 protein motif:CLUSTERS:PRK11088
protein motif:TIGRFAMs:TIGR02072 protein
motif:Pfam:PF08241.6"
/product="23S rRNA methyltransferase
%28guanine%28745%29-N%281%29%29-methyltransferase
[methyltransferase]"
/"ID=C83_00346
/protein_id="gnl|SC|C83_00346""""
/gene="rlmA"
/codon_start=1
CDS complement(80391..80687)
/ID=C83_00347
/color=8
/locus_tag="C83_00347"
/note="Molecular chaperone"
/product="Molecular chaperone"
/protein_id="gnl|SC|C83_00347"
/codon_start=1
CDS complement(80822..81013)
/ID=C83_00348
/color=9
/locus_tag="C83_00348"
/note="Transposon-encoded protein TnpW"
/product="Transposon-encoded protein TnpW"
/gene="tnpW"
/protein_id="gnl|SC|C83_00348"
/codon_start=1
CDS complement(81092..81943)
/ID=C83_00349
/color=9
/locus_tag="C83_00349"
/note="Site-specific recombinase  DUF4368

domain-containing protein [recombinase]"
/product="Site-specific recombinase  DUF4368
domain-containing protein [recombinase]"
/protein_id="gnl|SC|C83_00349"
/codon_start=1
CDS complement(81928..82941)
/color=9
/locus_tag="C83_00350"
/note="Serine recombinase [recombinase], protein
motif:Pfam:PF07508.7"
/product="Serine recombinase [recombinase]"
/"ID=C83_00350
/"
/protein_id="gnl|SC|C83_00350"
/codon_start=1
CDS complement(82988..83296)
/ID=C83_00351
/color=4
/locus_tag="C83_00351"
/note="Type IV secretion system protein TraG  VirD4
component [coupling protein]"
/product="Type IV secretion system protein TraG  VirD4
component [coupling protein]"
/protein_id="gnl|SC|C83_00351"
/gene="traG"
/codon_start=1
CDS complement(83293..83766)
/color=1
/locus_tag="C83_00352"
/note="PcfB family protein  DUF3801 domain-containing
protein, protein motif:Pfam:PF12687.1"
/product="PcfB family protein  DUF3801 domain-containing
protein"
/"ID=C83_00352
/"
/protein_id="gnl|SC|C83_00352"
/codon_start=1
CDS complement(83803..84231)
/ID=C83_00353
/color=10
/locus_tag="C83_00353"
/note="Phage replication protein  Helix-turn-helix
domain-containing protein [replication]"
/product="Phage replication protein  Helix-turn-helix

domain-containing protein [replication]"
/protein_id="gnl|SC|C83_00353"
/codon_start=1
CDS complement(84342..85589)
/color=9
/locus_tag="C83_00354"
/note="ISL3 family transposase [DDE recombinase], protein motif:Cdd:COG3464 protein motif:Pfam:PF01610.11"
/product="ISL3 family transposase [DDE recombinase]"
/"ID=C83_00354
/"
/protein_id="gnl|SC|C83_00354"
/codon_start=1
CDS complement(85628..86089)
/ID=C83_00355
/color=9
/locus_tag="C83_00355"
/note="Conjugative transposon protein helix-turn-helix domain-containing protein"
/product="Conjugative transposon protein helix-turn-helix domain-containing protein"
/protein_id="gnl|SC|C83_00355"
/codon_start=1
CDS complement(86218..86436)
/ID=C83_00356
/color=7
/locus_tag="C83_00356"
/note="Helix-turn-helix domain-containing protein [transcriptional regulator]"
/product="Helix-turn-helix domain-containing protein [transcriptional regulator]"
/protein_id="gnl|SC|C83_00356"
/codon_start=1
CDS complement(86536..87927)
/color=8
/locus_tag="C83_00357"
/note="Group II intron-encoded protein LtrA [reverse transcriptase], similar to AA sequence:UniProtKB:P0A3U1 protein motif:Cdd:COG3344 protein motif:Pfam:PF00078.21"
/product="Group II intron-encoded protein LtrA [reverse transcriptase]"
/"ID=C83_00357
/"
/protein_id="gnl|SC|C83_00357"

/gene="ltrA"
/codon_start=1
ncRNA complement(87975..88046)
/ID=C83_00358
/color=1
/locus_tag="C83_00358"
/product="group-II-D1D4-1"
ncRNA complement(88086..88279)
/ID=C83_00359
/color=1
/locus_tag="C83_00359"
/product="group-II-D1D4-3"
CDS complement(88556..88876)
/color=7
/locus_tag="C83_00360"
/note="Transcriptional regulator containing a XRE family HTH domain [transcriptional regulator], protein motif:Pfam:PF12844.1"
/product="Transcriptional regulator containing a XRE family HTH domain [transcriptional regulator]"
/"ID=C83_00360
/"
/protein_id="gnl|SC|C83_00360"
/codon_start=1
CDS 89041..89514
/ID=C83_00361
/color=1
/locus_tag="C83_00361"
/note="DUF1273 domain-containing protein"
/product="DUF1273 domain-containing protein"
/protein_id="gnl|SC|C83_00361"
/codon_start=1
CDS 89547..90068
/ID=C83_00362
/color=1
/locus_tag="C83_00362"
/note="DUF4367 domain-containing protein"
/product="DUF4367 domain-containing protein"
/protein_id="gnl|SC|C83_00362"
/codon_start=1
CDS complement(90136..90483)
/color=7
/locus_tag="C83_00363"
/note="helix-turn-helix transcriptional regulator

containing a HxlR family HTH domain [transcriptional
regulator], similar to AA sequence:UniProtKB:P37486
protein motif:Pfam:PF01638.11"
/product="helix-turn-helix transcriptional regulator
containing a HxlR family HTH domain [transcriptional
regulator]"
/"ID=C83_00363
/"
/protein_id="gnl|SC|C83_00363"
/codon_start=1
CDS 90625..91020
/color=8
/locus_tag="C83_00364"
/note="Pyridoxamine 5'-phosphate oxidase family protein
[oxidase], protein motif:Cdd:COG5015 protein
motif:TIGRFAMs:TIGR03618 protein motif:Pfam:PF01243.14"
/product="Pyridoxamine 5'-phosphate oxidase family protein
[oxidase]"
/"ID=C83_00364
/"
/protein_id="gnl|SC|C83_00364"
/codon_start=1
ncRNA 91035..91149
/ID=C83_00365
/color=1
/locus_tag="C83_00365"
/product="FMN"
CDS 91227..91826
/color=8
/locus_tag="C83_00366"
/note="Riboflavin ECF transporter S component RibU
[transporter], similar to AA sequence:UniProtKB:E5QVT2
protein motif:Cdd:COG4720 protein motif:Pfam:PF12822.1"
/product="Riboflavin ECF transporter S component RibU
[transporter]"
/"ID=C83_00366
/"
/protein_id="gnl|SC|C83_00366"
/gene="ribU"
/codon_start=1
CDS 91842..92465
/color=8
/locus_tag="C83_00367"
/note="4Fe-4S dicluster domain-containing protein

[electron transfer], similar to AA
sequence:UniProtKB:P00195 protein motif:CLUSTERS:PRK07118
protein motif:Cdd:COG5015 protein motif:TIGRFAMs:TIGR04105
protein motif:Pfam:PF00037.21"
/product="4Fe-4S dicluster domain-containing protein
[electron transfer]"
/"ID=C83_00367
/"
/protein_id="gnl|SC|C83_00367"
/codon_start=1
CDS 92470..93249
/color=8
/locus_tag="C83_00368"
/eC_number=3.5.1.-
/note="O-acetyl-ADP-ribose deacetylase, similar to AA
sequence:UniProtKB:P0A8D6 protein motif:CLUSTERS:PRK04143
protein motif:Pfam:PF01661.15"
/product="O-acetyl-ADP-ribose deacetylase [hydrolase]"
/"ID=C83_00368
/protein_id="gnl|SC|C83_00368""""
/gene="ymdB"
/codon_start=1
CDS 93194..94069
/ID=C83_00369
/color=8
/locus_tag="C83_00369"
/note="Silent information regulator family NAD-dependent
deacetylase Sir2 [deacetylase]"
/product="Silent information regulator family
NAD-dependent deacetylase Sir2 [deacetylase]"
/protein_id="gnl|SC|C83_00369"
/gene="sir2"
/codon_start=1
CDS complement(94141..95541)
/color=4
/locus_tag="C83_00370"
/note="Relaxase/Mobilisation nuclease domain [relaxase],
protein motif:Pfam:PF03432.8"
/product="Relaxase/Mobilisation nuclease domain
[relaxase]"
/"ID=C83_00370
/"
/protein_id="gnl|SC|C83_00370"
/codon_start=1

CDS complement(95596..95862)
/ID=C83_00371
/color=1
/locus_tag="C83_00371"
/note="protein BART-1"
/product="protein BART-1"
/protein_id="gnl|SC|C83_00371"
/codon_start=1
CDS complement(95877..96161)
/ID=C83_00372
/color=1
/locus_tag="C83_00372"
/note="hypothetical protein"
/product="hypothetical protein"
/protein_id="gnl|SC|C83_00372"
/codon_start=1
CDS complement(96383..97816)
/color=9
/locus_tag="C83_00373"
/note="IS5 family transposase [DDE recombinase], protein
motif:Pfam:PF05598.5"
/product="IS5 family transposase [DDE recombinase]"
/"ID=C83_00373
/"
/protein_id="gnl|SC|C83_00373"
/codon_start=1
CDS complement(97940..99400)
/color=8
/locus_tag="C83_00374"
/note="Lsa family ABC-F type ribosomal protection protein
[transporter], similar to AA sequence:UniProtKB:P45127
protein motif:CLUSTERS:PRK11819 protein motif:Cdd:COG4586
protein motif:TIGRFAMs:TIGR03719 protein
motif:Pfam:PF00005.21"
/product="Lsa family ABC-F type ribosomal protection
protein [transporter]"
/"ID=C83_00374
/"
/protein_id="gnl|SC|C83_00374"
/codon_start=1
CDS complement(100062..100394)
/ID=C83_00375
/color=4
/locus_tag="C83_00375"

/note="plasmid mobilization relaxosome protein MobC [relaxosome]"
/product="plasmid mobilization relaxosome protein MobC [relaxosome]"
/gene="mobC"
/protein_id="gnl|SC|C83_00375"
/codon_start=1
CDS complement(100394..108523)
/color=10
/locus_tag="C83_00376"
/note="DEAD/DEAH box helicase family protein  Superfamily II DNA/RNA helicase  SNF2 family [helicase], protein motif:Cdd:COG4646 protein motif:Pfam:PF00176.17"
/product="DEAD/DEAH box helicase family protein Superfamily II DNA/RNA helicase  SNF2 family [helicase]"
/"ID=C83_00376
/"
/protein_id="gnl|SC|C83_00376"
/codon_start=1
CDS complement(108540..108749)
/ID=C83_00377
/color=4
/locus_tag="C83_00377"
/note="Putative tranposon-transfer assisting protein %28TTRAP%29"
/product="Putative tranposon-transfer assisting protein %28TTRAP%29"
/protein_id="gnl|SC|C83_00377"
/codon_start=1
CDS complement(108746..109339)
/ID=C83_00378
/color=1
/locus_tag="C83_00378"
/note="DUF4316 domain-containing protein"
/product="DUF4316 domain-containing protein"
/protein_id="gnl|SC|C83_00378"
/codon_start=1
CDS complement(109351..109962)
/ID=C83_00379
/color=1
/locus_tag="C83_00379"
/note="DUF3848  DUF4316 domain-containing protein"
/product="DUF3848  DUF4316 domain-containing protein"
/protein_id="gnl|SC|C83_00379"

/codon_start=1
CDS complement(109959..112034)
/color=10
/locus_tag="C83_00380"
/eC_number=5.99.1.2
/note="DNA topoisomerase III, similar to AA sequence:UniProtKB:P14294 protein motif:CLUSTERS:PRK07726 protein motif:Cdd:COG0550 protein motif:TIGRFAMs:TIGR01056 protein motif:Pfam:PF01131.14"
/product="DNA topoisomerase III [topoisomerase]"
/"ID=C83_00380
/protein_id="gnl|SC|C83_00380"""
/gene="topB"
/codon_start=1
CDS complement(112103..113113)
/ID=C83_00381
/color=1
/locus_tag="C83_00381"
/note="DUF4238 domain-containing protein"
/product="DUF4238 domain-containing protein"
/protein_id="gnl|SC|C83_00381"
/codon_start=1
CDS complement(113242..114324)
/color=1
/locus_tag="C83_00382"
/note="DUF4366 domain-containing protein, protein motif:TIGRFAMs:TIGR04211 protein motif:Pfam:PF08239.5"
/product="DUF4366 domain-containing protein"
/"ID=C83_00382
/"
/protein_id="gnl|SC|C83_00382"
/codon_start=1
CDS complement(114314..114571)
/ID=C83_00383
/color=1
/locus_tag="C83_00383"
/note="DUF4315 domain-containing protein"
/product="DUF4315 domain-containing protein"
/protein_id="gnl|SC|C83_00383"
/codon_start=1
CDS complement(114585..116339)
/color=4
/locus_tag="C83_00384"
/eC_number=3.4.-.-

/note="NlpC/P60 family pepidase, similar to AA
sequence:UniProtKB:P21171 protein motif:CLUSTERS:PRK13914
protein motif:Pfam:PF00877.13"
/product="NlpC/P60 family pepidase [pepidase]"
/"ID=C83_00384
/"
/protein_id="gnl|SC|C83_00384"
/codon_start=1
CDS complement(116343..118730)
/color=4
/locus_tag="C83_00385"
/note="Type IV secretion system protein TraC VirB4
components [ATPase mating pair formation], protein
motif:Cdd:COG3451 protein motif:TIGRFAMs:TIGR02746 protein
motif:Pfam:PF12846.1"
/product="Type IV secretion system protein TraC VirB4
components [ATPase mating pair formation]"
/"ID=C83_00385
/"
/gene="traC"
/protein_id="gnl|SC|C83_00385"
/codon_start=1
CDS complement(118690..119124)
/color=4
/locus_tag="C83_00386"
/note="PrgI family protein Type IV secretion system,
protein motif:Pfam:PF12666.1"
/product="PrgI family protein Type IV secretion system"
/"ID=C83_00386
/"
/protein_id="gnl|SC|C83_00386"
/codon_start=1
CDS complement(119141..120007)
/ID=C83_00387
/color=4
/locus_tag="C83_00387"
/note="Membrane protein trbL/VirB6 plasmid conjugal
transfer family protein"
/product="Membrane protein trbL/VirB6 plasmid conjugal
transfer family protein"
/protein_id="gnl|SC|C83_00387"
/codon_start=1
CDS complement(120033..120248)
/color=4

/locus_tag="C83_00388"
/note="Conugal transfer protein  Maff2 family, protein motif:Pfam:PF12750.1"
/product="Conugal transfer protein  Maff2 family"
/"ID=C83_00388
/"
/protein_id="gnl|SC|C83_00388"
/codon_start=1
CDS complement(120392..120703)
/ID=C83_00389
/color=7
/locus_tag="C83_00389"
/note="helix-turn-helix domain-containing protein [transcriptional regulator]"
/product="helix-turn-helix domain-containing protein [transcriptional regulator]"
/protein_id="gnl|SC|C83_00389"
/codon_start=1
CDS 121129..122430
/color=9
/locus_tag="C83_00390"
/note="Transposase IS116/IS110/IS902 family [DDE recombinase], protein motif:Pfam:PF02371.10"
/product="Transposase IS116/IS110/IS902 family [DDE recombinase]"
/"ID=C83_00390
/"
/protein_id="gnl|SC|C83_00390"
/codon_start=1
CDS complement(122539..124290)
/color=4
/locus_tag="C83_00391"
/note="Type IV secretion system protein TraG  VirD4 component [coupling protein], protein motif:CLUSTERS:PRK13876 protein motif:Cdd:COG3505 protein motif:TIGRFAMs:TIGR02759 protein motif:Pfam:PF02534.8"
/product="Type IV secretion system protein TraG  VirD4 component [coupling protein]"
/"ID=C83_00391
/"
/gene="traG"
/protein_id="gnl|SC|C83_00391"
/codon_start=1
CDS complement(124290..124775)

/color=1
/locus_tag="C83_00392"
/note="PcfB family protein  DUF3801 domain-containing protein, protein motif:Pfam:PF12687.1"
/product="PcfB family protein  DUF3801 domain-containing protein"
/"ID=C83_00392
/"
/protein_id="gnl|SC|C83_00392"
/codon_start=1
CDS complement(124810..125121)
/ID=C83_00393
/color=1
/locus_tag="C83_00393"
/note="hypothetical protein"
/product="hypothetical protein"
/protein_id="gnl|SC|C83_00393"
/codon_start=1
CDS complement(125131..125772)
/color=1
/locus_tag="C83_00394"
/note="DUF3846 domain-containing protein, protein motif:Pfam:PF12957.1"
/product="DUF3846 domain-containing protein"
/"ID=C83_00394
/"
/protein_id="gnl|SC|C83_00394"
/codon_start=1
CDS complement(125827..126342)
/ID=C83_00395
/color=1
/locus_tag="C83_00395"
/note="hypothetical protein"
/product="hypothetical protein"
/protein_id="gnl|SC|C83_00395"
/codon_start=1
CDS complement(126354..127280)
/color=10
/locus_tag="C83_00396"
/note="ParB/RepB/Spo0J family partition protein  ParB-like nuclease domain [partitioning system], similar to AA sequence:UniProtKB:Q83AH2 protein motif:CLUSTERS:PRK13832 protein motif:TIGRFAMs:TIGR00180 protein motif:Pfam:PF02195.12"

/product="ParB/RepB/Spo0J family partition protein
ParB-like nuclease domain [partitioning system]"
/"ID=C83_00396
/"
/protein_id="gnl|SC|C83_00396"
/gene="parB"
/codon_start=1
CDS complement(127237..128064)
/color=10
/locus_tag="C83_00397"
/note="plasmid-partitioning protein A [partitioning
system], similar to AA sequence:UniProtKB:P37522 protein
motif:CLUSTERS:PRK13869 protein motif:Cdd:COG2894 protein
motif:TIGRFAMs:TIGR03453 protein motif:Pfam:PF01656.17"
/product="plasmid-partitioning protein A [partitioning
system]"
/"ID=C83_00397
/"
/protein_id="gnl|SC|C83_00397"
/gene="parA"
/codon_start=1
CDS complement(128106..129008)
/color=10
/locus_tag="C83_00398"
/note="Replication initiation protein RepA [replication],
protein motif:Pfam:PF06970.5"
/product="Replication initiation protein RepA
[replication]"
/"ID=C83_00398
/"
/gene="repA"
/protein_id="gnl|SC|C83_00398"
/codon_start=1
BASE COUNT 30470 a 32944 c 29545 g 36049 t
ORIGIN
1 gtgatttccg atccctttct ggatcccgcc accggagaga tcattgtctc gatcgtaaac
61 ccggtttttt ctgaatcagg cagcgagatt gccggtctga ccggtttaga tgtctattta
121 agcagtctgg gccaggtaat ctcatcgatc aaagtcgggg aacacggtta tcttgaactt
181 atctccaata attcggatta catatacagc gacgatccga gcgcatcggg caggaatgtt
241 tccgaactcg atatccccca ggagtataaa gataaggttc aaagcaatta taatggcgtt
301 ctcaacttca gttatagcgg catccggtat acctccatgt tcagaaacag cgcaaccacc
361 ggctggctgg ctgtcgccac gctcccgatt tctgaggtaa atgcgacacg caaccggctg
421 ctcaccatac tgggcgtgct gtccctcata attctggccg gacttacaac tgttatcgtg
481 tttattgtac accgtaccat gaaacctctt gccggtatca gcagaagcat ggaagatttc

541 tcacagggta atctgggcgt cgacattcct ctcggccgga cagatgatga aatcggcaga
601 atggccggca gcatccgttc gtctgttgcc gctctcaagg atatgattca taatgtcaca
661 cgcgtcctgg aagaaatttc ccggggcaat ctgaaactgt ccgtagacgg taattattta
721 ggagattttt cttttatagg ggatgccctg gagcagatta taaaatccct gaattatacg
781 ctttcccaaa tcagctcatc cgcccagcag gtagcctatg gctccgaaca gggggcctgc
841 ggcgcgcagt cgatggctca gggcgctacg gaacaggcgg cggcggcaga ggaactggcg
901 gcggtcatag aggacatctc ccagcagatt atatccaatg tcagcagcac ttcaaaagcc
961 aatatgagcg taactacggt ggtaaatgaa gccgagataa gcaaccgccg catgcaggag
1021 atgctgtcgg ctatgcagga catccgacag agctccaacg agataagtaa aattatcagc
1081 gcaattgaag atatttcctt ccagacaaat atattgtccc tcaatgcagc cgtcgaggct
1141 gcccgcgccg gtgaaaacgg cagaggtttc gccgtcattg ccgcagaaat caggaatctg
1201 gccgcaaaga gcgccgaagc gtcccataac acgacgtccc tcatcgccgc ttcgttaaaa
1261 gcagtggaac acggaacgcg tatttctggt gaaacggctg tctctctcaa gaatgtcctg
1321 gacggtatta aagaggtgga gatgattatt gagcagatat cttcagcttc cgacgatcag
1381 gcccgctccg tggaacaggt acatcagagt atcacgcaga tttccgatgt tgtccagctc
1441 aactcagcta ccgccgaaga gagcgcggcg gccagcgaag aactgtccgc acaggcgctg
1501 ctgttgaagg aattgattgg ctcgttccgg ctcaggaacg aaccctgata aaatagtaaa
1561 taatcttttt tactttacag ccattgattg cttattcatc agtcagatag ctattacata
1621 tttcaccgcc tgcggcgggt cgctgtggcg acattattcc ataccgccgc aggcattagc
1681 tcacttagca agtgtaacga gcttagtgaa caagccctct ttagagttgt ggtcctccga
1741 ggagcggttt taatttaaca ggagaacttt cctgttaaat taaaaaaccg aaaacctctc
1801 acatcaaggt tttcggcatt gtattcagtg ttcctgagcg gattcgaacc gctggccttc
1861 cgcttaggag aaccattgaa catttgtcat ctgtgttcac ctatgtacag gaaagtcgct
1921 atccgtcagg cttttttcgt attccgagtc gctccgtgtt cgtccaagtt caccgatgtc
1981 gttagatgtt caccggttca ttttagcagt tttttggatt tttctaaaaa ggggaacaag
2041 aacatttttt atctaccgcc aattttagca ctttttcgag taaaaatcca tctgcagtcg
2101 aaccgcaagc ctgaacagaa tatctcacat ctccctgaaa agtactcgct ggggaagatg
2161 ataaaaaaga gagtattgca ttctcggttc aaatgagatc cgagtttaca atactctcta
2221 tgtaagtgac gggccggtaa tccggccctt attttagttc aggctcttcg ccaggcttgt
2281 cagcagcgct gccatttccg gattactgag tatctttgcc agaaggtcag gatccacacc
2341 tgaaggggct gcttttgctt cggatccggt ttcagccggt gactcaatac tgtctgaacc
2401 tttcccgctg taaaaggctt tctcaaacaa tgctgcgttg ttcttacggt catcatcaag
2461 aatgtgggaa tattgatccg taaccatctg tgcctgtgca tggccggaat cgccctgaac
2521 cgatttgata tcgccgccgt tcagcttcag cttataagtg atgctggaat gacgaagact
2581 gtgaaagacg acctttggca gatcattctt ctcaatcagt tctgcaaaga gggccgtgat
2641 gcgggattgc tctgtcggca tgcccaagcc attggccatg accagatcga aatctgcgta
2701 ttcattgcca agggcttctt tcgtgaaatc ctgctcccgt ttccactcta ccagcatctc
2761 agcgacggtc ttcggaagaa agatcttgcg ggtactggtt gccgtctttg gtttcttcaa
2821 tacaaggacg gttttattgc tttcccgcag ttccggaaag acgcgaagca cgtccttctt
2881 tcccaaaacc ttcataacac tcttatcgac acgctgcagc tccttattca cataaacata
2941 ggcctttccg gcggcgatgc tttccggaga gatgtctacg caatcccagg tcaggccaag
3001 cagttcgcca attcgaagag aacaggcaaa tgccaggttg atggccagtt tcaggcgctc
3061 atcctcacat acctcgaggg catgaaacag agtaggagcg tcccagatct cgcgcttctt
3121 tgcttcagat tttggaaccg tggcatacat agcagggttc ttctccataa gctcccattt

3181 cacagcctga tgaaaggcgc tgcgcagaat gttatgtatc tttcttacgg tcggcggagc
3241 cacataaatc acatttttag agtttgcttt ctgagtgatt cggtgcacgg aaggagtctt
3301 cagaagctgc atataatact tctccagaac ccgtgctgtc acgtcactga gcttcatact
3361 gccgatgatc ggagatatgt aatgctcgat cagagccgta ttgcttccgt agacggacat
3421 ggaccatgtg gtcttgccat agagagcaac atactctttc agcagctcat ccaccgtctt
3481 gcactgatgg atcaccatag tgcctatttc ctgccgatat tcaatctcag ctttccggcg
3541 ctttgcatcg gccatggtct tgaagctttc ccatttctgc tttcgcttgc cggttgcttc
3601 atcgtcatag aggtaaacga cattgtaacg gttatttctc tgtacaatgg atgccattgt
3661 ctctcactcc tttcctagtt ctcgcgttcg agtttctgct gaaacaggaa ccacttgata
3721 tcatcacggg ggatcagata cttgagtccg tattttttgg cttcaatttc gccggtctga
3781 atcagcctgc gcacctcatt gtatgtaagg tcaagaagta tcgccgcctc ctgcagggta
3841 aatgctgcct ttgtttcgtc caccttgaga cgaggcttca ctgcttcctg ctgcattctc
3901 tcaatcgcca tgatttcctc cggtgatctg tcgcaaagtt tgcgatattt gctctgaccg
3961 gcataccagc gctcaaagct gtcctttgtt acgcgccgtc ggtctgcaat atagacaaac
4021 tcaaactgat ctttgtctct gccggttaat aggatattgt agacttcctt ccgggtgatc
4081 aaaagcagtc ttgccatttc cggcatcgtc atggatgccg cttccagctc tgcatctcgt
4141 tcccggtccg cttgggttcg atacttggtc tgcgtcttgt accatgcctc aaaaacatcc
4201 ttccggatac gcatccaggt atccacctca aaggtctcaa tgtgatcccg cttgatcaag
4261 gtataaaccg tgtcatcgga tacaccgagc agcgccgcca tctcctgaac cgagtaagaa
4321 tatgccctga gctctgcacc gggcggcgat ccatccacct tcttgtactt gatctgattt
4381 gcataccagt gctcaaagct gtcgatcaca actcgcatct tgcctgcgac aatgatcgtt
4441 tcaaagcact gcctatgtac cagccaataa gaatccgttt ttccaagccc caacatccgc
4501 cgcatctccg gaacggacat tgtagtcctt ttcttttcat ccatatcggg tcctccttat
4561 aaagtgatag ccaggtctta agacctggct acatcgtagc ggttatccgt tcgaattgtt
4621 aggatgtcgc ggaacggctc gacatccttg acaacttaca gcttgttgtc caaccactca
4681 tcgaagctct tctttgaaat tcgataggcg gatccgatct taaaccagcg aaattcctgc
4741 tgttccaaca gcttatatac cgtgccgcgg ctgatgccga ggatgatctg cagatcctca
4801 acggtatagc agcgcttctc tgtgatgggt attgcatttt ctgacggtaa ctcaaaagtc
4861 tcaattgctt ccatatgtca ttcctcgctt tccgttttca tctttgccgc aaattccaaa
4921 tcagggaaac tccacgcatt ccggaatggg caaggtatcg tccggatcca gatgaaggct
4981 tacgcggatg ctgtcgtcaa tgcgttccat gtgcttttgg ctcatgcgcc ccagatactt
5041 gattacctga cacttgttga tggtgacgat ctgttctgcc tcactcatgg aggtttcatt
5101 cagaccgcgc tgcttttcga gaacgcagtg tgtaggctgc cccagcttct tcaattccgt
5161 ggtaagcggc acgacgatca atgtcggacc gaagaaattg ccgctgttgt tctgaatatt
5221 gatgaccggt cgaacgccgc cctgaatatt tccgcgatgc tcgcccaggt ctgccagata
5281 gatatcaccg cgtctatagt tccaatcttt tctaagcatt tttctcctcc tcatataggt
5341 tctatgtaaa agggcgggat atcgcttccc acccaagaaa gccgctaagg gcttaatctc
5401 aataatatgt tatccccaca tttgtcctcg acacccctgg gaactggcag gcgctctgac
5461 ctgccgggaa tacattcagg cgtctggctg ctaccagatc catgggaatc taacccctcc
5521 gtggttctca ccgagctgcg tacagaagta tcattgtgct gtgatggatc aatgctgtat
5581 atcggaagca atccgatact tacaggccgg ttcacgctcc cgccttctaa tggcgatcgt
5641 tgcgttcatc cgctgatgtg gctccccatc acagggttgt gcctgaagaa tggtatgtaa
5701 ttttcaaggt tcctgtagag gagtcgcttg gctccttaaa gtgacgtttt tcacatcttt
5761 gatacttttc tttaccggct tgccaaagcc ggattcccaa aatgcccgcg taccgccttt

5821 ggatgaaaag cggcacttct ggacgcgcgg tagatttcga tcattcgtcc atagacgatg
5881 acgaaaatca caacaccgag tgccttgatg gtaaactgaa cgaccatgca ctgcaggaac
5941 agtgcgaaca gaggcccgat ttccatggcc tccagcgtat cctgcatggt ttctaaggct
6001 gcagcatcca ccgccgtatt cccggcaatg agcccgcctg cgttattgat cacatgctgc
6061 gtcacatcaa agactgccat cacaatattg aaggtgttgg tgatgagtgt cacggctaca
6121 aaggtcttaa agacccattt gaagaagatc caggtttcaa aattggcaag gttattgtgg
6181 tcaatgatca tctgaatcag ctcatagcag gcaatgaagg tcaagatcaa gcctgcaacc
6241 gggataatca ctgattcgga aacattccgg atcatgacga agacgcccgg cgagaaattc
6301 gccggagtcg tccccacctg cgtcgcaata tcgccaacct gagtgtttac cgaatcaaac
6361 agtcccgtca gctggttcat gatgccggaa acaagcagtt ccttcagcca ttcaccaatc
6421 tgatcgagta cgctgtccaa cgattaacac tcctttctta gccgaagagg ccggaaagca
6481 ggggaaccag agtgataccg atgagggcca cgccgccacc tgccatcagc tgcttcatac
6541 cctgagactt tgcttatgtg tgataaagaa gcaatagaag tgcaaaaaat tttgccgttc
6601 ctatccgaca cttggccatt gtgtcggata ggtcgggcgg ttattcgggc aagtcaatct
6661 tgccgacaaa gctgtaataa atctcaatgt cctgcctgcg ggtgccgttc tcatcatagc
6721 tgcactcatg caccacaatt ttctcgatca tttcccgcaa gagagtgggg gtcagttctt
6781 caaaggcaag gtgcttgcgg acaatgccca taaatttctc ggcgttgacg gtagctgcct
6841 gtgacttgtc cagttcggct tgcagggcgg cggctctctt tttcagctcc gcttgctcgg
6901 cttcgtagtc agccgacagt tccatgaaac gctcatcgct gattttgccg tttacattgt
6961 cctcatacag ccgcttgata atgcggctga tttcagaaat gcgttcctgc gcctgttcaa
7021 gctgcttgat ggctgcggcg gtctttcgct tgccgccgat ctcgttctgc tggacaagta
7081 gtttcacaaa ccggctctca tgcttggctg cgtattcggt cacttgccgg agatttgcca
7141 ggacaccagc ggtcaacaga tcggtgcgga taaagtgcgc cgtacagttg cgggtgcgct
7201 tcttgtagct gccgcagatg tagcagtcct gtttgcggtc tttgttctga taccgctgct
7261 gatacagcac atggccgcag tcggcgcaga acaaaatccc ggagaacagc cccacttcat
7321 cgtagcggtt cgggcgtttg cgctgtttgc gtaactcctg cacccgttcc catgtttcct
7381 tgtcgatgat cggctcatgg tggttctcga aaatggcctg cttctcgacg gggttctcta
7441 tgctgtgctt gaccttataa gaaggctttt ccgttttgaa gttcaccaga catccggtgt
7501 actctcggtt ttcgaggata tgaacgacgg tgttggtcgc ccatttgcac tcatagcctg
7561 ggtgataacg gcgggtgctg cctgtccgct gatattccag cgtccccggc gtggggattt
7621 gctgctccgt cagcatacgg gcaatcttgg tcgggccgtt cccggcaagg caaagctggt
7681 aaatctgctg caccaccggg gcggcttcct cgtctataat aaaattctcg tcctcgtcca
7741 tcacatagcc gtaaaccggc ttgctggtaa caggcttgcc gctcatgcct tttgactttt
7801 tcactgcctt gattttcttg ctcgtatctc tcaccagcca ttcattgaac agatttcgca
7861 gcggggtaaa atcgttgtcc atgccctcgc tggcactgtc cacattatcg ttgacagcga
7921 taaaacgcac accctttga gggaacagca tttccgtgta aaaccctacc tgcaagtagt
7981 ttcgccctaa ccggctcatg tccttgacga tgaccgtacc cactttcccg gcttcaatgt
8041 ccgcaagcat ggcttgaaat ccgggccgct ggaagttcgc gccggaaaaa ccgtcgtcgg
8101 tgtaccagcg cagattggta aagccgttct gttttgcgta ggtttcgagt atcctttttt
8161 ggttcgatat ggaattactc tcgccttgca attcatcctc gtgggacaat ctcggataaa
8221 gggcggtaat gaggttttgg gtggcttgtc ttaacataaa atcctccgtt tccgacagcc
8281 agccccacta ttccgtggtt tcattgtacc acggaacggc gggggctgta cagcggcaaa
8341 agcgttaaat ttgcttcttt acagcttttc aattttccga tttttatggt atgggttgtc
8401 gtcccatatt tgcagtagaa aagttcataa ctccgtggta tactctgatt taacaaaaaa

8461 atcagaagtt aaaggagaaa acatgaagta tacaatagat gaattaaccg cggccaaaag
8521 gcaaattgat tcaactctgc acaagctaag agaaacagta aaaacatttg aatcaaagga
8581 taattccgag cgttataaat cacaaatcac attggcaaag agaagaataa aagcctttga
8641 aattgcaaat tattttatag agaatgagat taagaattgc taaagcactt ccgattttcg
8701 ttccagcggt catgctccct ggcatggccg cttttccatg ccccggttca cgccggataa
8761 gtcggctcct gtgtagcagc ggcttccgct tccagtaccc gcgccatttt gtcggcggct
8821 gtggttgtgg tgtctttctt gaaatagccg gacacgacaa ggacggaatt gcccatgcgg
8881 atttccgtca cacagtcagg gcggcgggtg gtgcgggtgt cgtgctgctt gttatctgcc
8941 ataggcaata ctcctttcag tcggtaatca gtttcttcat gctctccatt ttccgcttgg
9001 cggtttcctg ccggaagttg tcgccggtaa agcgtaccgg gacgcacatg gaaagcaggc
9061 ggtcataaat ccgggaatgg gcggtgtcct ccgggttgtg caggtcgtcc agcgtaaggt
9121 tggtcgtgac gatcagcggc ttgccgctgc ggtaacggct gtcaatcaca ttgaacacct
9181 gttccagccc gtattccgtc ccgcgttcca tgccgaagtc gtcaaggatc agcagcggat
9241 aacgacaaag gcgggaaatg tactcgttgc gcccctcaaa gctggcggca aggtcattga
9301 ggataagagc aaagttcgtc atgcggacgg ggatttcttt ctccatgagg gcgtttgcga
9361 tacagcctgc aaggtagctt ttgccggtgc ctacgccgcc ccagaacagg cagccgatat
9421 tgtctgtccg catatcttcc caatgctcca cataccggcg ggcaagcccg gtctgcgggt
9481 tcctgccgtt gtcgttctcg aaagtccagt cccgcatggt ggggtcggta aagccccggc
9541 gtttcagttc ttccacggtg tcaaggtgtc tgcgccgctt ttcggcggct tcccgttcct
9601 cgcgggcggt tctctggcag tcgcactctg ccgggtggcg gtcgcgcccg aagatagcgg
9661 ccttatccgg cgcaaaatag gcttctttcg gcttgcggca cttgccgcag tacagtaaac
9721 cgtcctcgcc ggtgtagtcc tccggctccg ggatggtggt cgtcatattc tccaaaaccg
9781 cgttgatttc attcttcata aactctcgcc ctccttgcat gagtagtctg gtatgccctt
9841 tttaggggct tcctttttgt cgttccccgc ccatatccgc agggcggcgg catagttctg
9901 gtagcttttc ccgttggcgg caaggtagcg gctcatttcc tcgatgaacc gttccagcct
9961 gtcagggtac tctgcctgca actcgtcgta ttcggtctgt gacagaaaaa tatttccata
10021 tcggccatag ggcgcggacg gcccccccact cactcccttt gtttggctct ctgtaaggtt
10081 gtttatagta gtttggttag gggacggttt tccgaccatc ataaggtcgg ttttctgacc
10141 atcagtagga cggttttccg gctctatgag gggcggactt ccggccatca gttggtctga
10201 aaactgtacc tgtggcactg gcggtacttt gacataaagc cggttcggtg cggagaagcc
10261 gccccgttcc cgttccaaca gccccgccgt gtccagttca ttaagcgccc ccttgatggt
10321 ggtgctgcct ttatccagta tttctgctat ctccgcgatg ggatagataa tataaatcct
10381 gccctcgtcg tccagccact tgtttttctg ggagagggtg gaacggtcta acagcagcga
10441 atacagcagc ttggcggtct gtgaaatctc cattttcagc aggaaacggg gatacggcag
10501 ataggcgggc agccgcgtgt ctgccctgat ataatcagcg ataggatcac ctccctgtgt
10561 tcgggttgat tttggcgtat tgtcacgcat taaaacgccg tttccggggg aaaaggataa
10621 atgtatcgcg taccctttga cacccacgaa aaaagccttg atttatgcgg gtttgaaagg
10681 ctctaaagcg tgacatctct gcctttgttt ccgctttctc tcggcggctt tgattttctt
10741 catgcgcccg gcgcagtcgg ggcagtattt tccccggttg gatttgggga caaaggccgc
10801 cccgcagacg gcacaccgtt tccggcttcc ccggcacaag agggctgcgg ccagctcccc
10861 gtcaaggggc aggacagcca cacggaacca gtggcacatg agggaaaagg aaatgctctg
10921 gacgcacacg caaggctccc cgtcgtccaa tagcaggcag ttcccctcat cgtagttaca
10981 gcactcatgt accagccgcc gcgcccgccg gtgctggcgg tagtccatgt ggggcaggtt
11041 atcgctcatg gctctgctcc tttctgcggt gcggctcgtc ccggcgcaaa atctggtcga

11101 tgttccgctt gacggtctgc aactccttga accgctgttt ctgttcgtga taggtgttat
11161 acaggccgtc tttctcggag ataagctgct cgatctcggc ctgcaacgct ttccgggcgg
11221 gcagcttggt aatgccatgc gccttgaaat accgggctgc tgcgtcggct atgataaaat
11281 cgctctcatg ggcctgccgg tatgcggcgc gggctttctc ggatttctgt gcccgcagcc
11341 cgtcgcgggc ggccttggtc tgggcgtagg ccaacacctg ccgctgcaac ttctttttcc
11401 cttgcagctt cgtttccagt gctttcagtt cgccgctggt ctggcgcatt ttctgatagg
11461 cggtgtcaac ggcggcttct aactgctccg gggaagtaaa gccgtattgc tggtagacgg
11521 acagcgtttt ggcggcctgc ttcagattgt gtatcttcgc ccagcgttca tagccccggc
11581 ctttgccctc ggccattttc tgctcaatgt ccacaagccg ctgcactccg tcctgtttcg
11641 gggcggctat ctcggctctg gcaacctgca agcggtcttt tatggtgcgt gggggatcgg
11701 gggatctggc tggcgcttcg gctgctctgg cggcgttttg ctctaaaacg gcaaagacag
11761 cggcgcggtc aaaatcgtcg cccagcttcc gggcggtaat tggctttgtc ctgtccggcg
11821 tgaggtagga aagccgcccc cggctctcct tgacggtcac accctcctgc agcaacagag
11881 aggaaaactc gtcaaagctg gcggcggtgg caagggcttt ccgtagggtc tgccgcagct
11941 tctccttgtt cgtttcaaac ttggtctgcc ggggcgtgat actatcggca atcatggggg
12001 cgttctgctt gtccagcgcg gcctgtccct ttttctgcgc ccaatactcc cggtcggtga
12061 cgcggttctt gctgccgttc aagaggtcga tttgataaag cccctcccgg tggcacatct
12121 ccatgacttc ggatttgaag tagcgcaggg cagcgtcggt acatcggtgc ttgcagccga
12181 ctttcgtatc ggccggcctg tccatgtagg gcaggaacgg cacttcctca atccgcaggc
12241 tgtttatgac gatatgcaca tgaatgttgc cgctgtggtt atgcccgtcc gggtgggtgc
12301 agacaagggc ctggtggccg ggaaaatgct ctttacagaa ttgttccccc aacgcctgcg
12361 cccggtctac ggtcaggccg ttgtccggcc cgtcccgcgg gtcaaagctg atgatgtagt
12421 ggtggctttt cacatcttcc cgccgctggt ttttctgata gcggagattg gcccgcatac
12481 acgcaacggc aaaatcctcc ccgtcgcagt tcagcgtgga cagccggtag tcctcgcggg
12541 ggataagcct gccggtttca tcaaggacgg gcttcatggt aaactcgtca tgctcaaaga
12601 gaagatattg ctcggcggct ccgtagtcgg cgtttttaga gttgatatgc ttgagtgttg
12661 ccaaccgcgt cacctacttt cttgaggact tcatacttga ggacggcaag gtcggatatg
12721 gcggcgcgta cctcgccgga aagggtgttg tagggtacgc cgtattcgtt cagataccgg
12781 gcaatctgat tgaggttgcc gccgatcctg ccgtactcgg ctgtcagctt cccgacagcg
12841 gaaagcaact cgtcattgac cgacgacaca tggacaaccg ggcgtatggt cgcccgccgt
12901 atcgcctgcc ggagaaattc ggactggctg ataccatagg gcgcaagccg cgctgtgaag
12961 tcggcatact catcttcgga cagccgggtt ttcacaacgc ggctgcggtg gggcgtgttg
13021 tatttctttc gcatggtgtg acctcctttc tcgcccgtaa gggcgcatga gcagggtttg
13081 gggaaggcac tccccaacaa gtttcccgcg aggggcaaaa ctgcagaatt gcggattttg
13141 gcaccgtggt agaaacttgc tctccgtgac tgcaaacttt ctctcacttc cgtccgccac
13201 tttttggaa aactgcaacc acaaaagttt gtttctcttt ttctgctact actaatcctg
13261 taaagggcat tcctgcccgc tttcgctaaa atgacaccgg acgcagtggt ttctacccgg
13321 tgttggagaa atcctttctc tttgccctct actacgggta aatgctccaa atgccaaaca
13381 ccagggcaga aaaattccct cctcgcttac tactacaacc ggcagggggc aaaatggccg
13441 ggagcggagc cggacaacaa aaaaagcagt aaatcttttt caagacttac tgctttcgct
13501 ggccgcgtcg gtggcggctg gtattcagtt tttatgactt gtccggtggt tcgtcaagcc
13561 ggaacttgaa ttgacagtca attaaccgct gctttacata gtcctccacc tcggcgttga
13621 cctgcccgtt cacttttgag aaatagcgga tatatccggc gtagtggagc aggacagcgt
13681 tcacggcttc tgggtcgccc tcacgggctt tgaggattgt ttcatagggg agaagtctac

13741 tcataccggc tcacccccat ttcttcgcgt agccgctgca aggcaagctg gatatgccgc
13801 cccgctgtgc tgcgtgaccg gccaatacac gcgccgatca cgcgctgcgg ctggcgcaga
13861 aagtaatagc gcaggatttc ttcccgcgtc tgctccggca aaacagagat cgcgtcggca
13921 agggcggcgt tgcagagcag gacggtctga ccgcagacgg taaatgggta ttcctcgtcc
13981 ggctccgacg cggcaaactg gacaaacttc tcgttcatca gatagtcaag ggaaacttgc
14041 cgttcccatt gccgtttaag cttcaaaatc ttgtccaagg cggcacagcg gataacagtc
14101 ttgcaaaagg cgttgtacct gcgctggata tattcttgat aggctatctg gtcggtcatg
14161 gaaattcccc tttctgaata atagtgcggg gcggtggcac ttcttgctgt cgccccttga
14221 ttgcctacca gcaaaggcgg gcggggctgt caacggctgc gcatatgcgc cgttcatctt
14281 gaccgttgac tggctcggct gggtttgcta tttacccgac aaacttgaat ttattttaag
14341 ctatcacgtt ttaccttctt aagccttact atcattacca taggaatttc tttgttggtt
14401 ttaaaacatt cttcataaaa tccatcaatg acaaatccag ctctaaaaca aaggttaaaa
14461 atatcttgta tggaacgatg ataataaatc tgctctttcg gttgcccttc aatcgctata
14521 tcatagtaac tgtgcggtgt catatatttt tcagtcaacg tgacaaaaca agggtgttgc
14581 gttgcaaaga caaaaattcc gctttcctgc aacagttcat aaacagccat aagaagtggt
14641 tcaatatccg taatatccat aattgccata ttagaaactg ctttcgtaaa ggctcgattt
14701 ctttttaatt ctaatatact ttttctatcg gtcgcatccg ccacacaaaa ttcaatttgt
14761 tttgcatatt gtgattgccg tcttttagcc aattctatca tttttttgct gtaatcaaaa
14821 gcgacaaccg aagcgcctct ttgtgcaaga tacgaagaat aatttccatt gccacacgca
14881 atatccaaaa tgtaatccgc aggattagga gatagaagtt ccgttacttt gggacgcact
14941 acctctctgt gaaattcatt agattcgtca cccattgcat tatcccaaaa ttgtgcgttc
15001 tcctcccaga tttttttact ttcctctgtt cccatgttct ctcccactcc ccaaatttgc
15061 ttttttgctt ccattaaatc ttccttacta tattccattg ttaccctcca taacttctga
15121 ttgttgccgt cttgacgatt atgtatcttt acattacctt ctgaaacata tggcgcacct
15181 tgtccaggcg gctgtttgga cggcggggct ggatgaccgg ctgaccgaca gcggcctgat
15241 atcctttcag ctctgtaagg catacgctcc gcccgttggt gtaaaaggcc agatcagtac
15301 ggtatgcctg tatacagcgg gcgggaatct cgccagtaaa gacaacttca tcctttttta
15361 cctgggccgt ttcgatggtg gcacagtatt tcggtgcatc atgataagcc ctggaaaggt
15421 attcctgggg cgcatagagg atgaaggaga gataaggttc cagcagctgc gtccccgatt
15481 ccttcaatgc ctgttccaat acaatcgggg ccaatgagcg gaagtccgcc ggcgtgctga
15541 ccggactgta ataaagcccg tattcaaagc aaatcttaca gtccgttacg ttccagccga
15601 acaagccctg ctccagcccg taacggatac catccctgac agcgttttga aaactctggt
15661 tcaagtatcc cagcgaaacc cggctctcgt attgtacacc ggagccaagc gagagtggtg
15721 taacagacag tcctatggat gcccaaaacg ggttgggcgg cacctcgata tggatggtgt
15781 ggctggctgc tttgagcggc cgctccatat aaatgacgga gggttccttt accactgttt
15841 caagcttgta tttttccgac agcaaagcgg aaacaacctc caactgcacc cggcccaaaa
15901 aagaaagaat gatctcatgg gtgatggaat ccacttcgca acgcaaaagc gggtcagtat
15961 ccgcaagttg cgtaagagcg tccagcagcc gttctctttg cgctgccgtt ttcggcgcaa
16021 tcgtcgtccg cagcatgggg agggggtcct cgcgccacct tttacgaggg agccgggttt
16081 ggtcccctaa tacatcgttt aacctcacgc tgtcgctggg aaggataaca atttcaccct
16141 gataagcggt gtctgtccga acaatttccc ctttggatgg aatacgcatc tctgtgattt
16201 tcagcttttc tctcccggcc agggccaccg tatcccgcag gcgcagcgtt ccgctgtata
16261 accgtagata gacacgccgc tggccgcaat cggtgtactc aaccttgaaa acgctgccgc
16321 atagggcggc gcccccctgt tccccaatcg gttggaacag ccctgtcacc gcatccatca

16381 acggttgaat gccaaggcca tttttggcgc tgccatgata gactgggaac agggaggcgt
16441 cttgaacccg ctgctgttcc tcccgcgcaa gtttttcccg gctgattggt tctcctgcga
16501 tatacttttc caataattca tcgttatttt cgatgaccgc atcccatgct tctatgtcgg
16561 tattttcctc caggactatt tccggggaca gcgacaccgt ctgcttgatg ataatatcgg
16621 cggagagctt atcccgaaca gactgaacca cgctctgcaa atcaacgcca gcctggtcga
16681 tcttgttgat aaagataacg gtgggaatgt tcattttccg cagggcatgg aacagaatac
16741 gggtctgggc ctgcacgcca tctttagcgg agatcaccaa gatggcccca tctaaaacag
16801 ccaaagagcg gtacacctcc gccaaaaaat ccatgtggcc gggcgtatcc acaatgttaa
16861 ctttacatct gtgccactgg aaggaagtga ctgccgcttg aatggtaatc ccacgctgcc
16921 gctccaaaaa catggtgtcc gtcctcgttg tcccttttc gacgctcccc ggttctgaaa
16981 tggctccgct ggcatatagc aggctctccg tcaaggtcgt ctttccagcg tctacatggg
17041 caagaattcc aatattgatt attttcatgt gattgtcctc cctttacagc cccaaagggc
17101 ataaaaatcc ccagcagtaa aatactttta ccactgggga ttataagttg cggacataca
17161 catatacagc atacacctgt ttgtgattgc tgtttttggg gatatgtcaa aattgataag
17221 gcaaaagtat tcttaaattg ggtacaaaaa actaagcccc tacaaaagga gctatcataa
17281 tcctttgttc ccactatttg attatagttt tatttaagaa taccttgccg catatttttt
17341 actccttttc tggattaaat cattgtatca catcagtttt aggaaagcaa gtacctaaaa
17401 gaaatttttc ttccccttat atgtaacaat cataccggct tcctagcgtt cagaatgttt
17461 tctgctgtct gctgtggtgt ttggttggaa ttgtccaacc aaaagccgat ccgtggtgtt
17521 gtctgcattt tactaaatac aaattcaatg tatacagaaa gatataagga gtgggaggga
17581 ttccgccgta gttggcattg taggaaaatc caaaagttta gattttccca caatgcttat
17641 cttttggtct ttggttcgga atagtgtagt gctggcggtc tatctcttgt tttcggttgc
17701 ttgcttcctt accgtacatg agcatttgca ccggggttat cgttgccgta gccttccaga
17761 aggttgatcg cgccccagat accaagacct gcaccgagag ctactaccag agtctgcagt
17821 acaccgactg cactgttgaa aaatgccata agtttgtcct cctttgcccg gatccgaagt
17881 atagttggtg gataaccgga ccatcagtaa tgaaaaatgt ggatataaca aagccctcct
17941 cgaagccgca tgtgcacctg cagccggaga agggcagaaa aatagccaga actgatgatc
18001 agtcagtcct ggctatgtaa aatgttattg tcgatttcta attttgttta tggccccata
18061 aatgaaatca gatcatatcc tttttcttta cctactttca tcatgtattc catacgatag
18121 acgaacctgc ggtaaaaatc aatgatcaga tcaatttctg tgttatcttc tgtcggcctg
18181 ttggcattat aggctttaat ctggtcctca ctcatatctc ttgcataaag caattgataa
18241 gtggcagtgc cccttttttc tcttaatttt gcggtgaact cgttatctct tcctgaatcc
18301 agagcatcaa ttgtgtcgtt aatagcattc aaaatacttg ttattgaatc gaaggtgaag
18361 aaattgtgtg tcaagtacca ttcaaaacct tgtgcatcgg ctctgttttt attagcttcg
18421 agttcatcat caaagaattt ataaaggaat ggggtaaggt attggctaac atcatcttcc
18481 tcaattgaga tttcagtact tctcatttca gcaacagaat cgagattttc tgtgtttgaa
18541 tagtcaatca cccttaccgg cataatccaa aaatatgagg aactatcatg accatcgaca
18601 atacgaattt cttccttgaa gatgtctctg ccaataaacg caaagcaaag attctttact
18661 tggtttagaa tttgctcgtg attttgatga aatgccaatg cgttgtcgaa cagttttttt
18721 aactcctcat atgacttgta gcatatatct gactctaaca aaaacctgtg tgttgtgtgg
18781 tatatttccc catctgacat atagttcggg taatcctcac gaagcaagtc cagcatatct
18841 ctaatgcaaa gctgttccgg attgcagaat ctcatgaaat ccgtttcttt ctgcggagcc
18901 tttccatcca atgcatccat gagcacttct ttgctgacat catcatagtt tctaacggtt
18961 atgtggtcat tacctgcatt tctgtatttt ctccaatgct tgtattcgac gtcataaatg

19021 tagtaaagat atccgccatg agtaataaag ataagatttt cttctggaaa tcttactgct
19081 tcaaaatcat caaataattt catctcattc tatctcctta ctcgctcaca cactatttca
19141 agcgaatata gcaggttgat tttccggctc cttcgcgttt cagttcgccg gatgccacca
19201 ttttccgcag cgcaccttca acagaactaa ggctcagtga aggacactgc tccctgatat
19261 cctgcttcgt gaaacgaccg attttattca aagcagcctt gcgaaccata tcaatggcag
19321 gaagtttctc ttccacgatg gcaaatctgt ccccgaagtc tttatacgca gccagaatgg
19381 taccaagcag atatttaata aacggaactg catcctcggt tccttcatgc cagccatgct
19441 gtgcacggtt cagtgcgtca taatacaaat ccttgttttt cgcgatctgt gcctccaggg
19501 agatatattt tccaacgtag aatccgctgc gatataagag caacgtcgtc agcaagcggc
19561 tcattctgcc attgccgtcg ttaaacggat gaatgcacag gaaatcgtgg atgaagattg
19621 gaatagcaat taagggctcc aattccatat tgccgatcac cttgttgtac tcctcacaga
19681 tccgatccag ggcttccggc gtctcgaacg gggccagggg agtgaaaagt atttcggtat
19741 gaccgtcagg gaaagtcgcg ctgatataat tctgtacatt tttcgtctgt ccggccattg
19801 ggttattcat atggctgtac attatcttat gaagctgcag gatatagttc tttgacagag
19861 gaatggcatc gtagctttcg tgaataatag aaagcacatc tcgatatcct gcaatttcct
19921 gctcgtctct gttctttggc gtggttttct cctctaccag ctgcctgatt cgtgtattgg
19981 tggtaacaat cccttcaatg gcattggatg cctctgtact ctggatcttt gcgatttcaa
20041 ccagcttctc caattcattc gggcgttgct tcaagtaaag ctcctgttta ccggcctctt
20101 tatagattgc agcaataagc ccaaggatct cagaatccca tttctgattc ttgatagctg
20161 agtaattaaa agttctcatt cccttagcct ccatcttttt cccttatatt atattcattt
20221 ataagggaat agtcaacatc tcggggggat ttatccttta taatacccaa caaataaggg
20281 aaagatgcta cgagtaaggg atagtattcc catatttcca gtaccttatt cgttacatgg
20341 aatagcagtc acacacgata cgtgcatgac tgctatatat tcgttcatgg gttataacag
20401 gcgttgccac attgagctgt tctcagtcag ttcatgtact cagagagagg ctttacctcg
20461 catcccttct ttgcataata atccagcatc tcgtcagcct ttttcaaatc gtagctggtc
20521 acgcagtcgg agatgacatg aacggtatat ccggctttgg tcatattaaa gcaggttgac
20581 ttcacacagg ctgtggcatc ggcaccggca atataaaact cgccgatccc gttatcggaa
20641 ataaacgcgg aaaaggcttc gctggtcaga gcattgctct tggtcttaac aaagatatgg
20701 tcggatacta ccttcagctc cggaacaagc tcggcacctt tggtatccgg cttaaaggtc
20761 cgtgttccgg cggtgatgtt atgatgcttg atatacacga tgtccatacc acgtttctca
20821 gcccattcaa tggctgtatt caaattgtga atgatatcct tataatgctt tgtgatgtca
20881 ttctggatat caatgacaac aagagcttga ttcataggtc tgcctccttt tttcgtgctt
20941 ccttgcgttt aatgggatgt cggatgaccg ttttccattt ttcgggagca accttgcgtg
21001 catcagacag gtagatttcg tgatgaaggc ggctgtctga gaaatcattt tcatagccgc
21061 tctccagaag atactcgttc ataatgctca ccgtagcagg ttcgtcgtca aatggaccaa
21121 tgtgcatgat ctgaacacaa aggccctcat ctatggtcag gaattccgcc ttgctgcagt
21181 ccagatgttt ctttttctcc gcttccccgg tggcccaatc aaagtctttc tgcgtgacaa
21241 aatcggggag gcggatcacg gagatccagt tgaaggtgga tttgtcgctg taatcaatgc
21301 cgtcaacgcc ttcctgccac cagaagcctt ccagcggagg caccacatag tcgaagaagc
21361 cctcgatgcg ataatctgtc ttatagctca tcttgagagt ataggccacg gcatacagga
21421 tgcccaccgc ctgctgatag gcaccgcctt cgtcgttggg atcacctttt cctctcacgg
21481 caatataatt tgccttcggt accgtcacaa tttccggctt actttttggc atataaaatt
21541 ctttatactc tttcttaaaa tcaaatgcca tcttcatttt tcctcctgtt cctgatactt
21601 tgctctaatt tcggaagaaa tctttctgag cttctcccga acagactgtg gttccagcac

21661 agtcacttta tccctgcagg acagcagcca agaaatgaga ccctcttcat ccatttccct
21721 ctcgaaaagt aaggagccgt ccgtttgaat ctgataggac gagacgccat aactttcgat
21781 cagatgccac ttcatggaag gatcaaacaa cgccttcagg cgaacattgg ccgggaagaa
21841 ctgtgttgtt tcaaattccg gcatgggtac tactctctct tccacaggtt caccggcatg
21901 caccagatga tccatgcggt tcagcttgaa catgcgaaag tcctgctttg ttcggcaaaa
21961 gccccagaca taccagctgc tccacttgaa gatcagataa taggtgtcga tcaggcgcgt
22021 gctttctcct tttggagagt aataatcaaa gcgaatcgcc tgctttacct cgatggcagc
22081 ctgtatcaac tcaatttttt cggagacagc ctctttgttc caggaagata gatcaatcag
22141 gatcgtatcg tttccatgaa caaaatcaga ggatccggct ttgatttttt ccataagctg
22201 actgtaataa tggctgccgc tgacactgtc cagactccga agccctgcca tgatcatctg
22261 catatctctt gaagtgagaa gggtacgatc catgcggtat ccgttcatga tctgaatacc
22321 accacccaca ccctgtgttg tgcagattgg gatccctgct ttgcatagat catcaatgtc
22381 cctgttgatt gttcttttcg acacttcaaa atgctctgcc agttcaggcg ctgtcgtttt
22441 ttctttttgg agcagaaggg atagaatccc tatcattctg tctatcttca taatggtttc
22501 acttccggtt atatgatatc aaggtaaaca cgacaactat atgtcatgtt tatattacta
22561 catccaaaat aagaatgaaa tagcaatatg taagcaggca gcttgcgccg cctggccgga
22621 tttactgctc cggcaagcag tctgacttcg gattaaccga agagtccgga aagcagagga
22681 accagagtga taccgatgag ggccacgccg ccaccggcca tcaactgctt cataccctga
22741 gactttgctt atgtgtgata aagaagcaat agaagtgcaa aaaattttgc cgttcctatc
22801 cgacacttgg ccattgtgtc ggataggtcg ggcggttatt cgggcaagtc aatcttgccg
22861 acaaagctgt aataaatctc aatgtcctgc ctgcgggtgc cgttctcatc atagctgcac
22921 tcatgcacca caattttctc gatcatttcc cgcaagagag tgggggtcag ttcttcaaag
22981 gcaaggtgct tgcggacaat gcccataaat ttctcggcgt tgacggtagc tgcctgtgac
23041 ttgtccagtt cggcttgcag ggcggcggct ctctttttca gctccgcttg ctcggcttcg
23101 tagtcagccg acagttccat gaaacgctca tcgctgattt tgccgtttac attgtcctca
23161 tacagccgct tgataatgcg gctgatttca gaaatgcgtt cctgcgcctg ttcaagctgc
23221 ttgatggctg cggcggtctt tcgcttgccg ccgatctcgt tctgctggac aagtagtttc
23281 acaaaccggc tctcatgctt ggctgcgtat tcggtcactt gccggagatt tgccaggaca
23341 ccagcggtca acagatcggt gcggataaag tgcgccgtac agttgcgggt gcgcttcttg
23401 tagctgccgc agatgtagca gtcctgtttg cggtctttgt tctgataccg ctgctgatac
23461 agcacatggc cgcagtcggc gcagaacaaa atcccggaga acagccccac ttcatcgtag
23521 cggttcgggc gtttgcgctg tttgcgtaac tcctgcaccc gttcccatgt ttccttgtcg
23581 atgatcggct catggtggtt ctcgaaaatg gcctgcttct cgacggggtt ctctatgctg
23641 tgcttgacct tataagaagg cttttccgtt ttgaagttca ccagacatcc ggtgtactct
23701 cggttttcga ggatatgaac gacggtgttg gtcgcccatt tgcactcata gcctgggtga
23761 taacggcggg tgctgcctgt ccgctgatat tccagcgtcc ccggcgtggg gatttgctgc
23821 tccgtcagca tacgggcaat cttggtcggg ccgttcccgg caaggcaaag ctggtaaatc
23881 tgctgcacca ccgggggcggc ttcctcgtct ataataaaat tctcgtcctc gtccatcaca
23941 tagccgtaaa ccggcttgct ggtaacaggc ttgccgctca tgccttttga ctttttcact
24001 gccttgattt tcttgctcgt atctctcacc agccattcat tgaacagatt tcgcagcggg
24061 gtaaaatcgt tgtccatgcc ctcgctggca ctgtccacat tatcgttgac agcgataaaa
24121 cgcacaccct tttgagggaa cagcatttcc gtgtaaaacc ctacctgcaa gtagtttcgc
24181 cctaaccggc tcatgtcctt gacgatgacc gtacccactt tcccggcttc aatgtccgca
24241 agcatggctt gaaatccggg ccgctggaag ttcgcgccgg aaaaaccgtc gtcggtgtac

24301 cagcgcagat tggtaaagcc gttctgtttt gcgtaggttt cgagtatcct tttttggttc
24361 gatatggaat tactctcgcc ttgcaattca tcctcgtggg acaatctcgg ataaagggcg
24421 gtaatgaggt tttgggtggc ttgtcttaac ataaaatcct ccgtttccga cagccagccc
24481 cactattccg tggtttcatt gtaccacgga acggcggggg ctgtacagcg gcaaaagcgt
24541 taaatttgct tctttacagc ttttcaattt tccgattttt atggtatggg ttgtcgtccc
24601 atatttgcag tagaaaagtt cataactccg tggtatactc tgatttaaca aaaaaatcag
24661 aagttaaagg agaaaacatg aagtatacaa tagatgaatt aaccgcggcc aaaaggcaaa
24721 ttgattcaac tctgcacaag ctaagagaaa cagtaaaaac atttgaatca aaggataatt
24781 ccgagcgtta taaatcacaa atcacattgg caaagagaag aataaaagcc tttgaaattg
24841 caaattattt tatagagaat gagattaaga attgctaaag cacttccgat tttcgttcca
24901 gcggtcatgc tccctggcat ggccgctttt ccatgccccg gttcacgccg gataagtcgg
24961 ctcctgtgta gcagcggctt ccgcttccag tacccgcgcc attttgtcgg cggctgtggt
25021 tgtggtgtct ttcttgaaat agccggacac gacaaggacg gaattgccca tgcggatttc
25081 cgtcacacag tcagggcggc gggtggtgcg ggtgtcgtgc tgcttgttat ctgccatagg
25141 caatactcct ttcagtcggt aatcagtttc ttcatgctct ccattttccg cttggcggtt
25201 tcctgccgga agttgtcgcc ggtaaagcgt accgggacgc acatggaaag caggcggtca
25261 taaatccggg aatgggcggt gtcctccggg ttgtgcaggt cgtccagcgt aaggttggtc
25321 gtgacgatca gcggcttgcc gctgcggtaa cggctgtcaa tcacattgaa cacctgttcc
25381 agcccgtatt ccgtcccgcg ttccatgccg aagtcgtcaa ggatcagcag cggataacga
25441 caaaggcggg aaatgtactc gttgcgcccc tcaaagctgg cggcaaggtc attgaggata
25501 agagcaaagt tcgtcatgcg gacggggatt tctttctcca tgagggcgtt tgcgatacag
25561 cctgcaaggt agcttttgcc ggtgcctacg ccgccccaga acaggcagcc gatattgtct
25621 gtccgcatat cttcccaatg ctccacatac cggcgggcaa gcccggtctg cgggttcctg
25681 ccgttgtcgt tctcgaaagt ccagtcccgc atggtggggt cggtaaagcc ccggcgtttc
25741 agttcttcca cggtgtcaag gtgtctgcgc cgcttttcgg cggcttcccg ttcctcgcgg
25801 gcggttctct ggcagtcgca ctctgccggg tggcggtcgc gcccgaagat agcggcctta
25861 tccggcgcaa aataggcttc tttcggcttg cggcacttgc cgcagtacag taaaccgtcc
25921 tcgccggtgt agtcctccgg ctccgggatg gtggtcgtca tattctccaa aaccgcgttg
25981 atttcattct tcataaactc tcgccctcct tgcatgagta gtctggtatg ccctttttag
26041 gggcttcctt tttgtcgttc cccgcccata tccgcagggc ggcggcatag ttctggtagc
26101 ttttcccgtt ggcggcaagg tagcggctca tttcctcgat gaaccgttcc agcctgtcag
26161 ggtactctgc ctgcaactcg tcgtattcgg tctgtgacag aaaaatattt ccatatcggc
26221 catagggcgc ggacggcccc ccactcactc cctttgtttg gctctctgta aggttgttta
26281 tagtagtttg gttaggggac ggttttccga ccatcataag gtcggttttc tgaccatcag
26341 taggacggtt ttccggctct atgaggggcg gacttccggc catcagttgg tctgaaaact
26401 gtacctgtgg cactggcggt actttgacat aaagccggtt cggtgcggag aagccgcccc
26461 gttcccgttc caacagcccc gccgtgtcca gttcattaag cgccccccttg atggtggtgc
26521 tgcctttatc cagtatttct gctatctccg cgatgggata gataatataa atcctgccct
26581 cgtcgtccag ccacttgttt ttctgggaga gggtggaacg gtctaacagc agcgaataca
26641 gcagcttggc ggtctgtgaa atctccattt tcagcaggaa acggggatac ggcagatagg
26701 cgggcagccg cgtgtctgcc ctgatataat cagcgatagg atcacctccc tgtgttcggg
26761 ttgattttgg cgtattgtca cgcattaaaa cgccgtttcc gggggaaaag gataaatgta
26821 tcgcgtaccc tttgacaccc acgaaaaaag ccttgattta tgcgggtttg aaaggctcta
26881 aagcgtgaca tctctgcctt tgtttccgct ttctctcggc ggctttgatt ttcttcatgc

26941 gcccggcgca gtcggggcag tattttcccc ggttggattt ggggacaaag gccgccccgc
27001 agacggcaca ccgtttccgg cttccccggc acaagagggc tgcggccagc tccccgtcaa
27061 ggggcaggac agccacacgg aaccagtggc acatgaggga aaaggaaatg ctctggacgc
27121 acacgcaagg ctccccgtcg tccaatagca ggcagttccc ctcatcgtag ttacagcact
27181 catgtaccag ccgccgcgcc cgccggtgct ggcggtagtc catgtggggc aggttatcgc
27241 tcatggctct gctcctttct gcggtgcggc tcgtcccggc gcaaaatctg gtcgatgttc
27301 cgcttgacgg tctgcaactc cttgaaccgc tgtttctgtt cgtgataggt gttatacagg
27361 ccgtctttct cggagataag ctgctcgatc tcggcctgca acgctttccg ggcgggcagc
27421 ttggtaatgc catgcgcctt gaaataccgg gctgctgcgt cggctatgat aaaatcgctc
27481 tcatgggcct gccggtatgc ggcgcgggct ttctcggatt tctgtgcccg cagcccgtcg
27541 cgggcggcct tggtctgggc gtaggccaac acctgccgct gcaacttctt tttcccttgc
27601 agcttcgttt ccagtgcttt cagttcgccg ctggtctggc gcattttctg ataggcggtg
27661 tcaacggcgg cttctaactg ctccggggaa gtaaagccgt attgctggta gacggacagc
27721 gttttggcgg cctgcttcag attgtgtatc ttcgcccagc gttcatagcc ccggcctttg
27781 ccctcggcca ttttctgctc aatgtccaca agccgctgca ctccgtcctg tttcggggcg
27841 gctatctcgg ctctggcaac ctgcaagcgg tcttttatgg tgcgtggggg atcgggggat
27901 ctggctggcg cttcggctgc tctggcggcg ttttgctcta aaacggcaaa gacagcggcg
27961 cggtcaaaat cgtcgcccag cttccgggcg gtaattggct ttgtcctgtc cggcgtgagg
28021 taggaaagcc gcccccggct ctccttgacg gtcacaccct cctgcagcaa cagagaggaa
28081 aactcgtcaa agctggcggc ggtggcaagg gctttccgta gggtctgccg cagcttctcc
28141 ttgttcgttt caaacttggt ctgccggggc gtgatactat cggcaatcat gggggcgttc
28201 tgcttgtcca gcgcggcctg tcccttttc tgcgcccaat actcccggtc ggtgacgcgg
28261 ttcttgctgc cgttcaagag gtcgatttga taaagcccct cccggtggca catctccatg
28321 acttcggatt tgaagtagcg cagggcagcg tcggtacatc ggtgcttgca gccgactttc
28381 gtatcggccg gcctgtccat gtagggcagg aacggcactt cctcaatccg caggctgttt
28441 atgacgatat gcacatgaat gttgccgctg tggttatgcc cgtccgggtg ggtgcagaca
28501 agggcctggt ggccgggaaa atgctcttta cagaattgtt cccccaacgc ctgcgcccgg
28561 tctacggtca ggccgttgtc cggcccgtcc cgcgggtcaa agctgatgat gtagtggtgg
28621 cttttcacat cttcccgccg ctggtttttc tgatagcgga gattggcccg catacacgca
28681 acggcaaaat cctccccgtc gcagttcagc gtggacagcc ggtagtcctc gcgggggata
28741 agcctgccgg tttcatcaag gacgggcttc atggtaaact cgtcatgctc aaagagaaga
28801 tattgctcgg cggctccgta gtcggcgttt ttagagttga tatgcttgag tgttgccaac
28861 cgcgtcacct actttcttga ggacttcata cttgaggacg gcaaggtcgg atatggcggc
28921 gcgtacctcg ccggaaaggg tgttgtaggg tacgccgtat tcgttcagat accgggcaat
28981 ctgattgagg ttgccgccga tcctgccgta ctcggctgtc agcttcccga cagcggaaag
29041 caactcgtca ttgaccgacg acacatggac aaccgggcgt atggtcgccc gccgtatcgc
29101 ctgccggaga aattcggact ggctgatacc atagggcgca agccgcgctg tgaagtcggc
29161 atactcatct tcggacagcc gggttttcac aacgcggctg cggtggggcg tgttgtattt
29221 ctttcgcatg gtgtgacctc ctttctcgcc cgtaagggcg catgagcagg gtttggggaa
29281 ggcactcccc aacaagtttc ccgcgagggg caaaactgca gaattgcgga ttttggcacc
29341 gtggtagaaa cttgctctcc gtgactgcaa actttctctc acttccgtcc gccacttttt
29401 tggaaaactg caaccacaaa agtttgtttc tctttttctg ctactactaa tcctgtaaag
29461 ggcattcctg cccgctttcg ctaaaatgac accggacgca gtggtttcta cccggtgttg
29521 gagaaatcct ttctctttgc cctctactac gggtaaatgc tccaaatgcc aaacaccagg

29581 gcagaaaaat tccctcctcg cttactacta caaccggcag ggggcaaaat ggccgggagc
29641 ggagccggac aacaaaaaaa gcagtaaatc tttttcaaga cttactgctt tcgctggccg
29701 cgtcggtggc ggctggtatt cagtttttat gacttgtccg gtggttcgtc aagccggaac
29761 ttgaattgac agtcaattaa ccgctgcttt acatagtcct ccacctcggc gttgacctgc
29821 ccgttcactt ttgagaaata gcggatatat ccggcgtagt ggagcaggac agcgttcacg
29881 gcttctgggt cgccctcacg ggctttgagg attgtttcat aggggagaag tctactcata
29941 ccggctcacc cccatttctt cgcgtagccg ctgcaaggca agctggatat gccgccccgc
30001 tgtgctgcgt gaccggccaa tacacgcgcc gatcacgcgc tgcggctggc gcagaaagta
30061 atagcgcagg atttcttccc gcgtctgctc cggcaaaaca gagatcgcgt cggcaagggc
30121 ggcgttgcag agcaggacgg tctgaccgca gacggtaaat gggtattcct cgtccggctc
30181 cgacgcggca aactggacaa acttctcgtt catcagatag tcaagggaaa cttgccgttc
30241 ccattgccgt ttaagcttca aaatcttgtc caaggcggca cagcggataa cagtcttgca
30301 aaaggcgttg tacctgcgct ggatatattc ttgataggct atctggtcgg tcatggaaat
30361 tcccctttct gaataatagt gcggggcggt ggcacttctt gctgtcgccc cttgattgcc
30421 taccagcaaa ggcgggcggg gctgtcaacg gctgcgcata tgcgccgttc atcttgaccg
30481 ttgactggct cggctgggtt tgctatttac ccgacaaact tgaatttatt ttaagctatc
30541 acgttttacc ttcttaagcc ttactatcat taccatagga atttctttgt tggttttaaa
30601 acattcttca taaaatccat caatgacaaa tccagctcta aaacaaaggt taaaaatatc
30661 ttgtatggaa cgatgataat aaatctgctc tttcggttgc ccttcaatcg ctatatcata
30721 gtaactgtgc ggtgtcatat atttttcagt caacgtgaca aaacaagggt gttgcgttgc
30781 aaagacaaaa attccgcttt cctgcaacag ttcataaaca gccataagaa gtggttcaat
30841 atccgtaata tccataattg ccatattaga aactgctttc gtaaaggctc gatttctttt
30901 taattctaat atactttttc tatcggtcgc atccgccaca caaaattcaa tttgttttgc
30961 atattgtgat tgccgtcttt tagccaattc tatcattttt ttgctgtaat caaaagcgac
31021 aaccgaagcg cctctttgtg caagatacga agaataattt ccattgccac acgcaatatc
31081 caaaatgtaa tccgcaggat taggagatag aagttccgtt actttgggac gcactacctc
31141 tctgtgaaat tcattagatt cgtcacccat tgcattatcc caaaattgtg cgttctcctc
31201 ccagattttt ttactttcct ctgttcccat gttctctccc actccccaaa tttgcttttt
31261 tgcttccatt aaatcttcct tactatattc cattgttacc ctccataact tctgattgtt
31321 gccgtcttga cgattatgta tctttacatt accttctgaa acatatggcg caccttgtcc
31381 aggcggctgt ttggacggcg gggctggatg accggctgac cgacagcggc ctgatatcct
31441 ttcagctctg taaggcatac gctccgcccg ttggtgtaaa aggccagatc agtacggtat
31501 gcctgtatac agcgggcggg aatctcgcca gtaaagacaa cttcatcctt ttttacctgg
31561 gccgtttcga tggtggcaca gtatttcggt gcatcatgat aagccctgga aaggtattcc
31621 tggggcgcat agaggatgaa ggagagataa ggttccagca gctgcgtccc cgattccttc
31681 aatgcctgtt ccaatacaat cggggccaat gagcggaagt ccgccggcgt gctgaccgga
31741 ctgtaataaa gcccgtattc aaagcaaatc ttacagtccg ttacgttcca gccgaacaag
31801 ccctgctcca gcccgtaacg gataccatcc ctgacagcgt tttgaaaact ctggttcaag
31861 tatcccagcg aaacccggct ctcgtattgt acaccggagc caagcgagag tggtgtaaca
31921 gacagtccta tggatgccca aaacgggttg ggcggcacct cgatatggat ggtgtggctg
31981 gctgctttga gcggccgctc catataaatg acggagggtt cctttaccac tgtttcaagc
32041 ttgtattttt ccgacagcaa agcggaaaca acctccaact gcacccggcc caaaaaagaa
32101 agaatgatct catgggtgat ggaatccact tcgcaacgca aaagcgggtc agtatccgca
32161 agttgcgtaa gagcgtccag cagccgttct ctttgcgctg ccgttttcgg cgcaatcgtc

32221 gtccgcagca tggggagggg gtcctcgcgc cacctttttac gagggagccg ggtttggtcc
32281 cctaatacat cgtttaacct cacgctgtcg ctgggaagga taacaatttc accctgataa
32341 gcggtgtctg tccgaacaat ttcccctttg gatggaatac gcatctctgt gattttcagc
32401 ttttctctcc cggccagggc caccgtatcc cgcaggcgca gcgttccgct gtataaccgt
32461 agatagacac gccgctggcc gcaatcggtg tactcaacct tgaaaacgct gccgcatagg
32521 gcggcgcccc cctgttcccc aatcggttgg aacagccctg tcaccgcatc catcaacggt
32581 tgaatgccaa ggccattttt ggcgctgcca tgatagactg ggaacaggga ggcgtcttga
32641 acccgctgct gttcctcccg cgcaagtttt tcccggctga ttggttctcc tgcgatatac
32701 ttttccaata attcatcgtt attttcgatg accgcatccc atgcttctat gtcggtattt
32761 tcctccagga ctatttccgg ggacagcgac accgtctgct tgatgataat atcggcggag
32821 agcttatccc gaacagactg aaccacgctc tgcaaatcaa cgccagcctg gtcgatcttg
32881 ttgataaaga taacggtggg aatgttcatt ttccgcaggg catggaacag aatacgggtc
32941 tgggcctgca cgccatcttt agcggagatc accaagatgg ccccatctaa aacagccaaa
33001 gagcggtaca cctccgccaa aaaatccatg tggccgggcg tatccacaat gttaacttta
33061 catctgtgcc actggaagga agtgactgcc gcttgaatgg taatcccacg ctgccgctcc
33121 aaaaacatgg tgtccgtcct cgttgtccct ttttcgacgc tccccggttc tgaaatggct
33181 ccgctggcat atagcaggct ctccgtcaag gtcgtctttc cagcgtctac atgggcaaga
33241 attccaatat tgattatttt catgtgattg tcctcccttt acagccccaa agggcataaa
33301 aatccccagc agtaaaatac ttttaccact ggggattata agttgcggac atacacatat
33361 acagcataca cctgtttgtg attgctgttt ttggggatat gtcaaaattg ataaggcaaa
33421 agtattctta aattgggtac aaaaaactaa gcccctacaa aaggagctat cataatcctt
33481 tgttcccact atttgattat agttttattt aagaatacct tgccgcatat tttttactcc
33541 ttttctggat taaatcattg tatcacatca gttttaggaa agcaagtacc taaaagaaat
33601 ttttcttccc cttatatgta acaatcatac cggcttccta gcgttcagaa tgttttctgc
33661 tgtctgctgt ggtgtttggt tggaattgtc caaccaaaag ccgatccgtg gtgttgtctg
33721 cattttacta aatacaaatt caatgtatac agaaagatat aaggagtggg agggattccg
33781 ccgtagttgg cattgtagga aaatccaaaa gtttagattt tcccacaatg cttatctttt
33841 ggtctttggt tcggaatagt gtagtgctgg cggtctatct cttgttttcg gttgcttgct
33901 tccttaccgt acatgagcat tagcaccggg gttatcgttg ccgtagcctt ccagaaggtt
33961 gattgcaccc cagataccaa gacctgcacc gagtgctact acgagagtct gcagaacacc
34021 gactgcgctg ttgaaaaatg ccataaagtt ttcctccttg atttgtccgc ctatgtggtc
34081 gggataatga aatgaaattt gccccgcgat tcccggcatg tgcacctgca ggaaaagaag
34141 ggaataaaaa tatccggggc cgatgatcag tcagtcccgg atatgtaaac aggcagcaag
34201 ctgccgcctg ttgcccgagc atttactcat tttgttctgc tttcctgcag ccgagtgttt
34261 ccagaatacc tttcagatgc tttgcgcttc gggcatattt gtctccggta tccaaccgga
34321 tccagatggt tgctgtaccc gggctggcat caatgctgat gatgtgatct gcattgatat
34381 actcctcacc ggtgtatacg aatgtcgcca ttttgcctcc tacctggcag cttcctgctt
34441 cctggccgcc ttaccggaag ggctggctgc cgctgtttct tccttccggt cagacaagtc
34501 cgagaggaca gagcctctgg atttatcttt ataggactcc atcagggcct tcagttccgt
34561 gttggtcact tctctggtct cacgacccca ggtggatctt ccggtttcat ccatgccggt
34621 cttcaccgaa cgggaaagcc gggtcgtacc atcctcgggg agcttcatcc agacgccttt
34681 tccgaattga ttatcgtgaa tcatcttcgg ggagatcaca aaggattccc agggacgggt
34741 atcggtctta tccggattcg ggatactgac ttccaccagc tccttgccgc ttttcgaagt
34801 aaaggactca cccacgagcc ccttaccaaa tttgatcgtt acctcggcca gcggctggcc

34861 aaaggtttcg ccacgcgcca tggcttcctc cgcttctctt acgaagtcgg gaacatccat
34921 cggcaacgga tttctttcat ctgccattgc ggcttacctc cttatttttc tgcatctata
34981 acaaaaaact catccttcgg atgaagcttt gccctggtac tcaggtactt ctcaatgtcg
35041 aaggtgagtt tcttatcata atcggcggtg tatttgtagt tcgggtgctg ggtcagatcg
35101 tatttatccg acaggaaggg acgaacgcct ctcagctgaa ggacacattt gctgccgtcc
35161 ataacagcca gttcatcacg actcatgagc tccttgccca gcttctggta gttggttccg
35221 tagcttggac tattgcctcg tgtatccgag gtattgaagg tatcgatggt ttctttgccg
35281 agtgcttcgg agagttcttt cagagccgcc caggaatacc tggcagtcca tgttgccgat
35341 gatggtattg gcattgtcct tgtaaatagc cttcagctga gaccttgcct gcagcaccag
35401 acaggcactg atctcacggg aacggatggt ggcaaccagc ttttccaggt tcgggatctg
35461 gccgatgttt gccgcctcat cgatcaggca ccgcacatga accggaagcc tgccgctgta
35521 aacatcatcc gctttttcac acagcaggtt gaacagctgc gtgtagatca tactgatcag
35581 gaaattgaag ctgccgtctg tatctgacat catcaggaac agggcagttt tttcatctcc
35641 cagagtatcc agctccagct cgtcatacat ggtgatatcg cggacctcct gaatatcaaa
35701 gggagcgaga cgggcaccgg cagacaccag aatacttttc agagtcttgc ctgcagccag
35761 cttgaacttg gcatattgcc ggactgcgaa atggtccggc ttcttctttt tcaggtcttc
35821 aaacatcaga tccaccggat tctgaaaatc ctcatcatcc tcacggacct ccatggcatt
35881 gataaattcg atgagcgtgg caaagttctg ttcctccttg ggtgcctcat aatggatgta
35941 gccgatgaga gcacaataga gcagtgtctc tgccttgtcc cagaacggat cgccgccctt
36001 accatcaccc ttggtatttg ccatgagtgt ggtcgtcagc ttcaggatat ccttttcgct
36061 gtggatatag gcgaaaggat tatagtgcat ggacttgtga aaattgattg tgttgaagat
36121 tttgatccgg taattcgcat ggacaaatgc atttccaacc tcgagcagag tcgtgccctt
36181 cggatccgtc agaacatagg aagaatgaag ctgcaacaga ttcggcttaa tccagaaccg
36241 ggttttaccg gaaccggagc cgccgacgat cagtacattc ttgtttctgg cattagcagg
36301 attcttcggg cggttgctca tcatcagtcg ttctgtcttt gtgaggatca cattgttctc
36361 aaagaccgga tcgatgaaag gctcgatgtc cttcgggcca ccccatcggg cagaaccata
36421 ctcaaggtta tgcttaaaat ttttggcatt ctttccccgg atgtagacag cagtcctcag
36481 acctgccccg cagatcaggc caaccatcaa gtcaatggga tgaaagctcg gaagaggatt
36541 tccgaaagca atgcccatcg tcgtgaagaa tgacagcagc ttttcggaag cattcgcgcc
36601 ctccgccatt cgccaggctt ctcccagatt ggtacagacc agaccaatca gaatataggg
36661 gagggcccgg atcagaagct ttttcagctt aacggtatcc gggctcatcg cgactgctcc
36721 ctctgcttgt ccctgcttac caccttcgcg gggatatctc taacggcatc gatcagctta
36781 tgcagcgtct tcagaacgct cggacgcttg ttctttgcca gtatctctgc agagtaggca
36841 ttgaatgccg ccgtcatggc atccgcgtct ctggccttaa agaacaccag gtacttcggt
36901 ggaagtacgg atttatcctt ggtgattgca taatcgacac cgtatttccg tgcatacttt
36961 tcaaagccct tcaggtctgt tttggaaatg tcgatattgg tgacgccctg attctgaccg
37021 atcaactcct gaatggtctg cttgccggtg ggcttctcat tttttattgc tgcttcctga
37081 gattttcgct tgatataagc gatacccttc aggccaagtc ggacaatact gcgagccgta
37141 agcttggtcg tgctgatggc caaattcact gtcctgtttt ctacttcctc ctgcaatgct
37201 caccacctcc ttcagttcgt ttttccggca tcatgccttt tttccggcgg agccctccac
37261 cttcataatg ccgtcaagag tcgtagcggc aatacggagc ctctttgcca gaccgatatc
37321 gtctttgacc tgtccatcca tcgagtcgcc gcaaactacg agaatgcggg atcttctgag
37381 aagctccgat gccctttcaa tcctgtcctt gtgctcctgg ggaatctcat ctcttaagag
37441 accttcgtac atgagtctcg gacagatagg agaaaaccct gcatcataca cctttcggca

37501 ataacgcagg gccgtgcttt ttgcttcaaa ttcgttatca ctccagtttg cagtgatata
37561 tgccagtggt ctaaccatgt gtacctcctt atttcaacat cattagtgac cgtcatgctc
37621 agaccttttg tgcttcggct taccgccaat ctcttcggca gctttcaccg cttcgcgtgt
37681 ctggagggcc tcttcgggaa tagggagcgc catgatgctg cgtcccattt tcacgaaggc
37741 ctccggctgg tgaaacagct cttcaaactt gccgacctgc tccggagtga gggaaccgaa
37801 gctgtcctcc gtaagaccgg tcaccaggaa ggaaccggca atcacatcat aaacctcatt
37861 gtcctcatca cgaagagcac ggttgagagg aagtccgttg attttgccct cctcattgac
37921 aatcagacca accggatcct cgaagggata aaccacctcg ataaaaccgc ctaccagctg
37981 ctgaagatcc tcaaggccgg tctttgcttc aatgaccttc gggtgctcgt tgggttcgac
38041 aagaagcatg gtaatcgtat ctgcctccac caatccagcg ggctcatcgg caagcatggc
38101 ttctactctg gcagcctcgc gggcctcaaa ggtacgggca ttttcaaaga tatgcgcttc
38161 actgtcgtag tggaacacat tgtcggagag ccggtccgca ggggcaacct ccgcctcatt
38221 gatgcttcgc accatttctt ccagatgact gagctccata cttccgtcat cggcaatgaa
38281 cagaacttca tggattgaag aagggagaac gtagaaatcg ccgcccatcg tttcagccgc
38341 ctgatccaga aaatccggat actgaatggt gcaggcaccg ttctgaccgc cctctaccgt
38401 agctacccag atgggagaag gatcatccgg aataaacata ccggcagcat cgcccatcat
38461 gtcactcatg acatcgttca tgttgcgaag ggtcgccgga tggttctgca cggcagcttc
38521 catggcatcc gaatgaagct gatccgcagt aatgtcatag gtctgaagca tgttgttggt
38581 tacgagaatg ctggcagagc cctgctcatt actttccatc tcgaagcggt aaacaagagc
38641 catgtcttcc acaggtctgt gaggaatctc ggaaagcttt tcctcgtttc cggcaatggg
38701 aatcatctga agggtcagct tttccttcat ctgatcgtag tccatcagcg cccgggtatc
38761 gaactgcggc atgcttccag caatctctgc cacctgcttt tcaatgttat gaagggccgt
38821 ctccataggc ataccgtcct caacacgttt gtaaacatag tccagattca tcgtggcggc
38881 aacgttactg cctgcagggc tgacagccag tccggtatag gattctccct gcagtttcga
38941 aacggcctga atgccaatat ccagatcggg gaagcgctca cgcatctcat cgacaacctg
39001 aatcttaaac tgatcaaaat ccatgtgtta gtcctccttt ttctctttcg gcagccaata
39061 accacagatg gtctttccgt cttcaccggt aatggtcacg aaatggtcat tcaaaaagat
39121 cattgcagca agaaagttca tagcggtaag ctcctcgcat tcggtttcat cgtcattgac
39181 tctggcaata taaaccggag ccgtgatggt gatatcacct tcgctttccc aagtgtgctc
39241 catatagtgg agaaacagat ggtcgccgat ctcaacacaa taaaacgtct gttcatcatc
39301 aaagcccggg aactttgaca ggacatcttc ttccgtcatt agctgtaccg gcttgccggg
39361 aaccgcaaaa aggacggtgg catccataca ggccatatca tcctcgtccc agtcatccat
39421 gtcttcgaaa ttcttcgtgt cccagggatt cttctttccc tgaagcaggc tggcaattcg
39481 ggaaaggaca atggcactgt cacagaacat ttctgccagc tcgtctacag aatctgcaag
39541 ctcaattgct gcagtctctt tttcggaagg ctttttccgg ataccaagca ggttctgcat
39601 ctgattcatc atttcttttt cagtcattga ttgatactcc ttttcgtatt ttgggtaaaa
39661 agaaagcact atcaaaaggc ttgcttctga cagtgctcgg ggtatgggta ttaatttatc
39721 gttcctgatc gcgagctttc tttttctgcc actgatccag aagcttgata atggtgtcct
39781 gcatctgctg tgtggtgtag gacctgggga aatacttcct cagctgttca tttcggaagg
39841 tgacccgttc aagatcacct ttcttcactt cattcatgat ttctctggca gcgtccatgg
39901 tgaaatcacc ttcctgcgcc agcttcttca agcgctgagc ctgggaaaga gaaggggaag
39961 cctgtacttc atccatggcc tgcatgaaga tttcctgttc ctccggagca aggtaggaaa
40021 gctcgaccgc cggattgaag gaaatctgct tctgatcgac catttccaga agttcgggaa
40081 tgaggttggt gagacggatg tagcgctgga cttgctttct actttcacca gcttcctgag

40141 ccaattgctc gtccgacctc aacttagggc caagttggcc cgaagttaaa tctgatcgtt
40201 taccctgatg tttcatcgct tccaacttca ttttgtacgc ataagccctc tcgctgggca
40261 gaatgttctc tcgctgaaga ttggaatcca ccatgagcac cgtggcggca tcatcatcca
40321 tctgccttac cagaaccgga acctccgtaa gccctgccag ttcagcagca tgggcacggc
40381 gatgtccgga gatgatctca taaccacctg cttccagcgg cctggcaatc agaggggtca
40441 gcactccgta ctgtgcgatg ctctccgttg ttcgaagcat ggcttcatca tcgaccacct
40501 tgaagggatg atccttaaag ggaaccagct cgctgatgct gatattctgc actcgttcca
40561 actgactctc tgcacggctc tcatccgttt caaagatatc atcgtagctg gtcaagctga
40621 tgtttccgcc tcttttcggc attcgctagc acctcctttg tcagactccg gtaggcctca
40681 gcgactttgc cgcccggatc gtgtttgaaa atactcacgc cctccgcgct gatttccgca
40741 gcacggacgg agcgagggat gtctgcgtcg tagaccttca gcttgctgcc gtaggtatcg
40801 cggatcagcg tactgatctc ctttgcaaag ttggtacgaa gatccaccat agtcagcaga
40861 attccttcga tccgaagctt gggattgatc tgacgcttga ccttgtttac cgtctgaagc
40921 agctgctcca gaccctttgc cgacagatag tttgcctgaa caggaatgag gacctgatcc
40981 gatgcggcaa gcgcattgac ggtcagcatg cccagggagg gcatacagtc cagcagaatg
41041 tagtcgtact gaaatttcaa agcatctaaa acctgcctta aaacagtttc gcggctcatg
41101 gcgttgacca gcgacacctc catacctgac agcgtaatat ttgcaggaac aagatccacg
41161 ccttcttcat gtgacaggat accctcctta ggtgcgatga gctgatcctg catgaccttc
41221 gccatcaggt cggtcagcgt gaccggaaga tcatccggtc tgggatgtcc aagggcaatg
41281 gttagggacg cctgcggatc cgtgtccacg agcagcactc tttttccctc attggcaagg
41341 ccaatgccca gattttcgca ggtggtagtt tttccggttc cgcctttctg attaacaaca
41401 gccatgattg tggctctcga cataatgatc acctcctaga tcaagccatt ggccatgtcg
41461 tgctgtaccc agttctgata gtacgaactg atggtagttg gtgcgttgta gaggactgcc
41521 agcatgtact gcttcatgtc cttaacgcgg gttgtgttgt ttgccagtgt catcagggca
41581 tatttgatat gctctgcatc cagcttcata aatcgacttt ttacgacttc gattggcata
41641 tcttcccggt tgatacgaat ggtctttttg cttgagcagc aaacatcgac catgagatcg
41701 accatagcct gaatggtatc ctgatcaagt gtcttatccc gggaaagata atccagctcg
41761 agagattctc tgaaataatc ctcatactga gctcggctcc gcatcgcttc gcttcctttc
41821 ccgacatttc tgtcggaata gaaggaatcg gtatcactat actcagtatt aataatatca
41881 gtcttatttc cgtgtccttt tgacacttca agagatgtga tttcgacact tctagaagtg
41941 tcattttgac atttcaagaa atgtctctcc acagggctgt ccaccgatat gaagttcttc
42001 acatagatca ggtttggttt tcccagcccc tgacgctttc tctcaatgag accgactttg
42061 tcctccaaat ccgataacaa ctggatcgct tttttgtttc cgcagttcat gtccgccatg
42121 atttcttcaa tcgtgcagat gatatacacg cggccctgtt catcgatcca tccgtttttc
42181 gcagagaggg gaacgcggtc cagtaagatt ccgtatagag tcttggcttc agcagacaga
42241 tatttgaacc ggtcattcgt aaaaagcacc ttgggaaccc ggtagaaggc aaactgttct
42301 gcctgggctc catagaaata atcaaacact catagtcacc acctgctttc aggagaattt
42361 atcgcgcttc agaggctttt tgcgaaacac atccttatcc ttgccatagg ggcatctggt
42421 gaactggcac cggcgatact tgaaatccgg gcgatagtag aggcagtgag cgcagcttcc
42481 gggtttatgg tccgatgcac tcccgggatt ttctctcttt gcagccctcg ccagcttctc
42541 atatctgcgg aagacctcat ccttcttact catttgccgc accttcttcc tttgcgcggt
42601 tagcgcggat tgctctgagc catgcgtcca attccttgta gcagacagcg ctgacacagg
42661 gatggccttt ggcgtaacag cagccctcac atttcttttc ctgctcggtc tttactgcct
42721 cagccaggta gtagcagttc tgcgtaccaa gcacacagcc gaccttgcga ttcttccaga

42781 agaagcactt gcggcaggta tccggcttgt catcatagta tctgatcccg tcgatcagaa
42841 tggttttgcg tttcactttt tttctcatca cgatatcctc catagttttg atccgccgcc
42901 ggggaagtcc ggagacgaca aaagccccgc agtgattttc atcaccacgg ggctatgtaa
42961 gatatcggtt ctatactgaa tgttggggtg tgctgaaaag cacacttcca acattctttt
43021 cgttgtaata aaatacatct ataagaagcc tcctaaacgc attatcccga ccaaagttta
43081 atgttacgga ggtgacccct tatagatgta ccctaattat accaaggctt tccttaactt
43141 ggaaggtgtt tctattaaaa aagtggttca ggcagactct ttcattaaaa ttttcatcca
43201 gtcccaaccg gtagaacaga cttgtccctg ctgtggggct aaaacaaaac ggattcatga
43261 ttaccgttta caagaagtcc aggacattcc cttactggga aaacaagtaa tcctgcttct
43321 tcgcaaacgc cgctacctct gcccatactg ccgcaaacgg ttcacggaac cctattcgtt
43381 cctccccagc taccaccgca ggacccgcag gctggcattt tacattgtct ccctgctccg
43441 gcagaccttt tccttaaaac agatcgcaga gcttaccggt gtttccgtcc agacggtctg
43501 ccgccttctg gacacgattt gctatcctcc gcctgaccag cttccacaag cgctttccat
43561 tgacgaattc aaaggcaatg cttctaccgg taaatatcag tgcattctgg ttgatccgaa
43621 aaagcgccgg atcctcgaca ttctcccaga tcggacccag agccatctgg ctgattattg
43681 gcggaacatt cccaggaaag aacgcctgaa ggtaaagttc ttcgtctgcg atatgtggcg
43741 cccctacacc gaacttgcac aaaccttctt tccaaacgct acaatcatcg tggataaata
43801 tcatttcatc cggcaggtga catgggcaat tgaaaatgta cgcaaacggc tgcagcgctc
43861 catgccggtt tctctgcgta agtattataa acgcagccgg aaactcattc tgacccgcta
43921 caaaaagctg aaggatgaga acaaacaggc ctgtgattta atgcttcact atagcgagga
43981 tctgcgtctg gcacaccgca tgaaagagtg gttctatgat atctgccaga tggaagcgta
44041 tcgtcagcag cagagggaat ttgatgactg gattgcgaat gcacagagct gtgggatcaa
44101 ggaatttgag gcctgtgcta agacctacag ggcctggcga aaagaaatcc tgaatgcctt
44161 taaatacggg ctgacaaatg gtcccacaga aggatttaac aacaagataa aggtattaaa
44221 acggagcagc tacggaatcc ggaatttcaa acggttccga acccggatac ttcactgtac
44281 atcataggat aaaaaggtcg gtaatgcgga ggcttatttg gcatgccaaa atcagtaaaa
44341 cctggaacta aacataaaaa gggccctttt catagaattg gggcgggatt ttaacaaatc
44401 ccaccccaac tattgacaaa gagccaagat atcgatattt agttgttcag gagaatgaaa
44461 aactgaaatt tatcggtgta atttttcaag ttcatgctgt atttcttcag catgtttaat
44521 ctgttcattt aagttccatt ttccaatatt ttctccatct ctgagaaaca tagaaccaca
44581 ttgatggaaa taaaattcaa tgtttcgctc tttggcttct aaatacaaat ctcttaccca
44641 ctcatacatt ataggtctga catctgcttt gggtgccatt tcaccgccta cattaacaca
44701 cttgattaaa cctgtatcca gatactttcc aagagatata ggagcaatta acggtgagca
44761 aaagacttca cggtgtttta ggggagcatc aagaaaataa ggcaatcgta tatctgccat
44821 ttcttgattt tcgatggata tactcaaatg gaaatgctcc cagccttcac cccaatccgc
44881 aggaacgcac tctttaattc tctcaggcct ttttgttgtt gtaacaaaaa tacaatcctt
44941 tcttcgtctg atgaaatccc agcagccgtc tctccattca tctgcttctt cgataaagaa
45001 atcagaagaa aaacacagtt taacaagaga gcccggcgga cagtccttat ccttcagctc
45061 ataggtagtc tttcccttgg taataatcgt agtatcacga ccgtatcttt tgtccatttt
45121 ataaacaaag caatttttgc atccggggct taccttttta cagccatgcc acggattcca
45181 tggaatcgct ttttcaatct gtgtgttttg cataatgtca cctccgcaaa ttccgatttc
45241 ctgcttcctc taatcccggc ggttgatttt agaaatctaa aatttcgcca cctgattttc
45301 taacgccttt agaaatccgc cctccggtct tttctttcgt catagcagtt ctgggtatca
45361 tatttcagat ttctgaggcg atagggggat ttatcacaga tttacaccca gatacaccgg

45421 tgaacacgca aacacgcgtc atttttagcg gaatcaccag ttcaggtcgt ttttcctcat
45481 tagataaaag ccctgttttt ttagcacttt atcaacgaag cctctttttt attagaagaa
45541 acgcggaaga tttcgtgaaa aatcttccgc gatagtgcgc tggcgtccct gactggattc
45601 gaaccagcgg ccttccgctt aggagtagac ccgatacgat gaaaccaagt gcaacacagt
45661 gcaagaaaat ccttggaaat acaaggtttt tcactgtttc aaatcctatc tggtattacg
45721 cttcataagc attttcatag ggattttaca ccgtccaatt ggctcgaaat tagcaaggct
45781 ttcgattttc ggtggacgtt gcaagtgcaa gcaaattcct ctatgtaact tctcgtgata
45841 cgaaatgata ggaaatccta ttcattctcg tgcggaggcg ctgatgtctt tagacaccat
45901 cttagtctgt ttaatatatc atttcaaaat agatttttca accagcatta ttaagaattt
45961 caaaggtttt ttcagactca atgattctat gtaatttaga cccgccgaag caggtttttt
46021 tttatttcag cataatctct ctatcctcag cgccgagcat ataagcaccc ttggcaaaag
46081 catcccttgc cagaagctct gctacatcaa ccctatcgaa cagcatccat atttctttct
46141 ctttcggagt cagttcatgg ctatggtcaa acaaatgctc aagtcccaat ccttcaagca
46201 atgccgactt tgctgctttg tattcctcag ctctgccaac atattcgtcg ttcctttcct
46261 tggaaaatgc ctcgatattg tcagctctca gatgctcgta atactcgtcc ataaagattc
46321 tatccatact cgttgacctc cataacaaaa aaataagggc agaccctttt acagctattg
46381 tagctatgta aaaggatctg cccttgcagt tactataatg tttttgccag cgacttcagg
46441 agtgctgcca tttccggatt ttgcagaagc ttcagaagaa gttctttatc atcagcgtag
46501 ctttccgttg cggctggttg aaccggctca ggcgttgcct gtctgcctga gtagaacgca
46561 gcttccattc tttcagcgtt cagtctgcgg tcatcgtcga tgatatggga gtaaacatcc
46621 gctaccatct tgacctgtgc atgaccggaa tcgccttgaa cggatttcat atcgccgccg
46681 ttcagcttca gcttgtaagt gatactggaa tgtcggagac tatggaaaac cactttcggc
46741 agtcccttct cctcaatcag cttattgaaa gcacggttga taacctgccc ctcaatcggc
46801 ttgccattgg aagaacagaa aaccagattg aagtctacga actcgtcacc gaaaaggtct
46861 ttcagttctt cgatgtcagc ctttcgctgt accaacattt ccgctacggt tttcggtaag
46921 aacaccttgc ggacacttgt tttggtctta ggagttttga gaaccaatgc ggtatgcgta
46981 ctggcgaagg tcggcgggaa cttgaacatg atgtccttac cgtccagatc tgccatcgct
47041 tcacgattta ctctctgcaa ctccttttca acgaagatgc ttgcctgacc aagttcaata
47101 ctggtagagg aaatgtcaat acagtcccag gtaagcccca gaagctcgcc cattcgcagg
47161 gaacaggaga aggcaaggtt aatagccaga cgaagaatat catcgtcaca ggcttccaga
47221 gccttctgga gcacttctgc tgtccagatg tccctggtct tgtgttcttc tttcggcagc
47281 gtagcgtgct ccacagggtt tctggtcatc aattccccact tgaccgcctg attgaaagca
47341 tttcgcaatg tcttgtgaac ctctcggaca gtatgcggag tcagatattc tgtacgggct
47401 tttacatact tagaagaaac cgctttgaca gaaagcaaat ctcggtaata cttgtccatg
47461 attctcggcg tgacatcctc cagcttcata tcaccgatga ggggacgaat atagtttgca
47521 atcaggctct ttctgctttc ataggtagac atcgcccagg tgttcacacc atagattgac
47581 atatactcat ccaaaagatc attgacggtt ttagcagaag ggggaataaa ggttccggac
47641 tcctgctcat attcaacctg caacttcctt ttctttgctt cggcattcgt ttcaaaggtt
47701 tcccacttct gacgcttgtt gccgttctca tcggtgtatg tataaaccac agaatacttc
47761 ttttttctct taacgataga tgccatgtaa tcagctcctt tcctgtctat cttccgctaa
47821 ctggtagcgg gtctgattgt gataccattt ttcaaagctt tctttggtag ctcgcttata
47881 cctgccaacc ttgataagct caaaatcttc attggcaacg atatagtaaa tggtctgacg
47941 atgtacccca agcattcttg ccatttccgg caggccgtag gtggatgcca taatatctgc
48001 atccagttcc tgatcctcta ccgtgcggta atcggtttga gaagcatacc atctttcaaa

48061 gctctccttt gtcactctgc gcttgcccag gacatcgacc acatcaaaat gccctttggc
48121 aaccagttcg taagcagtcg cttctttcag cccaaggcgc tgtccaagct cctccattga
48181 atatgtagtt ttcttcagtt cctcgccggg aggggttcca tcgaccttct ggtatttgaa
48241 ctgattggca taccattctt caaagctgtc gatcatcacc ctcatggtgt tgccgacaag
48301 aatggttttg aagtaattct tcttaatgag ccagtatgac tccgttttac caagaccgag
48361 catcctgccc atttccggaa cagacatact cgtgcgtttt ctcatttccg ccatcgtaca
48421 acctcctatg taaaaaatgt agccatgtgt ttcttgtcat ggctacattt tatcgaactc
48481 tcaatatgtt ttgaagtttg tcgccgggtt gtacgacacc cttgacaaat cttacatttg
48541 atgatccaac cactcatcaa aactcttttt ggagattctg atactgccac cgatgcggac
48601 gctatggaaa actttctttt tcacaaggtt gtatgcgctg gttctgctga tgcccagaat
48661 atcctgaatc tcatcaaccg tataggttct cttatcaaac tgaactgcgt tagcagccat
48721 cgcttcttcg gtgcgctgat tcatgacagc gattctttct tcaaacattt atgttcactc
48781 cttcgctctc tttgctttat gttttgggga acgctttttc agctgttcca ttgccaggct
48841 aacggataag gatttctcaa tcgctaccat ttcacggggc gttaccttac ctaagtaatt
48901 atcaatcctg cttttatcaa tggtacgaat ctgttccagc tgaacaacgg acgcttcttt
48961 gaatgcagga ttgtcgtaaa taacaaagtg cgtcggcata ttcgctttct tatagattct
49021 ggtggtgacc gttgccacaa tcaatgtggg agcgtgttta ttaccggtat cgttctgaat
49081 gacgataacc ggacggatgc cgccttgttc cgacccaact accgggtcca gattggcgca
49141 ataaatatca ccacggcgat atacccaatt ttctttcatc atgtgccatg acctcctttc
49201 gttgggtatg taataactga ctgccagaaa ggaatcaggc agtcagtcca aaagtctatg
49261 tgtcccatat ttgccacgca cccctgggaa agagggctta cgccctcagg gaatcaccga
49321 acgactgatt gcgaatcagt tccataggaa tctaacctct gccgggttct ccgccagctc
49381 cgtagagaag tatctttaat cctgttgctt tcatggccgt attgggagca acccaatatt
49441 tacaagccag tattgatcgc tggcatcggg agagagacaa gcgcttaatt tatagaagaa
49501 agtcgttctg tttctctgtc cgatacgtct cggcgtaccg ctgatgtggc tgatgctctc
49561 gcaccggcaa caggaacgta ttcgatgact gtgtattcag ttttcaagga tcaagcagcc
49621 gattttcttt ggctgtatct aaattgtacc tgttcctcat gccagttttt attgcctcgt
49681 aggtatagtt ttgaggttat tcacgaccta aaaatatgta atcaagggtt gtttcaaagc
49741 gcacaaccaa ttcgattgcc aaatcaatag agatacctct gttgccggtt tcaattcgtg
49801 caatggtgtt tgcggcaaca cctaactgct ccgcaagctg ttcctgtgta agcccatgtt
49861 cctttctcaa ggctttaatt ctttttccgc tttcaaccaa gtcgtaatac attttcgttc
49921 ctccgtgttc ttgaatttgt gaagttgcaa aatcaaaaac agagggagga gatcgtgccc
49981 gattcgcaat gctctttgcc atctgcctgt cctccttgca ggcataaaaa aagagccgga
50041 gtaaattact ccaagctctc gttcatgcct ttgtgttaaa atgaaggaag ataggcaggt
50101 ataaaaaaag ccccgtaccc attacaggta cgaggcgtaa ctctctctat tcaattttat
50161 gctactatct cccagcatag ccattttaac accggacaag tcggactttc agtaggaaaa
50221 ccgtcgtatt ttcatcggat tttagtcgga ctaaaatcct gtcttagcac tcacaataac
50281 acaaggcttc ttcctccctg gcactttcca ggaaatcttt caatttagga atggctgttc
50341 cccaaagtga cagaccaaat aacagtatcg cttctttctt tcgatcataa tagctgctgc
50401 gttccatatt cagaacctcc agcatttctg attccctgta cttgaagcgg ttgagatatg
50461 cctttgagat aatctcacag tacagagcgc ctttatccgg atattctctg attttcaaca
50521 tagccatatc taccagctca atgatccact ttgtctcgaa taagctgcga atacgttctt
50581 caaatcgttc tcttgcttca tccggagcaa agttctccag ataaatcaat gcaccatcta
50641 agctgtcacc gtaagtacaa ataagggtat cacatacatc attcgcacga tcaatggtag

50701 accagcacac ctctcgatat atggataaaa tcagcttcgc tctttcatac agaaccttct
50761 cgtcaatgtt ctgcatccgg caaagcattc gaatattgct taatacctgt gtattatatc
50821 tgctcattca gaatctcctt tcagcctaat ccataaaata agaattcctt tctaactcct
50881 gtcacttggc taaagggaaa caaataccat gggacaagaa tgactataac ttccgatatt
50941 tgattaaaat aacagaagtg aaatcggaag ttcttgactt tataaaagtt ccgatatata
51001 attaaatgga actaaaactt ctgttctgat tctcattata acattttcca gaagtttgtc
51061 aacaaataat cggaagttac gaataaatat tttttgaaag gaaggacagt accatgacat
51121 ttggagaaaa aattaaagaa gctcgtttgg ctatgaacct atctcaaacg gagcttgccc
51181 agatgaccgg catttccgaa agatcccttt acacctatga gcagctcgga acactcccca
51241 gaaaaagcaa catcagaaaa cttgctgaag cgctgcatat ctctgtgtcc tatcttttgg
51301 acgaatctga aacagacagc cagagccata tagatcagga tatgtttatt ttagaagcaa
51361 aggaaaactt cggttcaaag ggagcaaaag aagctcagga agtgctgggg cgtgtaaatt
51421 ctctgtttgc cggtggagaa ttggacgagg acgcaaaaga tgtattcttc caggcgatta
51481 tgtcagttta tatggactct aagaaaacag cacgggaaaa gtacactccg aagaaataca
51541 gaaagcataa agataaagag taaaatcctc aagcacagaa tatgttttct gtgtgcgatt
51601 cggaggtgaa gcaaatgaaa acggcagagc atatcattga aactgtcgat aagcttgtgc
51661 gaaaatatca caccagagac ccttacgagt tgtgtcagct gttaggtatc aaaatccatt
51721 attacgattt acagaagaaa ttgaagggct tcttctatta ccagtccaga cagaagaata
51781 tcgtgataga tcacaatgtg aacggtatcc tggaacgtat cttagtcgca catgaattag
51841 gacacgctgt cctgcatacg aaaatcgcca tgatgcacgg ctttcaagaa atggaagtgt
51901 ttgatgatag atcaatcgaa gaaaacgaag caaacttctt tgcggcagag cttctgttag
51961 aagatgagga ggttctggaa tgcctttcgg aacacacatt ttttgaaact gctaaaatgc
52021 tctatgtacc ggcagcatta ttagattaca aattcagcct gctccatgaa aaaggggagc
52081 ttgtaaattc gatgtatatt cgtaaagcag atttcctgaa gggagatctt cacgcctacg
52141 ataatgacat tggctatggc gatatatgaa tatctatttt tatatcttta caaaaaagtc
52201 caccagtacg atgcgcagtg catcatgact ggtggacttt ttagttatca tcaagaaatt
52261 gttcagatca agcaaaaata tcatgtccgc caacaggcag aagcacaagg cgcttatttc
52321 agatattccg caaagaacgc tttcagcttt tcaaacggaa tgatgtccat ctgatcgtaa
52381 aggtcagtgt gatttgcacc ggggataatg agcaattcct tgttatcgcc ggtcagcttg
52441 ctgtacgctg tttcgctgaa atagcgggag tgcgcctttt ccccgtgtac cagcagcacg
52501 gcagagcgga tttcgctgct gtattgcaag atcggcatat ttaggaacga tagcgaggag
52561 gtcacattcc aaccgccgtt ggagttcaga ctacgggggt gatagccacg tggcgttttg
52621 tagtaagcgt aatagtcctt cacaaactgc ggagcatcct ccggtagtgg atcgacaaca
52681 ccgcctgcca atgcataagt accactcctg taatcttccg tgcgctgtgc attcatggct
52741 tttttcttct caaagcgtgc ctgctcagaa tcctcactgt caaaatagcc gttggcgttc
52801 acacgggtca tgtcgtacat ggtcatagca gcggtcgctt tgatacgagt gtcaatggct
52861 gcggcattga tcgccatgcc gccccaaccg caaataccga tgataccgat gtgctccgga
52921 tcaacattct cctgtacgga gaggaagtct accgctgcgc agaaatcctc ggtgttgatg
52981 tccggcgaag cgacatagcg gggtgtaccg ccgctctcgc cggtgtagga cgggtcgaag
53041 gcaatcgtca agaagccaag ctccgccatt ttctgtgcat acagaccgga ggactgctcc
53101 tttaccgcac caaacggacc gctgacggca atggcgggca acttgccctc tgtgcccttc
53161 ggcgtgtaca tatccgctgc caatgtgatg ccatagcggt tgtgaaaggt tactttcttg
53221 tgatcaacct tgtcactttt cgggaacacc ttatcccatt cctgtgttaa attcagtttt
53281 tccatattca gacctccttg cctgttgcct taaaatatac tgtgttgtcc acggcttcca

53341 accattcatt ggaggtttcg tcccctggaa cctctaccgc cagatgagag aaccagctgt
53401 ccggcgcagc accgtgccag tgcttgaccc cggtcggaat gtttaccaca tcgccggggc
53461 acagctcctg tgccggtttg ccccattcct gatagtaacc tcgaccggcc acacaaacga
53521 ggatctgtcc gccaccgttt ttggcgtgat gaatgtgcca gttgttgcgg catccgggtt
53581 caaaggtcac attgtaaatg ccgacctgct gcgtagaaag aggtgcgaga tagctttggc
53641 caataaaata tttcgcaaag tcatcattgg gtgcgccgat gggaaagacc atttcattct
53701 gatgcttggc ttttgcgtcg gcagcatcgt cctccgccca tacctccttt gccatacgga
53761 aggccgccca cgccttaggc cagcccgcat aaaatgccgc atgggtcagg atctccgcaa
53821 tttccgtttt ggtcacaccg ttgttctttg cagttgtcag gtggtactga aaggacgaat
53881 ccgtcagtcc ctgcgccatc agcgccacca ctgttacaat gctgcggtct cgcagagaaa
53941 gcttgtcttc tcagctccac acctgtccaa acagcacatc atcattcaat tctgcaaatt
54001 taggagcaaa ttcccccaaa gcctctctgc ctgccgtttg tttaactgcc atggctcaca
54061 cctccttgcc catctgatag gcttgctcca gtacagcatg acctgcgatt tctcccacgc
54121 tgtttacgcc gcctgcaaac actgtgccag ccagcgcaca gcgggggaag cagtctaccc
54181 agccctgcac ggctttctct gtacccgcaa aggtttccgg tgcgtcctct gccgccgctg
54241 caagcagata ggctttcgta aagacgtagt ccgtatcgaa caatgggttg gcacgatcca
54301 gcatagtctt gagctgaccg ctgacgcagt aatagtagac gggtgttgcg aacaccagca
54361 cattggcatc atgcattttg gcggcaattt ctacggcatc gtcttggatg acgcagtgcc
54421 ccaacttcag gcaggcaaga cagcccttgc agaagccaac ctgcttttct cgcagataca
54481 cagtttctac ctgatggccc gcctcccgtg cgcccttggc aaacgcttcc gccagcgttt
54541 cggagttgcc gcccttacgg ggacttgagg aaatgattag taccttttta ttcatcgtag
54601 aacactcctt gcatttttat attctttgtg ttattctcat agtagcacct aaacctaact
54661 ttaggtcaag atattttttg aaaattttg gaggaaaatt tatgtattca attggacagg
54721 tggcagagat gttcggattg cccatctcta cgctgcgcta ttatgataaa cagggacttt
54781 ttccgaacat ggaacgggta tccggcatcc gcaagtttag tgaggcggag attgaagcgc
54841 tccgggtcat tgaatgtctg aaaaaggctg gcatggagat taaggatatt cggcagttca
54901 tggattggtg cgtagaagga ccggccacct acccgcagcg aaaggcaatg tttgaggcgc
54961 aacgggtgca catggaagca gagcttgagc agatgaaccg gacgctggat atgctgaaat
55021 tcaaatgctg gtactatgag caggcaatca aagacggcag tgaggacaga ctaaagtcac
55081 tgattcctga ccgcttgccg gaggaaatcc gaaaaacata tgaaaatgcg catcgctaaa
55141 ttcaatgatt tcaactggct gtacgttccc agtgctacca aagtccaaaa agagcccaca
55201 agggagtgtc gcaaaataac tgattagtca gttatagagc gaggctcccc gtggactctt
55261 ttagcgttta gtcaaataga ttcatcgtcc ggaaggtcaa aatcatcttc cgtacatttt
55321 ctgataatat gttttgtttt ctccatcggc aaaggcacat tttcctctgc aaagtattct
55381 ttctttacga cttcaactac tgcttcgtga ttatctttac cagcaggaac aagcacaaaa
55441 tcgccgacct caatggtatc gtcatccgaa atatagtagt agctcttgta tccatcttca
55501 aactctacac tgcaatagat aatatcattg tcacagcgcc gtacctttcc atacacagac
55561 gggttcataa tttcacccca cccatagaaa gacataaatt caaagacgga ctccataaac
55621 tccgcccaat cgtccggaag cccctttta tcgtaggttc cttgaatgag cttttcattg
55681 ccctttgag aaataacctt aatggtgtag tctatggttt caagcgggtc aaccactaca
55741 ttctccgggt ttccttctat atggtcaaag atgctttcta catccatatc ttcaagaagg
55801 ctttcaaccc cgccctcaac tttgtatttt ctgcttacag aacacccaga accgatattc
55861 tgaatgtgtt caatactttc gctctctcta tccacaatta aagactctgt ataatcccag
55921 acagcgtatt ctgcgtgttc tgaagtagga acttttggct tgatctttgt gatacgatga

55981 tagtttactt cgattcgatt caccatatca ggcttattat tcccatcaaa ggcatacaaa
56041 tcaggcattt tcaagctgtc acgcagcaga tctgacaggt caattccgtc cacttcaaaa
56101 gacgagcata acgaaccgct gaacttatag acctttcctg cggtattggt cagctccata
56161 ttccagtttc caatatcagt tgcaaaaact tcatcgtacc cattgctgaa atactctgta
56221 aacgcagaaa tgattctggc ggcaaccgct ttatcaattt tcagattctg actgtgggct
56281 ttttcgtatc gaccatctct gcgctgtccg aaaacatacg aagaaaacca tacttctcct
56341 tctgcattga gggttatgtg ctgttcaact tcttcgtccg gttccgggca aggcccataa
56401 ccaagccgat tggaaacaag ccgaatcttt ctaagttctc ctttgaacag gaagggattc
56461 tcgccggaaa gcagggcaag gcggctcaat gccagaataa accatgaacg attttcagtg
56521 ttcaagatgg ctgctgcatc gtatgcccaa tgattaaaat atcgccattg ggaatagatt
56581 gccgacccaa gcagtttaat gtccgtgacc tcgtcaatga tacgatttag cgtatcacaa
56641 ctgcttgctg catcgccata tttttctgaa aaggcatggc cgcaatccat ctggaatccc
56701 aatgcctgac aatcatcagc caagtaatgg tcaaccagct ctatgtagct gattttctga
56761 tctctaaatt tctcaatcta ctttgtggcg aataagtgta cttgctctaa atactccatt
56821 gcgacctcct tccgaatcgg attaaaactc ctcccaagtc cgaccgttca gatcatttgc
56881 gattacatcg gtgttaggca ttgttttata tccgaacata tcagcaagat tgaagaactt
56941 gcttttttga atcgggttct tactgctttt tctaaagtcc atcgggtaga acttttcaac
57001 gatgaaatat ctatggctaa tcgtccgaag gttatagcca taattatctg cacggcggat
57061 tgcctcgttg atgcgttcaa tctcatctgt ggtaggttcg gctccgcttt cggcacgata
57121 agtcatgctg ccgttatttt caactgcgat gatggtcttt ttcagcttac caatggcacg
57181 gatactcttg tccttataca agccgatata tccatgttcc gagaacttgc gggattcctg
57241 gtcatagaac aaatccaaat ccatattgtc ctgaaaagtt gtaccggcta cgattgctct
57301 catccattta tagccatccg gtatcagctt ttcatcaaag caatacttct tgaaatcgtc
57361 taaaacggcg atgatttccg tatctctgtc gtcaacaata tcctccatag cgtccacaag
57421 ctgctcgaag gtcaagttga cgtgcttgat cggatggata agccgggttg cattttcgct
57481 gttatacttt ttcagttcaa tcgctaattc atcaaacagt tgcggtttca tcggcttggg
57541 gtcaagggtg agcaaaactt ttatatcctc agcgccaaac tgctccagat gatttacaag
57601 ctgctgctta taaaactgat tgtagagctt tgtttccacc actattttga agctcttttg
57661 gctgataact gcatctggaa cactatcgtc accaacgacc tgcaaatcaa aggtgatttc
57721 cgggctttca tcctctccaa gaataaggct gttcagcatg gaaaagaact tatcagcgtt
57781 atagttatac aaacgagaga gcatcagcat ggtgtttgag gtatccacat tctctttgga
57841 gtggtatctc tgaaaataat gtatctgcat atcgcacctc ctaaaaccca tctgccggtt
57901 cgtttgcgtt accgggtcag ttgataactc aattccagca agggcttgat ttcatctgcc
57961 gagactgtac catcgagaag aatgctgatc caattctctt tgttcatgtg ataggcgggg
58021 aagaatccct ctttcatacg cagtgaacct atcatcatcg gatcgcactt gatgttcagg
58081 acatcgaccg ttttatcttc tgccaatccc agctttcttg cattgatttc aactaccaac
58141 gcaaaccact ttttgttatt cttatggcgt aaaacgcaat tccagtcatt ccacgggtaa
58201 tccggttctg cgccatagat ttcttttgcc cattcaaata gttcttttcg ttccatcgtt
58261 tccatcccta tatgataagg cgatacacca tagattaggc ttccatcacg ttatctgtcc
58321 aaagctcgca ctgtgtcatt acagtctgaa cagcatcatc catacccctcc ggcggatatt
58381 tatgtttctt caacagtcgt ttaatgagag tacgcatttt ggcacgggca gactctttac
58441 gctgccagtc aatcgtttta ttcttgcgaa gtgtatctgc caattctttt gtaatcgcta
58501 tcagttcttc attttcgtaa aaatccttga ttgcttgcgg tttcgtaagt gcatcgtaga
58561 atgccaactc atcagctgta aggccgagct gttcgccctc tttgtgggct tctgccagct

58621 gttttgcgag tttcagcatt tctgcaatga cttcttcatt ggtgagcata ccattcagat
58681 atgcgttgat agctcgctgc atgatctcgc tgaacttttc ggacttaacc acattcgttc
58741 tacggtaaat agatacctgt tccgctatca gtttttcag cagctcgatt gccaggttct
58801 tctctttcat tttactgatt tcctcaagga acttcggatc aaacaaagaa aattcctctt
58861 gcatatcaga gaacaggtta ataacaccct cgctcttgat gctctgcttc aaaagctcat
58921 tgatacgagc gttcatttca ggcagagaga tcttcttacc ggttcccgca ttggtcagcc
58981 gcagcaccag gacacgcacc gactcaaaga acgcagcttc aaagcgatca gtttcacaca
59041 ccagagaaga acaaagagaa agagcctgat gaagcatgag ggcttccctg ataaaggtgt
59101 ctttttcttc ttctttttca cgggcgatga tatagtttac cgcaccgctg atggtctttg
59161 ctctttccag atcggttccg gtagtgaact tggtgtaatc aaatccatgg aacatatcac
59221 ggcagacacc gagcttttcg ataaacttcg gataagcgac tttggcaacg tccgtatcac
59281 catagttttt cttatcacgg gtggtgtagt cattcattgc ctgttttaat gcggtagcaa
59341 tgcctacata gtcaacgacc agaccgcctt ccttgtcacg gaacacacga ttgacacggg
59401 cgattgcctg catcagatta tatccgctca tcggcttata cacatacata gtagccaggg
59461 acggaacatc aaatccggtc aaccacatat caacgacgat ggcaatcttc atcgggctgt
59521 tgttgtcctt gaatttttc gccaactcgt ccttgtgtcg cttattgccg ataatatcat
59581 gccattcttc agggtcgtta tttccgccgg tcataacaat accgactttt tccgtccagt
59641 cagggcgcag ctccagaatc cgcttataga tcttcattgc aatcgggcgg gaatacgcca
59701 cgatcatcgc tttgcctgta agcagacctt cacggtaatt ctcgtaatgg tcaagaatat
59761 cgctgaccag agagttgatg gtattatcgt tgcccaatac ggcttccatc tgtccgagag
59821 ttttcttgct cttttcaata acctccggat cagcattatg tgccatgatt tcatactctg
59881 tatcaatcat gtgcagcgta gcttcatcca ggttgagctt aatgacacgg ctttcgtaat
59941 agaccggacg ggttgcgcca tcttcgacag cctgcgtcat atcataaatg tcaatgtaat
60001 caccgaacac ctcacgggtg ctgcggtctt tcgaggaaat cggagttccg gtaaagccaa
60061 tgtatgtagc gttaggcaag ctgtttcgga taatgcgggc agtaccaaca acacgcttgg
60121 cgatttcttc gccgtcctca ttcttggtca tcttaatctt ttcggttaag ccatactgtc
60181 cacggtgggc ttcgtctgcc ataacgatga tatttctgcg ttcagaaagg gcttcgtgag
60241 attcctcaaa cttctgcatg gtcgtaaaga taataccgtt tgcctgacga ccggcaagga
60301 gatcacgcag attctcacga ctctgagcgt gcatcgggtc ttgacgcaag aaatccttgc
60361 actttgcaaa ctgtccgaaa agctgatcgt ccaggtcgtt tctgtcggtc agcacaacaa
60421 tcgttgggct atccaaggct tcctgaagca gatgggcata gaataccatt gacagcgatt
60481 taccgctgcc ttgcgtatgc cagaacacac cacccttacc atctgtaacg gtagctcgtt
60541 tggtcgattc gattgccttt cttacggcaa aatactgatg ataaccagcc agtattttga
60601 aggagttcac accttcgttg gaaaagcaga taaagttctt tatcaggtca agcagtcgtt
60661 ccttctggaa catacccctca aagaaggtat caaactgtgc gtactgcgta ttctcgtagc
60721 ttccatcgac cgttttccat tccatgaagc gatcttcacc ggaagtaatt gtaccggctt
60781 tggatgtcat ctggtcactc ataacgcaga tggcattata gatgaacata gaagggattt
60841 ccttcatgta gttgcggagc tgccggtatg ccgcagaagc atctgtttct tctctggatg
60901 gtgatttcag ttctacaata acaaccggta taccattcag gaaaagcaga atatccgggc
60961 gtttgttgct gttttcgatg aaagtccatt gatttgctac gataaaggag ttgttgtcag
61021 gattcttgta atcaacaaga tagacgatgg aggagcgttc ctcgcctttc tcaaaatagc
61081 gcacaggaat accattttgc aggtagtcca taaaaacagc gtttttctgc accaactcgc
61141 cgttttcaaa gttcttcagc ttaaaaagtg catcactaat agcgtcctcc ggcagagaac
61201 ggttcaaacg atatagggaa gcaacaagtt cttcctcata cagaggattg cggaaatccc

61261 tctctatgtc gggtccatac aggtaactgt accccatgtt ccggaataat tcgattacgg
61321 agttttcata gtccgcttcg gtatataatc ctggcataat atcgctcctt ttctgtggta
61381 ttcatactcc atttttatat catagcattt tctttatata ttaggtcgct ttgccggcga
61441 cgatttccgg gtaaacgaaa atttagcggc ttaaatgtca aggctggaaa catccagttc
61501 tccagacatc aaccgaggta gcaaagtatc acgagtggcg gcaagtctat cattttcagc
61561 acgaagattg cgaatctgcg caaaaatcgg ttgtacgatt gagttgaaag cactcaattc
61621 cgcttccgat ggtatgacga cagacacttt gttcagattg ccttggctaa tctttggctg
61681 aacggctcca gtaacaattg actggacatt ggttaaactg aacagcaaat aaagcgtttc
61741 gacagtaaag ccattcttac ccgtgataat gtgggcatga ttgttcaccc agaactttcc
61801 ttccacatat tgcagaattg gaaaaccctt gctgtctaca acagtgccat cttcaccaag
61861 gagtagatag attccgtcaa agagatacct atccacatag tccattactg atgtagcacc
61921 atagtacgga taaatctttg cgaggttggc acgttcacgg ctcgaaaagg gtacgcgttt
61981 tgaatcgtgt agttctatga tttctgaaac tgtgccaagt cgccaatccg ggggcatctc
62041 accatcgaag ggtttcaaat caacaaaacg atccttataa agagcctgtg cctgctgaag
62101 taaattttcg tttatcgcat tatttatttg tattttggca tcaatcgctg agcataggtt
62161 tgtaattttt ctttgctcct caatgcttgg gaaagagaag ggcattctct taaaatcttt
62221 caatacaatt cttggtatcg cagagcctga accgtaatta tcacgaacat ctttccttac
62281 ggctggcgat tttaatgtat agtataagaa ttcgggcgtt atcattctgc aatcagccct
62341 caatattgct attgaagata ataatgccgg gcgttcatct tgcttataaa tgataatatc
62401 ttcaaaatat cttgccccgt cttttcctac caaaatatca tttttctctg ggacacaccc
62461 ttgattccga agtttttcat aatcttcttc gccaatctgt ttgcaatttg agaagtcaaa
62521 accaaattcg gtaaaatctt taacagaagc catataatgc ccttgttcta cacttttagg
62581 gctattatgt gctccgtcag tcacaacggt acacacatca tctatcgttg ctgtaatcca
62641 attagatttc atagccaatc gcccccagct tcttacggat ttctgcttcg agttcatggg
62701 acttggcaaa catattagaa agctccgaag tcaggcgagc cattttttca tcaaacggtt
62761 cgctatcttc ttcctgttct tcaatgccaa catagcgccc aggagtcaga atgtagtcct
62821 gcttttcgat agtctgcaaa tcggctactg cacaaaatcc cttcacattt tccaaggttc
62881 cgttctggaa tgcttcaaag gtgtccgcca gtttctgaat atcttcatca gtgaagtctc
62941 tgtgtttgcg gtcaaccatg tagcccatct tgcgggcatc aataaacagc gttttgcctt
63001 tctgcttttt gttcttgcta atgaaccaga gtgtaactgg aatcgtcacg ctgtaaaaca
63061 gctgcgtagg cagggcaact ataccttcca ccagatcggc attgataata ttctttctaa
63121 tctcaccttc gccggaggac tgcgtagaca gtgcgccatt tgcaagcacc agaccaatct
63181 tgccgttcgg tgcaaggtgg tggatcatgt gctgaatcca tgcgtagttg gcattgccgg
63241 caggcggcgt tccgtacttc caacggacat cgtccagcag tttatcctgt ccccagttgg
63301 acagattgaa aggcggattt gccataatga aatctgcacg aaggttagca tgaaggtcat
63361 taaagaacgt gtcagcctga tacgggccga aatctgcatc aatgccacga atcgccatat
63421 tcatctttgc catcttccag gtatccgcat tggattcctg accataaacg gaaatgctct
63481 gacgattgcc gctgtgtgcc tggatgaact tcacgctctg aacaaacata ccaccggaac
63541 cacagcaagg gtcgtacacc cggcagttgt tgaacggctt caggattgca acgattgtct
63601 taacgatgct ggacggtgta tagaactcgc cgcccttcac accttcgtaa gctgcaaact
63661 gtgcgatgca atactcgtaa gttctgccca gaagatcttt gccttcctct gtatcagaca
63721 tatccatatt ggtgaacaga tcgacgacat cgcccagaac tcgcttgtcc aagtccgggc
63781 ttgcgtaatt cttcggaagc acatttttca aagatttgtt ctctgcttca atctctcgca
63841 ttgcttcatc aagaactgtg ccgatttccg gagtgtgagc cgcagctgca acggtagacc

63901 atcttgcttt ctccgggaca aagaaaatat tctcgcctaa atactcgtca cgatcttctt
63961 caaagccttc gccttctgcg acaagctcgt tatattttct ttcaaatgca ctggaaatat
64021 agcgaaggaa aatcagccca atgattacct ttctatattc agcggcaggg atgtgtcccc
64081 ataacacaca ggcagcatcc caaatttgtt tttcaaagcc aatgttggca ttgttctttt
64141 cagccatttc cgattcctcc actctaacag tctattgtag ttatgcttct tcagcttata
64201 ttctaccaaa gaggctatcc cataaaacgg acatcttact tttcgtccgg aactatttcc
64261 ataatatcgt caaaggtgca gcccagggta gaacagattt tcacaagcgt atctgtagtt
64321 acattctctc ctcgatgcat tttattgatg gtatagttac tcagatttgc ctgttcagcc
64381 aaatcttttt tcttcatatc tcggtcaatc aatagcttcc atagtctctt atagctgaca
64441 gccatcattt caccattcct ttcttccaga tattttgttc aaaaagtacc tgtacttata
64501 attatagcac gcaaataagt aaactcaaca agaaagtgag ctatcactat attttatttt
64561 ctcattccaa agaattatcc gattttttaa gtgacaaggc ttttagagca tcgagaagcg
64621 gaattttggg gttttacaaa ttggcaagca gtgtaaacct gtacaggttc acttattttc
64681 tcgattttcc ctccggtgca atcttcgcaa atttgcttgt tggagaagca tccccaatcc
64741 cctttgaacg gtatctgcga taccggctac tcgttcattc tgaacgcttt ctggagaaag
64801 aaaaaacctg gactaccatt tgctggggtt agatagcgat actttttaca tctaccatgt
64861 tgcggcggtc atggttgaca acttgcgtcg gctcctgctc ggtatcaacc tgtccgacaa
64921 agctataatg gatttcgatt ttccgggtgt tggtttcccg gtctaattca tgtatcaaaa
64981 tacggtcgat aaactcatgg acgttttcat aggtcaattc ctgtatgtcg ctgtatttcc
65041 cgacaatctg aataaacctt gatatatccc tttttcgctc ggtaacggtt gcgatctgct
65101 gttgtaactc gtctatcctt gcggcaaggg atttcttttc gtcctcatag ccggaagtta
65161 gaaacacaaa ccgttcatct gtcagtctgc ctaatgcgtt gtcctcatac agcttgcgga
65221 aaagcgtgtc aagttcggtc atacgcgcct gtgctttgaa aagttccttt tgctgctgcg
65281 ctaatgcctt tctcgcttcc atgtcgccgt actcggtagc tttacagata aagtcctgtt
65341 cgtgttcctt aacataccgc aaaacccgcc gcatatcttc cagcactaaa tcaagaagta
65401 cgtttttgcg gatataatga gaggtacact cgctgcctgt ccttgccctg ttgcgccatg
65461 cgccgcaaga ataggaaaac ttgcgcggct cgatatttac gccctgttgg taatacatct
65521 tatgtctgca accagcgcaa aacagcaagc cggaaaaaat atcaatctcc gctgcttttg
65581 tcgggcggtg gcgggtagca atccgctttt gcgcaagttc aaaggtttcc tcgtcaacaa
65641 gcggctcatg ggtgtttggg aagaaatagc gttgttcttc ttcgttcttc cgggtctttt
65701 tcgacttgta ggataccttg tgagttttcg cggttatggt atgccctaaa tattcctgcc
65761 ttgctaatat gtcatagagc gttttgtccg gccaagtata gggggcgatc attgcccgct
65821 gataccgcgc ctgtcctgta cgctgatagc gcaacgctcc aatcgtcaag actttgtttt
65881 ccgcaagcca tttttggatt tcgcacatac gcacgccgct aacaaacata gcgaaaacct
65941 gtttgacaat cggcgcggtt tccgggtcgg gtataaggtg gtggcggttg ttcgggtctg
66001 ggatatagcc gtatggatat tccccgttga cgcgctcgcc tttttgcgcc tgtgcctgtt
66061 taactgcgcg gattttcttg cttgtgtcgc gggcgtaaaa ctcgttaaac cagttccgca
66121 agggggtaaa ctcgttatcc tcccgcgctg tgtcaactcc gtcgttgatc gcgatatagc
66181 gtactccgtt ttcgggaaat acaatctcta tgagttcgcc cgttttcaga tagtttctgc
66241 caagacggga aaggtctttg gtaatgaccg tcgccactcg tccggcttcc acttcctgca
66301 acatcgcttg cagcccctcg cgctcaaaac tcacgccgga aaatccgtcg tccacaaaaa
66361 actttgtgtt caaaaaatgg tgttcgtccg catagcgttg aagtatcatt ttttggtgct
66421 gtatgcttcc gctttctccg gcttgcatat cctcttgact taaacggcag taaagggcgg
66481 tgattttatt cgactgtaac attagtcctc ctttccgaca gccgaataaa caggtataat

66541 gtgttgctat cattatacca tatcccgccg cctaatgcac tactcggacg ctaaaaatcc
66601 cggatttccg ggctttttct gcaaatcctt tacaatgagt ttctcaatct ttttatggat
66661 tgtatcggtt gccttttcac tcggcaacgc ttgaacaaga taggtgattt tccctatttt
66721 gcgttcggtt gtaattgtct gttttttatc cattagaaaa acctcctttt cctgtatgga
66781 taaactatat tcgtcaaaag gcaaaaacgg gcggttgtta gctgcaacct cacgggagtt
66841 tcaccccggc ccatgcttat gggtgcgccg cgcaatactt tgagggtgtt caacgcggga
66901 gtatcattgt gccgccttgt atgtcgtcgc ccacaagctg ctaaacctgt tttacggcgc
66961 ggtagttctc gctcggcttt tcccctatgg ggaaaagctg tcatggcgta cccgccgact
67021 ggctcggtac agacggaatg tacccgtttg cctgttatgc tgcgcaccaa aggccgcttt
67081 ctttgtcaaa gaacaatagg ctttgtcggc ctgcaaaagc acatcaacag cggatatgct
67141 tttgcggaac gacaaatagc ccataacggg aagcgtggaa aggggacacg ctccccgcat
67201 gggctaaaag attatcctac atgaaaagca agggtgcggg taataaggcg cacttgcagc
67261 ctgttacgca tttcctcgtc cacataggaa taggtattgc ccgcgtcgtc tttcagcgtg
67321 cgggtacaaa gtttcgctat gtaaccctcg tagtgcttca agatcgcgca catggcggtt
67381 gtgtcgccgt ttgccgctgc ggaaatgaca gggaacggca acagattttc agacttccta
67441 acggtattca tcatctgctt ttccctccaa atactgtttg attttcgcaa gcgcggattt
67501 cctgtgccgg aaaaccgtcg tgcgtacaac attcagcagt tcgccaattt ccgcgtcgct
67561 catatccaag aagtaggaca aaaggataat gtcgcgtttc ctttcgggca aagcgttaag
67621 ggcttcggca agcagttcat ttttgacgag tacatcaaag ccggacactt gaaaacggaa
67681 ataatcgctt tcgtattcgt ccgttgtgaa aagctgcgcg agttcgcttt cgctcaaatc
67741 cgaaaaagta acttcgcgtg ctgcgcgttt cgcaagagtg cggcggtggc ttttcgcttc
67801 gccgaccaaa gttttctttg ctaatgcgtc gtactgatgt tgtattcttt ccttgtcgga
67861 agaagatagc tccatagagt tcacctcctt cccgcgtgga gtagaggacg ggagttaagg
67921 gcttgtcctc tctgcccctt tcgctccgta cccgttcggg agatggctct tgagtgcgct
67981 tttcagaaaa aagttgaaaa aagcaaaatg cgcccgtccg caagacgcgg acgggcgcaa
68041 aggaaatgta agcagtagct aaatgtattg acagttcaca ttcatagccg gaatatcccg
68101 cttgcggagc atagcttttt ttgaccgcaa agcattaggc gggttacagt aaataaaaat
68161 ggacacggcg atacctcccc gataccgccg tatccttaaa aaagggcgac tttaatacac
68221 tacccgcaat ccattctata atagggaaca gcggcttttg cgcaatatta aataaaaaag
68281 ccgataacac ataggttttt tgacctaatg cgttatcggc tctgcgtcta tgcgtctggc
68341 acttttaatc attttttttg aatttattat atgtgttagc taactgtcca caagccgcgt
68401 taatctctct gccatgagaa actctcatag taacttcaag tcctgcttgc tctaattgat
68461 gtttgaatgc aactatttcc cgtttctgtg gtgctttaat tcttgaattg cttgttgggt
68521 tgtattgtaa cacgttaatc ataacttttt tgccccgaaa ccattttgca agttgtctta
68581 catctgaggg ccggtcattt atacccggta aaagcaaata cgcaaagaca attttgcgat
68641 tatgcctttc agaataggat aaggcttgct taacaacatc ttcaatagca tatatgcgca
68701 tgtgaggaat aatacgattt cttgcagatt gtgttgctgc gtgtaaagat attgtcaact
68761 gaattttaag atgttcctcg cgcaattttt ttaattgatc gaccggacca actgttgata
68821 tggtaatgcc gtcggttgga aagttgagcc catatctatc tcggagaata tggattgctt
68881 ttatcaagtt gtcataattg aataaaggct ctcccatacc cataaaaacg atacggttta
68941 cttttcgacg caacaatata atctgttgta cgatttctga cgatgttaga ttacgaacaa
69001 agccatttcg cccggactca caaaaaatac aaccaacagg acaaccgact tgtgtgctca
69061 cgcaaacagt tccaccatct cgccgcttga taaaaaccgt ttcaatgtat ttgttgtctt
69121 tcagttcgta aacatacttt tcagtatcac tgcttttgca aatatttttt accaacattg

69181 atagattttt tttgcgtggt aattgcttat ataattcttc atataaggct tttgcttcat
69241 tctcgccaat aacttccgac atttctttgt aagtaaatcc gtatggatca ttccgtactt
69301 ccgcaggcgt atatttaggt aaacgtttca tttttatatc ctttcttcta aataataaaa
69361 gtttgttttt ctgtattttt gattacaatc tctaattttt ctacatcaat gatatgcgtc
69421 aatttgcatt tagggcagaa aagaggaaaa tcttttaata cagtattatg aaatacttga
69481 actctcgtct ttccgttaca aatcggacat tttatccaat attttggctc tactccaaaa
69541 gtagtggata agtacctttt gtgatatact tgtcacgggg tgatgaaaca ggtggaaaaa
69601 aacttcacag cgatgattgc aggttctctc aatctaagtg aaccgtggta cgtggaggga
69661 gctgagttcc gtccggaaga gcaagaagtg catatctacg tcaatgtcag aaagactgca
69721 gccctcgtct gtccatgctg tggcgggcct actttacgta acggctatga gccaacagag
69781 cgtgtttggc ggcacggaga ctgcatgttc tatccgacct acgtccattg caggcgtcca
69841 cgggtgaagt gtccaaggtg cggcacgaag caggtgaacg ccccatttga gcggaagaac
69901 agccgcttca ctctgttgtt tgagggctat gccatgctga ttcttgccga catgccaagg
69961 gctcgcgcag cagcagtgct tcggtgcgat gagaaatctc ttgcctctat actcaactat
70021 tgggtagacg atgcggtgag taatcgctca ctgagcggcg tgaagcggct cgccatcgac
70081 gagacctctc gccttcgggg acatgactat gtgacagtcg tgatcgatgc cgaccaacgc
70141 tgtgtaatcg acgtagaggc aggcagagat aaagccacgg tgtccacctt cgctgagaaa
70201 ctggagatgc gtggtggcaa tgcggcggcc atctgtgcgg taacaagcga tatgtcgaag
70261 tcttacctgc cagccattga ggaaaacttc cctaacgccc tgagcatcat cgacaaattc
70321 cacgtcaaga aggtgctgat ggacgccctt gacgatgttc gcaaggagga gcagcgaacc
70381 gctgcgtcca agaaagagct ctttcttgga aggcgtctat tcatgatccc caaagccagg
70441 atgaccagcg aacaggcgga aaaactggga tctctgtcca agctgtatcc caagactggt
70501 aaggcttaca gcattgtagc aacccttgat gacttctata cctgccaaac ggtggaggaa
70561 gctgaacagg ctttcgaccg cctctactca tggatgcgtc gttgccgttt gaagccaatg
70621 aaagaggccg ccaagacact ccggaaccac aagacgaaga tactggctta cttctaccac
70681 cgcataacga atgctgtctg tgaggggatt aactccatga tacaggcggc aaaacggaag
70741 gctcgtggct ttaatacatt cagaggattt gcctcgatga tataccttgt tgctggcaag
70801 ttacagctgg ctgtgccacg cccctttga cccttatcca ctaaaattgg agttgaacca
70861 ttattgcgta ttgtcaatct ctctcaaacc gggtagtaaa aatacggcaa gcgcaacgat
70921 gataatgcca attccgcaaa tgacaaacca tttctcaact cccagccgct ccgccagagg
70981 cccggaaatg acaaggccaa acggcatggc gagggaagcg gcgcttgtca gtagagaaaa
71041 aactctcccc agatattctg gtttgaccgt ttcttgaaaa atcgcatttt gcacaccgta
71101 aaatggagcg gaaattccca taacagtaca gcacacaaca aagacaagaa atgcgtctgg
71161 cggcaaaagc ccggagagca tattgctaac gcccatcagg agaacggaaa gaccgatggt
71221 gtaccgccgt ttcttaaaac cgccccaaat gctcagaatg actccgccaa gcagcatacc
71281 aaccgcaaaa gcaatttcag cggcagaggc atgggccggt gttcctttga agtatgacat
71341 acagatgaga ggaaaaagcg tactgattgg catataaaag aacatataaa ttacaccaat
71401 ccagagcaga gcaaagaggc ccctgttttg ctttaatacc acatacccct ctttcatatc
71461 ctgtaaaaac tgttgtcttt tcgtttctgg acatagttca ggcgttggta tggacgaaat
71521 cgcaacagtc acacaggcaa ggattgcacc cacaatatct aacaatataa ttgcattaag
71581 aggccaaaca gcatataaaa acgctgcggc agctggactg ataatcgcac ttactgcttg
71641 catggtctgc gtgtaaccag cgcattttgt aagttcctct tttggcacaa tcataggtgt
71701 tgctgcgctg aatgctggag aatgaaaagc agttcccgca cttcggatta acaggacaac
71761 cataatagac catacgggca attccatgta aaacgccacc agcgccagaa taccgccagc

71821 ggcggcaatt atcaaatccg caccaatcat tacgctttta cggctgtgcc ggtcaacaaa
71881 agcacctgca aacgggccta aacaggcttg cggcaaaaat ccaatcagtg ttgccgccgt
71941 caatatgatt gcagaattag ttttcgcaac caaataaaag ataatggcca tttgcaaaat
72001 accgctgcta ataaaggata ttgcttgtcc ggcaagaagt gtaaaataag tttttctcca
72061 tgaattgttg ttttgggtca taatgcgaac ctcctttttt attgcattgt tcctttgttt
72121 ctgcaataaa aagcaggcgt tgtcccacag agagggcaga cgcctgcata acaacaaagc
72181 acgaaaaaac accacgatac agttcaaaga ctgctcgtgt caaacctacg tgttcctttg
72241 catacgcacg caaaaaaagc ccaccatcat gtggaaaggt acttctactt tttattgctt
72301 attagcgtac acaaattaac acacgtaagc ctccttccaa tctcaaactg tcgcacagta
72361 taacacacat ccgatagact tgtcaaccat ttttaacctt gttttccatc ttgtttttcc
72421 atctcttacg ggcagtagca aagccaggcg agccagtcaa cggtcaagat gaacggcgct
72481 ttcagcgccg ccgttgacag tctcgcccgt ctttgctaat gggtaatcaa ggcgggaaag
72541 ccattttagg gctttcccgc ccatttcaat tttgggagaa gaaaggcccc aaaatgaacg
72601 agtgcggcca aacactttta gggattggcc accttgtcca cccatccagc ggggcggcgg
72661 ggaacggtca aggccgggcg tcagcccatt catttcagcc ttgacggttt cctgccgtcc
72721 tgctactttt cccggagtgt ggccacacat ctggtcacgg tgtccagagc tttgggactt
72781 tttgtcccat aggccggggg cggaggacga gggacggggg ctttcataat acgccctgtc
72841 tgctgctgaa atcagccctt tgtttccggc ttaccagccc caaacaaagc cctgctttcc
72901 tgccgcgtca aatgcccttg ccctccccgg cggcaacggt attttcaggg caacgccgac
72961 agagcgtata acacactaca ctttgctccg caaagtcgtg tgccaaatgg ggcgctgccc
73021 cctttggaaa cccccgcaac aaacaggtgc tatgcaccca gccgggagca tagcgccgtt
73081 tgttgtcagc agcccgtttc acggtctgcg tgtagttcct tgtaaactga cctaaaggta
73141 atgataacgc ccgcaatcgg cacaatgacg ttttccgata ggctgcgtat catattgaat
73201 atgcccgcgt tccacccggc gggggttgtg ccgacttgcg ttgcgatttc tccaacctct
73261 gtatttacgc tgtcgaacag cccggagaga ttagagagta ttccgtctgt caaaatgccg
73321 cgaaaccaat cttcaatcat ctgccatatc aaaagctaat ccctcctttc ccgcgccctc
73381 ccgcgtgaag cgggagagcg cggcgggtag tgttagtttg tggcattatc caaacaggcc
73441 ggagagcagc gggacaaggg taaggccgat aagggcaacg ccgccgcccg ccatcaactg
73501 tttcatgccc tgcgacttcg cgccggggtt atcgttgccg taccсctcca tgaggttaat
73561 gccgccccaa atgccaagtc ctgcgccaag cgcgattacc aaagtctgca atacgtcaac
73621 tgcgctgttg aaaaaatcca tataaacctc gctttctgcc gctaaaacgg ctcaaaaaat
73681 gttgttggaa ataaaaatgc cgccgttttt attcggcggc tgcgtcctcg tcggacaaat
73741 ctatttcgta tatgtcaaag ggttcgtcgg gctttacgat tgccggacgg gtggacatat
73801 accgttcaac gtcaaaagcg tttttcttgt cgaagtcgga aaggtatttg tagttcgggt
73861 gctgcgttat atcgtactta tcgctgaaaa aggggcgcac gccgcgcaac tgcaaaatac
73921 attttcctcc gtccataact gcgatttcgt cctgtgtcat caactccttt cccaatttct
73981 gataattgag gccatgagaa acctgcgtgc cgcgattttc ggaagtgttg aaactgtcga
74041 ttgtttcacg tcccaagttc tccgaaatat cctttgcatt tttcccgcgc cctcccaaaa
74101 acaaggtgct gtcggcattg tcaagtatga tttctgccgc gtccttgtaa atggctttta
74161 gctggctctg cgattgcaga atgatagaag ccgagatttc ccggctgcgg attgtggcaa
74221 tcaatttttc aaactgcggg atttgtccaa tgtttgcaaa ctcgtctaac aggcagcgaa
74281 catgaaccgg cagcttgccg ccgtattcgt cgtcggcctt gtcgcacaag agattaaata
74341 actgcgattg cagcatagcg ataacaaaat taaacgtgct gtccgtatcg ctcataatca
74401 aaaacaaagc tgtttttctg tcgcctaacg tgtcaagttc taattcgtcg taggacataa

74461 gatcgcgcaa ttcctttata tcaaaaggag cgagcctcgc accgcaagaa atgaggattg
74521 attttgcggt tttgcccgcc gccaatttat attttcggta ctgcttcacc gcgaaatggt
74581 cggggtcgcg ttcttccaag tcggcaaaca gcaaatcgac ggggctttga tattcctcgt
74641 cgtcctcgcg ggcttcgctg gcgttgataa gttccaacag ggtaataaag ttcatttcct
74701 cggcgggggc ttcgtaccag atgtagccca ccaacgcgca atacaaaagc cgttctgcct
74761 ttacccaaaa atcctccgcg ctttttctc cctcaccttt ggtattcgct atcaacgtat
74821 ttaccagctt caaaatatct ttctcgctgc ggatatacac aaaggggttg tagtgcatag
74881 attttttgaa gttaatcgta ttcagtacct taatgcggta gcccgcccgt tgcagcagct
74941 tcccgcactc aatcaaaacc gtacctttcg ggtcggttaa gacgtaagag gaatgaagct
75001 gcattagatt aggtttgaca aaaaatcttg tttttccgct gccgctgccg ccgatcacga
75061 cgacgttctt atttctcgcg tacttcggct gcttcgggcg gctgttcatg gtaagccgtt
75121 ccgtctgcgt caaaatgatg ttgttttgga aatccgggtc tatgtagggc tttatatctt
75181 cggcgtttcc ccatgaagcg ttttgtcaag tgcttttttg agttattgac gcaaattctt
75241 gtgaaagtgc gaaagtgcaa agtgcaaagg gtctgcgttt tgcactttca gcttgcgggt
75301 tcgggcggct gcttcttctt gatgaagtta taccattgac cgccgacacg ccttgcgccc
75361 aaaatgctgc ccatgttgcg cttggcgttc tgtaccgtcc tctcggatat tccggcttcg
75421 gctgcggcct ttacaatgtc ctcgctggca agctctttcc cgtctgcaag caggtcaagt
75481 atcagcttct cggcttgttc ggtcttggtg gcggtgttcc cgcccgcgcc ggatagcagc
75541 tcgtcggcgg ttatgtcgta ttcgcctatc catgaaaagc ctgtttccgg gtcaaggcaa
75601 aaggcaacgg gcttgccctc cggcgccagg gaagatttgt catggacgat aacacgcaca
75661 ttcggctccc gcttcacgcg cccgatcagc aggacgctcc ttgctgcggc gcggaagtcg
75721 atagaaccta agccccggta tgcgctctgc cctccggcag ctttgttcaa gtgtccgata
75781 aggataacgg cgcacccggt acgctcggca acatcggcaa gacggcggaa catggggcgc
75841 acttcgtttg cccggttcat gtcggtcttt tcgcccatgt acgcctgtat tgggtcaagg
75901 ataatcaacc gcgccccgtt ctgcgtgatt gccttttcta tgcgctcgtc ggagagggta
75961 agctcccgct tggcttcatc aatcacaagc acgcggtcaa ggtcggcttc ggcttctatc
76021 aagcggggct tgacggtatc gcccagcccg tcctcggcgg tctggtaaat cacttggaat
76081 ggcggcaggg gcttcattcc cggcagcgtt cccccggtgg tgcaggcagc ggcaaggcgt
76141 agggcaaagg tggtcttgcc ctcgccgggg ttgccttgca caatggttac tttcccgaaa
76201 gggatatacg gctcccatag ccattcaacg gtctgcgtgt caacctcgct catgcggata
76261 atctcgacgg tttcttcctg cggcggctct ttcaggccgt aaactgcttc ctggatatac
76321 ttcccgtctg tgatttctgc ccggtgctgc aatacctcgt tccagtcctt aaacagcggc
76381 acaaggcggt ggacggtcaa gccctccggc acaagctccg caaggcggct acaagcgtcg
76441 tttccggctt ggtcgctgtc aaggcagagg taaacggtct tgatgttcgg acggtcagaa
76501 aggaaacgca atagggcttt ttctcccacg ccgcccaatg acaaatagct ctgcttctgc
76561 cattcctttt tgaacaggca gaggaaagag agcaggtcaa cgggggcttc aaaggcaaaa
76621 agcctgtcgc cctcgccccg gtaacagaaa ttaaaggctt tgtcgctgcc tttcacatcg
76681 agccggaagt ttcccgccgt tcccttgctg tgggcgtagc gcggtattcc gtcctcgtcc
76741 cgccccacaa atacggcgtt gtggtgggct gcgtcctcgt aaatatcgcc gcgggcaaag
76801 aaaaagcccg tcacatcttc atcaatgcgg cgggctgctg tgaggtagtt cctcgctgtt
76861 cggttgtcgc tgtttggggg tggtagccgg aagtcggaaa gggacgcggg gcaaggtctg
76921 tccggcggcg gtgcggctcc cttttctcct gtgagaagtt ccacggcttc ggtgaagctc
76981 ttgccgaaaa actctataac aaaatcaacg gggccgccgc ccttgctctg gctgtgcctg
77041 taccatttgt ttccccggac ggtcaagctg tcgtgccgtt tccagcggta ttcattcccg

77101 gcgcgggtaa gctgctcgcc ctgtgattgc aggaaagaaa caaggtcggc ttgattcgcc
77161 cggtctatct gttcttgggt gtagtacaaa aatctcaact ccattcatcg ttctaagtcg
77221 tgccgcttgg cgcgggtctg ctccggctgg tcggctttca gtgcaatatc aacgcacttg
77281 cggatattgt gcagctcggc aacctcggcg cgggtttctt tcagcttggt atattccgct
77341 gttctctgcc cgctgagagt ggcgtactct ttttcccatt cagagatagg caaggtcttt
77401 gtccctttcg gcagatttgc atggagatag cggctcgctg cgttccatag cgtaagctcg
77461 gcgcggtggg cttcctcgta cttgtcacgc ttgctcgtcc agccattttt cagctttttc
77521 agctcgtcgt gaatgggctt gtactcggtg tagttcttcc cgtattcaat caatttctgc
77581 aattctttca tgcgcttctc ggcggttttc atgccctccc ggatagcgtc ggcttggtcg
77641 ctgacagagg aaagggcagc gtccagctcc gcaagggtag aaataccatg ctcggaaaga
77701 tagttgaccg cctttgcaac ggttttcagt tcatcggcgg cgtgctgcct ttgccaactc
77761 tgcgaatact tccggctctt ttctctctga acgcttaaat acttcatcag cagatttgca
77821 aggccggggg attgcggggg ctgctccggg gcggtttccc gcgccttgaa cagttcgcca
77881 atccattctt tgagctttcc aatctgcgcc ctgatttccc ggataaggcg gtttgccttt
77941 cggatattcc ggttcagctc gcctttctcg gtggctatgc ctttcttctc catttgacag
78001 gccgctacgc ccatgtggac ggtgggcagc tcgtcaatgc cgcgctcggc gttgctgcgg
78061 tagtcgatac gctccgggtg tccggcgcgt tcaaggtagg cgtttgaaat atccgcccat
78121 gccttgcgcc ataaaagcgc gttgcccttg tcgttccagc ctgtgaggtc aactttgtgt
78181 gtcttgtatc tgccgcttgg taagcgtatg cgctcgccgt tctcgtcaag gtcatattcc
78241 tttttggact tcgccgccca tgcgccgcgc tcgtcaaggg ggcgcatggt aagcatgata
78301 tgacaatggg ggttgccgct gtcggtgtcg tgaatggcaa aatcaacgca cattcctctt
78361 gaaacaaatt gagaggaaca gtattcacgg acaagccgga tctgttcctc tctggataat
78421 tctatgggga gtgctgcgtc aatctctctg gcaagctggg cgttcccggc tttctcgtaa
78481 agctccacgc tgttccacaa ggttgaacgg tcagagaaag agggcggggc atggggcggc
78541 agtaggattt ccgtatggac aacgccgcgc ttgcgggtgt agtcatgggt cattccgtcc
78601 cactcgtttg tgattttctc gccgcttcga taggcggctg cggcaacggc tgacttgcct
78661 tttcctcggc tcacaatgcc gatgttacaa tggtaaatgg ctatgggtat cacctcccgg
78721 ataggataac caaacccgca aaaatggtac agacgccaac ggcgggtgta ctgtttttgt
78781 gggtgtgcag ggatagccgc gtaagcggcg caaggggtgc agcccccttgt tgcggcaaag
78841 ccgccatatc ggagcgcggg gaaagttccc tgtgcggaga tacgccctcg gcagagcgca
78901 cacgcccgca agggtgtata agtgcgccct ttgttccaaa gggattattc gcttttcccg
78961 ccctcggtgc gttttttcag ataagcccgc gcttcctcgc tcgtcagggc aagccggaga
79021 aaggcggcgg cttcctcatc ggtcatggtc ttggcttccg gcacaatgct ctcaaagact
79081 gcgcctcgga cgaccagccg gtggctgcgc gtccggcgtt cctcttggga taacttttgc
79141 cgcaacacct tttcccgatt ttcaagctgc cggattttct tcttcccgtc ctcaatctcg
79201 gcttgcaatt cctcgcgttt ttttctctcg gtttcgtcat gggtggtcac tcctttctac
79261 ctgtcggcat agaaaaagga cagtcgattt cctcggctgt ccttttgatt agggtgtttg
79321 gtttactccg ttctgctcaa taaatgggaa tattaaatat caatacgccc tttaatacat
79381 tcctattaaa acatcagctt cgattgtcaa atgtgtgaga gtacgccaat caacgcaatc
79441 tttttcaacg caaaagagaa gcggcgtcat ttcaataaac gacattcttt gttctgatga
79501 caactgtacg gtggctgaaa ccgtttctct tgaaatggtt ttaagatatt tctcaaaata
79561 ttcaacgaca gactcatttg aataatctgg atttttcaag tgtgcttgca ctttagtcct
79621 gatttccctc aaatgatttt tcgtaggaat tactttcaca acatatccgt taggagatag
79681 caatcgctga aattctttgt agtgtgcagg cgaaaatatg tctaaaatac aatccatact

79741 tccgtctttc aaagggattt ttgataagtc agtaacaaac cacttcactg ctttgcgctt
79801 gtcttttttt gatgcaatct gtattgcctc ccttgacaag tcaaaggcaa agatgtttcg
79861 ttctgttctt tgctgtatct gccttgcata gaaaccctcg ccgcaaccaa catctaaaat
79921 gtttcttata gatggcgtat cagaaataaa ctgtatgatt ttctccaaca caacatcata
79981 catgccatac tccaaaattt ggtgccggtt gtcaaaagac cgcttgctgt agttggtttt
80041 gggcgatgat tttaacaaca aatttacata cccatatctt gaaatatcaa agcaatgtcc
80101 atgtctacaa attaggctat tgccttgtat atccattgat tttgtgcaaa ttgggcattg
80161 aaaataaacc ttgctgtctg caaagcgtga taaattatta ttcatttgtt tttcctccgt
80221 aattacatct ctgcttcaat aagcgagtta tgccatacgg ataaccgcaa cataactcgc
80281 ggttttatct taatctcata aatgtaacca ttgtgttgtc ctccaatcaa ttccgcagta
80341 ttactctgct tttagaaaaa gtatagcaca aaactatgaa catttcaact ctatttcagc
80401 ctgtcggcga cgttgctctc tctggcatac cgccgcgccg cttcccgccg ttcctcgctg
80461 tatggggcgg tcagacggaa agagaagcgg cctttctcaa tgtcaaactc catgcagccc
80521 gtttccgggt ctgcgtcagt ctgctggcag atcgccggat aacggcggct gtatgcaagc
80581 agacgctttt tcaaggcggt gttgtgggtg cggatatgga taaggggggtc tttctcgtca
80641 aaccaaatat cggtggtctt ttcctgcttg gtaagccctg ttctcatgca aactcctttc
80701 tgtgcccctg ttacgcaata ggggcatttt tcgggtgttt tctccggctc ttgacttgga
80761 gaaaacggcg tttttgacga taaaaaccgc ccggacgtgg aatgtatcgt cggggcggct
80821 gttatcgcaa gatttccgat ttttccggct cttgaccgtc agacgcagat aaacacgaac
80881 tgtcggcaag tatcgtcttg agcagcttgt cgcggaatgt ttctgtgctg ctgtgattga
80941 aaaataactc cgcaacaata gtctgcccgt ttctctgcgt cgtgatgata ctgtcggggg
81001 tctgttctgc cataggcagc tcctttctcc gggcacaaaa gagggacttc ccgtagccgg
81061 aaaatccctc ttggttgtcg gttattctgt tttactgtgc gggctgtgcg gcggcggtct
81121 gcgcctttct ccttgcccgg tattcccgcg ccttgatacg ctcatactcc cgctgctttt
81181 caaggtttcg cgcccggtac tcccttgaat acttccggtg gtaggcgcgg ctcttttcct
81241 ttttggcttc ttcgatttct tcccgcattt gccggatttc ctgctctgtc ggctctgcaa
81301 gctgcgttac ttcgttctca aatctgccga tatagttgaa atatatgctg atgtgctgga
81361 ttgcgtatct cgcccttttc tggtcgcgct catgcacttc aatcctgctg ataaactcgt
81421 tgagaatggc gggggtgagg tcagtaaagg cggcatggcg ttccgtcagc ttcaaaaact
81481 tctgcgcccg tcctcccgcg ttctcataag cggatagctg ctcttggagt atggaaagct
81541 ccgttttcaa tgcgtagtat tcttctgaat acttctgcga catttgctcg tagcggtctt
81601 gcgggatagt gccgagggcg ttgtcctcat agagcttgtt cagcaccttg tcgatctgtt
81661 caaggcgcgt cgtgatttgt gggatacgct tctgctgctt tttggtctgg tcggtctgct
81721 gcatggcaag gttctttttc actaaggctt caaactccgc ccggttgctg atagaatagt
81781 cctcgatttt cttcagcact tccgcgacgg tctgcatgag caagtccgcg tcaatgatgt
81841 gcggggaatg gcatttgggg tttttggctt ttcccttgtg gtactcgctg caataggcaa
81901 cgtgccgctt gccgccgttt ctgtaatcta tgcggatgtg cattttgcgc cgcaatcctt
81961 acagaaaagc aagcccgaca aagggtggat ttccccgtcc ccgttggggc gtctgacggg
82021 cgcgttttcc aaaatccgct gtgcggtttc aaagtcggcg cggtcaataa tcggctcatg
82081 cacattttcg gttatctgcc attggctccg gtctacatag tggttccgct tgtcccggaa
82141 atgctttgta gtcttgaagt tgactacatc gccgcaatac tcctgccgtg taaggatatg
82201 ggtcaaagtg gctttgttcc acttgtaacg gttatcctca ttgagcgccc tgtttttgca
82261 cgttccccgc ccgcgctctt tcatgtagaa tgtgggcgtt gggatttgcg cctgcgtcag
82321 atatacggcg atttggttgc ggtttttccc gtccagaaac aggcgaaaaa tcaaacggac

82381 aactccggcg gcttcctcgt cgattatcca aaaatccttg ttgtccgggg atttgacata
82441 gccatagaga gcttcggtgg caatcggctt tccactcatg cccttcgtct taatgccggt
82501 ctttactttc ttgctgatgt cctttgcgta ccactccgac atgatgttga taaagggcgc
82561 aaactccaat gtgtcgggct tctcgctgtc tatcccgttg ttgaccgcga taaagcgcac
82621 attgttccgt ctgaatatct ccatagcgtt gccgacttgg agatagtcgc gcccccagcg
82681 ggtcatgtct ttcataatac acacgccgat tttaccgttt tctacgtcct ctatcatgcg
82741 ggagtaggcg gagcggtcga aaaatctgcc gctttcatcg tcgtcaatgt agtgccggat
82801 attcgtcaat tttagctgtc tggcgtagct ttccaaaaac ttcttctggt tctgtatgga
82861 gttgctttcg ccgccgtccc tgtcctcgtc gcctacggaa aggcgggagt aaagggctgt
82921 gattttgttg taatcattca tgtgcatatc ctcctgtgcg tccatgtaga gttctttaca
82981 cttaagatta tgcactccat cgggcagagc cgtactccgc gcccttgcgg tatttcttcg
83041 cgtttttccc tttgacatag actatcaagc ggataatcac cgcaccgaca atgccgacaa
83101 gcaaatccaa cgggtaaacg ctcggcaagg cgttttcaaa ggctgcggaa aatccttgtg
83161 tgatattcag tatctttgca gaaatatccg cgccgggggc aagccgcacc gcctgtccga
83221 ctttatcaaa gagatagaca aagaacacat aggggatatt cggaagtatc agctttttgt
83281 agttgatctg cttcatatct cgcgcccccct gtccattttc tttaccttgt cgcgggcggc
83341 gttaagctgc ttcgccttgt cctttgcagc ggcaagggct ttgcggatag agggcttttt
83401 ctcccggttc agcttctttg cggaaaactc ttgaaacgct gcggtcataa cgtccgcgtc
83461 ccggcctttg aaaaacacaa ggtaacgggt ctgctcgccc ttaacctttt ttagcgaaaa
83521 atctatgttg tatttcttcg ccgtactctc aaaggcttta atattgctgt cggtgatctc
83581 gatgttggaa acacccgcgt tctgcttcat aagctgccgc atggtctgtt tcccatgcgc
83641 gggtttgctt ttctgctcca aaaacttttg gattgctttc tgtaacgcct gttcggtaag
83701 ccgcgccgcg ccttttccca ctttgacgta tagggcaatc gttttttggg ttacttcctc
83761 ctgcaaggta tcaactcctt tcccggtaaa gtgcggctat gctcattcgc cgccgtataa
83821 gtcgtgattg acaagggcgg tatagtagct gtcaatggta ctcggcgcgt tgaacagcac
83881 cgcttttaga tactgcttga tgttgcgcac tttggttgtg ttctccctca tgcaatccaa
83941 aacaaactca atatggctgc tgttcagctt caaaaacttg gatttcacca attcggcggg
84001 gtagtcgtcc ccggcaatac ggattgtctt tcgggctgtg cagacggttt ccagcataag
84061 gtcaacaatc tcgtccaaac ggtctttgtc aattttgcag ttttgaatga gatggtcgta
84121 ttcgatattg tccttgataa tctccctgta aatctctact gcgctattgc ttttcgcttc
84181 cgttcttttc ctttccggcg gcgtagccgc ttcgccatgc gcaaagggca aggggtttag
84241 ggaatggaaa ggaatggaat cggtacttga taaatcagta attgattgtt ctttacttaa
84301 tatatcttta ttggttcaac tccaatttta gtggataagg gtcaaaaggg gcgtggcaca
84361 gccagctgta acttgccagc aacaaggtat atcatcgagg caaatcctct gaatgtatta
84421 aagccacgag ccttccgttt tgccgcctgt atcatggagt taatcccctc acagacagca
84481 ttcgttatgc ggtggtagaa gtaagccagt atcttcgtct tgtggttccg gagtgtcttg
84541 gcggcctctt tcattggctt caaacggcaa cgacgcatcc atgagtagag gcggtcgaaa
84601 gcctgttcag cttcctccac cgtttggcag gtatagaagt catcaagggt tgctacaatg
84661 ctgtaagcct taccagtctt gggatacagc ttggacagag atcccagttt ttccgcctgt
84721 tcgctggtca tcctggcttt ggggatcatg aatagacgcc ttccaagaaa gagctctttc
84781 ttggacgcag cggttcgctg ctcctccttg cgaacatcgt caagggcgtc catcagcacc
84841 ttcttgacgt ggaatttgtc gatgatgctc agggcgttag ggaagttttc ctcaatggct
84901 ggcaggtaag acttcgacat atcgcttgtt accgcacaga tggccgccgc attgccacca
84961 cgcatctcca gtttctcagc gaaggtggac accgtggctt tatctctgcc tgcctctacg

85021 tcgattacac agcgttggtc ggcatcgatc acgactgtca catagtcatg tccccgaagg
85081 cgagaggtct cgtcgatggc gagccgcttc acgccgctca gtgagcgatt actcaccgca
85141 tcgtctaccc aatagttgag tatagaggca agagatttct catcgcaccg aagcactgct
85201 gctgcgcgag cccttggcat gtcggcaaga atcagcatgg catagccctc aaacaacaga
85261 gtgaagcggc tgttcttccg ctcaaatggg gcgttcacct gcttcgtgcc gcaccttgga
85321 cacttcaccc gtggacgcct gcaatggacg taggtcggat agaacatgca gtctccgtgc
85381 cgccaaacac gctctgttgg ctcatagccg ttacgtaaag taggcccgcc acagcatgga
85441 cagacgaggg ctgcagtctt tctgacattg acgtagatat gcacttcttg ctcttccgga
85501 cggaactcag ctccctccac gtaccacggt tcacttagat tgagagaacc tgcaatcatc
85561 gctgtgaagt ttttttccac ctgtttcatc accccgtgac aagtatatca caaaaggtac
85621 ttatccacta cttttggagt agagcctctt tatttaattg cgttggattt tccaacgtag
85681 gtttttcctg cgtaggtttt tccaacgttg gattatccaa tgttggattt tccaatgtag
85741 gtaaatccaa tataggcggc tgtccgccgc tggtagctgc ttccggctct cgcggctgcg
85801 gctgctcgta tatgacgtaa tctgcgcccc gcaagcgtcc cttttcgtcg cgctcccggc
85861 tgcgcacgat atatccggct ttttcgagtt cctttaccgc ttcgcggatt gcgtcgatct
85921 tctcccggtt gatatgggat aagcccgcaa gggtataatc ccaatcttcg ggcagcgaaa
85981 gcatttgcga caacaagccc tttgccttta aggacaattc cttgtttcgc aagtggtggt
86041 tgctcataac ggtatatccc gtatttcgct ccactctgaa aactgccata tttcctcact
86101 tcctctcttg gtatgcgtgg acaattagaa agccgttttc ggctgttttg gtattgcaat
86161 atgccctttg ccgttttcgc gtctgcaaag ccgcatggca caagggcttt tgcccgtcta
86221 tttgtccatg ctggcgtgcg ttttccgcgt ccttttgccg gggcaatagg gacactcttt
86281 gaaaacgcaa aactcatatt tccacccggg gcggtagaaa cggcaagtgc cgcaatcttc
86341 atcatctccg gcacaaccgg attgatagcg gtcaaatccc ggcttctgct gcatgagcaa
86401 ttcaaacgcc cgctgcttgc cctttgtgaa atacattccg gctgcacctc ctttcctgcg
86461 ggcaatcgag taggaggcgg tgattaaccg ccgtcctctc acaccaccgt gcgtaccgtt
86521 cggtacacgg cggtttcaac agttgacgtg cacagactga taagctatgg ctaaatcata
86581 aaagccactg cttatcagtc tttcttttgt cattgccata tttaccgcta ccgtttgcgt
86641 ggtaaaccaa tatcctctcc ggctgtttcc tgcctgccat gccaaatctt ccggcactcc
86701 cattttccgc aggttcttca ccttggttct cggtttcttc cattgtttcc atatacacat
86761 tcggattctg tggtatagcc acttattgag ttcttctatt cggttcttca tttccgcaat
86821 cccgtaataa ttcagccagc cacacatata cacttttatc tttgctaatg cagggcgtat
86881 gctctgaact gaccttcggg aactaagttc tttcagcttt gacttcattt tcttccatga
86941 ctttccatga acccggacat atatgccctt tccgttcttt cccaatgtaa agccaaggaa
87001 tttaaaagtt cggattgcaa acacgctgac tacacggctt ttctcttggt tcactttcaa
87061 tttcagcggc ccttccagat atttcgtgct cgtctccagc agtctcttgg cagctctctc
87121 gcttttcgcc agcagcacga tgtcgtccgc gtaacggaca aatacaactc cacgcttctg
87181 atactcctgg tcaaactcat tcagatagat gtttgccagc agggggaaa gatttcctcc
87241 ttgcggcgaa ccttcttctg tttccatgac tactccgttt tccatcacgc cactctttaa
87301 gtacctctta atccactgga ttaccctttc gtccctcaca tttctgcgca ggatattcag
87361 gagtatctca tggttcaggg tatcaaagta tttcgacaag tccaatgcta ccgcatggat
87421 atatccccgc tctgcatact ctttcacctt ctgaatggca tctttcgcgc ttcttcccgg
87481 gcgataaccg taactgttct ccgagaacaa cggctcgtag accggtgtaa ggcgctgtgt
87541 tatggcttgt tggaaaatgc ggtcttttac ggttggaata cctagctttc gtattccgcc
87601 gtctggcttg ggaatctcca ctcgtcttac cgggtcgggg gtgtactttc cccttctgat

87661 tctctcgata agttctttct ggttttccct caggtatgcg cctatttcct ccacggtaat
87721 cccgtcgatt cctggcgctc ccttgtttgc ctttactctc ttaaaagctc tgttgaggtt
87781 gcccttctcc agtatctcct ccaagagttc cggctgtgca ctgtcccttt ccttccatat
87841 ccggcggaaa gaacggtgcg ctctcacata cccttcacgt tccgcgctat ctctttgcga
87901 gcagctatct ttgttttctg tacccaatgt tctttcctcc tttcccagtt gtactaaaga
87961 ctcctgttga ttcggtcctt catctctcga ctactacgac ctcggctgac ttctgtacgt
88021 tcagcactat ctttcgataa tggttatccc tttcaggaca ttccgcacag acctccccgg
88081 gtaccacacg tctctttctc tccatctatc tgccacattt accatacgtg attccgtgtg
88141 gttattgggc ttcgacttgt tcagcagcct taccctcacg catggcctga tgtgatttct
88201 gtccgtcaga ccagagattt gcccatgggt tagtatgttc cccacatccg gcttccttca
88261 gatttcacct cacgatgaac acccttgcct tcagctatat ccttcccacc accgggcgga
88321 ttccggactt tcaccggtta gagacgtgcg ccgccgggcg cactaaaaaa acggcgttac
88381 tttctccccc aaaggggtta aggtaacgcc gtttgtgtgc gtattcagtt tttaacacat
88441 tccctgtcta tccgctgtcc gcaaaaccat tgattttatt agctttttgc cgctatcgct
88501 atcacacctc attatttggc atccaggtta taatcttaat tccatttgat tattcttact
88561 tctcaatcca gccaataaag gtactgacct tttcgagatc ctcctcattc atggttctca
88621 gcttctcgca taattcacga tacggggttt ttgcatcgtc cagtggagcg aaaaattctg
88681 ccggtgtcat atcaaaatac gagatgatat tgaacagttc cctgagagac ggcaaggccg
88741 ctccgctggt tataccacga atgtatgagc cactcttgtc caaatccaaa ctcatcttgt
88801 gttcggaaat attctttgct attctcagtt cagtaattct gatacgaata aattcatcat
88861 actgcgtgtt ggccacttct gtccctcctt cctcaattat tgtagtatga attggaaggc
88921 agtgttggta gcaagaaaat gaattttcgc tttatccttt cttgtattgg tgatataatt
88981 accaatatcg ggttttgatt ggcgatacca gtatgaagta tctccaaagg agagattttt
89041 atgaatttat tcgctgtcag tttctttgga cacagagaag ttgatgatcc gtttgtgatt
89101 gaacgacaat tagaagcaat catccgtgag ctgcttcttg ccaaagaata tgccgagttc
89161 cttgtagggc gagacggaga atttgaccag cttgtttcat caacgatccg cagatgcaag
89221 cgaacgatcc gggatgacaa cagttctctc gttcttgtac tgccatatat gacggcagaa
89281 taccggaaca atgaacagtc atttcatgag tattatgatg aaatcgaaat ctgtgtcgaa
89341 tccgcagaaa aacacttcaa gtccgcccat caagtcagaa atcggtccct ggttgaccgt
89401 tcggatttgg ttattttctg tgtcgaacac gattctggcg gagcttatca gaccatgcga
89461 tatgcaaaaa aggcaggggc aaagattatg aatctgccag atgtacctcg ttgactttcg
89521 cattattttt ctcggaaagg aagaacatga ataaaaagaa acgaatagcc attttatttg
89581 ctgtcattat tgttgtggca gttggaatta ttattgctgt gacttcacat cagaaaaagg
89641 aaatggactt cgataaattt gctgaccgtt ttgaaaagag cgcagaagaa gttcttggag
89701 aagatgtaaa gaaggtctcc aaaacagaac tcgcagccac acttacctac ggagatcaaa
89761 acgacaagga taataacatt tactaccgta tccttaaaac cacctttttac gagggagacc
89821 cggcagaaat caccggatta catacggaag ccctgggagc gcttttcccg gttgacagta
89881 tggacagctg tgaggaaatg atgattcagg actggcccgg cgctctatat aagaagggcg
89941 acacggcatt tctgtgctgg acgtattctc cagaagttac ttatgttctg gaatacactc
90001 caagcaaaat tgatgattca gaaattatca aaatggccaa aagtgccgaa cttgtagatg
90061 aagaataaat gttaaagagc cacgactgtc actcacatct aatggagcaa tagcggtggc
90121 tctttcttaa ccatactatc tatattgtgc tttgtagttt tccccccaag accacatggc
90181 atccagaatg ggttttagac tgtaccctaa ttcggttaat gtatactcca cacgaggcgg
90241 cacttctgca taaacttttc ggtttaccag tccgcttgcc tccatatctc gaagctgagc

90301 agtcagaacc ttttgagata catgaccgat tgattttttt aattcaccaa agcgtttggt
90361 accaggcatc agatcacgca ggattaaaac tttccattta tccccgatca gcgttaaagt
90421 tgtttccacc gggcaagccg gcaaatcttt ttccaaagaa acggtcgttt tctgctcatc
90481 cattgtatta cctccaatag tttcctttta gtaactacgg cacaatattg tgcttacttt
90541 acgaattatc cttacagcgc tattataata tcagcgaaag aaaaaagcaa tctttccaaa
90601 gaaagcattc aggaggtaat caccatgaac aaagtcgttg aatttttaca ggcaaaccct
90661 gtccagtatc ttgctaccgt tggtcgtgat ggcaaggcca aatgccgtcc tttcatgttc
90721 tgctttgagc aggacggaaa gctgtggttc tgcaccaaca acaccaagga cgtttacaag
90781 gatatgcagg aaaatcccta tgtggaggtt tctgtatcca gtcccagcta tgaatggatt
90841 cgtctgagtg gcaaggcagt gtttgaaaac aacatggctg tgaaggaagg ctgcatgaac
90901 aacccgattg tcaaaggtca gtaccagact gccgataacc ctatcttcga ggtgttctat
90961 ctggaagatc cccacggcgt gattgccgac ttctccggca atcctcctta cgagttctaa
91021 gaagaaccga aaacaataat cttcggggca aggtgaaatt cctgaccggc agtaaaagcc
91081 tgcgagcaga aacgctgatg cggtgacatt ccgtagccga cggtatagtc cggatgaaag
91141 aagatgtgat aaggcttacc ccgcagatga taaatccatc tgcggggttt tgtcttaaat
91201 ccaaatcata tctctcaagg agaattatgc gtaacgttac caatgccaaa atttctgttc
91261 gcaccattac catgactgca ctcttgtctg cgattgccta tattcttgca ttcgtagagt
91321 tcccgattcc tctgtcgcca gcttttgccc gaatggattt ctctgatttt ccagctttga
91381 tcggagcctt tgcttttggg ccaatagttg gtacactgat tgagcttgta aagaatacat
91441 tgcagcttct ttccacctcc acaggaggaa tcggagaatt tgcaaacttt ctcatgggcg
91501 catcttttgt ttttactgcc ggattgattt atcaatatca taaaaggaaa aaaggtgcat
91561 ggattgccgg gattgtcggc agtattgtga tgggaatcgt agccgctgct gcaaactatt
91621 ttattcttct tccattattt gaacagttta tgcctctgga acagttgatt gctactttta
91681 gagaatttat tccatttatc cataccaaac tggatgtggt tctttataat gcgttccctt
91741 tcaattttct gaaagggctt gtaatcagtt ctgttacgat gttagtctat aaaaggctgc
91801 gcccagtgtt aaaaggtgcg gtctgaaagg aggggtccga aatggaaagc aataaatatt
91861 tgtcctatct cgttgaagaa atacacacca ccgtagtcgc aacagtagat gattcgggtt
91921 tgccggttac agctgcgatt gatatgatgg atgctgatga aaacggtgtt tattttctga
91981 ccgcaaaggg gaagggtttc tatgacaggc tgaaaaagcg tggggctttg gctttgaccg
92041 gcatgaaggg cgaagacacc atgcactgcg ctgcgctttc tattcgggga aaggtgcggg
92101 aattaggcag tgagtttctt cccagactgt ttgaaaaaaa tccctatatg aatgaaatat
92161 atccggatgc agcgtccaga aatgcgttga ccgtttttgt catttacgaa ggaagcgggg
92221 aatggtttga cctgtccaaa aagccgattg agcgtgccag ctttactttt ggcagcgcag
92281 aggaagtgcg ggaaggttat tttatcacag atgcctgtat cggttgcaag ctctgctatt
92341 ccaaatgtcc tcaaaggtgc attgatattt ccgaaaagcc tgcggtcatt catcaggaaa
92401 attgcctgca ttgcggtaac tgctatgaga tctgtccagc cagagctatt gtgaaacggg
92461 gataatgtca tgaatcaaga cgatagaaga agatttttaa ttgaagaact gttacgagaa
92521 cgtccttcct gccgccgtca ggagataccg gaagaaccag acttccagaa gatgctgttg
92581 cgtgggctaa tgaatgtccg gcagccgagc gttgtcagcc gtcacttcct ggaagtacag
92641 gacgcctatc tgcaggaaga agtaaagaga aaaggaatta cccggctgtc cgatctgtca
92701 ccgatacagg atagcatcta cctttggcgt ggagatatta ccttgctgga atgcggagca
92761 atcgtcaatg cggccaacag cggtatgact ggctgttacc aaccctgcca caactgtatt
92821 gacaactgta tccataccta tgccgggctt caacttcgcc tgaaatgtgc tgagatcatg
92881 gaacagcagg gctatgaaga acccaccgga caggccaaaa tcacgcctgc ttataacctt

92941 ccctgcaatt atgtcattca cacggtaggt cccatcgtgc ggggagaatt gacacggcag
93001 cactgtgaac ttttgaattc ctgctatcgt tcctgtctgg aactggcgga caaaaacggc
93061 gtgggtagta ttgcgttctg ctgtatctcc accggtgttt ttatgttccc aaatgacaag
93121 gcagcggaga ttgctgtgcg aactgtgcgg gaatataaga agaaaaccgg aagcaggatc
93181 gaggtgatat ttaatgtttt caaagaacag gatgaacgaa tttacagaga actgctccag
93241 acagattgac cgtctgaagc aggtgctggg aacagcggat gctgtcctga ttggtgccgg
93301 tgcggggtta tctgcctccg caggacttac ttatagcgga gagcgttttg aacgctattt
93361 ttcggatttt catcagagat atggaatcac tgatatgtac tccggcggtt tttatcctta
93421 tgacacgctg gaagaatatt gggcatggtg gagccgtcat attttttaca accgctatgc
93481 ggatgcacct gtaccagtat atcaggtgct tctgtcactg gtgaaaaaca aagattattt
93541 tgtgctgacc acaaatgtgg atcatcagtt tcagaaagcc ggctttgaca aacagcgatt
93601 gttttataca cagggagact atggcttgtg gcaatgctca aaggcttgtc ataataaaac
93661 gtatgacaat gaatcttctg tccgtcggat ggtagaacag cagcaggata tgaaaatccc
93721 gtcagaattg attccccatt gtccggtatg cggagcgcca atgataatga atctcagatg
93781 cgatgatact ttcgtacagg atgagggatg gtatgccgcc gcagaacgat attctgactt
93841 tgtaaagcag cataggagcg gaaaaatctt gtatctggag ttaggtgttg ggaataatac
93901 gccagccatt attaaatatc ccttctggcg tatgactctg caaaatccga acgcaattta
93961 tgcgagcatc aatctcgaaa atatgatgat tcctcaaatt gaaaatcagt ccattttaat
94021 aaacatggat attgcttccg ctgtgaaaag gctgtcccaa aaggaatgaa tattgtcaaa
94081 aatagcaaaa agacgccacg atctacactc caatcggagc atagaccgtg gcattttgat
94141 ttaccggaga ggtttctttt cctctttttc ctgttctttt ttcttgtcga tattcagaat
94201 cgtttcaacc gtctgcttgg cgatcagata ttcctgcatt tctttcttca cctgacggta
94261 ctccgcatag gcttgttttt tctcaatcag gagccggtta aactcctcat gcagaacctg
94321 acgggaagga accttttttc catccagctg gtcaaaggct tgcttcgccg ctttatgcag
94381 catcagctca tcacgatgtt cttcaaaaaa cgttttgctg tacccggact tgcgatacgc
94441 ttcgtaggtc tttctggtct tggagtaatt gttgatgtgg gtctgcaatg ctccaatctc
94501 aaccattcgt ttctcagcca ccttaatgga atcggaaagt tcagagaatc gagtggtcag
94561 cttgtctgtg agagcgacca agtcctcata gctgtccaca cctttttctt tcagaaggca
94621 gactgtcctg gcagcttctt tacggttgta tttctttgcc cacttctcat agccaacaga
94681 ttttccttcc gccatcttgg actgaatatc aatgagaagg tggaactgct tctgttcagg
94741 agcctcccgg aaaccaccct tagacttgtg caggcttttc ccggagataa cggcacggag
94801 ttcttcttca ctgtatccct cgccaagaga acgcagacga ataaaccgct tctgttcctt
94861 tccccgaaac gctggctgct ttccatcctt atactcgaat cccatatcag caagagcctg
94921 aacgaaattc tcaaaactct tgggcttctc tttgagtacc gagtcgatag cttcacaaag
94981 ctcgtcacgc tgggttcgct tttttgggaa aaccgtccgt ttctgacgct cccggtaggg
95041 cttcggcgtg ataatggaaa gtccatgctc cgcacaaaca cggtcactca gtttctgtac
95101 tgccagcccg gaaaggaaga agttattgaa cttccgggag caatcaacgg aagtagaatt
95161 ataaatgata tgattgtgta tatgggcttt gtcaatgtgg gttgccacaa taaaagcgtg
95221 tttacccttc gtccagcgca tcgccgtctc atagccaacc tgatttgctt cttccggtgt
95281 tatttcttcg ggcttaaaag attggcgaat ctgataagcg atgatattgt tcttctgctg
95341 tctgcccgtg atgtgctgat attgccgttt ggaaagcaga aattcctcat cagcagtctt
95401 gggatcacat tcataggaac tgataaattc gccgttgttg gtctttgctg cgttttggga
95461 gtagtcggta cgatcagcca gacactgagc aaccgatttg cctttgttta catggagtgc
95521 aatcagcctt gtagctgcca ttcattcacc gtcctttccg aaagaaaaag ccacccgtag

95581 gtgacttcct caaaattatc taagcatctt ttcccgatct tccgcaccac gcagatatgc
95641 cagttttacc agatgcagct gaagggtata caggctgttt atggtgtgtt cgcattgttc
95701 ccaaagggct gtattcgatt ttcccacgat accagccaaa gtatgaacag cttcatctgc
95761 cacactttgc agttctgcat accgttcatc atttccggca atctctccga tggcatcatt
95821 aaaagtaaca tcaaagtaat gctccataaa tcctctgtcc atccgtttct cccatcttaa
95881 taacaaataa attccttgat atgatttccc aaagtttcaa gttttacaac cgcccatacc
95941 aacagaaaga ttataaggat tgccgctaat cccataccgg caaggatagc cagggaagtc
96001 cacagtgcct ttacgatgca gtagatggag cagcctgcaa tcaccaggag caacagagcg
96061 atcacataag aggaaatgtt ggtgatcaga tttccgagga tacaagcaac agttacaatg
96121 aacaacaacg gcagtgccaa tactttcaat actatcttca taggaacaac ccctttcttt
96181 tgaccatatt gtacattgag aagtgacgaa ctgcaatgat gtcaccacct atgtaacagc
96241 caaaaagcgt tgagcataaa aaagggcgca ctggctattt gtgaaccaat gcgtcccgtt
96301 caatctgttt tcagttcgac aaactgaaat ttgagtctct gtaaaactgg aattaaccca
96361 tacaattatt ttaatttaat ggctactgaa caatagccca tttttcggcg ggcacgaaaa
96421 tagccagcag atatgtaagc agcttcaaaa agacgcactg aatcttgcgg agattcagca
96481 ccaaaacgga catagcaatc acatgggcag cggtttcccg cagttttgcg gtcaccagcc
96541 ccatgccgca tttgcgcttc gccaagctaa atctgcgctc cacttctaca cgctcacact
96601 catcccggta atcctgcgcc ttatctctgg tttcgccctt ctttgggcga cccagtgccg
96661 gaccggatag ccggattccg tgttccttgc aataactgag attctcacgg tttcggtaga
96721 tcttgtccgc cagaatgcgg ctgggataat gcccttcccg ctcccggaac cgttccgtca
96781 tctcctgcag attcccggct tcattgtaag catcaaagga acagcattcc aaccgtgtcc
96841 acccatccac aacactgatg tccaacttcg caccaaactc cactggcttt cccacttttc
96901 cccgtacaat gggacgaacg aagggctgac ttatactgac aattcggtca gacacgctgt
96961 gggtgcggtt atcgtacatg tacttctgct gttcatagat ggtgcggatc gtatccagac
97021 gttccatctg tcgcgtggtc agagatttcc caagggtcag tttgccatct atggcatcga
97081 ggtcacgtcg gagataagtc agctgttttc caattgcttt gcgggtcatc ttgacggtat
97141 gcttgcggca tcttgtgtat ttcaggtaat ccttccgggc acgctggcgg taagtccggg
97201 gcttcttccc atctgcggga tcatggagaa catccagaat cttctcggca ttctctctgg
97261 cctcattgag cagagaaacg tcctgggggt aacggatatt ggacggtgca caggtggcgt
97321 ccacgatcat tgtgccgctg tttccgccag ctcctgggtg gtcatcagag tcatcattgt
97381 cgtcctcgga ttctgcatct tttgcctgac gctcctttgc gtcccgcacg atcatctcat
97441 tgatctcacc cagaacttcc ggcgtcaacc gcttgcggaa atacaccatc agggatggat
97501 caaagggcag cttttcgtca tcatagccgg gatacccaca gaagtactga agatacgggt
97561 tctcctggat ctgaagtgca gtctcctcgt cggagtagcc atactccgcc tgaatgatgc
97621 aggcgcccag ggccagccgc agtggtttgg ctacattacc tttgcggttg gtaaacagag
97681 cggcgtaccg cttctcaatc tccagccagg ggatcgtctg ggctttcttt acccaacggt
97741 tgctttcctt tagattcatg cccactggct gtttgaaatc cgccaagctg atctgaccat
97801 tgctgtaacg atacatagct gtcctccaag tgcaaggctt ttgcctgatt cttgcgtttt
97861 tcgatactct tattatatca cgaatcagcc tcaacagctt gcactgcaag gatttttcaa
97921 ttattcagca gacattaatt tagatattca caatattact tgccacagta tctcgaaaaa
97981 cattatcatg ttctaccagg agcatggtcg gggcaaactc cgttatgagc tgttcaatct
98041 gcatacgtga gtacacatca ataaagttga gcggttcatc ccatacatac aagtgtgcct
98101 tttcacaaag gcttttcgca attaggactt tcttcttttg cccgccagaa aagtctttaa
98161 tatccttttc aaactgtaca cgctcaaagt ccattttcct aagaagtgcc ttgaacagac

98221 tttcatcaag gttgttttct tctgcaaact cggaaagggt tccacataaa tacgatgtat
98281 cctgtggcac ataagaaatt acaagcccag agccaagtgt tactgtaccc gtataatcta
98341 tggactgccc gactaccaac tttaataggc tacttttgcc actgccgttc ttcccgtcaa
98401 gcacaatgcg ttctccctgt cttatttcaa acgaaacggg ttcacaaact gaaatgccat
98461 cataacaaac tactacattc gataatgatg caatcaaatc tgtatggtaa ttaagcggtt
98521 gtatcttgag tgcttctgcg gtttccacat ttttcagcaa tgctgatttt tgttcgattg
98581 tctgttgttg tctcgcttct atagattttg aacgcttcat cattttggcg gctttatgac
98641 cgacataacc tttatctgct gccccaattt tggaagcttc tacacgttca gaccatactg
98701 cagagcgttt tgccgactgc tgtaatcgtc tgatgtcctt ttgcaaacgt tcgttctgag
98761 ccagttcaaa ttcttgctgt cgctcaaaat ttgacatcca tgatgaaaag tttcctcttt
98821 gcacttcgat attcgctcgg ttaatcgata gaatatggtc aacacaatcg tcaagaaagc
98881 gacgatcgtg agaaactaag atgaatccct ttttcttttt cagatatgtc gccacacttt
98941 ttctcgcttt ggcatccaaa tggttggtag gttcatcaat cagtaggaaa tgcccttcat
99001 taagaaaaag agcggcaagc aaaacctttg tctgctctcc gttggaaagt gtttcaaaag
99061 gtcgccaaag cacatcgaca tcaacatcaa gataagaaag ttctcgcata agttcccatt
99121 cttccgcaag tgggcaaacc tcctgcaata tatcctctgt aatacgattc ttatctgaaa
99181 cgggataagg aaaataatca aactgtacgg atgataggat ttttccgcta tactcatatt
99241 tcccaagcag aagattcaga aaggttgttt tacctcgtcc gtttctgccc acaaaaccaa
99301 gcttccaatc ggtatcaacc tggaaactga cattttcaaa aacattatca taacttgtcg
99361 gatatgaaaa cgtaaggttt tcaattttaa tcattgacat aaatattatc ctcctttgcg
99421 tgtcatacga tacaaagtcg gttcggtgaa tgtttactcc gtatagattt caccaaaagg
99481 aaccgacact atacggagtg aatcatgtgt aactatctgt tctgcactat tcataaagca
99541 cctccaaaac aaaataagag ccgcaagaaa gtttatcctt gcagctcgtc atagcataat
99601 aacaccacta agcaaaagtg gcaacagcac tatattgagt aagatagaca tataactttc
99661 ttgcaataga cacaataatg ccaaagtata gcactaccgc atattttgat tttatttgca
99721 agaaaagtta atcatctttc cacctctttc taaatttcaa aacattatac cataaatcgc
99781 aggcaaatac aagtggttac aagaatttaa caaatcgctt ataatacaat catataactt
99841 tactcattcc ttactgaccg taaaattcaa agtttctgtt ccctataaaa ttagtttatc
99901 gggatttacc aaattgccaa acaacgattg atcgaactat caagttctct acttgccatg
99961 attatagcac atggatatga ataattctac cagctaaaca cggttgtttt tccctatgta
100021 aagccgcagt caaaactgcg acttcccgac aaattatcgg attactgaat ggaggacaac
100081 ctgacaagga cttctttcat gtccgtccag atttcttcca gacgttcctg caagtcctta
100141 atgtcggctt catagatgtt gccggtctca ttggctcgtt ttgcgtactg attcaggttg
100201 ttggaacaga ttcgcagaag cgaaaccagt gctttcacat cctgcaaatc cagattgata
100261 cagtagccgt ccagagccat ttttctcaga tatgccccca tactgcggat accaagctcg
100321 tccatcttct gatggattcg ctcgacctcc tccttggatg ccaggaaatg aacatcctgg
100381 tctcgctttg ccattacaac acctcctcgt agccctcaga gcgttttgcc ggcgggatat
100441 gctcggactt agatttcaga tcagccagaa cagaggggcg ttccgcttct cctttttcag
100501 cacggctttc caccgtgtcc tccatattga gtgccgcatc cagctccgca agccgctgac
100561 tttttgccac aagttccgct tcctggggga atggttttcc aagttctgct tttgccgctt
100621 cctgctgatt gtagagattg gtcagttgtt cattggctct ttccaaccgc tcaggaatgc
100681 cagaaagtgc gttatccagt cgggtgatat ttcccctggc atccgtgcca agtgctacac
100741 ggtggctcat agcccctttc agaacgatgt caaactcact tcggaagctg tcataggaaa
100801 gctccatccg gaaacctcgg tagctgccca acggaaccgg atcagcactt tttgtctcct

100861 tgcaggcggc aagaatgatt tcaccggcat cggctttttc cgcataatgt ctgcccatga
100921 tttccatacc acagaatccc tcggagatct gcggatgtgc ctccacggtc ttgatgtccg
100981 cttcaaagcc atggatatag cctgcctgct tttcaatctc agccgggaag tatttcagga
101041 gcttatcctc catgcggtac tgctggctct ggtggtcagc tttcagcact ttcagccttg
101101 ctacatcaat atccaagtcc atcttttcct tgatgagcgg attaccggcg cacagtgctt
101161 tgatctccgc ataggagagt gcctgctcat ccacatcgtc acaggaacga accggagatt
101221 tggaggtcat gatctgagag ataaatttct gcttattctc cagggtctga tacagatatg
101281 cgtcgaatgt tccctcagtc acatactgat agacctgaac ctctttattc tgattgccct
101341 gacggatgat acgaccgttt ctctgggtca tatcggacgg acgccagccc acatccagat
101401 gatgaaccgc caccagtttg tcctgcacat tggttcctgc gcccattttc tgtgtgctgc
101461 caagcagcac acgcacctgt ccggtacgaa ccttggcaaa gagttctttc ttttttgctt
101521 cagtatccgc atcgtggata aaggcaactt cctcggcagg tacaccgttt gccatcagct
101581 tggttttgat gtcgtcatag acattgaagg tgccatcgtt ttttggtgtc gaaaggtcgc
101641 agaacaataa ctgtgtcagc ttgtctgcct gacctttctg ccagatacgg aggacatttc
101701 ccacacaggc attgagcttg ctgttcgggt catccggcaa cagtgggttc atcagccgct
101761 gatccagtcc cagcttccgt ccatcgctgg taatcttgag catattgtcc acagacgggt
101821 caatcatacc ggcatgaaca tccgcagctc tttcggaaag ggaagcaacc atatccttct
101881 gatactcaga gggctgcacc acaatcgtct caaatttcgc ttcgggaacc ggaagatgca
101941 gctggtctga ggtcttgatg tcggctactt ccttgaacat attcatcagt tcaggaaggt
102001 tgaagaactt ggcgaatcgg gttctggcac gatagccggt tccttcagga gccagctcaa
102061 tggccgtcgt tgtctcgcca aaggtggatg cccaggaatc gaaatgagtc agatttttct
102121 tctggagcgt accgtactgg agatagcgca tgaccgtgta aagctcagtc atggagttgc
102181 tgaccggagt gccggtggcg aagataacgc cacgaccgcc ggtgatctcg tccatatagc
102241 ggcatttcat aaacatatcc gaagatttct gggcttcgga ggtggagaga cctgccacgt
102301 tcctcatttt tgtgtagagg aacaggtttt tgaacgcatg gctctcgtcc acgaacagcc
102361 ggtcaacacc caactgttca aatgtgatga catcatcctt tctgtcggcg gcaagaagtt
102421 tatccagtct ggcttccaga gattttctgg ttttctccat ctgcttgatg gtaaaattct
102481 cgccacgctg atatttcagt tctgcgattg caccctcgat ctcatcaatc tgttcctgca
102541 actgccgttc ctgccgctcg gcagacagcg gaatcttctc aaactggctg tgaccgatga
102601 taacggcatc atagtcgccg gttgcaatac gagcgcagaa cttctttcgg tttgcggttt
102661 caaaatcttt cttcgtggca acaaggagct tggcgctcgg atagaggtgg aggaactcat
102721 ttgcccactg caaggtgagg tggttcggaa ccacaaacat ggacttctga ctcaggccaa
102781 gtcgtttgct ctccatcgct gacgctgcca tttcaaaggt tttgcccgca ccaacttcgt
102841 gtgcgagcag ggtatttcca ccataaagca catgagcgat agcgtttctc tgatgttcac
102901 ggagggtaat gtccgggttc ataccgccga agcggatatg gctgccatca tactcacggg
102961 gcctggtgga gttaaaaagc tcattgtact tcgtaaccag tgcttcccgg cggcgagggt
103021 ctttccaaac ccaatcctga aacgcatcct tgattgcctg ctgtttctgc tgtgccagag
103081 tggtttcctt tttattcagg acacgcttct gctttccgtc cgcatcttcg atggtgtcat
103141 agatacgaat gtctttcagg ttcagtgttt cttccagaat ccgataggca ttggcacggt
103201 cagtaccgta ggtcacatat gccgccacat cattgtagct gggagagctt ttcccattga
103261 tacgccactc tgcggtcatt tcggagaatt tcacctcaat ggaacggcgc aggtagtatg
103321 gggtctcaaa ggtttcctcc atgaactgct gaatatagtc tgcatcaatc catgtggcac
103381 ccagacggac atcaatctca gacgcatcca aatccttcgg ctgtgccttt tgcagtgcct
103441 ccacattgat attgaaagcg gaatcacggt tcgccgccat ctgtgcgata cgcagcttgc

103501 ttctcacatc accagaaagg tattcgtccg cagtctgcca accagcttcc acatcatcag
103561 gtgccatcgg gtctttgaag atcaccccac gcagttccgt ttgaatggct tcgtaatcct
103621 cctgggttcc caaaagctgt gccatgaaag gaagatcaac tttaccacgc tcaccaatag
103681 agattgccag agcctccgat ggggtatcca cgctggtcac tttgcgttcc ggcttgatcg
103741 ttctcttggt gaacatatcc gccttgctct ttaagttgcc gtcctcatcc acattttcca
103801 gagaacacag aaggtaatag gaggaatcat cagagaaagc ctgcccattt gcacggttgt
103861 tgatgagacc gttctttgcg gtaaacgcat cgtaggcatc attcagttct gcctgcttct
103921 gagaaatcat ctcatccggg taatcctcca gctgatattc gatcagctca ttgacgatac
103981 cacggagttc caccatgcct ttcacacggt cttttgcttt ctcgttcagc tccacatggc
104041 gcatgatgga gttttcacgg aagtacactt cgccgtccac tacggcatag gaatagttct
104101 ttacatctgg agtagctgga atggtttccg cttccttgcc cttgctgttt tcaaccggca
104161 gctccacggc ctgataggtt ccctgaatgt gggaaaccgc ctctttcaga agatcggaca
104221 gctccgcacc cggaatcgga cggacggtaa tatcatcctt gccgtaaggg gtactttcca
104281 gaaccgtgtc gcccagaacc atttccgggt gatcgacaaa gtagctgttc atggtgtgtc
104341 cttcttcagt gcggttcaga tgcacccaat caggaacgat gtcaatcgga tggtcacgct
104401 tttgcaggaa gatgatgtcg gacacgactt ccgtaccggc atttttctta aaggcatcgt
104461 ttggcagacg aatggcacca agcagttccg ccctctgtgc cagataccgc cttgcatccg
104521 agttcttgga atccatggta taacggcttg ttacaaaagc gaccacgccg ccaggacgaa
104581 cctgatccag agctttggca aagaaatagt tgtggatgga gaatcccaac ttatcataag
104641 gcttatcgga taccttgtag ttaccgaaag gtacattgcc cacagccaga tcgtagaagt
104701 ctctgcggtc tgtcgtttca aaaccggcga ctttgatttc tgcgttggga tagagcttct
104761 gagcgattct tccggtgatg gaatccagct ccacaccata cagacggctg ctctgcatct
104821 caggggcag cataccgaag aagttaccga tacccatggc aggttccaga atgttgccgg
104881 tctcaaagcc catctggccg actgcatcgt agatggagcg aatgaccgta ggactggtat
104941 agtgggcatt cagggtggaa gcccttgccg cagcatattc atcctcggac agcagatttt
105001 tcagcgtggt gtactcctgc gcccaatttc ccttatcagg gtcaaaggca tctgccagac
105061 cgccccagcc gacatagttc gacaagatct cctgttcttc gggagtagcg tttcggttct
105121 cgctctccag ctggaacagg gtcttgatgg cttccatatt tcttacaaat ttctgtttcg
105181 gtccaccttc accgatgtgg tcatctgtga tgtggaagtt caccgcagga acgggtggaa
105241 cttctctggt cgtaatggac tgctcaatct gagcgatgat gtcgatgacc tcaggattct
105301 gccagccgtc caccttggta tagtcccgaa cgatggtgtc ctgaccgtcc ctgcaatgaa
105361 tgaggtcaag actgcccggt aacggtcgcc cggtattcat tgcacccaga gcctttgcct
105421 gggtggaagg gagggcgtga taggctctca gggcatcttc aaaggaactg tattccttaa
105481 tgtccagcgc accacgcttc tggatgtcct ctgctacata gaagtggtca atgaagtgtt
105541 cctgctgtac cggttcggca atgccatata cctgtgcctg tgccttttca tcaggatagt
105601 caccatagat acggatgcac ttatgctcct gctgatcgta aacggcagca ccgtcatact
105661 tgaacccgtc atcctccata gttccgtcta catagccctg ggcagctttt tcggcttcct
105721 ccagagtgta gtaatccagt ttatcatcaa agccgttttc aaagtgatga taagcggcaa
105781 ccagataacg acgttccggc tgtgctgttt cggtgcggat ttcgccctga ttttcccctg
105841 taacggagtt tttttcggta taggcatcac cccttactgc atggtttctt tcgtcctgag
105901 aaagcaaacg ctcgaaattc tcacggcttt ctgcacggaa gatcggataa agcagagagg
105961 ggtctctcag ctgcacttcc cggtcggtga tctgctccac aagaaaagcg gtgtcgtcca
106021 gatagaccgt atctccaacc tgataattcc acaccggtgt ttcctcaacc ggctctgcga
106081 tattatctgc ggattcttcg ggctgtactg cttccatcgc cgccacagtc tgacggactt

106141 cctccaggta gtcattggca agccactcat cgtcgaagta ttccgtaacg ccctcatggt
106201 ctacatagta ggttccggtc tgttcgtccc agatggcata ctcgccatcc tcggtatcga
106261 ccacagcaaa acgctcatag acatcttcca gaggatttgt tttggcatct tcgattgcct
106321 gccgctcggc ttcctcctca ggggtaagaa agcgaccttt tttcatcaga gcatcaattc
106381 tctgtgccac ctgtgtccag gacatctgca ccttatcgca gccgcttttg gtgtagcgaa
106441 cgcccttggc atcgtgatcc tgaccgctgt gggttgcacc ggaacaggca tgggaatgac
106501 cgccggtgcc gtattcatgc ttcagaaact cggctttgtc cttaacagaa tgattttcct
106561 gccaatattc atagatacgg cgctttccac cggcgaagcc gctgccacgg gtcaggtctg
106621 cattgacctc atcatcggta atgaactgac ggacaagggg catctgcatc atatcgctct
106681 gatattccct gacaggggca tccagttcat tcaaccgctt ttcaatgcca tccagcttgt
106741 gataatgaaa gcgaagcaga gaacggtcag catcatgggc aatcataaac ccacgatact
106801 cctctttcat tgccgctaaa aactcaggct gtttgagctg ttctgccagc catgcggttt
106861 cttccggaaa accattgctg cggatttccg acaaagaggg gaagaatccc attgcctttg
106921 cttcatcgga taggtcatgg taaagatacc aaatggactg agccatcttt tcacgctcaa
106981 agccgggagc ttccaccagt tccacattgg tagcgtactg accgttttcc agcaactctc
107041 caatccgctt tgcgacatcc tcccatggga tgatctgcgc cctggggctg tcaatggcag
107101 aggaaccttt ggcaaggtga atgccgtccc cggaatacca tgcggaaata ttgccgctgt
107161 cctccttcag accaaaacca ccatggtaga cctgcttcaa ggtctgtgcg atttcttcca
107221 ccggcttctg cttcatatag tccagcacta cgattttcct tgcatcatcg gtattactgc
107281 caagcaggag aaaatggtcg atttcttcct gagaaaaaga aaaagcagag ggctttcggc
107341 tctctgcttc gtcgataaat cggatttgct gttcttctgt gaggaaaagg ttcagggtca
107401 gctgttgata agctccgtca tcacgatttc ctccgcctgc gctttcaggc tgctcatgtg
107461 tcttgcccag cccagctgat ccgtctcttt ctccggtgcc gggttgttct tcagatactg
107521 gttcatcaga agttccactc tggctttcgc cgtctcctcg atctcccaca ggtgggggaa
107581 caggctctcc gtcagaatca tgtcgttcag gagcatcgga ttgttctgct ccagaaacgc
107641 tctgcgcatc ctgccatatt tgcccaaagg tcttgtttcc ttgttctcca acctgatgtt
107701 cgggattctg taatccccgg cctgtctgta agtcatttcg ctcataatct gcgctccttt
107761 cagcagctct gttctgctcg tattcaccaa tcgctttctc gatctcctga aacatccgtg
107821 aactgattgc attgactgcc gtgccgaggg tgtttaccgt ctggcgggtg ttaaaatcaa
107881 agacattcag gaagtcctca tgctcgaagt agttatcagg gttctcaacc agtcggcagt
107941 acatcgcata ggatacgctg tttgcaaccg ctctcttgaa agccacttcg atgttaagtt
108001 catcatactc ctcaaggaag ctattggcaa cgatgtcgag gatcggcttc ttgtaatcct
108061 cccaatactc cgcagaaaga gacagcgcaa tctcctccag ctgcgcctcc aacagtccat
108121 ctgcggcaac gccaaaggaa cgctccatag cttccatcac atacgcacgg ttttcctctg
108181 tgacttccca cggtttcagt tcaggggatg aacggcggac accagtatca gccacatcga
108241 atacataccg cagatagggc ttatccctgt tggtcacaaa cagtgcgatg ccggtggaac
108301 ctcgtctgac atatctgcgg tacttctcgt cgttccactc gtcataaggc ttacagaagg
108361 tggcattggg gcgctgggca taaatcatca cctgttccgg ataagtgaac tcatagtttc
108421 gggacatcgt taccagaaag gaagtccagt tatccttgtt cgatgtgatt tttccgatgg
108481 tctctcttgc caacccggaa taaaatcgct cttttgcgtt catcatttcc ctccaatcct
108541 taaaaatcag gataaagatt cagcctatca aaagtctcat catccatacc ggacaactta
108601 ttgagaacag aatcagccag ggcactaagc tcggcttcct cctcggaaag ctgctccttc
108661 atctgtatga gtgcttcgca cagccccgac tttgtgccag ggctgtaaag catcatcaag
108721 ttccgttcat cttgtgtgaa attcatcatt ccatatccct ttccgggtgg cttctgggtg

108781 gtttgtgggg tgtatcgaaa accgacattt ctttcagctg ttccaacacg gaaggtcgat
108841 cttccgcagc gggagccttc ccgttgttaa tgatgccatc caacatccca taatcgtcct
108901 ccatagccat ctctgcattt ttcagatagt tttcgggttt cataaagtcc tgaagctcca
108961 cgaaaccgaa ggcatcaacg taataacagg atacttcacc ggcgaccttg agcgccacga
109021 tgtcgctgac cgacatactg tgaccacgga agtcctccgg gtggtcaata ttgaacctgg
109081 tataaagagc ttccagctgc tcggcagcac cggcggactg ctcaatcgga ccattgtaaa
109141 ccacctcata atggtcaatg gtcggctgat tttccagctt ctgaatgtgc caatagggca
109201 taaaacgaaa ctcgtcggtc tcatcaatgt gtttcagctg aaggactgcg taagagtccg
109261 tagagctttc cagaaacgct accattctgt catcggaagt ggtatttcca tgaacatcat
109321 cccatttttc aattttcata ggcgacctcc ttatcgttgc ggttccttgt acttatgcga
109381 gggcgttgca ggggcgattt tttccgtagc ctgcgccaat ttctccataa cagattcttt
109441 ccttgcaggg gaatctacca caaccgcctg cctgccaacc tcaaaagcct gttccatcac
109501 atcttcatag accatggcac gaagctccct gcgttccagg ccaaaggatt ccagaatttc
109561 ttttctggct tcgatcatag ttaaatccgg gttatccagc tgaccgccat ctatctcgtt
109621 gtagttctca tccaacagag tgtaatcgta gccatcctca caggtctgca ccgccaggat
109681 tctgtcccga ccgaccttcc atgcggcttc atcacccatg atttcaggct ctgccatata
109741 ggttgcgccg ttgcgctcca tgatttctgc gaactggcag atgtggaaca cattgtaacc
109801 aacctgggta tggtactcgt cgatgaattt gcagggtttc acgacagttt catcgggata
109861 gtgaatctgg acacagccac cgtccggaat acggaacaag tccttatagt ggctgtcaat
109921 aaatcggatg tccttgccgt ccgggttata ctcattcctc atttgcgttt tcctccgtta
109981 tcaaattcca tgttaaacct ggtctttccg tcctcgaata cctccatata gaccgtggca
110041 tcaaaggtgt ttccggtctt agctgattta cagcctttca ggttgatttt cccagagcca
110101 agcagtttct gtgccgtggc actggtcatg ctcttgccga tactctcgaa gtagcggttg
110161 ttcttccaga gggcaaactt gcagcttcgg ttggaacaga aaaatgcttt cgccttttct
110221 tccacagaac tgccgcagac agggcatttc cccacggcat tgctttcctt tgacatcaga
110281 acttcagagt cctgaatgac ctcgtaattc cgaatcagac cggcaatcac atctttgatc
110341 tcatccataa aatcctcgct ggcgtattcg cctttctcga tgagcaacag cttttcctcc
110401 caatccgctg tcatagacgg ggactgaatc tgttcgggca tgactgttac cagagccgta
110461 cccttgtggg tcggaatcag atgcttagtt ttcttatcgc ctttccgttc caggaaaccg
110521 atacgcacca gtttttcgat gataccggca cgggttgcag gggttcccag acctttgcgt
110581 tccacctcat cgggcatctc gtccgccccg gcagtttcca tgctctgcag gagcgtatct
110641 tcggtgtagt gcttcggagg agaggttttt ccctccttaa tctccgtgct gatgacagcc
110701 atttcatcgc caacggaaac ggaagagagc acctgttcct tttcgtcaga ttttgtcgaa
110761 ccactttttc tggcaagtgc tttccagcca ttatgcagga tggtttttcc cttggcagag
110821 aacgctgtgc cgccgcagtc cagttcaatg actgtctctg cgtatcggtg cggatctccc
110881 acagccacga aaaggcgggt ggaaatcaac tgcaaaattg caagctcgcc cttgggcaac
110941 tcattcagat tgcatttctg caattctctg gtagggataa tggcatggtg gtctgtgacc
111001 tttttgttat ccacaacctg ctccgcatgt acaggtactt tatccacacc tggggcgggg
111061 aatgtgtgga aagccatatt tacgaggtct ggcagattgt gtgccatatc gtctgtcagg
111121 tatctgctgt cggtacgagg ataagtgacc agcttctttt catagaggtt ctgggtgtaa
111181 tccaaggtct gctgtgccgt atagccaaac acacggttgg catctctctg taaactggtc
111241 agatcgtaaa gcgaaggtgg cttttcattt ttctccttgc gttccgcctt tgtgaccgca
111301 gcaatagaac aatctttgct gacagcttcg gcatccgact tctttttgaa gcgttcgctt
111361 gctgccgtaa acccatccag tccaatctga accgtataga agggttccgg tttgaaagcg

111421 gcaatggcag cttcacgcat gactgccatt gcaagggtgg gagtcatgac tcgaccaaca
111481 ttcagggtct gcccataaag ggtcgaaaaa agtcgagtag cattaatgcc aacaatccag
111541 tcagctcgct ctcgacacag ggctgcttca tacaaagcat catattcagt tccagacctg
111601 agattttcaa agccctcccg gatagcggaa tcttccatgg aagaaatcca aagccgctca
111661 aagggctttc tgcacccggc tttatggtac accagtcgga agatcagttc gccctcacga
111721 ccggcatctg tcgcacacac aaggctggcg acatcatctc tggccatgag ttttttcagg
111781 ataccgaact gcttcttggt tccggaagaa acctcatacc gccagtccgc cgggaagatc
111841 ggcaggtcag catacgccca tttgctgtat tttacatcgt aggcttcggg ttgtgccagc
111901 tccaccagat gaccgacaca ccagctgacc acatagccgt ttccttcaag gtagccatct
111961 tcacgcttgt ttgccccaag cacctttgcg atactctggg cgactgaggg tttctcagcc
112021 aatactaatt tcattcgcat ctgctccttt ccaaaaaaat aagcagccac acaccgtagt
112081 gcatgactgc tatgtatgaa acttatttaa tgtattgtct cagtatttct ttctcgtttc
112141 cgattatcaa atgacttgat gatctggcta ttgcagaatt gtattctctt acagcttcct
112201 ccgaaacatc aatctgtctc tctgtattat ctccgtttat caccagtgct tttgtaggtg
112261 atataggata aaaaaatgtc agctcagttg tctcagctaa atcattatcg taatcacatt
112321 tgaggttaat aataggttga tccgatgtga taaacgggat tgctgtattg ttattaagca
112381 gtgcaatcgt agtctctttg ctcatcaacg aatatgccaa cgatgtttga atatgccaaa
112441 agaaaaatat cgacaggttt tctaaattca ccttacttaa gtcaattccg acttttgcaa
112501 aatcgagttt tttgattgaa tccccaaact gcttcatcca gcgctctttt aatggcaatg
112561 ttctgaaata ctggatacat atatcactca aaaagttgaa ccggtcattt ctatccagat
112621 tatagtagaa tcctattccc tgctcatcta ttattctgaa atactgaact aattcaccct
112681 ctatatcaca ataaaaatct tccatattgt tattcttttg aacacggata atctgttcgc
112741 acttagagat ttcatgatgc aaatcatcag gaatattatt ttccgttcca aatcgcttta
112801 tcgcagcctg tttcattaca tcaagaattt tttgattttc aatcgggaga agataagtct
112861 caatgaaaaa ctccatctgc ccacgaacaa ctggactcat agtttgcaaa taacaatcca
112921 atagcttttt ttcgtctaaa tttaagctgt tgattgtata aaatgaattt ttgaaggcaa
112981 tggcttcagt tccagtcgga aatagcttct ttctatctct cagcacccat actgcattgt
113041 tttctttcca ctttgataag taagcttgaa aaacaaaatg atgtttcttt tttacatttt
113101 ctatcctttc catgactttg cacctcatat tattctatgc tttcttttat tatagcggtt
113161 tgtctcccaa tatcaaaccc aaataaaaga gctatgtaaa ggcaatccga agattgcccc
113221 aggcatcatg cctgttacgg attataccgg ttcgttgtcc tcctcatcag tggcaaagtc
113281 aacatcatcg tcgtcctcaa attcctcgct gtagccgtag tcctcatcat cgtctacata
113341 gtccgcatca gggtcaggct tcgcagcttc tttttctttc ttcttttcct gtaacttctt
113401 aaaggcgaag aaaccaccgc cgccagcaag ggcaagcaga gcaaggatag ccggaatcat
113461 attcgtctta ggcttctcct caggcgcagg ctctgtgacc tcaggctccg gttccgtctc
113521 ggtcggttct ggagtggttc caaccacagg ttcggtatac ttggcagctt cttcctcgct
113581 catcagcatc aaaaggtcgg cttcgtccac ctgattcaag aagtgaacgg tctcgtttcc
113641 ttcatcgtca cggtcgatga tgatataaaa gtaattgccg ttctttgtga ccgcagtaat
113701 gaattgtttg ccggttttgt caggcgagcc gacatcatca accagtgtca ggttgccatc
113761 gggagtgagg gcagattcgc tggcatccgc cccggtgtcc atattctgca tgaacatatt
113821 catgaacatc ataagcagtg ccatatccat ttcgctgttc tgctcggctt cctcaatcag
113881 ctccagatag tcaccgtgta catagccatg cttttcggga acggtgattt cataccagtc
113941 tccctcggtc ccgatcacct ttacttcctc gccgggacga agctggtcga tgatctcata
114001 atccatgcct gcaccggtac gcacatttag ccgggaaccg ttcgtgacca ccttgccggt

114061 tcccatgcca tcgtcctcgg ttttcccggc atcatacttc cagtcctcgc cgttgaaaga
114121 aaataccaca gtaccatctt catctacgga tacgctgtaa ggaagctctt ccgcagtctg
114181 aaccgtggtt tccgcagggg cagtttccac ggtttcgata ctggaagccg ctgtctcatc
114241 cgtataggcg aaagcggttg tcgaaaacac gctcatgcaa agcacagccg ccagaaaagc
114301 ggcacacagt cttttaatct tcattctgca tctcctcctc gttcatagcg ttgattgtct
114361 caggcttcgg agccatatcc gcagcaaaca tacggagcag ctcagaaagc tggtcagggg
114421 tcaggttcgc agcgtgaacc atttcatgga tttcggagtt ttcctcctca cggtatctgc
114481 gttccaggtc ttttactctg gcatcccatt cagcccgttt cctgcgggct ttctcaaggt
114541 ctgcaccgat tctgtctaat ttggcaccca atagaatgtc ctccttaatt ttggattctg
114601 ccgaagcagt aaaaatgctg ttgccagtaa cttgtttcaa tcgaagcata agagatcggg
114661 tttccacagt gaatcatcat cccattgccg acataaatac caacatgcga tgcaccggag
114721 gtatcatagg tgtcttggaa gaagatgagg tctcccggcc ttgcagagct tcgagggata
114781 atatcacata cacccatcaa accgtttgcg gtcaatctgc caacgctcca gccattgccg
114841 cagttattga tgacccacga aacaaaaccg gagcagtcaa aggaagtgga gggactgctt
114901 ccaccccaga catacgggta tcccagatac ttctcagctt cccggatcat gttggcaaag
114961 cgggtatccg tcagagcttc gcccggaata tcatagtcgg tgtattcgcc ggtgactgca
115021 taaatgtcgt ccccgaagat gtccgggcgg ttcccgcttg tctggaggag tacacggtat
115081 cgctccaatt catcctctgt cagaccggag cttccgatca cacggttcaa tcccttgttt
115141 accagcttca cattcaggat ataatactca tactcaacct caacttcgta ctcgtattcc
115201 tccgtcgtgg tctcgccggt ttccgggtct gtggatgtcg aggtgccggt acgggtctct
115261 gtacgggttc ggatttccac ttcttctgtc aaaattaact catactgctg atcgaacagc
115321 caccgcagag tgctttgcac ttcttcacgg gcgtaatcct caaacttcac agtcagatat
115381 gccgccagct catagggatt gtggttgatc tcatcaaggt cataacggta ttcatcgtaa
115441 ccgctgtgag tgcgctcgat gttgttgatc tgatttctca gggcagcttc catcgcctta
115501 taatcattgt ctgccccgat gatgtcctcg tcctcggcag taaaggaagt ccccagaatc
115561 acctgcgtcc ctccctgaaa catggcagtg caggacgaga acatactgca aataagcata
115621 aaaagcagtc ccgccgccag aacaagcaga atgactttgg agtgcgtggt gcaaaactcc
115681 gtggctcttt ctgtgattgt cttggctccc tgggctgctt tttccgcacc cttggcagct
115741 gttgtctttg ccgtactgtt accggctttg gcagctgcat attctttctt gattgcctgt
115801 ttctgctgcc agcgggaaaa cgggttggaa cctgcattgg gattatcctt cttgaacttt
115861 tcataaaggg catccacatt ggctttgtct gacttttgca ccaacttatc ggccttatca
115921 tgggctttca gtttgtggct gtagatggcg tgatcgacgg tgtaggcagt ggattccact
115981 gcctttgtcg tttcctgtgc ggactgaaca ccgacattat catcttcata ctccgatacc
116041 gcccgatgca ccttggatgt gagggtcaca gccgtaccac gggtagcaag ctgttttgcc
116101 gaagatgggc gatccgtagc cacgctgttg tccgccttgc cgaaacggag ctttgccttc
116161 cgggaaccgg cggcatccgt ttcctgacgc agcttggttg ccttcttttt ggaagccagc
116221 ttatctaccg ctttttctgc tttgtcagcc gctttatccg cacggtcagc agctttcttc
116281 acctgagaat ccgcaaggtc atcttcggtg aagcggagtc gtgtcggttt cttcgccatg
116341 ccttacgcct cctttgttgc caccacctca ttcagcttcg tggtcataat gcggtagagt
116401 tccgtatcgg tcgggaaccg gtcaatgaac ggcaggatca cattgccata aaacagaagc
116461 ccttcgccct cgccggaatt ggtcacatag gacagctggt ggggagagat attcagacgc
116521 ttcgccagaa tccctctgtc ctcgcttgcc tgattcagca tataaatgaa gtcggaattt
116581 tcgaggatat tggcaatctc cggggaacga agcaggtctt tgacattctg ggtgatgccg
116641 gtgggcaggc cgccccattt acggaatcgc ttccagattt ctacggaata ggttgctgtc

116701 tgctcctcct tcaaaagcag atggaactcg tcaatgtagt atctggtgga tttaccggca
116761 cttcggttgg ctgtcacacg accccaaacc tgatcctgaa caatcagcat accgattttc
116821 ttcagctgat tgccaagctc cttgatgtca tagcagacca ggcggttatt gatgtccaca
116881 ttggttcgat tgttgaacag cttcaaagaa cccttcacat agatttccag tgcagtcgcc
116941 acatgatggg cttctttctc gtcctgtttg agaagtgcgt tatacaagtc ctcaaggaca
117001 ggcatatttt caggaaccgg attgtcaaaa tatctctggt aaatctgatg cacacaacgg
117061 tcaataactg tcttttcaat gggctgcaag ccgtccttac ttcccatgat caactcgcag
117121 agggaaagaa tgaagtctgc tttcagggca acagggttat catcctcaga gtagttgata
117181 ttgatgtcca tgggattgat atagttggta ctggaagaac tgatcttgat tacctgacct
117241 tcaaacttgt gaaccagcgc cgcatattct gcttccgggt cgcagatgat aatgtcatca
117301 tccgtcacga ggaacgcatt ggtaatctca cgctttgcgg agaaggattt accggaacca
117361 ggcgtaccaa ggataagccc attgggattt ttcagtttct ttctgtccac cataatcagg
117421 ttgttggata aggcattcag accgtaatac aggctttcgg caccattctg gaacagctcc
117481 tgtgtggtga acggcacaaa aatagccgtg gaactggtag tcaggccacg acgaatctca
117541 atcagattct gtgccagcgg cagactgctc atcaagcctt gttcctgctg gaaatccagt
117601 cggcgcagat tgcagttgtg cttctgggcg atactctgtg cctggaacac attgttttcc
117661 aactcctgtt cggtgcggcc ggtgttcatc acaagaaagg tcagcatgaa catccgctcg
117721 ttctggctct gcaattcttt cagaagtgct tttgcatctt tgccgtaggt ggcaaggtca
117781 gagggaatga tgtccatgtc ataaccggaa cggacagcct ttttctgttc ctcaatcttg
117841 gaacggtcaa gctccgtgat ggttcgcttg atggtcttga ttgccgccgt ctggtctacc
117901 gactggatgt gcatggtcac gatctggctg gattccatat ccagaaagtc agccagcata
117961 cggtcggaaa ggtcagaagc gataatttgc agataggaca tcgaaccgta gatgctgccc
118021 atctggaagg tacggctgtt ctgaaaggca aacgctgtcg gcgcaataaa gtcctttacg
118081 gacagaccgg attccacaag atacttccag tcaaagaaga acttatcatt gtcacccatg
118141 tggaacatgg agtgcatcag acgaagccgt tccttgccgt tcatcgtgga tgccatcaca
118201 cccaaacggc ggaagttgtt gagcaggtca ttctcgatat ggttcaatct cggttttgcc
118261 tgacgcatgg actctgcttc gatgccgaag gtcagatact tggtcttggt cagaccgtta
118321 ttgccctgag aaagctgctt tttcaggatc tgactgtact cggcacggga agcattgaag
118381 ttgtcctttc ggaacgggat acggatgctc ttttcaaagc tctccgcatc ggtactcatg
118441 ttcataaagg acagctcaaa ctgaacagag ctgtcgaaga aattcaggaa actgcaccac
118501 tcatcgaaaa tggcggtttt atcctcctgc tgtgccagct gatagttgat gtcctgaaac
118561 tgaatggttt tggtgtagta gctgtctgtg acccggcaga taccgtccgg gaacattctc
118621 tgaaagggaa tggactgctg ggcggtctgg ggaatcccat tgtctttctt tgcccgttct
118681 acaacagcct taatttctct ctgctgcttt cgattcagat ttgcaggcat ttttatttcc
118741 cttgcctgcg gtttcttctt gaaaaacaat tcgctcaacc tccttttcaa gttgatattg
118801 acgcaccaga gcgtcgtaat agttgttcgt ctgataaggt cgaaccttgg gacgaataaa
118861 ttttgcctga atgaaatgtc ctgcaatgac ttccagcggc tgaccgtcct tttcgtacat
118921 agccagaaag aacaggggaa gcatgacaat catcatggtc agtgccgcaa ggcttacatt
118981 tcctgttcct ttgaccaaaa agaaaatcgg caccccaatg agtgccgcag agccgaaaca
119041 gatcagctgt cgttttgtca gattgaagaa cacctttgtt tttaccctcg tcaggtcacg
119101 gggtacgggg atatatgctg ccactgaaaa cctccttcct ttagtgggca gagaacacgc
119161 cctttgcaag agaaccggtc ttgaacaggg tgaagcacaa aagaaccgtg tatcccatga
119221 caccccagat tgaagcgata atgtcctcac tgaaagcgac cgactgcacc agaaccgcat
119281 agataccgac acagatcata atcaggaaac cctggaaccc gattgcaaag agtgacctca

119341 gatagttctg tccaacgctg ctctgttctc tgttcccgaa ggtagcaaaa ggaatcggag
119401 caaggctgac catgagataa atctcaatca ttcgtccgta aacaataacg aagatcacga
119461 ttgccaatgc cgacattgtg acctgaacaa tcagggattg caggaacagc cctaatagcg
119521 gaccgagatc cattgccatc agcgtatctt ccattgtttc aagcgccgcc gcatctatgg
119581 cggtgtctcc cgcaatgata ccggcgctgg cattgatcac gtgctgtgcc acatcaaaca
119641 cagccattgt gatattgaat gtgtttgtga ttaacacgac cgcaacgaag gtcttgaaca
119701 cccacttgaa aaaaatccat gtctcaaaat tcgccagatt gttgtggtcg ataatcattt
119761 gaatcagctc gtaacaggcg ataaatgtca gtatcaagcc tgctatcggg ataatcactg
119821 actcggagat gttgcgtacc atggcaaaga cacccggcga gaaattcgcc ggggtcatgc
119881 caacggaggt tgctatctcg ccaacctggt tgttgacgga ctcaaacatc cccgacagat
119941 ttcccatgat ggctgaaacg agcatttctt tcagccagtc tgttatctgt tggagtatgc
120001 tgtccaaagg cgattacctc cttttttacg gattaaccaa agagaccgga aagcaggggt
120061 acaagggtgg ttccgatgag agcgacaccg ccgccagcca tgagctgctt catacсctga
120121 gactttgcgc cggggttgtc gttgccgtag ccttcaagca ggttgattgc accccagatg
120181 ccgagacctg cgccgagagc gataacgaga gtctgaagaa cggttactgc gctgttgaaa
120241 aatgccatat acgtttcctc ctgatatttg aaatggattt tgaaatgaaa aaagccctcc
120301 acaataccgg agaaccggtc ggagggcaga gacaacgaga ctattcaact gcaatcatat
120361 ctatcaccac cacctttcaa gggaaatagg gtcaaccatc tttgccggtg cgattcacca
120421 gaccatccct tgcttttgca atcttctgga tgcggcttgt cagctgggca atcgttgcat
120481 tgatttcatg ctgattgtcc gcatagaagt agccatcata atcgctgcaa ataggttcac
120541 ctttgcttcg cagggaattg acgattcttc tgatttctga ccсttttaca tgaaaaacgg
120601 cttcaagcgt tttacttgaa accggattgt tggaacctct gtaatacttg gacaggtatt
120661 cacagaatgt ttcctcaata ttgaattggg gagttttttc cattaaggcc ctcctcgtat
120721 aataagtttg taagcaagaa aaaccgctta cagacttatt ctttttttggg tataaatggt
120781 aaggatatct aagaggtact cccctcatcc cgattcccat ctatacagtt gatagttaca
120841 tttcaatata ggtagatcct gtggtatgat atcaggtaga gtctgatgtc cgaaggcact
120901 gcttctcctc ctttcagccg gcttcgtccg agactgcttt gaaagcattc agcctggcac
120961 atacggcaca ttccgctgac acagaatttt catccccagc cgttagcccc agcaaaaaag
121021 gctgactggg tgcatgctca ttgcgcgagg gttcggtcac cgtatgcggc acaggataaa
121081 ccttttgctt caggcttcac aaatccccat cggaatggag gtgatatcat gttaaagctc
121141 aactatatgt ccaccttgtt tgtaggtatt gatgtaagct caaagaccaa tgctgtctat
121201 gctatggact tcgaggaaaa caaatacatc tcatcatcgt ttggcaataa ccagccgggt
121261 gccgataagc ttgttaatat gattgctgag tgtatgcaca agcataaaaa cctgaatacg
121321 attcttatcg tgctggaatc cacttctgta tacagcgtcc atatttcaaa cttcctatcg
121381 accagcgagg ttctcatgcc ctacaagcct tacgtcttct gtgtaaaccc aaaggccact
121441 tcgaattacc gaaagtccta tatcggaatg gaaaagaccg atcccacaga tgcatatctc
121501 atcgcagact ttggcagggt aggccgtacg aaaaaactgg agccatggcg cggcgggcag
121561 ttcatatccc tgaaacgcct tacaaggcac cgtatgcacc tttctgagtg catcaccagg
121621 gagaagactt atatggtttc gaatctttat ctgaagttca gtgaacttca gctccttgaa
121681 ggtgatgaca agccttttgg cagtctctat ggtgccacgt cctcagctgt cctgacggaa
121741 tttttgtctc cgcaggagat tatcgatatg ccggaagagg agctgcttac cttccttgcg
121801 gggaagagca ggaaccgcat ctctgaccta tcaaaaactt cggaactcct gcgcaaggca
121861 gccagggatt cctaccggtt agacaagtgc atgtatgaac cactgagcat atcactggca
121921 agctcgttta actgcatcca cgcctatcag aaagaaatca agctgatcga gcaggcgatt

121981 gagaaatgca tcaacggaat gaaccctaac gccctgctca tccttcaatc catacctgga
122041 ataggtccga tatgggcatc cggaattcta tcagaaatag gcgatataac cgtgttccat
122101 tcatccgatg cccttgccaa atatgccggc ctttactggc ctaagggtga ttccggagat
122161 ttcacttctg aggacaacaa aatgtcgaaa gccggaaatc cctatcttcg ctgctatctt
122221 ggcgaagcgg caaacagcgt caggaagcac atccctgaat atgctgattt ctatgccaga
122281 aaatatgctg aggtaaccaa acatcagcac aaaagagcac tcgcgcttac atctcgtaaa
122341 ttagtacgac tcgtttttgg attgctggtc aaaaaccaac tgtacaccgg cgaaaagctg
122401 gacaccgaat ataatattga gtcgaactga cattcggtta tacgtaaaat ataccgaagg
122461 tctgttaaag ttaccctttt ttctgaaaaa catatcaaaa tcttcttcaa aaagttcttg
122521 acatatcacc agaatgcttt aatcggcatc gactactaca aattcatcac ctgcctttaa
122581 tttcaatctg cggctcagaa actgttcgat gttgaactca ttctttttgt cgaagtctga
122641 tgtgtactta tagttcggat gctgtgtcag atcgtacttg tcagacagaa acggtctcac
122701 accacgcagc tgcaagatgc acttaccacc atccagaact gccagctcat ccacggtcgc
122761 aaggtctttc cccagctttt gataattcat gccgtaggat gggctgttgc ctctggtatt
122821 tgaggtgttg taggtgtcga tggtctcttt ccccagcgcc tgattcagct ctttcagtgt
122881 ggtaggctca gaaccgccga ggaaaatacg ggaatccata ttgccgatga tggtgtccga
122941 attgtccttg taaatggctt tcagctgaga ctgcgcctga aggacaaggc aggcagatat
123001 ttcacggctt cggatggtcg caaccagctt ttccagattg ggaatctgac cgatatttgc
123061 cgcttcgtca atcagacagc gcacatgaac cggaagtctg ccgccgtaca catcatctgc
123121 tttctcacaa agcagattga acagctgggt ataaatcatg gagattagga aattaaaagt
123181 cgcatccgta tccgacataa tcaggaacag tgctgttttc ttgtctccga gagtatctaa
123241 ctgcaattcg tcataggctg taatatcccg gacttcctga atatcgaagg gagcaagcct
123301 tgcaccacag gaaatcagaa tactcttagc tgttttgcct gccgccagct tgtatttctt
123361 atactggcgt accgcaaaat gatttggctt tttcttttcc agtgcttcaa acatcaggtc
123421 aaccggattc tggaactcct catcgtcctc acgcacctcc attgcgttta agaactcaat
123481 cagagtggag aagttctgtt cctcaaccgg cgcttcatag tggatatacc caatcaaggc
123541 gcagtacaga agggtctctg cttttgtcca gaactcgtca ccggctttcc cgtcgccttt
123601 ggtattggca atcaatgtcg tcaccagctt cagaatgtct ttttctgaat ggatataggc
123661 gaacggattg tagtgcatcg acttcttaaa gttaatggtg ttgaaaatct tgatgttgta
123721 accatgtttc aacaaagcat tgccgcactc gatgacaata ctgcctttcg ggtctgtaac
123781 aacatagctg gagtgcatct ggagaagatt cggtttcagc cagaaccttg tttttcctga
123841 acctgaacct ccgataatca gaacattctt gtttctggca ttggcaggat tcttcggtcg
123901 gttggacatc atcagccgtt ccgttttcgt taggataata ttgtcctcaa actttggagc
123961 ggtgaacggc tcaatgtctt tggctgtacc ccaacgggca gaaccatact ccacattgtg
124021 ccggtacttc ttggcgtttt tgcttttcag ataaaccgca agccgaagcc ctgcgccaca
124081 ggcaattccc accagcaaat ccagcggatg aaagctgggg agtggattgc ccagggcaac
124141 cggaagggca taaaaaaagc tgaggatttt tgcagaagaa tctgtaccct cagccaaccg
124201 ccatgcttct ccaaaattcg ttgcaaacaa tcccaccaga atatacggga tgttcagaat
124261 caggagcttt ttcaaattga ttttgctcat cacaaatcac gctcctgttt cttttcccga
124321 accttcctgg ggatagctgc taccatttct ttgaacttct tcagctgtgc cagaacgctc
124381 ggtttcttat ccttcgtcaa tgaccttgcc tgatgttcct tgcagatagc atccagggca
124441 tctgcgtcct tggaacggac aaagacaaga taacggggag gatcaacgga cttatctttt
124501 ctcacggcaa agtccacacc gtatttcctg gcgagccttt caaaatcccg gatgccatct
124561 ttatcaattt ccacattggt ggcaccctga ttctgtccca gcagctgttt cacggattgt

124621 tttccgtgaa ccgccgtatc tctggttcct tttctcgcct tgaccttttc ccggtgctgc
124681 aaatatttgc gaataccggc aatgatactt cgagcagtca gccttgtcgt gctgattgcc
124741 agatttatcg tcctgttttc aacttcttcc tgcatgaatc atcacctcct gcaagcagta
124801 gagtgagggt catcgtgcca tatctctgcg gtcgtaatac agatttcctc tggcaacatt
124861 ggcatgactg ataatcttga tgttcccggc tttctggtta tccagagctt tctgataccc
124921 ggcttcgctc agatacagac ggtttctttc gcctttccag ccaacaggag attcattggt
124981 cagcacatca aagataatca tatgctttgt ggcttcatcg aatcttgcca ggtccatata
125041 ggtgtgtccc tgatactcct tggctcctgc ctgtatcctg cggcgttcca tcaattctgc
125101 aatcgttgag cttctttcca tgatgcctcc ttacagcgta tctcgatcag ggctgctctt
125161 gcgtggggtg gggtcagcct tgataggcgc atcttctttt ttgaccatat catcgggcaa
125221 cggcaaagcc atgatactgc gccccatgcg aacgaaggtt tcaggctgat ggaaatgttc
125281 ttcaaactgc ttcatcagct ccggggaaag ggaggaaaaa tcctcctcac ccaaaccgac
125341 aaccagaaag tcaccggcga tgatgtcgta gatctcgccg tctccgtccc gcaacgcacg
125401 attcaggtct ttccccatga tttttccctc atcgtgcatc accaatgcga caggctcatt
125461 aaaaggataa acagcttcaa tgtcgccacc gacagctttc tgcaaatcct ccaaatcgca
125521 accgatctga acctgctggg gatacatata gggctttacc agaagcaccg tcatggtttc
125581 tgcctgttcc tgctgaatct cacgttcctc cgtagtgcca aactcaacgc cggtaaagtt
125641 ctcggagtcc agaatgtact cggaattgta gtaggctgtt ctgacctgtt cttctgcatc
125701 agcacgggac ttggcttcca ccgttaccgt cttttccagt gtctcggtga tctttacatc
125761 gtattccttc aaacttcatt acctccattt cgtcagacat aagaaaagcg accatccacg
125821 aacctgttac agttcataaa tagtcgcttc atgttcccac tgggggacac acatagattc
125881 cagttcctcg atcagttcct ctggggaaaa gaacaaatcc tccgcacttt tcaaaatctt
125941 tttgaaagcc cggtactccg tcacatagcc ggtatcatca gagccgatca gcattgacac
126001 acagcggaca atggcaaagt tcatatcctc cataagccac agttcggttt cataggtggt
126061 tttgatttca tccaaaaccg cctgctccac atcggaaata atgaggcagg cacgctggtc
126121 aaacagttcc atgccacggt aattgaagca tttctcatag ccgccggcta cggaatattg
126181 atataccgtc tgtgccgcac ctctcaatgc ggtgcaaacc tccggcagac acaaaccacc
126241 ggcatagccg atctcatctt catcataccc ctcataatct tcgggggaaa acagctcaaa
126301 cagccagtcg attgctttac tcatttctct ttcgttaaac atcgggattc accttacctt
126361 tccatagatt tctctcgttt cttctgccat tgctccagca gctgaatgat tttatcctcc
126421 atctgcttgc tggtgtagga cttcgggaag tatttccgca gggagtcggt tttgaatgtc
126481 accctgtcca gctcgccctt tttgacctca ttcataatct cgcacatggc atccagagta
126541 cactcgccct cctgtgccag ttttttgatt ctctgtgcct gtgaaagaga aggcgcagcc
126601 tgggcaaagt ccatagcaga aagaaaatcc tgctgttctt ctcttgccag ataagaaagt
126661 tccaccgccg gattaaattt aatctgaccg ttatccacca tatccagaag ttcggggata
126721 agctcggtta aacgaatata gcgttgcact tgtacacgac tatcgccgct ctgttcagca
126781 acttgattta ctgcccacga cttctgccca agttgggcag aagttaaatc acttcttgca
126841 ccttgatgtt tcagggcatc taacttcatt ttataagcat aggctctttc actcggcaga
126901 atggtttctc tctgcaaatt cgcatcgacc atcaaaataa tggcggcatc atcatccata
126961 tcacggacga taacgggcat ggtttctttc ccggcaagct ctgatgcacg gtgccgcctg
127021 tgaccggaga tgatctcata accgccatcc gggtcgggac ggacaatcgc cggggtgaga
127081 acaccgaagt cacggacgct ttcaaccgtt ttcatcatat cttcatcgtc taccacccgg
127141 aagggatgtc cctcaaaggg gtgcaattca gaaagcggaa tttcctgcac ccgctcctgc
127201 tgggcttcgg ctcgtgactg gtcggtagag aaaatatcat cgtacccctt caagctgatg

```
  127261 tttgcgcctt ttttctgcat tttttaacac ctcctttgtc agaacccggt aggcatcagc
  127321 gacttttcct ttgggatcgt gcttaaagat actggctcct tcggcactga tttcctctgc
  127381 acgaacagaa cggggaatgt ctgtcttgta gactttcagc tttcctccat aactttcccg
  127441 gatcagattg ctgatgtcct tggaataatt ggttctgctg tccaccatcg tcagaagaat
  127501 accctcgatt ttcaattttg ggttgatttg tcgtctgacc ttgttgatgg tttccagcag
  127561 ctgttccaag ccctttgccg gaagatatgc cgcctgaaca gggatcagca cattgtcggc
  127621 agctgccaaa gcattgactg tcagcatacc caaagagggc atacagtcca gaagaatgta
  127681 gtcataattc tgcttgaccg tatccaggta ctgacggaga atggtctccc ggctcatggc
  127741 attgaccagg gaaacctcta agcctgacag ttcaatattg gcgggcatca agtccacacc
  127801 ttccgggtga tgaagaatgc cttcgccggg ggcaatcggc ttttccatca taatctttcc
  127861 catcatgtcg gaaagggtgg aagataaggt atccgggtac ggatttccca gactgactgt
  127921 taaagaagcc tgcgggtctg tgtctaccag cagaaccttt ttgccctcca tcgccagtcc
  127981 cacacccaaa ttttccgtgg ttgtggtctt gcccgtgccg cctttctggt tgactacggc
  128041 aatgacggtt gcttttttcg acaaatatct ttacctcgct ttctattctg tgtatactgc
  128101 attgattacc acctccctcc aagtttcccc gaagccatat catgcctgac ctttgcgtca
  128161 tagtgggcgc ttatggtgta gggggcattg tatatcgaag caagtaaata ctgcttgata
  128221 ttgcgaacat cggttgtgtt ctctttgaaa gaactcatta caaatctgat gtgttcagaa
  128281 tccagcttca taaaactgct tttgacaatt tcaatgggct tatcatcact ggcaattcgt
  128341 atttgcttcc gattactgca catgacttca ataatcaaat gcagaattag atcgagtatc
  128401 tcatggtcat acggataatc ctgtttcaga tagtccatag aaagctgttc ataaaaatac
  128461 tgataatatc gggtaaattc gtcagtctca tcgacttcct ttccacaaaa accggaagga
  128521 aagaaaggat ctgtatcact aaaatcaata taattagtat cagtattatt accttgtgct
  128581 tttgacagtt cttgaattgt gtttttgaca actccagaat tgtccttttg acaattcaag
  128641 aattgtcttt ccacagaacc ggaaatgaag ttctttacat agatcaggtt cggtttccca
  128701 agtccctgtc gcttgcgttc tatcagaccg catttctttt ccagctcatc cagcagcttc
  128761 accgctttct tttcagcaca gcctaacgca cctttgattt cttcaatcgt gaagatgata
  128821 tagacacggc cttcctcgtc cagccaaccg ttccgggctg agaggttcat gcggtcaagc
  128881 agaataccgt aaagcagctt ggcatcggtt gatacattcc agaagcaatc ctcggtgaat
  128941 agcaccttcg gaacacggta gaatgcgaac atctccgctt gctgcccata gaaataatca
  129001 tacgtcac
//
DEFINITION     Integrative and mobilisable element IME_6
    fasta_record    1..7027
                /ID=D90|SC|contig000001
                /label=D90|SC|contig000001
                /note="D90|SC|contig000001"
                /colour=5
    CDS             356..634
                /ID=D90_00915
                /color=1
                /locus_tag="D90_00915"
                /note="hypothetical protein"
                /product="hypothetical protein"
                /protein_id="gnl|SC|D90_00915"
```

/codon_start=1
CDS 738..1505
/color=8
/locus_tag="D90_00916"
/note="Nucleoside phosphorylase, protein
motif:CLUSTERS:PRK07115 protein motif:TIGRFAMs:TIGR01721
protein motif:Pfam:PF01048.14"
/product="Nucleoside phosphorylase"
/"ID=D90_00916
/protein_id="gnl|SC|D90_00916"""
/codon_start=1
CDS 1559..2431
/color=2
/locus_tag="D90_00917"
/eC_number=2.7.7.-
/note="Aminoglycoside 6-adenylyltransferase Streptomycin
adenylyltransferase, similar to AA
sequence:UniProtKB:P17585 protein motif:Pfam:PF04439.6"
/product="Aminoglycoside 6-adenylyltransferase
Streptomycin adenylyltransferase [antibiotic resistance]"
/"ID=D90_00917
/gene="aadK"""
/protein_id="gnl|SC|D90_00917"
/codon_start=1
CDS 2573..2926
/ID=D90_00918
/color=1
/locus_tag="D90_00918"
/note="hypothetical protein"
/product="hypothetical protein"
/protein_id="gnl|SC|D90_00918"
/codon_start=1
CDS 3205..4746
/color=4
/locus_tag="D90_00919"
/note="DNA strand transferase  Plasmid mobilization system
relaxase  MobA/MobL family , similar to AA
sequence:UniProtKB:P07112 protein motif:CLUSTERS:PRK13889
protein motif:TIGRFAMs:TIGR02768 protein
motif:Pfam:PF03389.9"
/product="DNA strand transferase  Plasmid mobilization
system relaxase  MobA/MobL family [relaxase]"
/"ID=D90_00919
/gene="mobA"""

/protein_id="gnl|SC|D90_00919"
/codon_start=1
CDS 4812..4973
/ID=D90_00920
/color=1
/locus_tag="D90_00920"
/note="hypothetical protein"
/product="hypothetical protein"
/protein_id="gnl|SC|D90_00920"
/codon_start=1
CDS 5050..6687
/color=9
/locus_tag="D90_00921"
/note="Serine recombinase, protein motif:Pfam:PF00239.15"
/product="Serine recombinase [recombinase]"
/"ID=D90_00921
/protein_id="gnl|SC|D90_00921"""
/codon_start=1
CDS 6947..7027
/color=1
/locus_tag="D90_00922"
/eC_number=5.99.1.2
/note="Type III DNA topoisomerase TopB partial, similar to
AA sequence:UniProtKB:P14294 protein
motif:CLUSTERS:PRK07726 protein motif:Cdd:COG0551 protein
motif:TIGRFAMs:TIGR01056 protein motif:Pfam:PF01131.14"
/product="Type III DNA topoisomerase TopB partial"
/"ID=D90_00922
/protein_id="gnl|SC|D90_00922"""
/codon_start=1
BASE COUNT 2402 a 1135 c 1722 g 1768 t
ORIGIN
1 ttatcgagtg aaaataatgt ggaaatctca gtatgaggaa atgttttcat aggcattttt
61 taatgtggtt ttcacaataa gtatcgttgg atgtcggaag atagagttgg ctaaagagca
121 tacttacctt ccctttcctt ctcttcctct tcctagagaa aaaagagaaa aagaaaagca
181 gaaaagaaag aaccaaagaa aagaataaaa gaaaaagata aaaaagagat ataccatctc
241 tatcttatcc tatcccatcc tatccgcctg cctacggcag gctttattgt ttgcccgaca
301 aaagtacgag tcaagagtcg tagtaaatac gagccaagga gaaaggagtg atttaatggg
361 cagtttatcc aagtatgagc aggaaacagt tatcaatttt aatgcagggg aacagacagc
421 agtagtttat acccgagata aggcagttat gaggaaactt gacgcattag ttattgcgtt
481 tccagaggtt tataagctgg tgggtgaaac agttattgat aagacctatt caatgccgaa
541 atcatgcgtt agttatcgta agccgagaat actggatgat aaatatagag aactaaaaag
601 aaagcaaatg acacagtata atatgcaaag ataatccatt gaaaaaaagt gtgttcggtg
661 atataatttc gttgcaatat aagtactgat tttgaattaa atcagttcga caaacttgaa

721 tttgacggag agatagtatg gctattatta aaaacgaaat accgattttg gagtttgata
781 cagagcagac agcagttctt aatcccactc atgaaaatct tggtttgaac ttacccaaga
841 aatgtgtgtt cgcatttctt ggtgcttaca tagatgaata tgccagtaaa agtgatacca
901 aacaggtatc cacttttgag agtgcaacga agcattatcc catttacatc accaaataca
961 agggtgagga cattgtgctt tgtcaagcac ccgttggtgc ggcagcagcc acacagatac
1021 ttgattggct gattggttat ggtgtaagag aggtaatctc ggcaggtagt tgcggtgctc
1081 tggaaaagtt ccccgaaagt acattcctcg taccaaacaa ggctctgcgt gatgagggaa
1141 catcttatca ctatgcgccc ccttctcgat ttgtggaaat cagcgagaag gcaagaaaag
1201 caattgaaga aaccattctt gagcatagta tgaaatatca ggaggttata acctggtcaa
1261 cggatggatt tttccgtgaa accaaagaaa aggtggctta tcgcaaaagt gaaggctgtg
1321 cagttgtaga aatggaatgt tctgctctgg cggcctgtgc tgatttcaga ggggcaacct
1381 ggggtatgat actttatacc gcagacagtc ttgcagatgt tgacaaatac gacgaaagaa
1441 actggggcgg taatgcatac gaatatgcac ttacgctctg ccttgatgcg gttatcaaat
1501 tataagaaca gttagaatca atcttttttt gtgttataat tttattgagg tgaatgaaat
1561 gaggactgaa aaagaaatat acgatttagt tttgaatttt gcttttcaag atgagagaat
1621 tagaattgta actcttgagg ggtctcgcac taatgttaat ataccaaaag acaatcttca
1681 agattatgat attacttttt ttgttataga tatgggagaa tttttgaaaa gcgatgactg
1741 gcttagtgtt tttggtaaca gaattatgat gcaaaaacct gaagatatgg aattatttcc
1801 tccagaagaa aaaggatttt cttatataat gctttttgat gatggtgtaa agattgattt
1861 aactttgtta cctgtttcaa tgcttgagga gtatcttact cgcgataaac ttgtaaaaat
1921 catgttagat aaagataata tgattaaaac agaaatagtt cccacagatg aagattatta
1981 cattaagtgt ccaacagaaa gaaagtttga tgactgttgt aacgaatttt ggaatgttgc
2041 aacatatgta tccaagggct tacttcgtgg tgaaatacta tttgcaatag accatatgaa
2101 tgaagtatta cgtcatgaac ttcttaggat gattagttgg tatgtcggta ctgaaaaagg
2161 atttaatttc agtttgggta aaaactataa atttttagat aagcatattt caaaagaatt
2221 gtgggataac ttactaaata catattcaat gagttcgtat gaagaaatgt ggaaatcgtt
2281 tgatttgtgt ctatgtttgt tcaaaaaaat atctaaaaaa gtggcagatt cacttggtta
2341 taattatcct gattatgatg aaaacgttac aaaatatttg gaaacacaaa agagaatttc
2401 aaataaatat ctttatggta ataggtattg acaaattcca atttgtctaa cagataagtt
2461 ttattttaac gagtaattta ataaagtgtt gagcaccaaa aggtggcatg gatattaaaa
2521 ccatgaccgc ctttttgtt accaaaaatc aaggaaaaag gagaatgaaa tcatggtaca
2581 gaataacgaa aaaacagcag aggttttaaa agcagagagg attgtggagc aggctaaagc
2641 aagactggca gaagccaaga gaaaggcaag ccagcagaag cgaaaagagg aaaatcagca
2701 taagtacatg ataggtggca ttgttcacaa atattttccc gagtgctacc agtatgacga
2761 gctggagctg aacagaatta ttgcttcggg gattaaatca gagcagtgcc agcgcattat
2821 agaaattgtg aaaaaggaaa gtgctgaaaa gagtgttggt gttgcttcaa atgctgaaag
2881 tgaggtggca gggaatgaag gaactggcaa gggcgaaaat gtctaaaacc agtcaacgag
2941 attttctcgt tgcgcacaga tataccatca cggatggtat gtgcgctctc cgaggggtct
3001 tgcagacagc tattgcctgc cgagggcagt caaaagggaa cactgcgttt tcccttcggt
3061 gcagagggcg cacctaccca tttgcatact ttgcgttcag tgaagctcca aagagcggag
3121 ctgcactgcc cacaaagtca gccgataccg cacaatccac agtatcggtc tgccacagcc
3181 tatgcaggag agtgaggtgt ggctatggca gttaaaaacc taagtacccg agttgccatt
3241 atcggaagag gtgcaaagtg cagaggtggt tcatccgcag ttgacaagtc tgcatatatc
3301 agccgtacca ccatgtactc tgaatatgac ggaacaacct attacccgaa gtataccgag

3361 gatttggtgc ataatgaggt catgcttccg gtcaatgcac cagcagaata ttcagaccca
3421 tcggtgctat ggaactccat tgaaatgagg gagagtaaat ctgccaaagc acagctggcg
3481 aggtcatata aaatcaatct gccgaatgag tggagttatg agcttgccat tgaggtaatg
3541 agggattatg tgaaaagaaa ttttgtggat gatggaatgt gtgcccagtt tgccatccat
3601 gactccgaaa acccgaatac gcaccagcgc aatcttcact gtcatattat gcttaccatg
3661 cgcccgattt tggaagatgg ttcatgggga gataagcaga agaagattta tgcactggac
3721 gcagacggaa acaagataag aaaaaagaat gggcaatata aatgcaccac gcaggatgtg
3781 actggatgga acagcaggga gaatgccagg aagtggcgca aggatttagc tgacaccata
3841 aatgcggtca atgataaaaa cggattaagg gagaattttt gggagcatag gtcatttgcg
3901 gagcaggggc ttgataccat accacagata catttgggag agaaagccag tgcgctggaa
3961 agggcaggaa tacagacaga gagagggaat gtcaaccgca ggataatgga gcagaataag
4021 gctattctta ctgcaaaaat gcttgtggca aaagcagagg aacagttgca gaagcttgtt
4081 aattccaagc cagttgaagc agtacgcaat actgcaaatg aagtccttga tatgattagg
4141 gcagttgtcg gcaaaaaggg cagactggaa ttgccagtag tcaagagcaa atatttgagg
4201 aagatttatc aaagacagtc attgcaggag caggagagaa tggagcgttt tgttgtggca
4261 aatcagattg agagctttga gcagttgcag gagtttaaga cacaacgaga gccagcctat
4321 gagaagctgt cagcggagag acaacagctt gcggagcaga tagcaagact ggagcagttg
4381 cttgtggcat acagtgatta tgagccttat atcgtgtatc ataagacgag taattcgctg
4441 aaaggatttg ccaagcgtaa atatgataaa gagcatgaaa cggagcttag agaatacaac
4501 agttatcgtg caaaattaaa gaaaatgctg tcctctgatg agaagattac tccgaagaaa
4561 tggacagcgg aaaaggcgaa actggaagaa agattgcagg agagcaatcc cgactatgca
4621 agagtggtta cggagcttgc ctcaacagag gttattgagc ataacagaaa aatgcttgcc
4681 atgacccgag aagccgagca gaatcagcgc agtcagagca agaagaaaaa tgaacagtca
4741 ttataacagc gagagggcag tttgcaagca atggtttgta gctgtccttt ttgcatgaag
4801 ggagaaaatg catgaaggaa aaaaagatga aagggagcag gagtgctcca agacccacac
4861 gcattgttgt agagcgtaaa tatgtcgggg atgtgaagat ggaagaggta ttccgacaca
4921 tcgcagaaga ggaaatacag aataaggcaa acaaatttat tgaaagtgca taaattgttc
4981 tggaaaaggg cttcattaag aggtataata ctccttgatg aagtcccttt tcattatggg
5041 aggtaaccta tggaaaggga aaaaatgaat aaagaaatca ctgctttata ttgcagacta
5101 tccaaagagg acaagcagag tggcgagagc atgagcataa gcacacagaa ggaaatgctt
5161 atggattatg caaggcagaa tggttttagg aattgtcggt attatgtgga tgacggatat
5221 agcggtacta attccgacag agaagcattc caagaaatgt tggatgatat tcgtgatgga
5281 aaggttgcgg ttgtaattac caaagaccag tcaagactgg gaagaaatta tattgaaacc
5341 ggtacttaca tggaaatctt tttccctgag catggtgtca gatacattgc aattaacgat
5401 ggttatgact ctgataagga cgaccatttg ggaatgatgg cacccatccg aaacatcatg
5461 aatgaaatgt atgcgagaga tacttcaaag aaaatcaaat ctgctttgca tacaaggaaa
5521 atgcagggca agtacttgtg tacaacagct ccttatggtt acaaaaaaga cccgacagac
5581 agcactcatc tgataattga tgaagaggtt gcaccagtgg ttaggtacat ttatgatttg
5641 gcagaggaag gactgggact gcactgtatc tgtaaaagac tgcacgatga aaaaatcctt
5701 aaaccaagtt gtcacaagaa ggaaatgttt gagcggtttg tagacgatgg taaaatgtat
5761 gactgggata atgcgtatgt cagccagttg ttacacagtc ctacctatgc aggacatctt
5821 gcagtggatt gcaagccgac aatttccatg aaaaccaaga agcgcaagta tattgcattt
5881 gcggataggg aaatcatacg gaacacgcat gaagccatta taccgcaagc acaatgggac
5941 agagtacaga caatcatatc cagccgaagc agcagtttta acactgaaaa aacagagtat

```
    6001 gacaatatct ttcggggatt gttaaggtgt gcggattgtg agaggacaat gcttgtgaag
    6061 gtggagcata aacggattag ggataatgtt ctagacaaga ccttttattg ttgcagtacc
    6121 taccgtaagt acggaagaaa tgcttgtgga gcgcataatc ttgaagcgag aatgctacat
    6181 gaagctgtcc ttgcagacat tcagagacac gcagagcttg cagttacgga tagagaggaa
    6241 ctggttaggc acattgctaa gagcttagat attaagctgt cagagggcaa ggagaagcat
    6301 aagagagcaa taaagcagtg taaagcgaga attaacgaga tagatgactt gtatgcgcag
    6361 ttgtacgagg atttgtcaaa ggggcttatt cccgagcgta ggttcaagat gatgaccgac
    6421 agactggata aagagcaaga agaactgttg gaaaagctgg aacagtacga gagggagaaa
    6481 aacagtggga aagagcagtt ggcaaatatc caagacttca ttgatgaggt cagcgagtat
    6541 gctggtatta cagagttgaa tttcaaaatg cttcatcagc ttatcgacaa aattctagta
    6601 tttcccgccg aagagattga cggcgagaaa gtccaaaaaa tccaaattca ctacaaattc
    6661 atcggtgcat taggcatgat tgagtagtga aatctattat taaatcacct acgaaaacta
    6721 accaacagat gctggtcgtg aaggtgagtt agtagcaaga tggataatag aaaaagcagg
    6781 ggtaaggaag ccactaaaga gattatggat ttcctcacag acagataagg ctattcttga
    6841 tggatttaga aatcttaagc caggggcagc ttacgagaat ttatataagg cagcagtatg
    6901 cagggcagaa gcagattgga ttgtagggct taatgtaact cgtgctttga cttgtaagta
    6961 taatgctcag ctttcagcag gaagagtaca atcaccaacc cttgcaatga ttgtgcaaag
    7021 agaagaa
//
DEFINITION     Integrative and mobilisable element IME_7
    CDS           complement(1..1524)
                  /color=9
                  /locus_tag="F41_01422"
                  /note="Serine recombinase, protein motif:Pfam:PF07508.7"
                  /product="Serine recombinase [recombinase]"
                  /"ID=F41_01422
                  /protein_id="gnl|SC|F41_01422""""
                  /codon_start=1
    fasta_record    1..7451
                  /ID=F41|SC|contig000001
                  /color=5
    CDS           complement(1626..1820)
                  /ID=F41_01423
                  /color=1
                  /locus_tag="F41_01423"
                  /note="hypothetical protein"
                  /product="hypothetical protein"
                  /protein_id="gnl|SC|F41_01423"
                  /codon_start=1
    CDS           complement(1893..2837)
                  /color=9
                  /locus_tag="F41_01424"
                  /note="Plasmid recombination enzyme Mob_pre, protein
                  motif:Pfam:PF01076.13"
```

/product="Plasmid recombination enzyme Mob_pre"
/"ID=F41_01424
/protein_id="gnl|SC|F41_01424"""
/codon_start=1
CDS 3393..4172
/ID=F41_01425
/color=1
/locus_tag="F41_01425"
/note="hypothetical protein"
/product="hypothetical protein"
/protein_id="gnl|SC|F41_01425"
/codon_start=1
CDS 4318..5013
/color=8
/locus_tag="F41_01426"
/eC_number=3.2.2.9
/note="5'-methylthioadenosine/S-adenosylhomocysteine
nucleosidase, similar to AA sequence:UniProtKB:Q9KPI8
protein motif:CLUSTERS:PRK05584 protein
motif:TIGRFAMs:TIGR01704 protein motif:Pfam:PF01048.14"
/product="5'-methylthioadenosine/S-adenosylhomocysteine
nucleosidase [nucleosidase]"
/"ID=F41_01426
/gene="mtnN"""
/protein_id="gnl|SC|F41_01426"
/codon_start=1
CDS 5057..5857
/color=7
/locus_tag="F41_01427"
/note="helix-turn-helix transcriptional regulator immR,
similar to AA sequence:UniProtKB:P96631 protein
motif:Cdd:COG1426 protein motif:Pfam:PF12844.1"
/product="helix-turn-helix transcriptional regulator immR
[transcriptional regulator]"
/"ID=F41_01427
/gene="immR"""
/protein_id="gnl|SC|F41_01427"
/codon_start=1
CDS 5894..6409
/color=1
/locus_tag="F41_01428"
/note="hypothetical protein, ab initio
prediction:Prodigal:2.60"
/product="hypothetical protein"

/"ID=F41_01428
/protein_id="gnl|SC|F41_01428"""
/codon_start=1
CDS 6583..7449
/color=2
/locus_tag="F41_01429"
/eC_number=2.7.7.-
/note="Aminoglycoside 6-adenylyltransferase Streptomycin adenylyltransferase, similar to AA sequence:UniProtKB:P17585 protein motif:Pfam:PF04439.6"
/product="Aminoglycoside 6-adenylyltransferase Streptomycin adenylyltransferase [antibiotic resistance]"
/"ID=F41_01429
/gene="aadK"""
/protein_id="gnl|SC|F41_01429"
/codon_start=1
BASE COUNT 1862 a 1752 c 1760 g 2077 t
ORIGIN
1 cagcgtattc aaaagttcgg cggtcagttc tgttggattg acgcactctt tcatcaaggc
61 aatccatttt tctgcgtctg cggcgttctg gctttcagtg gcgatagcgg attgaagctg
121 ttcaatcttt gtgtccagtt cagcctgttc gctctggtac ttcccggaca gcatagagaa
181 gttgtattct gtgatacgcc ccgccgccca gtcctcatac atccgggcaa acagggtatc
241 gacttcggct ttccgcttct ccgctttgtt caactctgcg gcttgcttct ttcttgcggc
301 ggcctgtccc ttgtcattgg catttaacag ccgcttcaaa agccgttctt catcatgctg
361 taccagcctc gaccagtatt gcagacggga gaggacatag gcgtaaagca catcatagcg
421 gatataatgc atggagcatt ggcgtgtgcc ctgtccgtac ttgctacaat gataatggcc
481 gtaaggcttg ctgttctgcc tgttctcccc ataggacagc gaccagccgc aatccgcaca
541 ttttaccaat ccggaaaaaa tctgcgtcgt accatccttg cactttctgc ggcggtttgc
601 tatctgctcc tgtacctgcc ggaaaacctg ttcgctgata atcggctcct gcgtgttctc
661 tacccgcacc cattcacttt ggggcttgcg tatcctccgc ttgttcttaa aggaaatgtt
721 tgtttcccgg tagtggatgg tatggccgat ataggtttcg tctttcataa tgctcttgac
781 ctgggctatc gtccatgcgt agcttttttc ctctggcgca cccgcataga tgtttgcaaa
841 agtcccgtca cgctggaaat tgataaatcc cggtgtggga accttttcca taatcagtat
901 tcttgtgata ctggccgccc ctctgccatg aatggcaagg tcaaaaatct tctccactat
961 ccagcgggtt tcctcgtcga ttatcagctt gtttttgatt tccgggtgtt tcctgtaccc
1021 gatgggagca taggcaccga tacgctgtcc tgtggcaaac ttcgctttga aagcggcctt
1081 taccttgcgg ctcgtatctt tcgcaaacca ttcattaaac aggtttttga acgggacaaa
1141 atcggaaagc cctttctctg tgtcctcatt ctccgcaacg gcgatgtagc gcacccgttt
1201 ttccgggaac agaaattcca gatagtagtc catcatcaca tgttcccgcc cgaaacggga
1261 aaggtctttc gtgacaatgc agtttacttt tccggcttcc acatcgtcca tcatgcgctg
1321 aaaatccggg cgttcaaagt tcgtccccga ccagccatcg tccacatact caccgactac
1381 atgaagtccc tgttcctttg cgtactgttg caggattgtc cgctgtgttt caatgcttac
1441 gctgtcacca tagttttcat cgtcccggct taatctcata taaagcgccg tgttgtaaat
1501 cgtagtattg taaggttgtt tcaccgttaa aaatcctcct tcttaaagaa acaacccacg

1561 cttacaatac ttttgctcta tggcaattat atcataagcg tgggcgtgtt atcaatgatg
1621 ggctatcagg ttgaagcggc tttttctgcg gtatgccgca ccacatctac aatcagatca
1681 ccgaggggct tcccgcctgc ggcgaagtgt tcggagattt ttatacggtt gttcccacat
1741 acaaaatact gcgtgccgtt ctctgtttgt atgacctgcc ctgtccgggg cgtaaaaata
1801 tcgctttcgc tttttgccat ccagtgtcct ccatattcag ttttcaaggt acaatgccgc
1861 acaggggcgg cgaaaatttc tgcttatatc tatcaccttt cctttactgt gtgccgctgc
1921 tgccgtttca cctgcgccag cacttccggc gggatacggt caacgaaatt gcggagattg
1981 tgtaattcgc tttccagcgt tgcccgttcc atgcggtctt tcatcttgcc ttgctcactg
2041 gcctttgccc ttgcttccag ctttttgttt tcttccagca ggtcattgat tgtgaccttg
2101 tactttttca actgcccgga gaagttctcc atctgcggga accacttttt cagcatggag
2161 agggcttcct cttttttctt tccggcgttc agcgggttaa tgccgtcaag ggcggcttca
2221 atggctctgg cctgtttgga gagggaaacc gcctgtttga aaagccgggt ggggatatgc
2281 ttccggcctg tcttgctggc actctcccca cgctccaagt cgggatattt ctccaccata
2341 taggcgtgaa aatcgtcctg ccacttcgtc aggtttgccc ggttgccgat aatctcctta
2401 gcgcacaggc ggttgtcctt tgtcagcgga acaaaggtca aatgcaggtg gggcgttttc
2461 tcgtccatgt gtaccaccgc cgacacgata ttttcccgtc ctacccggcc aatgaggaaa
2521 tctgccgccc tttggaagaa cgcctgtatc tcctttgggg atttcccctt gaaaaactcc
2581 gggctggcgg ttatcagcgt atcgacaaac cgtgtactgt ctttccttgt ccggcatccg
2641 gcctgctcga tacggctctg aatgaagtgg taatagcgtc cctccggctt gacgatatag
2701 aaattgtatt tgctccggct tgtgtcaatg tcggggttgc tggcgtattg ttctttctgt
2761 ctttcgtgat gggcttccag cggccttgcc gggttgccct tatgcttctc aaaccgcaaa
2821 attgcgtgtt gtgccattct gctcctttcc ccattccttt cctatccggg gttttccaca
2881 gggaaaatcc cgccggaatt atctctgata cgataggaat ggaatggata taaataaatc
2941 attttaatct ttgtatttct atataccgtc tggttttcgt acttctcaag attgatttcc
3001 gcacttcata agtacggttt tcagtcctcc ggcgtaccag aacgggattt cttgaagtcg
3061 gtgtttggaa ccacttcata ggattttggg aaaatgcggt tgggctttcc acagccctgc
3121 ttctggattt ccaccagtcc ggcgtattgc agttcccgca gggtgttcac cgctttctgc
3181 cgcccacagt ggagcaggtc aaccacttca cagatggggt aatacaggaa aatccgtccg
3241 cagtcatccg cccacccatt cttgcgggat aactctgtcc ggcgcaggat aaaggcgtac
3301 agaacctttg cctcgttgga caggggcttg aatgtagggg cttcaaagag gaaattgggg
3361 agccgggtga agctgaacgc cttttccggc tgatggatat agatggtatt tgtcatagtg
3421 cgttttgtgg acggttaaaa gcccgttttc cggggcggac gatactttat accacctgcc
3481 ccggtttggg gcttgcggag cctgataaat caaggctttt ttcactccta actgtccaca
3541 gcagacctcc ttttccgttc actttctgtt ttctgccgcc tgtgaactct gacggcacag
3601 ccggggcagt attttgcccg gttggatttg gggacgaaca cgccgccgca gaccgcacag
3661 catttcaagt ccttatctcg gaaaatctcc gcttccagcg ttccgtccag cggcaagacc
3721 gcccagcgga accacttaca gcagaccgag aaagaaaccg tctgcgggca ggtgcaggtg
3781 tccccatcgt caaggacaat gcagttgccg tcctcacagc aacagcactc ccggcgtatc
3841 aggctggccg cctgtttcct ctgcgccggt gtcatgcggt aaagggaacc gtccggcctg
3901 cgctccagtg gcggcaagtc tttatagggg ttatctctca tgcgttccct ccattttgct
3961 ttccgttgcc gttctccccg aaaaggtttc ctgccccatt cctgcggcgt attccggcgt
4021 tggcacgctc cccgcaactt cgggacaact tgtcccgaag tgacctctgg acaaagtgtc
4081 cagaagctgg ctccttgcag tgcttcccca gccggggtat tccttttcgg acggtcattc
4141 tgttttcaag gtgccgttca tcgatgaact aaaataagtc tacacctaaa tgaccgtccg

4201 tcccgtatat ccataaagtc ggagatccgc ccggaaaact gcctatttcc acgctctcgg
4261 aattgtggta aaataaacat aaaaattggc tttcaagcca taggaggaca ataatctatg
4321 aaaatcggaa ttttatgtgc gggtgacgaa gaactcgccc catttttgcc cttgataagc
4381 aactgtaaga taacggaaaa agctatgctg aaattccatg ctggacagat tgatagtgta
4441 gaagttgtcg ccctgttttc cggcgtgtgc aaagtaaatg ccgccatagc cgctcagctt
4501 ttgatagatg tattttgcgt ggatataatt atcaattcag ggacagcggg cggtatggag
4561 ccagaacttg aaatattcga tacagttatc tccacagaag tttgttacca cgatgtagcg
4621 ccagacattt taacggaatt tcatccgtgg atgaattctg tgttttttgt agcagaccca
4681 aagctgatag atatgtcaaa gtctgctgtt gacaagcttt caacttctgg aaaggtagtt
4741 tggggccgta tggtaacggg agaatcgttt ataaccgatg agagccggca aaaaataaac
4801 gatgaatttg ctcccttgac ggttgacatg gaaactgcca gtatcgctca tgtctgctat
4861 gcaaatgaca taccgtttat cgccattcgg tgtgtgaccg atactgaaaa acacagcgga
4921 gttgataatt ttgacaaaaa ctgtgcaaaa gcatcaagta ttgccaaaga tattacggtt
4981 gctttattaa gagaaatcaa aaagtcgtgg taagattgga tgatatttta gaacagagga
5041 cagagggagg gagagtatgg ccctaactat tcaagaacga ctaaaagacc tgcgtgtgga
5101 gcgcggcttg acgctggaac agcttgcgga gcagacccac ctctccaagt ctgctttagg
5161 cagttatgag gggacaact tcaaggacat cagccactat gcccttatca agctggcaaa
5221 gttttacggc gtgaccgccg actatctgct gggccgttct caaacaagaa atcacccaaa
5281 cgccgatctt gcagacctgc atttgagtga tgatatgatt gaactattga aaagcgggct
5341 ggtggacaat tctcttttgt gtgaactggc gacccacccg gattttcccc ggctcatggc
5401 cgaccttgaa atctatgtga acggcgtagc gggaaagcag gtgcagaacg caaacgccat
5461 tgtggacgct gtgagtgcaa cgattatgaa acagcacaat cccggcttga ctgacccgca
5521 gttaaggcag cttatcgccg cccatattga cgacgacagc ttttgccgct atgtgataca
5581 gcaggatata aacaagatag ccctcgacct gcgggaagcc cataaggacg atttttttcag
5641 tgtcccggag gacaacccac tggaagattt cttgcagacc gtagaggaaa ccaccagccc
5701 ggacagcgac ccggagcagg cggctatgat gtttatctgc aagcggctca aactgaatta
5761 taagaagctg tccgaggaag aaaagaagtg gctgaaaaag attgcacaga agtcggactt
5821 gctgaaaaat ccaaagccac agcgaggacg gaagtaagag aataataaat cagattatat
5881 ggaggtattg gttatgaaaa aatggttact gataatttca gcgactgtta tttgcgttgc
5941 tgccgctttt ttatatttct ttagtttaac tcccaaagga gataccatta ccagcagaga
6001 aagcatactg aacaccgcaa tttcaaaggg gaatgaatgg actattgcaa aagaactgga
6061 acttggcggt tatatcgtga gtggagcata tagtgcagat aataaatcta cacttgccat
6121 ctttgaacca acaggaaatg gagattacaa atttagtaca tcaacaaacc gaaatagtga
6181 tgagattatt gttggtggcg ttgcaataaa cggagaatgg tatgatttaa tttggtttaa
6241 tggtgctaaa acagagtatg ccgaaatcac ttatacaata aatggacaag tgcaagatac
6301 attgagatac agcacagatg atatggatat tatctcaatt aaaaacccag aaaaagagta
6361 ttccattcat gtggtttact atgacaatga tggaaataag tacgaataaa ttgtttttca
6421 tcgagaaaat agcaaagcca gccgagccag tcaacggtca agatgaacgg cgcatttcat
6481 gcgccgccgt tgacagtccc gcccgtcttt gctaaagggt aatcaaggcg ggaaagccct
6541 aaaatggttt cacacctatt tcaataattg gaggagataa aaatgagatc agaaaaggaa
6601 gtttatgata ttgttttgaa ttttgcaaaa acagacaaac gcattcgcat ggttactttg
6661 gaaggatcta gaacaaatac aaatattccg cctgatgatt ttcaggattt tgatattact
6721 ttttttgtta cggatatgga cagcttcaca agtgatgata aatggctaga tatatttggt
6781 gaaaggttga ttctgcaaaa gccggaagat atggaattat ttccagctgt agaaaaggga

6841 ttttcatatt taatgctgtt tactgatgat gttaagatag atttaacttt gctgccgctg
6901 gaactgatag acgagtattt tacatgggat aaactggtaa agttactgtt ggataaagac
6961 aaccgtatcg taaagccgcc aataccaacg gatatagact accacttgca gaagcctact
7021 caaagaatgt ttgacgattg ctgtaatgaa ttttggaata ctacaacata tgtagtaaag
7081 ggcttatgcc gcaaagaaat tctttttgct attgaccata tgaatgatat agtacgaaaa
7141 gaattgcttc gcatgatttc ctggctgatt ggtatcaaac agggatttca tttcagtttg
7201 ggaaaaaact ataaatttat gaagcaatat gccccagagg aattgtggga acgacttatg
7261 tccacttata atatggattc ctatccccat atgtgggaat cctttgaaca atgtatggca
7321 ttgttccggg aggtttcgtc agaagtggca tgccagttgg attaccagta tccactatat
7381 gatgaaaaaa tcagtaatta tgtgattcgg caaaagaaaa aatatggcat tgaagatgat
7441 aacaaataaa a
//
DEFINITION Integrative and mobilisable element IME_8
CDS complement(1..1920)
/ID=F41_02236
/color=9
/locus_tag="F41_02236"
/note="protein motif:CLUSTERS:PRK13413 protein
motif:Pfam:PF00239.15"
/product="Serine recombinase [recombinase]"
/protein_id="gnl|SC|F41_02236"
/codon_start=1
fasta_record 1..24717
/ID=F41|SC|contig000001
/color=5
CDS complement(2314..3156)
/color=10
/locus_tag="F41_02237"
/note="DNA replication protein DnaC ATP binding protein,
similar to AA sequence:UniProtKB:P0AEF0 protein
motif:CLUSTERS:PRK08116 protein motif:Cdd:COG1484 protein
motif:Pfam:PF01695.11"
/product="DNA replication protein DnaC ATP binding
protein [replication]"
/"ID=F41_02237
/gene="dnaC"""
/protein_id="gnl|SC|F41_02237"
/codon_start=1
CDS complement(3153..3680)
/color=10
/locus_tag="F41_02238"
/note="Replication initiation protein RepA , protein
motif:Pfam:PF06970.5"
/product="Replication initiation protein RepA

[replication]"
/"ID=F41_02238
/protein_id="gnl|SC|F41_02238"""
/gene="repA"
/codon_start=1
CDS complement(4417..4860)
/color=8
/locus_tag="F41_02239"
/eC_number=3.1.-.-
/note="DNA mismatch endonuclease Vsr  Very short patch repair protein, similar to AA sequence:UniProtKB:P09184 protein motif:Cdd:COG3727 protein motif:TIGRFAMs:TIGR00632 protein motif:Pfam:PF03852.9"
/product="DNA mismatch endonuclease Vsr  Very short patch repair protein [nuclease]"
/"ID=F41_02239
/gene="vsr"""
/protein_id="gnl|SC|F41_02239"
/codon_start=1
CDS 5409..5738
/color=7
/locus_tag="F41_02240"
/note="Helix-turn-helix transcriptional repressor RghR, similar to AA sequence:UniProtKB:O32236 protein motif:CLUSTERS:PRK08154 protein motif:TIGRFAMs:TIGR03070 protein motif:Pfam:PF01381.16"
/product="Helix-turn-helix transcriptional repressor RghR [transcriptional regulator]"
/"ID=F41_02240
/gene="rghR"""
/protein_id="gnl|SC|F41_02240"
/codon_start=1
CDS 5758..6810
/color=8
/locus_tag="F41_02241"
/eC_number=2.1.1.37
/note="DNA cytosine-5--methyltransferase, similar to AA sequence:UniProtKB:P20589 protein motif:CLUSTERS:PRK10458 protein motif:TIGRFAMs:TIGR00675 protein motif:Pfam:PF00145.11"
/product="DNA %28cytosine-5-%29-methyltransferase [methyltransferase]"
/"ID=F41_02241
/gene="haeIIIM"""

/protein_id="gnl|SC|F41_02241"
/codon_start=1
CDS complement(6857..7468)
/color=8
/locus_tag="F41_02242"
/note="TIGR02646 family protein  HNH endonuclease,
Pfam:PF01844.17"
/product="TIGR02646 family protein  HNH endonuclease
[endonuclease]"
/"ID=F41_02242
/protein_id="gnl|SC|F41_02242"""
/codon_start=1
CDS complement(7452..9101)
/color=8
/locus_tag="F41_02243"
/note="AAA family ATPase  Predicted ATP-binding protein
involved in virulence, protein motif:Cdd:COG3950"
/product="AAA family ATPase  Predicted ATP-binding protein
involved in virulence"
/"ID=F41_02243
/protein_id="gnl|SC|F41_02243"""
/codon_start=1
CDS complement(9193..11049)
/color=9
/locus_tag="F41_02244"
/note="Serine recombinase, protein motif:Pfam:PF00239.15"
/product="Serine recombinase [recombinase]"
/"ID=F41_02244
/protein_id="gnl|SC|F41_02244"""
/codon_start=1
CDS complement(11104..11367)
/ID=F41_02245
/color=8
/locus_tag="F41_02245"
/note="Stage II sporulation protein R"
/product="Stage II sporulation protein R [sporulation]"
/protein_id="gnl|SC|F41_02245"
/gene="spoIIR"
/codon_start=1
CDS complement(11367..11597)
/color=8
/locus_tag="F41_02246"
/note="Seryl-tRNA synthetase ,protein motif:Cdd:COG4443
protein motif:Pfam:PF09901.3"

/product="Seryl-tRNA synthetase [RNA synthetase]"
/"ID=F41_02246
/protein_id="gnl|SC|F41_02246"""
/codon_start=1
CDS complement(11643..12497)
/color=10
/locus_tag="F41_02247"
/note="ParB/RepB/Spo0J family protein Plasmid
partitioning protein parB, similar to AA
sequence:UniProtKB:Q83AH2 protein motif:CLUSTERS:PRK13832
protein motif:TIGRFAMs:TIGR00180 protein
motif:Pfam:PF02195.12"
/product="ParB/RepB/Spo0J family protein Plasmid
partitioning protein parB [partitioning system]"
/"ID=F41_02247
/gene="parB"""
/protein_id="gnl|SC|F41_02247"
/codon_start=1
CDS complement(12506..13471)
/color=9
/locus_tag="F41_02248"
/note="Plasmid recombination enzyme Mob_pre, protein
motif:Pfam:PF01076.13"
/product="Plasmid recombination enzyme Mob_pre"
/"ID=F41_02248
/protein_id="gnl|SC|F41_02248"""
/codon_start=1
CDS complement(13996..15021)
/ID=F41_02249
/color=10
/locus_tag="F41_02249"
/note="Replication initiation protein RepA"
/product="Replication initiation protein RepA
[replication]"
/gene="repA"
/protein_id="gnl|SC|F41_02249"
/codon_start=1
CDS complement(15480..15650)
/ID=F41_02250
/color=7
/locus_tag="F41_02250"
/note="RNA polymerase sigma factor Sigma-70"
/product="RNA polymerase sigma factor Sigma-70
[transcriptional regulator]"

/protein_id="gnl|SC|F41_02250"
/codon_start=1
CDS complement(15734..16159)
/color=7
/locus_tag="F41_02251"
/note="RNA polymerase sigma factor Sigma-70, protein motif:CLUSTERS:PRK12522 protein motif:TIGRFAMs:TIGR02937"
/product="RNA polymerase sigma factor Sigma-70 [transcriptional regulator]"
/"ID=F41_02251
/protein_id="gnl|SC|F41_02251""""
/codon_start=1
CDS complement(16254..16766)
/color=7
/locus_tag="F41_02252"
/note="RNA polymerase sigma factor Sigma-70 region 4, protein motif:CLUSTERS:PRK08583 protein motif:TIGRFAMs:TIGR02937 protein motif:Pfam:PF04545.10"
/product="RNA polymerase sigma factor Sigma-70 region 4 [transcriptional regulator]"
/"ID=F41_02252
/protein_id="gnl|SC|F41_02252""""
/codon_start=1
CDS complement(17479..19398)
/color=2
/locus_tag="F41_02253"
/note="Tetracycline resistance protein tetM from transposon Tn916, similar to AA sequence:UniProtKB:P21598 protein motif:CLUSTERS:PRK13351 protein motif:Cdd:COG0050 protein motif:TIGRFAMs:TIGR00484 protein motif:Pfam:PF00009.21"
/product="Tetracycline resistance protein tetM from transposon Tn916 [transcriptional regulator]"
/"ID=F41_02253
/gene="tetM""""
/protein_id="gnl|SC|F41_02253"
/codon_start=1
CDS complement(19765..20139)
/ID=F41_02254
/color=9
/locus_tag="F41_02254"
/note="Transposon-encoded TnpV"
/product="Transposon-encoded TnpV"
/protein_id="gnl|SC|F41_02254"

/gene="tnpV"
/codon_start=1
CDS complement(20120..20368)
/ID=F41_02255
/color=10
/locus_tag="F41_02255"
/note="ATP-binding protein  AAA family ATPase  helicase"
/product="ATP-binding protein  AAA family ATPase  helicase [helicase]"
/protein_id="gnl|SC|F41_02255"
/codon_start=1
CDS complement(20401..22245)
/color=8
/locus_tag="F41_02256"
/note="Restriction endonuclease  Helicase-associated Z1 domain, protein motif:Pfam:PF10593.3"
/product="Restriction endonuclease  Helicase-associated Z1 domain"
/"ID=F41_02256
/protein_id="gnl|SC|F41_02256"""
/codon_start=1
CDS complement(22257..23408)
/color=8
/locus_tag="F41_02257"
/note="NgoFVII family restriction endonuclease, protein motif:Pfam:PF09565.4"
/product="NgoFVII family restriction endonuclease [endonuclease]"
/"ID=F41_02257
/protein_id="gnl|SC|F41_02257"""
/gene="ngoFVII"
/codon_start=1
CDS 24052..24363
/ID=F41_02258
/color=1
/locus_tag="F41_02258"
/note="DUF4258 domain-containing protein"
/product="DUF4258 domain-containing protein"
/protein_id="gnl|SC|F41_02258"
/codon_start=1
CDS 24369..24596
/color=8
/locus_tag="F41_02259"
/note="Type II toxin-antitoxin system MqsRA family

antitoxin MqsA, protein motif:TIGRFAMs:TIGR03831"
/product="Type II toxin-antitoxin system MqsRA family
antitoxin MqsA [toxin-antitoxin]"
/"ID=F41_02259
/protein_id="gnl|SC|F41_02259"""
/gene="mqsA"
/codon_start=1
BASE COUNT 6783 a 5586 c 4132 g 8216 t
ORIGIN
1 gccatcataa caatgtaaat aatcgttcga tagctgtttt gcactcgttt gagccggtaa
61 attcggtaaa tattggtgta tcagcgccat gatttcctct ttcgatttta caatcttgta
121 tggtttcagc gaattatccg tatcttccag aatccccagt tctttcatca gatggatata
181 aggcaagata ctttgtttcg ttttcttaaa tcccaatgct gtgatatgtt ctgacaagta
241 ttttgcagat gtatatccac gttcatcttc tataatcagc ttcattacaa gagcgattac
301 agtattttcc cggcgattca tctcatcagc aggactgtaa ttgattaaat cagacaaacc
361 atattttaaa gtgatttctg gcattccctc tttatctacg tcaattcgct caatcaacag
421 ttcaatatct cttcggtcaa gagttccacc tgcaatgatt ttatcaacta catccaatgc
481 gtttctcaat ttgtccttta catctggtgt ttctatagta gtttcgttca gttctttttc
541 ctgtatctcc aaaccatgaa tctgtgccag taaatccttt tgcatggaat cataggtttc
601 gtggatcaaa tcttcatttt cagatgctat ggataaatct tttatcttct gtgtcaaaag
661 taccttcaac tgatttttc gattttgaat ttcttcttga aggtcctgtc gcttttcttc
721 cagcgttttc ttttctgcct caaaatccct catatcgtac gtggaaatca cttcacaaag
781 cgcctctctg cacatcttaa tataattcag cacatcttca atcaaatcat tttcttcaat
841 cagatgtgct ttttcacaat atcgtctgcc ttttgtatta taggtggtac aaatataata
901 ttttctttct ctgctggatg tttgacgctt aatcggtgtc aaacgacttc cacaatcttt
961 acaaaacaaa cagcttccaa aaggattggg aatttctgaa ccgatccatt ctcccctgct
1021 tccacgatag tttgtgcgat tacgcttttc ctttaattcc tgtacaagag aaaacgtggc
1081 tttatcaata attgtcggat gatgatcttc aaaaatacat tgttcttctt ttggaacacg
1141 tttatccttg ccatgtaccg tgtttctggc acgttttctc aaacgaaatg ttccaatgta
1201 aaaatcatta tctaataatt ctttcaccat gctgtcagac caattggtgg cgactttctt
1261 tttgctcacg cgaccttctt caatttctcg ttcccgctgt accatagagg gtgtcggaat
1321 accctgatcg gtcagataat ttgctatttt acgataaccg gaaccggaga tataaagatc
1381 gtaaaccatt tttacaaatt ctgcctcttt gggaacaatt cggataatgc tcttatcttt
1441 cgcatcccga caatatccga aaggtggttt tgtcataaga gtgccttctt tttgtctagc
1501 tccgattgct cttttgattt tcttactggt atcttttaca taacgctcat tgaaccatgt
1561 tttaattcca atggtgtcgt cgctggaatc aaaagaatcg taattatcat caatgacaat
1621 gagatgtttt ccgcgttcct gaaattcatc caaaagcaaa agaaccttcg cattatgtct
1681 gccaagacga gagaaatctt tcacatatac ggtatcaata tctttatcta aatcagccat
1741 catctgttga aatcctggtc tgtcaaaaat ataacctgat actccgtcat cttcatacca
1801 tcggtcaata accattccat ggtcagatgc atactgattg ataattaact tctgattttc
1861 tatagaaaca taatttcgct tatcgtcatc tcttgaaaga cgcacatatc cagcattcat
1921 gtcctcatcc tttctgaaat attatttata gaataatgta atataattat atatcgaaaa
1981 atctttttcg taaacggatt atcatataat tttacaatga attatattat tgtgaatagt
2041 aaatatgggc gagagaatgt ctcccgccca tagaaattac tgattgattt cttttatatg

2101 gtttcgaatc aacgctttta ttccattatg tatattttta tccggctgtc catctgcaaa
2161 cgtgacaacc cgaaaaccta ttttcgtata tttctcctta tctgcctcaa agtcttctga
2221 agatgtataa tctgttcgcc acaggtaata tgtaggtttc tgcagagcta tcacactctt
2281 tctatttctg atgtctatac taagatataa acatcagatt atttttgtgc ctgtctgctc
2341 tataatattt tcagtctgac ttgattcggc agtagcgaaa aaggttttca ttttctccat
2401 tttttcatgg gcaatatcac tccgatagct agaaccgcca acatataacg gaacacacat
2461 ttctaaaata cgatcgtaaa ttcgtttctt ttctatgttc gtttcttctt ttaattgttt
2521 tatcggaaga ttggtagtta cgattaatgg tctgccggag cgatatcttc gatcaattac
2581 gctgaataca attccaagag catactcgga acttcgttct gtgccaaggt catccagaat
2641 taaaagttca tattttgcca gagcattgat atattctgtt ttatcctcta atggaaacat
2701 atcatcaata attgtattaa aatttgtcat cttcacagta atttcctgat tcaaaagtgc
2761 gttggcaatg caaccggcta tgtaactctt tcctgttcct attggtcccc acaagagaaa
2821 accaatatga tttttcttca tatcgttcca gttcgctgta tattcttttg ccttctcgat
2881 ggtggtagtt tcttctgcca gttcaaaatt ccaatcatac attcgtgatt cttcaaaaca
2941 gatttttcta tttctctcca ctaattcgca atgttctctt tctttatatg ccttttcttc
3001 tgcttcacgt ttttgacgga tacatgcaca ttgtctgaaa tgtttttttca taccaagaat
3061 actgtctgga ggaaaatatg cctctacagg ttctccacaa atcttgcaat atgctaatcc
3121 atctgcacct aaatatgtgt tttcctgctt tatcatagac tgtcgccctc ctcatattcg
3181 taatttactc caatcttatt actaaaatta tatttttctt ttactttact aggtggcaca
3241 tttccgacca tagaagaggc tgttttccga ctgaatgaag gacagctttt catcctcatg
3301 gaagttgtat tttcgtctgt cggctttaca tccggttcag aaatcaactc tatacagggg
3361 attcgcatat ataaatgatt tggttttgaa aatcctccgg aactgcgaat caacaatcca
3421 tactcatcta attccttcat ggcagctttt acagaagtac aacattttcc ccattttaaa
3481 gaaatttctg taattggaaa gacgaaatat acatacccgt cttcattgag ccaatggttt
3541 ttctgtgaaa ttctggctct gtcatataac aacatataaa gtaattttgc attttgagag
3601 agtggtagct ctaacaaaaa tttagggaac tgataatagg gaagtatctt ttccgtttcc
3661 atcatatatt caaatttcat attttctttt acctttctga aaaattaagg agacggatgc
3721 cggctttccc aaagagattt tatgctttat aaaaattctc ttttgatatc tcctcatata
3781 acgaaaggaa aaaggctcat tttacagggt acaaagaaaa aaatttataa gagcaagatc
3841 cacccatgtt taacgttatc gcgtttccgc ctccgttcac cttatgggtg ttctcttgat
3901 ggggagtgac aattccccat acccttgctt atataattgg cttacaaaag aatttgttcg
3961 catcaccatg tgtattttca aaaagaaaaa aacatccaat tacttattga atgcctcaaa
4021 atagaattat gttaattatg atattttgct gcgtacaccg ccggtgtagc taataaaaaa
4081 gtgggtgtgg ggaggcgaca gttccccagc aagataaaac tccgaaatat tacttatatg
4141 gcagaatggc gatattgtaa tacggggtgg atcttgctct taataatttg aaaaagaaaa
4201 aatgctgaaa atactatttt caaagatacc tgatattcag agtgtctcag attttttatg
4261 atgcttttac aatatgattc tgataatttg ccttatgaga aggtgggcaa gaatgagccg
4321 gtttgcattg cagatgaggt tccgttgata tctaaatatg acaacaatgc atatccacca
4381 aattcattta tacttggcgg atgtgcatca actatattat tcaagttctt gactttgcat
4441 ttttaattca aaaatagttt cttcaataac ccgtacacat tcatctgtat tattttttat
4501 atcttttccc cagaatctga ctacagtcca tcccataaat aacaatttct tattaatctc
4561 atcgtccctc tcacggtttc tggaaatctt ctccacccaa tactcactgt tatttccttt
4621 ttcaagtctt ggttttaaaa cctcccagtc ttttccatga aaaaactccc catcacaaaa
4681 aatagcaatt ttatactttg tcaatacaat atccggtttt ccaggaagat tcttataatt

4741 ctttctatat ctatatcctt tattccataa cgcctttctt aatattgttt ctatttttgt
4801 attatcagac cgtatatgtt gcatattttt ctttctttgg cctggtgtca aattatccat
4861 atcgattacc tcaaaaagac ttcatcatta tgatatccaa gaaatgattt accctgcaaa
4921 aatttatctg gaagattaat ttgttttctt tcatatccat agaactattc tcttgtctgt
4981 ttatccattt caacattttt catttttagt ttactatcta atgttattaa tcctttatcg
5041 aaagccttat catgtaatga atttaaacat aaacaattac atagattcgt tctttcagtt
5101 ttttcgtcac agactctcca cagtttaata tggcttgcaa ttaacaagtc tgattttttc
5161 attcccatta tgtaacattt actccattgc gatttgccat attattttcc ttttcataat
5221 gtatattatt atctgaaatt tatatctcac atttacaata tactatatca catttttaca
5281 gcaagttatt ttttaatcac acaaattctt atagttcata ttttcgctgc tttttggcag
5341 cgttttatta attttttatt gacattaggc agcgcacgtg ctatactttc acaaggggag
5401 gagagaatat gtccaattta ggccaattta taaaatccaa aagagaatct tctggattat
5461 ctctgaaaaa attaagttcc gcaactggaa tcagtgacac tgaaatatta aaaatcgaaa
5521 atggtacccg taaaaaaccc agctgggaac atctttgtca aattgcaaat gcattaaatt
5581 ttcacccatt tgagatttta cttaacgctg gatacatatc tgaaaatgat attcatccac
5641 cattgctaat cagtggacta gaggatttga atgaacaaca actcaataca gttcaattat
5701 tcgttgattt tatcaaaact caggaaaaaa ctgtctgaac aattaaggag gctacttatg
5761 caatatagat taggagaatt attctgtgga ccaggcggaa ttgcatgggg tgccactcac
5821 gccaatatag gcaattctaa tttttcaatt atccatgaat gggctaatga ttatgatgcc
5881 aatacatgcg aaacatatag acacaatatt tgtccaaacg aaccacagac cgtataccat
5941 gaagatataa gaaaattaga tatgaataag ctttctccta ttgacgcatt ggcttttgga
6001 tttccatgca atgattatag tgttgtaggt gagcaaaaag gaatggatgg cacatttggg
6061 ccactttact cttatggtgt caaagcttta aaactgttca gacccatgtg gtttcttgca
6121 gaaaatgttg gtggattaag aagtgccaat gaaggtaagg cacttgcaaa aatattagaa
6181 gaactgcgag gcacaggata tacaattact cctcatttat ataaatttga agagtatggt
6241 attccccagg caagacatcg tttaataatt gttggaatca gaaatgatat tcctgtggaa
6301 ttcaaggttc catccccagc accttatgct aacgttaata acacatgtcg aaatgctatt
6361 gaaaatccgc cgattcctca tgacgcttat aataacgaac ttacaaaaca atccaagcaa
6421 gttactgagc gattaaaaca tataaaacca ggacaaaatg cttttactgc tgatttacca
6481 aaagaattac aactaaatat aaagggtgca aaaattagtc agatatataa acgtcttgat
6541 cccacaaaac catcttatac cgtaactggc agcggaggcg gtggtacaca tatctaccac
6601 tggtcagaac ctcgtgcact tacaaaccgt gaacgtgccc gattacaaac attcccagac
6661 tcttatgaat ttcaaggacc aaaagaaagt gttcgcaagc aaattggtat ggcggttcca
6721 tgcaagggtg cgcagattat ttttgaagca atacttaact gttttgctgg tattccttat
6781 gaatatgttg acgcaaatat taatgaataa gtcaaagggc aagtagtatt tttaattaaa
6841 tcttgccctt ttcttattat acgtaagtac ttatgcactc taataatatc tcatttgcct
6901 tgtccaatgt tcgttccgtc cttaatgttt cccataatat ttgttttgcc ctttcttcta
6961 catgacaatt ttcccctaag ccaaacatat ccattgtatt tcttgctttc atttcctcac
7021 ttctagttaa acctgaagat atacgaatta ttgcaccctc actttcaaaa taatgatcta
7081 catcatcaaa ataaggatgc acgattgtaa atgtacactg attgtagtaa atatttttca
7141 attctattgt atccttcctt cctttttcg aagaactatt gcagaattga caagacattg
7201 caagattttt tgttgtatat tcaaactctg gatataaatg tctcggagca atatgttcaa
7261 tttcccctct tgatgtccct cctagtttta atccacaata tacacatttt ccattctgcc
7321 ataatagcaa ttgcttcttg atactttcct taatttcttt atttatcttt tttccagaat

7381 tccagtcctc atacttggga attccaagtc ttcttatata ttttttatgt actctagaat
7441 atttgacatg attattcaga ttcaacattt cttatgtact ccttcgcttc ctcaataacc
7501 ttattcaaag gatccgtatt atccaaaata tatttagaca atttatcata taaatctcga
7561 atatgatcta actcttcttt tttattatta cttatggcat ataataattc ccgcacatcc
7621 atatcaaaat aataattcct aaatgtcctt acaccaaaaa tattatacaa gatattttca
7681 gcagaccatg catatgtact ataatccaaa gtcaagaaat gctttccccc ttcatctaca
7741 tttaccgtaa tcaatgaaga attttcagga ttcaaatccg aaaccatata atgcgaatga
7801 gaagcaataa taaaatggca agaattatat tctgcaaaaa cctttttcaa aaaactaata
7861 tacttaatct gccaattagg atgtaaactg atttccggtt catcaattag aactatagaa
7921 tttttgctta tattattaaa tatattagta aacgaataca atatatgttt ttccccagaa
7981 ctagcctctt caaatgaata tcttttccca tttttgtata gatttaatgt catattctta
8041 aataccttca aattagataa tcctttgaaa tatatgtagt cactggctat ttttttgagc
8101 tcatcttctg atgtattcat atcaaaggat actccaaata taattttact tttgtattta
8161 acataggaac tattcttctt aatcccatta ataaaatctg ctaattctcg atacgtttca
8221 tccgtatacc gctttaccgc atcgcttcta taatcttctt tctgtactat tttctcgcca
8281 taccctttaa gttttttatt caaatcatct gcaaagaaat ctctccccga catttcacac
8341 tcaatactta tttcagcatc aatacataaa aactccgaaa gcctattaat aaaattccat
8401 atttttccat ttactgctaa gttcaaaata gaatctgcca ttttattatt cattgaacta
8461 gtccacgatg cattagatgt cattctcacc cccaaatatt catatatact attcgcctct
8521 gtagtacttg actgtttgaa tggaaatttg tcatttaaca taaatgatac tgcaagagcc
8581 ttggttggca aaatagactt atcatatttt tctaacaaaa caccattctg atgtataaaa
8641 atttgtctat ttagaatttg tagctgaatc aattccccat ttatagtata tgtaatctca
8701 taacgagagt atttcatttg tcctaaagca ttttgacttt caatggcatt aaaaatttct
8761 gctaaaattt tcaatataaa actttttcca gatccatttg ctccaatgat caatgatgta
8821 gttggttcat tacttttatc attcaattct tcagagaaaa aacgtatatt acaatgttct
8881 ttttcaccat ctcgtaccaa taatgagata agtttgaatc ctgacatttc cacttcatca
8941 atcttttgaa cctttccatt ttcgtccgaa atattgaatt ttcttctttt attatcatca
9001 tccccagata tgaactcaat ttttatagtc ttatcatcat atatactaac tacaacataa
9061 tctccctctt ttgcattgtt taactcgtac cattcataca atacttttaa agaaccatta
9121 ccctgaaact gcttagcaaa atccctttta ctttgtgaac cagctgcctt taattcatat
9181 tcttgtccat tatcagtcgg tcattttttg cggacagttc atacatgcta tccttttctg
9241 ttctctgctc ctgcgtttgc tctcgtttga tttctccctc cgtatcttct tggcacaagc
9301 aggacaatac cttgatgcac cagagcccgg gacatattca acgccgcaca ccgtacaatt
9361 tttagcgggc attgctgccc accgttttgt aggtatgata tgtaattcat cgacaaagcc
9421 gatgaattta tagtaaatct tgatttcctg cttcactgtt ccgtctgcca tcttctcacg
9481 ctccgaaaca agtatcttgt ctatgagtgc gtttataact gttgcatcca gttcttttaa
9541 gccttgataa tttcggataa gggcgaggaa gtcacggatt ccccgtgatt tctcgtagct
9601 ttcattaaga gtttccgtca cctctttcag ccttgcttca atttcaagct gctctttctg
9661 gtatttcccc gacatcatct caaaattccg ctcggtaata cgctccatga ccttatcctc
9721 gtagagagag gaaaacagcc tgtccagttc cgcaaggcgt ttgttcagct tcttacgttc
9781 tttctctaat gctttcgccc tgctctggtc tgtttccgtg agccgctttt ctatggccct
9841 gaccgccttt tcatcattca ctgccatatc cgcaaaacag ttgatgtcgg caagaacggc
9901 attgaataaa tcccttgctt ctatattgtg ggcactacac tcgcttcttc cgtttcttgc
9961 ataattatta catgaatact gtacacagtc gataatctct gggcgtttcc tcctgtgtac

10021 gttcattgcc cgcagagcac atccgcagtc cacacacttg ataacgcctg caaaaatatt
10081 tacaaatcct cccttgttct gtggaagcct acgacttgta ataagctgtt ggacaatatc
10141 aaattcctcc tgcgtgacta ttccctcatg ggtattgggt atcacttccc attcttcggg
10201 cagcttagag gggcgtttct tgcttttcat attggcggca atccgtttgt agcctacaag
10261 atttcccgca tatatcgggc ttcttaaaat gctcctcacg ctgttcccac tccaaatata
10321 gcgtttgtcc tcgttcccct caaaatgacg ttcaaagcct gtttcgccac gctccgccgc
10381 ataagcggca gggcgtagga tatgctgttt attaagatgt ctgcaaattt tggcaactcc
10441 attccctttt aatgcaaggt cgaatatctc ttttacaaca tgtgccactt tatcatctat
10501 cagcagatgg ttgtggtcgg cagggtcttt gatatagcca taaggggcgg tagttcccat
10561 gaatttcccc tgttgaaacc tcgcccgata tgccgatttt atcttaacag atatgtcagc
10621 ggcatacatt tcgtttaaaa tgttgcggaa aggcgtgatg tccatagcag atttattcaa
10681 ggtatctacg ccgtcattga ccgctatata cctcacgtta tgctctggga agaaaacttc
10741 cagatataac ccacaatcaa gatagtttct ccccagacgg gataaatctt tcgtaatcac
10801 gcagtttatc agaccgcttt caatgtcttt tatcatattc tggaaacttg gtctttggaa
10861 atttgtacca gaataaccat cgtccacata cgtttttgct atgtgccatc cctgcttttt
10921 cacataatcc gtgaggatgg atttctgtgt cgcaatgctc gcactctcgt tatccgtacc
10981 atcgtcttta gataagcggc aataaatgcc gactaaatag attttctttt cttctttgat
11041 tcctgccata ctgtaaaacc tccgtatctg tcctatctgt tttcatgtcc cattgcgtac
11101 attctaagcg gacagcccct cattgtcgaa ggtgtcgccc tcggcaatct tcttccgaat
11161 atcctcggag ataatcggga caaacgcttc tgtaacggtc tgtgtgccga catattcacg
11221 gctgattatc atctgcactg gtgcttttgg cacgatacgc tttctctttt tatctgcttt
11281 cttatcctcg cccatactta aatcttcctt tccagacagg gcagggaaga gtttggaaat
11341 gtccccgccc tgccaatcag ataccattaa tcctcgctgt ttaattcttt ctgtaatgct
11401 ttaaggagtg ccgccgcttc atcagcgttc agcgtgattc ccttgccgca cttttcacgg
11461 ttcggggaaa agctgcggat gtcatacttc ggctctttcc cattccatga aatgagattg
11521 atttcctttg tgtagccact gtcgcccgta gacaatactg cgatttcctt tacgatttca
11581 tactggattt ctctcattct ccatacctca actctctgta tttacttccc gaaaaaagaa
11641 gattagcggc tgtcacggtt gcgtttcctc tgcaactcac gctccaaaag tttgatgatg
11701 gtttcctcca tctgcttcgg cgttgtgttc ttcggaaagt attttttcag cttgcttgtg
11761 ttgattttca aggtttcttt ctggttgccc ttttcctccg tcataatcgc aaatatcgta
11821 tccatgtcaa gccgcccgct ctggcttaac tgtttcatcc gctgtgcctg tgagagtgag
11881 ggcgttgctt cttcgctctc catcgtggca aagaggtttt cctgctcgtc tttcttcaag
11941 aaggacagtt ccaccgcagg cgtgagggcg attttcccct cgtccaccat ctgcaaaatc
12001 ggcggtatca gttccgtcag gcggataaaa cgctgcacgg tcatcctgcc cacgccgaag
12061 ccctgtgcca ccttgtcgtc cgtccgcaac ttcgtcacaa cttgtgacga ggttaagtct
12121 gtgcggaaac cctgccgctt catggcttcg gatttcatct tgtaagcaaa cgcccgctcc
12181 gatggcagga tattctcacg ctgcaaattg ctgtctacaa gggtgatgat ggctcggtca
12241 cggtctaagg gcaggacaaa cgcaggcacg gtatttatcc ctgcaagttc agaagcacgg
12301 acacgccgct gtcctgcaat cacttcataa ccgtccccgt cctctttcgg gcgtgtgatt
12361 atcggcgtga caatgccaaa ttccttgatg ctttccgcta actctgacag tgtttcatct
12421 tctgccacat gaaacggatt gtcggggaac gggtacaagt ctttggtctt taacacctta
12481 aaatcctgtt tcttcattca catatctacc tttctttcgc tctgcttcta tgatttttct
12541 gcaattcttc caacacttcg ggcggtattt ttgacagcag ccgccccatc tgctcgttgg
12601 ttctctgcaa ttcaaatatc ttctgattcg ctttctgcac ctttagttcc tgctcgtact

12661 tttcatcacg catacgcccc gcatagtctg attcctgccc aattctctct ttcaaactgt
12721 cgatatacgc ctgctgtttc ccgatttcct tagagaattt ctccacgtct ggcagccatg
12781 cagagagtaa atctaacgct ttatcccttt tcttccctgc gttaaaggcg ttgatgtcgg
12841 atagggcaga cacgatttct tcatactgtt tatcaagcct gccgcctaat ttatagagcc
12901 atgtggggac gtgtttccgc ttggtttcca ttgaggactg cccccgttca agctgattcc
12961 accgtgagga catccgctca tggtaggcgg tctgccactc ggataatgat ttctggttgc
13021 ctaagatagc tttcgctgac agcttattgt ctggcgtaat cggcacaaag cagaggtgca
13081 tatggggcgt tctctcgtcc atatggacga cagcggagag gatattctgc tttccaacac
13141 gctccgaaat gaagtcaaga gccgtctgga aatacgcttt ttgttcttcg ggcggtaact
13201 ggttcataaa ttctggtgaa gctgtgatga gcgtttccac catcatcacg ctgtctttcc
13261 ttgtcctgca ccccgcttcg gctaccatgc ggttaatctc tttcttgtag gtgtactttg
13321 gtggtgctat gagatggtaa ttgtttttag agcgttccat atctatatct gggttgcttt
13381 tgtaggcttc tttcttccgc tcgttgtggc attcacaagc cgcaacgccg cccgcttttc
13441 gtttctggaa acgcaggatt gcataaggca taggcggatt cctcctttct ttcggcgggt
13501 gaccgtcctt tggggtgaca gcggtgacgg tggtgacagc agttttggga taccccctgc
13561 cgactgccgc aactgtcacc ctcacccccct gcatgacggt catgtcgatg atgacggtgt
13621 ttttgggata cccccctcgc aagagccgtc accgtcatgc cgccattctt ttatccgaag
13681 ccgtaaggcg taggatagca gggcacagcc ctgccttaag ggagtccaga gggaacgtct
13741 ggcacacgac tttgcagggc aaagtgtagt gtgttacacc ctgtaaacgc agtcggaaaa
13801 atcggaatga tttttctgac cgcaggggtg gttttacacg accgaaaagg gcgagtaaat
13861 ctcacgcctg cattttgcgt ttacggggtg ttctcccgta gggatgacag cggcagggta
13921 tcctaaaatc actgtcacca ccgtcactgc tgtcacccgc agggcagagc cgccagcttt
13981 acatggcttt aagcgtcaat gacagtaaca ggggggtatc ctaatatcgc tgtcaccacc
14041 atcaccgctg tcacccgccc tccttgcaag ggcaatccgc ctgccgtctt tcctgcggct
14101 gtactggtag cagatacggt tctcgcttaa aaatgtggtg cggtattcat tcagccattt
14161 cgtaatcacc gtgggtatgg tttccgtttc ccccatagcg gctaacagtt ccgttgccgt
14221 gcctatccat tcttccttat ccctcataaa atccaccaac cgaaaaagga catctggtat
14281 cgtttctttc gcaagctgct cctgcgtttt ccgctccaca agctcccaac ggcaatcacg
14341 gaaacgcagc gtgtattcct gataaggcgt gtccctgccc gtcacataca gcttggcggt
14401 gtcagacgca cgtttctcct tttccagaac aaaggtagcg tccgcactcc ccgttaatcc
14461 tgtcgtccca gacaccttgt tgaacacgtc gctgtcattc tgctttcgga tgtggtgtac
14521 gacaatgacc gccagagagt gcctgtcggc aaagtctttg atgagggaga tgtccccata
14581 gtcgcttgca taggcattgt cttttgaagc tgtacggact ttctgcaagg tatcaatgac
14641 aatgagcctg ctgtctgggt aatctttcag ataatcttca agctgcacga taagaccgtc
14701 tgacagcttg cagcttgcca cggcaaagtg gagccgcccg cttgcttcgt ccgtcaaacg
14761 aaataacctg tcctgtatgc ggcagaacgt gtcctcaagg cagaggtaaa gcacatcgcc
14821 ctccattgtc ggcatatccc ataaagggat tccctgcgac acgcataagc atagcttcag
14881 catgagccag cttttgccta tcttctgtga gccgcagaac agcgataagc ctgtcgggat
14941 aaggctgtcc accacaaagg atggtttctc aagcggttca taaaggagcg tttcggcgtt
15001 gactgtctgt aacttctgca tggctttcct cctttcgggc ggtgtctttg ttttcgttgc
15061 acataggcgt tgacctcctt aaaaatagat ttactcccac ggaaaaaagt gagagtatgt
15121 aatgccgcca atcccacaca aataaaaaac agatttcttt ttcgggcggt gtcggtcacg
15181 gttggagata tttcttcaat ttccgccttt actttctcct ttgcaatcgc aacagatttc
15241 tgtattgccg aagcggtgca atgctccata gcggcgattt ggtaaaaatt gaactcatac

15301 tcgtagtaga gcaggaaacg ccgcctttgt atctctggaa ggctcccaac cgccttatag
15361 agcgtttcat tccgttcttc ctcaaccatg cgttcatcaa ggctcttagg cacacgcaac
15421 gcccgtctgt aaagggtttc gtcccatacc tcgttaaact ccctgtgccg ctcgtcccat
15481 tagaaaagat tcctgttcct gcgctccatc tgccgaaact ccataaagaa ctgttccgac
15541 acttccaact cgtgggattt gccctgcccg tccttaaagc tgataaaata ccttgtgccg
15601 ctttccgtgg attcctcccg aagcgtgtat gccttaaccc tgtatgccat cctgttgtcc
15661 tcctgcaaaa aatgagtgag gggatttcca tccctcaccc agtagcccgc aggatggcag
15721 gctgttttag aagctataaa attttccgca gcttcttccg cagatggtct aacctcgcat
15781 agatggcacc agtcgtcaaa tgcacaagcg gggcaatctc ctttgtggaa tacccctgca
15841 ttttcagcag gacgattttc aaggtacgcc cgtccaccgt gactaatact tgatagagat
15901 tttcgctctc aatctcttcc agtaactccg caaccgtacc cacttccgcc tgccgctccc
15961 tgcctgccat gtcctcaaga tattccgcaa cgtcattcgt ccatcggtaa aaccgcctgt
16021 tggaattgaa gtctgccctg tccgccatgc gtatctgctc aatggtcgct tcatcaacgc
16081 cgcactcacg cagcagcttt tcctccgctt ctttccagat acgccatttc ctgtcctccc
16141 gtccgtggtt gtatgccatt cttacttcct ccaatcagaa ttgattgaga gggcagagaa
16201 aagccccctc atcttcccaa aggaaaaaac aagggggctg aacgccttaa attttaattt
16261 tctattatct ttcttagttt tattaggatt ctggctttgc gtttgtttac aacgctctgg
16321 gttatgccca acctcgcccc gacttcacgc tcggaaagtc cgtcaaaaaa gattgcctgt
16381 atcagctcct gttcactatc cgacagcaaa ggcagggcgg ctttcagcct gtccaccatg
16441 accgcattga caacggtttc tgcaatgtcc accgcttcat cagtgataaa gtccagagga
16501 ttcccctcgc tgtccgtaaa tccgtcgaga gatagcagtc tattctttgt atctaatttt
16561 tgcagataac gccaccgttc cttgtcacgg tagaagtctg tgtattgctc cctcacgact
16621 tcaagcagac agccttgaat ggggataaac agcttgtcca tataggtctg gtcggattcc
16681 ctgcaacggc agaactccgt gtaggataat tccacatagc cgccactttc tctgatatat
16741 acctttcttg gtgcatattt caccataaat acctcccatc tgaatttttg aaatgtaaaa
16801 aatccagatg gagaggcgga gaacgacacc gcatatcaga aacagcccta cggcactttc
16861 caacaaaaat cgacaaaaga aaaaccgcaa aggctctgtg acctttacgg ttataggaaa
16921 tagtaattct attgtatgga gattgtactt ctactttcaa ggtgagaagt accgttgcac
16981 tggcagaaat ttttttaact ggtttcctgc cgttctctga tatgctatgt attgataaat
17041 tttgcttcgt atgagcagat agataactgc ccgaaaaaag gacataaaaa aatccctcca
17101 aatttaaaaa caaattcaga gggtaattta gggtataaca aaataaccca cccgaatatc
17161 gttttttttta tttggtgagt ttgttctttc aaatacgaaa caaaaagagc cgatgattcg
17221 tgaattgttc cacaatttca tcggctctgc gtcttaagcg tctggctctt tgatgacagt
17281 tatcttcaag ttgtcaactt taatcaatct ttcttgtctg cattttggac aataaagggg
17341 ataattctcc aaaacagtgt ccttcctaat tttattacgg gttttgctcc cacaaacagg
17401 acacaatatc cattcgcatt tcatcataat tagtctctaa tcctttcaaa tctcatttta
17461 tatgactttt gcaagctgtt aagctaactt gtggaacata tgccgaacct tatctatacg
17521 gctattcggg cggcggggtt ggcaaataaa tttaccaata gctggctggt atccttttaa
17581 ctctgtcaag cagactccct gcccatttgt gaaataagtt aaatcgttcc tgtattcttg
17641 aatacatcta gcagggattt ctcctttcag aatgacctcg tcattcttta tctgagtact
17701 tacaatatct gcacaatacc ttggagcatc atgatacgcc cgtgagagat attcctgcgg
17761 tgcataaatt tcaaagtgga gatatggctc taatagttct gtccctgctt tttttaaagc
17821 ctgctccaat acgatagggg aaagcagccg aaagtctgcg ggggtactta caggactata
17881 atacaatcca tattcaaaac agattttaca gtctgtcact ttccatccat acagcccctg

17941 ctcgcagcca taaagaaccc cctccataac cgcattttgg aacgattgat ttaaatatcc
18001 aagtgaaact ctgctttcat actgcactcc gcttccaata gggagcggct ctatggacaa
18061 cccgacagaa gcccagaaag gatttggcgg gacttctatg tggatggtat attctgcttt
18121 tctaagcggt ctttccatat atataacagt aggctctttt atttctgcct ccacatgata
18181 tttttcctca aggatggcac aaatgacttc catctgcaca ttccccaaaa aagaaagtat
18241 aatctcatgc gttgtagtat ccacataata ttttaaaaga gggtcgccat ctgaaatttc
18301 tgtaagtgcc ccaagcaata tttcccgctg ttcagatttc tttactgcaa tcgttgtttg
18361 gagcataggg agaggatttt caataaattt tctctgcggc aacagtattt cgttccccaa
18421 aatactgttt agctgcaaaa catcatttgg taaaattaca atatcaccag agcaggctgt
18481 atcggatgaa tataattcac cgtttgtcgg aacacacatc tctgtgattt ttattttctc
18541 tttttcagat attctaataa catccctcaa atgcaatgtt ccgctatata tacgcacata
18601 aacaaaacgc cgccttttct ctgaatattc aatcttaaaa acctgcccgc atagttcaga
18661 ttgaccttca ggcgttgatg aataaaattt actggcaatt acttctataa gctgccgaat
18721 ccccagattg tttttagcgc ttccgtgata aacgggaaat aacgttccgt tttggaatct
18781 cctgttttct tcctgttcca gttctgacat tttaaacggt ttccctgaca tatatttctc
18841 taatagttca tcgtttccca taattaccgc atcccactgt tccatatcgt cattgtccgt
18901 tacatttata tggggatgct gcccaacctt ttgcttcact ataatttccg aagaaagctt
18961 tgctttcatt tcccgatata ccattggcaa atcaatcccc tcttggtcaa ttttattgat
19021 gaaaaaaatt gtcggaatct tcattatctg tagtgcatga aacagtatac gggtctgtgc
19081 ctgtatgcca tcctttgcag aaactaataa tactgctccg tctaatacgg ataaagaacg
19141 gtatacttcc gccaaaaaat ccatatggcc tggcgtatct ataatgttga cttttacatc
19201 ctcccactga aaagatgtca ctgctgtctg gatagtgatt cccctttgac gctccaaatt
19261 cattgtatct gtccttgttg tgccttcatc tacgctccct agttctgcaa ttgcaccact
19321 ggtatacaat aaactttccg ttaatgttgt ctttcctgcg tcaacgtgag ccagaatgcc
19381 taagttaatt attttcatgt gattttcctc ctatcaacac ccaaaaaagg gcataaaaat
19441 acccagtgat aaatactcct atcactgggt aaataactcc aatagcccca aaacacttat
19501 atgttttcgg gcatataaaa ttacatgata aaagtattct taaactgggt acaaaaaact
19561 aagccccata ttaaaagtga aacgggactg ctacttttg ttcccactat caaattgaca
19621 gtttatttaa gaataccttg ccgcatattt attaactcct tgtataatac tgaatctaat
19681 tatattcctt aaccctttat ttgtcaagct gacaaactaa agcagaaaaa gcggcaggat
19741 ttccccctgc cactaatcat ctgtttatgc aaaaataatt tccttttcca caatctccct
19801 cgcacaagcc cttatgttat tgaggcatcc tgtccattct aaggcgtttt ctgcctttag
19861 ctgttccgtt atgccctgtg cctgtttcat accctctatg agcctttcaa agcgttcctg
19921 tgcctgtctg ttgatgtcgg caaggtaggc gtttagcctg ccgcttgtaa gaagattggt
19981 gtatgtaact ttacggtact gttttagata atctaaatgc cgttgccccc agatgcctat
20041 tgcctgttct tcttcggcgg gtacagttaa gcacggtatc aaataatccc cttgcctttc
20101 gtatttgccg cccagttcct caaataatga ttttgccatt gtctgttacc tccacattct
20161 tttttatttt gaatgtccgc aaaatccgtc ctacatcagt cgtacattta tatatttctc
20221 cttttctaat attatattca tacgcatagc tagccggtat cgcgatatac gattccttct
20281 ttcgaatctt tcttgatgtt gtctttcgtc tctctcccca atgaatgtgt gcatctttta
20341 tctgaacata atactttcct aataacatta ttttctccat tctctaaaaa tcatttatat
20401 ttactcctct aaaacatcgt aatatactgt gttttctggc aacttaatat taggtatcca
20461 aagaggtcta ccacccccagc ccttacttcc agttacctga tacattgtca atactatctt
20521 ggaaggaaac tcacttccca atctccaatc atcaggagaa agcagtgcac ccgttccttg

20581 cgcaacatct ctgtttctgc ggacgattag tattccttgt cctgctggtt gttcagataa
20641 tattgtatca ataactgaaa caaatgcttg taactcaaaa tctggacttg gaataatatg
20701 ccctaatatt tcttttacaa ttctaagact aacttgataa tataactctt tctcatcaaa
20761 tggttcaagt aattttgaaa tattttcgat tgagtcattg tctggataat atggataata
20821 atttgttcca cctgaaagca tctctacatg atttctatca agcacatttt ttctcgttgg
20881 attcaaacca tttggataat atatttttat atcctcaatt cctctttcga cctgagcaat
20941 aatagaatta ttagttgcat ttatatccgc aaacagttta tacagatttt catcaatgta
21001 caccttcatc atacccggat cacgatcata tccaaacata cggctatgtt gccacattgt
21061 gtctgcttgt ggttttttac tcgttcttgt gtaataaata gtttgtagac caggaaatgt
21121 gactcctctc cccaatgtat ttccaccaat tacaaaatta catcctgatt cataataaga
21181 actctccaca tcattttttc cattcataac gaaaactttg aacatacttc cagttaataa
21241 ttctttaacc ctatcataca aatcttctaa agaacatttt tgagatcttg acggattcat
21301 atctacatac tgtctgctca attcattctt taactctaca tcgagatttt caacacacca
21361 ttttagttct ctttgtatat ccccttcaat ctttttatgt acagccactc ttacacttgg
21421 atgaaataaa caattagata ctttcccacc tgacaaaaga atctgagcag ataccaacag
21481 atgcctgaca acaacttccc tcatgggcat tttcagtttt cccaaaaatg acacacaatc
21541 cggctttcca cttgaaggaa agaaaaaatc tcctccgaga tacgaatccc ccggctgaaa
21601 ataatatgta aaataaggat gccacccaga cgcaagtgtc tgtaaaaaga ttgcctgcgg
21661 cgttcctgtc acctgcaagt ataaactgct tgatgctcca ttttttatgc tgtccaaata
21721 tttattgatg gaagactggc gattcctgtt gactaatgta tttaatgaag ccgcatctgc
21781 ctcatcatca atgatgaaca acggatttcc tttcataaag ccagttgaat taagaatatt
21841 cgcccacaat ttcagtatcc gtacgttctt cttcaataca ataattactg gatccattag
21901 actattttcg gtaaaaagtg ccgcatcatt ttctccacaa atacagaatc cttcaagatc
21961 tgcttttact cggtctagag tctgttgctg aagaacaacg ttatccgtag tcagtaaaac
22021 gaataccgga aatcctaaat ctgtagcctt acatatgatt ccaaacatct ggcccgtttt
22081 tccagactga acgtttccca gcaagaggcc aatttcatga ctggaaaaag aaaatgtttt
22141 tatatgctta tctccaacat catttgccgt tttttcaata gattctgcaa gttttttatt
22201 tcctctatca gttatcttct taagatatgt tttcaaatac cgcatgtact tttctcctat
22261 tcctctttgg atccagttgt aaatgaaagc atccagacat ccaacggatt tccatcttca
22321 tctaatgctg tctgacctgt cttcttaaat tccagtctgt cacatccata ttcctgtagc
22381 atttcttctg taatcattcc cagtcggtcc gtatcagttg ccgtattatt taccggtttt
22441 acaatcccag ctgccgccag ccttcctttc agccatcttc ccattatcaa ttcatcgcca
22501 actgcactaa actgtttatt attatcactt gttgtatgtg ctttaaacca atacccatca
22561 tctgtcacta caaaaaaacgg tttattcttc tctggataac cttccatacg gtacacatcc
22621 gcactaactg tcaactgcgt ttcataccag tctcttgcct ttttcttact tctcggtgcc
22681 gcatagcaga cattaatatt tgactttgta taatgctttc catcatccaa atgacgttcg
22741 tctttcttag gaactttcag tttcagaaag aatgaaacat acgctttaca cctttcataa
22801 aactcaacgt ttgattttgg tacagatgta accagctcta ttccatttaa tgcagtatta
22861 acttccctca caagaggcat attctttaca agtgcaatat tctctgaaca gtttggcatt
22921 ttcaacttct caatatgttc tgcaatctca gccacttcct gttcatttgt tgtaatagcg
22981 gacatttcat actgtcttct attgcctgct tccaatttaa taactccaag atttgccgat
23041 ccaataattg ctgaaaatgg tttctcgtct ttgtaaaaac aaaataattt cccatgatat
23101 ttaaagctct ttacaattcg gatctctcca gtatttgatt cctgccactt ttcattaatc
23161 tttaatgccg tatgatacgc cgcttccggc ataccttcaa taaagtacat tccaatattt

23221 aaacatactt ttgaaatatt atcttcattc acaatcctat ccaattcaac aagtgatgca
23281 ttagaaatat atccaactgc aatttcgact ctatctgcta tttttatctg ctcattaaag
23341 cagtctatta ttgtttcctg attatcctct gttcctaatg gcagaatatt tgagtataat
23401 aatttcatac tgcacccct ataatacttt agtgttttta ttcaatttta ctacaaaaca
23461 actaataatg ctaggaaaga aataccattc tcaatcacgt gacatcaaat tcaatttcct
23521 ccaaatacaa gtattgtgaa acaaaaatac ttatactata attgacttat gtcaaagcag
23581 gggaatacta atatcagaat acggttttac ccaaaagaat tgatctgtgt tcagagaaaa
23641 aatacatgat ggcaagaatc ctatatggat aaattgaata aagaatatat tgagttactg
23701 agtagtgaag gaaaaccttc tgaaaaattc tgagctttag aagagcgaat tcgcaacgat
23761 aaaaaaaata caggtgtaca gttaaaaatg agtcgttcta attgcattcc taacattatt
23821 tcacttttaa atgaaggagc aatcacgatg aaagatttag aagaatttag cgatgaattg
23881 aaagaaacaa tacatttcat cacaagataa aattcctatc tattcagaac agagcattca
23941 caatttatag ctaaagagat tatttggcag gcgtcgtagt tcttatgtgt tttcgataat
24001 aaaattgatg tataatattt atagattgta acaaaattaa ggagggagcg aatgaattta
24061 gaaatatatc aaaaattgtg ttcagaatca aaaattctct ggacacagca ttgtctgcaa
24121 cgaatgcagg aaagagatat cagtcgtgct gatgtaaaaa atggaattgc aaccggtgaa
24181 attatagaag actaccctga cgattatcct aatccaagtt gtctgatttt tggatataac
24241 gtaaatggac gaattttgca tattgttgcc gggtgtgata atataaatat atatattata
24301 acggcatatt acccagatac aaagaaattt gaagatgatt taaagactcg aagaaagagg
24361 tgatggatat gtgtatgtat tgtaaaaata gtacaactgt caatagcaca acaacacacg
24421 tcgtaaatta taaaaattgt attattgtaa ttaaaaatgt tccatgcctt gaatgtgatc
24481 agtgtggaga aaaatattat acagacgaag ttgctgaacg ccttgaattg attgttgata
24541 tggcgaaaaa attaatgcag gaaattgcag ttatcgatta tccacaggtg gcttaataat
24601 tttggcggaa atatcatttt cgatatttcc gccacttctt tttatatact cttcgcacta
24661 tttattctgt ccacccagtt acataaaatt acaatactct ctaaatttac attaata

//

DEFINITION     Integrative and mobilisable element IME_9

CDS          complement(1..62)
    /ID=F66_02590
    /color=1
    /locus_tag="F66_02590"
    /note="hypothetical protein"
    /product="hypothetical protein"
    /protein_id="gnl|SC|F66_02590"
    /codon_start=1

CDS          207..995
    /color=8
    /locus_tag="F66_02591"
    /note="Heat shock protein  META domain-containing protein, protein motif:CLUSTERS:PRK10449 protein motif:Pfam:PF03724.10"
    /product="Heat shock protein  META domain-containing protein"
    /"ID=F66_02591

/protein_id="gnl|SC|F66_02591"""
/codon_start=1
CDS 1323..3236
/color=8
/locus_tag="F66_02592"
/note="Molecular chaperone DnaK, similar to AA sequence:UniProtKB:P0A6Y8 protein motif:CLUSTERS:PRK00290 protein motif:Cdd:COG4820 protein motif:TIGRFAMs:TIGR02350 protein motif:Pfam:PF00012.14"
/product="Molecular chaperone DnaK [chaperone]"
/"ID=F66_02592
/protein_id="gnl|SC|F66_02592"""
/gene="dnaK"
/codon_start=1
CDS 3438..4004
/color=9
/locus_tag="F66_02593"
/note="Mobilizable transposon protein TnpA , protein motif:TIGRFAMs:TIGR01764 protein motif:Pfam:PF12728.1"
/product="Mobilizable transposon protein TnpA [DDE recombinase]"
/"ID=F66_02593
/gene="tnpA"""
/protein_id="gnl|SC|F66_02593"
/codon_start=1
CDS 4001..5248
/color=9
/locus_tag="F66_02594"
/note="Tyrosine recombinase, similar to AA sequence:UniProtKB:P0A0P0 protein motif:CLUSTERS:PRK00236 protein motif:Cdd:COG4974 protein motif:TIGRFAMs:TIGR02224 protein motif:Pfam:PF00589.16"
/product="Tyrosine recombinase [recombinase]"
/"ID=F66_02594
/protein_id="gnl|SC|F66_02594"""
/codon_start=1
CDS 5572..6213
/ID=F66_02595
/color=1
/locus_tag="F66_02595"
/note="hypothetical protein"
/product="hypothetical protein"
/protein_id="gnl|SC|F66_02595"
/codon_start=1

CDS 6347..6715
/color=9
/locus_tag="F66_02596"
/note="DNA binding domain  excisionase family
Helix-turn-helix domain , protein motif:TIGRFAMs:TIGR01764
protein motif:Pfam:PF12728.1"
/product="DNA binding domain  excisionase family
Helix-turn-helix domain [excisionase]"
/"ID=F66_02596
/protein_id="gnl|SC|F66_02596"""
/codon_start=1
CDS 6759..7100
/color=9
/locus_tag="F66_02597"
/note="DNA binding domain  excisionase family
Helix-turn-helix domain, protein motif:TIGRFAMs:TIGR01764
protein motif:Pfam:PF12728.1"
/product="DNA binding domain  excisionase family
Helix-turn-helix domain [excisionase]"
/"ID=F66_02597
/protein_id="gnl|SC|F66_02597"""
/codon_start=1
CDS 7103..8209
/ID=F66_02598
/color=9
/locus_tag="F66_02598"
/note="AAA family ATPase  Sms DNA repair protein"
/product="AAA family ATPase  Sms DNA repair protein"
/protein_id="gnl|SC|F66_02598"
/codon_start=1
CDS 8410..9309
/color=10
/locus_tag="F66_02599"
/note="DNA primase, protein motif:CLUSTERS:PRK05667
protein motif:Cdd:COG0358 protein motif:TIGRFAMs:TIGR01391
protein motif:Pfam:PF01807.14"
/product="DNA primase [replication]"
/"ID=F66_02599
/protein_id="gnl|SC|F66_02599"""
/codon_start=1
CDS 9466..10437
/ID=F66_02600
/color=7
/locus_tag="F66_02600"

/note="MarR family transcriptional regulator  winged
helix-turn-helix domain-containing protein:Prodigal:2.60"
/product="MarR family transcriptional regulator  winged
helix-turn-helix domain-containing protein
[transcriptional regulator]"
/protein_id="gnl|SC|F66_02600"
/codon_start=1
CDS 10425..11204
/ID=F66_02601
/color=1
/locus_tag="F66_02601"
/note="hypothetical protein"
/product="hypothetical protein"
/protein_id="gnl|SC|F66_02601"
/codon_start=1
CDS 11210..11971
/color=8
/locus_tag="F66_02602"
/note="Serine/threonine protein phosphatase, protein
motif:Pfam:PF00149.22"
/product="Serine/threonine protein phosphatase
[phosphatase]"
/"ID=F66_02602
/protein_id="gnl|SC|F66_02602"""
/codon_start=1
CDS 12193..12645
/ID=F66_02603
/color=1
/locus_tag="F66_02603"
/note="hypothetical protein"
/product="hypothetical protein"
/protein_id="gnl|SC|F66_02603"
/codon_start=1
CDS 12642..12974
/ID=F66_02604
/color=4
/locus_tag="F66_02604"
/note="Conjugal transfer protein TraE"
/product="Conjugal transfer protein TraE [mating pair
formation]"
/gene="traE"
/protein_id="gnl|SC|F66_02604"
/codon_start=1
CDS 13033..13398

/color=4
/locus_tag="F66_02605"
/note="Conjugal transfer protein TraE, protein
motif:Cdd:COG3451"
/product="Conjugal transfer protein TraE [mating pair
formation]"
/"ID=F66_02605
/gene="traE"""
/protein_id="gnl|SC|F66_02605"
/codon_start=1
CDS 13447..13746
/ID=F66_02606
/color=1
/locus_tag="F66_02606"
/note="DUF4133 domain-containing protein"
/product="DUF4133 domain-containing protein"
/protein_id="gnl|SC|F66_02606"
/codon_start=1
CDS 13869..16574
/color=4
/locus_tag="F66_02607"
/note="Bacteroides conjugation system ATPase TraG, protein
motif:CLUSTERS:PRK13721 protein motif:Cdd:COG3451 protein
motif:TIGRFAMs:TIGR03783 protein motif:Pfam:PF12846.1"
/product="Bacteroides conjugation system ATPase TraG
[ATPasse mating pair formation]"
/"ID=F66_02607
/protein_id="gnl|SC|F66_02607"""
/gene="traG"
/codon_start=1
CDS 16795..19335
/ID=F66_02608
/color=10
/locus_tag="F66_02608"
/note="Smc-like chromosome partition protein"
/product="Smc-like chromosome partition protein"
/protein_id="gnl|SC|F66_02608"
/codon_start=1
CDS 19350..20075
/ID=F66_02609
/color=1
/locus_tag="F66_02609"
/note="hypothetical protein"
/product="hypothetical protein"

/protein_id="gnl|SC|F66_02609"
/codon_start=1
CDS 20103..20867
/ID=F66_02610
/color=1
/locus_tag="F66_02610"
/note="DUF5045 domain-containing protein"
/product="DUF5045 domain-containing protein"
/protein_id="gnl|SC|F66_02610"
/codon_start=1
CDS 20911..21741
/color=8
/locus_tag="F66_02611"
/note="Phage antirepressor KilAC domain-containing protein, protein motif:Cdd:COG3645 protein motif:Pfam:PF03374.8"
/product="Phage antirepressor KilAC domain-containing protein"
/"ID=F66_02611
/protein_id="gnl|SC|F66_02611""""
/codon_start=1
CDS 21757..22158
/ID=F66_02612
/color=1
/locus_tag="F66_02612"
/note="hypothetical protein"
/product="hypothetical protein"
/protein_id="gnl|SC|F66_02612"
/codon_start=1
CDS 22158..23288
/color=4
/locus_tag="F66_02613"
/note="Bacteroides conjugative transposon TraJ protein, protein motif:TIGRFAMs:TIGR03782"
/product="Bacteroides conjugative transposon TraJ protein [mating pair formation]"
/"ID=F66_02613
/gene="traJ""""
/protein_id="gnl|SC|F66_02613"
/codon_start=1
CDS 23354..23755
/ID=F66_02614
/color=1
/locus_tag="F66_02614"

/note="hypothetical protein"
/product="hypothetical protein"
/protein_id="gnl|SC|F66_02614"
/codon_start=1
CDS 23759..24373
/color=4
/locus_tag="F66_02615"
/note="Bacteroides conjugative transposon TraK protein, protein motif:TIGRFAMs:TIGR03781"
/product="Bacteroides conjugative transposon TraK protein [mating pair formation]"
/"ID=F66_02615
/gene="traK"""
/protein_id="gnl|SC|F66_02615"
/codon_start=1
CDS 24386..24805
/ID=F66_02616
/color=1
/locus_tag="F66_02616"
/note="hypothetical protein"
/product="hypothetical protein"
/protein_id="gnl|SC|F66_02616"
/codon_start=1
CDS 24789..25931
/color=4
/locus_tag="F66_02617"
/note="Bacteroides conjugative transposon TraM protein, protein motif:TIGRFAMs:TIGR03779 protein motif:Pfam:PF12508.2"
/product="Bacteroides conjugative transposon TraM protein [relaxosome]"
/"ID=F66_02617
/gene="traM"""
/protein_id="gnl|SC|F66_02617"
/codon_start=1
CDS 26001..26846
/color=4
/locus_tag="F66_02618"
/note="Bacteroides conjugative transposon TraN protein, protein motif:TIGRFAMs:TIGR03780"
/product="Bacteroides conjugative transposon TraN protein [mating pair formation]"
/"ID=F66_02618
/gene="traN""""

/protein_id="gnl|SC|F66_02618"
/codon_start=1
CDS 26846..27391
/ID=F66_02619
/color=1
/locus_tag="F66_02619"
/note="hypothetical protein"
/product="hypothetical protein"
/protein_id="gnl|SC|F66_02619"
/codon_start=1
CDS 27400..28053
/ID=F66_02620
/color=1
/locus_tag="F66_02620"
/note="DUF1566 domain-containing protein"
/product="DUF1566 domain-containing protein"
/protein_id="gnl|SC|F66_02620"
/codon_start=1
CDS 28120..29028
/ID=F66_02621
/color=8
/locus_tag="F66_02621"
/note="Fimbrillin family protein"
/product="Fimbrillin family protein"
/protein_id="gnl|SC|F66_02621"
/codon_start=1
CDS 29109..31298
/color=4
/locus_tag="F66_02622"
/note="TraM recognition site of TraD and TraG  Type IV
secretion-system VirD4 components , protein
motif:Cdd:COG3505 protein motif:Pfam:PF10412.3"
/product="TraM recognition site of TraD and TraG  Type IV
secretion-system VirD4 components [coupling protein]"
/"ID=F66_02622
/protein_id="gnl|SC|F66_02622"""
/codon_start=1
CDS 31352..31849
/ID=F66_02623
/color=8
/locus_tag="F66_02623"
/note="lysozyme  Glycoside hydrolase family protein"
/product="lysozyme  Glycoside hydrolase family protein
[hydrolase]"

/protein_id="gnl|SC|F66_02623"
/codon_start=1
CDS 31938..33146
/color=9
/locus_tag="F66_02624"
/note="Tyrosine recombinase, protein motif:CLUSTERS:PRK00283 protein motif:Cdd:COG4974 protein motif:TIGRFAMs:TIGR02225 protein motif:Pfam:PF00589.16"
/product="Tyrosine recombinase [recombinase]"
/"ID=F66_02624
/protein_id="gnl|SC|F66_02624"""
/codon_start=1
CDS complement(33396..34607)
/color=9
/locus_tag="F66_02625"
/note="IS1380-like element IS612 family transposase, protein motif:Pfam:PF01609.15"
/product="IS1380-like element IS612 family transposase [DDE recombinase]"
/"ID=F66_02625
/gene="tnpA"""
/protein_id="gnl|SC|F66_02625"
/codon_start=1
CDS 34872..36089
/color=2
/locus_tag="F66_02626"
/note="macrolide efflux MFS transporter Mef A 29  Major Facilitator Superfamily, protein motif:CLUSTERS:PRK10489 protein motif:TIGRFAMs:TIGR00900 protein motif:Pfam:PF07690.10"
/product="macrolide efflux MFS transporter Mef%28A%29 Major Facilitator Superfamily [antibiotic resistance]"
/"ID=F66_02626
/gene="mefA"""
/protein_id="gnl|SC|F66_02626"
/codon_start=1
CDS 36109..36963
/color=8
/locus_tag="F66_02627"
/note="Nucleotidyltransferase domain-containing protein, protein motif:Pfam:PF01909.17"
/product="Nucleotidyltransferase domain-containing protein [nucleotidyltransferase]"
/"ID=F66_02627

/protein_id="gnl|SC|F66_02627"""
/codon_start=1
CDS 36960..37694
/color=8
/locus_tag="F66_02628"
/eC_number=2.1.1.-
/note="Class I SAM-dependent methyltransferase ycgJ ,
similar to AA sequence:UniProtKB:O31474 protein
motif:CLUSTERS:PRK08317 protein motif:Cdd:COG4976 protein
motif:TIGRFAMs:TIGR02072 protein motif:Pfam:PF08241.6"
/product="Class I SAM-dependent methyltransferase ycgJ
[methyltransferase]"
/"ID=F66_02628
/protein_id="gnl|SC|F66_02628"""
/gene="ycgJ"
/codon_start=1
CDS 37727..38635
/color=2
/locus_tag="F66_02629"
/eC_number=2.7.7.-
/note="Aminoglycoside 6-adenylyltransferase Streptomycin
adenylyltransferase, similar to AA
sequence:UniProtKB:P17585 protein motif:Pfam:PF04439.6"
/product="Aminoglycoside 6-adenylyltransferase
Streptomycin adenylyltransferase [antibiotic resistance]"
/"ID=F66_02629
/protein_id="gnl|SC|F66_02629"""
/gene="aadK"
/codon_start=1
CDS 38644..39174
/color=2
/locus_tag="F66_02630"
/note="Streptothricin N-acetyltransferase Sat4
Acetyltransferase GNAT family , protein
motif:CLUSTERS:PRK10975 protein motif:TIGRFAMs:TIGR01575
protein motif:Pfam:PF00583.18"
/product="Streptothricin N-acetyltransferase Sat4
Acetyltransferase %28GNAT%29 family [antibiotic
resistance]"
/"ID=F66_02630
/gene="sat4"""
/protein_id="gnl|SC|F66_02630"
/codon_start=1
CDS 39335..40342

/color=9
/locus_tag="F66_02631"
/note="IS5 family transposase, protein
motif:Pfam:PF01609.15"
/product="IS5 family transposase  [DDE recombinase]"
/"ID=F66_02631
/protein_id="gnl|SC|F66_02631"""
/codon_start=1
CDS 40535..41320
/color=2
/locus_tag="F66_02632"
/eC_number=2.7.1.95
/note="Aminoglycoside 3'-phosphotransferase, similar to AA
sequence:UniProtKB:P0A3Y5 protein motif:Cdd:COG3231
protein motif:Pfam:PF01636.17"
/product="Aminoglycoside 3'-phosphotransferase [antibiotic
resistance]"
/"ID=F66_02632
/protein_id="gnl|SC|F66_02632"""
/gene="aphA"
/codon_start=1
CDS 41596..42168
/color=7
/locus_tag="F66_02633"
/note="DeoR-like helix-turn-helix domain protein, protein
motif:CLUSTERS:PRK11886 protein motif:Pfam:PF08279.6"
/product="DeoR-like helix-turn-helix domain protein
[transcriptional regulator]"
/"ID=F66_02633
/protein_id="gnl|SC|F66_02633"""
/codon_start=1
CDS 42273..42548
/ID=F66_02634
/color=1
/locus_tag="F66_02634"
/note="WYL domain-containing protein"
/product="WYL domain-containing protein"
/protein_id="gnl|SC|F66_02634"
/codon_start=1
CDS 42677..43318
/color=8
/locus_tag="F66_02635"
/note="5-amino-6-%285-phosphoribosylamino%29uracil
reductase, protein motif:CLUSTERS:PRK05625 protein

motif:Cdd:COG0117 protein motif:TIGRFAMs:TIGR00227 protein
motif:Pfam:PF01872.11"
/product="5-amino-6-%285-phosphoribosylamino%29uracil
reductase  RibD family protein [reductase]"
/"ID=F66_02635
/gene="ribD"""
/protein_id="gnl|SC|F66_02635"
/codon_start=1
CDS 44164..45477
/color=9
/locus_tag="F66_02636"
/note="IS21 family transposase, protein motif:Cdd:COG4584
protein motif:Pfam:PF00665.20"
/product="IS21 family transposase [DDE transposase]"
/"ID=F66_02636
/protein_id="gnl|SC|F66_02636"""
/codon_start=1
CDS 45440..45802
/color=10
/locus_tag="F66_02637"
/note="DNA replication protein DnaC, protein
motif:CLUSTERS:PRK09183 protein motif:Cdd:COG1484 protein
motif:Pfam:PF01695.11"
/product="DNA replication protein DnaC [replication]"
/"ID=F66_02637
/protein_id="gnl|SC|F66_02637"""
/codon_start=1
CDS 45854..46189
/color=10
/locus_tag="F66_02638"
/note="DNA replication protein DnaC , protein
motif:CLUSTERS:PRK09183 protein motif:Cdd:COG1484 protein
motif:Pfam:PF01695.11"
/product="DNA replication protein DnaC [replication]"
/"ID=F66_02638
/protein_id="gnl|SC|F66_02638"""
/codon_start=1
CDS 46259..46552
/color=9
/locus_tag="F66_02639"
/note="IS3/IS911 family transposase, protein
motif:Pfam:PF01527.14"
/product="IS3/IS911 family transposase [DDE recombinase]"
/"ID=F66_02639

/protein_id="gnl|SC|F66_02639"""
/codon_start=1
CDS 46813..47427
/color=9
/locus_tag="F66_02640"
/note="IS3 family transposase, protein
motif:Pfam:PF00665.20"
/product="IS3 family transposase [DDE recombinase]"
/"ID=F66_02640
/protein_id="gnl|SC|F66_02640"""
/codon_start=1
CDS 47564..48235
/ID=F66_02641
/color=8
/locus_tag="F66_02641"
/note="Nucleotidyltransferase domain-containing protein"
/product="Nucleotidyltransferase domain-containing protein
[nucleotidyltransferase]"
/protein_id="gnl|SC|F66_02641"
/codon_start=1
CDS 48232..48972
/color=8
/locus_tag="F66_02642"
/eC_number=2.1.1.-
/note="Class I SAM-dependent methyltransferase ycgJ,
similar to AA sequence:UniProtKB:O31474 protein
motif:CLUSTERS:PRK08317 protein motif:Cdd:COG4106 protein
motif:TIGRFAMs:TIGR02072 protein motif:Pfam:PF08241.6"
/product="Class I SAM-dependent methyltransferase ycgJ
[methyltransferase]"
/"ID=F66_02642
/protein_id="gnl|SC|F66_02642"""
/gene="ycgJ"
/codon_start=1
CDS 49010..49481
/color=8
/locus_tag="F66_02643"
/note="HD domain-containing protein  phosphohydrolase
[hydrolase]"
/product="HD domain-containing protein  phosphohydrolase
[hydrolase]"
/"ID=F66_02643
/protein_id="gnl|SC|F66, protein motif:Cdd:COG1418 protein
motif:Pfam:PF01966.16"

/protein_id="gnl|SC|F66_02643"""
/codon_start=1
BASE COUNT    15339 a  10040 c  11861 g  12241 t
ORIGIN
1 tgagctgtga taggaggcgt aacacagcag aatagaatca aagacaggac aattattctc
61 atatttgttt actttaggtg aactatgtaa attgttcgtt tgttgtatat ttattggcaa
121 atgtatatat tttatgttgc attggatatt atttcccaat cttttgatat ctttgccccc
181 gattttaact taaacgttat aacaatatga agaaagtatt tgtttcaatt tgcattgctg
241 gtgcagcact tgcaatgtct tcgtgccgtt cggttgagaa ggcaattccc ctgtcatcta
301 tcaatggtga atggaatatc atagaagtca atggctcgaa aatcactccg ggcgaaagta
361 aatcccttcc ttttatcgct tttgatacag caaccggtcg tgtgtccggc aatagcggct
421 gtaatcggat gatgggtagc ttcgatgtaa atgcgaagcc gggaagtctt gaacttacag
481 gaatggctag tacccggatg atgtgtccgg atatgactac tgaaaataat gtattgaatg
541 cttttgcgca agtaaaagga tataagaagg caggtaaaga caaaatgtat ctttgcaatt
601 cgtccaaccg cccggttgtc gttctacaaa agaaggaggc tgatgtgaaa ctttcggttt
661 tgaatggaga atggaaaatt aaagaagtga atggtgaggc aattccttcc ggaatggaaa
721 aacaaccttt cattgctttt gatgtgaaga aaaagacaat tcatggcaat gccggatgta
781 atctgattaa tggcggtttt gaaaccagta ccagtaatgc caaatctatt tctttcccgg
841 gagtagcgag cacaatgatg gcttgtccgg acatggaaac agaaggtaag gtgctgaaag
901 ctatcaatga ggtgaagtca ttcgatgtgt tgtccggtgg tggcattggc ttgtatgatg
961 caaatggcgc attggtcatc gtgctggaaa agtaataaat agattatcta tataatagga
1021 tatgcctgtg tcgcacatga tgacacaggc atattttttt tcttttcatc gaaccagagc
1081 cgtctgcatt tgtttaaaga aacagttgct gactggcttt gaatatgact atgtcagcag
1141 tgtctattta ttaaaactgt ctttttggca gatattatgt catgcattcg tataatttcc
1201 ttttggcaca cggtttgctt tttagtaagc gtccgttcgc gacgaggctc gctgataaga
1261 gccggagaaa caaccggaca aaaagaaatc atagaataaa taataaataa aaaataacga
1321 tcatgggaaa aattattggt attgacttag gaactacaaa ctcttgtgtt gccgtatttg
1381 aaggtaacga acctgtagta attgcaaaca gtgaaggtaa acgtacgact ccttctgttg
1441 tcgctttcgt agatggcggt gaacgtaagg taggagatcc tgcaaaacgt caggctatta
1501 cgaatcctac acgtacaatc ttctctatca aacgtttcat gggtgaaact tgggatcagg
1561 tgcaaaaaga ggtgactcgt gttccttata aagtagtgaa aggtgacaat aacactccgc
1621 gtgttgacat tgacggacgt ctgtacactc cgcaggaaat ctctgctatg attcttcaga
1681 agatgaagaa gactgctgaa gactatcttg gacaggaagt gacagaagca gttatcaccg
1741 ttcctgcata tttctccgac tctcaacgtc aggctactaa agaagccgga cagattgccg
1801 gtctggaagt aaaacgtatt gtaaacgaac cgacagccgc tgctcttgct tacggtctgg
1861 ataaggctca caaagatatg aagattgctg tattcgacct tggtggtggt acattcgata
1921 tttctatcct tgaatttggt ggtggtgtgt ttgaagtact ttctacaaat ggtgatactc
1981 acttgggtgg tgatgacttc gaccaggtaa tcatcaactg gttggtacag gaattcaaga
2041 atgacgaagg tgctgacctg actcaggatc cgatggcttt gcaacgtctg aaagaagctg
2101 cagaaaaagc taagattgag ctttcttctt ctacaagtac agaaatcaac ttgccgtata
2161 tcatgccggt aggtggtgtg cctaagcact tggtgaagac tttgactcgt gcaaaattcg
2221 aatctctggc tcacgaattg attcaggctt gtcttgaacc atgtaagaaa gcaatgagcg
2281 atgcaggttt gaacaatgct gatattgatg aagtaatcct tgttggtggt tcttcccgta
2341 ttccggctgt tcagaagttg gtagaagatt tcttcggcaa gactccttct aaaggtgtga

2401 atccggatga agtggtagct attggtgctg ctgtacaggg tgctgttttg acagatgaaa
2461 tcaaaggtgt agtattgttg gatgttactc cgttgtcaat gggtatcgaa acactgggtg
2521 gtgtgatgac taagttgatc gacgctaaca ctaccatccc ggctcgtaag agtgaaacat
2581 tctctactgc tgccgataac cagacggaag ttactatcca cgtgttacag ggagaacgtc
2641 cgatggctgc acagaataaa tcaatcggtc agttcaactt gacaggtatt gctccggctc
2701 gtcgtggtgt tcctcaaatt gaggttacat ttgatatcga tgccaacggt attttgaaag
2761 tatctgcaaa agataaggct accggtaaag aacaggctat ccgtattgaa gcatccagcg
2821 gtttgagcaa ggaagaaatc gaaaagatga aggctgaagc tgaagctaat gctgaagcag
2881 ataagaaaga acgtgaaaag atcgacaagc tgaaccaggc tgacagcgta atcttctcta
2941 ctgaaaatca gttgaaagaa ttaggtgata agttgcctgc tgacaagaaa gctccgattg
3001 aagctgcttt gcagaaattg aaagatgctc acaaggcaca ggatttggct gctattgata
3061 ctgctatggc agaaatcaat actgctttcc aggctgcaag tgctgaaatg tatgcacaga
3121 gcggtgcgca aggtggagca caggctggtc ctgatatgaa tggtggtgct ggccagcaag
3181 ataacagcaa gcacggagat aacgttcagg atgctgactt cgaggaagtg aaataagcac
3241 atagagaaat aagaatacaa tatcaaatgg aaagcgtgta gctcaatgag ttacacgctt
3301 ttcttgtttt tggccgtaca caataatatt gctatctttg cgttattatt ataaacatct
3361 gtaccttatt tgtacctgta agaaaaagta gatatgatga gaagaataga tattaagaac
3421 tacaaaatga atgaattatg gcagtagcaa agtataagat agtacggaaa tgtcctgtgt
3481 gcggagaaga gttcttcgca aggactttgg agtcctggta ttgttcaccc aaatgctcca
3541 aggtggcgtg gaagcgtaag catgatgaag aaaaacgcca acttgaactg gacaagattg
3601 taagcaatat gcccaagtct aaagagtata tcagcataac ggaagcgtat gctatgtttg
3661 gtgcaagccg ctcaaccatt tataggctca tctacatgaa aaagatttcc ttcattgagc
3721 cagagaaagg aatacgactt gtatgcaaag gagaactaat gaatctgttt ccgttaagac
3781 agtctccgct tgacaccaag ccaaggaaac cagttaccat gtaccgcatg gaaccagagg
3841 attgctatac aataggtgag atttccaaga aatttcatct ggatgacagc accgtgtatg
3901 ctcatatacg aaaatactca ataccgaccc ggcaaatcgg taattatgtg tatgctcaca
3961 aggaatcaat agacaaactt tataaagata taaaaccatt atgaaaagat tagattatac
4021 taaagtctcc gtcaaattaa agaaaagtga gtggagagat gagtggttca tctatcttga
4081 agcttatcct gtttatgaga cgggaaatga caaacccaaa cgagtgcgtg aatatctcag
4141 gaggtctatt actaccccta tatgggataa acgacgaagc gagcgagccg ttgctggtag
4201 aataaagtac aagcccaagc gagatgacaa tggcgttatt cagtgtaaat ccaagcaaga
4261 tatggaaact tgcctttatg cggatggcgt tcgagtcctg cgtcaaaaag agtacgacaa
4321 catggctctc tttactgacc agcaaatgga aatggcagaa caaagcgaac gctcaaaatg
4381 caatgttctg gagtatattg agaagttaat caaagaaaga gaggaaactg cgtcagaatc
4441 gatagttgtc aactggcgca gattgcacac actgttgtca atgtttgcca agtgtgacta
4501 catacagttc tcgcagattg acatgaagta tattgaggca ttccgttcct tcttgataaa
4561 ggcaccgcag ggtggttcta agaaaggaac tatttcaagg aatacggcat cgacgtactt
4621 ctctatcttc aaggcggcac tcaaacaagc tttcgtggat ggctatctga attgtgatat
4681 tgctgccaag gctaaaaata tcatgtttca gagtgctcga agagagtatc tgtcactgga
4741 agaacttaat atattggcaa agactccgtg tgatgatata ctaaaacggg cagccttgtt
4801 ctctgcactg acaggaatgc gacatagcga tattcaaaag cttaaatggt cagaggttga
4861 agaatacaat ggtgggtacc gtctcaattt cactcagcag aagactaagg gcgttgaata
4921 catgcctata tcgccacagg catacaagct ttgcggtgag cgcaagaaag acggagaact
4981 gttggtgttt gccggactgc ctgacccttc atggatatct aggcctttgg aacggtgggt

5041 aaaagcatca gggattacca agcacatcac ctttcattgc ttccgccaca cctatgcaac
5101 gttacaattg gctaacggta cagacatata taccgtaagc aaaatgttag ggcataccaa
5161 tgtaaaaacc acgcaaatct acgcaaaagt aattgacaag aagaaagatg aagctacaga
5221 agcttttaag ttggacatag atgaatagtg attatcattt gtaagcactt agtatagaag
5281 accacttgta tcaataagtc ctttattgta caggtggttt ttagtgcctt tcattttcaa
5341 cattctatgt aatgatgcta ttgttttggc actttaacct ttcgtggaaa ccaaacattt
5401 ataacaataa acagataaaa atgaaaaaag acagcctttt ttggcttctc tctttttgaa
5461 gagagcaagc gtgtccacaa acatgtggag tgtacagaag acttctacag agaactgtat
5521 gagttgtttg agatcgcaag aagttggtat cttcaagctg aaaagaatca tatgaagaca
5581 gagtacttca tagaactatt tgagcgcagt caccaatata ttgactgtga ttgtgcacgt
5641 tgtaagaact cgcgacagat ggcaattggc attattggta ctttgtttag agactctaag
5701 tgtgtgtcaa taatcaagaa aaaagaagat tattccaagc aggaacttat tgagttgttt
5761 cttcaacatg tgggtactgg tctgccaatc cttacccgaa agaaatcgtc catcctcact
5821 ttgggatgcc aattgtcaga cagacagatg gatttgttgg tagaacttgt ccaaagtcac
5881 gacatatttg attttgccga caacagtgat gtgcgtagcg agttgtgtcg cctattcaag
5941 tgcgacttgg acgcatctat ccgtgtaaag aatgtacgta atgtagccgt actgtttgac
6001 gctatggctc aataccattt gataaacaat aattggcaat atgtcatggg agaaggtcgc
6061 ttcttaacca gtatcaaaaa ggatggaaca gagaaattca taacttcaag ttgtctttcc
6121 tcttctcttt caaggataag aagaaacgtt tccatgacag caagtcaata tgccatctgc
6181 aaatccatcg aacagatatt aagagaggaa taaatcacaa taattctgct aaatgaaagt
6241 aaaaacgtga tagtttggtt gatactgcct tactatcacg ttactttttc tatttagtga
6301 cagtctcccc ataactttgc atcaaaaatc ataaaaaata attagcatgg aagaaaatag
6361 tataagattt gaagatattc ccaacgctat aacgggggta ctgaaaaagc tgtcatcatt
6421 agaagacaag attgatggta tatatgaact tgttcaatca gaaaaagaag aaacatggtt
6481 tacggttgct gaattatgtg cttatctccc aacacatccg gtagagcata ccatatattg
6541 ctggacaagc aatcgggaaa taccgtttca caagagaggc aaacgcatca tgttcctcaa
6601 atctgagata gatgaatggc ttcaaggaac caagggcaaa tcgaagaacg agattcaaag
6661 agaagcagag gaatatgtac tatctacaca aagaaagaat agacggctaa tataaccgtg
6721 ctactgtctg taaacaattt aattaaggta cgtgacacat gaatgaaaga cattttagac
6781 tctatgagcg aattgtcgca atagaagaca gtttggaggc tttgggacct attgacaaac
6841 taatagagcg aatcgaggaa ttggagaaga tggtcaagca gaccaaaacc gtcttggggt
6901 ttgacgaagc atgtaagtat attggtgttt ctgaaagcct gctatacaaa cttacagccg
6961 caaaagaagt tccacactat aagccaaggg gaaaaatgct gtattttaat agagaagaaa
7021 tcgacaagtg gctgctccaa aacaaacaag aggtcattgg aatggtaacg aaaatagaaa
7081 ttgataatcc gaaagaatga aaatggaaga aaataacaaa catgtacagc ctaactcaaa
7141 agaggaagga gtacagcgtc tcaatcgcat attgagcgaa tcgttgataa aggcaacaga
7201 tacttataaa acacctcctc agattatatg ggttgataat tccagcatag ccacattggg
7261 caacttcagt gcttctactg gtaaggcaaa ggcgaaaaag accttcaacg tgtctgcctt
7321 ggtagccgct tccctcgcca acggcaaggt actgaactat cgcgccagtc tgccagaggg
7381 gaaacgcaag atactgtatg tagatacgga gcagagccgc tatcactgcc ataatgtgct
7441 ggagcgcatc ctgaagctcg ctggtctacc tacaagcatc gacaacgaga accttgactt
7501 tatctgtctg agagagtaca cgccctccgt caggatagag gtcatagact atgcgcttgc
7561 ccaagaccaa agctatggac tggtcatcat cgacggaatc agagacctgc ttctcgacat
7621 caacaatgcg ggagaatcgg tcgaggtcat caacaagatg atggaatggt catcaaagta

7681 tgacctgcac atccattgcg tattgcatca gaacaagggc gacaataacg tgcgcgggca
7741 tatcggtacg gagatgaaca acaaggcaga gacggtgctt gtcatcacca agagcaccac
7801 caatcccgac atcagcgagg tgaaggcgat gcacatccgt gagaaggagt tcaagccgtt
7861 tgcctttaca gtcaatgagg aagggttgcc agagattgtc gagcacacgc cggagaagga
7921 agaaggtgac aagcaaccgt caaggtttac ttaccaagac ttgacatccg aacagcataa
7981 cgaggcgctg acggctgcat tcaaggagaa acctatcaaa ggattcgatc gcatggtgga
8041 ggaattaacg caagcctatg cagacattgg ctttaagcgt ggcagaagcg tcatcatcaa
8101 aatgctgaaa tacctgatta acgagcaaaa gctcatagtg aagcgggata atcattacta
8161 tttcggatat acacctgccg agattgacct gttccatgaa gaggaataaa aacagttcag
8221 aaaccagttt agtttacgtc cgtacctata tataagaaat aaggccaaag taaaccgaaa
8281 tcggaggctg aattgggaag cggaaaagta gtttagcaaa tggtttagtt tacgggtgtg
8341 cctatatagg agaaagggta aagtaaactg tttccatgga caaagtaaaa ccgaacaaat
8401 gaaagtacaa tgaatattga acaatcgaag aaactgagca ttatagactt cttagacaag
8461 gagaatgtaa cgctgaaaaa gaagaagggc aatgcctact ggtatctgtc acccttcagg
8521 gacgagaaga ccgcatcgtt caaggtcagc aagaaggaga acctgtggta cgactacgcc
8581 attaaggagg gcggtgacct tgtggagctg gtaaagcgta tgtataacaa acaatccgtt
8641 tccgatgccc ttgcctacct tgcgtccaag agtattgcca ctgtggataa ggcaatcgag
8701 accgctatcg cagccaagga atacacgaca accaagatga acgatgtgaa gctgctccca
8761 ctgagcaacc attctctgct gtcctatttc agtagccgac ggatagacat caccattggc
8821 aggatgtatt gcagggagat acactacaag gttgagcaaa aacattacta tggcattgca
8881 tttggcaatc tgagtgaagg acatgaggtg cgcaatccct atttcaaggg ctgcatcggg
8941 cataaggaca tcaccctgct tgcccatacg ttcaacgagt ggcaaaacgg ttgtctggtc
9001 tttgagggct tcatggactt ccttgcttat atgaccctcg ttaaacagca agaccgatgg
9061 tttgtcgttg aaagcccttg cgactatatg atactgaact cagtcgccaa cctcaaacag
9121 gcattgcatt accttgaccg ttacacccac atccattgct tcctcgacaa cgaccaggca
9181 ggacgcaaga cggttgagag catcagcaat gtattcgagt acagggtgac ggacgagtcc
9241 ttcagatatg ccgactacaa ggatgtcaac gactatctga tgaggaagaa acgaacggca
9301 tctgaatgac attataaaga cgaaaacacc atcatttgat gcaaaaatgg ctcaaatgca
9361 ttgaaattac atatcaaatc tgcacttttt cgcccaaaca tgcttgattc tcaattatta
9421 tccttatttt tgcatcgttt taatacatta ttaaaatgta gcagaatgga attgaaaaaa
9481 gaaataagga tattgggcag acgctttgag gtggaacctc gtgccaagga aagcctgaaa
9541 ggcatcacgg ttggcgagaa actctcatac cgtttctatg atggcacata ccaaggcacg
9601 cccctgctgt tcgttgagcc caagaagggc aatccctctc cacggacttg cgccatcacg
9661 ggcaagcggt tgaccgaagc cctgggactg cctgcggtgt ttatccttgc gcccggaccc
9721 acctacgagc gacacagact ggcagacaaa ggagtcttct ttgtcatgtc cgaggaatat
9781 gcccatctgc caggcatcat agcattggag aagacaagca acaggaagat tgccgaagta
9841 ttgacccctg tggcgcaata catactactc tatcacctac aggtgggcag tattgaagga
9901 atgtcgccaa gggatattgc accgttgctt ccatattctt acgaaagcgt cacgttgggt
9961 gtcacctgtc ttgaagatgt aggactgtgt caaaagattc agaacggaca gcgcagcaag
10021 gtggtacact tcgagttgaa gggcaaggaa ctgtgggaca aggctcagaa cgtcttgctg
10081 tcacctgtgg agaaccgcat cttttgcgat gacatacgct tggatgcgga ataccccgta
10141 tgcggcatca atgccctggc acattattcc atgcttaatc gtgaccggga agagatgata
10201 atgatgacaa gcaaggaata tcgtgccgtc aagtcggctg atgtcatgga gaatcccaac
10261 atctatgacg gcaactatat catagaggta tggaaatacc ctgttgtcag caaaatgggc

10321 gacaaaagcc agtgggttga tcgcctttcc cttgtccttt cgttgagaga tgatgatgat
10381 ccgagagtag aaaaagaagt agaacgaata atcagtgaac agaaatggaa ggattagtca
10441 agttccgtga agcatttgcg gaatactcag aaaactatgt ggtgataggc ggagcagcct
10501 gtgacatcac catgaccaat accgtggtgc gtccacgcgc cacacacgac atcgacatga
10561 tagtcatcgt agagaacatg accgaggctt tcgccaatcg cttctggcag tttgtgcggg
10621 aggctggcta tcgacctgaa aagaggaaac aggaggctgg tgagccacca aggtatgaaa
10681 tgtatcgttt ccttgatggt aaagacggct atcccgagat gatagaactg ctgtcacgcc
10741 accccgacgt gcttggagag ccaaagggat ttgtcataga acctatcccg accgatgaag
10801 atgtgtcgag tcttagtgcc atcatcatgg acgatgatta ctatcacttt actatcgcac
10861 acagccaact gactgatggc atacgccatg ccaattccgc tgccttgata gcactgaagg
10921 caagagccta cctaaatctc atggcagaca agcgggacgg caagcatgtg aacaccaaag
10981 acatcaagaa gcaccgttcg gacatcctga aaaatgtggt cataatgact gaggacaaca
11041 ttgaggctcc tgcttcaata gtggcatgta taagggagtt tgtcgcctcc atcagagccg
11101 actggtcaac tctcgcagaa ccattgtcaa agtcactcgg tcaagacgag gcatttgtga
11161 ccggactgtt agatcaactg gatgaattgt ttatagaagc atgaccaata tgaagataca
11221 atacgcatcc gacctgcatc tggagtttgc cgacaattgg cggtatctga aagcccaccc
11281 cttggaggtg acgggcgaca tactgctgct tgccggcgac atcggctatc tcggcgatga
11341 caactactcc aagcatccct tttgggactg ggcatcggag aactacaagg aggtgcattg
11401 ctgcatgggc aaccatgagt tctacaaata ttatgatgtg gcgacactgc ccgacggcta
11461 tctcttggag gtacgtccca atgtattcag ccactacaac ggtattgcga ggattggcga
11521 caccgacatc atcctctcca ccctttggtc gaggattcct ttggaggacg cttacttcac
11581 cgagcaagtg ataagcgact tccgccgtat attatataaa ggtgagctga tgacccatgc
11641 ccaattcaat gctgagcatg aacgttgcct cacctttatc aaggatgcag tggcatattc
11701 gcaagcagcc cataagattg ttgtcaccca tcatgtacca tcgttccgta tgctccaccc
11761 aaagtttcaa ggcagcaagg caaacgtggc gttcaccgta gagttggagg actatatcac
11821 cgatagcggc atagactatt ggatatacgg acattcgcac accaacatcg atgccaggat
11881 aggcaacacc caatgcttgt ccaaccagtt gggctatgtt ttctctaacg aacatcagga
11941 cttctcccat ggcaagtacc tgaccatata gccatccaca ttttttcat ccaccctctt
12001 tacaagctct tccctatcgg aagggctttt tttgtgcctt tcgagtttga tttttgagta
12061 tgattgcgag tgtaatcaaa attagagtta agatttggct ccgcggaaag tcatttcggt
12121 ttaagattga agaaaaggta gttcctttgc atcggcaata agccaaacca tagtaaaatc
12181 aaacttgaaa atatgaagtc atttgtaata ttcgccattg tcgtgaccat aatctatgtc
12241 atctactata cggtcatcat cgtgcaggat ctgtacggaa agcccaagga cgagaagtca
12301 cagggcgagt cgtttgatgt cagtgacatg accgacgagg aagagtcgat agcggtcagc
12361 gagagcgacg gtggtttcag cgtaggcgac aaccaatatg agacagccta cgaggagaag
12421 cagcttgcag agcctacaga ggaggcagcc gccactgcgg aagagagcaa gcctcatgtg
12481 cttgaaaaga tacaaagcgc aatcgagaag aagatggagg aagtgaatac cacctattcc
12541 gaccccatgt attcggaaga attgaacaac accatcatcg cccgcggtct tcgtcacaat
12601 ggacgtgaca tggtcaaggt ggaatcactc aataatgaga tatgatgtca aaaacaagaa
12661 gtgcatttct atcattatat gccgctttaa ccccagctct ggccagtgct aagtgtggca
12721 acgtggacta cagctgggga gcggacgcac ttgcgagtgc ccatgactat gccgtgacga
12781 tgatgctcta tatcgtttat ctgtgctatg cagtggcggg catcgtggtc atagtcagcg
12841 ccttgcagat atatataaag atgaatacgg gcgaagaagg agtgaagaag aacatcatga
12901 tgctggtggg agcctgcctg ttcctcatag gagcgaccat cgtcttcccc gctttcttcg

12961 gttatcagat ttgagataca ggtgatacga gataagataa ctcaaagtgt aacaaattaa
13021 aaggtaaaaa gaatgtttga aaggattaac aagaaggtga agggattctt ctcttcacag
13081 cgcatgaaaa cgctcgccct gatgttgctt gtcggcacca ccgcagcaat ggcacagaac
13141 gcagccggcg actactcggc aggaacgacc gcgcttacca cagtgacgga ggagattgcc
13201 aagtatgtac ctatcgtggt gaagctctgc tatgccattg caggtgtcgt tgcaattgtc
13261 ggagccataa gcgtttacat cgccatgaac aacgaggaac aggacgtgaa aaagaagatc
13321 atgatggtcg tcggcgcatg tattttcctc attgcagcag cccaggcact gcctctgttc
13381 ttcggcattg gaggttaaac gcagcgtgag attgtgagta tgtagaacgt taaaacactg
13441 acagttatgg cagatacaaa gcaagagcgt tttccggatt atcccatttt caagggactg
13501 caacgtccat tggagttctt cgggttgcag ggacgctaca tctactgggc ggcagccacc
13561 gcaggaggag ccgttgcggg tttcatcctc ggctactgca tcttcggttt cgtggcagga
13621 ctcgtgctgc tagtcctcgc cgtagctgtc ggtggcgttc tcatcgtcat gaagcagcgc
13681 aagggcttgc acagcaagaa gagtgacaac ggcatattca tctatgccta ttcaaagaaa
13741 gtgtaaaaca aaaaagagaa tccattatgt ccaagtggaa agctatcaat gattgtaagt
13801 gaatggaggt gggggcatca cgcatgaggg tgcgcccgcc tttttgcgat taaaaaagaa
13861 aaagaacaat gaccctatac atcatactca tattctcagc catctgcgtg ggcatggcca
13921 tcagtgtcaa ggcattcggc acgggaggga agcgcaagcg catcttccag gacatctact
13981 tctccatcga ggaagtggat ggcatcggcg tgctctatac caagacgggc gaatactccg
14041 ctgttctgaa gatggagaac cctgtgcaga agtattccgc caacatcgag gcctactatg
14101 agttcaccca cctcttcgcc gcccttgccc agaccttggg cgaaggctat gcgctgcaca
14161 agcaggacgt gtttgtgagg aaggcattca aggaagagaa cggtggaaat catgagttcc
14221 tgagcgagag ttacttccgc tatttcaacg gcagaccctt taccgacagc gtctgctacc
14281 tcaccatcac acaggagaac aagaagagcc gtctgatgtc gttcgacaac aagaaatggc
14341 gtgacttctt ggtgaagata cgcaaggtgc acgaccagct gagggatgcc ggcgtgaagt
14401 cccgattcct cggcaaaacg gaagcctgcg agtatgtgga ccgtttcttc tccatgaact
14461 tccgtgacaa ggtggtgtcg atgaccaact tcaaggtcga tgacgagacc atcggcatgg
14521 gggaccgtag ctgcaaggtg tatagccttg tggatgtgga ttatgccaac ttgccgtcgg
14581 tcatccgtcc ctacgcaaat atcgaggtga acaacaccag tatgcccgtt gatctcgtga
14641 gcatcgtcga tagcgtgccc ggagccgact gcgtggtgtt caaccagatg gtgttcgtgc
14701 ccaaccagaa gcgtgagctt gccctactcg acaagaagaa aaaccgccat gcctccatgc
14761 ccaaccccag caacctgatg gctgtagagg acatcaagcg tgtgcaggag gtgatagccc
14821 gtgagagcaa gcagctcgtc tatacccact acaacctcgt ggtggcagtg agcggcgaca
14881 ccgacataca gaaatgcacc aaccatctcg aaaactcctt ctcgcgcatg ggcatccaca
14941 tcagcaagcg ggcctacaac cagttggagc tgttcgtcaa ctctttcccc ggcaactgct
15001 atggcatgaa tgccgactac gaccgattcc tcaccctcgg cgatgccgcc acctgtctga
15061 tgtacaagga gcgcatcgtg cataatgagg acaccccgct gaagatatac tataccgacc
15121 gccaaggtgt gcccgtagcc atcgacatca caggaaagga gggaaaggag aagctgaccg
15181 acaactcgaa cttttttgt ttgggtcctt cgggcagcgg caagtcgttc cacatgaaca
15241 gcgtggtgcg tcagttgtgg gagcagaaca ccgacatcgt catggtcgat acgggtaact
15301 cgtatgaggg actctgcgag tatgtgggcg gcaagtacat cgcctatacg gaggacaagc
15361 ccatcaccat gaaccccttc aacatcagca agcgggagct gaacattgag aagatagact
15421 tcctgaagaa cctgatactg ctgatatgga aaggctcgga gacgcagata cccgaactgg
15481 agttccgtgt cgtggagcag ttggtgacgg agtattacga tttctatttc aacggtgtgc
15541 agccctatcc ttcctcacag aaggagacac tgagaaaaaa cctctccacc atggagaagc

15601 ggcgcggcac ggagctgacg cagatacacg acaaggtgga gaaactcatc aagggactgg
15661 aggaacggcg catggcattg agcgtgaaga ccctctcctt cgactcattc tatgagttcg
15721 cctgtgagcg tctcgaccag atctgcatcg agaacaacat caccaccatc gactgcgaca
15781 acttcgccta tatgctccaa aacttctacc gtggcggcaa gtacgacaag attctcaacg
15841 agaacgtgga tagcaccttg ttcgacgaga ccttcatcgt cttcgaggtg gatgccatca
15901 aggagaacaa gcagctcttc cccatcgtga cgctcatcat catggacgtg ttcctgcaga
15961 agatgcgcct gaagaagaac cgcaagtgcc ttgtcatcga ggaggcatgg aaagccatcg
16021 cctcgccgct gatggccgag tacatcaagt acctatacaa gacagcccga aagttctggg
16081 cgagcgtggg cgtagtgaca caggagatac aggacatcat cggcagtccc atcgtgaagg
16141 aagccatcat caacaactcc gacgtggtga tgctgctcga ccaaagcaaa tttcgtgagc
16201 gattcgacga gatcaaggcg atattaggat tgacggatgt ggattgcaag aagatattca
16261 ctgtcaaccg cttggataac aaggagggaa gaagtttctt ccgtgaggtg tttatccgac
16321 gtggatcgac gagcggagtg tatggtgtgg aggagccgca cgagtgctat atgacctaca
16381 ccaccgagcg agctgagaag gaggcattga aactctataa gcatgaactg aagtgcagac
16441 atcaggaggc tatcgagcgg tattgcagag attgggatgc cagtggcatc gggaagtcgt
16501 tggcatttgc gcagaaggtc aatgaggcgg ggcatgtgct caatttgacg gacgatggag
16561 ccactcgccg ctgacacggc gagcacgggg gctgattgat ggaaacggaa atcccggaat
16621 agacgtaaag cccagatgcc cgaaagcaaa actttcggga acggtggctg ggtagtattt
16681 acgattagta gtaaatacga ttagtagtga atactaccaa tagtaaatac tacaagtagt
16741 aaagtagtgg aatagtagaa actgaaagta gtaaaaacgt aaagtagtga agcaatgaga
16801 cgtttatacc ttattataat atatatgtgt cttacacttg tcgtgggaag ggttcatgcg
16861 cagtattaca gcgtgaacta cgacaagcag atcgttgccg ccatggctgc cgcattcggt
16921 gctggagcta tggcagagag ctattataac gagcaggtgg gcgaaatctt gaagcactac
16981 aatgccgcag aggttgccac tgccggcatc ttcgctgcca agttcttgga gcgcaaggca
17041 ttcaccgacc tcggcatttg gagcagtagc acggagaact actattaccg ccgcatccac
17101 aatatggtgg cgaacaagat tatgcccaag atatggacgg tggcaggaat gatgctgaaa
17161 agtccgcaga ccgccctcta ctggggcagt tacctgatga agatatgcgc cgacacaaag
17221 tcgctgtgta tgcagtttga gagtgtcgtg accaacagta gcttgacctt tgccgacatt
17281 cagttcttgg aaatcaacca agagatagcc gccatcctca aactctcgga gataggcaat
17341 gtagattggc agcagctgtt ggatgacatc gccagtgtgc ccggcaactt caccgtggag
17401 aacctcaccg ccgacatcga gacgctttac aatatgggag catcgttggc gacggcaggt
17461 atcaacgggg catcaggcgc actcttgcag cagagtgcct tcaacgacct tttcaatggc
17521 aaggtgagta aaatcatcga catctatgac aactaccatg acctctatga acaggcagag
17581 aacggcatcg gcagcacact ccttagtctg gtgggtggcg aggacaacgt ggcagccctc
17641 ttcgacctct ccaactacaa catcacctcg tggatgacag actatctgag cgagacgcag
17701 ggcaactact atacacagcg ttggtacatt gcccgacgag accaaggcag cgtaagcctc
17761 tgtgactact atccgccgac cgatgacaac agtattctga acggtgacca ttggtatcgt
17821 atcaacacct ccgaccccaa cttctaccct aacagcacgc agcgagagca gatactggcc
17881 aactcggaga accatgccgg atggagccgc agccgtgtgc agcagctcaa cgcccagagc
17941 gacggcttca cctatacgat gaactattgg atgtcgtcgt acatcatcag ccgtggcggc
18001 aagcagacca agaaggccta cgcctacgag atacacgtta ccaagagctg gaacaacgtg
18061 gaagaggtgt atgaggatgt cttcgactcc tacagcatgg acttgaatac cttcaagcgt
18121 cagttgcagg cacgtctctc ggagtataac gacaacgagg agggctatgt ctattacatc
18181 ggcagtgacg cacgccgtta ctatcaagcc accgatgccg ccaaactgaa gggcgtggag

18241 agcgtgacca tcagcgtcac ctgttcggac ggtgtcaccc tcggacaggg cagcacgcag
18301 tacaagtgcc gtgagtgcgg cggttcgctc aatgcccaca gcaaggaatg cgccatgagg
18361 acaagcgtat cagaaaataa tctcgaccta tcggaactgg acgagaagga acgtgaggcg
18421 aatagcaaga tagccctgtt ggaggcgcag ataagccagt tggagacaga gaacaggaac
18481 ctgctcacgc agatagccaa tgcctctgtg gaggaagccc ctgccttgcg gcagcaacat
18541 aatgtcaaca agacgaagat agacaacctg aagaaggaat tggcgacatg gcagcagcag
18601 ttggcagaca tacagaatgc caagagcgag gcggctaacg acaatgccac ctccaccgat
18661 gactactacc gcatccctgc catcatgcag gatgtgaagt cagcctacaa tctcacgtgg
18721 caaggtgccg gttcgtggaa cggctacacc tacgtccgaa ctgccaccat gcccaacatc
18781 aacggcatta tcaccttcaa ggcgacactg tccattgcaa ggaagcccaa gtacttcctt
18841 ggcatcaaga tacaccgtgc gatactgcaa atatcgtggg agctgacctc ggaatacacc
18901 gacacacagg tggtggatgt gctgaccctt gacccgaatg cctctgacga ggagaagacg
18961 aggcaggtga acagccgtat ctcggagata gcccaggagt tcccgtcgtg cagcatcacc
19021 acagaatatg ccaagagcga ggcaacccag accgccgaca atgacgatgt aatacacctg
19081 ctctggtcga gcgaccgtct ggagatagcc cgtgaagtag atagccgtat caccaagatt
19141 tacgccgact tggtatcatt agaaaaaatg atgcactaca agcgcagcat cattgacgtg
19201 ctgaaggaca ttcttccttc aatcgacgca gaccaaggtc gcagaaggac actcattgaa
19261 gagtgctacg agcgatggag agagaatgca gaccctaaca gagggaatgg tgaagataac
19321 gagaacgaaa actgagggag gaacggagaa tgagacatac catcatcata gctctcctgc
19381 tgattaccct cttccccaat acagccttgg cgcaatgggg attcgatgcg gtatcggtgg
19441 aagcatacat caacgaccac aagaagcagc gcagcctctt gctggcacgc agcactttgg
19501 agtacagcaa tcaactgctg catgaataca gcagtgaaga ggtgggcaaa tacaaggaac
19561 tgaatgtgga tttggacaaa tacacccggg ccttcgacat catcgatgtg atgtaccagt
19621 cgttgcgcac ggtgctgaac gtgaagagca cctaccagac ggtgagcgac cgcatcagtg
19681 actacaagaa cctgttggag gacttcaatg ataaggtcat caagcggaaa cacatcgaac
19741 tggctgacac gatgctcatc agcatcaaca agaaagccat cgacaacata tacaacgaag
19801 gaagccaact ctaccatagc gtcaacgacc tcgtgttgta tgccacagga gcggcagcct
19861 gttcaacttc cgacctgctg atggtgttgg acaatatcaa caattcgttg gacctgatag
19921 aaaaacatct gaaccaagcc tacttccaga catggcggta tgtgcagctg cgcatgggct
19981 attggaagga gaaggtgtat cggacgagga cgaaggaaga gatactggaa gatgccttca
20041 gccgatggaa agtggcagga aaacttgact attgacggat tgtaaacaaa agaagaacgg
20101 acatgaagaa gacaatgata ctcatcatgg cggtagcaac caccatcaaa gccacagccc
20161 agagcgtgac ctacaaccac gactcctcca agcagaacca gataacggtg atggagacag
20221 gtggcggttc tctcacccct gagttctatt attggctgtt acataacaac tataagaaga
20281 cagcggcaga gaagaacaag ttaggtttcc gcactcttgc cggcatcaac ctctacaacc
20341 aagtagatga tgcggagaag atagactctg ccctcaccaa gcgtgctgag gtggaagcct
20401 tgaacgtagc cgaccgacag atagacctcg cctggcttgc cgaaagcagc aagatcaacg
20461 gtcagttgga caagatgaag gcgaacatag accatatcat ccctacagga ggcaccatca
20521 acgacaagcg tcggtgggag gaactgtaca atatgtacca gtgtgccgtg aaagccacca
20581 aggatgccta tatgcccaat gcacagagaa agcggcagta tcttagcatc tatgccgacc
20641 tgacgacaca gaacgagaca ctactgaaat accttgtgca gctcaacacc caaagccaga
20701 ctaccgccct cttggcagcc accaacgacc gtgtagtaca taaaggcagc atcatcagcg
20761 atgccaagag ccgttggcag gaaaacatga agggcgtgag gagctcgaca ggcactgacg
20821 gcgatgatgc caacagcggt gagggtgaag aaagtgtgaa cagatagaaa taagggaaga

20881 gaagaaataa tcacaaaaca gcatagagat atgaacgata ttcagatttt caacaacgag
20941 gaatttggag cagtccgaac cacaggaaca ccagagcaac ccttgttctg tttggctgat
21001 gtggcaagag ttcttggact aaaaaccagc aagttggtac aacgactttc ggatgatgta
21061 ctttcaaagt accccatctc ggatagtctt ggaagagaac aggttacaaa ctttatcaat
21121 gaagacggac tgtatgatgt catccttgac agtcgcaaac ccgaagccaa acgtttccgc
21181 aagtgggtga catcggaggt gctcccctct atccgcaagc atggtgccta tatgacacag
21241 cagaccatcg agaaggcatt ggcagagcca gacttcctga tacggttagc agtcaatctg
21301 aaggaagaac gacaaaagcg tctgcttgtc gaacaggagt gtgaacacca aaggacacgc
21361 atcgttgagt tgggatccaa ggtggatgac ctgcaacagg aggtgacgga gatgaaggac
21421 aaggtgagtt accttgacat catccttgcc acgaagagca gtgtcctggt cacacagata
21481 gcacaggact atggcgagtc gtccatccgc ttcaaccgac gtctgaagga gatgaacatc
21541 cagtatcagc gtggcaagca gtggattctg tatgccgact acaaggactg cggttatgtg
21601 accagcgaga cctacctcat taaacacaag gacggtacgg aggatgtgcg catgaacacc
21661 aaatggacac agaaaggtcg ccgcttcctc tatgagaaac tgaagagcgt gggagtcatt
21721 cctgttatcg aaagaacata aaataaagaa atagctatgg aatcaatatc aaaaatacag
21781 ttacgtctct atgccgccaa gaggaaaaac ggcaagtggc agttggagat gtcaaggatg
21841 ccgaagcgaa tcagcgttat cggcaggaca cccatcgttg atgagcatta tatgccatct
21901 gatttagagg tagtttcaat gtcaaagctt cacaaatatg tcggaagtta ctacggcaag
21961 attgtgaaga ccctcaaaga ggaaggcatc atcaccaagg aatatggaat gtggaaactg
22021 cgtgaagacc ttcaggacaa gggcattgcg gtatatgtca caggcaggat gcgctgcttc
22081 taccacttct atctgtcatg gacaccgaaa ggcatagagt ttatcaaaga gattatcaac
22141 aatagaacca ggcactgatg gcagacaaca tactcagcga tttcggcatc aacatcctcg
22201 aagaggagat tgacgatgtg atattccaga ccaacgagtt cctgaccgat gccaccttca
22261 caggctcaca agggccgttc tggtggatcc tccaaatgtg tatggcattg gcggcactgt
22321 tctctatcat cgtggcggca agcatggcct acaagatgat ggtgaagaac gaaccgttgg
22381 acgtgatgaa actgttcaag cccttggtgg tgagcatcat cctctgttgg tggtacccac
22441 ctgctgacac aggcatgacc aacagcggca gcagctggtg cgtattggat tttctatcct
22501 atattcccaa ttgcatcggc agttacaccc acgacctcta tgaggcagag gcgacccaaa
22561 tttccgacaa gtttgaggag gtgcagcagt tggtatatgt cagggactcg atgtacactg
22621 acctccaggc acaggcagac gtggcgcata agggcacatc agaccccaac ctgatagagg
22681 cgacgatgga gcagacgggc gtggacgaag tgaccaacat ggagaaggat gccagcaaac
22741 tgtggttcac ctcactgacg gcaggggccg tggtgggcat cgacaagata gtgatgctga
22801 tagccctgat cgtgttccgc atcggttggt gggcgacgat atactgtcag cagatattgc
22861 tcggaatgct cactatattc ggtccaatac aatgggcatt ttcgttgctg cccaagtggg
22921 aaggagcgtg ggcgaagtgg ctaacccgtt atttgacagt tcatttctac ggagccatgc
22981 tctacttcgt cggtttctac gtcctgttgc tgtttgatat tgtgctctgc atacaggtag
23041 agaacctgac ggccatcacc gccagtgagc agacgatggc agcctatctg caaaactcgt
23101 tcttctcggc aggctatctg atggcagcga gcatcgtggc gttgaaatgc ctgaacctcg
23161 tgcccgacct tgcggcatgg atgataccag agggcgacac cgccttctca acccgaaact
23221 tcggcgaggg cgtggcacaa caagccaaga tgacggctac gggaaccatg gcgaccgtga
23281 tgagataaca tgtgagtgtg cggtgagtca ccgcacctcc tattgaaaca aataaaagta
23341 accatcaaga aaaatgaaag caaaaaatga aattgagcgt tggctcaaag atgagaaatt
23401 catggcgttt gccaacaagc gtgcaaagga agagtttttc aattccgaaa acaactacat
23461 cgacccacag tatgaggaga tggcagaagg ctttgaggat aacgacgagt atgtagtgcc

23521 tatggtagat tacctgagtt atcgcttgca tcgtgccaaa atctacagaa atcgacggag
23581 acgtgaacgt gacatctggt gggtgtggat ccaactgaaa tacgaaggca tctacgtaga
23641 ggcttgcatc aagtattacg ccaaactggt ggaagaagtt gagaaggaca tatacaccat
23701 cctgcaccgt gaatatgtga gaatgaagag aaatcagacc tcaaacaagc agtaacgtat
23761 ggtaatcaag aacttagaaa acaaaatcaa gctggtgggc atcatctgct ctgcgttcct
23821 catgggctgc atcatcatct cggtgtcgag catctggacg gcgcgctgca tggtgacgga
23881 cgcgcagcaa aaggtgtatg tattggacgg caacgtgccc atactggtga accgcaccac
23941 gatggaggaa acgctggatg tggaggcaaa gagccatatc gagatgttcc accattactt
24001 ctttacgctt gctcccgatg acaagtacat ccgctattcg atggagaagg cgatgtattt
24061 ggtcgatgaa actggtttgg cgcagtacaa tacgctgaag gaaaagggct tttacaacaa
24121 tatcatgggt acgagtgccg tgttcagcat cttttgcgac agcatcaagt tcaacaagga
24181 gaagatggag ttcacctact acggtaggca aaggatagag cgaaggagca atattctgat
24241 gcgtgaattg gtgacggcgg gacaggtgaa gcgcgtgccg aggacggaga acaatcctca
24301 tggactgctc atcacgaact ggagaacctt gctcaacaag gatattgagc agaagtcgaa
24361 gtcaaacttc taatatccta gcggtatgag tgtaaaagga atcagacgga tgcttgtggg
24421 cgaggatatg cccgacaaga acgaccccaa gtacaagaag cggtatgaga aggaggtggc
24481 ggcagggcgc aagtttgcca agaccgcgag gatagaccgt gcggcggcta aggtgcaagg
24541 ctttgccaat gcccacaaga cgctgtttct cgtcatcgtc ttcggctttg tcgccacgtg
24601 cttcggcatc aacatctaca ggatggtgcg gtattacggt catcggacgg aggctgcaag
24661 tgccatcgag cggcaggaag ggcgcatgaa gcagatgatg aatgcagccc attgcatcac
24721 tcctcccgtg acgcatcgaa ccaacgagga gaatcgagac aagtcagtaa catcaaaatc
24781 agacagcgat gagaataaca gataagataa acttccgtca gccgaagtac atgctgccag
24841 ccatcgtgta catcccgttg attgccacag gctattttgt ctttgacatg ttcaatacgg
24901 aggtggcgga aacgcaggac aagaacctgc agaccaccga gtttctgaac cccaacctgc
24961 ccggagcgca gattaagaat ggtgacggca tcggtggcaa gtatgagaac atggcgaagt
25021 cgtatggtcg catagcagac tattcggctg ttgataacat tgagcgtgac aacgaagaag
25081 agaaagaagc gtatgaatcc aagtacacag aagaagacct tgcggaactc gtccgtcagg
25141 cagaagagaa ggacgatgca gccgaagtgg ccgatgccaa ggcgcgagaa gcggaagcac
25201 tcgaagcact caacaaggca ctcgccgaag cgaggctgag aggacaggcg gcagttgccc
25261 ccgtggccca ggataccaca caggctgggc ctcccaagga gcagattgag gtgaaaggtc
25321 agattgccga ggacagcaag gcggtgaagg cactggacga ggaggacaag gcacaggagg
25381 tggtgaagaa ggtgaaggtg acttcggact atttcaacac cctgacgcag aacgagcatg
25441 agccaaagct catcaaggcc atcattgacg aggacgtgaa ggctacggac gggtcgaggg
25501 tgcggctgag actgttggac gatattgaga ttggtgagac cgtggtgaag aaagggtcgt
25561 acctctatgc caccatgagc ggtttcggtt cgcagcgagt gaagggtagc atcaactcca
25621 tccttgtgga tgacgagctg ataaaggttt cgctttcgct ttatgacacg gacggactgg
25681 agggactgta tgtccccggc agccagttcc gtgagaccac gaaagatgtg gcgagcggtg
25741 ctatgcagag caacatgaac atcgaccaga gtggagcgaa caacagtttc tcgcaatggg
25801 gaatgcaagc cgtgaccaat gcctaccaga agacaagcaa tgccatcagc aaggctatca
25861 agaagaacaa agtgaagctg aagtatggga cgtttgttta ccttgtaaat gggaaagaga
25921 aaaggaaata gacactgccc gacaacggaa ttgtcgggaa cggtggctca tagagtataa
25981 acatcaaaaa atcaactaat atgaattgtt ttcagaaaat gaatgtctgt gcattgcttt
26041 cgctgagcgt gctttcagcc aaggcgcaga ccacctacat ggagatggag cagctgacgg
26101 tcaacgacca agtgaccacc gtcatcaccg cctccgaacc catccgcttt gtcgatatat

26161 ccacagacaa ggtggtgggc gaccaaccca tcaacaacac catccgcctg aagccgaagg
26221 acaacgtgta tgccgacggt gaggtgttgg cgatagtgac aatcgtgacg gaacgctacc
26281 gcacgcagta tgcactgctc tataccacga ggatgcagga ggcagtaacc gataaggaaa
26341 tagaatgctc ggagcgcaat gcctacaaca atcccgctgt aagcctgtcc actgccgaca
26401 tgaccaagta tgcacgtcag atatggtcat cgtctgccaa ataccgcaac gtggcgacga
26461 agatgcaccg catggtaatg cgcctgaaca acatctacag cgtgggcgaa tatttcttca
26521 tcgacttctc cgtggagaac aagaccaaca tccgcttcga catcgacgag atgcgcatca
26581 aactatcgga caagaagcag agcaaggcta ccaatgcgca gatagtggaa ctgaccccat
26641 ctcttgtgtt ggacgatgcg aagaccttca agtatggcta tcgcaacgtg attgtggtga
26701 agaagatgac cttccccaac gacaaggtgc tgaccatcga gctgtcggaa aaacagataa
26761 gtggacgcac catctcgctc agcatcgact atgaggatgt gctgtctgcc gattcgttta
26821 accatatact gttggaggag gaataatgag aattaggaat gaggaattag gagtgaggaa
26881 ttggcttgtg cataccttac tcgttgtcat ctgttttatg gcaagcatga ctccggcaat
26941 ggcgcagcag aatagcgacc gcatatcctt ggggttcgga tgcctgtatg agcgaggact
27001 ggacgtgacg ctgtcctacg agtatgagac gaagtaccac aatgcgtggg agtattttgc
27061 aaacggatat atcaagtgga acgagtgcgc atcgtgcggt catgtctgtc ctgagagttt
27121 ttggaacaac taccgcagtt atggtttcgg ctttgcctat aagccctgtg tggcaagagg
27181 acgcaaccac catggcaaca tgcgcatagg agcatcggca ggtagcgata ctgacaggtt
27241 cttgggtggc atccacctcg gctatgagca aaactataca ctcagacacg gctggaagct
27301 cttctggcag gtgaagaccg atgtgatgat caagggtgaa gacctgttca gaacgggtat
27361 tgtcctgggt gtgaaacttc ctgtaaaatg actatgcgta tgaagaaact gttttcaatc
27421 ttggcagttg cggcaacggt gttgctgact gcctgtgacg agcatcagga ctttcccgac
27481 acagccatga aggtcaccca tatcttgacc actgacggca aggtgatgcc ctacgaaacc
27541 tacgagcaat cgtgcaagca agccattgca gtggtgttca acgtcaatca gagggaagaa
27601 atggagggaa acggatatgc cgtctatctg tgggacatcg ctcccgaagc ctttgctgac
27661 agtatcggtg tggaacagga cacatcctgt gacctcaccg cctacgacgg caacaagaac
27721 acttttgccc tttatggcac gaccgatgtg aagtcgcctt tggctgagcg tgtcttcgac
27781 atgtggcgtt acgggcaaag tgcctatatc cccagtgtgg cacagatgcg tctgctctat
27841 catgccaagg acatcatcaa cccatacatc ctgaaatgcg gcggcacacc tatccccgat
27901 gaggatgatg actgttggta ttggacttca acggaggtag aaggacaagc gaccgccaag
27961 gcatggcttt actcactggg aagcggtgcc atacaggaga ctcccaagtg gcagccgcat
28021 aaggcgagac ccatcatcac gattatggat tagcccatat acagacatcg agtatcaccc
28081 gtatcatcat caaagtacaa caattaaaaa agaaaaagca tgaaaaagaa tttgtttcta
28141 atgttgggac ttggagccat gctcctcacg tcctgtgacc aagacatcgt caatgagatt
28201 tacctgcccg gtggcggagg ttctgtggtc gtgaccgaca gcacatcaac caagcagctt
28261 caggtattca ccaatctgga gatgctgcaa cctgccgttt ccacacgagc cgttgacaac
28321 cagtgggaac tcaatgatat gattggcatc tcttccacag ggatgatcaa caatatgaag
28381 ttcacccgaa gcggcggcac gaatcagttt gtgtcagcca acaaggtgtt tttcacggac
28441 acagacactc acaccttcaa tgcctactat ccctatacgg caaatcctac caacgacctc
28501 atagagttcc aagtgccgga ggcagctgta cagagtgagc agaaggtcaa tgattttctg
28561 tttgccagtg gagcggcatc atacgccaat ccgagcctta cattgaactt cacccatcag
28621 atggtgcgtg tcatcatcaa ggtctatacc tcaccagaat acggtttcac caccaatgat
28681 tttgcaggaa gcctactcac gttcgtcggt tattttgaca gcggtacttt caacatcaag
28741 accggaaagg tagaatgcag tgacgagtcg gtgacgacct attatttcgg cgacccacaa

28801 agtgaccaca tggaatatac gctttatgtt cctgcacagg ttatgccaga catgacattg
28861 caactcagca atggggagaa cttcgtattc ctcacaaacg acacgtggaa agcgggtcat
28921 tcctacagct attcattgaa gcctgtacgc aacaccttgg aggtcatatc tgcaactatt
28981 tctccgtgga cagtagagtc tgagaagacc atcaacggct acgaataaaa ctcataagag
29041 gattggaggg tgtccaccgt ggcatccttc atcttttctc attatcaccc aacatcatcg
29101 taaagtaaat ggaagaaagc aaagagttac agggttttta caagatattc agggcggtga
29161 tttatgtctc catcctgatg gagttcttcg cctacgcact ggagccggag cagttggact
29221 tcttgggcgg cgtagtgacg gatatccaca ctcgcatcaa gcggtggatg atttaccatg
29281 acggcaacct cgtgtatagc aaggtggcta cgttcctgct gatatgtatc acctgtgtgg
29341 gaacgcgcaa caagaagcat ctggagtttg acgctcgcag gcaggtgctc tatccgctgc
29401 tgtcgggagt gctgctgatt gtgcagtccg tatggctgtt cggtcattcg ataatgccct
29461 gtttctatac cctccgtttg aacacttggc tctatatgct cacttccatc gtcggtgtcg
29521 tattggtgca tatcgccctc gacaatatct ctaagttcct gaaggaaggg ctgctgaaag
29581 accgcttcaa cttcgagaat gagagctttg agcaatgcag ggagttgcag gagaacaagt
29641 acagcgtgaa catccctatg cgctattatt acaagggcaa gttccgcaag ggatgggtga
29701 acatcgtcaa tccgttccgt ggcacatggg ttgtgggtac tcctggttct ggtaagacct
29761 tctccatcat cgagccgttt atcaggcagc acagcgagaa gggctttgcc atggtggtgt
29821 acgactataa gtttcccact ctcgccacca aactctacta tcattacttg aagaacaaga
29881 atgccaagga cagcaagatg ccccatggca tgaagttcaa catcatcaac tttgtggatg
29941 tggagtattc gcgccgtgtg aaccccatcc agttgaagta catcaacaac cttgccgcag
30001 cgagcgagac cgccgagacc ctgcttgaat ccctgcaaaa aggcaagaag gaaggcggcg
30061 gtggcagcga ccagttcttc cagacctctg ccgtcaactt ccttgccgcc tgcatctatt
30121 tcttctgcaa ctacggcaag gagccctacg acaaggacgg aaagatgctg attgccgagc
30181 ggcgtgaaga cccaaagacg aaaaggctga tacccacagg acgggtgttc gaccattcgg
30241 gcgcggaggt gcagcccgct tattggttgg gcaagtacag cgacatgcca catatcctat
30301 cgttcctcaa tgagagttat cagaccatct tcgaggtact tcagaccgac aacgaggtag
30361 ccccctttgct tggcccgttc cagactgcct ttgccaacaa agccatggag cagttggaag
30421 gtatgattgg cacgctacgt gtctatacct cccgtctcgc caccaaggaa tcttactgga
30481 tattccacaa ggacggcgat gacttcgacc tgaaggtcag tgacccgaag aatccaagtt
30541 atctgctcat tgccaacgac cccgagatgg agagcatcat cggtgcactg aatgccttga
30601 ttctcaaccg ccttgtaacg agggtgaata caggacaggg caagaacgtt cctgtcagca
30661 tcatcgtcga tgagttgccg accttgtatt tccacaagat agaccgactg ataggcacag
30721 cccgaagcaa caaggtgagc gtggcacttg gctttcagga acttccgcag ttggaagccg
30781 attatggcaa ggtgggtatg cagaaaatca tcaccaccgt gggcaatgtg gtgagcggtt
30841 cggcaagagc caaggaaacc ctcgaatggc tgtcgagtga catcttcggt aaagtagtgc
30901 aactcaaaaa gggcgtgacc atcgaccgtg acaagacctc catcaacctc aatgagaata
30961 tggacagcct ggtgcctgca agcaaaatct ccgatatgcc cacaggttgg atatgtggac
31021 agaccgcccg tgacttcgta gtcaccaaga ccgggatgaa cggaagcatg aacatccagg
31081 agagcgagga gttcaagaca tccaagttct actgcaagac cgacttcgat atgacggaaa
31141 taaagaagga ggaatcggag tatgtccctc tccctaagtt ctataagttc aagtcgaaag
31201 aagagcgaga gcgcatcctc tacaagaact tcgttcaggt tggggaggac gtgaaagcca
31261 tgattaaaga gattcagcag ttccgcacta tgatgtagtg atattgatta gagtgcggtg
31321 agacatcgca cctcctatat aaaataatgt aatgagaaga tacatcatcg tactattgag
31381 catgttggcc atctgtagcg tgccctgtca aagccaaacc attgacaggg cactgtttga

31441 acggatggta gccattacca aacacttcga gggttggcac gacccaaaga ccacaccggg
31501 ctatgtgggc tatggtcacc agctacagaa gggagagcgt ttcccgaaaa cactgacacg
31561 tcagcgtgca gatttgctgt tacggacaga cctcctccgg cacctgcgcc tatatgcccg
31621 ttatggcagg gatgcttatc tgcttgccac cctctcgtac cagattggac cagccaagtt
31681 gttaggcaat ggacgatacc cgaaagcgtc cctgctcacc cgattggaac gtggcgacag
31741 ggacatcctg ccactctatc tgagctattg caaatggagg ggaaaagcag tggcatccat
31801 taggaaacgg aggtgggtgg aatatcagct gttgtatgcc gagcggtgag gcaccatccc
31861 tcctatacct gtttgtctgt cacttatatt atatataaag gtaataattg caatttcaat
31921 gttaaagtca gagcattatg gcatccatca aattaaagtt cagaacatca tctgttcaag
31981 agaaagaggg tcgcctgtat tatcaggtga tacacaatcg tgtggcacgt cagatccata
32041 cagagtacag aatctattcc tcagagtggg atgcggacca ttcaaacatc attttgccaa
32101 cttccgtcac tccacaacgt caagcctatc ttctttcctt gaaagataca ttggatgctg
32161 acaggaagaa actgctgttg gtcattgcac ggttggacaa ggaagggcag tcatatacag
32221 cagacaccgt cgttgacaac ttccatgagg gaaaagaatt acatggcatt atcggctata
32281 cgctggagtt gaacgaaaaa ctgagacgca tcggcaagaa gcggatggta gcaaggtaca
32341 aaaccaccct caacagcctc cagcgttacc tgaaaggagg cgatgtgcca ttggaggaag
32401 tcgatgggac aacgatacaa ggctatgaac agtggctgaa agacagcgga ctgtgccgca
32461 acaccacctc tttctacatc cgcaacctgc gcaccatcta caatcatgcc gtagatgacg
32521 gactggtcat ctcttcctca ccgttcaagc atgtatatac cggcatcgac aagaccgtca
32581 agcgggcact ccccttggaa atcatcaagc aactgaaaga gctggacttg agtctgaacc
32641 cccgtttgga gcttgcaagg gatatgttcc tgttcagttt ctatacccgt gggatgtcgt
32701 tcatcgatat ggcacaactt acaccaagca atctgcatgg cagtacgctc atttaccgcc
32761 gtcagaagac ctcacaacag ttgcacatca aatgggaacc tgccatgcag gagatagtgg
32821 cgaagtacaa aacagatgat tctccctacc tgctccctat agcaaaagga gagggagcca
32881 tattctggag acagtacaag aatgcctata gccgtatcac caagcagttg aagaaaatcg
32941 gagaaatgat aggcctgtcc gtacccctga ccacctacgt agcccgccac tcctgggcaa
33001 gcatagccaa gagcaagaac gtgcccgtat ctactatcag cgaggcattg ggcatgact
33061 ccgagaaaac cacacagata tacctctcct cattggacac ctctgtcgtt gacaacgcca
33121 acaatctcat catcaattct ctttaggagg tgcagcgtct cactgcacct ttatttcaat
33181 atttataact attgtattgc ggtgtgatac cgcaactata ataaacctaa aatccgcaac
33241 tatcatccaa tacacgatta agggtagatt tatgagtcgt tagtagcagc tttcagtaaa
33301 aagcaacagc acaacatcaa tatgtagcca ccatcccctt accccacgat gcgacttgag
33361 gtaatacgca gattcaaacc ggaaagaaag gattcttatc cgaattctgt tttgaagggt
33421 tttgcataag ctcggttctc tgtgtagata ttcagcacgt attgccttgc agtcatgatc
33481 cacttggcag gtacggagat gaatctgaag acaaaagcct ttatgcgact cgttttcttg
33541 agcccaaaag ccttggtgtc aagcctgctc atgatggtct tgtagaaatt gtgtatcaat
33601 gcagtaagca gaagaaagac agtattctcc gccatgaatg acttggggag cctgctccaa
33661 ccgaatccgt tgttcatgtc gtcaaagata cgttccttgc cgccacgcag attgtagaat
33721 tcaacaatgt cccttgtcga tgacttgtaa tcgttggtca gaatacaacg gtaagtgtat
33781 tcgccttccc acaggtcaag gtcgccactg ttgcgtcttt gtctctggat gacaagacga
33841 tagcacttgc cttcccattt ctcaacgaga atggaattga gttcgaactg gatgccgtta
33901 atctcctccg tcttccatcc tctcagagca aagatgtcat tgtagagcga actgcatcgg
33961 ttggcacgga tgtagaaatg tttgcaatgc ttctctatct cactgacgat ttccttcgag
34021 caggaaccgc agtctgccct gaagcgattt acacggatgt tctgggattc cagaagagcg

34081 aagaatctct tatgggtgtc tgcctgatga aaacgcacat tcgtgttacc atcgctgttc
34141 tcgatataga ctatcttgtc accgataaca tatacgccag gcctgtagcc gaggaacttt
34201 ttgtaggtcg gttttgcatc atacttctcc gtttcaagga actgatggtc aaagtcaaca
34261 tcgtattcct caatttcctt caactcgcct gtagaaacca aagcgtttat aagcaatgtg
34321 ttgagtttgt ctgcagtatt gaaatcatag gtcttgcctt ggtcggaagt ataggagatg
34381 ttttcctgtg tcagttcctt gatggctctg aggatggtat cagagctgca tgtacgaagg
34441 gtaggatgat acgagagatg gcgcatcagt tgtgacgtta catcttccac gcatgagccg
34501 ccacagaaat aaacgctcat cagcgaacgg actatctcgc tgaactgata tccgaagata
34561 ctgctgcatc tctgacccag tgttgagtcg ataacgggtg aaagcatgga gtcaaatttc
34621 tccatgattg aaaaaattcc tccaaaaggt gtgagttttt cgaattttat ttgtaccttt
34681 gcagcgcctt gttacgattt tcgcttgttt tctaattgca acactaagat aagtgaaaaa
34741 tctgacacgg caaaatccta ggcaacttat tgttgctcag gaacttataa ataaagttaa
34801 attatagtgt tgcggaatta agggtgttaa aatatacgct gatacaaata agaatttagg
34861 agataatata aatggaaaaa tacaacaatt ggaaacttaa gttttataca atatgggcag
34921 ggcaggcagt atcattaatc actagtgcca tcctgcaaat ggcgattatt ttttacctta
34981 cagaaaaaac aggatctgcg atggtcttgt ctatggcttc acttgtaggt tttttaccct
35041 atgcggtctt tggaccagcc attggtgtat tagtggatcg tcatgatagg aagaagataa
35101 tgattggtgc tgatttaatt atcgcagcag ctggggccgt gctagctatt gttgcattgt
35161 atatggagtt acctatctgg atggttatgg tagtattgtt catccgtagc attggaacag
35221 cttttcatac cccggctctc aatgcggtta ctccactttt agtaccagaa gaacagctta
35281 cgaaatgtgc aggctatagt cagtctttgc agtctataag ctatattgtt agtccggcgg
35341 ttgcagcact cttatactcc gtttgggaac taaatgctat tattgccatc gatgtattgg
35401 gtgctgtgat tgcatctatt acggtagcaa ttgtacgtat tcctaagcta ggtgatcaag
35461 tgcaaagttt gaaaccaaat ttcataagag aaatgaaaga aggaatggct gtactacggc
35521 aaaataaagg attatttgct ttattactcg ttggaacatt atatatgttt gtttatatgc
35581 ccataaatgc attatatcct ttaatcacta tggaatattt taatggtaca ccgatgcata
35641 tttctattac ggagattgct tatgcctctg gtatgttgat agggggtcta ttattagggt
35701 tatttgggaa ttaccaaaag cgaatcttat taataacggc atcaattttt atgatgggga
35761 taagcttaac catttcagga ttacttcctc aaagtggatt tttcatattt gtagtctgct
35821 gtgcaataat ggggctttcg ggtccgtttt acagcggtgt gcaaacagct ctttttcagg
35881 agaaaattaa gcctgaatat ttaggacgtg tattttcttt aactggaagt atcatgtctc
35941 ttgctatgcc aattgggtta attctttctg gattctttgc tgatagaatc gatgtaaatc
36001 attggttttt actatcaggt attttaatta tttgcattgc aatagtttgc ccaatgataa
36061 ctgagattag aaaattagat gcaaaataaa ataaaggagt gtgttcgtat gatatataat
36121 attattcaat cagtgacaga aaaattatcc tctttgcctt ttatagaagg catcgtatta
36181 gggggatccc gtgcaagagg tacccataca gaggattctg atatagatat cggaatttat
36241 tacaatcaag aatcatttga cctgacagcg attaaccaaa ttgctacaga gttggatgat
36301 gagaatagaa acaaccttgt tgtacctccc ggagcgtggg gtgattgggt caatggaggc
36361 ggatggttag ttattaacgg gtatcatgtt gatctgattt tacgtgatat aaaacgggtg
36421 gaacaaataa tcaaggatac ggagcaagga attgttactg caaattatca gaccgggcat
36481 ccccatggtt atataagtgc tatgtatcgt ggagaattgg ctatcagcaa gatacaatat
36541 gctaaaaatg aaagcttatg cgaattaaaa aatcaggcgg aaatttaccc tggtgctcta
36601 aagaagagct tgataaactt ttttttattt gaagcagagt tctctttaat gtttgtaaaa
36661 gcaaatgcgg gagcagagga taaatattat attgcaggcc atgtttttcg tattatttca

36721 tgcttaaatc aagtactatt tgcatgtaat aatgcttatt gcattaacga aaagaaagct
36781 ataaaactgc ttgaaacttt tgaatataaa cctaaaaaat atgccgaaag ggtaaatcat
36841 atttttgaag tactcggtct ttcacttttt gaatgctacg atatgaccga gaagctttat
36901 aaagaagtga aaaaaattgc aacggagata aataactttt taaacgaggg ggaattcaga
36961 tgaaagaaaa caaatatgat gataatatat tttttcaaaa atacagtcaa atgagtcgct
37021 cacagcaggg actagccggt gcaggagaat gggagacttt gaaaaagatg ctacctgatt
37081 ttaagggtaa gcgtgtgctt gatttaggat gcggctatgg atggcactgt atatatgcga
37141 tggaaaacgg tgcttcctct gtagtaggtg ttgatatttc tcataaaatg ctcgaagtag
37201 caaaaggaaa aacccatttt ccacagattg aatatgaatg ctgtgccata gaagatgtgg
37261 atttcccaga ggagagcttt gatgtaatac taagttcgct tgcgtttcat tatgtagcag
37321 actatgagaa tttaataaaa aagatatata ggatgctgaa ggctggtggc aatttagttt
37381 ttacagttga acatcctgtt tttactgctc atggaacaca agactggtat tataacgaaa
37441 aaggagaaat actgcatttc ccggtggaca attattatta tgagggcaaa cggacagcta
37501 tgtttttgga agaaaaggtt acaaaatatc atagaacact gaccacatat ctaaatacac
37561 tgctttcaaa tagttttata ataaatcaga ttgtggagcc acagccgcca gagaacatga
37621 tggatattcc ggggatggcg gatgaaatgc gacgcccaat gatgctgatt gtatcggcaa
37681 aaaagaagat gtaataatat agaaaaaata aacgaggagt atgtaaatga gatcagaaaa
37741 agaaatgatg gatttagtac tttctttagc agaacaggat gaacgtattc gaattgtgac
37801 ccttgagggg tcacgcgcaa atattaatat acctaaagat gaatttcagg attatgatat
37861 tacatatttt gtaagtgata tagaaccgtt tatatctaat gatgactggc ttaatcaatt
37921 tgggaatata ataatgatgc aaaagccgga ggatatggaa ttattcccac ctgaagaaaa
37981 gggattttcc tatcttatgc tatttgatga ttacaataaa attgatctta ccttattgcc
38041 cttggaagag ttagataatt acctaaaggg cgataaatta ataaaggttc taattgataa
38101 agattgtaga attaaaaggg acatagttcc gactgatata gattatcatg taagaaagcc
38161 aagcgcaagg gagtatgatg attgctgcaa tgaattttgg aatgtaacac cttatgttat
38221 taaaggattg tgccgtaagg aaattttatt tgctattgat cattttaatc agattgttcg
38281 ccatgagctg ctgagaatga tatcatggaa ggtcggcatc gaaacaggct ttaaattaag
38341 tgtaggcaag aactataagt ttattgaaag gtatatatcc gaggatttgt gggagaaact
38401 tttgtccacc taccggatgg attcctatga aaacatatgg gaagcattat ttctatgcca
38461 tcaattgttc agggcggtat ccggtgaggt ggcggaaagg cttcattatg cctatccgga
38521 gtatgatagg aatataacaa aatataccag ggacatgtat aaaaaataca ctggtaaaac
38581 cggctgcctg gatagcacat atgccgctga tatagaagag aggcgggaac agtgattaca
38641 gaaatgaaag cagggcacct gaaagatatc gataaaccca gcgaaccatt tgaggtgata
38701 ggtaagatta taccgaggta tgaaaacgag aattggacct ttacagaatt actctatgaa
38761 gcgccatatt taaaaagcta ccaagacgaa gaggatgaag aggatgagga ggcagattgc
38821 cttgaatata ttgacaatac tgataagata atatatcttt actaccaaga cgataaatgc
38881 gtcggaaaag ttaaactgcg aaaaaattgg aaccggtacg cttatataga agatatcgcc
38941 gtatgtaagg atttcagggg gcaaggcata ggcagcgcgc ttatcaatat atctatagaa
39001 tgggcaaagc ataaaaactt gcatggacta atgcttgaaa cccaggacaa taaccttata
39061 gcttgtaaat tctatcataa ttgtggtttc aaaatcggct ccgtcgatac tatgttatac
39121 gccaactttg aaaacaactt tgaaaaagct gttttctggt atttaaggtt ttagaatgca
39181 aggaacagtg aattggagtt cgtcttgtta taattagctt cttggggtat ctttaaatac
39241 tgtaggtaac ctctaaaaat tcataaaaaa ttgttgaaaa aatcgtcaga ttatggtata
39301 ataaagatac tataataaag gcggttgaaa tgaaatgcag atcaatcttt ttgcggaaga

39361 aacacaatta cagagattat ccgaactcgg agattcattg gaaaaactga atgttataga
39421 ctttgaatca ttcagacctt tgcttgaaaa ggtaaattgt aaagaacgta aaagcaaagc
39481 cggcagaccg tcgtatgatg ttgtaatgat gttcaaaata ctcgtattgc aacgattgtt
39541 taatttatct gatgatcaaa ctgaatacca gataactgat cgaatgagct ttcagcgttt
39601 tattggctta tcattgggag agcgtgtacc tgatgcaaaa acaatctggc tttttaagga
39661 taccattacg caaaaagaaa taattgagcc tttatttaat cagttcacag cgcaattgga
39721 atcgcaaggt attataacgc atacaggaac aattgttgat gcgacttttg tggatgcacc
39781 gcgtcagcgc aacactcgtg aggaaaacaa tcaaattaaa aatggcgaga tacccaaaga
39841 atgggagaaa aatacccata aacttgcgca aaaagataca gacgcccgtt ggacaaagaa
39901 aaatgatgaa actcattacg gatataaaaa tcatgtaaaa gtggatgccg acagtaagat
39961 tattacggat tacgcaacca catcagcaaa tatacatgac agcaatgaat tttcagaatt
40021 tctcggagaa gctgacaagg taatatacgc tgacagcgcc tatgtgggaa aagaaattcc
40081 gaaacatgta gaaaaccgca tatgtgaaaa aggatatcgc ggcaaaccgt tgactgatga
40141 acaaaaagaa agcaatcgaa gaaaatcaaa aattcgatgc agaattgagc atgtgtttgg
40201 atttattacc gtatctatgc atggtcttac tgtgcgttcc attggcataa agcgagcaga
40261 gtttaatata ggtcttacaa accttgttta taacttttgc agatatacta ttttaaaacg
40321 aaaagaggtg tacatgggat aactatgccc aaatgagaat aattctgaat atttagggct
40381 tatcctccct gaatatttag cgagaatcaa tggcaatcgg ctatttcttg ccttgatttt
40441 tgaaaaaata atcgaaatag caaaatatta tgattgggaa tcaattttta gaggttccct
40501 gtagaaaaga ggaaggaaat aataaatggc taaaatgaga atatcaccgg aattgaaaaa
40561 actgatcgaa aaataccgct gcgtaaaaga tacggaagga atgtctcctg ctaaggtata
40621 taagctggtg ggagaaaatg aaaacctata tttaaaaatg acggacagcc ggtataaagg
40681 gaccacctat gatgtggaac gggaaaagga catgatgcta tggctggaag gaaagctgcc
40741 tgttccaaag gtcctgcact ttgaacggca tgatggctgg agcaatctgc tcatgagtga
40801 ggccgatggc gtcctttgct cggaagagta tgaagatgaa caaagccctg aaaagattat
40861 cgagctgtat gcggagtgca tcaggctctt tcactccatc gacatatcgg attgtcccta
40921 tacgaatagc ttagacagcc gcttagccga attggattac ttactgaata acgatctggc
40981 cgatgtggat tgcgaaaact gggaagaaga cactccattt aaagatccgc gcgagctgta
41041 tgatttttta aagacggaaa agcccgaaga ggaacttgtc ttttcccacg gcgacctggg
41101 agacagcaac atctttgtga aagatggcaa agtaagtggc tttattgatc ttgggagaag
41161 cggcagggcg gacaagtggt atgacattgc cttctgcgtc cggtcgatca gggaggatat
41221 cggggaagaa cagtatgtcg agctattttt tgacttactg gggatcaagc ctgattggga
41281 gaaaataaaa tattatattt tactggatga attgttttag tacctagatt tagatgtcta
41341 aaaagcttta actacaagct ttttagacat ctaatctttt ctgaagtaca tccgcaactg
41401 tccatactct gatgttttat atcttttcta aaagttcgct agatagagtt ctatattaga
41461 gatgtaaaga gttttggtga agaggtttgt gagtatggct tttataacca gggtgcaatg
41521 agaagcacta acagcagcta gcttaagaac gcataggata tactttttat ggaagtcaca
41581 gaggaggaat aaaatttgaa caggattaac agagtaacct ctattcttat tcagctgcaa
41641 tcaaaaaaaa taattcctgc caaagaaatt gcacagcgat ttaatataag cttaaggaca
41701 gtttacagag atatccggac acttgaagaa gccggaatac caattggatc tgaggccgga
41761 aaaggatatt ttcttgtaga gggcttccta cttccgccgg taatgtttac agcggcggaa
41821 gtgggtgcat tgattacagc agggaaattt ttaaattgcc atggagatga atcatttata
41881 aaggattttg attcagctat gtataaaatc aaatctattt tgaagcatgg tgaaaaaaac
41941 tatgcacagg agttggagaa cagtattaac gtgtacagca catctggaca gaaaaataca

42001 ttggcggata acgttattgc agcaattcag actgcaatct gcaataaaag ggtaatatca
42061 attcaatatc ctgcatcagg agggcaagaa ccggaaagca gaatgataga gcctatatca
42121 ctggggtttt atgaacagaa ctggtactta atcggttttg caggctaaga aatgaatata
42181 ggaattttcg tgtggatcgg attaaaagca tttgcataga agaagagctc tgtcaaagcc
42241 atgatggctc tttggaacaa atattaaaac aaatgctttc atataaaaaa ctatataatg
42301 taattcttag agctgaaaaa ggggagacat acaattctat caagaacagg tattctctcg
42361 gcttccttga agaaaccgat ttgggaagta aaatggaaat cgaattccag acagattcct
42421 ttgagatatt gagcaagcag ctgattgaat atggatcagg cattgaaata gtccaacctg
42481 atgaattaaa atgcataacc cgtaaacacc tggcacaaat tacgaatcac tgcctaaact
42541 tgatttagag aaagcgattg gtctgttatg tattttttgt aaggtttcat atcccctgct
42601 gacatacagc tgtcagtgta ttatgatatt ctactatcaa aaaggaggat tgaaaaatat
42661 gaaaccaaag ataattatgc acacacaaat tagccttgat ggccgcatta aaggctttga
42721 taatccggaa gtttactacc aggttgccgg gggaattcat tcggatgcgg tgctgtttgg
42781 gtccaatact gtttttactg catttgaaaa atatcctgct gaaacagagg ctgattttga
42841 gaaaatcata acaagcccgg aggaccccag accaatagga gtcatacctg acagcagagg
42901 tatcttgaga agtcttcatt gcttaagaaa tcttggatat ctgaaggaaa ttgtcattct
42961 ggtatcaact gccactccga aggaatattt gaattatttg gaggaaaggc attacccata
43021 tattgtgtca ggaaatgatc atgtggacta tgaaaaagcc tttcagatcc tgcatgagca
43081 atatgggtgt aaatacatgc ggacagacag tggcggtgga ttaaccaaca aattattaga
43141 aaacggactt attgatgaaa tcagtcttgt gatatcacca tgttttgtag gtaataaaga
43201 aaaacaacta tttgataacc ttctgttatc tgaaaaagtt aatttggaat tgcagggaac
43261 agagaatgca gaacatggct gtctgtcatt gcgctatgct gtgaaaaata aggattagtg
43321 cagcatactt taaaaagtgt ctgttgggac agtgtcctaa aggaagcata agctcatggc
43381 tagcggcggt gttaaagtaa cggcactcgg ccccaatata tagttgatat gcccctaagc
43441 ctttgagggc aaggcggtca accgcgcctt ttccggtccg catacagtcg ttcaatgagg
43501 gtatcaagct cggcaatgcg gcggtgcttc tcggtggttc gtttcttcgc cgccgttttc
43561 ggctaattct aattgatccg aactgtaagg ataagcaatt atccggggct agttaagtat
43621 cttagagagg aagataaaga taagtaaacc aattttaaag ttgtaccgaa taaagccttc
43681 aagaggacaa catgcttttg aaggctttat tgcttacaat tacttggatt gtaacgcaaa
43741 agcaggcacc aaatcaatga aaaatagcct atccaggcga ctgtttcaga aatatgggct
43801 tccgccaagg ggataccagt ggggtgcgac atctttttg cccactcaag gcacgttgag
43861 tgtgttgtct tgatgtcgag ggtggagaaa tatgactaaa caaaatactg ataaataaag
43921 ggtttcctga cattgaactt ttcccatcgc tactttgccg aggatgttaa tgtcgggaaa
43981 cccttatttt tatttttaga ggaaatatat caactctcct taaagtgata gggttgagtt
44041 gatagattag ataacttcgt ggggttgtgg agatataaat gctaagataa taatgtacag
44101 agttgctaat atgtctatgt ataatataag gtaaaattta gcacagtatt aatggtgatt
44161 aatatgattt atatgataaa cattaatact gaaatatttt taagatcagt aaaagattta
44221 aataagctaa aacttttagt cgaggttaat aattgggata gacctaattt tagtgccata
44281 gctagagaat taggagtgga tagaagaaca gttaaaaagt attatgatgg agatataaag
44341 aaagttagaa aatctaaaaa atcaaagata gatgatttct atgatatcat ttcgagtttg
44401 ctttcagcag aaacggacca aatattttat tataaatctc atctctatag atatttggtt
44461 agagaaaaag gattagattg ttcaagaagt aattttaatt attatatttt aaaacatgaa
44521 aaatttgcag aatactttaa accaaacaaa aagaaagatg ctgtaaaatc tgaaacttcc
44581 tttggaaaac aagcacaatt tgattggaaa gaaaaattaa aattttcttt taaaagcgga

44641 gaagagttta tctttaatgt tggcagtttg atattatcag catctaggtt caaatattgg
44701 gcaatttgtc catcaacatc tcaagcatat ttatttgatt tcttaactaa tgtttttgaa
44761 gcattaggag gtgtgcccaa agaaatagtt atagataatg cttcaacaat gatggataaa
44821 gctagaacgg aacgttctga cggtaaagtt aatcctaaat ttcagcagtt tgctgatgac
44881 tttggattta atatcgtacc ttgcattaga gcaaggccaa atactaaagc aaaagttgaa
44941 aatccaatgc gagttattga tgaaataatg acttataatg gattgcttaa taatgaggaa
45001 gagttattcg agaagatgca agaaataaca aatgaagcta attcaagagt ctgtcaggca
45061 ataggtattc ctcctatatt agtatttaaa aaagagaaag aacatctttt accactacca
45121 aatgataaaa tatgttctta ttacaaaaat acaacaatac atgcgaaagt caattccaat
45181 ggactatttc gctataaggg aaatctatat tctgtccccg ttgattttat taataaaaat
45241 attgttgtta aagttattga taataaccta tatgtttatt atgaccataa attaataact
45301 ttacatacag tttctaataa aaaaattaat tatcatgaaa atcatcactt agcaatgatg
45361 agcttaactt ttagaaactc ggataaagaa gatgttaaaa actatgcagc taaacattta
45421 gaggagatga aaaagtttaa tgaacaatta tcaacagtta cagaagaact tacatgaact
45481 taaattaaca caaatcagcc tacaaataga tgactatatt aataaaataa atgagcataa
45541 actaagcatt gtagacgctt tgtgtgagtt aactagtaaa gaggttgaag ttaaaaattt
45601 taatgctgcg cacgctatgg ttaaagtagc agcattcccg cacctgaaag aattgaagga
45661 ctttgatttt gagtttcaac ctaaaataaa caaacaacaa tttttagatt tcgaaagctt
45721 acgtttttta gataataata gcaatattat attgataggc aatagtggcg tgggtaagac
45781 ccaccttgcc acgtcaattt gaattgctgc agcaaagaaa agagtaagta cctattttat
45841 aaaatgccaa gatttgatag agcaattaaa aaaagcttat attgaaaaca aacttgatga
45901 tagaataaaa cattttagta aatacaagct tcttataatt gatgaaattg gatatttgcc
45961 aattggagaa caagaagcaa aaatgttttt tcaacttata gatagacgtt atgaaaagaa
46021 aagcacaata gttactagta atattaatct atccgactgg aatcaaatat ttttagacaa
46081 tatgctagct agtgctattt tagacagact tgttgatcat tctactattg taaatattct
46141 cggaaactct tacaggacgg ctactgcatt atccaaaata tggtgttagt ataaagtagt
46201 ggacacatta aatcgaacta agtaaaataa tatttagtta agaaaggatg tgtccattat
46261 gggaaagggc agaagatacg atcaagaata taaagatatg atagtagaat tatttaagtc
46321 aggaatgagc ttagcagagc taagtagcga atatggcatt gcaaaatcaa caattaacgg
46381 ctgggttaaa gatgtaaaag aaattaaagt agatgaaaat gaagttatga cattaaagga
46441 agttaaggct ttaaaaaaag aaatggcaag aattaaagag gaaaatgaaa tattacacca
46501 aaggcgggct ttagcccaaa aagctatggc catatttgca acaagaaatt agatgaatta
46561 atagaattta taaatagcaa taaaaacaaa catgatatta agactatatg taccgttcta
46621 ggcgtagcca gaagcacata ttataaattt tttgataaaa ctaaatctgc aagagaatta
46681 gaaaatgagg aattaaaatc agctattaaa aggatatata aagagaataa aggcatatat
46741 ggtgctccta ggattcatca tatacttggt atagaaggat tcaatgtatc attgaagcga
46801 gttcaaagac gaatgactga acttggcctt tgtgcagtaa ctgtaaaaaa atacaaacct
46861 cattcaagta aaaaggtagc agaagattta gaagacgttt tgaaaagaga ttttactact
46921 acttctatta acgaaaaatg ggtaggagat attacatata tttatactat caaagatggt
46981 tggtgctatt tagcttcagt tctagactta tattctaaaa aaataattgg ctacgctttc
47041 gatcagagaa tgactaatga tttagttgtt aaagccttaa aaaatgctta ttacagccaa
47101 tttcccgata agaataagca attaatattt catagcgatt taggctctca atacactagc
47161 aatgatttaa gggagctatg caaagagttt aatattatac aatcgtttag taaaaaaggt
47221 tgtccatatg ataatgcttg tatagaatct tttcattcat taataaaaaa ggaagaaata

47281 tatagaaata cataccgtac cttcgaagaa gctaatatgg ctatttttaa atatattgaa
47341 ggttggtata acagaaaaag gctacactct tctattaatt acatgacttc agatcaatgt
47401 gaattattag ctagaagtgc agcataaaaa agtttgactt tttgtgtcca aaatattgac
47461 ataagaccaa ggtacccata cagaggattc tgatatagat atcggaattt attacaatca
47521 agaatcattt gacctgacag cgattaacca aattgctaca gagttggatg atgagaatag
47581 aaacaacctt gttgtacctc ccggagcgtg gggtgattgg atcaatggcg gcggatggtt
47641 agttatgaac gggtatcatg ttgatctgat tttacgtgat ataaaacggg tggaacaaat
47701 aatcaaagat acggagcaag gaattgttac tgcaaattat cagaccgggc atccccatgg
47761 ttatataagt gctatgtatc gtggagaatt ggctatcagc aagatacaat atgctaaaaa
47821 tgaaagctta tgcgaattaa aaaatcaggc ggaaatttac cctggtgctc taaagaagag
47881 cttgataaac tttttttat ttgaagcaga gttctcttta atgtttgtaa aagcaaatgc
47941 gggagcagag gataaatatt atattgcagg ccatgttttt cgtattattt catgcttaaa
48001 tcaagtacta tttgcatgta ataatgctta ttgcattaac gaaaagaaag ctataaaact
48061 gcttgaaact tttgaatata aacctaaaaa atatgccgaa agggtaaatc atatttttga
48121 agtactcggt ctttcacttt ttgaatgcta cgatatgacc gagaagcttt ataaagaagt
48181 gaaaaaaatt gcaacggaga taaataactt tttaaacgag ggggaattca gatgaaagaa
48241 aacaaatatg atgataatat attttttcaa aaatacagtc aaatgagtcg ctcacagcag
48301 ggactagccg gtgcaggaga atgggaaaca ttgagaaagc tgctgccgga ttttaaagat
48361 aagcgtgtgc ttgatttagg atgcggctat ggatggcact gtatttatgc gatggaacac
48421 ggagcgtctt ctgttgtagg tgttgatatt tctcataaaa tgctcgaggt agccaaagaa
48481 aaaacacatt ttccacaggt tgaatataaa tgctgtgcta tagaagatgt ggaattccca
48541 gaggagagtt ttgatgtaat attaagttca cttgcgtttc actatgtagc agattatgag
48601 attttagtaa aaaagatata tagaatactg aagtctggtg gtaagctagt ttttacggtt
48661 gaacatcctg tttttactgc ctatggaaca caagactggc attataacga aaaaggagaa
48721 atactgcatt ttccggtgga taattattat tatgagggca aacggacagc tgtgtttttg
48781 ggagaaaagg ttacaaaata tcatagaaca ctgaccacat atctaaatac actgctttca
48841 aatggtttta taataaatca tattgtggag ccgcagccgc cggaaaacat gatggatatt
48901 ccgggggatgc aggatgaaat gcgccgtccc atgatgctga ttgtatcggc gaacaaaaaa
48961 gtggatagat agaactaaaa cacctataag gtataaatgg aggtaattta tgtttaaaga
49021 aaaaattatt atagagatga aagaagtatt caaagaaatt ccttttggca tagagcatac
49081 tctcaaagtt ttgaaaaatg cagaagatat aatgaaagga gaaaatatcg gagaggaaga
49141 aaaagaattc atcagtatta ttgctatact acatgatatt ggtgctgttg aagcacaaaa
49201 aaaatacggt tctattgatg gtgtctatca agaaaaggaa ggaccagaag tagcgaaaga
49261 aatattaaaa aaggtaggct ataacaaaaa tattgataga atatgcttta taataggcaa
49321 ccatcatact ccatctaaaa ttgatggact tgattttcaa atacaatggg aagctgattt
49381 gcttgaaaat ttaacgatta tggataaaga aaaagaacag gaaaagataa aaaagtgtat
49441 agatgaaaac tttaaaacaa acacaggaaa aaggatagct t

//

DEFINITION Plasmid PLASMID_1

fasta_record 1..4724

/locus_tag="B77|SC|contig000002"

/label=B77|SC|contig000002

/note="B77|SC|contig000002"

/colour=5

CDS 10..1062

/color="4locus_tag=B77_03324note=Relaxase/Mobilisation+nuclease+domain,+protein+motif:Pfam:PF03432.8product=Relaxase/Mobilisation+nuclease+domain+[relaxase]'ID=B77_03324protein_id=gnl|SC|B77_03324'"
/codon_start=1
CDS 351..548
/locus_tag="B77_ORF80"
/color=8
/note="Type+II+CRISPR-associated+endonuclease+Cas1"
/product="Type+II+CRISPR-associated+endonuclease+Cas1+[endonuclease]"
/codon_start=1
CDS 653..1318
/locus_tag="B77_ORF42"
/color=1
/note="Cpp3+protein"
/product="Cpp3+protein"
/codon_start=1
CDS 1362..1595
/color=8locus_tag=B77_03325note=Sodium+ion-translocating+oxaloacetate+decarboxylase+beta+subunit+membrane+protein,+Cdd:COG1883+protein+motif:TIGRFAMs:TIGR01109+protein+motif:Pfam:PF03977.7product=Sodium+ion-translocating+oxaloacetate+decarboxylase+beta+subunit++membrane+protein+[transporter]'ID=B77_03325protein_id=gnl|SC|B77_03325'
/codon_start=1
CDS 1653..1796
/locus_tag="B77_03326"
/color=1
/locus_tag="B77_03326"
/note="hypothetical+protein"
/product="hypothetical+protein"
/protein_id="gnl|SC|B77_03326"
/codon_start=1
CDS 2025..3374
/locus_tag="B77_03327"
/color=10
/locus_tag="B77_03327"
/note="Replication+initiation+protein+RepA"
/product="Replication+initiation+protein+RepA+[replication]"
/protein_id="gnl|SC|B77_03327"
/codon_start=1
CDS complement(3532..3636)
/locus_tag="B77_ORF194"
/color=8

/note="Molybdenum+cofactor+biosynthesis+protein+MoaD"
/product="Molybdenum+cofactor+biosynthesis+protein+MoaD"
/gene="moaD"
/codon_start=1
CDS 3800..4195
/locus_tag="B77_ORF67"
/color=9
/note="Transposon+protein+TnpV"
/product="Transposon+protein+TnpV"
/codon_start=1
CDS complement(3851..3976)
/locus_tag="B77_ORF147"
/color=9
/note="Transposon+protein+TnpV"
/product="Transposon+protein+TnpV"
/codon_start=1
CDS 4396..4716

/color=4locus_tag=B77_03329note=Plasmid+mobilisation+relaxosome+protein+MobC,+protein+motif:Pfam:PF05713.5product=Plasmid+mobilisation+relaxosome+protein+MobC+[relaxosome]'ID=B77_03329gene=mobC'protein_id=gnl|SC|B77_03329
/codon_start=1
CDS 4638..4724
/locus_tag="B77_ORF107"
/color=4
/note="Relaxase/mobilization+nuclease+domain-containing+protein"
/product="Relaxase/mobilization+nuclease+domain-containing+protein+%28partial%29+[relaxase]"
/codon_start=1
BASE COUNT 1500 a 823 c 1340 g 1061 t
ORIGIN
1 atagattatg tgacaaaaaa agaaaagaca gaggagaagc ttgtcagcgg tctgcattgt
61 gagccggaga cggtcaagga agaaatgcaa gccacaaagg agctgtgggg caagacggat
121 ggaagaacct ataagcatta tgtgcaatcc taccatgagg acgaggaaat aaccccggag
181 caggctcaca agaacgctgt cgagctggca gagcatacaa aggcatggaa agggcatgaa
241 gttctgatag ccacgcatat agacaagggg catatacaca cgcattttat tgtcaattcc
301 gtaaattatg aggacggtca taagctccaa tggagtaagc acgaccttaa agacctaaag
361 gaacggtgca acgagcagag ccgggaacag ggcttgcatg tgccggagaa aggaaagacc
421 tttgctggag aagtccggga agaaacggtg gcatggagca aagacaccta ccagcttttg
481 aaacaggcag agcagggaaa ggtcaaaagc tatgtgcagg atattgccct tgcggtgctg
541 gactgtaagg aaacagcgac cagccgggaa aactttatcc ggatgatgaa tgaaagaggg
601 tacggcgtag actggcagga cagccacaag tacatcacat ataccgactt agccagggaa
661 caggcaggag aaaaggcttg taaaatcagg gataacaagc tggaaaaata ctacaatatg
721 gattttggaa aggagagtat ggagcatgag tttgagagaa atgcacgaac agcagaagca

781 gagcagtcca agagagcagc ttcaagagtt aaaccgacag agccggacag agctggaaaa
841 cagtctgctc aaagaagcgt tggcgatgtc gagcgagaac tgcgaggaat tgatgaagca
901 gtcaaatcaa gaacaagtaa gggcagagca gaacaggcag agagacgcag agcagaacag
961 gaagctcatc agcgagctga agcagagcgt agagctgctg aagaacagca acgaattgct
1021 tcaagacgct atatcgggcg agattgggag cctgaaagat gaggttaagg aaaacaccgt
1081 ccaagaggtt agaaaggcgt tacaggcgaa tatagaggct atacagaggg ctacaaagct
1141 tcttgagaag cagtctgata cacttacaag cgtgatgaag cagaagatcc gggagctgga
1201 ggaaagcaag aacagctttt tcagatatga gggcttaaag ctgtacctgt tttggggcgg
1261 tatggtctgt aatattttag tgtttcttat gctgttatat ggtatgcttg gtaagtaaaa
1321 agaaatcgac aaatcggaag gttgagaaag gagattagtc aatgaaaaaa atattaggaa
1381 tcgttgctgg tattgttgga ataatatctg ttattgtggg tgttgcattg aagatgaata
1441 ataatgctat tgctataatc ggaggtgcag atggtcctac ctcggtattt gtcgcaggaa
1501 agttgaatag tgttcttgtc agcatcttgc ttattgcgat aggtgttgtg atgttgatag
1561 tagcgataat tttttatttg aagagaaaaa ggtaaaataa tgtagtgata agataagcgg
1621 tcaaggcgag agggagtgag gtgtaaaaga ctatggagaa agaaaaaaga gagcaggaga
1681 gctgcacaag ctgcgcttat aggaaagatt ggtattgtcc cgttgccgga gaccgtgtaa
1741 agagagaggg gcactgttct gaatacgaga aaagtgagat tgcgaaggaa aagtagcagt
1801 gctataatgg cagggaaagg gagagaaaac cagccctttt ttaagaggtc aaaaatgtgc
1861 aataaactgt attgataata taagccccct ataggtgttt ttaggggtgg ggaaagggag
1921 agaaaaccag cccttttta agaggtcaaa aatgtgcaat aaactgtatt gataatataa
1981 gccccctata ggtgttttgg aggggagaaa aaggtggtgg gaatatggca aaaaaaccga
2041 ctgaaaagca gttgtataag ctgaaaaatg aatggctggg gcagttcttt gaggaagtga
2101 agccgaaaga gttttatcgg gctgtttttc cagagggaag ttttgagaga aaggggcatt
2161 ttgaggacgg aaaggcaaat ggcattataa ccatagttga gaatgataag gcgaagaatc
2221 ggattgtatt tgatgatttg tcggtcattg atgaggtcaa gggggcagag tttgctgtca
2281 tgtcccctgt ggcgtattcc ggcagaaatc ggacggcagc caatgcaagg tggctgtatg
2341 ggatagcgat agaccttgac ggggtggaga tgccgcagct ccgtgatgtg ttccatcaaa
2401 tgaaccatga tattatcccg aaatgcactt actgtattaa cagcggtcac gggctgcatt
2461 tatattatct gctggaaaag ccggttccct tgtataagca tttgcaggat aagcttcgag
2521 agttcaagta tgagctgatt gcgaaagtct ggaaccgtta tacaagcact ttcacagaac
2581 gggaacaggt tcagtatcag gggatttttc agggcttccg tatggttgga acgcagtcaa
2641 agctgggaaa gagataccсg gtaaaggctt ttgaaacggg cgagagagtg acgattgagt
2701 atttgaacgg gtatcttatg gataagagca aggcggtcac ggattttcac tataagagcg
2761 atttgagtct tgccgaagcc cggaagaagt acccggaatg gtatgaaaag cgaattgtgc
2821 agggggagcg ccgtgagcgc tggcatgtca aaagggactt atatgattgg tggctgcgga
2881 aaataaaaga gggggcttct gtagggcaca gatacagctg cctttgcgtt ttggcgtcct
2941 atgccgttaa gtgcgatatt ccggaagatg agctttttac agacgctttt tccctattgc
3001 agttccttga tgatatgtca gatgatgagc ataaccgctt taccaagcgt gatatactgg
3061 acgccatgca gttttatcaa gagaattatg tcacatacag tcgcagggaa gccgaaagag
3121 tatcggctat tgctatgcca gcaagcaaac ggaacggaag aaaacaggct gaacatttag
3181 agcgtgcaag gttggtacaa acattggatt atccaaatgg agcatggaga aataaagagg
3241 gtagaccaaa aggcagcgga gaaaagtcga agattgtcga agaatggaaa cagcagcatc
3301 cggacggaaa gaagatagat tgtgagagag aaactgggct tagccggcat actgtattga
3361 agtggtggaa gtagggtata cttgaaatgc cgaaaggcat atgctataat accagtaggc

3421 gaaaatgaca gaagtgctca tgtggcacag caaaaagccc cggaggttgc gactccgggg
3481 cttttctatt ccgttctggg tgggtggctt aatcccaagg ctggctacca gttattcttc
3541 tccgtccaac catttgcaga tgtagtaggc aactacactt gccaaaacag agagaaaaaa
3601 tgacagaagt tcactcatgc ggttcacccc ctttctctac ctgtctgggg gcggtagcgt
3661 cgttcattat atcacagggg cagaaatgcc acaagaagcc atttcccctt gtttattatc
3721 aaagccataa ggaaaagagt gtcaagggcg gtcatagacc gcagcgttta cccttgatac
3781 tcttttttgt tggctataat gccggagggg aaaacggctg tccgtccgtt tcgttacttc
3841 cccctgtgct tcattgtccg ttccggcgga agaaagtaaa gggtggtctg cgaccatttt
3901 ttatctgctg tactcgctga tcggcaagcc gaagctcata ctgcagataa aaaacaagcc
3961 cttgactgtc tccctcctac actttcaaag tcgcccaaga ggaagtaagc tccactatat
4021 tcttggcaga atatacgcac gaatatatcc gcttgatata gggtgtctat catgctataa
4081 tatgtgtagc ggatatattg caagttccct aaagtgttac cactttagaa cttgtgaggg
4141 tggcaagccc cccttcaatc cccctttttaa acgagccgga aggctcataa tatgacggtg
4201 gtagcaccga agaaagagag gtcaaaggca tgaaaaaaga tatggataaa attcagtttg
4261 agtacagata cgaggtgcag gagctgatga aagtaattga taaatatgta aaacagaatc
4321 cggcagaaaa ggaaaataag acgctggagc gtttctttga cctacttgat gtcatggata
4381 tggagtggta agggtatgaa caagacaaga ccaaagcagc tatcattccg ggtaagtgaa
4441 gaagaatacc agcagctaca ggagaagatt tcagagagtg gaaagaacca gcaggaatat
4501 atcctttcct gtgtgctgga gaagcaaatc gtgaatacgg acggtatcaa ggaacttatc
4561 ccggaactga aacggatagg gaacaacctc aaccagatag caaagcggtg taatgagggg
4621 ggaatgttgc cgagtgaagc ggaagtgcgg aagcatggag aggagctgaa caaagtatgg
4681 cagtcattaa ggcggtatct tcaaaggcgg gcatagggca cgcc
//
DEFINITION Plasmid PLASMID_2
fasta_record 1..70011
/ID=F51|SC|contig000002
/label=F51|SC|contig000002
/note="F51|SC|contig000002"
/colour=5
CDS 773..979
/color=7
/locus_tag="F51_03060"
/note="Helix-turn-helix domain-containing protein, protein motif:CLUSTERS:PRK09943 protein motif:Pfam:PF12844.1"
/product="Helix-turn-helix domain-containing protein [transcriptional regulator]"
/"ID=F51_03060
/protein_id="gnl|SC|F51_03060"""
/codon_start=1
CDS 1076..1450
/ID=F51_03061
/color=7
/locus_tag="F51_03061"
/note="Helix-turn-helix domain-containing protein"

/product="Helix-turn-helix domain-containing protein [transcriptional regulator]"
/protein_id="gnl|SC|F51_03061"
/codon_start=1
CDS 1512..2615
/color=1
/locus_tag="F51_03062"
/note="CAP domain-containing protein  Cysteine-rich secretory protein family, protein motif:Pfam:PF00188.20"
/product="CAP domain-containing protein  Cysteine-rich secretory protein family"
/"ID=F51_03062
/protein_id="gnl|SC|F51_03062""""
/codon_start=1
CDS 2685..3059
/color=8
/locus_tag="F51_03063"
/note="type II toxin-antitoxin system PemK/MazF family toxin, similar to AA sequence:UniProtKB:Q7A4G9 protein motif:Pfam:PF02452.11"
/product="type II toxin-antitoxin system PemK/MazF family toxin MazF [toxin-antitoxin]"
/"ID=F51_03063
/gene="mazF""""
/protein_id="gnl|SC|F51_03063"
/codon_start=1
CDS 3199..6762
/color=8
/locus_tag="F51_03064"
/note="Uncharacterised surface anchored protein, similar to AA sequence:UniProtKB:P18481 protein motif:Cdd:COG4932 protein motif:Pfam:PF05738.7"
/product="Uncharacterised surface anchored protein"
/"ID=F51_03064
/protein_id="gnl|SC|F51_03064""""
/codon_start=1
CDS 6833..6961
/ID=F51_03065
/color=1
/locus_tag="F51_03065"
/note="hypothetical protein"
/product="hypothetical protein"
/protein_id="gnl|SC|F51_03065"
/codon_start=1

CDS 6977..7234
/ID=F51_03066
/color=1
/locus_tag="F51_03066"
/note="hypothetical protein"
/product="hypothetical protein"
/protein_id="gnl|SC|F51_03066"
/codon_start=1
CDS 7237..7623
/ID=F51_03067
/color=1
/locus_tag="F51_03067"
/note="hypothetical protein"
/product="hypothetical protein"
/protein_id="gnl|SC|F51_03067"
/codon_start=1
CDS 7625..8269
/color=8
/locus_tag="F51_03068"
/note="Class D Sortase  cysteine transpepidase , protein
motif:Cdd:COG3764 protein motif:TIGRFAMs:TIGR01076 protein
motif:Pfam:PF04203.7"
/product="Class D Sortase  cysteine transpepidase
[transpepidase]"
/"ID=F51_03068
/protein_id="gnl|SC|F51_03068"""
/codon_start=1
CDS 8280..8582
/ID=F51_03069
/color=1
/locus_tag="F51_03069"
/note="hypothetical protein"
/product="hypothetical protein"
/protein_id="gnl|SC|F51_03069"
/codon_start=1
CDS 8586..8804
/ID=F51_03070
/color=1
/locus_tag="F51_03070"
/note="DUF4236 domain-containing protein"
/product="DUF4236 domain-containing protein"
/protein_id="gnl|SC|F51_03070"
/codon_start=1
CDS 8868..9632

/color=8
/locus_tag="F51_03071"
/eC_number=2.1.1.72
/note="Modification methylase DpnIIA DNA adenine
methylase, similar to AA sequence:UniProtKB:P04043 protein
motif:CLUSTERS:PRK10904 protein motif:Cdd:COG0338 protein
motif:TIGRFAMs:TIGR00571 protein motif:Pfam:PF02086.9"
/product="Modification methylase DpnIIA DNA adenine
methylase [methyltransferase]"
/"ID=F51_03071
/gene="dpnM"""
/protein_id="gnl|SC|F51_03071"
/codon_start=1
CDS 9659..9943
/ID=F51_03072
/color=1
/locus_tag="F51_03072"
/note="hypothetical protein"
/product="hypothetical protein"
/protein_id="gnl|SC|F51_03072"
/codon_start=1
CDS 9944..10255
/ID=F51_03073
/color=4
/locus_tag="F51_03073"
/note="TcpK chain A:2.60"
/product="TcpK chain A"
/protein_id="gnl|SC|F51_03073"
/gene="tcpK"
/codon_start=1
CDS 10631..11431
/color=9
/locus_tag="F51_03074"
/note="Tyrosine recombinase, protein motif:Cdd:COG4974
protein motif:TIGRFAMs:TIGR02249 protein
motif:Pfam:PF00589.16"
/product="Tyrosine recombinase [recombinase]"
/"ID=F51_03074
/protein_id="gnl|SC|F51_03074"""
/codon_start=1
CDS 11493..13082
/color=4
/locus_tag="F51_03075"
/note="DNA translocase FtsK DNA segregation ATPase

FtsK/SpoIIIE and related proteins , similar to AA
sequence:UniProtKB:P64165 protein motif:CLUSTERS:PRK10263
protein motif:Cdd:COG1674 protein motif:TIGRFAMs:TIGR03928
protein motif:Pfam:PF01580.12"
/product="DNA translocase FtsK  DNA segregation ATPase
FtsK/SpoIIIE and related proteins [ATPase  coupling
protein]"
/"ID=F51_03075
/gene="ftsK"""
/protein_id="gnl|SC|F51_03075"
/codon_start=1
CDS 13096..13266
/ID=F51_03076
/color=1
/locus_tag="F51_03076"
/note="hypothetical protein"
/product="hypothetical protein"
/protein_id="gnl|SC|F51_03076"
/codon_start=1
CDS 13259..14338
/color=4
/locus_tag="F51_03077"
/note="Conjugative transposon protein TcpC, protein
motif:Pfam:PF12642.1"
/product="Conjugative transposon protein TcpC [mating pair
formation]"
/"ID=F51_03077
/gene="tcpC"""
/protein_id="gnl|SC|F51_03077"
/codon_start=1
CDS 14350..14685
/color=4
/locus_tag="F51_03078"
/note="Conjugative transposon protein TcpD, ab initio
prediction:Prodigal:2.60"
/product="Conjugative transposon protein TcpD [mating pair
formation]"
/"ID=F51_03078
/gene="tcpD"""
/protein_id="gnl|SC|F51_03078"
/codon_start=1
CDS 14697..15065
/color=4
/locus_tag="F51_03079"

/note="Conjugative transposon membrane protein TcpE,
protein motif:Pfam:PF12648.1"
/product="Conjugative transposon membrane protein TcpE
[mating pair formation]"
/"ID=F51_03079
/gene="tcpE"""
/protein_id="gnl|SC|F51_03079"
/codon_start=1
CDS 15123..15290
/ID=F51_03080
/color=1
/locus_tag="F51_03080"
/note="hypothetical protein"
/product="hypothetical protein"
/protein_id="gnl|SC|F51_03080"
/codon_start=1
CDS 15341..17863
/color=4
/locus_tag="F51_03081"
/note="ATPase conjugation protein TcpF, protein
motif:Cdd:COG3451 protein motif:Pfam:PF12846.1"
/product="ATPase conjugation protein TcpF [ATPase mating
pair formation]"
/"ID=F51_03081
/gene="tcpF"""
/protein_id="gnl|SC|F51_03081"
/codon_start=1
CDS 17863..18867
/color=4
/locus_tag="F51_03082"
/note="Conjugation-related peptidoglycan hydrolase TcpG,
similar to AA sequence:UniProtKB:P40767 protein
motif:CLUSTERS:PRK05684 protein motif:Cdd:COG3951 protein
motif:TIGRFAMs:TIGR02541 protein motif:Pfam:PF01832.14"
/product="Conjugation-related peptidoglycan hydrolase TcpG
[pepidase]"
/"ID=F51_03082
/gene="tcpG"""
/protein_id="gnl|SC|F51_03082"
/codon_start=1
CDS 18970..21468
/color=4
/locus_tag="F51_03083"
/note="Conjugation pore-forming membrane protein TcpH,

protein motif:Cdd:COG5027"
/product="Conjugation pore-forming membrane protein TcpH
[mating pair formation]"
/"ID=F51_03083
/gene="tcpH"""
/protein_id="gnl|SC|F51_03083"
/codon_start=1
CDS 21471..21965
/color=4
/locus_tag="F51_03084"
/note="Predicted membrane-bound metal-dependent hydrolase
TcpI, similar to AA sequence:UniProtKB:P64481 protein
motif:CLUSTERS:PRK11648 protein motif:Pfam:PF04307.8"
/product="Predicted membrane-bound metal-dependent
hydrolase TcpI [regulator]"
/"ID=F51_03084
/gene="tcpI"""
/protein_id="gnl|SC|F51_03084"
/codon_start=1
CDS 21974..22732
/ID=F51_03085
/color=4
/locus_tag="F51_03085"
/note="Type 4 prepilin-like proteins leader
peptide-processing enzyme TcpJ"
/product="Type 4 prepilin-like proteins leader
peptide-processing enzyme TcpJ
[pepidase/methyltransferase]"
/gene="tcpJ"
/protein_id="gnl|SC|F51_03085"
/codon_start=1
CDS 23681..23986
/ID=F51_03086
/color=1
/locus_tag="F51_03086"
/note="hypothetical protein"
/product="hypothetical protein"
/protein_id="gnl|SC|F51_03086"
/codon_start=1
CDS 24001..25020
/color=8
/locus_tag="F51_03087"
/eC_number=2.1.1.37
/note="DNA cytosine-5--methyltransferase HhaI, similar to

AA sequence:UniProtKB:P05102 protein
motif:CLUSTERS:PRK10458 protein motif:TIGRFAMs:TIGR00675
protein motif:Pfam:PF00145.11"
/product="DNA %28cytosine-5-%29-methyltransferase HhaI
[methyltransferase]"
/"ID=F51_03087
/gene="hhaIM"""
/protein_id="gnl|SC|F51_03087"
/codon_start=1
CDS 25037..25210
/ID=F51_03088
/color=1
/locus_tag="F51_03088"
/note="hypothetical protein"
/product="hypothetical protein"
/protein_id="gnl|SC|F51_03088"
/codon_start=1
CDS 25207..26229
/ID=F51_03089
/color=1
/locus_tag="F51_03089"
/note="DUF4417 domain-containing protein"
/product="DUF4417 domain-containing protein"
/protein_id="gnl|SC|F51_03089"
/codon_start=1
CDS 26324..27730
/color=8
/locus_tag="F51_03090"
/note="Putative amidoligase enzyme  SWIM zinc finger
domain protein, protein motif:Pfam:PF12224.2"
/product="Putative amidoligase enzyme  SWIM zinc finger
domain protein"
/"ID=F51_03090
/protein_id="gnl|SC|F51_03090"""
/codon_start=1
CDS 28318..28719
/ID=F51_03091
/color=10
/locus_tag="F51_03091"
/note="Putative C-terminal domain of type IA
topoisomerase"
/product="Putative C-terminal domain of type IA
topoisomerase"
/protein_id="gnl|SC|F51_03091"

/codon_start=1
CDS 28817..29128
/ID=F51_03092
/color=1
/locus_tag="F51_03092"
/note="hypothetical protein"
/product="hypothetical protein"
/protein_id="gnl|SC|F51_03092"
/codon_start=1
CDS complement(29440..29673)
/ID=F51_03093
/color=1
/locus_tag="F51_03093"
/note="hypothetical protein"
/product="hypothetical protein"
/protein_id="gnl|SC|F51_03093"
/codon_start=1
CDS complement(29698..30600)
/ID=F51_03094
/color=1
/locus_tag="F51_03094"
/note="hypothetical protein"
/product="hypothetical protein"
/protein_id="gnl|SC|F51_03094"
/codon_start=1
CDS 30730..31137
/ID=F51_03095
/color=1
/locus_tag="F51_03095"
/note="hypothetical protein"
/product="hypothetical protein"
/protein_id="gnl|SC|F51_03095"
/codon_start=1
CDS 31201..32058
/color=9
/locus_tag="F51_03096"
/note="Rpn family recombination-promoting
nuclease/putative transposase, protein
motif:TIGRFAMs:TIGR01784 protein motif:Pfam:PF12784.1"
/product="Rpn family recombination-promoting
nuclease/putative transposase [DDE transosase]"
/"ID=F51_03096
/protein_id="gnl|SC|F51_03096"""
/codon_start=1

CDS 32075..32164
/ID=F51_03097
/color=1
/locus_tag="F51_03097"
/note="hypothetical protein"
/product="hypothetical protein"
/protein_id="gnl|SC|F51_03097"
/codon_start=1
CDS 32813..34075
/color=2
/locus_tag="F51_03098"
/note="Tetracycline efflux MFS transporter TetA , protein motif:CLUSTERS:PRK11646 protein motif:Cdd:COG2814 protein motif:Pfam:PF07690.10"
/product="Tetracycline efflux MFS transporter TetA%28P%29 [antibiotic resistance]"
/"ID=F51_03098
/protein_id="gnl|SC|F51_03098"""
/gene="tetA"
/codon_start=1
CDS 34059..36017
/color=2
/locus_tag="F51_03099"
/note="Tetracycline resistance protein tetM from transposon Tn916, similar to AA sequence:UniProtKB:P21598 protein motif:CLUSTERS:PRK13351 protein motif:Cdd:COG0050 protein motif:TIGRFAMs:TIGR00484 protein motif:Pfam:PF00009.21"
/product="Tetracycline resistance protein tetM from transposon Tn916 [antibiotic resistance]"
/"ID=F51_03099
/gene="tetM"""
/protein_id="gnl|SC|F51_03099"
/codon_start=1
CDS 36209..36487
/ID=F51_03100
/color=1
/locus_tag="F51_03100"
/note="YcxB family protein"
/product="YcxB family protein"
/protein_id="gnl|SC|F51_03100"
/codon_start=1
CDS 36938..37807
/color=7

/locus_tag="F51_03101"
/note="AraC family transcriptional regulator
Helix-turn-helix domain-contining protein, similar to AA
sequence:UniProtKB:P28816 protein motif:CLUSTERS:PRK10219
protein motif:Cdd:COG3449 protein motif:TIGRFAMs:TIGR04094
protein motif:Pfam:PF12833.1"
/product="AraC family transcriptional regulator
Helix-turn-helix domain-contining protein [transcriptional
regulator]"
/"ID=F51_03101
/protein_id="gnl|SC|F51_03101"""
/codon_start=1
CDS 38355..38507
/ID=F51_03102
/color=8
/locus_tag="F51_03102"
/note="Cytochrome c551 product=Cytochrome c551 [eletron
transfer]"
/protein_id="gnl|SC|F51_03102"
/codon_start=1
CDS 39587..40321
/color=9
/locus_tag="F51_03103"
/note="Serine recombinase, similar to AA
sequence:UniProtKB:P20384 protein motif:CLUSTERS:PRK13413
protein motif:Pfam:PF00239.15"
/product="Serine recombinase [recombinase]"
/"ID=F51_03103
/protein_id="gnl|SC|F51_03103"""
/codon_start=1
CDS 40342..40449
/ID=F51_03104
/color=8
/locus_tag="F51_03104"
/note="Signal pepidase"
/product="Signal pepidase"
/protein_id="gnl|SC|F51_03104"
/codon_start=1
CDS complement(40579..40758)
/ID=F51_03105
/color=1
/locus_tag="F51_03105"
/note="hypothetical protein"
/product="hypothetical protein"

/protein_id="gnl|SC|F51_03105"
/codon_start=1
CDS 40950..41747
/ID=F51_03106
/color=8
/locus_tag="F51_03106"
/note="Beta2 toxin"
/product="Beta2 toxin [virulence]"
/protein_id="gnl|SC|F51_03106"
/codon_start=1
CDS complement(41758..42399)
/ID=F51_03107
/color=1
/locus_tag="F51_03107"
/note="hypothetical protein"
/product="hypothetical protein"
/protein_id="gnl|SC|F51_03107"
/codon_start=1
CDS complement(42403..42726)
/color=7
/locus_tag="F51_03108"
/note="PadR family transcriptional regulator, protein
motif:CLUSTERS:PRK09416 protein motif:TIGRFAMs:TIGR03433
protein motif:Pfam:PF03551.8"
/product="PadR family transcriptional regulator
[transcriptional regulator]"
/"ID=F51_03108
/protein_id="gnl|SC|F51_03108""""
/codon_start=1
CDS 42949..43419
/ID=F51_03109
/color=1
/locus_tag="F51_03109"
/note="hypothetical protein"
/product="hypothetical protein"
/protein_id="gnl|SC|F51_03109"
/codon_start=1
CDS 43486..44016
/color=7
/locus_tag="F51_03110"
/eC_number=3.4.21.88
/note="LexA repressor  SOS-response repressor and protease
LexA , similar to AA sequence:UniProtKB:P31080 protein
motif:CLUSTERS:PRK00215 protein motif:TIGRFAMs:TIGR00498

protein motif:Pfam:PF01726.10"
/product="LexA repressor  SOS-response repressor and protease LexA [transcriptional regulator]"
/"ID=F51_03110
/gene="lexA"""
/protein_id="gnl|SC|F51_03110"
/codon_start=1
CDS 44302..44529
/ID=F51_03111
/color=1
/locus_tag="F51_03111"
/note="hypothetical protein"
/product="hypothetical protein"
/protein_id="gnl|SC|F51_03111"
/codon_start=1
CDS 44595..45089
/ID=F51_03112
/color=7
/locus_tag="F51_03112"
/note="DNA-binding protein  Helix-turn-helix domain-containing protein"
/product="DNA-binding protein  Helix-turn-helix domain-containing protein [transcriptional regulator]"
/protein_id="gnl|SC|F51_03112"
/codon_start=1
CDS 45220..45309
/ID=F51_03113
/color=9
/locus_tag="F51_03113"
/note="IS256-like element ISCpe3 family transposase"
/product="IS256-like element ISCpe3 family transposase [DDE recombinase]"
/protein_id="gnl|SC|F51_03113"
/codon_start=1
CDS 45473..45769
/color=8
/locus_tag="F51_03114"
/note="Histidinol-phosphatase  PHP domain-containing protein, protein motif:CLUSTERS:PRK07328 protein motif:Cdd:COG1387 protein motif:TIGRFAMs:TIGR01856"
/product="Histidinol-phosphatase  PHP domain-containing protein [phosphatase]"
/"ID=F51_03114
/protein_id="gnl|SC|F51_03114""""

/codon_start=1
CDS 45899..46255
/color=10
/locus_tag="F51_03115"
/note="Replication initiation protein RepA, protein motif:CLUSTERS:PRK07328 protein motif:Cdd:COG1387 protein motif:TIGRFAMs:TIGR01856"
/product="Replication initiation protein RepA [replication]"
/"ID=F51_03115
/protein_id="gnl|SC|F51_03115"""
/gene="repA"
/codon_start=1
CDS 46826..47104
/color=8
/locus_tag="F51_03116"
/note="Acyl carrier protein, protein motif:CLUSTERS:PRK07639"
/product="Acyl carrier protein"
/"ID=F51_03116
/protein_id="gnl|SC|F51_03116"""
/gene="acpP"
/codon_start=1
CDS 47116..49683
/color=8
/locus_tag="F51_03117"
/note="Alpha/beta hydrolase, protein motif:Pfam:PF00561.1"
/product="Alpha/beta hydrolase [hydrolase]"
/"ID=F51_03117
/protein_id="gnl|SC|F51_03117"""
/codon_start=1
CDS 49716..50666
/ID=F51_03118
/color=1
/locus_tag="F51_03118"
/note="hypothetical protein"
/product="hypothetical protein"
/protein_id="gnl|SC|F51_03118"
/codon_start=1
CDS 50898..51485
/ID=F51_03119
/color=1
/locus_tag="F51_03119"
/note="hypothetical protein"

/product="hypothetical protein"
/protein_id="gnl|SC|F51_03119"
/codon_start=1
CDS 51502..52305
/color=8
/locus_tag="F51_03120"
/note="ABC-2 type transport system permease protein,
protein motif:Cdd:COG4587 protein motif:Pfam:PF06182.5"
/product="ABC-2 type transport system permease protein
[transporter]"
/"ID=F51_03120
/protein_id="gnl|SC|F51_03120"""
/codon_start=1
CDS 52307..53101
/color=8
/locus_tag="F51_03121"
/note="ABC-2 type transport system permease protein,
protein motif:Cdd:COG4587 protein motif:Pfam:PF06182.5"
/product="ABC-2 type transport system permease protein
[transporter]"
/"ID=F51_03121
/protein_id="gnl|SC|F51_03121"""
/codon_start=1
CDS 53104..54090
/color=8
/locus_tag="F51_03122"
/note="ABC-2 type transport system ATP-binding protein,
similar to AA sequence:UniProtKB:O07190 protein
motif:CLUSTERS:PRK13636 protein motif:Cdd:COG4586 protein
motif:TIGRFAMs:TIGR01188 protein motif:Pfam:PF00005.21-"
/product="ABC-2 type transport system ATP-binding protein
[transporter]"
/"ID=F51_03122
/protein_id="gnl|SC|F51_03122"""
/codon_start=1
CDS 54096..55139
/color=1
/locus_tag="F51_03123"
/note="hypothetical protein, ab initio
prediction:Prodigal:2.60"
/product="hypothetical protein"
/"ID=F51_03123
/protein_id="gnl|SC|F51_03123"""
/codon_start=1

CDS 55136..56626
/color=8
/locus_tag="F51_03124"
/eC_number=6.2.1.3
/note="Long-chain-fatty-acid--CoA ligase, similar to AA sequence:UniProtKB:P69451 protein motif:CLUSTERS:PRK07656 protein motif:Cdd:COG4340 protein motif:TIGRFAMs:TIGR01923 protein motif:Pfam:PF00501.22"
/product="Long-chain-fatty-acid--CoA ligase [ligase]"
/"ID=F51_03124
/gene="fadD"""
/protein_id="gnl|SC|F51_03124"
/codon_start=1
CDS 56619..56864
/color=8
/locus_tag="F51_03125"
/note="Acyl carrier protein, similar to AA sequence:UniProtKB:P80918 protein motif:CLUSTERS:PRK00982 protein motif:Cdd:COG0236 protein motif:TIGRFAMs:TIGR00517 protein motif:Pfam:PF00550.19"
/product="Acyl carrier protein"
/"ID=F51_03125
/gene="acpP"""
/protein_id="gnl|SC|F51_03125"
/codon_start=1
CDS 56883..58142
/color=8
/locus_tag="F51_03126"
/note="phenylacetate-CoA ligase family protein, protein motif:TIGRFAMs:TIGR02155"
/product="phenylacetate-CoA ligase family protein"
/"ID=F51_03126
/protein_id="gnl|SC|F51_03126"""
/codon_start=1
CDS 58221..58841
/color=8
/locus_tag="F51_03127"
/eC_number=3.1.3.5
/note="5'-nucleotidase YjjG HAD superfamily HAD-IA family hydrolase, similar to AA sequence:UniProtKB:P0A8Y1 protein motif:CLUSTERS:PRK09449 protein motif:Cdd:COG5610 protein motif:TIGRFAMs:TIGR02253 protein motif:Pfam:PF00702.1"
/product="5'-nucleotidase YjjG HAD superfamily%29 HAD-IA family hydrolase [hydrolase]"

/"ID=F51_03127
/gene="yjjG"""
/protein_id="gnl|SC|F51_03127"
/codon_start=1
CDS 58871..60739
/color=8
/locus_tag="F51_03128"
/note="Non-ribosomal peptide synthetase  amino acid
adenylation domain-containing protein, similar to AA
sequence:UniProtKB:P0C061 protein motif:CLUSTERS:PRK12467
protein motif:Cdd:COG0147 protein motif:TIGRFAMs:TIGR01733
protein motif:Pfam:PF00501.22"
/product="Non-ribosomal peptide synthetase  amino acid
adenylation domain-containing protein [peptide
synthetase]"
/"ID=F51_03128
/protein_id="gnl|SC|F51_03128"""
/codon_start=1
CDS 60912..61979
/ID=F51_03129
/color=1
/locus_tag="F51_03129"
/note="Radical SAM protein"
/product="Radical SAM protein"
/protein_id="gnl|SC|F51_03129"
/codon_start=1
CDS 61999..64239
/color=8
/locus_tag="F51_03130"
/eC_number=4.1.2.9
/note="Phosphoketolase, similar to AA
sequence:UniProtKB:Q937F6 protein motif:CLUSTERS:PRK05261
protein motif:Cdd:COG3957 protein motif:TIGRFAMs:TIGR00232
protein motif:Pfam:PF09364.4"
/product="Phosphoketolase"
/"ID=F51_03130
/gene="xpkA"""
/protein_id="gnl|SC|F51_03130"
/codon_start=1
CDS 64236..65384
/color=8
/locus_tag="F51_03131"
/note="PHP domain-containing protein, protein
motif:Cdd:COG1387 protein motif:TIGRFAMs:TIGR01856 protein

motif:Pfam:PF02811.13"
/product="PHP domain-containing protein [phosphatase]"
/"ID=F51_03131
/protein_id="gnl|SC|F51_03131"""
/codon_start=1
CDS 65381..66169
/color=8
/locus_tag="F51_03132"
/eC_number=2.1.1.163
/note="Demethylmenaquinone methyltransferase UbiE, similar to AA sequence:UniProtKB:P67062 protein motif:CLUSTERS:PRK08317 protein motif:Cdd:COG4106 protein motif:TIGRFAMs:TIGR02072 protein motif:Pfam:PF08241.6"
/product="Demethylmenaquinone methyltransferase UbiE [methyltransferase]"
/"ID=F51_03132
/gene="ubiE"""
/protein_id="gnl|SC|F51_03132"
/codon_start=1
CDS complement(66705..66806)
/ID=F51_03133
/color=9
/locus_tag="F51_03133"
/note="IS256-like element ISCpe3 family transposase"
/product="IS256-like element ISCpe3 family transposase [DDE recombinase]"
/protein_id="gnl|SC|F51_03133"
/codon_start=1
CDS complement(66925..67176)
/ID=F51_03134
/color=1
/locus_tag="F51_03134"
/note="hypothetical protein"
/product="hypothetical protein"
/protein_id="gnl|SC|F51_03134"
/codon_start=1
CDS complement(67179..68024)
/color=10
/locus_tag="F51_03135"
/note="Plasmid segregation protein ParM ParM/StbA family protein, protein motif:CLUSTERS:PRK13917"
/product="Plasmid segregation protein ParM ParM/StbA family protein [partitioning system]"
/"ID=F51_03135

/protein_id="gnl|SC|F51_03135"""
/gene="parM"
/codon_start=1
CDS 68533..69345
/color=10
/locus_tag="F51_03136"
/note="Plasmid replication protein, ab initio
prediction:Prodigal:2.60"
/product="Plasmid replication protein [replication]"
/"ID=F51_03136
/protein_id="gnl|SC|F51_03136"""
/codon_start=1
CDS complement(69612..69911)
/color=7
/locus_tag="F51_03137"
/note="LexA family transcriptional regulator
Helix-turn-helix domain-containing protein, protein
motif:CLUSTERS:PRK00215 protein motif:Pfam:PF00717.17"
/product="LexA family transcriptional regulator
Helix-turn-helix domain-containing protein
[transcriptional regulator]"
/"ID=F51_03137
/protein_id="gnl|SC|F51_03137"""
/gene="lexA"
/codon_start=1
BASE COUNT 28359 a 7014 c 11223 g 23415 t
ORIGIN
1 gaaacttcac caactacgtc tattggaatg aaattcaaat tacttattaa gtctttaaga
61 ttacttgaaa gaaaatcttc tttttcagta gttgaaaaat tataatttat atcttgatac
121 tcaagagcat cagataaatt aatttcatgt tctttatcta aagtattagt aaattcttta
181 gataattcct gaaatagttt aatatcattt tcagctctat tacgagttct ttttagctca
241 ttaaacattc tttcagtacg taatttagaa gattctagtt ctttgaaact ttgattattt
301 ttttctatta gtttatttag tataaagatt ttttttgaaa gatctgaatt atttaaatca
361 tttgtatttt cagttgaatg atttagtcct aatagagaat caatagaacg gttcattaac
421 ttagataatt gtattaatgt atatatgttt ggttctactt cagctctttc ataataacca
481 atagtagttc ttgctacacc taatttttca gcaagttcag cttgagaaag tttatttttt
541 ttacgaaatt cttttatgtt taaagctaag ttgtgtagtt caatttcaat catgtataaa
601 cctcctttca atatagtttt agcaatttta gaagtaataa tctaataaaa cctcacttag
661 aaatgcaaaa gaatcttttt gtaaatacaa attgatatct atattgacat tgatgagtga
721 gcgtgataat atataggtaa attaaatgaa aatatggata aaggtgataa atatggaaaa
781 caagttaagc tttctaagaa agaataaagt gctttctcaa cgagaagttg ctagcttaat
841 tggtaaagaa gcacgtttta ttcacaaatt agaaaatggg aaagtaaaaa accctagtgc
901 aagtgaagtc tataaattat ctagaatata cgatacaagt atggaagaaa tttatagagc
961 aatagtatta aacaactaaa taaattaaat ttaagttttg tccagtagga gaaaaggaga

1021 gcgaactacg cccttttttta atactctttt aggacaaact ttaggagttg atgctgtgag
1081 aggtaaagga gaacttaata aaaataaagt tggaagtctt tatttagatg gtttttcggg
1141 gaaagaaata gcaaaaatat taggggctaa ggaagatact atttataagt gtcttaaccg
1201 aaattttagt catttgaaag aacataataa agctaaaagg gaattgaaaa aactacaaga
1261 tgaagaaata agaaaaatta ctcatagaga atgtaaaaag tatatgagtg atagaaactt
1321 tgttaagact aatagttcca tttacaaaca taatggacat ggaaatttta gtgttaaaaa
1381 agaagaagaa attggatgtg tagttccttt cgacgttcct aaacatttta gttttaagaa
1441 aaagttttag acagcttggt acatatgcga gatatattta aaaaatatta tttcaaaggg
1501 agagagttaa aatgaataag agaattaagg ccataataga atttgtgaaa gtacataaaa
1561 aagaaactat aggagttgca gttgctttag gaatagcagt cgttggtgta ggtggatatg
1621 caataagtaa gcattatgca aacactgata ctaacaatat agctttagct gagaaaaaag
1681 aagatcctaa ggcagatgaa gataaaaagt tagaagaaaa taaagctgca gacaatgaag
1741 aagtaactgt agaagagcaa gaagatggat cattaattgt taaagataaa gatggtaata
1801 tcatcgcaga cagttctaaa ggagacgatg tttctaagat agttgaagaa aagaaagaag
1861 ctggaagcga tgttaatata aaaaataaaa aaggtgaagt tgaaaaagta gagagtgttg
1921 ataatggaag tataaaagta accagtggtg aaactatagg ttctaaaaca gaaaataaag
1981 ttcaagcttc acaacaacca agtggtgaga aggttgagaa agtggaggat ataaaaggca
2041 ctactgcaag tagtgaagga agtattacaa ttgtaaaacc tagtaaggaa gaaagctcta
2101 gtaagcctga aagtaacaaa caagagcaac ctgaagagtc aaagcctgaa gaagctccac
2161 agccttctca aaaaccagaa gagaataaac aagaaaaacc tgcagaaaat caacaagaaa
2221 caaaaccagt tgaacctgca aaacctcaaa agcctcaaag aacttgggaa tatcaaagtg
2281 gtatgagcca agagctttgg aatgatttta atgcttatag acagtctcaa gggttaaatg
2341 ctttaaactg gagtggtaaa tatgctggat ggacaaagag tcactgtgaa gaaatggcac
2401 aaaagcaaag tggcttccat aagagttatc cggaaggtgg tcaaatagtt tatgatggaa
2461 ctgcatatac ttcagctagt tctattttaa atggatttaa aaattctcca gctcataata
2521 agaatatgtt agatccggat ttaacagaag gagcttgtgc agtatataaa gacagtaatg
2581 gaagatatta tacagttata ggttttgact attaatttat atataaagat ctagagtttt
2641 actctaggtc ttttttgata ctgttttttt cggggggaga acatatggga agagtgagga
2701 aagattttta taaaggagaa gtctataggg gtgatatttt ctatgcagac ttaggagaca
2761 taaagagtaa gtttggttca gtacaaggtg gaattaggcc agttattgta attcaaaatg
2821 atattgggaa taggtatgga gaaacttttg tagtgacttc tgtaacatct caacctaaga
2881 aagaaatgcc cgtacatact aaaataataa aagaatgttt agaaaagcca tctactattt
2941 tattagagca aataacaact atttctaggt atcagatgag agagaaaata ggaaagctaa
3001 atggatatga aataaaagaa ctaaatagag cattaggaat aagtattggt ttaagttaga
3061 aagttcaaga gtgttgcaat taagcagcac tctttttaaa tacaaatttt tagaaaggag
3121 aaaagaaaat gaattatttt gatacatctt aaattaattt atcaaaataa aaaataaata
3181 atatggagga gaaagaaagt gagtaaaaga aaaagattta aattaattac aacgataact
3241 ttaattttta cattcttgtt gactaacatc aaagtatttg cagtagaaat aaattctaca
3301 gacgcagaga gttatctaaa ttacaattct ccaacttggg gaaaggttct tcctataggt
3361 aatcatagat actatgctcc agatctaagg acctgttatt gtctaaatac aggtgcactg
3421 aatccaactg gacaagatta cacagaagaa ataccagttg atggaggaat agaaactatt
3481 atttattggg ggtaccctgc taaagatggc tcagaatggg ggatatcagc tgatgaatat
3541 agatattgta ctcaacttgc tatttgggct tatcaaaaag aagctggttt aagtagggga
3601 attgatagaa caagacttca aaatggaaca gtatctttaa gtagattaaa accagtaata

3661 gactttttag tagaaaaagg attaaacaaa gagttaccaa cattctttga agttactcct
3721 agtaatatag ttgctcatca agaaggtgat tatttcgttt ctgagcctat aaaaataaaa
3781 tctgactatg agtttaaaga tgctaaggtt actataaaaa gttcaagtaa tccaggatta
3841 aaagatgtag ttaagataaa agatatggat ggagatgaaa gaaataccta taattctaac
3901 gaaagtttta aagtttatat tcctatagac gcagaaacag gagatattaa ggttgatgtt
3961 aaagcaactg tagaattacc agcttcttta gcttatgcaa ctccagtagc aggtaaacaa
4021 gacatggcac ttgttgattt aagataccaa aatatgaata aggataatgt tactgtatcc
4081 tggactggtt taaatggagc tatagaactt gttaaaaaag gagacgatgg atctctttta
4141 actggtgcta aatttgtttt gaaaaacaaa gaaggcaatc aaatagctga agctacatca
4201 gaaaatggta gagtagtatt caatgacatt aagccaggag aatacatcat cgaagaagtc
4261 gaagctcctt taggatattt aattactaat ccagttaata ttactgtaaa acctaataaa
4321 gtaactactg ctgaaatggt tgatactcaa ataaaaggaa gagtagaaat aacaaaaatt
4381 gatgaagaaa ctggcgaacc aatttcaggt gctgaatttg agatggtgaa ggctgataat
4441 aacgaagtag ttgagaaaat gactacagga gaaaatggaa aggtattaag tgggttacat
4501 ccttttggaa actacattgt aagagaaaca aaagccccaa gtaaatatgt gcttaatggt
4561 aaagaatatc cagttactat taatgaacat atgaaaacaa tagaaattac tcatgctaat
4621 agaaaaatta agggtagagt ttcaatacac aaaatcgatg aagaaatgcc agaacttagt
4681 ttaaaaggtg ctgtacttgg tatttatgat gatgcaggta atgaagttga tagattaaca
4741 actgacgaaa agggagctgc aacaagtaag tatctagatt atggtcacta tattggtaaa
4801 gagctacaag ctccagaagg ttacaagtta agtgataaaa ctatagacat aaacataact
4861 gaagatgata aagtttattc ttatgacttt actaacaaag ttatgaaagg tcgcattcaa
4921 atagttaaag ttgatagtga aaacgaagaa aaaccagttg ctaacgcagg ttttaaagtt
4981 gttgcagtta atgttaatgg tatagaacca ggaactactg tagataaagt taaaactgat
5041 gaaaatggat ttgcatttac taaggatctt agatatggag tgtacaaatt cattgaagat
5101 gaaactccag aaggatattg ggctagtgat aaggaataca caattaacat aaatgaagat
5161 aataaagttt atgttaaata cgttaaaaac gctcctatcc aagctaagtt aagagttgtt
5221 aaagttgact ctaaagacaa taaaccacta gaaggagtta aattccaagt tagaaataat
5281 gatactaata agcttgtaga gtttacaaac tttgtaggaa taattcctca taaaacaact
5341 actttaacaa ctgataagaa tggagaaata ataactccac aacacttagc atacggaaat
5401 tatcaattgg aggaggcaaa acctaaggat ggttatataa gcataaagcc aatacctttt
5461 aaaatagacg aaaatgccgc tctagaagat attaaggact taggtactgt ttatactatt
5521 aaggtatcta acgaccgtat aacaggcgat atggagcttt taaaggtaga cagtgaaact
5581 aaagaaccat tagctgatat tgaatttaaa gttacagctt tagatggcct aatgaaagga
5641 caaacttggg acttaaagag taatgacaaa ggaatagtta gtttaaaagg tttagaatat
5701 ggccattaca aaatagaaga agttaaaaca ctttggaatt acgttattaa caaagagcct
5761 attttctttg atgttaaaga aaatggtcaa gtaattaagc ttgaaatgga aaataaaaag
5821 ataagagcta atgcagaatt aatcaaggta gattcagaaa ctaaaagacc acttgaaggg
5881 gctgagtttg aactttggaa tggtgaaaag cttgtaggaa attacacaac taacaaagat
5941 ggaaaaatag tagttgaaaa cttagaagca ggtaactatt atttcaacga aattaaagct
6001 ccagatcatt acgaaataaa tcaagatcaa gatttaagtt ttgtaataga taaagacggt
6061 gaaactaaga ctgtaactgc agaaaataaa gttaaaactg gagaagtaga ctttactaaa
6121 actgatgtta caaatggaca aaatattgaa ggggctaaaa tagagatagt aggaatagaa
6181 gaacaaaaca aacatgttaa atttgaattt acttcaagtt tagctggcaa taaattcact
6241 ttaccagaag gtaaatatga gtttagagaa actcaacaac cggaaggata cgagctttct

6301 actgaagttg gaacttttga aattaaaggt ggagaaatta taaaggccaa cttaaagaat
6361 gaaagaacta caggaaaact tatctttact aagacagatg ttgctacagg agaagttctt
6421 gaaggagcta aaataaaaat tgaatgtctt tctggcttag ataaaggaaa ggttattgag
6481 tttatttctt caaagaatgg aaatgagttt gaattagcag ctggagaata caaaatctca
6541 gaaactcaag ctccagaagg atatgagctt acaactgaaa ctggaacctt taaaataggt
6601 gaacatggtg aagtagttaa gtgtaactta actaacaaga aatttgaaat tgttaaaact
6661 ggttcaagca tagatgttaa tgcattaatt ccatttggaa ttatcttagt agcaggaggc
6721 ctaggagcat taatactttc aaagaaaaga aaattatcat aattttttta attaagagag
6781 tgccataagg tactctctta agtttacata agaaaataag gagatgggat gtatggttat
6841 aactattatt ttaggattga ttattataaa tatttttatg tatatgagtt tatttaaagt
6901 ggctaaaaaa gcagatgaag aacttgaaat attatttaaa aaacataatt ttcaacaata
6961 aaagggagtg agaaaaatga gtgaaaactt attatgtgaa atagaaaaat tagatttaga
7021 atttagttct ttatcaaata gaaaattaaa taaaaaggac ctagaataca gaaaatattt
7081 aatttcaaaa ctacaaagat tatcgaaaga gtatctaaga gcttgtggaa taagaaataa
7141 atataaactt gagaagatcc taagaaagta ttactttgaa tatcatataa aaacatattt
7201 taagtttttt aatttcagta acatagctgt ttaaaaatgg tttttattaa tttttttact
7261 ttcataggaa ttcttttaag tattttagga cttttttttc ttatattagg tttggataat
7321 tatattccga ttgtggcaag gatgatgttt ttagggatat ttatattggg aatttttatt
7381 ataggacttt cttctcattg ggtattaatg agtataaaga tactgatatt agaaagtata
7441 ggattattta tattaaatat atttttaata ttcatatttt ctaggtatga agttaaaaag
7501 aagtttataa aagtattttt gatagcttta agagttttat taggtctgtt aaatatatat
7561 ttaatttttt ttagcttaat aaatttaatt aaagcttatt ttacagtgat agggtgtgtt
7621 taaaatgatg aaaaaagcct ttataacctt aatcataatt ggaatattgt taataggagc
7681 agctttattt attgagttta aatcaaagtc aatccaggat gatttagttc aaaaatacga
7741 agaaaaaatg tatagtaatt cttccgtaaa gtatagtatt tctgaaacac ctgaaaatac
7801 agagaaaaaa gaagatataa aagatacttt tactcaacct aataacaata agaaaaatga
7861 aaaaaatgat gtgattggaa tattagaaat agatagtata ggggttaaag ctcctatagt
7921 tttaggagaa gaaaatctag attatgtagt ggctaagtac aggtcaagtt ctgattttgg
7981 agaattagga aatgtaatat tagctgcaca caataatatg cgtggatcta tatttagaaa
8041 ccttcataag cttaaaatag gatctacagt aaaaataata tctgataata aggaacttga
8101 atataaaata acaaaccgat atatcgtaga gcctaatgac acaagcaaaa taaatttttc
8161 taatgacaag aaagaaataa ctttaataac ttgtattaat catgctaaac aaagattaat
8221 tttaactgga gaattagtta attttaactt ggtacttaaa ataaaatagg aggaaataaa
8281 tgtgtatgaa taagaatact agaaagatag gcgaagttag aaagaaagtt gaagctataa
8341 gagaagaact agaaaaggat gttcttttta ctgctaaaac acaggagtca aaggagagaa
8401 ttaatgtaat aataactcaa gatgccaatg ataaaataat ggactttatg tatgataatg
8461 caaaggttct taaagaacaa tatggcatac aatctaggaa taaattatta gctgcaatta
8521 tagaatttga agcagatcat atagaaaata tatttgaaaa gttaaaaaga aaggagaaat
8581 aagatatggg atttagttta agaaagtctt ttaaagttgg taaaaattct aaagttaatt
8641 tatcaactgg agggggtgtg ggatttagta caggtaagaa aggagcgaga gtttctgtaa
8701 ataaaaatgg agtaaactta tatggtggta aaggtatttt acgatttact aaacatatta
8761 gttttaagaa aatatttagt tttttaaaca atctaaacaa ataaaataat catataaaac
8821 gagttgttct ccaagaactt aaataatgtt caagggggaa tttttttatg caaaaatcaa
8881 tgttaaaacc accaattaca aggctaggtg gaaagagtaa attaagaaaa gaaataataa

8941 gtatgatgcc agaccatgtg tgttatgtag aaccattctt tggagctggg tgggtgtatt
9001 ttggaaagga aaaatcaaag attgaagtaa taaatgacat tgaaaatgaa ataccaaacc
9061 tatttaaaat tataaagtac catgctccgg aaatgaaaag acttctgtct tatgagatat
9121 ctggaaggag tatattcgat gaatataaaa atgctacttt ggagcattta accgatatac
9181 aaagagctat aagatttatg taccttataa ctcaaagttt cggcggtcaa ggtaatcatt
9241 atggttatgg aactaataca ttgccatcac aaaaaatctt tgattgtaac ttagatgagc
9301 tacaagaaag attaaaaaac acctatatag aaaacttaga ttttcaaaaa ataattgaaa
9361 aatatgatag ggaatataca ttatttttct tagatccgcc gtactatggt acagccggat
9421 atgaagcaaa gtttggaata aaagagcaat taaaacttag agatattctt aaaaatatca
9481 aaggaaagtt cattctaaca atcaatgatt gtaaagaaac tagaagctgg tacaaagatt
9541 ttaatataaa agaagttgaa gtacattatt cagtttcaag acaatcagaa ggaagaaaaa
9601 gaaataaaga attgataatt acaaattact aaaaagtaga tggaaggaga ttaaaattat
9661 ggctgtaaag ttagataaaa attcaaaaga attagcacaa gatttattaa aattacaagg
9721 agtagattat atagagtggt taaataaaaa gcaccaagaa ttaataacta agaatacaaa
9781 gattataaga aattgtgtta aaaatgaaat aaaaaactca agtaaaaaag atgaagttat
9841 agaagaaatt aatcaagaaa gtaatgaaaa tcttgaaagt tttgaagatg aaaattacaa
9901 tgaggatgaa gtagaaagtg aacggaagga gttaatttat taggtgaaag atttaaataa
9961 atatgctttg gagtatagaa aagttgttag gtatgctctt aaagaaggcc ttgcaattta
10021 taacaacaac ttatcagaat tatatataaa tcatgagatt gtaaagaaac ttttagagaa
10081 ggataacttt gattcagtta atggcaaatt atcaatttgg agaagtttaa aatggataga
10141 agtttactct aaaaatagat ttataaagaa ggtaaaagta gaaaaaaaag ttataaatgt
10201 aatagcaatt aatgtagaaa tatttaatac tttaaattta cttttagaag attaaaaaat
10261 tcaaaaatgt gtgcaacttt atgtacaact tcttgtgcaa ctttgtgtgc aaccttaacg
10321 ttttaaaaaa tattaaagtt gcacacttat attattgatt atacaaggat ttatagcaat
10381 ttggttgcac atcgtattat agagtatata ttatatatat aatatataca tataatgtat
10441 taggagtaaa ttgtgtttta atttgcgact aatattttta tatgtatatg ttgtatagat
10501 aattttatac tcttaggcta tagatgtata tctatagtct atacagtaaa taacgtatag
10561 ttatttactg tataatataa taagtaattt attacttatt atattatttt agataggagt
10621 attttcattt atgaatatat ataatcttaa aaaagaaatg aaaaatatta tttattctga
10681 ttcttcttta agtaaaaatt ctattagaga tagaaactca gctcttaata aaatatttga
10741 aaattttaat aaccttgatg atgtttctgt tattactaaa gataaaattt tagataactt
10801 aaattcaatt aaatttaaaa ctaaattatc tgcaggtgtt aatgctatta gatttttaaa
10861 agagcataat cttgatgttg attttccttc tgaagatgaa ctaaaagaga ttattaaaaa
10921 taaaaagaag aataatcgta aacctactgc agagaaaaat ttaattgata ttgaaagaaa
10981 agttaatcga gttagaaaga aaaattatag attagcttat aagttaatgt tagagtcggg
11041 acttcgtgtt cacgaagttg cagctttaaa gaaagatgat ttttcttttg ataaaaactt
11101 aataactatt aatcttagag aagctaaggg taataaactt tgttctattg aacttgaaaa
11161 taaatattta tctaaaaata ttagtgagtt tattgaaact aaaagtgaag atgaaagagt
11221 atttccacac atgagatact tacaagaaca agctactaaa ataggaataa agtgtcatga
11281 tctgcgaagg ggatttgcta aaagaacata caaagaagaa ctagaaaatg actctccaag
11341 aagtgaggct ttagataaaa ctagaataaa gttacgtcat actagaagtg atactactaa
11401 gatttatttg aatagtaaga ttaaagttta aataagagtg ctttttataa gtgctcttat
11461 ttttattatt tttttagaaa ggagtgattt ttttgagtag tgattgggag aaaaaagtta
11521 atcaggaaat agagaatgga gcaaatgcaa ttataaatga gattgcaaat gttctaagaa

11581 ttttcttttt tagagtggga ttaggtttta aaaaatcatg gaagaataaa aagcttttta
11641 ttggtttctt tttaagtttt ttaattccta tagctgcaag aattaaaagt gattattttc
11701 ttgtagatac taaattttat tttaaaataa tatatttttc aacctttata gctcctttat
11761 tttatatggc tataatatcc tttgttaata gaaaaggaga taaggttgag gaagaattta
11821 agaaagcttt tgaagaaatg ggctttatag atatttcagg taatgcacct aaattaatta
11881 atattaaaca tgatgaatta actggaattg accaatactt ttttaaaaca tttattccat
11941 tagatgattg gcaacctaaa gctagattaa tggagtctag attaaataaa actatctttg
12001 aaattattca aggtatagac aaacagacag ttcaaatgaa tgctttggag ggaagtgtta
12061 aagtttctga taagctttta tggagtaatg atttaattcg tgaggaaggc gtattaaatt
12121 ttggtaagag tatttcaagt gaaataaaaa ttgatttaga tgaacaacca cacgttttag
12181 ttgctggtca aactggatca ggtaaatctg tacttatgag atgtcttttg tggcaagtat
12241 attgtcaagg tgcagatgtg tatatgatag attttaaagc cggcgttgag ttcggtttag
12301 actatgaaaa aatcggtaag gtaatgaccg aagtcgatga aactttaaag cttttagaaa
12361 ctttaatagt tgagaataca gaccgtctaa agaaatttag agaagctaaa gttaaaaact
12421 taaaagaata caataagaag tttaaaactc atttaaaaag aaaagttgta tttatagatg
12481 agatggctca acttatggat agctcaggag ttgataaaga aactaaagct aagttagaaa
12541 aaataagtta ttgtattgct acattggcaa gaacagctag agcatcaggt ataaatttag
12601 ttttaggagc tcaaagacca gatgctaatg ttatgaatgg tcagattaaa aataacgtta
12661 ctgttagagt atgtggtaga tttgcagatg gaccagtttc agaaatagtt ttaggaaata
12721 ataaagctaa aaagcttcca gagataaaag gtagattttt atttaaggta gatgcaaaca
12781 ctattgagtt tcaagcttac tattttaatg atgataaaga ttttaagcca gagcaagttg
12841 aaagacaaaa gtctcaatat gaatttattt caatcaaatc ttcagatgat gatttaataa
12901 ttgaggatgc tccagttgaa aatatttcag ttgttgagaa taaaactttt gatgacataa
12961 aacaagaaat agatgtcaat aaagatttaa aacaagaaaa gcaagatgta aataaagaat
13021 taaaatctaa aaataaaaca gatattctat caagtaaaga ttgggatgct agtaactttt
13081 aattaaaagg gtgtgatgaa aatgaaaaat agtgcaatag aaaattttga aaaggcatta
13141 gcagagcttt gtcgttctgc aagaaaagaa aatttaaatg taaatcttga tgtttacaaa
13201 gataaagaaa agaaaaagaa ttactatggc atagtaatta gagataggag tgaggattgt
13261 gtttagaaag aataaagaaa atactaagcc taaaaaagag aagaaagaaa aaaagtcttt
13321 ctttaagaaa aaagataaac ttaacatgga tcttgaggat tcagttgcta atttagataa
13381 gcaggttaat ggtgttaagg agaaaaaatt agtaagtgta atgccattta gaatagcaag
13441 agcttttata tgggttctta taggtattct ttgttttagg ggtggagttt cagtctttaa
13501 aggtaatcaa tctgaagctt tgaagaaaga aaatgaaaag atattaaata atatggagaa
13561 gcaaagttca cttgaaactc aagccttttc ttttgctgaa gaatttgctt gggattattt
13621 ttctagatat ccaattaata ctcaagattt tataaataga ataacaaagt acactacaga
13681 gcaactagca aatgagatga ataatggaac ttattcagat gtcatttata ctagtgcttt
13741 ttattttgaa aaatattctg aaaatcaagt taacgtaaat gttaaagcta gagtaagagt
13801 ttatacacct aaagctggtc aagagcaaac tccacaggac caacttcagt atgatactaa
13861 tttagttgat tactatttag aagttcctat agttttcgat aaagatatga atatggctgt
13921 agatgcatta ccagtaatga cagctccacc agagaaagct tatttcaaaa ataaagagtt
13981 ctccggaact agcgaaaatg atgctgataa aactaaaaag attacagata gtgtatctca
14041 attctttaag gcatattatg agcaaaatca aacacaaata gattatttct tagttgatgg
14101 agcagatatt aagggagcag gtcaaaagtt ctctttcaat aaaatcgata gaataaatat
14161 ttataagctt agtgataaag aattcttggc aattgtagac ctaaatatag attcctttgg

14221 aaattcaata aaacaaggat ttaaccttac tgtagttcaa gaaggagaca agttcttagt
14281 taaaacttta gagccaagaa cttcaaacat agatttaaat aataagatca ataattagga
14341 ggagttatta tgaaaaattt taagaattta aaagagaaag taaaattaca caagttaaga
14401 atacaaggtt tttttgtagc attattatcc tttatgacaa cttatagcac ttatgcttat
14461 gcagatataa atggtgagag tataaaaact aatcttataa gtaatatttt attacctata
14521 tctttcgtag ttgttggatt cttattaata aaagaacttg ctaaaaagag cgttgcaggt
14581 atattagttg ttctaattgt aggtggtttc gtaattataa ctatagctca accagattta
14641 ataaaatcct ttggtaactt tgtaaaaact atattaggtt tataggagtg atttttatgg
14701 aggaagatag aaagatttta agaagctata ataagcttaa aaactttgaa attaaaattt
14761 atagattagg aaatactaaa ataccctttc cagttcctat aacaactaca gtttatttcg
14821 ttgtggcatt aataataatt gctattatag atgctacaat tggaatacac ataaatttta
14881 tatataagta cgttgcttta cctttaggta tagcttatat agccaaaaat aaaaaaatag
14941 atggaaaacc actgcataag tatttactaa gacttataca gtttcaaaat atcaagaaat
15001 ctaaaataga aagatttgag attcgagaga atttagaaag ctttaggttt atagaaagat
15061 tttaggttga ctataaaatc ataaaatttt atattgaaaa tgtataattt tgaggtggtt
15121 ttatggaaaa taaatttaaa gattttttg atgaggaaat aattacaata agtaaatcag
15181 acgctattaa aagattagcg agaagattta aagttgataa tttagaagct caaaaaatat
15241 ataatgagtg gagaagaaat tggtgtgatg taaccaaaaa agagagttag tttttactaa
15301 ctctctttat tttattccaa gtttgaaagg agtatttaat atgaaaagta acaaatttaa
15361 ttttcctata aattttgtaa aggaaaattt gatttttaat aataaagcct gctatgcagg
15421 atttaaactt aatggatttg aatatagttc ttgtaataat agtgaaaaaa ttaggatatt
15481 agatagttta accgaactta taaaagaaat acctagtgaa gctcagattt tattaattcc
15541 aagagcagtg gattgtgaga gatccataag acctatgata aataaaatcc aggaagatga
15601 tcctttgaaa gaagtttcaa caattttagc agagaaaact atagaaattt tagaagaaag
15661 agctgaggat agatatatat ttaatcctgt tactgaagaa aatgaattca ttgctggaga
15721 taaaacatta gaatatgaca catatatatt tatttccctt ctaaaaggcg acgacaagga
15781 ttttatttct aaaggagggg aaatattcga gtatatgata aaaaaaccta aaaaggcttt
15841 aaatgagttt ttaggcatat ctcagaaaac catatctgaa agagatttta gaatttataa
15901 gaaaaagtct acagaatttt tagaaataca atctatgaga tttgatatac gtcctttaag
15961 aaaaagtgaa atagatagtt tgagtgccag ggttactaaa cgtggtcatc aatcagcgga
16021 tgagcttgtt aagtggagtg ataatttctt agaagtagag caagatgaag aaaaagttat
16081 tattccttta agaaatgcct ataaaaatag agttactggc cttatagagc aaggtgataa
16141 aatattaaaa atacaaaatg atggatttac aagttatcag actttcttgg ccataaacaa
16201 tattccaagc atggagtttc caggcgttga gtatataaag ttaatccagg atatgaatat
16261 tggtgcagag atatgcattc acataaagaa gttttctgcg gaagaaagca aaagtaagat
16321 aaaaaataaa gctgctaaga taaatgcaca ggtaaatgaa gcacttcaag ggggacatat
16381 accatctgat gaagaatacg aagcgaaggc ggccgcagag tcgttagaaa aggaagtaaa
16441 gagaaataaa catataatcg aaacaagtat aagtatttgt ttagcttcca caaaggagaa
16501 aatagttaga aataacacca aagtccttat ggaatatttc aataaatcac taaaatttga
16561 gcttataaat ccatatacag accaatacag aattttctta gattttatac cagctaactc
16621 taattataat agagacttca tacagctagt tccaatggaa acattagcgg gtggagtatt
16681 tggagctagt gaccatttag gagatagcgt aggagcctat ataggctaca cagtaaatgg
16741 caatagaaag gtttatttat acatgggaag agctacaaag ttaaataaat cacctgctat
16801 gggcatatac ggtaacttag gtggtggaaa atcttttaat gctaatctaa ttgtagtttt

16861 acatgttctt ttcggtagct cagcactaat tttcgaccct aaaagtgaaa ggtcacattg
16921 ggctgctaaa ctagatttca tgggagattt aataagtata gttagattaa cacctaacga
16981 tgaagataag ggtaaactag atccatttaa tctttataac gataatatag acgaagcttg
17041 tgacttagct tttaatataa tagttgagat tgctaataca aatgaagaag aaaagataat
17101 tttaaaagaa actttaaata aaatgaagga tgaaaaaaga ccttcaatga agcgattaat
17161 tgaaatgatt aacgatgtat cagatagtga tgcttataaa aaagaagcag agcgtttggt
17221 aagaaaacta aatgcatcaa aggatgtagg tttatctaaa cttatttttg gagatggtac
17281 tgaaaaagct attaatctag acaatagatt aaatattctt cagatagata atctaacaat
17341 tccagatcaa ggtacaaaga aagaagaata ctctgaagaa gagaaattaa gttcttgttt
17401 aatgatgctt atgggaagtt ttacgaagaa atttgctatg aaaaaaagaa ataccttcga
17461 cttaatactc tttgatgaaa gttggttttt gaaaaattct gctgcaggta gaaaattata
17521 tgacttttta gcaagacagg gaagatcctt aaatgttgga tgtatattta acgggcattc
17581 agttcttgac ataccaactg aagaaattaa aaatactata acatataagt tatgtttcaa
17641 aacaaataat agagatgaag caaagagaat gcttgaattt atgaataaaa gcattactga
17701 agaaaatatt aaaatgctta tggaattaga aaataggcaa gctgttttcc aggacttaga
17761 tggaagggtt ggagtaatag aatttgacgc tgtttttgaa cagtttatag agtgttttag
17821 cacaacacca gaagatacct taaactatga ggatgtttct taatgaaaaa gctttttata
17881 tggagtggaa taatatcttt gatatttcca cttctatttt ttttcttaat tatagggga
17941 agtggtgaac agccaactcc agtaaatcct aatcctaatt taacggaaga gcagcttaat
18001 tttataagtc agattgtttc cggagcgaag caatcctatg aagaaacagg aatatttcca
18061 tcaataaccc tggcacaagc tatattagaa agtggttggg gaagatctgg gttggccata
18121 aaagctaata atttatttgg tataaaagct gactccggat ggaaggggaa agtattagaa
18181 atgccaactc aggagcatgt taatggggga ataataacta ttatagcaag gtggagagtt
18241 tacgaaagct ggaatgaatc tgtaatagat catggtaaat tctttgtaga aaattctaga
18301 tataaagaaa atggggtcct ggatgcaaaa aattatgtag agcaagctca gtgcatacaa
18361 aaagcgggtt atgcaacaga tccaaactat gcaaaccaat taatccaggt aataaatgat
18421 tttggactaa atctttacga tatgaacgga gatgttgtag ggaatgatgt tatagaaaaa
18481 gcaatagaag ctggaatgaa gtgggttggt aaatctccat atgtttgggg aggcggtcgt
18541 aaccaagctg acgttgatgc aggacgattt gattgtagtt ctctagtgca ttattgttat
18601 gcatctgcag gaatacagtt agggccaaga gaaagtgtaa caacatggtc cttaataaac
18661 atgggtaaac ctgtacctgc aagtgaaatg aagcgtggag acttaatatt cttcgatact
18721 gctggaagga acggccatat tggaatctac cttggaaatg gaaaattctt gaatgacagt
18781 tctacccaag gagtaagtat tggagaccta aatagtgctt attggagcag atattttaac
18841 ggaaatgtaa gaagagtcgt agaataaaaa ttcggcttta agaagaaaag tacctggata
18901 attatcaata tttttaaaaa tatccaggta ctttttata aatttacgaa cattaatcgg
18961 aggtgtgaaa tggagaatgt gaagattaat atatatagta aatttctgta tttgaaaaat
19021 aaattaactt ataaaaatat attaaaagca atagttataa ttgcaactat cgttatagct
19081 tacaactgtt taactaaaat agcttatgca gatggatttt taagctatgg taatagtgat
19141 gatccttata aaaacattcc tgagggatgg ggaatgttta tttctgaaaa gtacagaaat
19201 aactatgcct tagatattat agatactcat ggttgggagc ttatagataa ggctataaca
19261 ggactaacta atatagtatt tagcggaata gtaatgatta gttggtttgg agtaaatata
19321 tttagatttt gcttcacaaa tgatatagct gcaggctttg caaatcaatt aaatagtgtg
19381 ttaggctctc taaataatgg aattttcaac aatttcttta tgatagtatt tatgatttca
19441 ttgttttcaa tagtttataa tctatataaa aagaatttta gtgcaatagg atcacaaata

19501 cttgcagtag ccataatata tgcaactaca gcttttttaa gcaatggagc agttggcttc
19561 ttatctaaaa caacggagtt ctctaagact ataggaacta gtgccatagt tgctataaat
19621 ggtaaagata ataatgaggg ttccttagaa gattattcta aaactgttgt aggaacctta
19681 tggggaaatt tcgttcaaca accttgggta atgctagaat tcgatggaaa ggttccctta
19741 aatgatgaaa atatgacaac taaaaattct ataaatgaag aagcgttaaa ttactcgaaa
19801 gatatactat cagagcctta tggtagtgat gaaagagaaa aagctttaga gtcaataaat
19861 gaaaaattta ataaggagtt gttttcaaag aatggcataa atactaagtt ggtatccgta
19921 tgtgtattat tcgtaataac tctaataaaa atgattattt taatagctgt aggaatgata
19981 caaataggat ttcagcttat gtcaattctt ttaatattgt tacttccatt tatcatgctt
20041 attgccatag tgcctttctt tggtggtaca aatattataa aaggaatagc aagaaagtat
20101 ttaggaactc agctcggtat aattgttagt agctttgtat tagccttact ggttcttgta
20161 gatactgtta cacttacaat gtttcaagct tggggagcta acttaataat agctctttta
20221 atgcagtgta cttgttggat catggttatt gcctttagaa aacagttaat gaacgggtta
20281 ataaaccttc aaaatagaat tgctccagga acagcaggag catttaggtt tggagctaaa
20341 ccactagata aaggtgctga tattgcaaag gaatatgcag taaatccagc tatggataga
20401 gcgtatgaca taggagataa tttaataact aaagctcaat atgctaaaga atatacaaag
20461 ggaaatttaa gtctttacaa agacagaaaa gtagcacaag ctatagattt cttaggttct
20521 aaatacgata atttaaaaaa taataagcct gaagatatag agaatgctaa taatccaaaa
20581 gaaaatatag acggggttaa agatacaaaa aataaagtta actcaaatga aaatatcaat
20641 aaaaaacctg tagaaaaatt tgatttaaat ggagttaagg ccactaaacc tagtaatata
20701 acttctgaaa atattcctaa taagaatatt aatggcgaaa ataaacctaa tattgatgaa
20761 gacgcagtta ctaaaaatga tataaaagat cctatagaaa ataaggatat agctcctaaa
20821 actatagatg aaaacattaa aaataatgga aatgataatg gaactatcca ggataaaagt
20881 ccaggaagta aagaaaataa aactatttct gaagatgagg aaataactac aaaaccagaa
20941 gaaataaaaa ctcctgaaaa gactataaat aaagaaaatc taaatgaact aaaaaatgaa
21001 aatacatctg cagataagaa aaataatgat agtaatagta gtaatataag tgttactgaa
21061 attcctgaag atttagtgga ggaaaaatct gttgaagcta taaataaagc caaaaatgat
21121 gttaaagcaa ataaagaaaa aataaataat aatataaatt ctaagaatgt tgaaagtaat
21181 cctgaagtaa aagaaaatat agaagtagct tcatcaccag aagttacaga aattaagaaa
21241 aaatctacta agaaagttga aaatatcaaa aataatggac aaccaaaaga aaaatctgca
21301 gataagaaag aaattaaaga agtatcaact aaagaaagag ttgagagagt aaaatcattt
21361 atgggagtaa gtaataagga tataaaatct aaagaaaagc atttagaaaa taaattagaa
21421 tacattaaga aaggaaatag caaaaacttt aagaagaagg ggagctgatt atgctatata
21481 aaacacattt atcaacatca ctaaatgttg caataccagc tttatatcta agcaatgagc
21541 tatcactttt aacattaacg ggagtaggct tggggagcct acttccagat ctagatactc
21601 caaatagttt tattggaaga agattaaaga taatctcata tcccttaagt acaatattta
21661 agcatagggg atttcttcat tcaataactc ctgtaatatt aatatttatt ttaggcatat
21721 tcttaaaatc actatttata aaatctatcg ctttaggtta tttacttcac ttagtagggg
21781 ataccttctc agaaagcggc ataaagtggt ttttaccttt ttataataaa gatattaaaa
21841 tgccattagg tattctaaga tatagaaccg gtggcctaaa agaatatgtt atactaggaa
21901 taagcattgt aatattaatg ctagaaatac aaaaatttca tttaataaac ttattttcaa
21961 tataggaggt aatatgaaga agaaattagt atttttaggt ctaatatgtt taagtttatt
22021 tgcttttgta ggttgcggaa ctgaaaaact agatataagt aactgcatag atgtacgcta
22081 tggagaattt aatggttcag ccaaaatata tgaaagtagt ttagatttag gaaagttaca

22141 agatattcct aagttaaagg gacttactcc tgatatgcta aaaggagact ataaaataac
22201 tttagtagga gataaaactg atttaaaaaa tggagataaa gttaaacttc acctagagta
22261 taacaaagaa ctatataaaa gagactttaa tgtagaattt aaatttgaac caactgaagt
22321 aactatagaa ggtctgccag ataagttaac agatataaat caaatttcta aagagcaatg
22381 ggataaaata tatgctgaag ttaataaaaa agctgaaatt aaagctaaag ataataaata
22441 ttctgactta aaattagaaa aagttttaga atttaataaa aaaaaatctt ccggtggaat
22501 tacaatagag tttatttatt catataaaaa taaacttaat gaaactagat atataagttt
22561 ttatcattta tttaaattcg ataaagaaaa tactttttct aatatagaat ttttagaatc
22621 atctatttgg aatgataaat tttcaattaa ctatagcaat acattaaatg aaatattaaa
22681 acaggtatat cctgataata attataaagt tttatggagt gcagaaaatt aatttttttaa
22741 ttgtagatta taaaataacg atgataaaat tatattatca tcgttatttt tataaaaaat
22801 tttggtaata tattgtctaa ttttaggaaa tattgtataa ttaatttact caaggaaaaa
22861 gataaaaaat actataagaa attaataata agttatatac tcgtttaaat tacatatata
22921 aatttttcat agtttaaaaa ttttgatttt tactatgttt atttgttatg taatttaagc
22981 gagttctaaa gtccaagctc aaggaatttg agtttgggct tttttatatg cttttttaga
23041 gtatagacta caaatctata cagttgccta taacttaaat aggagcatgg taaagccgtt
23101 gtgataacgt aaatatttac cggagtgagc agatacacta taaactaatc ttagcgaaag
23161 ctaacaaggg tttgattgag ccaaggaaga aagttgatcc ggctattctc atattacaca
23221 tatatgaggg tataacgtaa gtcttattga aactctgatg aagatgttta gtctaagtag
23281 ctctataaga gctatatgac aaaattcagc ggttacgtgg agtacctgca gcttaaaaat
23341 aagctgcaac gaaaggaacg tatagagcag tagaaatact ggacaggtag gggtacaatt
23401 cctgttttga tgtacggatt aggtgaaata ctaatcacta atgtgttgtt accgggacca
23461 taggttaatg tggcgaggct ctgcataggg atcttaattc ggtgacatag ttatcctgtt
23521 cttttttttaa ttaaatcgga caggataact atgtacattt tgagggtact cgggatctaa
23581 tcattaaaaa cttaaaaagt caatatataa attaaatgta aatttgaatt tatttagata
23641 ttaatagatt ttataagaag taaagtgatg gtgaagtaag atgaataaga gaaagtgtac
23701 cttatgtaag aattttgata aatataaaga attcaattgt ttgtatttta agataaagat
23761 agaagatcct gaagtagcaa caaaatgtac taactttatt ccaaagagaa aggaaagtag
23821 aaagccttgt aaggattgta tatattacgt acgtaatcca aaggtaattc aagttaagaa
23881 gaataaagtt gtttttgata aaagatttta ttattgtcta cgccttaata agcgagggac
23941 ttattttaga aaggttaatt gtaaatactt ttcagaagaa ttataaagga ggaagagttt
24001 atgaaggtat taagtttatt tagtggaata ggagcttttg aaagagctat tgagaataaa
24061 aatattgagc atgagatagt taattattgt gaaaaggata ggtatgcaag ttatgcctat
24121 agtaaacttt ttaggttatc tgaagctaag aatttatggg atgttaattg ggttgatgga
24181 acacatttta aagatataga tcttgtaacc tatggcttcc catgcaccga tataagccta
24241 gctggaagct gtaagggaat aattaaaggt gaaacaaaat ccgggctgct atttgaagca
24301 ctaaggatta tagaagaagc tagacctaag gttgccatag ccgaaaatgt tagaaattta
24361 gtaagtaaaa tgtttatttc tgattttcat aatttaatta atgaattaga taggttagga
24421 tataactcct actggaaaat actaagtgga attaactata attcggccca ttctagagaa
24481 agagtattca tagtcagcat caggaaagat atagacaacg ggagctttga gtttaaagaa
24541 ggaaataata atttagttac cctagaaagc attactgaaa gctttgtaga tgaaaaatat
24601 tattgtaaag ataattctta tttaaaaagc ttcttaaata aggttaataa aagagtttgc
24661 aaggataaag agcctagtaa attcggatta atgagagttg gaacaattaa aaatccacat
24721 atgttacaaa tgaatagaag agttttctca gagctaggag ctagcccaac ccttgtaact

24781 ggatctgata gtattcctaa gattattcaa tgcaaggtta gaaaacttac tcctttagag
24841 tgctggagac ttgtgggatt ttcgtctgaa gattactggc tagttagaaa agctcttgaa
24901 gaacaatttt ataacgggaa agactgtact gatacgcaaa tgtataagat ggctgggaat
24961 agtattgtaa tccaggtagc agaaagtata attgaaagct taaaaggaat attatattaa
25021 aaattgaggt gagaatatgg gtgaaatatt attagaattt ctaaaagaaa ctcgtattat
25081 aaatagacaa gctaaattat taatagaaga taagaatagt ttatcttctt cagataaaga
25141 aatgttgaat aaaattatat taaacacatc taaatcttta agtaagttag gttcagaaat
25201 taatttatga ttagtttatg tggcagagat tgtaatagct gtgttatgaa aaaagaaaag
25261 atgtgtaatg gatgttccat gtgtgatgtt tctttttgta aatgtggtga aaaaagaaag
25321 cgttgtatgg ttgtttgtcc taataaattc ggttctttta ctttagtaaa aaatactata
25381 gtaaaggaac ccttaatagg aaataaacct ttagatttac caatttatat acctgtaatg
25441 cctgataaaa tcaaagaaag ttttaacttt aaggccaata aaaatataat tgctgttcat
25501 ggagagttct ttttaaatgc tgctggaagc aaaataacgg gggcatataa tccaggattc
25561 agagctgcat taaatttaaa agaagatcta tccggaatat tagaatttta tataaaggat
25621 agaacattag aaggcttttg ggataatcga aaatctattt ataaagactt aaaacgccag
25681 gactttttag gaataatagc tcctaatttt tctgtttatg aagatgctcc acgcctggaa
25741 catatatata atattcaacg ttctaaaatt gtttataatg aaatgatacg tgagggcctt
25801 ccggctattc cggatatatc ttggtactca aaagaagatc taaacttttg gataagagaa
25861 attaaggcca ataatataaa aactatatct tttagtttca tgaatgtaga tactaaactt
25921 aaagcaagta actcatggaa acactacttg ctaggattta aaattttaaa ttttaaaata
25981 cctttagatg tagaaatagt ggttgtaggt atttcttcag tccaaagaat tgaggaaatc
26041 cttaaaataa gtaaaagtag aaaaatatct tttatgcatc aagctgcttg ggttaatagt
26101 agaaatggag tatctgtaaa agataaaaaa caactagata aatctatttc taaagatgat
26161 attttcaaaa ataatctaga attttatact agtgagtaca ataaattata tgaaaagtac
26221 agtaaataac tgtacttctt tttttattgc tcctaggacc aataaaaata atagataagt
26281 aaagaagtgt gttaaataat cttatgaaag gggagattaa tttatgccta aaagaagaag
26341 caatactatt gcaaatacaa cggaaggatt aattggtggc tcaggatcta atctttctga
26401 aggaataata gaagatttag ttaatgctgt taatggtaga ttagataata ttgtaactgt
26461 acaaggatta tctaggacca gaaacagaat tgaagatata aatatagttt ctttaggtaa
26521 taatcgatat gaagtaagtt caactgaagg aggaacttac attgttgata tgaatgaaaa
26581 tagctgtgaa tgtggccaat atgtttatag aaatgaatgc tgcagacact taatagcggt
26641 cagagaagct ttaggcgaag ttgcagaagt agatactata aacaatgtag aaactactag
26701 aagagctagg gatataagag atgaaataaa tagaaatatt cgtggggaag aacaagacga
26761 taatcacttt tatacagaca atgaagttga tttcgataac gattttgcag aaattaatga
26821 cgattctata gagtatgagt acgaaaacgt actaaacgga aataagagta catttggagt
26881 tgaacttgaa tttgttggtg gtgatgctaa tgctatagca cgagaactct atgatttagg
26941 tatagttagt tctcctagaa gattaggcta tcactcaaga agtgagccag gcaaatggaa
27001 gcttgagaga gatggttccg tatccgatgg agatgccggt ggagaaatag tttctcctat
27061 tcttaaagat actccaaaga catgggaaca aataaaagta atatgtgatg ttgctaaaag
27121 acatggggct agagtagatc aaaggtgtgg tggccatgtt catataaaca tggaaaaact
27181 agatactgca agacaaagat ggagaagatt ttttaagact atagaagtat atgaggattg
27241 tatttataga gctgcaggtg gagatttagg aagagtaaga agtaatgcta ggcattatgc
27301 cactccattt agtcctaggg ccgatgaaag taaatatata agatttaata tggataatga
27361 tgaagatgta agaagaatgg cagcagaagt ttcaaaagga aatagatatt atggtattaa

27421 cttaactaat atagcaagag atagagctcc tacagttgaa tttaggcatt ttaacggttc
27481 attaaatgag aagcaaatcc aggctaatat aaaaatggct gccggtataa taaatgcttc
27541 agaaaaagct agatttagag atactgaaga tgaaattttc aaaaagagag gaaatattct
27601 aaagaatact tcaaggctag atggaacaca aactaaaaag aaaatgatgg agtttttaga
27661 cttaacattt ccacgtagaa aagataaaaa tgcaatacta aatgttttta agaaaaatga
27721 atggagataa tatactttta tagaaaataa tattataata tttacaagta ttacaataaa
27781 taatcatttc tctttttatt aaactaaaat aacgaaagag ctctacttag taaatctaag
27841 cagagctctt tcaatgactt ttattgtgat acaaaatgaa tatattataa ttataaaaaa
27901 tttataatta taaataaaga aaaaattaaa cgttttctta ttttaaaaat ttcttaacat
27961 aatagcctaa atttatatta taataatttt aagcttttac tatgtaaata gccatttttc
28021 cttacgtaat ttaagaaaac aaagaaagtc ctacttaaaa ataagtagga ctttctttgt
28081 ttaacttagg cacttataca taaagtcttt taaatatttt aacttaaatt tcaagaaaaa
28141 taaagagtaa tatattgttt atttattaag tttaaagtta taattaaggt gtatctttta
28201 agagttttta tatcataatt atagcttttc acatttcaac ttataactaa gaaagtatct
28261 atttttagat actttctttt tactataata agtataatat tttaggagag tgattatatg
28321 aaatggtcaa aagaaaagat taaggataaa aaggaatact ttttaagcca aagaaaaaat
28381 cctactggaa aatttacaga aatggttgtt agaacatatt ttttaattta taaacaatgt
28441 tctctaaatg acaatttttg tcctagtaat aagataaatt ccaggatttt agtcagcgat
28501 gtgtatgaaa atttttatga taataaaaca tcaaaccatg ttcctagtgt ggttgatgac
28561 tgtataaata atttaaataa aatgggctat ataaaattta ttaaaactaa ttcatcacca
28621 aagtggatgg tgaaaattga aaaaaattta gattttattc tagatggtga agaagatttt
28681 tacttcaaaa aatataatat aactcaaaaa atagtataaa aatatgtcta taaattaaaa
28741 atttaaattt ataaaaaaga aagtattatt gataatactt tcttttttgt ttttaacata
28801 taggaagtga tgatccttgg atggtttaat aagaagaaag atactagcct ttttacaatg
28861 gaatgataaa aatggttatt atactgacga acgatgtgac ttggaagaag ttcaaaaatt
28921 atcattagaa gaaagtataa aatatttttt tggagtgata aattcagagt tttattattc
28981 aatagctgat aatatctttg agttaagctt ttacgaaaca ataaaatatg ctaaagatta
29041 taagttttat aatcaaactt ataaaaagtt aaaattattg atagataata atcctaatga
29101 aaatttatat agaaatttac ttgaataaaa aaataaagtt acttaaaaat aagcaacttt
29161 atttttagtg tttatatata taaaggcaaa aagcaagaaa cttatagctt aacaatatta
29221 tactttaaat actaaaatta ataaagccag ttaataaata actggctacc tagatagatt
29281 caagaatctt caaatagtag aagatataat tattttacaa taaaaattaa taattatcaa
29341 aaatattaat ttttgataaa ataaaattgt taacattttt tgaattatat ttttaaagag
29401 cagttcaatt ttcgggattt tgagctgctc tttttactat taataatatc tgcgatttgt
29461 ttttaaaaca aatataaatc ttaatatctt tggtaaaaat aaaagcataa acattcctgt
29521 taaataagct attccactat aattcccaac aatataataa ttgtgagcag cataaaaata
29581 atacactaac attacatatc taccaagctt aattaatttc atatttgaaa aactaaatat
29641 acataaaact attaatacta aaataaaatc cataatatct caatcccctt ttatttatta
29701 ttcatgtaaa attttatgaa ttactccaaa gctgcaatta tttaaagctg caagctcttt
29761 tattgttttt ccttcttttc tttgagctct tataatattt ttttcttctt cagtaaatct
29821 aggttttcta cctgctccac gactattctt tttagatttt tctttaagct ctaaatattc
29881 tcgacttact ctcattaaat ccattctact atctacatta aatttcttat aaattcctga
29941 taatacatca cgataatatt cttcagtttc tttattttta tttttaaac tctcaatttc
30001 tgataaaaat ttattttgtt caaactcact attattcaat agtttaactg taaaataagt

30061 gttataacct tcatttcttc tttcatatat ttgaggtgtc ttttctattt taaaaagctt
30121 tcttaattca ttaattaata tttcaacttc atcttttgag ttagtttcaa ttctaaattt
30181 aaatccttct tggccataag tttgattata tgcttcttct aattcttctc ttgaatattc
30241 taaaacacta actggatatg tattaactgt ataatatgat ttactagatc tgctgtaata
30301 ttctctttta acgggtattt cttttattgc tccaatttta tgaagctgat ctattctttc
30361 agtattaaca ccataaaaat cagcaattgt ccatgtagga atgtatccaa acttttcaaa
30421 cttattaaaa atagtatcaa attctccagc tacaactaat ttatctataa ttttagattt
30481 agcttctcta ggtttaaaat ctattccata ttcttcagca agttttatta tttcttgttt
30541 tggagtgtcc ttattattct taagttcatt tagaaaaatt tcttttaaat tatccttcat
30601 atcatcaccc tttatattta atatgttaaa tatattatat cataaattaa ttaatgaaaa
30661 aaatcttttt tcattaatta atttatgata taatcaaaat ataaagataa gtggggagga
30721 aatggcagta tgatagtgat tgataaatct ttaggtgaaa taaatcctga aagttattta
30781 ataaaaaatg ctaaagataa tacatatctt ttagcattac ctaataacct taatggatat
30841 aattattttg aagtgtacat agataaacta aatagaagta ttcatgtttt tgatagtcta
30901 gagaatagaa aagggggaac tagtgctatt aatagtgtag atagaatatt agaaattagg
30961 agacctttaa atttagatct tgactataag cttgtgatat attacccaga tcatagtata
31021 tttaaagctt gcataacaac ttatcatgaa agaaaaggat ttaataaaaa tagagactat
31081 gtaacctata taccttttct gaagaaagct gaattatttc taaaaaatag attttaatta
31141 gagcttttag ataaaactat attataatta gagtataact attgaaagga gtaaattaaa
31201 atggtgtata gaacattact aaatccacaa atagattttg tgtttaaaaa gatatttggt
31261 acagaaaaga ataaacctat attaattaac tttttaaacg cagtaataaa accaacaact
31321 cccataaaag atgtagaaat caaaaataac gatatagata aagattttat agaagataaa
31381 tttagtagat tagatgtaaa ggctacaact agcaataaag agcatataaa tattgaaatc
31441 caggtaaaga atgaatataa catgatacaa aggaccttat attattggag taaaatgtat
31501 tcagaacaaa tacaaaatag agataactac agtaagttag aaagaactgt ttgtataaat
31561 atcttaaatt tcaaatattt aaagaatgat aaatatcata atgcatatag gctaaaagaa
31621 ataacttcaa atgaagagct gacagatctt caagaaatac atttcattga attacctaaa
31681 tttaatgaaa tcggtaataa agagtatgtt gaaaatgttg agaaaatgga tgctttagaa
31741 aaatggttag aattcttagt tgaaccagaa agtaatactg taaggcagtt agaattaagt
31801 aatgaagaaa ttaaactagc aaaatctgaa ctatatagac ttagtatgga tagtaatgaa
31861 agagaacagt ataatatgag agaaaaggca atatacgata gaatatctgc cttagaaaat
31921 gctgaagcta aaggaaagat agaaagagaa ttagagttaa taaaggagtc tttaaatcaa
31981 ggattagaaa tatccttaat ttctaaaata acgggattaa gtgaagaaga aatattaaaa
32041 ataaaaaaag atatttaata aaggaggaaa tagtatgaat ataattataa tatgtgctat
32101 tataggtgga ttaatgggat acttctatat gaatagaaag cttgagaaaa aaacaaataa
32161 ttagtaatat ttttgtgata actaaaaaag actctataat ttagagtctt ttttgatttt
32221 aattaaattt ggataaatat aaatataaaa atattgaaaa aaaataaaaa aaaatatata
32281 atgtagttat gctaaaatat aaacatgaaa aatataaatg taaaggtgtt gtaaatatgc
32341 gcaaagaggt aatttttaat taaattaact gaataaggat tatttttaaa agtatatgaa
32401 aaatcttgaa taactcaagg tttatttgtg gtggaatcaa ttatttataa tattgatatt
32461 accatgttta tttgtgtact ctaaaaataa tcagtattat aatgagcatg ttacatcttt
32521 atttgcagga taggaaaacc gtgttctaaa cacggtattt ttatgtctat ttttgctata
32581 tatagatatg gtataagatt ttaatttgaa tagcttaatc ataagttata taattaaagt
32641 tctatatttt atgtttatat ttaggttata tgattatgag gctgtaagtg aattatgttc

32701 atttgcagct ttttttgtt taaggagatt attttgatga tttatatatt taaatatgaa
32761 gattatgaaa aacaaacaaa ttttttaaga aatactagga tggtgatgat taatggttaa
32821 taaactttca gcatataaaa cttatttatt attttcagct attgcagcaa tgtgtttttc
32881 gttagtagct acagttatgg tagtgtatca cattgaaata gttcatttaa atccacttca
32941 acttatactt gttggaacta ctttggaatt agcatgcttt atatttgaaa ttcctacagg
33001 tatagttgca gatgtgtata gtcgtaaact atctattgtt attgggggag ttttaacagg
33061 agtgggattt attttagaag gttctatttc tagttttgtt ttcgtacttg tagcacagat
33121 tgtatgggga ttagggtcta cttttatcag tggctcgctt gaagcttgga ttgcagaaga
33181 agagaagaat aaagatttag atgaaattta tataaaggga gcacaagcag ggcagatagg
33241 agcatttatt ggaatagtac taagcactgt aatagctaat ttctctgtaa ggcttcctat
33301 tatagttagt ggagttttat ttataattct tgcattattt ttatggttat atatgccaga
33361 aaataatttt aaaccatctg ctcctgggga tttaaataca ttcaaaaaga tggtatatac
33421 atttaaatct ggtcttaaaa ttgtaaaaag taaatctata attatgattt tacttgcagt
33481 aactttattt tatggattat caagtgaagg ttatgataga ctttctaatg cacatttttt
33541 acaagatact acacttccta agcttggaaa ccttagttca gtgacttggt ttggaatttt
33601 tggaatttta ggaatgatat tgagcttcat agtaatgcat tttatggcaa agaatcttaa
33661 gaatgaggat aataggaaaa atggaaaact attattatgt ataaatatac tttatatatc
33721 gtctatgttg atatttgctc ttacaaaaaa ctttagttta atgttaatag cttatttggc
33781 aacaaatacc tttagaatta taaatgaacc tatattcagt gcgtggttaa atgggcatat
33841 agatgataac tctagagcta ctgtactttc tataaatgga caaatgaatt ccttaggtca
33901 aattttaggt ggaccaatta taggaatcat agctacaaat atttcagtaa gtattggtat
33961 agcatgtact tcgttattag taacaccggt attagtgtta tatattgttg ctatgataat
34021 tgataaaaag gtggatgata gagttggagg tattgattat gaagaaaata attaatatag
34081 gaatcgtagc acacgtggat gcaggaaaaa caactataac agaaaactta ttatattata
34141 gtggagctat aaaatcagtt ggaagagttg atttaggcaa tacacagacg gattctatgg
34201 agcttgagcg taagagagga attaccatta aatcgtcaac catatctttt aattggaata
34261 atgttaaggt gaatattatt gatactccag gacatgtgga ttttatttcg gaagttgaac
34321 gttcattaaa tagcttagat ggagcaatac tagttatatc aggagtagag ggattcagt
34381 cacaaacaag aatattattt gacacattaa aggagttaaa tattccaaca ataatttttg
34441 taaataagct agatagaatt ggggcaaatt tcaacaaagt atttgaagaa ataaagaaga
34501 atatgtccaa taaagtagtt agattacaag aagtatatga tgtaggaagc aaagctgttt
34561 atataaaaaa actatttgat acatgcataa taaatgatga tgctattaat gttttatcag
34621 acttagacga agcattttta gaaagatata ttggtggaat agaacctgat aaagaagaaa
34681 tacaagaaaa gctttcatta tatgcaagag aaggaagtct atatccagta ttttgtggtg
34741 ctgcagcaat tggacttgga attgaagatt tattagatgg aatttgtagt tattttccat
34801 ttgcaagtaa tgattgtgaa agtgatttat ctggggtagt atttaaaatc gaaagaacaa
34861 gtaaaaatga aaagaaggtt tatgtaagat tatttggagg aaaaatatct gtaagagata
34921 aaattcaagt acctaataag gagatagcag aaaaagtaaa gaaaattaat aggttagaaa
34981 atgggggagt tgttgaagca cagaggatag aagcagggga tataggtatt ttatatggac
35041 ttacaagttt ccaagtggga gatgttattg gaatttcaaa tgataaaatt aaaaatatat
35101 ctatagctaa accagcatta aaaacaacaa tttctgcaat tgataaagaa aaaaatccag
35161 agctatttaa agcattaaca ttacttgcag aggaagatcc actactagaa ttagagatga
35221 atgacataga taaagaaatt tatgtaaact tattcggtga agttcaaatg gaaatactaa
35281 gttccatgtt agatgattta tatggaataa aagtagagtt ttcgaatatt gagactatct

35341 ataaggaaac acctaaaggt tttggagcgt caataatgca tatgcaggaa gacttaaatc
35401 cattttgggc gacagtaggc ttagaaatag aaccagcagg gagaggcgaa ggtcttaggt
35461 atatttctaa tgtttcagta gggtcattgc caaaatcttt tcaaaatgca attgaagaag
35521 cagttattaa gacaagtaaa caaggattat ttggatggga ggttacagat gtaaaagtca
35581 ctcttagctg tggtgaattt tttagtccag ccagcactcc agcagatttt agaaatgtga
35641 cacctatggt attcatggaa gcattatata aagcacaaac tgttttatta gagccattac
35701 atgagtttga gttaaagatt cctcaaaatg ctttaagcaa agcggtatgg gatttagaaa
35761 ctatgagggc aacctttgat aatcctattg ttatagggga tgaattctca ataaagggat
35821 taattccagt agaaaattca aaagaatata aaatgaaaat agcttcatat acagaaggta
35881 gaggaatgtt tgtgacaaaa ttttatgggt ataaggaagc ttcagctgaa ttttcaaaag
35941 cacgcaaaaa aacaacgtat gatccattga ataaaaaaga gtatttgctt cataaactaa
36001 acgcaattag agattaaatt taaatagata agttaaattt ttatatgtag gatttaaata
36061 taagacattc aaagcctaaa agtaaaggaa gggtcgtctg aagaagatga agacccttcc
36121 tttttcatg ttactatgct taattgtaag tgcttgaaat ttgtaatata atgtaaataa
36181 agtttttaat aataagggga cggcacaaat gaaaatcgac tttaaaataa caaaagatga
36241 ctatataagc ttcaatttaa atcatttaga aaactcgaaa tcacagaaaa gtacatttaa
36301 catacttaga tatgctgtac ccatagtact ttctatacca atctacttta caggaacggg
36361 gatatttaat caaccgagta tatattggat tatagtagcg atagtttttt tagtcatttg
36421 gattttaaca tatcctaagc aatacaaaaa attagtagct aaagaaactg ataaactcat
36481 aagttaagga gataattcat caatatttgg aagtaagact ttagaaatta ttgatgggac
36541 aatagtagta aaaggggatt ctatatcaga aacagtatcg ttagaaagta taaaagatgt
36601 aaaagtttac gaagatatga tccttatata taagtggatt tgtagctcat ataattccaa
36661 ggagatattt agataaagaa atagagaaag aacttatgaa agagcttaaa aaatgtaagg
36721 ctttagaaca attaaagtaa aacgtgctcc tttggaggtg taacagagaa ccgtaccaca
36781 agtaaggtag cgcggatgat aagaataaaa atttatatat ataatttaga tcttagttat
36841 acttaaaaaa taattagtac gataatatat gtatagataa tatcgataaa agtatattac
36901 taaaaattat aattgttagt atgaggagtg attatacatg gatatgctta ccaatttcaa
36961 taaagttatg gagtatatag aaacgcattt agaagaagaa atagacttta atatggtttc
37021 aaagattgca ggagtatctg agtatcattt tagaaagatg ttttcgtatc tttcaggtat
37081 gacactttcg agttatataa gaaaaagaag gctatctaaa gcaagttttg atttaatgca
37141 aaaagatata aaaataattg atatagctat aaaatatgga tatgattcag cagatggatt
37201 tagtagggca tttaaagaat ggtttggtgt gaatccttca gagattaaag atgctaagaa
37261 tcttaaagta tttccttgta tgacttttca attaactatt aaaggaggaa gtaatatgaa
37321 ttatcgtatt gaaaaaaaag aagcttttaa attagtagga gttaaaggtc aagttcctat
37381 tgtgtttgaa ggtataaatc aagaaataat gaaaatagca caaagtataa cagaagaaca
37441 aagaaaatta ttacaatcat atagaaatga agatgttaaa actgttgtta acgcttcatt
37501 tgattttgat gataaaaggt atgaagaaaa aggtaactta aatcacttaa ttggatctat
37561 aactacttta gatatagatt ttggtgaatt tgatgttgta gaagttcctg cttgtacttg
37621 ggcaattttt ccttgtaaag gaccatttcc aactgtattt caagatacat ggggaaaaat
37681 tttatctgaa tggttgccat cttcaaatta tgaattaata aacgttccag aaatatcttt
37741 caatagagat atgtcagata tgcaaaatgt ttatagcgaa atttggattg gggtaaaaga
37801 aaagtaatta gaactgaaat tttattctat atcacagtta aaggtacctt gaataaatta
37861 aggtgccttt tgatacttat ttagaataac tattaaggaa ccttatcata aataaatgag
37921 tttttttgta aagtattgat attacttcaa tagtagctat tttatttcag aaagtatagt

37981 atcataatca ataattaagt attaagaaaa ttagattcaa taaatctaaa tataactgat
38041 aattagtaaa atatataagt aaataagtat tctaatcagt gtattaaatt tggatatttt
38101 tatatagaaa gtattgttaa taatgaaact atacttaaaa taaggtaaag ataacatggt
38161 taaataccaa atattgtgtt aattttaatt atttataaaa ataattaaaa aaaggttgca
38221 attaattaaa tactatgtta ttataaaaga gtagcaaagt tgctaacaaa aaaagtttca
38281 tttaggtaaa tagtttctcg aaaattagag attattatat caatttagaa tctttaattt
38341 taggaggaat gtttatgtca gataaaacat taaaatgcag agattgtgga agtgaattcg
38401 tattttcagt aggagaacaa gaattctaca aagaaaaagg atttgaaaat gagccaacaa
38461 gatgtatatc ttgtagaaga gctaagaaag aacaaaatag aagataattt tagattgtta
38521 taggttgtag tttaaactac aacctttttt agtataaaaa aataaaataa ctattgtaaa
38581 aacctcactt aatgctgata cggagaaccg tatcacaagt aagtgcgaac tttaccaaaa
38641 ttggatatat aaatgaaagg taggtctgct ttaattctgc aatgaaactc acaaaaatct
38701 aaaatttgaa tttttgtagg ttattgtagt tttattcata ggttcttcgt ttttcaccta
38761 tttcagcaaa ttagaatttg ttttttctaa tctatagtat tgaaaaaaat aaaagtttat
38821 atataatgta attatactaa aatataaaca tgaaaaatat aaatgtaaag gagatgtaaa
38881 tatgcgcaaa gagttagttt ttaactaaac taactgaata aggattattt taaaaagtat
38941 atgaaaaaaa tcttggataa ctcaaggttt atttgtggta gaaacaatta ttaataatat
39001 tgatattacc atgtctattt gtgtactcta aaaataatca gtattataat gagcatatta
39061 catctttatt tgaagaatag gaaaactgtg ttctaaacac ggtattttta tgcctatttt
39121 tactatattt aaaaaaatat caagtttaga ttatatatgt ctacctaagt acacttaata
39181 agtttagaac aatttattga attatcaaag tttaatataa cttacaatgt atctataaat
39241 ccagaaatat taactcctac aaatagatta aaaaaattta tagatttggt taaagataaa
39301 aatataaatt gtataatggt tccttagtgt aattcgttat aagaaaaagg gatattgtta
39361 aaaaatttca ttaaaaacaa taaataaaag taagttttta tataattttt ttcttacttt
39421 cattttgtag tttataaaag ttcaccttta taaactttat ttaatttatt tatttttcgt
39481 ttttaaatgt atataatttt atttttaaac gtttattaaa atctattgat tttaataaac
39541 gtttttctta tacttaaagt aataatttaa atatggagtg ataaaaatgc gttataagtt
39601 gacttatgta tatggtgact cagatcaaaa atttactcaa acctttagca gtaaagttct
39661 tatggaaagt tatattgaaa caggtaagga taaagattta cgggtaatta atatagaaag
39721 tagtaaactc tatggctatg caagggtaag ctctaaagag caaaatttag atagacaaat
39781 agaagcctta aaggagtatg gtgtaaatga aagagacata ataactgata agcaaagtgg
39841 aaaagacttt aatagagaag gatataaaac acttaaagag cagctattaa gaagtggtga
39901 tgttttagtt ataaaagagc tagatagact aggtagaaat atggcacaaa ttaaagaaga
39961 atggaatgat ctgcaggcaa aagagataaa tatagttgtt attgatacac ctatactaaa
40021 tactgaaggt aaaagtaatt tagaaaaaac cttaatatca aatatagtat ttgaattatt
40081 aagctatatg gctgaaaaag aaagagtaaa gattaagcaa agacaagcag aaggtattgc
40141 taatgcaaag gctaagggaa aacatttagg taggccaaga gtagaatatc caggtaattt
40201 taaagaagta tatgacaaat ggaaagctaa aaaaatcact ggtgtaaaag ctatggaact
40261 aatgaattta aagaaaaatt ctttttataa cttaataaaa aaatatgaaa ttgaaaaata
40321 agaaaattaa ggtaatttta tatgttttca aataaaaagc ttagaaactt atttagtaat
40381 tggttaattc caataatttt agcaatttta cttgctgtaa ttattaataa attattttat
40441 acaaaatagt agttcctagc caattaatgt ccccaacatt aaaagttgga aatcaactat
40501 ttgttacgaa agtttataat acttcaaata taaagagagg agatatttta gtctttaatt
40561 ctgcaagaaa aaataaggtt aatatttttt attaacctta ttttttcttg cagaatataa

40621 aacccaatta atattactta taataactaa tattataagt aaaggaatcc ataaaaattt
40681 aggaatcatt attaacataa taaatcctat aacccaaatt attattgaat atatcaaata
40741 tctaagaaat agtttcaaat aatcacctac ttaaaattaa atttattatt aatttaatta
40801 atctaaaaat caacctttct aatattataa tcttcctttt gataaattaa cataattcta
40861 gataataatt cttaaaaagt atctttattt aaaaaatgtt aaaaatttga tataatagaa
40921 ttgtaaaaaa aattcagggg ggaaaccaaa tgaaaaaatt aatagtaaaa agtacaatga
40981 tgcttttatt ttcttgctta ttatgtttag gaatacagtt acctaataca gttaaagcaa
41041 atgaagtgaa taaataccaa tctgtaatgg tacagtattt agaagctttt aaaaaattatg
41101 atattgatac gatagtagat atttctaaag atagtagagc tgttactaaa gaagaatata
41161 aaaacatgtt aatggaattt aaatatgatc ctaaccaaaa acttaaatca tatgaaataa
41221 caggttcaag aaaaattgat aatggcgaaa tttttctgt gaaaacagag tttttaaatg
41281 gtgctatata caatatggaa tttacagtat catatattga taataaatta atggtaagta
41341 atatgaatag aatatcaata gtaaatgaag gtaaatatat tcctacacca agtttcagaa
41401 ctcaagtttg tacatgggat gacgaattaa gtcaatatat tggagacgct gttagtttta
41461 cacgttctag taaatttcaa tatagttcta atacgattac attaaacttt agacaatatg
41521 caacttctgg atcaagatcc ttaaaggtaa aatacagtgt agtagaccat tggatgtggg
41581 gggatgacat tagagcttct caatgggtat atggtgaaaa tccggattat gctagacaga
41641 taaaattata tctaggttca ggagaaactt tcaaaaatta tagaattaaa gtagaaaatt
41701 atactccagc atcgattaaa gtatttggtg agggttattg ttattaaaaa aatgaggcta
41761 cgcctcattt tttcttttga tttttaatat tttaataata taaaaaccta gaatggatga
41821 tattattcta aaaattgtta aatcaaaatt aaaagaatga aaatgtaaca atgaaataat
41881 aacattaaat aataaaaaaa taattaaact tggttttaga gaattatatt tatttataac
41941 aataggtatt aaaataccaa aaggtataaa taataataga ataaataatt gatctaatac
42001 aaatacatta tctaataacg gtttattact attaaataaa ttatataaaa atttaaaagg
42061 taaatatgca ccttttgata aatagccacc accattagta gtattggaaa tacaaattaa
42121 aataaatata gtatatattc ctagaaataa agatgataaa atcctagtat ttattagatt
42181 tgctttaaga taagactctt tagtagaatt tataagttca tctatctgaa gagtgttaat
42241 tactttctca ttagcttctt tttcagaaaa accttcattt aaaaaatcct ttttcatttc
42301 ttctgcggta tcaagtaact ctaattttaa ttctgataat tctctagatg aaatatctaa
42361 tttccttaat tttttattta aaagtctttt caatgtcata ttttattcct ccttaatatt
42421 ttctattaaa ttagatataa gataaaagtc tttaagtttt aaatttaatt caattaaacc
42481 agcttcagta attttataat attttctttt tttctttgaa ctattagttt caatgaaata
42541 atcttgtatt aaattttttt cttttaatct tttaagtgct ggatataagg ttccttcccc
42601 taatattaaa acattatcag ttatgttttt aatttcttta gatatctcat atccatacat
42661 atctttattc tttaataaag atagtaaaat tatttctaat gaacctttta atatctctct
42721 atccaaaaaa tcaactccct ttaagttaat ttagtgttct ataatttaat tttacgaaaa
42781 catacattgc tttacaaggt attaattaaa aaatatatta tattttttaa ttaatttcag
42841 aaaatactta aagtattata atttatataa aaaagttaaa gcaaaaagct ttaacttttt
42901 tatttctgag ttataatgtg agtaaaaata ttgagaggtg tattaagtat gaatttagtt
42961 aaaacaagag atgatttaga aagagaagct cctagactaa aaaaagagtg gatacaaaaa
43021 attgatagta tagatgatgc aaatagaaaa tatgtattag tttttgatga tctagtattc
43081 gaggcagacc atgaacagga tattacttca agactaataa gagattatat agaaactgat
43141 gatagaaata tgcagcttct atttagaatt gactttgcaa gagctttaag tatgtatagc
43201 ataatgaatg gaattaatgt tgaagtttat aataacggta agaaagtaag agataactat

43261 gctgtatctg aagatgatcc ggattatgaa aaggactatg aaattcctga tgtaatcctg
43321 gatgtatttg atgaatttac tctttttaag ggccttaatg aacttaagta tgctaaaata
43381 tactataagt ctgatgatgg agagtacaag ctattttaat ataaattaaa ataaagaagt
43441 agctttaagt aataaaaagc tgcttctttt ttataaggtg aacatatgaa aaaaagagaa
43501 caaatatata catttttaat agattttata aaagaaaaag gatatcaacc tacggttaga
43561 gaaattgctc atgcagtaaa tttaaagtca tcgtcttctg tttatagaca tttagaaatg
43621 ttagaaaaag atggactcat aatacttgga aaaagtgagc agagaggtag aaaaagaagt
43681 attcatatta ttgatttaaa caaaaagatt aaaaaaaatg tatcagaaga aactattaaa
43741 aatttaatta aattttcaaa aaaacaagtt aatactaata atgatgtatt tcccaagaaa
43801 gttactaatg attatatagt aactgtagat aatatacttc ctgatgacat tattattatc
43861 gagaaagtaa attctctaga aaaggatagt ataggattat tattatttga aaatcatacc
43921 tttataaaaa aaatattttc agattatgaa aaggtatatg atactgatcc tgtagtttta
43981 gggaagatta ttggtgttta tagagatttt ttctgatatt acttgaaaaa acataaaaga
44041 aacgttaaaa taggttaaca atattaaaat ttactgtaat aatgtattag aaactttcga
44101 gtttatcagt cactagttaa accagtatta actattaaat taattgcaaa cttaatttga
44161 tctgtatatc tattaacaca aaaaagatta tctctttaat tttagagata atcttttttg
44221 tgtttagaat taaccaagaa tattatacaa tttaattata tcctatactc atagtaatta
44281 taaatttggg tgagaaaatt tatgagatta tcagaagaag ataagaaatg tataaaaatt
44341 ctttatagaa atggcttaaa tgtaaaagaa attgaaaaat tattaaataa tgacttttct
44401 aaagacgcta ttcaaaaaaa tatatataga catttaaatg aatttaggaa tgaacatatt
44461 ataaatagat ccttaaataa aaataaatta aagaaagaat ttaaaaatat tattaagaat
44521 aatttttaat aaaggatttt ccataatttt caagaatatt atttatatat tcttaaatta
44581 gggggagtta agatatgaaa acagctcaag aagtagccaa tgaattagat ttaaccagga
44641 ctgaagttta taatttgctt agaaaaaaaa gattttctcc attagttaaa aagaacggtg
44701 gtcaagtagc gataagcaat gaacttttca aattattaga agaagaaatt aaaaataaaa
44761 gaaataatat taaaccacaa gagaataaaa atatagatat aaaacaagaa gataaatcat
44821 tagaagaaaa aaatactata gttgatgtaa aagctgaaga aaatcctgat tttaatacta
44881 tagaaattct acaaaatcaa ataaaattaa aagataatca aataactatt ttaaataatg
44941 ttataactaa taatttaaag agaataaaag atcttgaaga aaaaaatcat gagtttaaag
45001 aattacatga agaagtcaat aatttaagga tggttattga ttcaatacaa actaaaaaaa
45061 agagaaaaag acttttagga atattttaaa agagaaggtt aattataacc ttcgagactg
45121 tagggaaatt tgtgtaaaaa caaatttccc tacagttttt ttgaatattt agctggaaaa
45181 tttggaataa tattcttaat attttaagga ggatttcaaa tgagtaaaga aatcaattat
45241 caagaagaaa ttaaaaattg taaatcagtt gaagatttag ttggtaagaa tggattaatg
45301 aatagttaaa ttctttatga accataaaaa ttttttcttt agtaaaaacg attgcaacat
45361 attaattaaa gtggtattgt tttcaaggga taaatcttct taatatctac actttacata
45421 ataaagacta atgatctttt tacacttaat taataggagg tatattaaat ttatgttaaa
45481 aaaagttaat tttcatactc atacagagag atgtagacat gcagttggta aagataggga
45541 atatattctt gaagccatag aaaataatct tgatattttg ggttttagtg atcatggacc
45601 ttatccagat aatagattta gattaagaat gaattatatt gatcttttag aatacagcga
45661 tacaatgaat aaactaaagg atgaatttaa agagaagatt aaactaaaga ttggattaga
45721 aattgaatat atttttacag aggattttta ttataaatat cttctataag aagtaaaact
45781 tgattaccta gcattaggtc aacacgtatt tttaactaat tcaggtgaat taaaagatac
45841 ttatttttta aagagcacaa aagattatat agactatgct gaaagtattt gtaaaggcat

45901 gaaaagtggt tattttacat ttatagctca tccagactta atatttttaa atgattttcc
45961 atgggatgat aactgtgaaa aagcttgtga tttaataata gaaacggcta aaaagcaaga
46021 cttaatactt gaatttaatg caaatggatt aaggcgagga ttaagaaatt tttatgatgg
46081 aaaaagatat ccttatcctc ataaaagatt ttgggaaaaa gctgcttcta agaaaataaa
46141 aactttaata aatgctgatg ctcattcacc taatgatatt tgggataaaa aaatggacga
46201 agcttattct attgcaaatt cattaggatt gaatttagtt tataatataa aataattatt
46261 tatatttaat ttttatttag tatagataaa atcaaccatt ttacataata ttataaactt
46321 aaggaaatct gataaattat acgtgttata tcaagttttc ttaagtttat atatgtatta
46381 aaataaatat atgaaagtta cttttaaata tatacttagt taaaaaagaa cttattttta
46441 aaaaaatata aatttctttt aaatcattta ataattcaat tcaatattta tctaaatact
46501 catctgaaat tttcaaaaaa tgatctgtat tattaaaatt ataaaaaagt aattctagtt
46561 tttaagatac ttgaaataca atcatagaaa aatactcttt aatatcttaa aattattaaa
46621 tttactgata atatatgaag atataatcaa atttattttt aattgaggat ttattactac
46681 atttatggta tttcatcatt gcatatcaca atatttatta aatttacaac aatatacata
46741 aataataaaa aaataacatt aaagtttatt tattgtattt tttatggtaa tatattacta
46801 taaaaaaata ttaaaaggag aatttatgaa atctaattat gatgaaataa ggcaaatttt
46861 aaatgaaaca ggtatagata caaaaaagaa tacaaattat gacaatttag aaatagattc
46921 aattacaata atagaattga tattaaggat agaaaaattt tatggaataa ctatattaca
46981 aaacgatata aaaatagaaa atttttcaaa tgtaaacgct ataaataatt tgcttttaaa
47041 gtatttaaaa aaagaaactg atgaaagcat tgaagctaat gaagaattaa atttttatat
47101 ttaataggag gaagtatgtt tataaaaagt gattcctcaa aaggcattaa ttataatttt
47161 gaaaataaag atatatgttt aacagaatgt ttatttatag gtcctatcaa aaatcctata
47221 gagaataatt tttttaatta taaattagat aatatattca ttaataataa ttttaaaatt
47281 actgaaggtg agtacttata tttaataaat aaagaaaaat gtctattgaa atatttttct
47341 aaacttgaga gattcaatgg aaattattat agaaaatcat tagatttttc agaatttaat
47401 ttaaaaaaag atgaaagaat ttttgcgttt tttaatatta attctaatat tgatagaaat
47461 attattttaa aaataggtag atccaatgat tttaaattat ggttaaataa tactcttatt
47521 tataataaat catctataat atataatgaa ttttatgaat caactggatt tagtgattca
47581 atcttcataa atttaaaatt aaataaagga gataatttaa ttttaataga attaaataaa
47641 aattctcaaa tgatttcatt aaaattatct aaagataaat atgataatga attaaatgta
47701 tttaaagatt attatgaaag tacttgttta aatagaattg gtattattca tcacaccaac
47761 ttatgtatta atctagataa atacgaattt atgataatac ctaaggatta tataaatatt
47821 gatctaaata gtgaattaac aataaaaatt atatttaaaa ataatgttat tagtataatt
47881 aaatcaaaat tttataaaaa aaatataatt aatattttag aatttaaaaa tagtgacgaa
47941 tgtattatga acattgaagt tatatattta actaaaagta atattaatat gatacataaa
48001 ttaaaggttc tgttacctaa ttatttgttt ttaaagagta attcaaaagg atatagaaat
48061 aatattatta attatttgga ttttataaag aaggatattc ctttagaaat gattccatgg
48121 ataatttctg aagaatataa taaaataata gatttaagta gtattacaag ttatttatta
48181 gatactagtg gaattcatag gatatttttc acttcaaagt tagacaatac tcttcaaaat
48241 atttgtgtat ttctaccgga agattattct agagaaaaaa gttattcaat agttgtacat
48301 ttagcaccgg cagtatcggg aaattttata aagtttcata gtcagaagtc ttataaatct
48361 aatgtaattt ttgttgaaat ttcaactagg ggcgtaacac aaggaagtta tataggagaa
48421 tatgcaatat tagaaaatat aaaaaattta aaagagattt taaaattaaa aaaatatact
48481 ttcaatttta taggttattc aacaggtgca tttgcagttt gggcattagc agaaaattat

48541 cctgatatgg ttaattctat tgtaacaatt tctggttttc cttattatag aaacttgtgt
48601 aacttaacta atataccgat tttaaatata tctggagata atgatgatat gattaaaaga
48661 tcatttgaag aaccaaccaa tttctttaag aataaaaatc cacaatataa aggattatta
48721 cttaaagaag cagatcatct tacaatcatg aacactctat ttagtaagta tgttataaat
48781 acattaataa acagaaatcc aaatgaaatt atcaatgtat catttaggac tgaaagaatg
48841 agacataata aaacaaaata catagaaata ttaagttata aagattatta tggaaaaatt
48901 gatattgagt tttataatga acaaattaat ataaatattg aaaatattac tagttttata
48961 ttttatgttc cttcaaattt taaaaataaa aaatttgata tttatataat aaataataat
49021 ataaagcaaa aattcactat aaaagatact gataaattat tgtttaaata ttttaaaaat
49081 agatttatat taattgatga agaagtatta gaagttaatt taatagataa tattaaggga
49141 atgggactgt tagatatata tatggatagt attaaaatat tatttcggaa tactacaaat
49201 gatgaagttc gcaaaagtat gtataaagtt gcaaaaacaa tgtcaaattt aaaaacaatg
49261 ggatataatc cgaatgttaa tgttaattat aaaatagaat gtatagatta ttttcataaa
49321 gataattatt gtaacgataa cataattcta attaatgaag gagaaaataa tttttttgaa
49381 gaaataattc acaaacttcc tattaaaccg tattgcaatg ggttttatta taacgatgaa
49441 tttatagaag gtgattattg tatattattt ataactaaga atccatataa tgataagaaa
49501 aagattctta tcatttattc aacaaaatat gaactattta attctaatta ttttttaaga
49561 aattttataa taccaagtta ttcaaatggt ttaaatcaat ggttaaatgg agacgcctta
49621 atatataaag ataaagaata tatagtaatt aattcgatag gttgtccgat taaatttaaa
49681 taatttaatt agataaaatg agaggttttt taattatgga gaaacaactt aaatttctaa
49741 cttctggttt aataaaaagt taccaatatt atgcgtatcc tataaatata atactttcaa
49801 aatataataa acaggaagat aaactgttat taaatttttt tacgaatgta tttagaggaa
49861 agagtatgca atattatgaa tcaaattatc aaaaatggga ttgttttaat ataaaaaaat
49921 taactgtata taaaaatttg tttataaatc aattaatgaa attaattgat gataatattt
49981 atattcatat tacaaaagtt aatgagtatt atatacctaa tagaagggcg tttaataaat
50041 ataattatat acatgatttg ttggtaattg gatacaacaa attagaagaa acttttttaa
50101 ttgcaggatt taatgaaaat aataacttta tgaaaactga agttaaattt actcaaatgt
50161 tatcttcttg tttttatgaa tctaattata cagaattgat tttaatatct gtaaaggaaa
50221 attataatta tattataaat actaataaaa ttaaaaaaga acttaaaaga tatatatcat
50281 gtgaagtttt aaatatgagt gaatatcaat tagatgaata tacgtttgga tttgatgcat
50341 ataaaaaatt aaataaagat ttaaaactat attcagaagg taatacagat tcaatgccgg
50401 gaatcattca agatatatat tttatatatg aacataagca aataatttat tataaattac
50461 aatatttgtg tactaataac ataatccctt tagatatttt agaagagttt tcattaataa
50521 aagaagaaag taatataata aaaaacttat ttattaaatt tattattaca aaaaattcta
50581 attatataaa aagaataagt ttaagattgg atagcttgtt aaaaaaagaa gaaaaaatat
50641 taaaaaaact taatttaata atctagtaaa attttatata ttaatcaata aaataggtag
50701 aaactttaat ttttagattt attaaaagtt tataataata gaaattcggt ttagaaagaa
50761 taaagactaa agttttataa cagtaaagaa aaaatatgta gttaatacct tgattataat
50821 gaacaggaat actaggtaaa caaaatagag gaaatataaa aaattatttt cagagaaacg
50881 agtggtgata attagaaatg ttaaagaatt ctttaaaatt gaaaaatgaa gctaatacat
50941 tattagaaag gattgaaatt ataaaaatat tgagtagata tacaaatgaa tactgttttg
51001 tgggtagtta tgctaacaat actatggtat ggagagatat tgatatatgc ttttatttaa
51061 atgaaactaa tattgaaaaa ttttttaatt taggttgtga gttagccaat ctagatggag
51121 tttgtgctat gaaatttaat aatgagttta tacttaatcc aaataaacct gatggatttt

51181 attggggaat agatattatt gataaatcca atggggaatt atggaaaatt gatataaagg
51241 gagttcaatt agatacatat attgacatta aacgttatac agaagagttg attactaagc
51301 ttgattttga gatacaagag ttaatagtgg ctgttaaaaa taaattgttg attagagata
51361 gtagatataa tagatataga acacctcatg gaagtagttt ttatatttat gatttaatag
51421 ttaacaaaga gattagaaat ataaatatag tattaaaaga attgaaaaag attggaatat
51481 attagtagtg acggaggaag tatggaagta tactatcaat taataaaagt agcaataaaa
51541 aagcagttac aatatagtaa atcgattatt atttttttta ttgtaaattc aatgaaaatt
51601 atgttatatt atacgctttt taatgctatt tataaatata atgggacaag tgaaatttta
51661 ggttatacat caacgcaaat gatttggtat tttggattac tatcagctac atggtcacta
51721 atttggagtt ctattgattg gcaactttca acaagtatta taaatggaaa ttttgcaatg
51781 gagctcttaa aacctatatc tatattttcg agaggattat gtgaatctat agggacaagt
51841 ataattagtt tttttactga atttttattg ctacttttag ttcagagctt ttttatattt
51901 cctaactttt taaacttaat ttctattata aaatatttag tattgttagt atttgcattt
51961 attatgaatt atctaataaa ttttatttgt ggaatagctg catttgttac aactaatatt
52021 aaatcagtta taaacttcaa atttatgctt tttgcagttt ttggtggatc atatatacca
52081 cttaatttct tgccagaatc tatcaagaat atttcttgct atttaccgtt ccaatatata
52141 atttactggc caatacaatt tttcctcaat actgaagaga ctagtgaagt aaaagtattc
52201 ttatatgtaa ttataaatca aattttatgg atactagtgt tatatattat atatagaata
52261 ttatggaaaa aagcaatcaa aaaatatgta tcttttggag gctaatatgg agaagtataa
52321 gcgattaata caattattta tgcataatag taagattgag attcttagaa atatgcaata
52381 taatgtagac tttattttat attcttttat ttcattattt tatactggta ttggtcctgt
52441 tgttcaatta attattttta agaatactaa aggttatcca ggatggaata tagagtcaat
52501 aattttattt cagggaatgg ttattttaat atttggtatc aataagttaa tatttcatcg
52561 tgtacgttct aatattaatt atatttttaa aacaggttca tttgatcaaa ttttattaaa
52621 accatatcca ccaatagctg ttatattatc aaatggtttc tatttggaaa gtataacatc
52681 aattatagcg ggatttatta taattaatta tagtctagtt aagttaaata taaattttag
52741 tatttatcat attttaatat ttttagttac aattgtatgt ggattattat taacagcatc
52801 tattaatatt ttatatgcaa cttttactct tcatttaatt aatactgaaa agtttagtga
52861 gataatagat acattacttt cttttaaaga atttcctttg cagatattcc ctagagctat
52921 acaattacta tttactacta tatttccaat tgcaattttt gcctatatac caactcaagt
52981 tttattagat agaatggatt ttagtctatt attggctata ggatttagca taattgcttt
53041 tattataagt ttgagaattt ggaataaata tatgtataaa tatgtaagtt taggaggata
53101 actatgttag aaaattatat tattaaacta aataatgtta aaaaagaata tagggttaag
53161 gcatacaaac aaggaatatt aaataatata aaaatactgt ttgaaaaaaa gtataaaata
53221 gtaagagcag tagatggaat tagtattaat ataaagtatg gagagataag aggattatta
53281 ggaccaaatg gagcgggaaa gtcaactaca ataaaattac taacaggaat tttacatccg
53341 tcggatggtg aaatagatgt aatggggtat aaaccgtgga ctgacagaat ggaatatgta
53401 aaaaatatag gagtaatatt aggtcaaaaa tcacaactta tttgggatat tgcaccgcta
53461 gaaacatata aattgcataa atctatgtat aatatttcag aaagtaaatt caatcagaac
53521 ttagaatact tttgtaaaat tcttgatata gatgaagtga ttagaaaacc tgttaggttt
53581 ttatcactag gagaaagaat gaaatgtgag tttgtttgtt ctctattaca tgatcctaaa
53641 attgtatttt tggacgagcc tactatagga cttgacgtaa ttgcaaagga atccataaga
53701 agatttataa aagaaattaa caaaagtaaa agaacaacat ttattctaac tacacatgat
53761 ttatctgatg tggaggaact atgtgataat attacagtta ttagtaatgg gcatgttgta

53821 gtagatgatt caatagataa tttaaaaaat ttattttctg agaaaaaggt tatagacttt
53881 aaattttccg aggaagtaaa tatcaaaaaa tttagtgatt tagacgtagt ttgtattgat
53941 ccatactctg tatctataag agtagataat aaagaagaat atcttaataa aataattagt
54001 aaagtagtta atgagtttaa tgttatagat attaatatat ctaaggtttc tatagatgat
54061 attataaaaa gatgttatat ggagaagtaa agataatggt taagattaat gaatataaaa
54121 catatgaagg tagtgactgt tttttaggtt gtatacaaaa ttatttattg ttatatcaaa
54181 atttagttga tgagtgtaac attttttttg aagggaatgg ctttgattta acatatatat
54241 taaatgaaga tgataagtat tctattaaat ttaaggcaca tatcaacgaa tccgtttaca
54301 agtattgcga taaatatggt gtaaaattac aaaaaaagaa atttaaagat ataaaagtag
54361 ctgataaaga attattagag gatttaaaga aaaatcatat agtaataaaa ttaaatgcag
54421 agaacttgaa gtataatgat gttttcaaat actctttttc agttacacac tatataaatt
54481 tgttatcata tgactttaat aagaatagag catatattag tgatggtttt atacctaaat
54541 atccaagaga aatttatcaa ggatggtata actataagga tcttagatat gcacgagaaa
54601 aagtggaata tgaatatata gttgttgaaa aagattcttt acaggatatt tatgaaaaat
54661 taaacaaatg tgaggtaaat aaaagattat ataattgcgt aattgataat atcagaagat
54721 atataagagg tggaaaagat aaagatgcat atcatgggat tgatgctata gaacatttta
54781 tagataaaat ttcaggtcta gtagtagatg atgcagaaga gttttcaaag aatatctata
54841 aaattcactt aaagcttggt gttaatggat ttattaatat aaaatcgctt ctagcaaaga
54901 caattttaaa attagggcaa gaatttactg ataaaaattt aataaaaatg tcagaagagt
54961 tacaagagat atatgagaag tggatatata taaatcgttc atttattaag tgttctattt
55021 ccaaaagaca gaaagatatt gatattatag ccgaacgaat gaacaagctt tttcaaactg
55081 aatacaattt gtataaaaaa cttttactat atatggataa aaatgaggtg aaattatgat
55141 atataaccta tttaaaaatt cagttcataa gtatggaaat aatattgctg tgcaatgtga
55201 tggaaagaag ttaacatata ctgaactaga tatttcaata aagcatatag caaatgaatt
55261 aaagagaaaa ggaataaggc aaggagatca tattgctata tttatggaca attcagttga
55321 atacattaag gcattttatt ctattaattt gtgtaatgca gttataattc caatttatac
55381 acatatagga cgagaaaagt ttttgaggct catagaattc ttagatgtta agtatatcat
55441 ttctactagt gattataagg aatttttaag caatgaaact atagttaaat gtcaaaaatt
55501 atccaatata ttcttattat ataatgatga tgtcaaagat ataggcggat tttataaagg
55561 gagtattatt aataaatatt atgaagaaac ccctgcagtt ataatgctat catcaggcac
55621 tactaatttg ccaaaaggga taatgctatc aaataagaat attaagagta atattaattc
55681 tatatctagt tatctagaat taaataataa agataaagtt ttattggtaa aaaatcttac
55741 acattcatct agtattacag gagagatgtt aacgtctttg agtaatgggt gtactttata
55801 tttaacacaa aaaattataa caccatcaat gattattaag ttaatttcgg aattggatat
55861 atcaatattt tttggtacac ctaatttatt aactttaata ttagaaaata agaatatagc
55921 taaatataat ttcaaaaaat taaggaaaat taatttttat gggtcaaaaa tggatgtaaa
55981 tattatacag agattatgcg atacctttac tgattctgaa ataatatatt cttatgggtt
56041 aactgaagca tctccaaggg ttagttatat caaaaagaat gatttattaa ataaaaaggg
56101 atcatctgga aaaccaatta aagatgtaag tatatttata gaaaataaaa ataaaaaaga
56161 attaggccct ggtatggttg gagaaatagt agttactgga cctaatgtaa tgatgggtta
56221 ttataagaat ttagatttaa ctcgaaaggc acttaataaa tctaaactgt atactggaga
56281 tataggtttt attgatcaag atggattttt atttataaaa ggaagaaaag ataacatgtt
56341 aaatgttgct gggaaaaatg tttatgcaga agagattgaa gagatattat gttctataga
56401 aggcgtaaaa gaatcattag ttactggtaa aaaagataaa ttttatggtg ataaactaat

56461 tgcctatata gtaaaaaatg aaaaaattga tactgaagag atattaagct atcttaaatt
56521 atatttagat aattataaag ttcctcatga aattatatgt gttaaatcac ttgaaaaaac
56581 agttagtggg aaaattgtaa gaaaggaggt gaagaactat gaatagcgat ataatgaaaa
56641 gagtaataaa cattttatca aataatgaaa ggtttaaaag tagtgaagtt aactatgact
56701 taaatttaat taaagattta gggtttgact cacttgatat tatgaaattc atagtagaat
56761 tagaagatga atttaatatt atgtttgaac cagaagagtt agagtatgag aatataataa
56821 atattaagaa tttatgtgat ttaataaata gtaagattat ttaaattgat tagggagaag
56881 aaatgagaca tataaatgaa gagttattta atgaatgcta tggaaaaatc aaaaaaaatt
56941 ttatggaaat aagtaagaat aatttatatt acagaaattt atttaataaa tataaattta
57001 atattgagga tttgttaaat tataatgatt tctgtaaaat acctattatg actaaacaag
57061 attatttaga taatatagat tattttgaaa aagttattag taatgaaata tataatgttt
57121 tatatacaag tggatcaaca ggaaaaccag taaaaatatt tattactaaa aaagatgata
57181 ctagtcgtaa tatgattttg tgtatgtacc gtaatcgtat ttctaaaaat ataatttgtg
57241 agaaaatgct tttaatacat tatttgaata ttaaaggtaa tgaggagtat ttaattgagg
57301 ataaaggaaa acagtgtcaa catttttgtt atggatgcat aaatgatggt gtattagaag
57361 aaatatataa atttttaaaa caatttcaac ctaaatggat agttggaaat acatcaatta
57421 tttatgcttt ctcagaatat ataaagaaat caggaaaaag tttatcaggt atggatataa
57481 gatatataga gtgcatttca gaatatataa caacagagca aagaaatagt ataaaagaag
57541 tttttaattt aaatcctaaa agtgcttacg gttctaatga agtaaataca attgctatag
57601 aatgtgtcaa tggaaatatg catgtaataa aggaaagtgt gtttgttgaa atagtaaatc
57661 aaaatgaaca gggaattggg gatatattag ttacttcatt agataataga ttaactccat
57721 ttcttagata tagcttagga gataagggaa gatgggttaa tgcagaatgt aattgtaact
57781 tcaaatctcc tatattggag ataactagat atagaagcaa tgattttcta ttaaaaaaag
57841 atggaactag aatggaagta tggaacttaa atagtgctct aagagaatca actaataaaa
57901 ctaattttca aataaagcaa tatcagttta ttcaaattga ttataactta tttaaattaa
57961 agattttaga aaaagaagat ggaaataata tgaataaagc gatgtttgaa agttatttta
58021 tagaagagtt cagtagtatt tttcaagaaa aagtaatagt tgaatttgag tatgttaaag
58081 aaacaattcc agttgagagt aaaaacggaa aatttaaata ttttaaatca atgcttacct
58141 agagagtaag ctaagatgta aaataaagtt taaatgtata gaatataaaa ataagtaata
58201 tatgattgtt gagggaaaag atgattaaag gaatttgttt cgacatgggg ggaacacttg
58261 ttaaatttag ttcgccacat agtctagttg atattataat ggaacattca gagttatcta
58321 aatctgaaat taaagatctg gttcgaaaac attttatgtt gaagatttgt aactctgatt
58381 ctgatattag tgagtttttta aagaaatgca gatgtaataa tagagataag attgtagaaa
58441 tcattagaaa tcgtaaggta acatatgaag ttattgataa agcaatagat acactggata
58501 ttttaaaaag caaaggttat aaaattggaa taatttcaaa tacttcttcc tataatcaca
58561 tatctacaga aattattggt ataagaaagt atattgatta tgaaatatat tcgtttgaag
58621 taggatttgt taaaccagat ataagaatat ttaaaataat gcagaagaaa atgggatttg
58681 ataaccatga gttaattcat ataggagata gtattacatc ggatgtaatt ggggctaata
58741 gagcaggctg gaaaagtctt tggtttaatg aggctaataa aaattcgggt gatagtatgg
58801 aaattaattg tttatctaag atagtaaatt tattaaatta gtaaatatat tcagacaggg
58861 gaagagtata gtgatagata ttaataataa attttttaat cttttatcaa aagatcaaaa
58921 aattaagtta gagttttata ataatacaaa aaagttaatt aatgataatg caacaattat
58981 aaaattattt gatgagcaag tattaagaac acctaatgaa atagcactta tatttaatga
59041 tgaacagata tcatacaaaa ctttaagcaa gaaagtagat ttattagcaa atagattagt

59101 caatatgagt atacctaata ctcaacctat aggaatattc ataaatagag gttttaatat
59161 gattatttca atattaggta tattaaaagc cggtttttcc tacttgccta ttgatattaa
59221 cttaccgggt gataggatta aatatatgat aaatgatgca aaggtaaact atattattac
59281 tgagcacttt cttgttaata aatttgtaga aaatgagtcc ataaaactta tttttattga
59341 agacgaaaat ttaaacagta ctggaagtgg agcatatagt aatgataata aaatggccta
59401 tataatatat acatcaggat cttcaggtaa acctaaaggg attatgatag ggcaaagatc
59461 agtatgtaat tttatagaag gagttacaga aaaaattaac tttaaagcag ggagtgcaat
59521 tttaaatatt gcaaccattt cgtttgatat tttcttctta gaaacgatag ttgcacttac
59581 taaaggttta cgagttataa ttgcagatga aataactcaa aaaaatttta aacttataaa
59641 tagattaatt aataagttta atattaaaat gatacagtgt acgccttcac aaatgaaaat
59701 attggtagaa tgtaacaata ctggaaacac tttgaaaaat cttgaaacta taatgatagg
59761 gggtgaatct tttgagatag agcttctaac ctcaataaaa aagttaacta aggcaaaagt
59821 atacaacatg tatggaccta ctgaaacaac aatttggtct agtattaaag aagttactca
59881 tagcgagaaa attacgattg gaagaccgat acaaaatact gaatattttg ttgttaatga
59941 atgtaaccag tgtgttccat taggtgaaga aggtgaactt ataatttcag ggttaggatt
60001 agcacatggt tacgtaaaaa aaagtcataa taaaagtttt atagattgtg attgtgtaac
60061 aggaaataga ttttataaga ctggagattt ggtaaaattt gatgaatcag gagagataat
60121 ttttattgat agaatcgata atcaagttaa aattagagga catcgagtgg aattaaagga
60181 agtagagctt agtataagaa aaaacttttc agtaaaagat gtttcagtag ttcctttaaa
60241 aaataaattg gatataaata tgttatgtgc ttatatagta tgtgattatg atattaatat
60301 tagtaatatt aaagctgtac taagtaaaga actaccatca tacatgattc caagttgttt
60361 tgcagtactt gagagacttc catatactca taatggaaag ttagattata tatctttaac
60421 tgattatact ttttatagta aatgtgcatt agtaactaaa aataatttca ttgaattgga
60481 agatgaaata cagatgaaca actctattaa tgaaaaatta aaaagtttag taaaaaaagc
60541 agctaatata gatagttcta taaatataag cataaatgag aatttaaaca atttaggaat
60601 tgattcatta tcatttatat cactaatagt cgatattgaa tacgaattca atatagtttt
60661 tgatgatgat atgctgttgt tagaaaagtt taggagtatt aatgatttag ctaactatat
60721 aaaaatacta ataaaataaa ttagactata tagttattat gttaagaaaa gtttaaggat
60781 atatttaata gttgaatttt gtaattttat ttaaaaaaag taagtgattt atggatggtt
60841 gaaaattatt tataaaattt aattttagtt ttatatggta tttaaaataa tatttaaatt
60901 aaggagtaaa gatgaaaaag atagaggaat ttaaaagatt atcaaaaaaa tataatattg
60961 gagttgaaga tatagtatta atagcattaa atgtttatgg tgttaatatt gaagattcat
61021 ccccaagaat aaggttaaag ttagtcttag atactaaaaa aggagaggaa ttttattttg
61081 gattacctaa cagagttgaa tcttgttttg aaataaaaaa taatatttta tatttagaag
61141 gaaatagagt ggccataatc actaatattg aaaatgatga ttgtgcatca agctattata
61201 gaaaaaataa aactgtaata acgttgaact ctaattcaag aagtaaatgt aatggatgca
61261 agttttgtcc aaataattta gaattaaata gtgaagatga ttctttggtt tcaaatatta
61321 agttgaggaa acattttatt aatatgttaa ataaagcagg tcataaggat atgagtaata
61381 tcgaaagagt tacaatttgt acaggatgct ttggagatga aaaaagatta gtaaatcata
61441 ttcaaatagt aaatgatgtt ttagttgaaa tgaattttaa aggttatctt cattacatag
61501 gttcagaaat tacatctaaa gagtctttgg atgtaatttc tagaaatatt aataagttta
61561 tgattacatt aacagtagaa tgttttacac gaagagattt aatattaaag tcgaataaag
61621 caaaaataac tatagaaaaa tatattgaat taataaaaca atgtacaagt ttaggattta
61681 ttactaatat aatatatata ttggggctag atccattaga cgatgttata aaaaatatgc

61741 aattgttttc aaagtatact aactactttc catcaattaa tatatttcaa gttcacaatg
61801 gacaagataa aagtttatgt tgtgatgagt atgataaatt agaatatata ttaaaagcaa
61861 gaaagaaaat tgaagaaata tatataaata caaatttacg tccaagctct tgggagtgtt
61921 atagaccttt atggtacttt gagtttgctg atgaaaaatt atgcggtata agaatatagt
61981 taaaaaggtg attagactat gaataatgat gaaactttgg ttaatataga taaatactgg
62041 agagcttcta actatttaac attattatgt atgtatataa ataaaaattg tatgttaaaa
62101 aaaaatttga ggcctgatga tttaaaaaag atgtcaacag gacattgggg agcaagcaca
62161 ggaataaatt ttatatatgc taatttaaat gcttttatat cgaaatataa ggtagattct
62221 attttagttg taggaagtgg acatagtgca gcttctatat tagcaaattt atatttggaa
62281 ggtactttac aacattattt tgaggatatt actttagatg agaaaggaat tgaaaattta
62341 atagccaaat ttggaaaata taatggattt agaactgaaa attatcctgg tatacctggt
62401 aatatatatg atggaggaga acttggatac tctttatctg ttgcttttgg agcagctttt
62461 aatttgaaag agaaaatagt gttttgtata ttaggagatg gtgaatgcga aacaggatca
62521 atagctactg gatggaatgg aataaaattc ataaatccta cagagtctgg tgtggtatta
62581 ccaattataa atttgaatgg gtttaaaatg ggatcaaaga gtattctttc tttaaagtct
62641 aataaagagc ttagagatta ttttagtgga ttaggctata atgcttttat tatcaatagt
62701 tcacataaag aattatttaa tgcattagaa gaatcttacc gtataattaa atataatatt
62761 aaaatgttta aagagaataa aaagaataat ttgaaatggc cagttattat atttaagtct
62821 ttaaaggggt taacgggtgg ctatgatgat actaagcaat taatagaagg tagattaatt
62881 tcccacaaaa caccaataat atcaaaggat atagataaaa aaactatatt attagaagaa
62941 tggctattat cttataaacc agatgaacta tttagtgaag atggaaaatt agttccatct
63001 atatctagta ttattcctaa taaagatttg cgcttaggta tgttaaatag taaaattaat
63061 aaaataaaaa ataaattaat attacctgac ttaactaagt tctgggtaga tgaaccttat
63121 tatagtaata taaaagtatt agataaatat tttaatgatt taataatatt aaatccaaat
63181 atttttaaaa tattttctcc tgatgagtta gagtcaaatg gacttaatat tttggcatct
63241 aatcattctt atgatataag caataataat ggtaataaat gtgtactaga aatattgagt
63301 gaaaatactt gtcaaggttg gatgcaaggt tatcttatga tgggaaacca tggttttttt
63361 atatcatatg aagcatttgc tccaatagta tcaagtatga tgagtcaata ttttaaatat
63421 ttaaagcaag ctagcaaaat aaagtggaga aacgaaatta attcatttaa ctatatatta
63481 acaagtgtat gctggtctaa tacttattca catcaaaatc cagactttgt taataatgcg
63541 atttgtagag atctatctta tataaatgtc tattatccta gtgatgctaa tgttctcctt
63601 gcttgcgtaa atgaatgcaa tcaaattaaa aataagatta atattattat tgtatcaaaa
63661 cgaaaattac cacaatatat tgatagttct tatgctttga aagctgttga agatggatat
63721 tatgaaatta attttacaag taacaataaa gtaaagttag atataatact agcatgttca
63781 ggtgatcata tggtaaagga gtctattgaa tgtgcaaatt tattaaaatc ttatattaag
63841 aatataaata ttaaaattgt tgtaattata gaaattacac gtttatgtga aaaaagagga
63901 aattatagag gaattaatga aagtattttc aagaagatat ttccttatga tacaaatgtt
63961 ttatacaatt atcatggata tccatcggta ataacaacac ttctatttga tagaaagcat
64021 agtaatcgat ttaaagtatt aggatattct gatgaaggtg atacttcagc tccagaatta
64081 ataaaaatga ttttgaatgg agtaagtaga tataacctat gtcttgaggc aatagatgtt
64141 ttaataaaaa ataaaaaaat agattttgat tattacaatt atataaaaga tgatttaaat
64201 ttaaaaataa ataagacttt tgaagagtta gagaggtgat ttacattagt tcatatgaat
64261 tttggaaaca gtatataaaa gatataacaa aacaattaga aaagttattg aaatctaaaa
64321 aagaatatat tttagcagac ttacatatac attcaaatta ctcatgtgat ggaatgcaaa

64381 cattaaaaca aattttaatg tttgcatcag aaaaaaaatt agatgttata tcaattacag
64441 atcatgatag tatttcagtt tataatgagt tacctaaaga aatagagatt ttgaaaaaaa
64501 aaggatataa gctaccaatt ataataccag gagttgaatt ttcagtaagg tttgacgaat
64561 atgggagcat gtgtcatata ttaaagtatt ttattaatcc ttttgatcct ataattatag
64621 aagagattaa cagaaattat aaagcttact ggaatagagc tgaaattcag attgataggc
64681 ttaagtataa taaagcagta atgagctatt tcagacctaa tagtataaaa gtctctataa
64741 atgattttaa agagttttta tataataatt acaaaatacc aattccagaa tataatagca
64801 ttgcagaata tatattttca aaattaaatt cagtaggtgt aaatgtttct gatgttcttg
64861 aaaaatcgat attaataaac aatgaagaca aatgcatagt gagaagagac ttgaaaaaat
64921 tagcatttaa tagatttatc aataagtata aaggatgtaa cataaataat tcaaagtact
64981 tattacccat attatctgtt cctaatatag atgatgagga atttattaat tttgatagta
65041 gcggtaattt atcagtaaat aactatggac aaattcgctt tgaagattta aatagaaaat
65101 atattactgt atttgctcat ccaaatgaaa ataaattaga aattttatca aagtattttt
65161 atcagaatca aattgttgat ggaatagaaa ttaataagag aaataggcat tgtgaaaaag
65221 ataaaatact aaaatttgtt agatcgcata atatgtttta tacaataggc tcagacgatc
65281 atggtgatca tttatatact tatgaggata ttaattatta taaattaaat tcaatagaat
65341 ttaaaaaaat tttaaaatta gttatgagga gagaggaact atgaccaata ttaataaagg
65401 tagtcagtac attgactcaa gaaatctaaa tgtaagaatt aaattacatg aaatgtttag
65461 tactaataag caaggttttc acgaatggat atttagccaa ttgaatctca aagaaaatat
65521 aaaagtttta gaaattggat gcggtcctgg agcattatgg tataaaaatt tcaatgaaat
65581 tcctaaattt attgatatta ctttagttga tttatcagca ggtatgttag aagatgctaa
65641 aaaaaatatt gaagaggtta caagtagtaa aaactttaaa tttgtaaatg ctaatataga
65701 gaaacttcct tttaaaaata aagaatttga tatagtaata gcaaatcata tgctatatca
65761 tgttccagat atctcggcag gtttagctga agttaaaaga gtattaaaag ataatggtgt
65821 gttctattca tcaactttta gtaaagatca tttacatgaa ataaattgca ttacaagaaa
65881 atttgttgat ataaagaaaa atagaacatc agatcgtttt tgcatagaaa atgctgaaga
65941 acagcttaat aaacatttta gtagtattaa attatttatg catcaagata gtcttattat
66001 cacagagtct caaccgttaa ttgattatat aatgtcaggt agcaaagcta aggaaacttt
66061 agttggtgat aaatataaag aattttcaaa gtatattaat gagtgtgtat ttgaaacagg
66121 tgtttttaat gtacaaaaaa aagcaggtgt atttatatca aaaaaataaa aaaattaaat
66181 tatatgtata aaaagtttat agaaaattaa agggagcaat aatagagaat gtagaataat
66241 ttgtgtaatt taatattcta tatcggttat ataactggaa attttatttc cagttatatt
66301 tttatatttt tcttcaaata ctaataatga gttagtatgt taattttagt atgtgcaaat
66361 ttttgaaata tatactttaa atccaaatat gtacaagcta aattgctagt gcagtttcga
66421 ttctctccac aaaaattatt tcaaattgta gtagaatagg accccaattt gctattggtg
66481 aagtccattt tttcatcact tgatctgtag ctaaatataa agacttcctt agtgattcat
66541 ctgttggaaa tactgttctt atcttagtaa attttcttaa ctgtctattg aatccttcaa
66601 acaaatttgt agtataaatc atttttctta taccacttga aaaatcaaag aatgtactta
66661 aattattcca attagagtac catgaatcta tgacattagc atatttataa tcccattttt
66721 cttttaacta atccattctt accaactaaa tcttcaactg atttacaatt tttaattttt
66781 tcttgataat tgatttcttt actcatttga aatcctcctt aaaatattaa gaatattatt
66841 ccaaattttc cagctaagta ttcaaaaaaa ctgtagggaa atttgttttt acacaaattt
66901 ccctacattc tctaaccttc ttttttaacc aaactcattg ttaatatcat catcaaattc
66961 atcttctgaa atatcatttt tatcattatc agaattagca aaatttttat cagaattctt

67021 agtaatactt atttcgttac caacaatcaa attataaatt aaaaacttta tataggttcc
67081 ttttttattc ttcttatctt ttactaaatc ccatatggct gcatctctag gatcatctaa
67141 atttaatctt aagttaacac gctctatatt tgccatgact atttacccca tttagcttca
67201 gctatcaatt tattaccttt cacattacta aataaaggat ctggaacaaa atttatattt
67261 cctaagtttt ctcttaatgc atcttgtaaa tctaagctgc caccgccagc tccccaaatt
67321 ttataagtag aaaaattagg aactttaagt ctaaccttat ttaatttatc tttaacaaat
67381 ctctcttttg ctttacttgc atcaatagta tcattcttaa ttagtcttat tatttcttca
67441 acatcagcat tattacctag tttattatag tcttcttgta tttcagaata taaatctatc
67501 ataccaccag gaacagtaaa tttcataata catttaccac cagaagatac acatatattc
67561 catgtcctac cacctatatc caatatcact aaatctttta ccctttcttc tttaggtaaa
67621 gtataaaatg agcttagtcc ttctgcaact gtagcaactt tatttatttt tatattttta
67681 gtaacttcaa ataaattaaa attaaaggac ttattaagaa gcttttcttt taattcttca
67741 cttattccta aattagaagc tggtactcct aataccaaat cagtctctat attatctaca
67801 tcacatgatt ttgaaattgc ataaagtaag ttaggtataa agtttttctt agctttatta
67861 aactcattct caaaatctcc agattccatt ctataaaatt tattttctaa ttctaaaaga
67921 tctgtatcaa aatcttcttt ttctacttct tgatatctac tttcaaatat aattccttca
67981 catgttttga tattataatt tcctaaatca actgctatta tcaattttaa aacctccatt
68041 ttttaaaaca aatcataagt cacttaaaag taattttaaa gtgtttaaaa agttactttt
68101 aagtgactta tattattata atatcaaaaa gttacctaaa agtcaacttt taatcaccaa
68161 aaagttactt ttaagtgact ttagaaaaaa tataataaaa aagagagtaa aattttctat
68221 aattttactc tcttgaaaaa ttatttttaa catggtatta ttaactcaat taaattaaat
68281 agtacggaaa aggaaaaatc cactaagttt cctcgactcg ccaaagtaag taaacttaat
68341 ggatgaacct aaatccgaaa tagaacaaag ttctatttga aatattcact tttcatagct
68401 ttactatatc aaagctattg aaaaaagtca atgatatcta ttgacttttt aatagaaatg
68461 taatattagc aaatttttac ttctttctat ttaggattta gtgtactttc ctagatgaaa
68521 ggagttattt ttatggcata cacacactta ttaactaaag aacagttaga acaagagaaa
68581 aggtttaatc tcgaagaaag agattttcat gtcttaaatc aaatcgaata taaagagtct
68641 agaagaaaag ctcaaagtat cttaagattt attagaaaag gtattttatt aaataatgga
68701 tcttggagca tctcattttc taaaattcat aaagattata atgattgggt aaataaaaac
68761 aaaaagaaaa gaccagaatt aagaaatata agcttaaaac aaataaaaaa tatagtaaat
68821 aaattaaaag atttaggttt acttattatt gaaaatgtta aaaaaagaaa ctgttatttt
68881 ctaccattac ccaataaatt acccaataat gaaaacatta caatccctga tacaacaagc
68941 atagatgcta atcaagctac tcctaggtat attaggaata ataatataga tatagattct
69001 aatagtaact caaaagaatt taatgcagat atgtatgaaa aatgtacaag tcttgttgat
69061 gttagaaata aagttaaaga gttacttaaa gctgcaagag ttaagagctc ttggattaag
69121 aataaggttt taacaaaatt aagtgaaaac tatagaaata taactgttaa atttttagag
69181 agttatatta atacagtcat tgagaacact agaaatacat attattcaaa ttatagaaaa
69241 tatatcaaga acaatgcgaa caacagagct ctaccaaact ttacagaaag aaattattca
69301 aaagattact ggaaatacct agaagaaaat ctatgtttta attaataatt taacaattaa
69361 tagaatttat tgttaaaaaa tagattataa ataaactgaa aaatttccag tttattaaaa
69421 attgggaatt accatatttt tttatagata atttgataca atgagaacag aagttcgtat
69481 gtatatgaga acaatcgttt gttttggtta ttgtcaataa aataaaatgt ttttatatag
69541 agcaatgata tttattgctc tatataaaat ataaatcctt gagtatataa cttaatatta
69601 aattttatga tttaaatagt atttaatttt tcttccagat attcaatatc gatattttct

```
   69661 aagtcaagaa tatcagttaa agatcctaaa actactccga ttatattaat atcactattt
   69721 ttatcataat attgtatttc atatggaagt gtactacatg ggtgtagttc taatttatct
   69781 tttttcttat gtatttttt acaagtagct tcattgtttt cacttaaata agctattact
   69841 atatcgccat caataggagt tttatttcta cagcaaataa ttaattcatt gtccttgaat
   69901 aatttattca tactgttccc ttttactctt aaggcgaaac aatccttgta attgctaggt
   69961 aaagcaattg atttactaat ttcattaaat gcatatgatg gggttccgca a
//
DEFINITION Plasmid PLASMID_3
     CDS             1..897
                     /color=7
                     /locus_tag="D90_04613"
                     /eC_number=3.4.21.88
                     /note="LexA repressor  Helix-turn-helix domai, similar to
                     AA sequence:UniProtKB:P31080 protein
                     motif:CLUSTERS:PRK00215 protein motif:TIGRFAMs:TIGR00498
                     protein motif:Pfam:PF12844.1"
                     /product="LexA repressor  Helix-turn-helix domain
                     [transcriptional regulator]"
                     /"ID=D90_04613
                     /protein_id="gnl|SC|D90_04613"""
                     /gene="lexA"
                     /codon_start=1
     fasta_record    1..11010
                     /ID=D90|SC|contig000004
                     /label=D90|SC|contig000004
                     /note="D90|SC|contig000004"
                     /colour=5
     CDS             complement(930..1244)
                     /color=1
                     /locus_tag="D90_04614"
                     /note="hypothetical protein, ab initio
                     prediction:Prodigal:2.60"
                     /product="hypothetical protein"
                     /"ID=D90_04614
                     /protein_id="gnl|SC|D90_04614"""
                     /codon_start=1
     CDS             complement(1257..2192)
                     /color=10
                     /locus_tag="D90_04615"
                     /note="ParM/StbA family protein, Plasmid segredation
                     protein ParM , ab initio prediction:Prodigal:2.60"
                     /product="ParM/StbA family protein  Plasmid segredation
                     protein ParM [partitioning system]"
                     /"ID=D90_04615
```

/gene="parM"""
/protein_id="gnl|SC|D90_04615"
/codon_start=1
CDS complement(2389..2886)
/ID=D90_04616
/color=1
/locus_tag="D90_04616"
/note="Ribbon-helix-helix domain-containing protein"
/product="Ribbon-helix-helix domain-containing protein"
/protein_id="gnl|SC|D90_04616"
/codon_start=1
CDS complement(3019..3588)
/ID=D90_04617
/color=8
/locus_tag="D90_04617"
/note="Type III toxin-antitoxin system ToxN/AbiQ family toxin ToxN"
/product="Type III toxin-antitoxin system ToxN/AbiQ family toxin ToxN [toxin-antitoxin]"
/gene="toxN"
/protein_id="gnl|SC|D90_04617"
/codon_start=1
CDS complement(3902..5473)
/color=4
/locus_tag="D90_04618"
/note="Relaxase/Mobilisation nuclease domain, protein motif:Pfam:PF03432.8"
/product="Relaxase/Mobilisation nuclease domain [relaxase]"
/"ID=D90_04618
/protein_id="gnl|SC|D90_04618"""
/codon_start=1
CDS complement(5434..5763)
/color=4
/locus_tag="D90_04619"
/note="Bacterial mobilisation relaxosome protein MobC, protein motif:Pfam:PF05713.5"
/product="Bacterial mobilisation relaxosome protein MobC [relaxosome]"
/"ID=D90_04619
/gene="mobC"""
/protein_id="gnl|SC|D90_04619"
/codon_start=1
CDS complement(6001..6888)

/color=8
/locus_tag="D90_04620"
/note="Site-specific DNA methyltransferase, protein motif:Pfam:PF01555.12"
/product="Site-specific DNA methyltransferase [methyltransferase]"
/"ID=D90_04620
/protein_id="gnl|SC|D90_04620""""
/codon_start=1
CDS complement(6956..7249)
/ID=D90_04621
/color=8
/locus_tag="D90_04621"
/note="type I restriction endonuclease subunit M"
/product="type I restriction endonuclease subunit M [methyltransferase]"
/protein_id="gnl|SC|D90_04621"
/codon_start=1
CDS complement(7493..7696)
/ID=D90_04622
/color=1
/locus_tag="D90_04622"
/note="hypothetical protein"
/product="hypothetical protein"
/protein_id="gnl|SC|D90_04622"
/codon_start=1
CDS complement(7720..7827)
/ID=D90_04623
/color=1
/locus_tag="D90_04623"
/note="hypothetical protein"
/product="hypothetical protein"
/protein_id="gnl|SC|D90_04623"
/codon_start=1
CDS complement(7914..8348)
/color=9
/locus_tag="D90_04624"
/note="DNA repair protein RadC  RadC-like JAB domain , similar to AA sequence:UniProtKB:Q8KES1 protein motif:CLUSTERS:PRK00024 protein motif:TIGRFAMs:TIGR00608 protein motif:Pfam:PF04002.9"
/product="DNA repair protein RadC  RadC-like JAB domain [repair and recombination]"
/"ID=D90_04624

/protein_id="gnl|SC|D90_04624"""
/gene="radC"
/codon_start=1
CDS complement(9489..10034)
/color=8
/locus_tag="D90_04625"
/note="Plasmid epsilon-zeta postsegregational killing
system Zeta toxin AAA family ATPase, protein
motif:Pfam:PF06414.6"
/product="Plasmid epsilon-zeta postsegregational killing
system Zeta toxin AAA family ATPase [toxin-antitoxin]"
/"ID=D90_04625
/protein_id="gnl|SC|D90_04625"""
/codon_start=1
CDS complement(10234..11010)
/color=2
/locus_tag="D90_04626"
/eC_number=2.7.7.47
/note="Streptomycin 3''-adenylyltransferas, similar to AA
sequence:UniProtKB:P0A0D1 protein motif:CLUSTERS:PRK13746
protein motif:Cdd:COG1708 protein motif:Pfam:PF01909.17"
/product="Streptomycin 3''-adenylyltransferase [antibiotic
resistance]"
/"ID=D90_04626
/protein_id="gnl|SC|D90_04626"""
/gene="ant%289%29"
/codon_start=1
BASE COUNT 3303 a 1965 c 1264 g 4478 t
ORIGIN
1 ttgaaaaata attacttttg tgaaaatcta aaatatataa ggagttcttt aaatatcagc
61 caaaaagaaa tagcaaatca cttaaaatta aatcctagct catattctaa ctatgaaaat
121 ggtcgccgag agcctggtat agatgtttta ataaaaattt ccgatttcct taacgttgaa
181 atagataaac tcttgactga aaaccttagc aacaatgata aatttttgaa ggatattcaa
241 aacaaaattt caaatgatat taattatgat gaaaaaatta aggatattaa tttagatatg
301 cgtaaaaata tcttatttga attggagaat aaaaaaagaa aatgtttatc cctaattaat
361 tatgaaattc caaaaaaaat taaagaaatt gataatctaa ttaagtatat aaatctttat
421 actgatcaag atgtagagat tagtgatgaa atctcaccta atactagtga atttaaaact
481 aatgaaaaaa gtattgacta tagatctata aacctaattg gaaaagtatc tgctgggaat
541 ccttgttatg cttacgaaga aataataaac acttataaaa tcccttccaa tatcctatgt
601 ccttctaaaa attattttat tcttgaagtt aaaggagata gtatgaataa aatctatccc
661 cctggtgatt taatattaat tgaaagtaca aatttagtta ataatgatga tattgttatt
721 gctataattg atgatgaagc tacttgcaaa aaaatcaatt tttctcctaa tgaaattatt
781 ttaacccctc aaagcactaa cccaatacac caagttcaaa tatataaaga tattaatgtt
841 catatattag gtaaagttct tggtaaactt tctgattata taagaaaaga tgagtaatag

901 tatatattac tcatcttttt atattttaac taatcaggta tatcatttaa taactcatta
961 atagagttct cataatcagt attttcctta tctacagata tttcttcaca ttttttttta
1021 ttatcttctt gtaacctatc tagtactttt tctactattt tattaatatc tatattggaa
1081 tgtacgttac ttgttccatt attattccct tttagaactg cccattctaa tactattctt
1141 tctaaactag ctgaatcact tgaaaggttt ctatcctctt tattttccct aacaatgttt
1201 aaagtttctt cttctattgt ataggtctta cttatcttcg ccattttagt atctccttat
1261 ttaaagactt ttgtacccac aatgtaaaga cctttagcag ttgccatttg tggtgcttct
1321 actaaagtac cattatatat tttcattctt tgagtagttc ctccaccaaa ataaatatcc
1381 atttcttcag tattaaccca ataactatca acttcttgag ctactttttc tgaaagtatt
1441 ctgtatcctt tttcctttaa tttatcataa tcaccagtat tactatctat ttcaaataac
1501 tctttagata aattatattt atcgtttaat tctctttgca catattcaag agctgtctta
1561 tttcctaatt caagcgattt agatttttta tcattaaact tcattccctt atcaaagtat
1621 gtaaattctg tagttctaaa acctacagtt accattccca taggtttatt agtagttgtt
1681 ctatctctta attcccaaaa taatgctgca tctgactctc taaatattga tatatcttct
1741 atgaaaactt tcttataaca tttagttata ttattcatta tctttactgt tttaccctta
1801 taagcttcta tcacttctgt taaagttctt ttattatatg atttatttgg aactcctaat
1861 acaagtttaa cactattagc tactgctaac ttatctaata atgcatataa taatttttca
1921 gcagttaacg taatcttaga ttccatatta ttagaagttg gagaataacc ttctacttca
1981 gctaatatac cagcaaaata ttcttctcct tcaacttcta aatatatcgg atctttatag
2041 tcatcaaaat ctatttctct tcctagacta actattgatt taaagcaaca ttcactagct
2101 tctccattgt aaactgtata accttttgta tatccacgcc caatatctgc tcctattatt
2161 tgagattctt gattaataat atttttctcc atttttacac ctcttttttt atttcctaat
2221 tactatattt atatagtaac acataaaaat tgttatgtca atgctatatc aatttttcaa
2281 ctaaaaataa ttgatatagc attggtatat aaattcaaat ttaaatatta attgttatgc
2341 caatggtata tcaatttaac atgttttgaa aattgatata ccattgtttt ataaattttt
2401 taaatattgt tctaaagcaa tatttattat ttctccaata gtttcattgc ttttataaga
2461 taattttttt atctcttcat aaattccagt atctatccta aatgatctta catcatttat
2521 attacttctc aatcctgtaa tattacttgt aatattgttt gaattctttt tattaattaa
2581 tgataaaatt tcatctaaat tatctaataa taataatagt ctttcattat ttacttcatc
2641 tttaaaattc ttactatctt ctctttgggt atttgcttct tttttagtaa tattacttgt
2701 aatatcattt tgtaatacat actgattatc tatcttttta tatcctaatc taataagtct
2761 tttatgaatt actcgttcat tttctccaaa tgattctttt tcgattttca ccattggttt
2821 tccattattc aactctttat ttataaattc aactattttt attaaattat ttttgttcca
2881 ttccattttt acacctcaca cttacttgta atattacact aatattgtac cattattta
2941 catataattg taaaaataag cgatttatat attaaattaa tatacaaatc gcttacctat
3001 atatatcttt atgaactttt aaatcttttc ttttttttta ttatgttcat cagctttttt
3061 tgaagcatct ttaacaatat cttttattcc tacagttttt gttactgctg cttctctaat
3121 atctgaatta tattctaaag cttttttttc tataagttca aaattaacac atcttttagc
3181 tataaagctt cttccactta taaccttatt atataattta ttagctttat ttttaattac
3241 atcttcattt ttatttataa atcttatttg ctttgttaat aatattgaat atttacgatt
3301 atgtctttct tcaaaattat actctactat attttcttta tgtactggta tcatattatt
3361 taaatttata acacctaaat ctccattttc caatttaaaa aaatctatat tattcttcat
3421 agatttatgt ttttctttag gtgatgccaa cggagctata taattcatcc cattaacttc
3481 taataatatc cctatatatg gacgtgattc atttttatta tctctaatat tatcatcaaa

3541 tttctttaga taatttatat aatcctcttt aactatatat attctcattc ccattctccc
3601 cttataaaaa aaaagtaatc aattttgatt acttttttca ctcttttagt ttctcgctta
3661 ctggtggaga ttcaccgctc ttttagtttc tcgcttactg gtggagattc accgctcttt
3721 tagtttctca cttactggtg gagattcacc gctctttata ctgatttata atcattatat
3781 tctatataat atattatacc attctgcata taaatattca actacattgt acttttattt
3841 atgcctttat ataaagataa ccaatgatcc taataactaa tgatctttgt ataagattaa
3901 tttatctctc tttttctttc tctttagtta agtcgatttt ttcttgtatt atattttcta
3961 ttttactatt agcctttctt atactttcta atcttttttc tatattttta aattcattac
4021 ctaataatga aatttcttct ttttgttcaa ttattttttc ttttaactta tctatactaa
4081 cgtttgtatt tatacctaag ttttctagtt tttctttagc atgaaggaac attaatatct
4141 cattttcatg ttttttatat cccccaaata tagttttctt atttccatgt ttctcgtaaa
4201 tttctttata attattgtat gttatcaaat atttagatac ttgattaaat ttagcgttct
4261 tattttctaa ttctttaata gtattaatta attcatcttt cttatttaat acttcttgta
4321 ctttaccatc aaattcattt ctatttgata tattttcttt tctcaataga actattgcat
4381 tggcaagctc tttaacatct cttatcctat ttcttctgat agtaaataca ttttcatcaa
4441 ttttaacatc ttcaacttta atatcattag atttttcatc ttgactagta aagtaattat
4501 ttatttgttt accagttaat atttcttctt tttctcctga agaatctata aatttatatt
4561 catttttga taatgttact ttacattcaa actcattaat ccaattacat tctgatttat
4621 ttataaatat atagttttca cttcctggca ctctaacaaa ataatttttt tcattttgat
4681 taataatcat atctttatta attattatct ctttagcctt gttattttca ttaattctat
4741 tttctatagc ttctttagta tattcttcac ctatagtttt tcctctgcaa aatctttgct
4801 gattaggtgc tttaaatgct atatattttc cttcctttat ttcaaaacct ctattaatca
4861 tttcagtttt taattcatca aatgacttaa ctttatctat taattcatca agagtatctt
4921 taatgtaagc tttccaactt gtacccttt tattctccaa ccactcttta taatgcttac
4981 cttttccttt aggattttct acaatactta atccattttc tttgcatagt tcatcactta
5041 ttgctctaag ttctttaaaa gtatctttat tacaattaaa ctttctatag gtcgcataac
5101 tataagcatt aaaaataata tgattatgaa tatgaccttt atctatatgc gttgcaacta
5161 catactcata tttttcatct gttaatttct ttgctaattc ttcaccaatt ctatgagctt
5221 cttcaggagt tattttatct tttctatcaa aagattgtat catatgataa gctaaattgt
5281 ttgatttttt actataatct cctatcactt ctttacaagt ttcttctgtc attttaaact
5341 ctaaatatgc agtttcaggg gcacaatttc ccccacttac taataacttt ccatcagttt
5401 tgctttcatc cattatataa tctaatgctt tttttaatgt tgacttgata gatttgattt
5461 tagtaactgc catacctgtt caagctctcc ttttaactct tttatatcat cttcataaat
5521 tctatttgtt tcatttgttc ttttagcaat ttgatttata ttaaccccaa ttttattaat
5581 ctctttaatt aaatttttta ttgtagtata atccgttttt actacatacc catctattaa
5641 catctttctt gcatatgctc caaagttatc tgttcctaac tgtttcattt tttcttttat
5701 tattttatct tcttcttcat tcacaaataa ttttttttga atatcccttt ttctattagc
5761 cataagttta tttcctttct aaaaataaat catttatgat ttattctatt gggggtttgg
5821 gggtatccct caacaagtaa ttttaaaatt aaagtttcct tttaatttta aaattaaagt
5881 gatgtgtgta cacactcact atgcttgcta cccttctcta tacctcttat tatacttttt
5941 aggaaaaagc atttcaacaa aaaagactag taattttttt accagtcttt atctttatat
6001 ttatactaat tgccatactc cacgttctat atttacgaaa atattaggat atatctgcaa
6061 tgtttgtctt atcttagctt tccaatttgg attattatat gtctttttat atccttctaa
6121 actcttatat aacttctcta agtttaccct cccccctaaa ctttcaattg ttgaagcaac

6181 aaggtctttc caagtaacct ttaaactatc tcttaaatcc atttttcccc tttttgtaat
6241 catgtaatct aaaattaaag gctcatcttt tctaagtatt aaacaatatt catgtgctat
6301 ttttataaaa ttttcattaa aatattttct atgattgctc atacaattat gttgttcttt
6361 aattactatt tgctctacag ttcctaattt attcatatct aaaagcattg aatataattt
6421 atgattcttt tttacatcag ccatcaatat tgccattctt cctccaactt tcaaactcgc
6481 atattgtttt gataggcaat aattaatagc ttttataaac tcttgataat ctttaatatg
6541 agataaatca tttttcaaag gtgtatctcc atacatatga ccactatact taataatatc
6601 ccaataaggt ggatgccaat atatgaaatt gtttctttct ttaatatcat tttctattaa
6661 atcaaatttt gtttcttctc ctttcaatcc atttaaatca taacaattag ttattattcc
6721 tagttttttt cctacatcat cagttgtaaa acttccactc atataatcac ttatttcatc
6781 aaatttataa attttatgta ggtcaattaa taattttcct gttgcatttc ccctatattt
6841 attatctcca tattcccctc tagttggata agaaactata ctattcattg ttaaaatctc
6901 cttttatttt tttattattt aagaaaaggt gttcccgagg aacacctttt taaaattagt
6961 attcttcagg aagtaagata gttattaata aactctttat tgaataatca tactcactca
7021 taatccaaat tttatgtcca tttatcatat agactcccat aaatctatct tcatattttg
7081 aattaggatt ttttaataat tcattttgaa attctttatc ctcattacat aaaattcccc
7141 aatcatcaga atggaatttt aataaagctg acattaaatc tttgtcttta aattccccac
7201 tttcaattaa ataattaact ccatttgtca taactactct caatttcata aatagcacct
7261 ttcctttctt gttgtttgtt ttttttgttg agcctaagac tacatattgg agtgagttgt
7321 aaaggaaaat ggaataaaaa attttggctg ctattcctgg taaggatcat agccgacaaa
7381 aatttttatg gaatggcttt acaacttgaa cggaaatatg ataggctggt gcttcaaaaa
7441 aaacataaaa ataaaaaaga agtccctaag gacttcttac ctttttgatt tattagacta
7501 ataatttaaa atcaatccaa tcaccaaaca tattacttat atcgtctata gcttcattaa
7561 tagaattata tttataactt ctttcatttt taaaatttat ataccaccaa ctgctcttat
7621 aatattcaac ttctataaat acaagttcat ttttaataga atttcttgct ctaaattcac
7681 aaattttttt tctcataata ttaccttcct taactatact taatctctcc aatatttttt
7741 cttatatcta tgaagtcctt caaaagtcat tgtttcattt actttatcac aatattcttt
7801 ataggttaaa agtacatata tactcatttc tattctcctt tactttaggc aaacaatgag
7861 ttataatctt attgtgagta agggtgtaac ttcattgtta tgcctttttt tatttataca
7921 attccttttt cttttaaact tgtataatca tcaccaatta ttaaatggtc taataattca
7981 attcctaata aatcccctcc gtccttaatt cttcgtgtta ttgaaatatc ttctttactt
8041 ggtgtagtgt cgccacttgg gtgattatgt gccagtatta tacttgcagc attacttaga
8101 atagcaggtt taaaaacttc tcttgggtga accacactag catttaatga acctatacat
8161 atagtgctta tagatgttat tttattttta gtatctaatg ttaataatac taaatgctct
8221 ctatcaactc catctaagta ttcagatact attttataaa catctttagg tgaaccaata
8281 ttcttattat ccacaagcat acttttcact ttttccatct taatagttac aatattaatt
8341 ctgttcaaaa aatcaattcc tttcttttta aattaacacc ctggtttaaa atcccatgtg
8401 ttgttgtttt tttgttgagc ctaagactac atattggagt gagttgtaaa ggaaaatgta
8461 ataaaaattt ttggctgcta actgcagatc cagttggccg acaaaaaatt ttatggaatg
8521 gctttacaac ttgaacggaa atatgatagg ctggtgcttc aaaaaaacaa tataaaaaaa
8581 agaagtccca aaggacttct tacctatttg gatgcactcc atcgggtacc cgcttatagt
8641 aaaaataagt ttagttattc tttgctttag cttggtcgct cggtagatgg aacaagctaa
8701 ggcaatagaa tatcttaact tattgaaacg ttataagtgg gtgatggggg gttcacttag
8761 ataaagatac cacatcaatc ccacgctgac ttttgctcaa ccgatacagc cggagggctg

8821 gataacaaga aaaggcggta tgcgccccgt ggaggggtag cgtaccgcct tttcttgtgg
8881 gggtttccaa agggggcaac gcccccattt ggcacacgac tttgcgaagc aaagtgtagt
8941 gtgttatacg ctctgtcggc gttgccgtga aaatgccgtt gccgccgggg agggcaaggg
9001 catttgaagc ggcaggaaaa cagggctgtg cttgcgtggc gtagagccgt aagcacaggt
9061 ctggtttcaa cagcagactg ggcgtatatg aaagcccccg tccctcgtcc tacgctccga
9121 cttgtggaca aagtgtcccg aagttgtggc tacactcccg gaaaagtaac aggacagcgg
9181 gcaaccgtca aggctgaaat gaacgggctt acacccggcc ttgaccgcca cccgccgttc
9241 cgctgaatgg gtgaacaagg cgaccaatcc ctcaaagtgc ttggccgctc tctttctgat
9301 tttggagtgt ttcttttctc ataattgaaa tgggcgggaa gccattttag ggctttcccg
9361 ccttgattgc ccatcagcaa agccagacgg gacggtcaac ggcggcgcat gaaatgcgcc
9421 gttcatcttg accgttgact ggctttgcta tttctctgat aaatgaaatt tcgattttta
9481 gaaatcgttt aagatagcaa tgcggctcca cttgaaattt tttcttttac tgcaaaaaca
9541 gcctcttgga tagaataggt tgtcgtttct ataacattcg attcatatat ccccaaatta
9601 caaaattgct cccacattgt ttctactaat tcaacatttg tttttctgtc taattttgaa
9661 cgttccacag ctcgtttcat agtttcttct ttactggctc ttaaaacaat atagtgtact
9721 tcataatctt ctcgaacaag ggctctccat ggttctaaaa accacggccc gacaattcca
9781 tctacaatta catcatatcc gccgcgggca tagcgctttg cagcttctaa aaaggcttca
9841 atgacaacta aattttgttc gttggactct ggtaaatgcg gcggtattgc tcctttgctt
9901 aaataatgaa aaaagtcgtc tgtgtgcata tgcacagatt tatccatgtt cgattctttt
9961 gcgacaatcg acgcagttgt tgtctttcct gttcccggcg aacctgtaat cacaataatt
10021 cttccttgat tcatctacat tctccctcgc aattcagatt taccgctctc tttctgattt
10081 tggagcgttt cttctctcaa gattgaaatg ggcgggaagc cattttaggg ctttcccgcc
10141 ttgattgccc atcagcaaag acgggcggga ctgtcaacgg cggcgcattt atgcgccgtt
10201 catcttgacc gttgattggc cccgctaact ttgctactct gttctaagta attcctcaat
10261 ttgtttttc atatattcta caagtgccat cgcttccttt tctactgtct cccattcatc
10321 agacaaattt cccaaataag cttcctttgc cgttgttagc agggggggga atctctccgg
10381 caattttaaa attacccatt ttgcggcaac atcttttgtc gtgatttctt ctgttactaa
10441 agtaaaccac attcgtgata aggttaataa cacatttctt tcatcaccct taaagctgga
10501 aatcaaacca ggtaaagaaa accgaattgc ttttttaatt tcgtgaaatg ggatagcggg
10561 aataagctct ttgctttccg ccccttcaa agttatgcta ttttttcttg cttgccataa
10621 taaaatcatt atatctgggt cattgcaggc ttgcggatat tctcctgctt ccatctctcc
10681 ccttagccat tcaccataca tatactgaca ttttggcgga aattgcaacg gaacaatatc
10741 agattgattg ataatagtta cctctaaagg tcttttttca atacagccta cggagccaga
10801 aattttcaat aattgctttg ttagatctgc tctgattgaa ttactcaatt cttgtttagt
10861 aattatcagt atatctatat cactatctgg acgcagccca ttcattgttg ctgaaccata
10921 taaatatatc cccaatattt gaccttgtaa tatggtttct gctattgaaa taacttgatt
10981 tacttgttgt ggaaattcat ttatattcat

//
